# Characterization of Extensive Diversity In Immunoglobulin Light Chain Variable Germline Genes Across Biomedically Important Mouse Strains

**DOI:** 10.1101/2022.05.01.489089

**Authors:** Justin T. Kos, Yana Safonova, Kaitlyn M. Shields, Catherine A. Silver, William D. Lees, Andrew M. Collins, Corey T. Watson

## Abstract

The light chain immunoglobulin genes of biomedically relevant mouse strains are poorly documented in current germline gene databases. We previously showed that IGH loci of wild-derived mouse strains representing the major mouse subspecies contained 247 germline IGHV sequences not curated in the international ImMunoGeneTics (IMGT) information system, which is the most commonly used database that curates the germline repertoires used for sequence alignment in AIRR-seq analysis. Despite containing levels of polymorphism similar to the IGH locus, the germline gene content and diversity of the light chain loci have not been comprehensively cataloged. To explore the extent of germline light chain repertoire diversity across mouse strains commonly used in the biomedical sciences, we performed AIRR-seq analysis and germline gene inference for 18 inbred mouse strains, including the four wild-derived strains with diverse sub-species origins. We inferred 1582 IGKV and 63 IGLV sequences, representing 459 and 22 unique IGKV and IGLV sequences. Of the unique inferred germline IGKV and IGLV sequences, 67.8% and 59%, respectively, were undocumented in IMGT. Across strains we observed germline IGKV sequences shared by three distinct IGK haplotypes and a more conserved IGLV germline repertoire. In addition, J gene inference indicated a novel IGK2 allele shared between PWD/PhJ and MSM/MsJ and a novel IGLJ1 allele for LEWES/EiJ and IGLJ2 allele for MSM/MsJ. Finally, a combined IGHV, IGKV, and IGLV phylogenetic analysis of wild-derived germline repertoires displayed reduced germline diversity for the light chain repertoire compared to the heavy chain repertoire, suggesting potential evolutionary differences between the two chains.

## Introduction

Antibodies (Abs), encoded by the immunoglobulin (IG) loci, are critical components of the immune system that function as cell-surface and soluble receptors for antigens(1). The process of somatic V(D)J recombination within B cells governs the formation of a diverse Ab repertoire capable of recognizing a vast array of antigens through interaction with Ab variable domains(2). Abs are formed from two pairs of identical heavy and light (kappa or lambda) chains, encoded by genes at three different loci in the mouse genome. In mice, the IG heavy chain is encoded by genes at a single locus on chromosome 12 (IGH), whereas IG light chain genes are encoded at the IG kappa (IGK; chromosome 6) and IG lambda (IGL; chromosome 16) loci(1, 3). Abs are separated into two functional domains: variable (V) domains that bind antigen; and constant (C) domains that carry out effector functions such as complement activation and Fc receptor binding(4). The V domain is encoded by variable (V), diversity (D, IGH only), and joining (J) genes, while constant (C) genes encode the C domain. Together, V, D, J, and C genes somatically recombine in B cells to generate the Ab repertoire, the entire expressed component of Abs circulating within an organism.

The mouse IG loci are structurally complex and consist of repeated, highly homologous gene segments(5–7). For example, the IGH locus in the C57BL/6 strain comprises 102 variable (V), 9 diversity, 8 of which are unique (D; IGH only), 4 joining (J), and 8-9 constant (C) functional/open reading frame genes(5, 8). The IGK locus of C57BL/6 is similarly complex in this strain and spans 3.2 Mb, with 91 functional V segments, 4 functional J segments, and 1 C segment, representing approximately 95% of the C57BL/6 germline light chain genes(9, 10). In contrast, the C57BL/6 IGL locus spans 240 kb and includes only 3 functional V segments, 3 functional J segments, and 3 C segments(5).

At the genomic level, the mouse IG loci have only been comprehensively characterized in the C57BL/6 strain. However, the C57BL/6 mouse does not fully represent variation within the mouse IG loci. In 2007, Retter et al. sequenced and assembled bacterial artificial chromosome (BAC) clones spanning the IGH constant region and part of the variable region in the 129S1 mouse strain(11), which was predicted by restriction fragment length polymorphism (RFLP) to carry a divergent IGH haplotype compared to C57BL/6(12). They showed that the IGH^A^ haplotype of the 129S1 strain is genetically different from the IGH^B^ haplotype of the C57BL/6 strain, containing major germline gene duplications present in the IGH^A^ haplotype that are absent in the IGH^B^ haplotype. Though the light chain loci have only been characterized in C57BL/6, early RFLP experiments reported the existence of 9 IGK haplotypes in commonly used inbred mouse strains(13), and more recent Sanger sequencing identified significant IGKV polymorphisms in NOD mice(14, 15). In addition, early isoelectric focusing experiments of mouse light chains highlighted IGK haplotype differences by showing that SWR/J, C3H/HeJ, DBA/1J, A/J, CBA/J, and C57BL/6J had identical focusing bands, whereas AKR/J and C58/J had observed differences(16). More recently, genome-wide high-throughput single nucleotide polymorphism (SNP) studies have revealed inter-strain diversity across all three IG loci(17, 18).

The more recent application of high-throughput Adaptive Immune Receptor Repertoire Sequencing (AIRR-seq) studies has also led to the discovery of extensive variation in the germline IGHV genes. For example, AIRR-seq studies of C57BL/6 and BALB/c mice demonstrated that the BALB/c IGHV germline set consisted of >160 genes, only 4 of which overlapped with those found in C57BL/6. In a subsequent study of 5 additional inbred strains, including 4 wild-derived strains representing diverse sub-species origins, we observed significant inter-strain variation in IGHV germline sequences, and catalogued 247 germline alleles unaccounted for in existing reference databases(19). However, despite evidence of potentially similar levels of diversity within the IGKV and IGLV coding regions, these loci have not been comprehensively explored across inbred strains.

In this study, to better understand mouse light chain germline diversity, we conducted AIRR-seq analysis and germline inference in 18 different inbred mouse strains, again including 4 wild-derived strains from diverse sub-species origins, as well as an additional 14 strains commonly used in biomedical research. Consistent with our observations in the IGH locus, we observe significant germline sequence variation between strains. In addition, inferred IGLV genes across the classical and wild-derived strains reveal the presence of fewer germline genes in classical strains than wild-derived strains, which may result from the breeding history of classical laboratory mice. This level of germline diversity is unexplored in the study of immune phenotypes and unaccounted for in existing gene databases. Despite the diversity observed, we uncover evidence for the presence of shared germline IGKV/LV gene sets and haplotypes among subgroups of classical laboratory strains, indicating that the light chain loci reflect shared sub-species origins.

## Materials and Methods

### AIRR-seq Library Preparation and Sequencing

Whole dissected spleens, preserved in RNAlater (Thermofisher, Cat. No. AM7020; Waltham, MA, USA), were obtained from female mice from Jackson Laboratories (Bar Harbor, ME, USA; https://www.jax.org) for eighteen inbred strains [BALB/cByJ (Jax stock #001026), n = 1; NOR/LtJ (Jax stock #002050), n = 1; 129S1/SvlmJ (Jax stock #002448), n = 1; MRL/MpJ (Jax stock #000486), n = 1; A/J (Jax stock #000646), n = 1; AKR/J (Jax stock #000648), n = 1; CBA/J (Jax stock #000656), n = 1; C3H/HeJ (Jax stock #000659), n = 1; C57BL/6J (Jax stock #000664), n = 1; DBA/1J (Jax stock #000670), n = 1; DBA/2J (Jax stock #000671), n = 1; NZB/BlNJ (Jax stock #000684), n = 1; SJL/J (Jax stock #000686), n = 1; CAST/EiJ (Jax stock #000928), n = 1; NOD/ShiLtJ (Jax stock #001976), n = 1; LEWES/EiJ (Jax stock #002798), n = 1; MSM/MsJ (Jax stock #003719), n = 1; PWD/PhJ (Jax stock #004660), n = 1].

We extracted total RNA from 30 mg of spleen tissue using the RNeasy Mini kit (Qiagen, Cat. No. 74104; Germantown, MD, USA). For each sample, IGK and IGL 5’RACE AIRR-seq libraries were generated using the SMARTer Mouse BCR Profiling Kit (Takara Bio, Cat. No. 634422; Mountain View, CA, USA), following the manufacturer’s instructions. Individual indexed IGK and IGL AIRR-seq libraries were assessed using the Agilent 2100 Bioanalyzer High Sensitivity DNA Assay Kit (Agilent, Cat. No. 5067-4626) and the Thermofisher Qubit 3.0 Fluorometer dsDNA High Sensitivity Assay Kit (Thermofisher, Cat. No. Q32851). Libraries were pooled to 10 nM and sequenced three times on the Illumina MiSeq platform using the 600-cycle MiSeq Reagent Kit v3 (2x300 bp, paired-end; Illumina, Cat. No. MS-102-3003); per sample read depth is provided in Table 1.

**Table 1.**
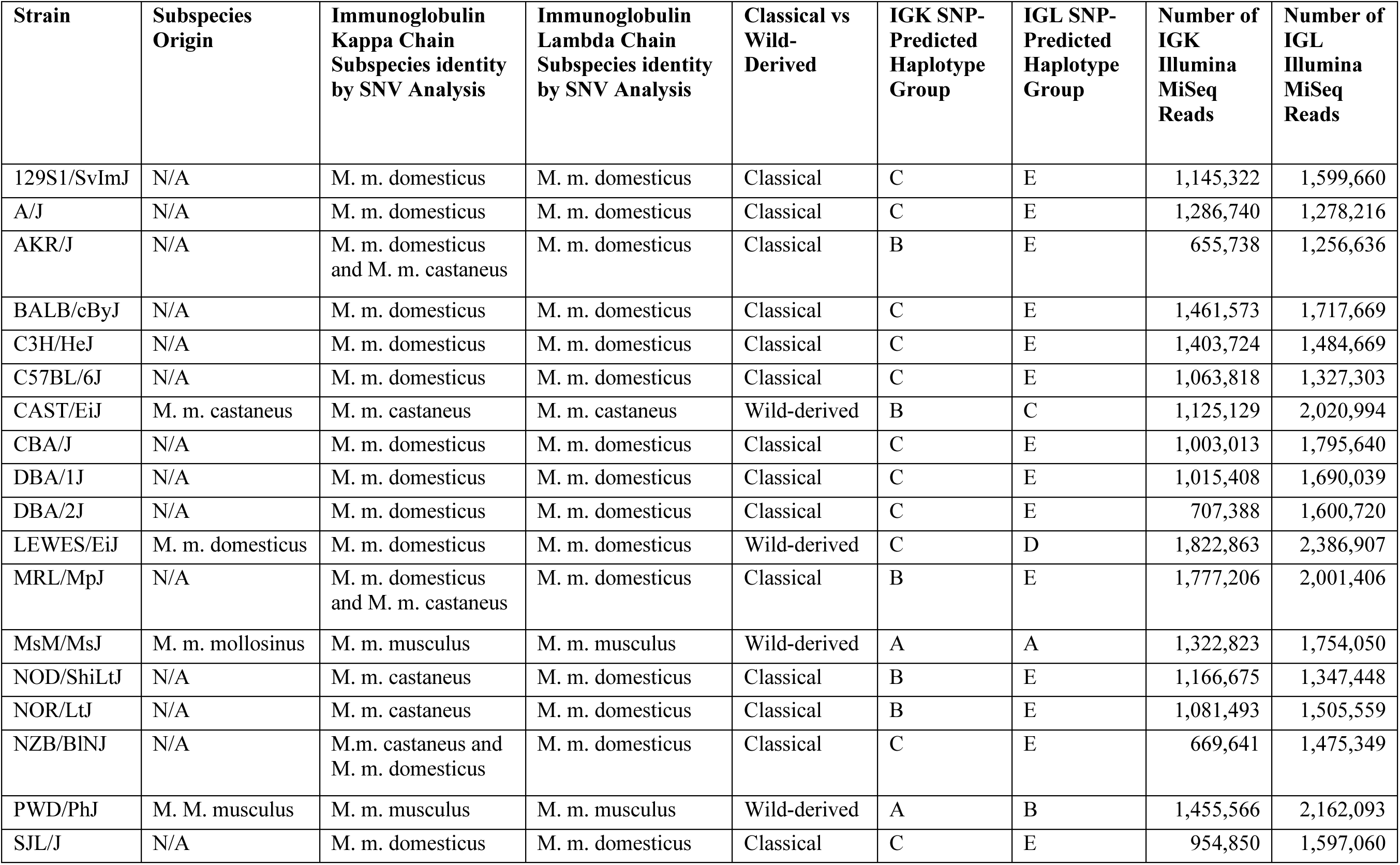
Subspecies origin and subspecies identity of the immunoglobulin kappa and lambda loci of classical laboratory and wild-derived mouse strains selected for IGKV and IGLV germline gene inference.

### Data Processing and Germline Gene Inference

IgDiscover v0.12(20) was used to construct a germline IGK database for each strain. We combined FASTQ reads from each MiSeq run and processed using IgDiscover v0.12(20) using the following parameters: (1) "barcode_consensus" set to false since samples did not have barcodes; (2) "race_g" set to "true" to account for the run of G nucleotides present at the start of the sequence; (3) "stranded" set to "true" since the forward primer was always at the 5’ end of the sequence; (4) "limit" set to false to process all reads; (5) "merge_program" set to flash; and (6) "ignore_j" set to "true" to ignore whether a joining (J) gene had been assigned to an inferred IGKV or IGLV gene. We used IGKV and IGKJ mouse sequences downloaded from the ImMunoGeneTics Information System (IMGT) (downloaded August 2021) as the starting database for IGKV inference.

Germline IGLV sequences were manually inferred using our previously established procedure(19). Briefly, IGL sequences were processed using the Immcantation Pipeline(21, 22) with the IMGT IGL gene database serving as the starting IgBLAST(23) database for germline gene/allele assignment. First, IGL primer sequences (IGLV1 5’-AGCTCTTCAGAGGAAGGTGG-3’; IGLC_var1 5’-AGCTCTTCAGGGGAAGGTGG-3’; IGLC2 5’-AGCTCCTCAGAGGAAGGTGG-3’; IGLC3 5’-AGCTCCTCAGGGGAAGGTGG-3’) were identified using maskPrimers align. Since primer sequences were not provided with the SMARTer Mouse BCR Profiling Kit, we manually determined the primer sequences by performing a multiple sequence alignment of the first 30 base pairs of the Illumina R1 reads. Next, read pairs were assembled using assemblePairs align, then duplicate reads were collapsed using collapseSeq, with the duplicate count of each collapsed sequence recorded as "dupcount". Downstream processing required that all sequences have a dupcount ≥ 2. Initial assignments to germline IGLV and IGLJ genes were performed using IgBLAST, with the resulting output parsed with Change-O MakeDb. Clones were identified by defineClones, with the clonal thresholds determined independently for each strain using distToNearest function in SHazaM(22, 24). Next, clones were clustered based on IGLV gene assignment and the percent identity to the nearest mm10 reference sequence. Finally, we determined consensus IGLV gene sequences using CD-HIT (cd-hit-est v4.6.8)(25), requiring that a given cluster sequence be represented by at least 0.1% of the total number of clones identified per strain. This process was repeated across all strains to create a unique inferred germline IGLV gene set for each strain.

We validated our inferred IGKV and IGLV germline sequences using TIgGER(22, 26). Briefly, Presto-processed reads were input into IgBlast(23) and Change-O(22), with our inferred germline IGKV and IGLV sequences as the starting gene database for IgBLAST(23) alignment. Upon generation of a Change-O table for each strain, the inferGenotype function of TIgGER was performed, and sequences were considered validated if they were successfully identified.

### IGKJ and IGLJ Germline Gene Inference

J genes were analyzed independently of V genes for IGK and IGL germline repertoires. First, IgBLAST(23) was run on all sequences using the IMGT germline gene database (release 202209-1; 28 February 2022) for IGK and IGL, and we passed the resulting IgBLAST output tables to Change-O. Then, we manually inspected Change-O tables for frequently occurring sequences in the J calls that were not exact sub-sequences of the IMGT reference J alleles.

Sequences present in a strain at a rate > 1% were added to a new J gene reference database that included novel J sequences and IMGT database J sequences. We then re-ran IgBLAST and Change-O’s MakeDb function with the new reference J gene set and inspected MakeDb output to ensure that no valid reads failed the pipeline due to missing J reference alleles. Lastly, we validated our candidate novel J alleles using OGRDBstats(27).

### Database And Inter-Strain Comparisons of Germline Gene Sets

We compared our inferred germline sequences to the existing IMGT gene database using IMGT HighV-QUEST v1.8.3 (7 May 2021)(28, 29). We also compared inferred IGKV and IGLV germline gene sets between strains in a pairwise fashion using BLAT(30, 31). For each inter-strain comparison, the germline set of the strain with the smallest number of inferred sequences was used as the ”query”. For each sequence in the query set, the best match from the alternate strain was assigned based on percent identity and match alignment length, requiring a minimum alignment length of 275 bp. The mean sequence identity of the best matches for all sequences in the query set was computed and used to express the average similarity of sequences between two strains. The mean similarity between inferred sequences across all strains in relation to their predicted SNP haplotypes was visualized using the Pheatmap(32) package in R.

### SNP-Predicted Haplotype and Sub-Species Origin Analysis

Each strain’s SNP-predicted haplotype and sub-species origin were determined using whole-genome SNP data from the Mouse Phylogeny Viewer(33). Data were viewed and downloaded for mouse IGK and IGL loci (IGK, chr6:67449994-70709994; IGL, chr16:19055093-19265093) to determine SNP-predicted haplotypes and sub-species origin. We performed a multiple sequence alignment of genotypes at SNP positions spanning the IGK and IGL loci to generate a SNP-predicted haplotype for each strain. We then used the multiple sequence alignment to construct a neighbor-joining phylogenetic tree to cluster strains into different shared haplotypes (Figure 1). To ensure that the SNP-predicted haplotypes were assigned accurately, we required genotypes to be present for all strains at all positions for which there was SNP data. For example, if the SNP array produced an N for a position in a given strain, then the position was masked across all strains in the multiple sequence alignment to prevent the Ns from contributing to the topology of the neighbor-joining phylogenetic tree, and thus biasing the haplotype groupings. In total, we masked 23% (74/324) of SNP positions for IGK and 39% (12/31) for IGL. Haplotype groups were assigned to strains that clustered together in the phylogenetic trees, with groupings annotated in Table 1 and Figure 1.

**Figure 1.**
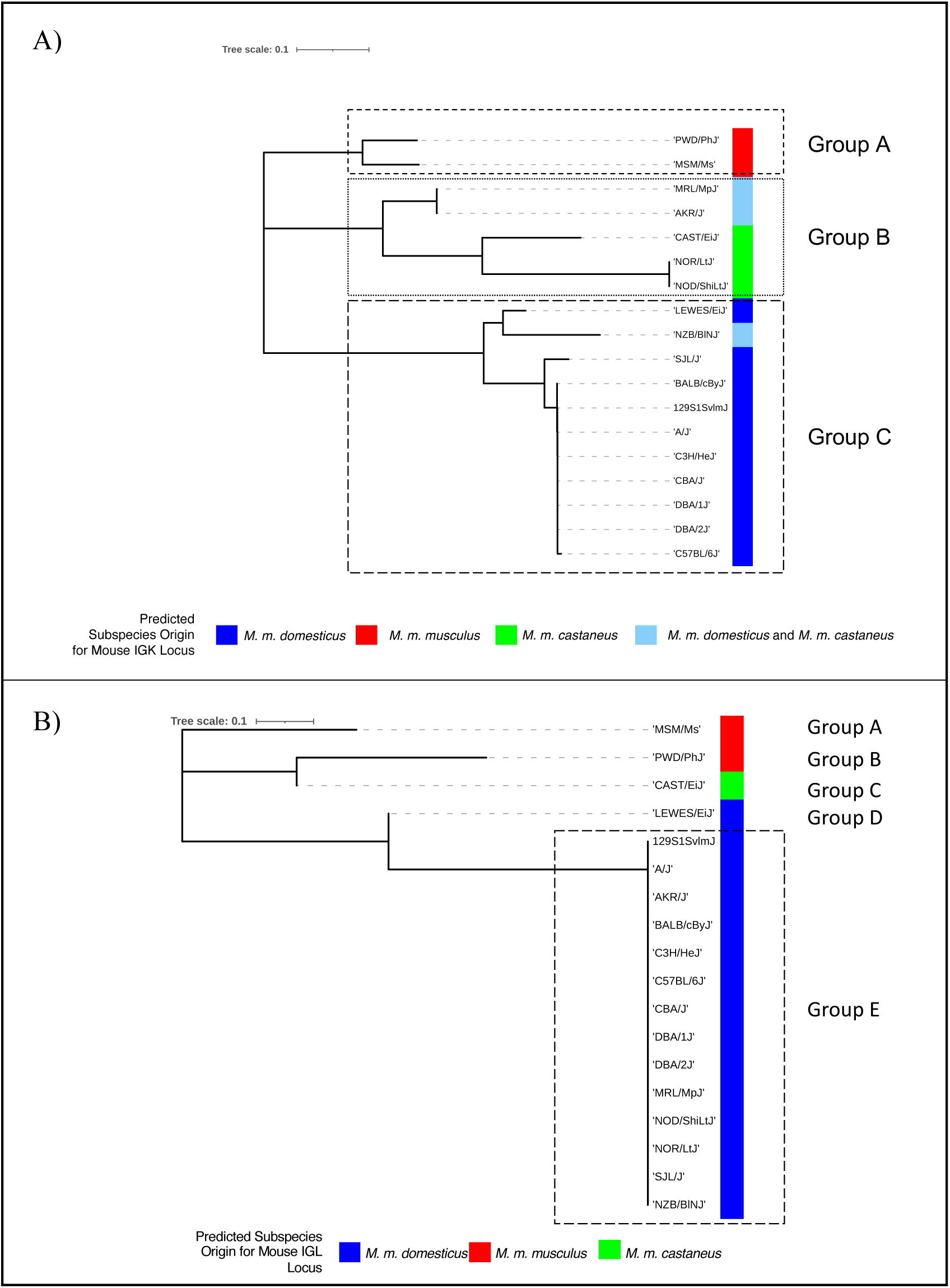
IGK and IGL haplotype phylogenetic tree generated from SNP data spanning the IGK and IGL loci. (A) IGK SNP-predicted phylogenetic tree. Color next to strains reflects the predicted subspecies origin for each strain’s IGK locus. Boxed clades represent three potential shared IGK haplotypes. Group A (PWD/PhJ, MSM/MsJ), Group B (MRL/MpJ, AKR/J, CAST/EiJ, NOR/LtJ, NOD/ShiLtJ), and Group C (LEWES/EiJ, NZB/BINJ, SJL/J, BALB/cByJ, 129S1/SvlmJ, A/J, C3H/HeJ, CBA/J, DBA/1J, DBA/2J, C57BL/6J). (B) IGL SNP-predicted phylogenetic tree. Color next to strains reflects the predicted subspecies origin for each strain’s IGL locus. Boxed strains (Group E) represent a potentially shared IGL haplotype.

### Evolutionary Analysis of IGH, IGK, and IGL V Genes in Wild-Derived Mouse Strains

For each strain and locus, the divergence of V genes was analyzed. First, V genes corresponding to the same strain and locus were translated into amino acids, and sequences translated with stop codons were discarded. Then, for each V gene, pairwise alignments against all other V genes were computed and the average percent identity was computed. For the three loci, phylogenetic trees were computed on amino acid sequences of V genes using the Clustal Omega tool(34). To analyze evolutionary relations between four wild-derived mouse strains, subtrees of the IGHV, IGKV, and IGLV trees with height at most 0.1*L* were extracted, where *L* is the height of the tree. Each subtree extracted this way represented a group of V genes from the same single family. For each subtree, the consensus sequence of corresponding genes was derived, and, for each V gene from the subtree, its divergence from the consensus was computed. The divergence was defined as the fraction of non-matching positions in the alignment between the gene and the consensus.

## Results

### Selecting Mouse Strains to Represent Diverse Sub-Species Origins and IGK/L Haplotypes

The mouse has been used in genetic studies, random mutagenesis experiments, the development of inbred lines, and the direct engineering of the genome through knock-in, knockout, and transgenic techniques for over one hundred years(35). As a result, many mouse strains are available for use in biomedical research. For example, C57BL/6, the most common and best-studied classical laboratory strain today(36), has been a popular model organism in immunology, with various knockout lines available and its genome sequenced by the Mouse Genome Sequencing Consortium(35). Another common strain, BALB/c, served as an early model organism used to induce plasmacytomas and monoclonal antibody production(37, 38). Other strains, such as NOD/ShiLtJ, SJL/J, and MRL/MpJ, are used to model autoimmune disorders such as autoimmune type 1 diabetes, experimental autoimmune encephalomyelitis, and systemic lupus erythematosus and Sjogren’s syndrome(39, 40). In addition, wild-derived mouse strains are often used to incorporate wild mouse genetics into laboratory strains by creating F1 hybrids(41). Given the diverse breeding history of these strains(42–45) we expected the genetic diversity of the IG light chains to resemble the diversity observed in the IG heavy chain(19, 46).

To select strains that captured IG light chain germline diversity, we examined the predicted sub-species origins and SNP-based haplotypes of the mouse IG light chain loci(17, 18) (Table 1, Figure 1) across classical and wild-derived strains. We leveraged early studies that reported the existence of alternate IGK and IGL haplotypes across mouse strains(13, 47–50), as well as genome-wide SNP data available for 62 wild-derived laboratory strains and 100 classical strains(18). To account for the different *Mus* subspecies, we included strains with IG loci predicted to represent the three major *Mus* subspecies, *M. castaneus, M. domesticus, and M. musculus,* which form the genetic background for classical inbred laboratory mouse strains(51, 52). In addition, we chose CAST/EiJ, LEWES/EiJ, MSM/MsJ, and PWD/PhJ to represent wild-derived mouse strains, in which we have previously inferred germline IGHV, IGHD, and IGHJ genes(19). In total, we sequenced the IGK and IGL repertoires of 18 different mouse strains representing three SNP-predicted IGK haplotype groups and five SNP-predicted IGL haplotype groups(33) (Figure 1).

### Mouse Light Chain Variable Genes Are Underrepresented In Germline Gene Databases

First, we inferred germline light chain repertoires across our selected mouse strains (Figure 2). We used IgDiscover to infer each strain’s IGKV germline sequences and our previous clustering method(19) to infer IGLV germline sequences. To benchmark the performance of our inference approach, we first assessed how well our C57BL/6J inferences compared to known IMGT C57BL/6 IGKV and IGLV germline sequences. 91/91 IGKV inferred sequences matched IMGT C57BL/6 IGKV germline sequences with 100% identity. Additionally, our three C57BL/6J IGLV inferences were 100% identical to IMGT IGLV sequences. However, since the three IMGT IGLV sequences that matched our inferences were derived from BALB/c(5), we also compared these three inferences to the mouse reference genome, mm10, derived from C57BL/6. The three IGLV C57BL6/J inferences matched mm10 with 100% identity.

**Figure 2.**
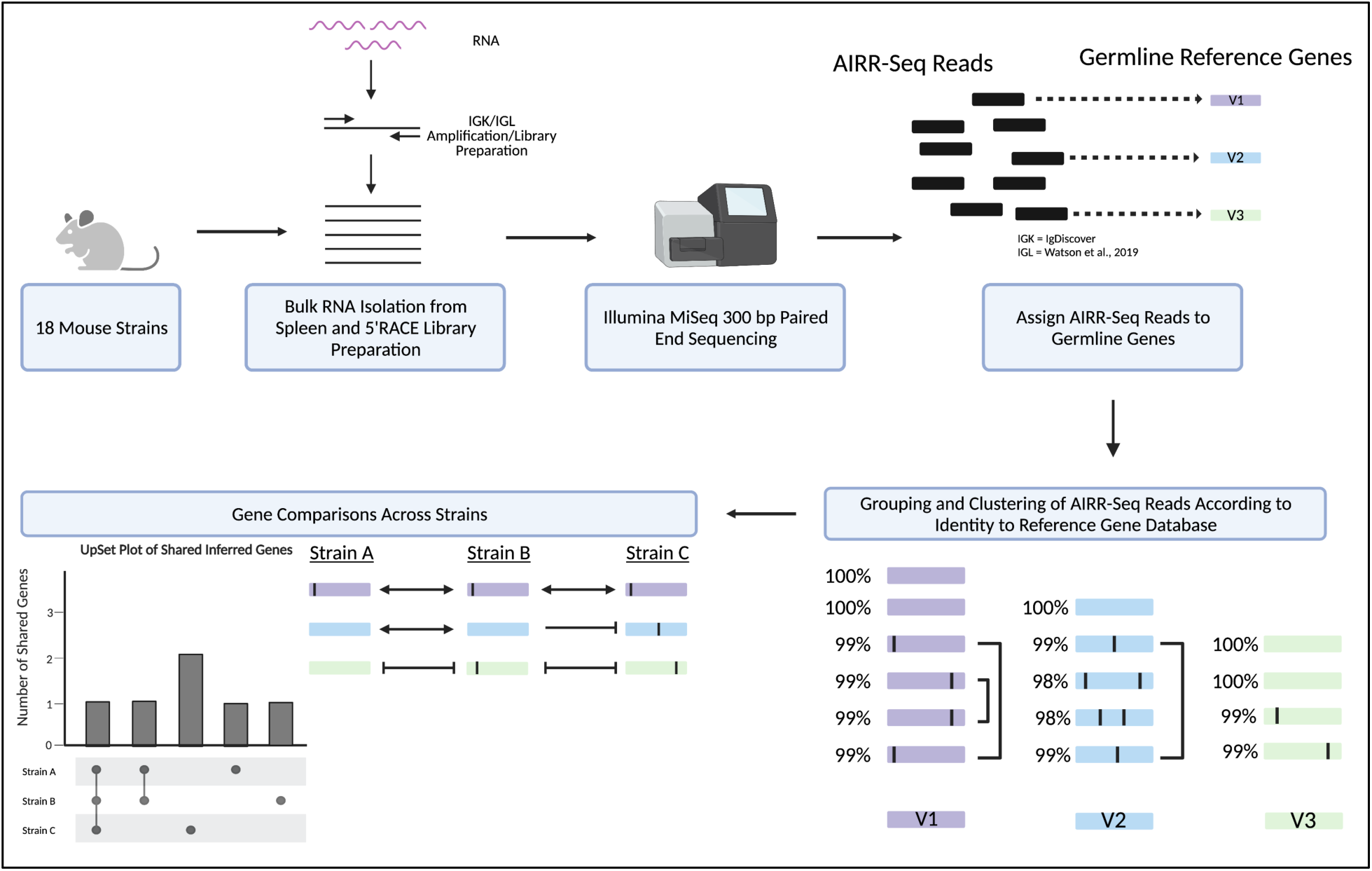
Experimental overview for IGKV/IGLV germline gene inference. Briefly, RNA was extracted from 18 different inbred mouse strains, then 5’RACE was used to generate IGK and IGL sequencing libraries. Next, libraries were sequenced on an Illumina MiSeq instrument (2x300 bp PE Sequencing), then the AIRR-seq reads were assigned to germline genes using either IgDiscover (IGKV) or our in-house pipeline (IGLV). After gene assignment, AIRR-seq reads were grouped and clustered according to their identity to genes in the IMGT reference gene database. Lastly, we visualized how the light chain germline gene repertoires were shared across strains using UpSet plots. Created with BioRender.com.

Across the 18 mouse strains, 1582 IGKV and 63 IGLV sequences were inferred, representing 459 and 22 unique IGKV and IGLV sequences, respectively (Supplemental Table 1). The sizes of inferred IGKV germline gene sets varied across strains, from 105 in NZB/BlNJ to 62 in NOD/ShiLtJ (Figure 3A). In contrast, the numbers of inferred IGLV germline genes were more conserved across strains. Three IGLV germline sequences were inferred for all classical laboratory strains and PWD/PhJ. However, LEWES/EiJ, MSM/MsJ, and CAST/EiJ had > three genes inferred from their repertoire data (Figure 3B). Inferred IGKV and IGLV germline sequences for all strains are available on OGRDB(53).

**Figure 3.**
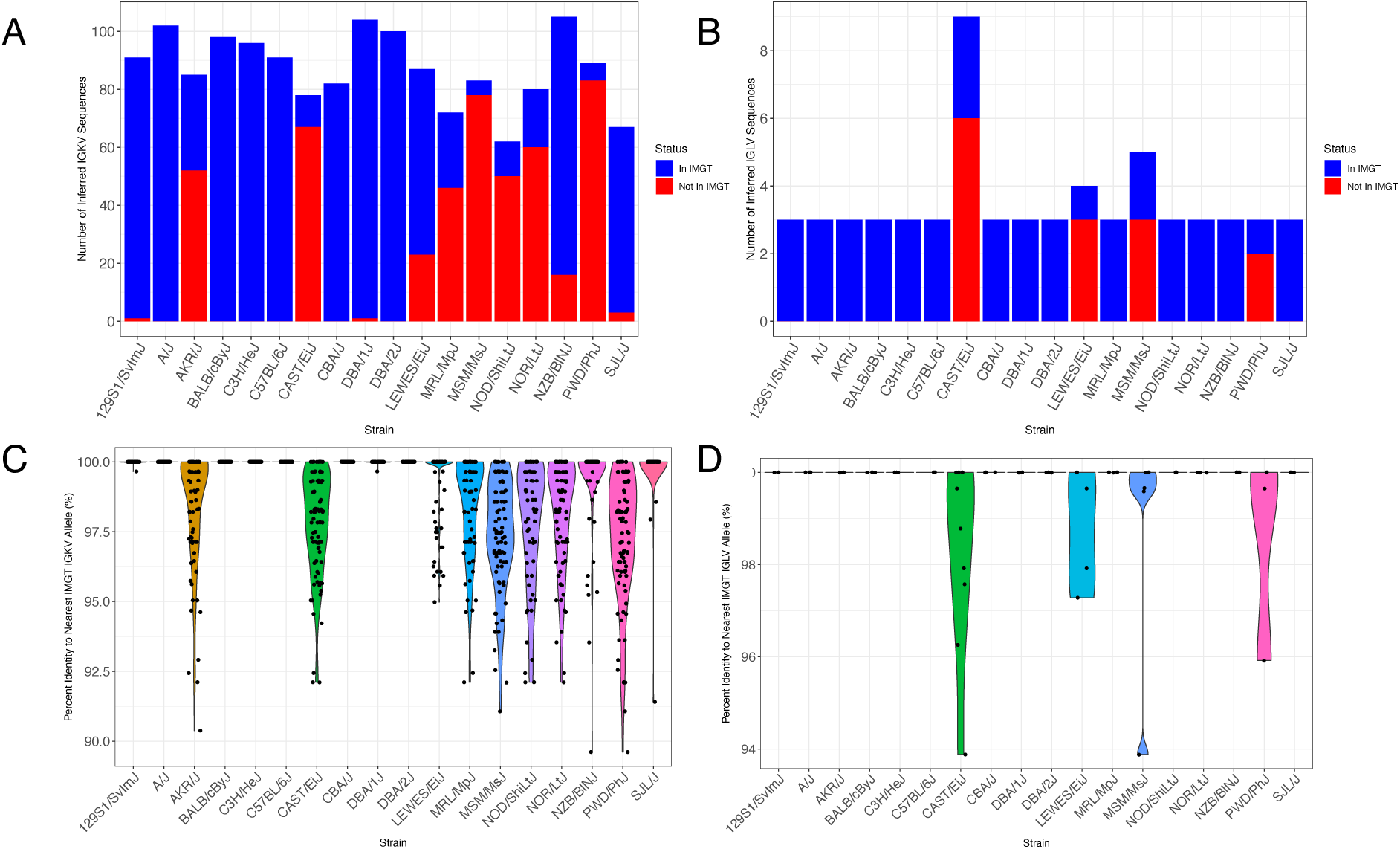
Representation of inferred IGKV and IGLV germline sequences in IMGT gene database. (A and B) Bar plots depicting IGKV (A) and IGLV (B) inference counts for each strain and whether the inferences were present or absent in the IMGT Gene Database. (C and D) Violin plots depicting sequence alignment percent identity of IGKV (C) and IGLV (D) inferences to the IMGT Gene Database.

Of the 459 and 22 IGKV and IGLV unique sequences inferred across strains, 67.8% (n=311, IGKV) and 59% (n=13, IGLV) were undocumented in IMGT (Figures 4A, B). A fraction of these non-IMGT alleles were identified in NCBI GenBank with 100% identity: 12% (n=37) of IGKV and 8% (n=1) of IGLV (Figures 4C, D). The number of undocumented (non-IMGT) alleles varied by strain (Figures 3A, B). For IGKV, while a significant fraction of non-IMGT alleles were inferred from wild-derived strains, there were several biomedically relevant classical strains in which the majority of sequences are not curated in IMGT (Figure 3A), likely reflecting divergence from the C57BL/6 haplotype, as has been previously noted(14). Strains such as NOD/ShiLtJ, NOR/LtJ, AKR/J, and MRL/MpJ, commonly used to model autoimmune disorders, have poor IGKV germline representation in IMGT (Figure 3A)(39, 54, 55). Sequence alignment of these strains’ inferred IGKV germline sequences to the IMGT database yielded percent identities ranging from 100% to 89.61% for IGKV, and 100% to 93.88% for IGLV (Figures 3C, D). Alignment percent identity was strain-dependent, as we observed significant IGKV sequence variation for the four wild-derived strains, and AKR/J, MRL/MpJ, NOD/ShiLtJ, NOR/LtJ, and NZB/BlNJ. Of all the strains investigated, MSM/MsJ have the fewest IGKV germline sequences (5/83) documented in IMGT, and CAST/EiJ have the fewest IGLV germline sequences (3/9) documented in IMGT. Collectively, we observe high levels of diversity currently unaccounted for in the IMGT gene database.

**Figure 4.**
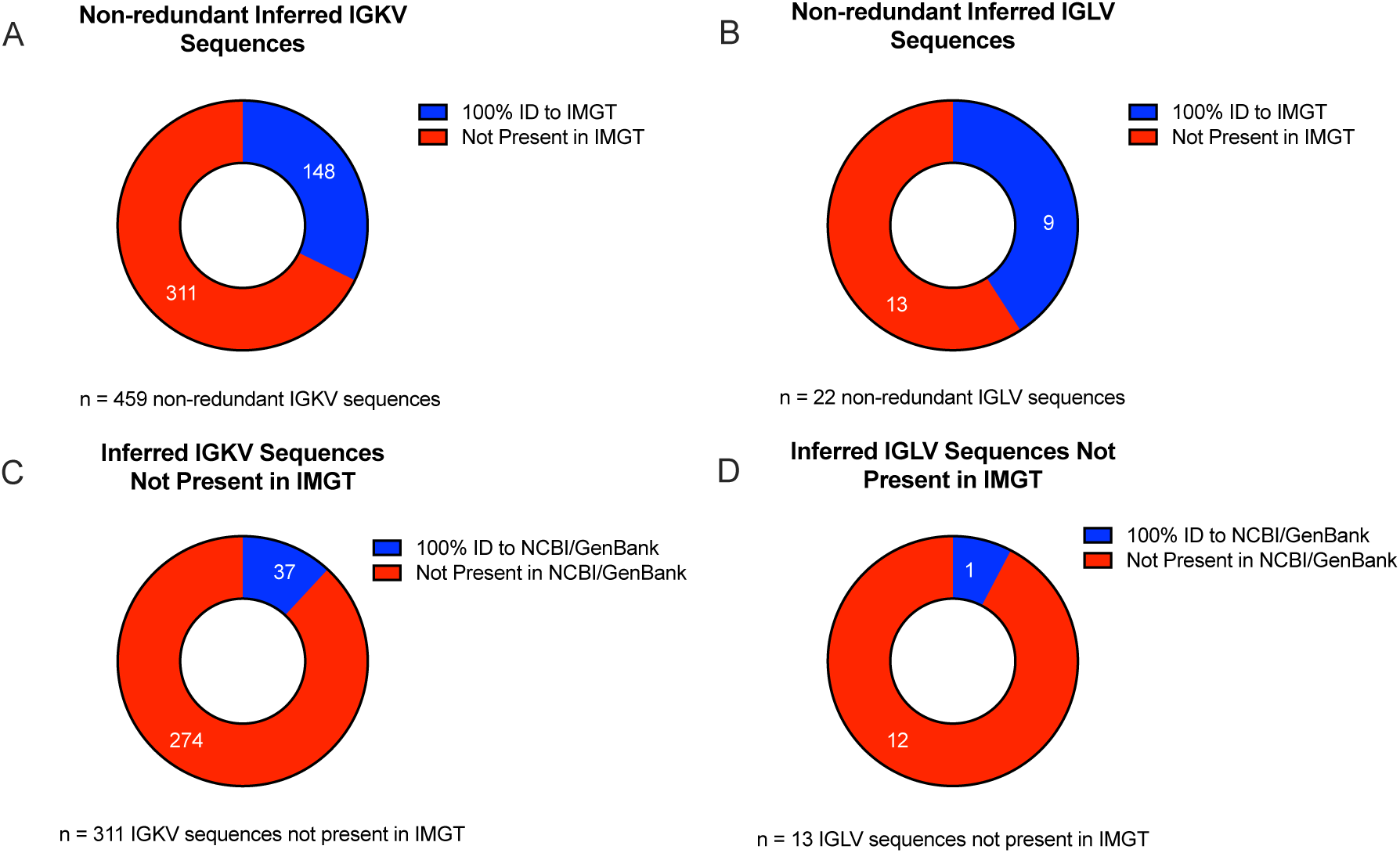
Presence and absence of non-redundant IGKV and IGLV inferred sequences in existing gene databases. (A and B) Non-redundant IGKV (A) and IGLV (B) sequences present/absent in IMGT gene database. (C and D) Donut plots depicting the 311 IGKV (C) and 13 IGLV (D) inferences missing from IMGT and their presence/absence in NCBI/GenBank.

IGKJ germline inference across strains revealed a novel IGKJ2 allele, with a single T to C transition shared between PWD/PhJ and MSM/MsJ (Figure 5A), strains of *M. m. musculus* subspecific origin, and members of Group A in our IGK SNP-haplotype phylogeny (Figure 1A). We inferred two novel IGLJ alleles among the wild-derived strains (Figure 5B). A novel IGLJ1 allele was inferred for LEWES/EiJ, and a novel IGLJ2 allele was inferred for MSM/MsJ. Both novel IGLJ alleles differ from the IMGT reference sequence by a single nucleotide. Interestingly, we did not infer any IGLJ3 alleles in MSM/MsJ; all MSM/MsJ J gene usage was restricted to IGLJ1*01 and the novel IGLJ2 allele. Overall, we found our SNP-haplotype groupings reflected in the IGKJ and IGLJ allelic variation across strains.

**Figure 5.**
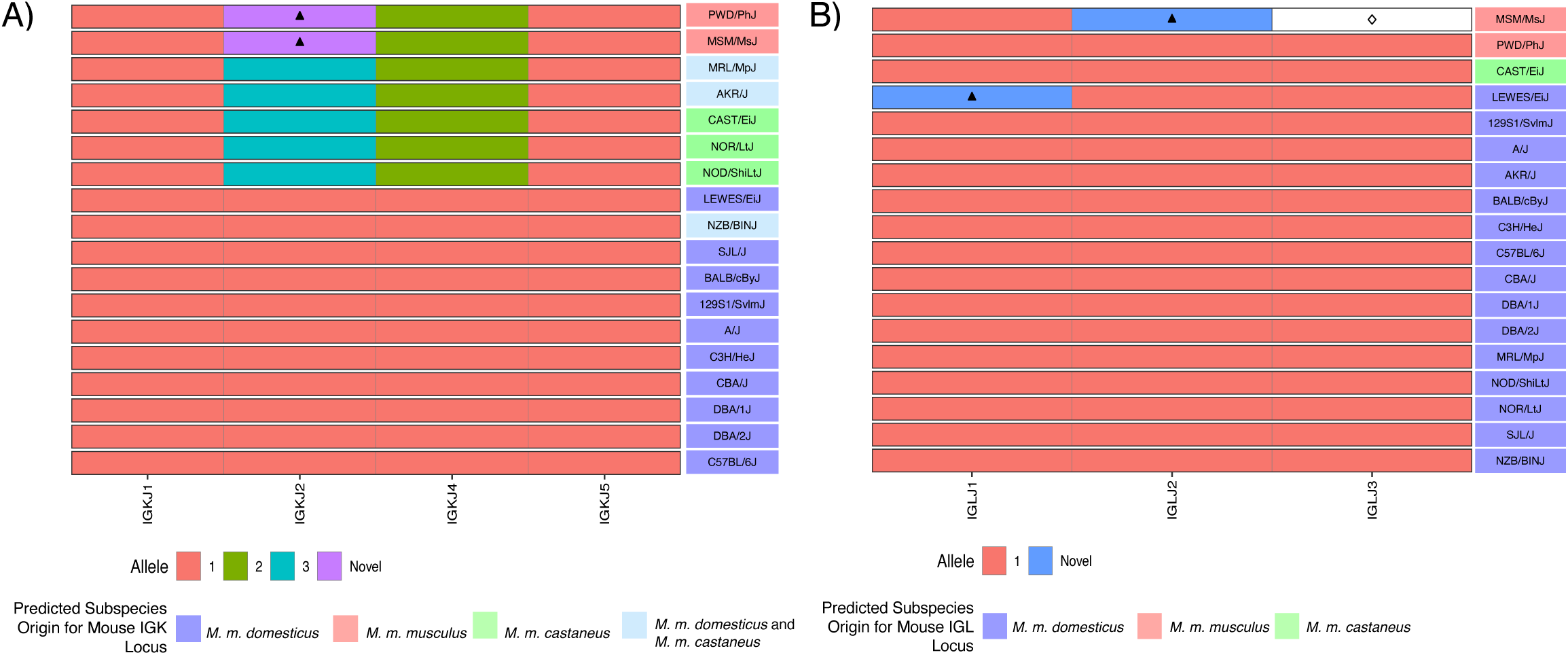
Inferred IGKJ and IGLJ novel alleles. A) A single IGKJ2 novel allele was inferred for PWD/PhJ and MSM/MsJ (5’-TGTA**C**ACGTTCGGATCGGGGACCAAGCTGGAAATAAAAC-3’). B) Two novel alleles were inferred for IGLJ: Novel IGLJ1 allele (5’-CTGGGTGTTCGGTGGAGGAACCAAA**T**TGACTGTCCTAG-3’) and novel IGLJ2 (5’-TTATGTTTTCGGC**A**GTGGAACCAAGGTCACTGTCCTAG-3’). Bolded positions are novel SNPs that diverge from IMGT reference sequence.

### Inter-Strain IGKV/IGLV Germline Diversity

We next considered the extent to which inferred IGKV and IGLV germline sequences were shared among strains (Figures 6A, B). Across IGKV germline gene sets, the most strain-specific sequences were observed among the wild-derived strains, CAST/EiJ (n=58), PWD/PhJ (n=57), MSM/MsJ (n=55), and LEWES/EiJ (n=21), with an additional 19 sequences uniquely common to PWD/PhJ and MSM/MsJ (Figure 6A). There were fewer unique sequences in each of the classical laboratory strains. For example, only nine unique sequences were seen in the NOD/ShiLtJ strain, which had the highest number of unique sequences amongst the classical inbred strains. Instead, we observed large sets of sequences that were identical across many strains (Figure 6A). The most extensive shared sequence set comprised 27 inferred IGKV germline sequences inferred from 11 different strains (SJL/J, CBA/J, LEWES/EiJ, C57BL/6J, 129S1/SvlmJ, C3H/HeJ, BALB/cByJ, DBA1/J, DBA2/J, A/J, and NZB/BlNJ). This degree of allele sharing was suggestive of the presence of potentially shared haplotypes. We assessed our IGK SNP-Predicted haplotype phylogenetic tree (Figure 1A) and found all 11 strains in Group C. NZB/BINJ was the only strain in Group C predicted to have a different sub-specific origin for the IGK locus. The Mouse Phylogeny Viewer(33) reports a *M. m. domesticus* and *M. m. castaneus* sub-specific origin for the NZB/BINJ IGK locus, which contrasts the other strains’ *M. m. domesticus* IGK locus sub-specific origin.

**Figure 6.**
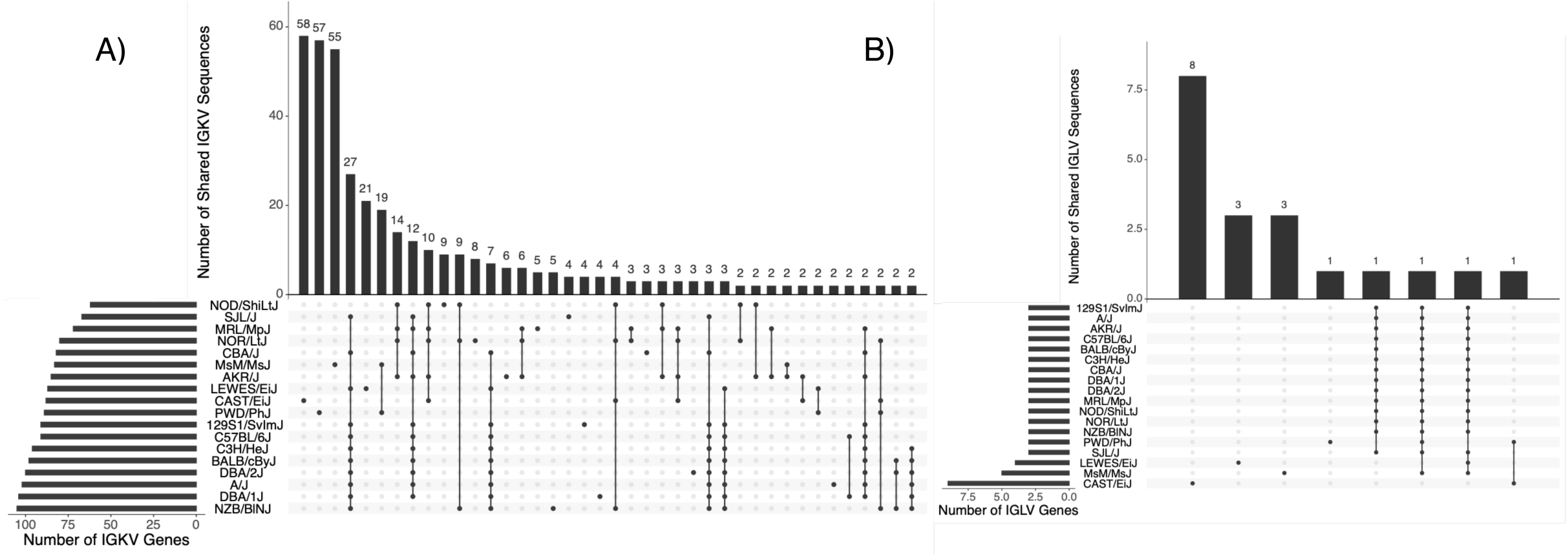
Shared and unique IGKV and IGLV germline repertoires. UpSet plots depicting the size of the germline IGKV (A) and IGLV (B) set from 18 mouse strains and the number of sequences that were unique to a given strain (dot) or shared among strains (connected dots).

Another group of strains, NOD/ShiLtJ, NOR/LtJ, MRL/MpJ, and AKR/J, formed a different cluster with 14 shared IGKV germline sequences (Figure 6A). These four strains fall into Group B of our SNP-Predicted haplotype phylogenetic tree (Figure 1A) and represent an IGK locus sub-specific origin of either completely *M. m. castaneus*, or a mixture of *M .m. castaneus* and *M. m. domesticus*. Finally, PWD/PhJ and MSM/MsJ, both wild-derived strains with a *M. m. musculus* sub-specific origin, were located in the Group A cluster, with 19 unique IGKV sequences shared between themselves (Figure 6A, Figure 1A). We also examined inter-strain diversity at the level of IGKV gene family by comparing the number of inferred IGKV sequences for each gene family across strains (Supplemental Figure 1). Overall, the IGKV4 subfamily was most variable in size and the most abundant family across strains, whereas IGKV20 was the least abundant and only inferred in PWD/PhJ, MSM/MsJ, and CAST/EiJ.

In each of the 14 classical strains, a total of three IGLV sequences were inferred, which is consistent with the number of genes found in the C57BL/6 mm10 genome(56). These IGLV inferences were identical across the 14 classical strains (Figure 6B), supporting the predicted sub-species origins and SNP-Haplotype phylogenetic tree clustering (Figure 1B). In contrast to the classical strains, additional putative genes were inferred in three of the wild-derived strains, CAST/EiJ, LEWES/EiJ, and MSM/MsJ, totaling nine, four, and five inferred genes for each strain, respectively. Phylogenetic analysis of inferred IGLV sequences revealed that inferred genes unique to CAST/EiJ formed an additional outgroup to all other IGLV1, IGLV2, and IGLV3 gene sequences characterized in the classical strains (Supplemental Figure 2). In addition to these IGLV paralogs, the IGLV2 sequences inferred in CAST/EiJ and PWD/PhJ, and the IGLV3 sequences inferred from CAST/EiJ, PWD/PhJ, and LEWES/EiJ, differed from those characterized in the classical strains, likely representing allelic variants. Thus, taken together, the majority of IGLV germline diversity observed in the animals studied here came from wild-derived strains.

To help validate the relationship between predicted IG haplotypes, sub-species origin, and the inferred germline sets across strains, we performed all-by-all pairwise comparisons of inferred IGKV and IGLV sequences across strains to group strains by sequence similarity. We reasoned that inferred IGKV/IGLV sequences within strains sharing predicted haplotypes would have higher sequence similarities. We found that mean pairwise sequence similarities varied considerably. For IGKV, the highest similarity (99.99%) observed was between C3H/HeJ and A/J, whereas the most divergent comparison was between NOR/LtJ and PWD/PhJ (95.30%). We used hierarchical clustering to group strains based on mean pairwise similarities, and the results corresponded to the three haplotype groups obtained from our SNP-Haplotype phylogenetic tree (Figure 7). IGLV inferences were much more conserved across strains. The four IGLV germline sequences inferred for LEWES/EiJ are consistent with Potter et al., who reported that wild *Mus musculus domesticus* had at least three IGLV genes(57).

**Figure 7.**
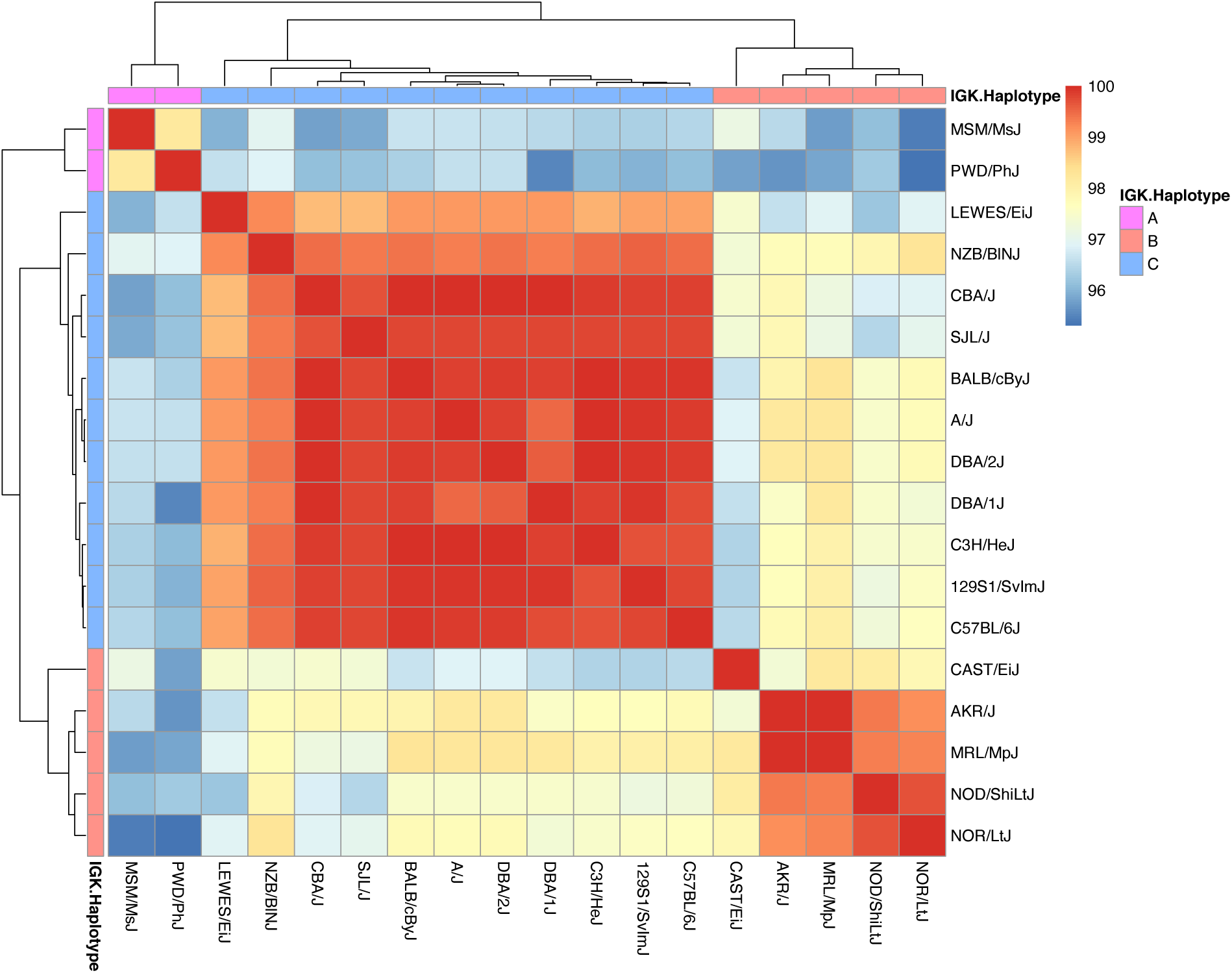
All-by-all pairwise comparisons of inferred IGKV germline genes show evidence of strain clustering according to the assigned IGK haplotype group. Heatmap depicting the mean percentage sequence match identities among inferred IGKV germline sets for each pairwise strain comparison.

### Diversification of IGHV, IGKV, and IGLV Sequences Among Wild-Derived Mouse Strains

Gene evolution through duplication and diversification events have helped shape the diversity of the mouse immunoglobulin genes. In mice and humans, light chain gene rearrangement begins on the kappa chain, but it follows heavy chain rearrangement. If this initial rearrangement is auto-reactive, then the gene organization of the kappa chain locus permits additional gene rearrangements through a process known as receptor editing to form a rearrangement that is not auto-reactive(3). While both the heavy and kappa chain repertoires have many V genes that have evolved through gene duplications, deletions, and sequence divergence(6, 58), the kappa chain repertoire contains less inherent germline diversity than the heavy chain(3). It has been hypothesized that the reduced germline diversity of the kappa chain repertoire has evolved to limit self-reactivity, while the heavy chain repertoire has evolved to increase diversity. Support for this hypothesis stems mainly from human AIRR-seq data, in which there is some evidence that there is less allelic diversity in IGKV compared to IGHV(3). Others have also suggested that the patterns of diversification and divergence are potentially different between heavy and light chain immunoglobulin genes. For example, Schwartz et al. analyzed amino acid sequences for human germline heavy and light chain genes and concluded that heavy and lambda V genes had higher diversities compared to kappa V genes(59). Our AIRR-seq dataset provides us with the opportunity to compare diversification and divergence patterns between germline heavy and light chain gene sets across multiple mouse strains representing diverse mouse subspecies origins. Therefore, we hypothesized that the mouse kappa chain antibody repertoire would have decreased diversity in germline V genes compared to the heavy chain antibody repertoire to minimize potential auto-reactivity.

One metric that can be used to compare germline sequence evolution of genes is the edit distance between two sequences, defined as the minimal number of mutations separating the two sequences(60). In 2019, we inferred germline IGHV genes among wild-derived mouse strains representing the major *Mus* sub-species origins(19). With the additional germline IGKV and IGLV genes from the same wild-derived strains, we compared germline variable gene sequence divergence rates for the IGH, IGK, and IGL loci of the wild-derived strains using phylogenetic trees constructed from multiple sequence alignments of germline IGHV, IGKV, and IGLV sequences. IGHV genes from all strains have lower average percent identities compared to IGKV and IGLV genes thus indicating that heavy chain V genes evolve faster compared to light chain V genes in mice (Figure 8A). Figure 8B shows that the pattern holds true for pairs V genes collected from pairs of strains. This data suggests that the mouse IGH locus contains greater inherent germline sequence diversity than the IGK locus despite having a similar number of germline genes. The resulting phylogenetic trees revealed that, as expected, V genes formed groups according to their families (Figure 8C). The only exception in the IGHV tree was IGHV12 family represented by three genes that did not form a single subtree but rather were broken into two groups, one of which was formed by genes IGHV12-2_1 (LEWES/EiJ) and IGHV12-1_1 (MSM/MsJ) and IGHV3 family and other one is formed by IGHV12-1_1 (LEWES/EiJ) gene and IGHV2 and IGHV8 families. IGKV16 family represents a similar exception in the IGKV tree: five IGKV16 genes did not form a single subtree and were found in subtrees corresponding to families IGKV11, IGKV13, IGKV15, IGKV17, IGKV19, and IGKV20. Finally, Figure 8D shows that, on average, the LEWES/EiJ strain has the highest divergence from the consensus. A similar analysis of IGKV genes revealed the same pattern (Figure 8E). These observations suggest that the LEWES/EiJ strain has branched from the common ancestor before three other strains (CAST/EiJ, MSM/MsJ, and PWD/PhJ) and accumulated more mutations in immunoglobulin V genes.

**Figure 8.**
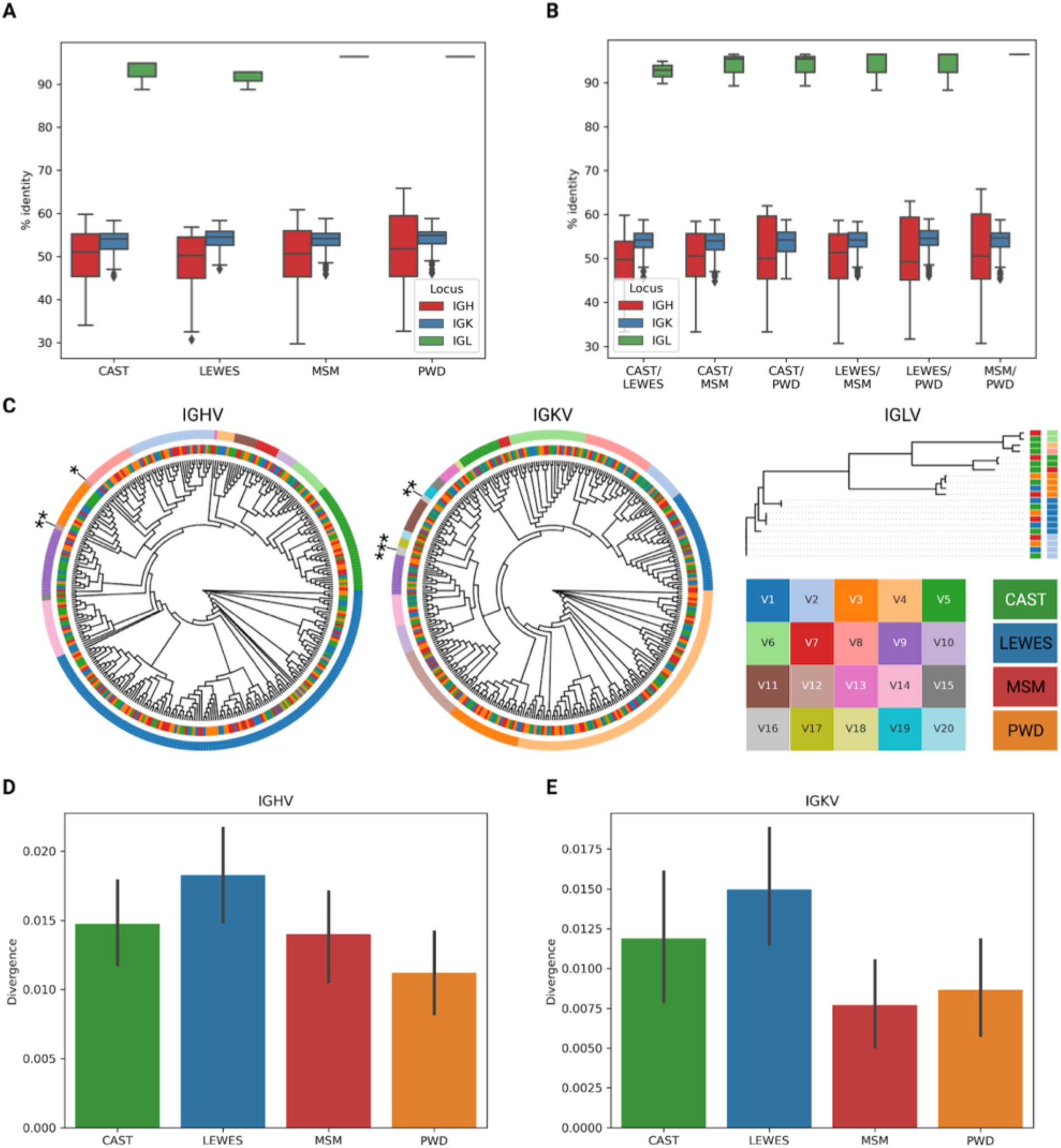
Phylogenetic analysis of mouse immunoglobulin V genes. (**A**) Percent identities of IGHV, IGKV, and IGLV genes of CAST, LEWES, MSM, and PWD mouse strains. Each bar represents a single mouse strain (CAST, LEWES, MSM, or PWD) and a locus (IGH, IGK, or IGL) and shows the distribution of the median percent identities of corresponding V genes. (**B**) Percent identities of IGHV, IGKV, and IGLV genes for all pairs of mouse strains. (**C**) Phylogenetic trees for IGHV, IGKV, and IGLV genes. Genes from families IGHV12 and IGKV16 that do not form single subtrees in the corresponding phylogenetic trees are labeled with asterisks. (**D**) The distributions of divergences of IGHV genes across mouse strains with respect to the consensuses of clusters computed as subtrees of the phylogenetic tree of IGHV genes of length 0.1*L*, where *L* is the height of the tree. (**E**) The distributions of divergences of IGKV genes computed in the same way as the distributions shown in (D).

## Discussion

This study was performed as a follow-up to our 2019 study in which we inferred IGHV, IGHD, and IGHJ genes of wild-derived strains representing each of the three major subspecies of the house mouse (CAST/EiJ: *M. m. castaneus*; LEWES/EiJ: *M. m. domesticus*; PWD/PhJ: *M. m. musculus*) and the *M. m. musculus/M. m. castaneus* hybrid strain MSM/MsJ. Overall, we found little overlap in germline IGHV repertoires among the wild-derived mouse strains and could not attribute all of the repertoire differences observed to variation in sub-specific origin. For example, while apparently *musculus-*derived strains did share IGHV sequences, many of these sequences were unique to each strain. Most importantly, the inferences were largely absent in existing gene databases such as IMGT. The diversity of germline IGHV sequences led us to hypothesize that extensive germline light chain variable genes would also be present. Therefore, in addition to sequencing the light chain variable genes for the wild-derived strains sequenced in 2019, we expanded the number of strains to include various strains used across the biomedical sciences that represented different SNP-predicted haplotypes for the light chain loci. The strains encompassed a variety of disease models in the biomedical sciences, including those used for the study of infection, autoimmunity, diabetes, cancer, and regeneration(54, 61).

One of the most fundamental tasks in analyzing AIRR-seq data is V, D, and J gene assignment, which is accomplished by performing alignment of AIRR-seq reads to germline sequences from a gene database. IMGT is the most commonly used database that curates the germline repertoires used for sequence alignment in AIRR-seq analysis. When comparing our inferred light chain germline sequences to those curated in IMGT, we observed that many strains’ IGKV germline sequences were absent from IMGT, and few undocumented sequences were found in NCBI/GenBank (Figures 3, 4). In contrast, the majority of genes characterized in IGLV among classical strains are curated in IMGT, including previously described wild-derived IGLV4 sequences detected in CAST/EiJ(62).

Critically, many of the inferred germline genes found to be absent from IMGT showed evidence of significant divergence from the closest curated allele in the database. For example, 59% (185/311) of non-IMGT IGKV sequences, and 62% (8/13) of non-IMGT IGLV sequences, had <98% identity to their nearest IMGT IGKV sequence. Thus, we would expect that these missing data could greatly impact the accuracy of germline gene assignment and SHM estimation in studies using these strains. Given that the IGK locus is similarly complex as the IGH locus, with different combinations of SNP-predicted sub-species origins and haplotypes, the IGK locus likely contains genetic variation similar to that observed in IGH between BALB/c, C57BL/6, and 129S1(11, 46). Our inferred IGKV germline sequences supported these three distinct haplotype clusters (Figures 1A, 6A, and 7). For example, 11 strains shared 27 IGKV germline sequences, and all 11 strains belonged to Group C in the IGK SNP-Predicted haplotype phylogeny (Figure 1A). We compared these strains to the historic IGK haplotypes identified using RFLP and observed that 9 of the 11 strains were previously designated the historical IGK^A^ haplotype(48). We also observed 14 IGKV sequences shared among 4 strains in our dataset belonging to Group B (Figure 1A), containing the historical IGK^B^ haplotype(48). Though CAST/EiJ does not share the 14 unique IGKV sequences with NOD/ShiLtJ, NOR/LtJ, MRL/MpJ, and AKR/J, the SNP-phylogeny data, and additional shared sequences between these strains, suggest that CAST/EiJ does indeed belong in haplotype Group B. Similar to the historic IGH^A^ and IGH^B^ haplotypes(6, 11, 63, 64), our results suggest that the IGK loci of strains carrying the IGK^B^ haplotype are similar to one another and different from the loci of strains carrying the IGK^A^ haplotype.

Strain clustering according to the sub-species origin was also apparent after performing all-by-all pairwise sequence comparisons (Figure 7) and validated our SNP-phylogeny groupings. Of note, the 5 strains in Group B (AKR/J, MRL/MpJ, NOD/ShiLtJ, NOR/LtJ, and CAST/EiJ) exhibited significant sequence divergence from the IMGT alleles (Figure 3D), highlighting the lack of representation for strains sharing this IGK haplotype. However, like the IGH locus, only a single complete IGK reference is available based on C57BL/6(10), illustrating an IGK haplotype not shared by all strains and only representative of a single sub-species origin.

Our cohort contained 5 SNP-predicted IGL haplotypes (Figure 1B); however, only one haplotype had representation by more than one strain. All wild-derived strains, MSM/MsJ, PWD/PhJ, CAST/EiJ, and LEWES/EiJ were clustered into single-strain clades according to the SNP-predicted haplotype, while the 14 remaining classical laboratory strains clustered into a single haplotype. Our data supported the predicted haplotype clustering, with each wild-derived strain containing at least one unique IGLV allele not shared with other strains and classical laboratory strains sharing IGLV alleles amongst each other (Figure 6B). Of note, we did infer more than the three canonical IGLV germline sequences in CAST/EiJ, MSM/MsJ, and LEWES/EiJ, which was expected based on previous data hypothesizing the existence of additional IGLV genes in wild-derived strains(57). A total of 13 novel sequences were identified among the wild-derived strains, with an allelic variant of IGLV2*02 shared between CAST/EiJ and PWD/PhJ. Inferred IGLV germline repertoires in classical laboratory strains were more conserved and composed of three IGLV genes.

Although expressed lambda chain genes account for only 3 to 5% of total serum IG, little is known regarding how the mouse lambda locus reduced in size(57). Furthermore, though gene deletion events could cause a reduced IG lambda locus in mice, it is unclear where it occurred in mouse phylogenetic history. The rat IG loci, similar to the mouse IG loci, have not been extensively explored and characterized across strains. Reports suggest that the IGL locus in rat only contains a single IGLV gene and two IGLC genes making the rat IGL locus even smaller than mouse(65). If we consider the rat and mouse to be distant relatives, this presents two possible scenarios that may have occurred during rat and mouse evolution. Either the mouse IGL locus expanded through duplications, or the rat IGL locus decreased in size through deletions.

Overall, in conjunction with other published studies, the data presented here demonstrate that the germline heavy and light chain repertoires are not conserved across biomedically relevant mouse strains(19, 46). Our phylogenetic analysis of heavy and kappa germline inferences for wild-derived strains revealed sequence divergence differences among the four major *Mus* sub-species origins. Furthermore, it showed that the heavy chain locus contained more inherent germline sequence variability than the kappa chain locus, which had previously been hypothesized but not explored across multiple strains representing different subspecies origins. If the function of the kappa chain locus is to limit inherent germline diversity to prevent B-cell receptor auto-reactivity as hypothesized, then it is crucial to properly characterize IGK haplotypes for mouse strains used in autoimmune research. In humans, we know that the light chain repertoire in autoimmune diseases like Myasthenia Gravis can become perturbed and result in errors during receptor editing during B-cell development(66). Although AIRR-seq data can help build germline gene databases for various mouse strains, this data lacks critical information on noncoding elements and gene positions that could elucidate gene expression differences between strains. Additional IG genome assemblies are required that reflect the haplotype and sub-specific diversity that we have presented in the mouse IG loci.

## Notes

### Competing Interest Statement

The authors have declared no competing interest.

