## Supplemental Material for "Characterization of Extensive Diversity In Immunoglobulin Light Chain Variable Germline Genes Across Biomedically Important Mouse Strains"

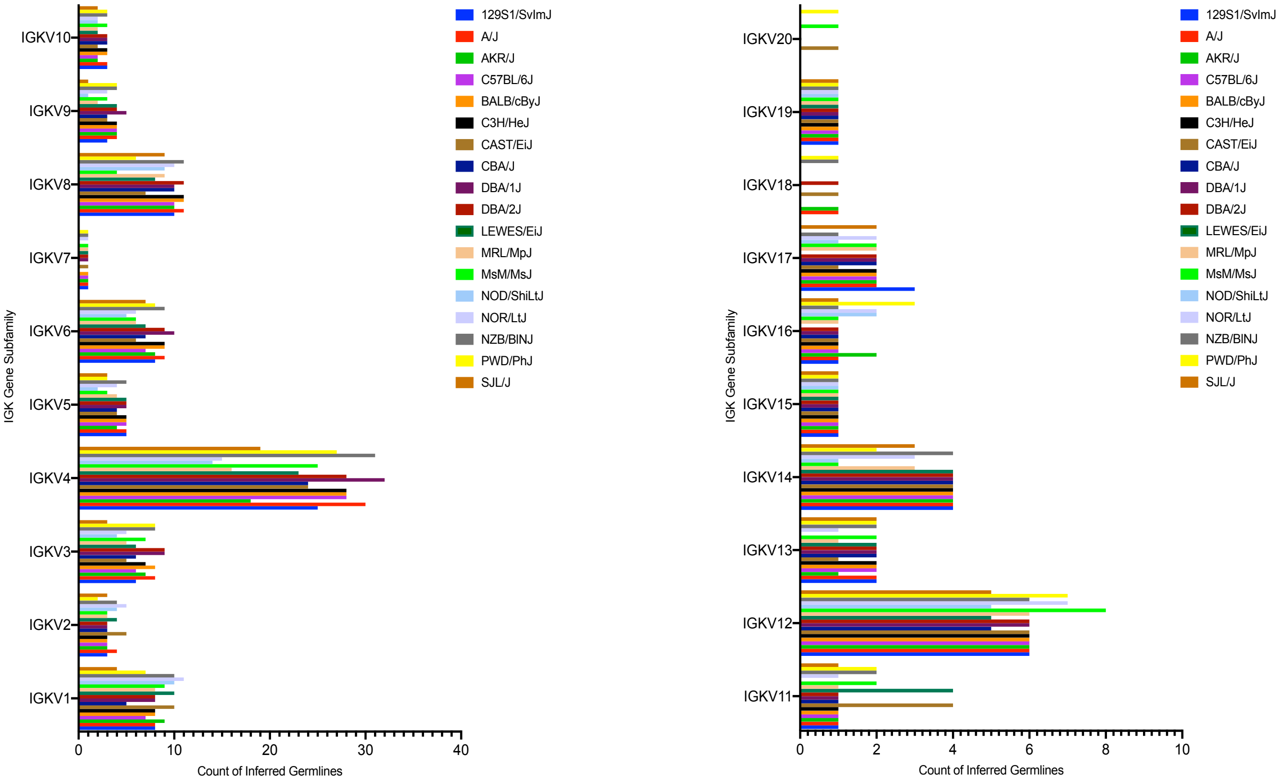


**Supplemental Figure 1. IGKV gene subfamily comparisons across strains.**

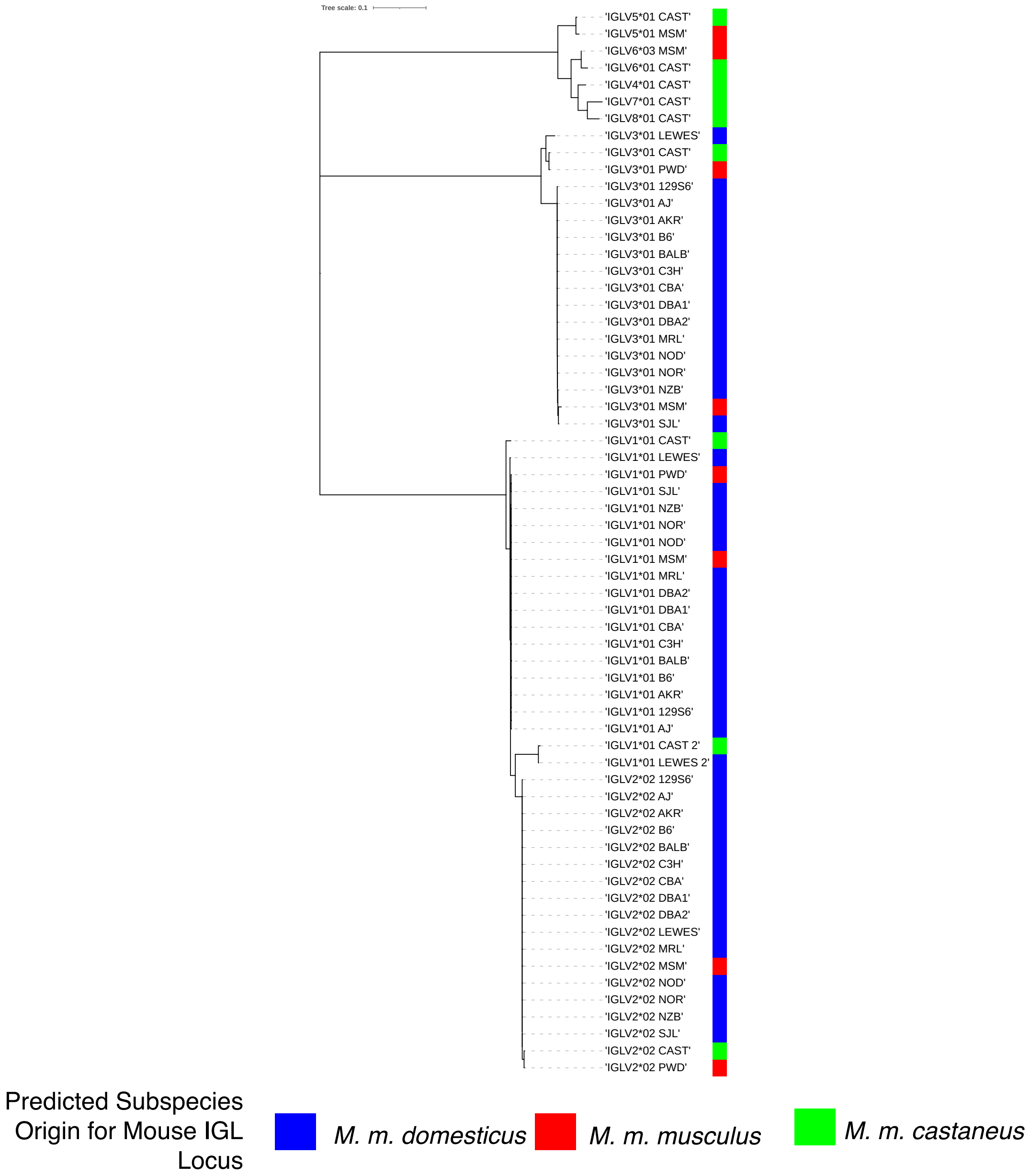


**Supplemental Figure 2. Phylogenetic tree of inferred IGLV sequences.** Branch lengths correspond to sequence identity percentage. If two branches are the same length, then the sequences are identical.

**Supplemental Table 1.** Inferred IGKV and IGLV germline sequences.

| Name | Strain | Closest IMGT Allele | % ID to IMGT Allele | Inferred Sequence Length | Sequence |
|---|---|---|---|---|---|
| IGKV6-15*01_129 | 129S1/SvImJ | Musmus IGKV6-15*01 F | 100 | 287 | GACATTGTGATGACCCAGTCTCAAAAATTCATGTCCACATCAGTAGGAGACAGGGTCAGCGTCACCTGCAAGGCCAGTCAGAATGTGGGTACTAATGTAGCCTGGTATCAACAGAAACCAGGGCAATCTCCTAAAGCACTGATTTACTCGGCATCCTACCGGTACAGTGGAGTCCCTGATCGCTTCACAGGCAGTGGATCTGGGACAGATTTCACTCTCACCATCAGCAATGTGCAGTCTGAAGACTTGGCAGAGTATTTCTGTCAGCAATATAACAGCTATCCTCT |
| IGKV8-30*01_129 | 129S1/SvImJ | Musmus IGKV8-30*01 F | 100 | 305 | GACATTGTGATGTCACAGTCTCCATCCTCCCTAGCTGTGTCAGTTGGAGAGAAGGTTACTATGAGCTGCAAGTCCAGTCAGAGCCTTTTATATAGTAGCAATCAAAAGAACTACTTGGCCTGGTACCAGCAGAAACCAGGGCAGTCTCCTAAACTGCTGATTTACTGGGCATCCACTAGGGAATCTGGGGTCCCTGATCGCTTCACAGGCAGTGGATCTGGGACAGATTTCACTCTCACCATCAGCAGTGTGAAGGCTGAAGACCTGGCAGTTTATTACTGTCAGCAATATTATAGCTATCCTCC |
| IGKV6-23*01_129 | 129S1/SvImJ | Musmus IGKV6-23*01 F | 100 | 287 | GACATTGTGATGACCCAGTCTCACAAATTCATGTCCACATCAGTAGGAGACAGGGTCAGCATCACCTGCAAGGCCAGTCAGGATGTGGGTACTGCTGTAGCCTGGTATCAACAGAAACCAGGGCAATCTCCTAAACTACTGATTTACTGGGCATCCACCCGGCACACTGGAGTCCCTGATCGCTTCACAGGCAGTGGATCTGGGACAGATTTCACTCTCACCATTAGCAATGTGCAGTCTGAAGACTTGGCAGATTATTTCTGTCAGCAATATAGCAGCTATCCTCT |
| IGKV12-44*01_129 | 129S1/SvImJ | Musmus IGKV12-44*01 F | 100 | 287 | GACATCCAGATGACTCAGTCTCCAGCCTCCCTATCTGCATCTGTGGGAGAAACTGTCACCATCACATGTCGAGCAAGTGAGAATATTTACAGTTATTTAGCATGGTATCAGCAGAAACAGGGAAAATCTCCTCAGCTCCTGGTCTATAATGCAAAAACCTTAGCAGAAGGTGTGCCATCAAGGTTCAGTGGCAGTGGATCAGGCACACAGTTTTCTCTGAAGATCAACAGCCTGCAGCCTGAAGATTTTGGGAGTTATTACTGTCAACATCATTATGGTACTCCTCC |
| IGKV8-27*01_129 | 129S1/SvImJ | Musmus IGKV8-27*01 F | 100 | 300 | AACATTATGATGACACAGTCGCCATCATCTCTGGCTGTGTCTGCAGGAGAAAAGGTCACTATGAGCTGTAAGTCCAGTCAAAGTGTTTTATACAGTTCAAATCAGAAGAACTACTTGGCCTGGTACCAGCAGAAACCAGGGCAGTCTCCTAAACTGCTGATCTACTGGGCATCCACTAGGGAATCTGGTGTCCCTGATCGCTTCACAGGCAGTGGATCTGGGACAGATTTTACTCTTACCATCAGCAGTGTACAAGCTGAAGACCTGGCAGTTTATTACTGTCATCAATACCTCTCCTCG |
| IGKV1-135*01_129 | 129S1/SvImJ | Musmus IGKV1-135*01 F | 100 | 302 | GATGTTGTGATGACCCAGACTCCACTCACTTTGTCGGTTACCATTGGACAACCAGCCTCCATCTCTTGCAAGTCAAGTCAGAGCCTCTTAGATAGTGATGGAAAGACATATTTGAATTGGTTGTTACAGAGGCCAGGCCAGTCTCCAAAGCGCCTAATCTATCTGGTGTCTAAACTGGACTCTGGAGTCCCTGACAGGTTCACTGGCAGTGGATCAGGGACAGATTTCACACTGAAAATCAGCAGAGTGGAGGCTGAGGATTTGGGAGTTTATTATTGCTGGCAAGGTACACATTTTCCTCA |
| IGKV1-110*01_129 | 129S1/SvImJ | Musmus IGKV1-110*01 F | 100 | 302 | GATGTTGTGATGACCCAAACTCCACTCTCCCTGCCTGTCAGTCTTGGAGATCAAGCCTCCATCTCTTGCAGATCTAGTCAGAGCCTTGTACACAGTAATGGAAACACCTATTTACATTGGTACCTGCAGAAGCCAGGCCAGTCTCCAAAGCTCCTGATCTACAAAGTTTCCAACCGATTTTCTGGGGTCCCAGACAGGTTCAGTGGCAGTGGATCAGGGACAGATTTCACACTCAAGATCAGCAGAGTGGAGGCTGAGGATCTGGGAGTTTATTTCTGCTCTCAAAGTACACATGTTCCTCC |

| | | | | | |
|---|---|---|---|---|---|
| IGKV6-20*01_129 | 129S1/SvImJ | Musmus IGKV6-20*01 F | 100 | 287 | AACATTGTAATGACCCAATCTCCCAAATCCATGTCCATGTCAGTAGGAGAGAGGGTCACCTTGAGCTGCAAGGCCAGTGAGAATGTGGGTACTTATGTATCCTGGTATCAACAGAAACCAGAGCAGTCTCCTAAACTGCTGATATACGGGGCATCCAACCGGTACACTGGGGTCCCCGATCGCTTCACAGGCAGTGGATCTGCAACAGATTTCACTCTGACCATCAGCAGTGTGCAGGCTGAAGACCTTGCAGATTATCACTGTGGACAGAGTTACAGCTATCCTCC |
| IGKV2-137*01_129 | 129S1/SvImJ | Musmus IGKV2-137*01 F | 100 | 302 | GATATTGTGATGACTCAGGCTGCACCCTCTGTACCTGTCACTCCTGGAGAGTCAGTATCCATCTCCTGCAGGTCTAGTAAGAGTCTCCTGCATAGTAATGGCAACACTTACTTGTATTGGTTCCTGCAGAGGCCAGGCCAGTCTCCTCAGCTCCTGATATATCGGATGTCCAACCTTGCCTCAGGAGTCCCAGACAGGTTCAGTGGCAGTGGGTCAGGAACTGCTTTCACACTGAGAATCAGTAGAGTGGAGGCTGAGGATGTGGGTGTTTATTACTGTATGCAACATCTAGAATATCCTTT |
| IGKV12-41*01_129 | 129S1/SvImJ | Musmus IGKV12-41*01 F | 100 | 287 | GACATCCAGATGACTCAGTCTCCAGCCTCCCTATCTGCATCTGTGGGAGAAACTGTCACCATCACATGTCGAGCAAGTGGGAATATTCACAATTATTTAGCATGGTATCAGCAGAAACAGGGAAAATCTCCTCAGCTCCTGGTCTATAATGCAAAAACCTTAGCAGATGGTGTGCCATCAAGGTTCAGTGGCAGTGGATCAGGAACACAATATTCTCTCAAGATCAACAGCCTGCAGCCTGAAGATTTTGGGAGTTATTACTGTCAACATTTTTGGAGTACTCCTCC |
| IGKV1-117*01_129 | 129S1/SvImJ | Musmus IGKV1-117*01 F | 100 | 302 | GATGTTTTGATGACCCAAACTCCACTCTCCCTGCCTGTCAGTCTTGGAGATCAAGCCTCCATCTCTTGCAGATCTAGTCAGAGCATTGTACATAGTAATGGAAACACCTATTTAGAATGGTACCTGCAGAAACCAGGCCAGTCTCCAAAGCTCCTGATCTACAAAGTTTCCAACCGATTTTCTGGGGTCCCAGACAGGTTCAGTGGCAGTGGATCAGGGACAGATTTCACACTCAAGATCAGCAGAGTGGAGGCTGAGGATCTGGGAGTTTATTACTGCTTTCAAGGTTCACATGTTCCTCC |
| IGKV8-19*01_129 | 129S1/SvImJ | Musmus IGKV8-19*01 F | 100 | 305 | GACATTGTGATGACACAGTCTCCATCCTCCCTGACTGTGACAGCAGGAGAGAAGGTCACTATGAGCTGCAAGTCCAGTCAGAGTCTGTTAAACAGTGGAAATCAAAAGAACTACTTGACCTGGTACCAGCAGAAACCAGGGCAGCCTCCTAAACTGTTGATCTACTGGGCATCCACTAGGGAATCTGGGGTCCCTGATCGCTTCACAGGCAGTGGATCTGGAACAGATTTCACTCTCACCATCAGCAGTGTGCAGGCTGAAGACCTGGCAGTTTATTACTGTCAGAATGATTATAGTTATCCTCC |
| IGKV4-59*01_129 | 129S1/SvImJ | Musmus IGKV4-59*01 F | 100 | 286 | CAAATTGTTCTCACCCAGTCTCCAGCAATCATGTCTGCATCTCCAGGGGAGAAGGTCACCATGACCTGCAGTGCCAGCTCAAGTGTAAGTTACATGCACTGGTACCAGCAGAAGTCAGGCACCTCCCCCAAAAGATGGATTTATGACACATCCAAACTGGCTTCTGGAGTCCCTGCTCGCTTCAGTGGCAGTGGGTCTGGGACCTCTTACTCTCTCACAATCAGCAGCATGGAGGCTGAAGATGCTGCCACTTATTACTGCCAGCAGTGGAGTAGTAACCCACCCA |
| IGKV12-46*01_129 | 129S1/SvImJ | Musmus IGKV12-46*01 F | 100 | 287 | GACATCCAGATGACTCAGTCTCCAGCCTCCCTATCTGTATCTGTGGGAGAAACTGTCACCATCACATGTCGAGCAAGTGAGAATATTTACAGTAATTTAGCATGGTATCAGCAGAAACAGGGAAAATCTCCTCAGCTCCTGGTCTATGCTGCAACAAACTTAGCAGATGGTGTGCCATCAAGGTTCAGTGGCAGTGGATCAGGCACACAGTATTCCCTCAAGATCAACAGCCTGCAGTCTGAAGATTTTGGGAGTTATTACTGTCAACATTTTTGGGGTACTCCTCC |
| IGKV4-91*01_S5760_129 | 129S1/SvImJ | Musmus IGKV4-91*01 F | 100 | 292 | GAAATTGTGCTCACCCAGTCTCCAACCACCATGGCTGCATCTCCCGGGGAGAAGATCACTATCACCTGCAGTGCCAGCTCAAGTATAAGTTCCAATTACTTGCATTGGTATCAGCAGAAGCCAGGATTCTCCCCTAAACTCTTGATTTATAGGACATCCAATCTGGCTTCTGGAGTCCCAGCTCGCTTCAGTGGCAGTGGGTCTGGGACCTCTTACTCTCTCACAATTGGCACCATGGAGGCTGAAGATGTTGCCACTTACTACTGCCAGCAGGGTAGTAGTATACCGCTCA |

| | | | | | |
|---|---|---|---|---|---|
| IGKV10-96*01_129 | 129S1/SvImJ | Musmus IGKV10-96*01 F | 100 | 286 | GATATCCAGATGACACAGACTACATCCTCCCTGTCTGCCTCTCTGGGAGACAGAGTCACCATCAGTTGCAGGGCAAGTCAGGACATTAGCAATTATTTAAACTGGTATCAGCAGAAACCAGATGGAACTGTTAAACTCCTGATCTACTACACATCAAGATTACACTCAGGAGTCCCATCAAGGTTCAGTGGCAGTGGGTCTGGAACAGATTATTCTCTCACCATTAGCAACCTGGAGCAAGAAGATATTGCCACTTACTTTTGCCAACAGGGTAATACGCTTCCTC |
| IGKV4-70*01_129 | 129S1/SvImJ | Musmus IGKV4-70*01 F | 100 | 282 | CAAATTGTTCTCACCCAGTCTCCAGCAATCATGTCTGCATCTCCAGGGGAGAAGGTCACCATGACCTGCAGTGCCAGCTCAAGTATAAGTTACATGCACTGGTACCAGCAGAAGCCAGGCACCTCCCCCAAAAGATGGATTTATGACACATCCAAACTGGCTTCTGGAGTCCCTGCTCGCTTCAGTGGCAGTGGGTCTGGGACCTCTTATTCTCTCACAATCAGCAGCATGGAGGCTGAAGATGCTGCCACTTATTACTGCCATCAGCGGAGTAGTTACCCA |
| IGKV6-17*01_129 | 129S1/SvImJ | Musmus IGKV6-17*01 F | 100 | 287 | GACATTGTGATGACCCAGTCTCACAAATTCATGTCCACATCAGTAGGAGACAGGGTCAGCATCACCTGCAAGGCCAGTCAGGATGTGAGTACTGCTGTAGCCTGGTATCAACAGAAACCAGGACAATCTCCTAAACTACTGATTTACTCGGCATCCTACCGGTACACTGGAGTCCCTGATCGCTTCACTGGCAGTGGATCTGGGACGGATTTCACTTTCACCATCAGCAGTGTGCAGGCTGAAGACCTGGCAGTTTATTACTGTCAGCAACATTATAGTACTCCTCC |
| IGKV8-21*01_129 | 129S1/SvImJ | Musmus IGKV8-21*01 F | 100 | 302 | GACATTGTGATGTCACAGTCTCCATCCTCCCTGGCTGTGTCAGCAGGAGAGAAGGTCACTATGAGCTGCAAATCCAGTCAGAGTCTGCTCAACAGTAGAACCCGAAAGAACTACTTGGCTTGGTACCAGCAGAAACCAGGGCAGTCTCCTAAACTGCTGATCTACTGGGCATCCACTAGGGAATCTGGGGTCCCTGATCGCTTCACAGGCAGTGGATCTGGGACAGATTTCACTCTCACCATCAGCAGTGTGCAGGCTGAAGACCTGGCAGTTTATTACTGCAAGCAATCTTATAATCTTCC |
| IGKV8-16*01_129 | 129S1/SvImJ | Musmus IGKV8-16*01 F | 100 | 305 | GAAATTGTGTTGACCCAGTCTATACCATCCCTGACTGTGTCAGCAGGAGAGAGGGTCACTATCAGCTGCAAATCCAATCAGAATCTTTTATGGAGTGGAAACCAAAGGTACTGTTTGGTCTGGCACCAGTGGAAACCAGGGCAAACTCCTACACCGTTGATCACCTGGACATCTGATAGGTACTCTGGAGTCCCTGATCGTTTCATAGGCAGTGGATCTGTGACAGATTTCACTCTGACCATCAGCAGTGTGCAGGCTGAAGATGTGGCAGTTTATTTCTGTCAGCAGCATTTACACATTCCTCC |
| IGKV4-80*01_129 | 129S1/SvImJ | Musmus IGKV4-80*01 F | 100 | 282 | CAAATTGTTCTCACCCAGTCTCCAGCAATCATGTCTGCATCTCTAGGGGAGGAGATCACCCTAACCTGCAGTGCCAGCTCGAGTGTAAGTTACATGCACTGGTACCAGCAGAAGTCAGGCACTTCTCCCAAACTCTTGATTTATAGCACATCCAACCTGGCTTCTGGAGTCCCTTCTCGCTTCAGTGGCAGTGGGTCTGGGACCTTTTATTCTCTCACAATCAGCAGTGTGGAGGCTGAAGATGCTGCCGATTATTACTGCCATCAGTGGAGTAGTTATCCA |
| IGKV3-4*01_129 | 129S1/SvImJ | Musmus IGKV3-4*01 F | 100 | 299 | GACATTGTGCTGACCCAATCTCCAGCTTCTTTGGCTGTGTCTCTAGGGCAGAGGGCCACCATCTCCTGCAAGGCCAGCCAAAGTGTTGATTATGATGGTGATAGTTATATGAACTGGTACCAACAGAAACCAGGACAGCCACCCAAACTCCTCATCTATGCTGCATCCAATCTAGAATCTGGGATCCCAGCCAGGTTTAGTGGCAGTGGGTCTGGGACAGACTTCACCCTCAACATCCATCCTGTGGAGGAGGAGGATGCTGCAACCTATTACTGTCAGCAAAGTAATGAGGATCCTCC |
| IGKV5-43*01_129 | 129S1/SvImJ | Musmus IGKV5-43*01 F | 100 | 287 | GATATTGTGCTAACTCAGTCTCCAGCCACCCTGTCTGTGACTCCAGGAGATAGCGTCAGTCTTTCCTGCAGGGCCAGCCAAAGTATTAGCAACAACCTACACTGGTATCAACAAAAATCACATGAGTCTCCAAGGCTTCTCATCAAGTATGCTTCCCAGTCCATCTCTGGGATCCCCTCCAGGTTCAGTGGCAGTGGATCAGGGACAGATTTCACTCTCAGTATCAACAGTGTGGAGACTGAAGATTTTGGAATGTATTTCTGTCAACAGAGTAACAGCTGGCCTCA |

| | | | | | |
|---|---|---|---|---|---|
| IGKV6-32*01_129 | 129S1/SvImJ | Musmus IGKV6-32*01 F | 100 | 287 | AGTATTGTGATGACCCAGACTCCCAAATTCCTGCTTGTATCAGCAGGAGACAGGGTTACCATAACCTGCAAGGCCAGTCAGAGTGTGAGTAATGATGTAGCTTGGTACCAACAGAAGCCAGGGCAGTCTCCTAAACTGCTGATATACTATGCATCCAATCGCTACACTGGAGTCCCTGATCGCTTCACTGGCAGTGGATATGGGACGGATTTCACTTTCACCATCAGCACTGTGCAGGCTGAAGACCTGGCAGTTTATTTCTGTCAGCAGGATTATAGCTCTCCTCC |
| IGKV14-126*01_129 | 129S1/SvImJ | Musmus IGKV14-126*01 F | 100 | 287 | GACATCAAGATGACCCAGTCTCCATCCTCCATGTATGCATCGCTGGGAGAGAGAGTCACTATCACTTGCAAGGCGAGTCAGGACATTAAAAGCTATTTAAGCTGGTACCAGCAGAAACCATGGAAATCTCCTAAGACCCTGATCTATTATGCAACAAGCTTGGCAGATGGGGTCCCATCAAGATTCAGTGGCAGTGGATCTGGGCAAGATTATTCTCTAACCATCAGCAGCCTGGAGTCTGACGATACAGCAACTTATTACTGTCTACAGCATGGTGAGAGCCCTCC |
| IGKV8-28*01_S7054_129 | 129S1/SvImJ | Musmus IGKV8-28*01 F | 99.66 | 305 | GACATTGTGATGACACAGTCTCCATCCTCCCTGAGTGTGTCAGCAGGAGAGAAGGTCACTATGAGCTGCAAGTCCAGTCAGAGTCTGTTAAACAGTGGAAATCAAAAGAACTACTTGGCCTGGTACCAGCAGAAACCAGGGCAGCCTCCTAAACTGTTGATCTACGGGGCATCCACTAGGGAATCTGGGGTCCCTGATCGCTTCACAGGCAGTGGATCTGGAACCGATTTCACTCTTACCATCAGCAGTGTGCAGGCTGAAGACCTGGCAGTTTATTACTGTCTGAATGATCATAGTTATCCTCC |
| IGKV3-2*01_129 | 129S1/SvImJ | Musmus IGKV3-2*01 F | 100 | 299 | GACATTGTGCTGACCCAATCTCCAGCTTCTTTGGCTGTGTCTCTAGGGCAGAGGGCCACCATCTCCTGCAGAGCCAGCGAAAGTGTTGATAATTATGGCATTAGTTTTATGAACTGGTTCCAACAGAAACCAGGACAGCCACCCAAACTCCTCATCTATGCTGCATCCAACCAAGGATCCGGGGTCCCTGCCAGGTTTAGTGGCAGTGGGTCTGGGACAGACTTCAGCCTCAACATCCATCCTATGGAGGAGGATGATACTGCAATGTATTTCTGTCAGCAAAGTAAGGAGGTTCCTCC |
| IGKV8-24*01_129 | 129S1/SvImJ | Musmus IGKV8-24*01 F | 100 | 305 | GACATTGTGATGACACAGTCTCCATCCTCCCTGGCTATGTCAGTAGGACAGAAGGTCACTATGAGCTGCAAGTCCAGTCAGAGCCTTTTAAATAGTAGCAATCAAAAGAACTATTTGGCCTGGTACCAGCAGAAACCAGGACAGTCTCCTAAACTTCTGGTATACTTTGCATCCACTAGGGAATCTGGGGTCCCTGATCGCTTCATAGGCAGTGGATCTGGGACAGATTTCACTCTTACCATCAGCAGTGTGCAGGCTGAAGACCTGGCAGATTACTTCTGTCAGCAACATTATAGCACTCCTCC |
| IGKV17-127*01_129 | 129S1/SvImJ | Musmus IGKV17-127*01 F | 100 | 287 | GAAACAACTGTGACCCAGTCTCCAGCATCCCTGTCCGTGGCTACAGGAGAAAAAGTCACTATCAGATGCATAACCAGCACTGATATTGATGATGATATGAACTGGTACCAGCAGAAGCCAGGGGAACCTCCTAAGCTCCTTATTTCAGAAGGCAATACTCTTCGTCCTGGAGTCCCATCCCGATTCTCCAGCAGTGGCTATGGCACAGATTTTGTTTTTACAATTGAAAACACGCTCTCAGAAGATGTTGCAGATTACTACTGTTTGCAAAGTGATAACATGCCTCT |
| IGKV3-5*01_129 | 129S1/SvImJ | Musmus IGKV3-5*01 F | 100 | 299 | GACATTGTGCTGACCCAATCTCCAGCTTCTTTGGCTGTGTCTCTAGGGCAGAGGGCCACCATATCCTGCAGAGCCAGTGAAAGTGTTGATAGTTATGGCAATAGTTTTATGCACTGGTACCAGCAGAAACCAGGACAGCCACCCAAACTCCTCATCTATCGTGCATCCAACCTAGAATCTGGGATCCCTGCCAGGTTCAGTGGCAGTGGGTCTAGGACAGACTTCACCCTCACCATTAATCCTGTGGAGGCTGATGATGTTGCAACCTATTACTGTCAGCAAAGTAATGAGGATCCTCC |
| IGKV4-57*01_129 | 129S1/SvImJ | Musmus IGKV4-57*01 F | 100 | 286 | CAAATTGTTCTCACCCAGTCTCCAGCAATCATGTCTGCATCTCCAGGGGAGAAGGTCACCATAACCTGCAGTGCCAGCTCAAGTGTAAGTTACATGCACTGGTTCCAGCAGAAGCCAGGCACTTCTCCCAAACTCTGGATTTATAGCACATCCAACCTGGCTTCTGGAGTCCCTGCTCGCTTCAGTGGCAGTGGATCTGGGACCTCTTACTCTCTCACAATCAGCCGAATGGAGGCTGAAGATGCTGCCACTTATTACTGCCAGCAAAGGAGTAGTTACCCACCCA |

| | | | | | |
|---|---|---|---|---|---|
| IGKV15-103*01_129 | 129S1/SvImJ | Musmus IGKV15-103*01 ORF | 100 | 287 | GACATCCAGATGAACCAGTCTCCATCCAGTCTGTCTGCATCCCTTGGAGACACAATTACCATCACTTGCCATGCCAGTCAGAACATTAATGTTTGGTTAAGCTGGTACCAGCAGAAACCAGGAAATATTCCTAAACTATTGATCTATAAGGCTTCCAACTTGCACACAGGCGTCCCATCAAGGTTTAGTGGCAGTGGATCTGGAACAGGTTTCACATTAACCATCAGCAGCCTGCAGCCTGAAGACATTGCCACTTACTACTGTCAACAGGGTCAAAGTTATCCTCT |
| IGKV6-13*01_129 | 129S1/SvImJ | Musmus IGKV6-13*01 F | 100 | 285 | GACATTGTGATGACCCAGTCTCAAAAATTCATGTCCACATCAGTAGGAGACAGGGTCAGCATCACCTGCAAGGCCAGTCAGAATGTGGGTACTGCTGTAGCCTGGTATCAACAGAAACCAGGACAATCTCCTAAACTACTGATTTACTCGGCATCCAATCGGTACACTGGAGTCCCTGATCGCTTCACAGGCAGTGGATCTGGGACAGATTTCACTCTCACCATCAGCAATATGCAGTCTGAAGACCTGGCAGATTATTTCTGCCAGCAATATAGCAGCTATCCT |
| IGKV3-12*01_129 | 129S1/SvImJ | Musmus IGKV3-12*01 F | 100 | 299 | GACATTGTGCTGACACAGTCTCCTGCTTCCTTAGCTGTATCTCTGGGGCAGAGGGCCACCATCTCATGCAGGGCCAGCAAAAGTGTCAGTACATCTGGCTATAGTTATATGCACTGGTACCAACAGAAACCAGGACAGCCACCCAAACTCCTCATCTATCTTGCATCCAACCTAGAATCTGGGGTCCCTGCCAGGTTCAGTGGCAGTGGGTCTGGGACAGACTTCACCCTCAACATCCATCCTGTGGAGGAGGAGGATGCTGCAACCTATTACTGTCAGCACAGTAGGGAGCTTCCTCC |
| IGKV4-72*01_129 | 129S1/SvImJ | Musmus IGKV4-72*01 F | 100 | 286 | CAAATTGTTCTCTCCCAGTCTCCAGCAATCCTGTCTGCATCTCCAGGGGAGAAGGTCACAATGACTTGCAGGGCCAGCTCAAGTGTAAGTTACATGCACTGGTACCAGCAGAAGCCAGGATCCTCCCCCAAACCCTGGATTTATGCCACATCCAACCTGGCTTCTGGAGTCCCTGCTCGCTTCAGTGGCAGTGGGTCTGGGACCTCTTACTCTCTCACAATCAGCAGAGTGGAGGCTGAAGATGCTGCCACTTATTACTGCCAGCAGTGGAGTAGTAACCCACCCA |
| IGKV5-48*01_129 | 129S1/SvImJ | Musmus IGKV5-48*01 F | 100 | 287 | GACATCTTGCTGACTCAGTCTCCAGCCATCCTGTCTGTGAGTCCAGGAGAAAGAGTCAGTTTCTCCTGCAGGGCCAGTCAGAGCATTGGCACAAGCATACACTGGTATCAGCAAAGAACAAATGGTTCTCCAAGGCTTCTCATAAAGTATGCTTCTGAGTCTATCTCTGGGATCCCTTCCAGGTTTAGTGGCAGTGGATCAGGGACAGATTTTACTCTTAGCATCAACAGTGTGGAGTCTGAAGATATTGCAGATTATTACTGTCAACAAAGTAATAGCTGGCCAAC |
| IGKV4-74*01_129 | 129S1/SvImJ | Musmus IGKV4-74*01 F | 100 | 292 | CAAATTGTTCTCACCCAGTCTCCAGCAATCATGTCTGCATCTCTAGGGGAACGGGTCACCATGACCTGCACTGCCAGCTCAAGTGTAAGTTCCAGTTACTTGCACTGGTACCAGCAGAAGCCAGGATCCTCCCCCAAACTCTGGATTTATAGCACATCCAACCTGGCTTCTGGAGTCCCAGCTCGCTTCAGTGGCAGTGGGTCTGGGACCTCTTACTCTCTCACAATCAGCAGCATGGAGGCTGAAGATGCTGCCACTTATTACTGCCACCAGTATCATCGTTCCCCACCCA |
| IGKV19-93*01_129 | 129S1/SvImJ | Musmus IGKV19-93*01 F | 100 | 287 | GACATCCAGATGACACAGTCTCCATCCTCACTGTCTGCATCTCTGGGAGGCAAAGTCACCATCACTTGCAAGGCAAGCCAAGACATTAACAAGTATATAGCTTGGTACCAACACAAGCCTGGAAAAGGTCCTAGGCTGCTCATACATTACACATCTACATTACAGCCAGGCATCCCATCAAGGTTCAGTGGAAGTGGGTCTGGGAGAGATTATTCCTTCAGCATCAGCAACCTGGAGCCTGAAGATATTGCAACTTATTATTGTCTACAGTATGATAATCTTCTACC |
| IGKV4-57-1*01_129 | 129S1/SvImJ | Musmus IGKV4-57*01 F | 100 | 292 | GAAAATGTGCTCACCCAGTCTCCAGCAATCATGTCTGCATCTCCAGGGGAAAAGGTCACCATGACCTGCAGGGCCAGCTCAAGTGTAAGTTCCAGTTACTTGCACTGGTACCAGCAGAAGTCAGGTGCCTCCCCCAAACTCTGGATTTATAGCACATCCAACTTGGCTTCTGGAGTCCCTGCTCGCTTCAGTGGCAGTGGGTCTGGGACCTCTTACTCTCTCACAATCAGCAGTGTGGAGGCTGAAGATGCTGCCACTTATTACTGCCAGCAGTACAGTGGTTACCCACTCA |

| | | | | | |
|---|---|---|---|---|---|
| IGKV17-121*01_129 | 129S1/SvImJ | Musmus IGKV17-121*01 F | 100 | 287 | GAAACAACTGTGACCCAGTCTCCAGCATCCCTGTCCATGGCTATAGGAGAAAAAGTCA<br>CCATCAGATGCATAACCAGCACTGATATTGATGATGATATGAACTGGTACCAGCAGAA<br>GCCAGGGGAACCTCCTAAGCTCCTTATTTCAGAAGGCAATACTCTTCGTCCTGGAGTC<br>CCATCCCGATTCTCCAGCAGTGGCTATGGTACAGATTTTGTTTTTACAATTGAAAACAT<br>GCTCTCAGAAGATGTTGCAGATTACTACTGTTTGCAAAGTGATAACTTGCCTCT |
| IGKV4-55*01_S6279_129 | 129S1/SvImJ | Musmus IGKV4-55*01 F | 100 | 286 | CAAATTGTTCTCACCCAGTCTCCAGCAATCATGTCTGCATCTCCAGGGGAGAAGGTCA<br>CCATGACCTGCAGTGCCAGCTCAAGTGTAAGTTACATGTACTGGTACCAGCAGAAGCC<br>AGGATCCTCCCCCAGACTCCTGATTTATGACACATCCAACCTGGCTTCTGGAGTCCCTG<br>TTCGCTTCAGTGGCAGTGGGTCTGGGACCTCTTACTCTCTCACAATCAGCCGAATGGA<br>GGCTGAAGATGCTGCCACTTATTACTGCCAGCAGTGGAGTAGTTACCCACTCA |
| IGKV4-68*01_129 | 129S1/SvImJ | Musmus IGKV4-68*01 F | 100 | 286 | CAAATTGTTCTCACCCAGTCTCCAGCACTCATGTCTGCATCTCCAGGGGAGAAGGTCA<br>CCATGACCTGCAGTGCCAGCTCAAGTGTAAGTTACATGTACTGGTACCAGCAGAAGCC<br>AAGATCCTCCCCCAAACCCTGGATTTATCTCACATCCAACCTGGCTTCTGGAGTCCCTG<br>CTCGCTTCAGTGGCAGTGGGTCTGGGACCTCTTACTCTCTCACAATCAGCAGCATGGA<br>GGCTGAAGATGCTGCCACTTATTACTGCCAGCAGTGGAGTAGTAACCCACCCA |
| IGKV6-25*01_129 | 129S1/SvImJ | Musmus IGKV6-25*01 F | 100 | 287 | GACATTGTGATGACCCAGTCTCACAAATTCATGTCCACATCAGTAGGAGACAGGGTCA<br>GCATCACCTGCAAGGCCAGTCAGGATGTGAGTACTGCTGTAGCCTGGTATCAACAAAA<br>ACCAGGGCAATCTCCTAAACTACTGATTTACTGGGCATCCACCCGGCACACTGGAGTC<br>CCTGATCGCTTCACAGGCAGTGGATCTGGGACAGATTATACTCTCACCATCAGCAGTG<br>TGCAGGCTGAAGACCTGGCACTTTATTACTGTCAGCAACATTATAGCACTCCTCC |
| IGKV9-120*01_S2628_129 | 129S1/SvImJ | Musmus IGKV9-120*02 F | 100 | 285 | GACATCCAGATGACCCAGTCTCCATCCTCCTTATCTGCCTCTCTGGGAGAAAGAGTCA<br>GTCTCACTTGTCGGGCAAGTCAGGACATTGGTAGTAGCTTAAACTGGCTTCAGCAGGA<br>ACCAGATGGAACTATTAAACGCCTGATCTACGCCACATCCAGTTTAGATTCTGGTGTC<br>CCCAAAAGGTTCAGTGGCAGTAGGTCTGGGTCAGATTATTCTCTCACCATCAGCAGCC<br>TTGAGTCTGAAGATTTTGTAGACTATTACTGTCTACAATATGCTAGTTCTCCT |
| IGKV3-7*01_S5296_129 | 129S1/SvImJ | Musmus IGKV3-7*02 F | 100 | 299 | GACATTGTGCTGACACAGTCTCCTGCTTCCTTAGCTGTATCTCTGGGGCAGAGGGCCA<br>CCATCTCATGCAGGGCCAGCCAAAGTGTCAGTACATCTAGCTATAGTTATATGCACTG<br>GTACCAACAGAAACCAGGACAGCCACCCAAACTCCTCATCAAGTATGCATCCAACCT<br>AGAATCTGGGGTCCCTGCCAGGTTCAGTGGCAGTGGGTCTGGGACAGACTTCACCCTC<br>AACATCCATCCTGTGGAGGAGGAGGATACTGCAACATATTACTGTCAGCACAGTTGGG<br>AGATTCCTCC |
| IGKV10-94*01_129 | 129S1/SvImJ | Musmus IGKV10-94*01 F | 100 | 287 | GATATCCAGATGACACAGACTACATCCTCCCTGTCTGCCTCTCTGGGAGACAGAGTCA<br>CCATCAGTTGCAGTGCAAGTCAGGGCATTAGCAATTATTTAAACTGGTATCAGCAGAA<br>ACCAGATGGAACTGTTAAACTCCTGATCTATTACACATCAAGTTTACACTCAGGAGTC<br>CCATCAAGGTTCAGTGGCAGTGGGTCTGGGACAGATTATTCTCTCACCATCAGCAACC<br>TGGAACCTGAAGATATTGCCACTTACTATTGTCAGCAGTATAGTAAGCTTCCTCC |
| IGKV13-84*01_129 | 129S1/SvImJ | Musmus IGKV13-84*01 F | 100 | 287 | GACATCCAGATGACACAATCTTCATCCTCCTTTTCTGTATCTCTAGGAGACAGAGTCAC<br>CATTACTTGCAAGGCAAGTGAGGACATATATAATCGGTTAGCCTGGTATCAGCAGAAA<br>CCAGGAAATGCTCCTAGGCTCTTAATATCTGGTGCAACCAGTTTGGAAACTGGGGTTC<br>CTTCAAGATTCAGTGGCAGTGGATCTGGAAAGGATTACACTCTCAGCATTACCAGTCT<br>TCAGACTGAAGATGTTGCTACTTATTACTGTCAACAGTATTGGAGTACTCCTCC |

| | | | | | |
|---|---|---|---|---|---|
| IGKV13-85*01_129 | 129S1/SvImJ | Musmus IGKV13-85*01 F | 100 | 287 | GACATCCAGATGACACAATCTTCATCCTACTTGTCTGTATCTCTAGGAGGCAGAGTCACCATTACTTGCAAGGCAAGTGACCACATTAATAATTGGTTAGCCTGGTATCAGCAGAAACCAGGAAATGCTCCTAGGCTCTTAATATCTGGTGCAACCAGTTTGGAAACTGGGGTTCCTTCAAGATTCAGTGGCAGTGGATCTGGAAAGGATTACACTCTCAGCATTACCAGTCTTCAGACTGAAGATGTTGCTACTTATTACTGTCAACAGTATTGGAGTACTCCTCC |
| IGKV4-86*01_129 | 129S1/SvImJ | Musmus IGKV4-86*01 F | 100 | 283 | GAAATTGTGCTCACTCAGTCTCCAGCCATCACAGCTGCATCTCTGGGGCAAAAGGTCACCATCACCTGCAGTGCCAGCTCAAGTGTAAGTTACATGCACTGGTACCAGCAGAAGTCAGGCACCTCCCCCAAACCATGGATTTATGAAATATCCAAACTGGCTTCTGGAGTCCCAGCTCGCTTCAGTGGCAGTGGGTCTGGGACCTCTTACTCTCTCACAATCAGCAGCATGGAGGCTGAAGATGCTGCCATTTATTACTGCCAGCAGTGGAATTATCCTCTTA |
| IGKV17-127*01_S0118_129 | 129S1/SvImJ | Musmus IGKV17-127*01 F | 100 | 287 | GAAACAACTGTGACCCAGTCTCCAGCATCCCTGTCCGTGGCTACAGGAGAAAAAGTCACTATCAGATGCATAACCAGCACTGATATTGATGATGATATGAACTGGTACCAGCAGAAGCCAGGGGAACCTCCTAAGCTCCTTATTTCAGAAGGCAATACTCTTCGTCCTGGAGTCCCATCCCGATTCTCCAGCAGTGGCTATGGCACAGATTTTGTTTTTACAATTGAAAACACGCTCTCAGAAGATGTTGCAGATTACTACTGTTTGCAAAGTGATAACATGCCGCT |
| IGKV1-133*01_129 | 129S1/SvImJ | Musmus IGKV1-133*01 F | 100 | 302 | GATGTTGTGATGACCCAGACTCCACTCACTTTGTCGGTTACCATTGGACAACCAGCCTCTATCTCTTGCAAGTCAAGTCAGAGCCTCTTATATAGTAATGGAAAAACCTATTTGAATTGGTTATTACAGAGGCCAGGCCAGTCTCCAAAGCGCCTAATCTATCTGGTGTCTAAACTGGACTCTGGAGTCCCTGACAGGTTCACTGGCAGTGGATCAGGAACAGATTTTACACTGAAAATCAGCAGAGTGGAGGCTGAGGATTTGGGAGTTTATTACTGCGTGCAAGGTACACATTTTCCTCA |
| IGKV1-122*01_129 | 129S1/SvImJ | Musmus IGKV1-122*01 F | 100 | 302 | GATGCTGTGATGACCCAAACTCCACTCTCCCTGCCTGTCAGTCTTGGAGATCAAGCCTCCATCTCTTGCAGGTCTAGTCAGAGCCTTGAAAACAGTAATGGAAACACCTATTTGAACTGGTACCTCCAGAAACCAGGCCAGTCTCCACAGCTCCTGATCTACAGGGTTTCCAACCGATTTTCTGGGGTCCTAGACAGGTTCAGTGGTAGTGGATCAGGGACAGATTTCACACTGAAAATCAGCAGAGTGGAGGCTGAGGATTTGGGAGTTTATTTCTGCCTCCAAGTTACACATGTCCCTCC |
| IGKV16-104*01_129 | 129S1/SvImJ | Musmus IGKV16-104*01 F | 100 | 288 | GATGTCCAGATAACCCAGTCTCCATCTTATCTTGCTGCATCTCCTGGAGAAACCATTACTATTAATTGCAGGGCAAGTAAGAGCATTAGCAAATATTTAGCCTGGTATCAAGAGAAACCTGGGAAAACTAATAAGCTTCTTATCTACTCTGGATCCACTTTGCAATCTGGAATTCCATCAAGGTTCAGTGGCAGTGGATCTGGTACAGATTTCACTCTCACCATCAGTAGCCTGGAGCCTGAAGATTTTGCAATGTATTACTGTCAACAGCATAATGAATACCCGTAC |
| IGKV3-10*01_129 | 129S1/SvImJ | Musmus IGKV3-10*01 F | 100 | 299 | AACATTGTGCTGACCCAATCTCCAGCTTCTTTGGCTGTGTCTCTAGGGCAGAGGGCCACCATATCCTGCAGAGCCAGTGAAAGTGTTGATAGTTATGGCAATAGTTTTATGCACTGGTACCAGCAGAAACCAGGACAGCCACCCAAACTCCTCATCTATCTTGCATCCAACCTAGAATCTGGGGTCCCTGCCAGGTTCAGTGGCAGTGGGTCTAGGACAGACTTCACCCTCACCATTGATCCTGTGGAGGCTGATGATGCTGCAACCTATTACTGTCAGCAAAATAATGAGGATCCTCC |
| IGKV14-111*01_129 | 129S1/SvImJ | Musmus IGKV14-111*01 F | 100 | 287 | GACATCAAGATGACCCAGTCTCCATCTTCCATGTATGCATCTCTAGGAGAGAGAGTCACTATCACTTGCAAGGCGAGTCAGGACATTAATAGCTATTTAAGCTGGTTCCAGCAGAAACCAGGGAAATCTCCTAAGACCCTGATCTATCGTGCAAACAGATTGGTAGATGGGGTCCCATCAAGGTTCAGTGGCAGTGGATCTGGGCAAGATTATTCTCTCACCATCAGCAGCCTGGAGTATGAAGATATGGGAATTTATTATTGTCTACAGTATGATGAGTTTCCTCC |

| | | | | | |
|---|---|---|---|---|---|
| IGKV2-109*01_129 | 129S1/SvImJ | Musmus IGKV2-109*01 F | 100 | 302 | GATATTGTGATGACGCAGGCTGCATTCTCCAATCCAGTCACTCTTGGAACATCAGCTTCCATCTCCTGCAGGTCTAGTAAGAGTCTCCTACATAGTAATGGCATCACTTATTTGTATTGGTATCTGCAGAAGCCAGGCCAGTCTCCTCAGCTCCTGATTTATCAGATGTCCAACCTTGCCTCAGGAGTCCCAGACAGGTTCAGTAGCAGTGGGTCAGGAACTGATTTCACACTGAGAATCAGCAGAGTGGAGGCTGAGGATGTGGGTGTTTATTACTGTGCTCAAAATCTAGAACTTCCTCC |
| IGKV4-50*01_S8668_129 | 129S1/SvImJ | Musmus IGKV4-50*01 F | 100 | 286 | GAAAATGTGCTCACCCAGTCTCCAGCAATCATGTCTGCATCTCTAGGGGAGAAGGTCACCATGAGCTGCAGGGCCAGCTCAAGTGTAAATTACATGTACTGGTACCAGCAGAAGTCAGATGCCTCCCCCAAACTATGGATTTATTACACATCCAACCTGGCTCCTGGAGTCCCAGCTCGCTTCAGTGGCAGTGGGTCTGGGAACTCTTATTCTCTCACAATCAGCAGCATGGAGGGTGAAGATGCTGCCACTTATTACTGCCAGCAGTTTACTAGTTCCCCGTACA |
| IGKV4-63*01_129 | 129S1/SvImJ | Musmus IGKV4-63*01 F | 100 | 286 | GAAAATGTTCTCACCCAGTCTCCAGCAATCATGTCTGCATCTCCAGGGGAAAAGGTCACCATGACCTGCAGTGCCAGCTCAAGTGTAAGTTACATGCACTGGTACCAGCAGAAGTCAAGCACCTCCCCCAAACTCTGGATTTATGACACATCCAAACTGGCTTCTGGAGTCCCAGGTCGCTTCAGTGGCAGTGGGTCTGGAAACTCTTACTCTCTCACGATCAGCAGCATGGAGGCTGAAGATGTTGCCACTTATTACTGTTTTCAGGGGAGTGGGTACCCACTCA |
| IGKV12-89*01_129 | 129S1/SvImJ | Musmus IGKV12-89*01 F | 100 | 287 | GACATCCAGATGACTCAGTCTCCAGCTTCACTGTCTGCATCTGTGGGAGAAACTGTCACCATCACATGTGGAGCAAGTGAGAATATTTACGGTGCTTTAAATTGGTATCAGCGGAAACAGGGAAAATCTCCTCAGCTCCTGATCTATGGTGCAACCAACTTGGCAGATGGCATGTCATCGAGGTTCAGTGGCAGTGGATCTGGTAGACAGTATTCTCTCAAGATCAGTAGCCTGCATCCTGACGATGTTGCAACGTATTACTGTCAAAATGTGTTAAGTACTCCTCC |
| IGKV4-91*01_S5997_129 | 129S1/SvImJ | Musmus IGKV4-91*01 F | 100 | 292 | GAAATTGTGCTCACCCAGTCTCCAACCACCATGGCTGCATCTCCCGGGGAGAAGATCACTATCACCTGCAGTGCCAGCTCAAGTATAAGTTCCAATTACTTGCATTGGTATCAGCAGAAGCCAGGATTCTCCCCTAAACTCTTGATTTATAGGACATCCAATCTGGCTTCTGGAGTCCCAGCTCGCTTCAGTGGCAGTGGGTCTGGGACCTCTTACTCTCTCACAATTGGCACCATGGAGGCTGAAGATGTTGCCACTTACTACTGCCAGCAGGGTAGTAGTATACCACTCA |
| IGKV14-100*01_129 | 129S1/SvImJ | Musmus IGKV14-100*01 F | 100 | 287 | GACATCCTGATGACCCAATCTCCATCCTCCATGTCTGTATCTCTGGGAGACACAGTCAGCATCACTTGCCATGCAAGTCAGGGCATTAGCAGTAATATAGGGTGGTTGCAGCAGAAACCAGGGAAATCATTTAAGGGCCTGATCTATCATGGAACCAACTTGGAAGATGGAGTTCCATCAAGGTTCAGTGGCAGTGGATCTGGAGCAGATTATTCTCTCACCATCAGCAGCCTGGAATCTGAAGATTTTGCAGACTATTACTGTGTACAGTATGCTCAGTTTCCTCC |
| IGKV8-34*01_129 | 129S1/SvImJ | Musmus IGKV8-34*01 F | 100 | 305 | GACATTTTGATGACCCAGTCTCCATCCTCCCTGACTGTGTCAGCAGGAGAGAAGGTCACTATGAGCTGCAAGTCCAGTCAGAGTCTTTTAGCTAGTGGCAACCAAAATAACTACTTGGCCTGGCACCAGCAGAAACCAGGACGATCTCCTAAAATGCTGATAATTTGGGCATCCACTAGGGTATCTGGAGTCCCTGATCGCTTCATAGGCAGTGGATCTGGGACGGATTTCACTCTGACCATCAACAGTGTGCAGGCTGAAGATCTGGCTGTTTATTACTGTCAGCAGTCCTACAGCGCTCCTAC |
| IGKV4-53*01_129 | 129S1/SvImJ | Musmus IGKV4-53*01 F | 100 | 292 | GAAATTGTGCTCACCCAGTCTCCAGCACTCATGGCTGCATCTCCAGGGGAGAAGGTCACCATCACCTGCAGTGTCAGCTCAAGTATAAGTTCCAGCAACTTGCACTGGTACCAGCAGAAGTCAGAAACCTCCCCCAAACCCTGGATTTATGGCACATCCAACCTGGCTTCTGGAGTCCCTGTTCGCTTCAGTGGCAGTGGATCTGGGACCTCTTATTCTCTCACAATCAGCAGCATGGAGGCTGAAGATGCTGCCACTTATTACTGTCAACAGTGGAGTAGTTACCCACTCA |

| | | | | | |
|---|---|---|---|---|---|
| IGKV5-39*01_129 | 129S1/SvImJ | Musmus IGKV5-39*01 F | 100 | 287 | GACATTGTGATGACTCAGTCTCCAGCCACCCTGTCTGTGACTCCAGGAGATAGAGTCT CTCTTTCCTGCAGGGCCAGCCAGAGTATTAGCGACTACTTACACTGGTATCAACAAAA ATCACATGAGTCTCCAAGGCTTCTCATCAAATATGCTTCCCAATCCATCTCTGGGATCC CCTCCAGGTTCAGTGGCAGTGGATCAGGGTCAGATTTCACTCTCAGTATCAACAGTGT GGAACCTGAAGATGTTGGAGTGTATTACTGTCAAAATGGTCACAGCTTTCCTCC |
| IGKV4-61*01_S2955_129 | 129S1/SvImJ | Musmus IGKV4-61*01 F | 100 | 286 | CAAATTGTTCTCACCCAGTCTCCAGCAATCATGTCTGCATCTCCAGGGGAGAAGGTCA CCATATCCTGCAGTGCCAGCTCAAGTGTAAGTTACATGTACTGGTACCAGCAGAAGCC AGGATCCTCCCCCAAACCCTGGATTTATCGCACATCCAACCTGGCTTCTGGAGTCCCT GCTCGCTTCAGTGGCAGTGGGTCTGGGACCTCTTACTCTCTCACAATCAGCAGCATGG AGGCTGAAGATGCTGCCACTTATTACTGCCAGCAGTATCATAGTTACCCACTCA |
| IGKV2-112*01_129 | 129S1/SvImJ | Musmus IGKV2-112*01 F | 100 | 300 | GATATTGTGATAACCCAGGATGAACTCTCCAATCCTGTCACTTCTGGAGAATCAGTTT CCATCTCCTGCAGGTCTAGTAAGAGTCTCCTATATAAGGATGGGAAGACATACTTGAA TTGGTTTCTGCAGAGACCAGGACAATCTCCTCAGCTCCTGATCTATTTGATGTCCACCC GTGCATCAGGAGTCTCAGACCGGTTTAGTGGCAGTGGGTCAGGAACAGATTTCACCCT GGAAATCAGTAGAGTGAAGGCTGAGGATGTGGGTGTGTATTACTGTCAACAACTTGTA GAGTATCCT |
| IGKV8-23-1*01_129 | 129S1/SvImJ | Musmus IGKV8-16*01 F, or Musmus IGKV8-23-1*01 ORF | 100 | 284 | GAAATTGTGCTGACTGTGTCAGCAGGAGAGAGGGTCACTATCAGCTGCAAATCCAATC AGAATCTTTTATGGAGTGGAAACCAAAGGTACTGTTTGGTCTGGCACCAGTGGAAACC AGGGCAAACTCCTACACCGTTGATCACCTGGACATCTGATAGGTACTCTGGAGTCCCT GATCGTTTCATAGGCAGTGGATCTGTGACAGATTTCACTCTGACCATCAGCAGTGTGC AGGCTGAAGATGTGGCAGTTTATTTCTGTCAGCAGCATTTACACATTCCTCC |
| IGKV9-124*01_129 | 129S1/SvImJ | Musmus IGKV9-124*01 F | 100 | 285 | GACATCCAGATGACCCAGTCTCCATCCTCCTTATCTGCCTCTCTGGGAGAAAGAGTCA GTCTCACTTGTCGGGCAAGTCAGGAAATTAGTGGTTACTTAAGCTGGCTTCAGCAGAA ACCAGATGGAACTATTAAACGCCTGATCTACGCCGCATCCACTTTAGATTCTGGTGTC CCAAAAAGGTTCAGTGGCAGTAGGTCTGGGTCAGATTATTCTCTCACCATCAGCAGCC TTGAGTCTGAAGATTTTGCAGACTATTACTGTCTACAATATGCTAGTTATCCT |
| IGKV12-98*01_129 | 129S1/SvImJ | Musmus IGKV12-98*01 F | 100 | 287 | GACATTCAGATGACCCAGTCTCCTGCCTCCCAGTCTGCATCTCTGGGAGAAAGTGTCA CCATCACATGCCTGGCAAGTCAGACCATTGGTACATGGTTAGCATGGTATCAGCAGAA ACCAGGGAAATCTCCTCAGCTCCTGATTTATGCTGCAACCAGCTTGGCAGATGGGGTC CCATCAAGGTTCAGTGGTAGTGGATCTGGCACAAAATTTTCTTTCAAGATCAGCAGCC TACAGGCTGAAGATTTTGTAAGTTATTACTGTCAACAACTTTACAGTACTCCTCT |
| IGKV4-69*01_129 | 129S1/SvImJ | Musmus IGKV4-69*01 F | 100 | 282 | CAAATTCTTCTCACCCAGTCTCCAGCAATCATGTCTGCATCTCCAGGGGAGAAGGTCA CCATGACCTGCAGTGCCAGCTCAAGTGTAAGTTACATGCACTGGTACCAGCAGAAGCC AGGATCCTCGCCCAAACCCTGGATTTATGACACATCCAACCTGGCTTCTGGATTCCCT GCTCGCTTCAGTGGCAGTGGGTCTGGGACCTCTTACTCTCTCATAATCAGCAGCATGG AGGCTGAAGATGCTGCCACTTATTACTGCCATCAGCGGAGTAGTTACCCA |
| IGKV5-45*01_129 | 129S1/SvImJ | Musmus IGKV5-45*01 F | 100 | 287 | GATATTGTGCTAACTCAGTCTCCAGCCACCCTGTCTGTGACTCCAGGAGATAGAGTCA GTCTTTCCTGCAGGGCCAGTCAAAGTATTAGCAACTACCTACACTGGTATCAACAAAA ATCACATGAGTCTCCAAGGCTTCTCATCAAGTATGCTTCCCAGTCCATCTCTGGGATCC CCTCCAGGTTCAGTGGCAGTGGATCAGGGACAGATTTCACTCTCAGTATCAACAGTGT GGAGACTGAAGATTTTGGAATGTATTTCTGTCAACAGAGTAACAGCTGGCCTCA |

| | | | | | |
|---|---|---|---|---|---|
| IGKV11-125*01_129 | 129S1/SvImJ | Musmus IGKV11-125*01 F | 100 | 287 | GATGTCCAGATGATTCAGTCTCCATCCTCCCTGTCTGCATCTTTGGGAGACATAGTCACCATGACTTGCCAGGCAAGTCAGGGCACTAGCATTAATTTAAACTGGTTTCAGCAAAAACCAGGGAAAGCTCCTAAGCTCCTGATCTATGGTGCAAGCAACTTGGAAGATGGGGTCCCATCAAGGTTCAGTGGCAGTAGATATGGGACAGATTTCACTCTCACCATCAGCAGCCTGGAGGATGAAGATATGGCAACTTATTTCTGTCTACAGCATAGTTATCTCCCTCC |
| IGKV14-130*01_129 | 129S1/SvImJ | Musmus IGKV14-130*01 F | 100 | 287 | GAAATCCAGATGACCCAGTCTCCATCCTCTATGTCTGCATCTCTGGGAGACAGAATAACCATCACTTGCCAGGCAACTCAAGACATTGTTAAGAATTTAAACTGGTATCAGCAGAAACCAGGGAAACCCCCTTCATTCCTGATCTATTATGCAACTGAACTGGCAGAAGGGGTCCCATCAAGGTTCAGTGGCAGTGGGTCTGGGTCAGACTATTCTCTGACAATCAGCAACCTGGAGTCTGAAGATTTTGCAGACTATTACTGTCTACAGTTTTATGAGTTTCCTCC |
| IGKV4-79*01_129 | 129S1/SvImJ | Musmus IGKV4-79*01 F | 100 | 292 | CAAATTGTTCTCACCCAGTCTCCAGCAATCATGTCTGCATCTCCTGGGGAGAAGGTCACCTTGACCTGCAGTGCCAGCTCAAGTGTAAGTTCCAGCTACTTGTACTGGTACCAGCAGAAGCCAGGATCCTCCCCCAAACTCTGGATTTATAGCACATCCAACCTGGCTTCTGGAGTCCCTGCTCGCTTCAGTGGCAGTGGGTCTGGGACCTCTTACTCTCTCACAATCAGCAGCATGGAGGCTGAAGATGCTGCCTCTTATTTCTGCCATCAGTGGAGTAGTTACCCACCCC |
| IGKV1-88*01_129 | 129S1/SvImJ | Musmus IGKV1-88*01 F | 100 | 302 | GATGTTGTGGTGACTCAAACTCCACTCTCCCTGCCTGTCAGCTTTGGAGATCAAGTTTCTATCTCTTGCAGGTCTAGTCAGAGTCTTGCAAACAGTTATGGGAACACCTATTTGTCTTGGTACCTGCACAAGCCTGGCCAGTCTCCACAGCTCCTCATCTATGGGATTTCCAACAGATTTTCTGGGGTGCCAGACAGGTTCAGTGGCAGTGGTTCAGGGACAGATTTCACACTCAAGATCAGCACAATAAAGCCTGAGGACTTGGGAATGTATTACTGCTTACAAGGTACACATCAGCCTCC |
| IGKV10-95*01_129 | 129S1/SvImJ | Musmus IGKV10-95*01 F | 100 | 287 | GATATCCAGATGACACAGACTACTTCCTCCCTGTCTGCCTCTCTGGGAGACAGAGTCACCATCAGTTGCAGGGCAAGTGAGGACATTAGCACTTATTTAAACTGGTATCAGCAGAAACCAGATGGAACTGTTAAACTCCTGATCTATTACACATCAGGATTACACTCAGGAGTCCCATCAAGGTTCAGTGGCAGTGGGTCTGGGGCAGATTATTCTCTCACCATCAGCAACCTGGAACCTGAAGATATTGCCACTTACTATTGTCAGCAGTATAGTAAGCTTCCTCC |
| IGKV4-92*01_129 | 129S1/SvImJ | Musmus IGKV4-92*01 F | 100 | 291 | GAAATGGTTCTCACCCAGTCTCCAGTATCCATAACTGCATCTCGAGGGGAGAAGGTCACCATCACCTGCCGTGCCAGCTCAAGTATAAGTTCCAATTACTTACACTGGTACCAGCAGAAGCCAGGATCCTCCCCTAAACTTTTGATTTATAGGACATCCATCCTGGCATCTGGAGTCCTAGACAGCTTCAGTGGCAGTGGGTCTGAGAGCTCTTACACTCTGACAATCAGCTGCATGCAGGACGAAGTTGCTGCCACTTACTATTGTCAGCAGGGGAGTAGTAGCCCACCA |
| IGKV9-123*01_129 | 129S1/SvImJ | Musmus IGKV9-123*01 F | 100 | 285 | GACATCCAGATGATTCAGTCTCCATCGTCCATGTTTGCCTCTCTGGGAGACAGAGTCAGTCTCTCTTGTCGGGCTAGTCAGGGCATTAGAGGTAATTTAGACTGGTATCAGCAGAAACCAGGTGGAACTATTAAACTCCTGATCTACTCCACATCCAATTTAAATTCTGGTGTCCCATCAAGGTTCAGTGGCAGTGGGTCTGGGTCAGATTATTCTCTCACCATCAGCAGCCTAGAGTCTGAAGATTTTGCAGACTATTACTGTCTACAGCGTAATGCGTATCCT |
| IGKV12-38*01_129 | 129S1/SvImJ | Musmus IGKV12-38*01 F | 100 | 287 | GACATCCAGATGACTCAGTCTCCAGCCTCCCTGGCTGCATCTGTGGGAGAAACTGTCACCATCACATGTCGAGCAAGTGAGAACATTTACTACAGTTTAGCATGGTATCAGCAGAAGCAAGGGAAATCTCCTCAGCTCCTGATCTATAATGCAAACAGCTTGGAAGATGGTGTCCCATCGAGGTTCAGTGGCAGTGGATCTGGGACACAGTATTCTATGAAGATCAACAGCATGCAGCCTGAAGATACCGCAACTTATTTCTGTAAACAGGCTTATGACGTTCCTCC |

| | | | | | |
|---|---|---|---|---|---|
| IGKV5-37*01_129 | 129S1/SvImJ | Musmus IGKV5-37*01 F | 100 | 287 | GACATCCTGCTGACCCAGTCTCCAGCCACCCTGTCTGTGACTCCAGGAGAAACAGTCAGTCTTTCCTGTAGGGCCAGCCAGAGTATTTACAAGAACCTACACTGGTATCAACAGAAATCACATCGGTCTCCAAGGCTTCTCATCAAGTATGCTTCTGATTCCATCTCTGGGATCCCCTCCAGGTTCACTGGCAGTGGATCAGGGACAGATTACACTCTCAGTATCAACAGTGTGAAGCCCGAAGATGAAGGAATATATTACTGTCTTCAAGGTTACAGCACACCTTC |
| IGKV7-33*01_S6039_129 | 129S1/SvImJ | Musmus IGKV7-33*01 F | 100 | 305 | GACATTGTGATGACTCAGTCTCCAACTTTCCTTGCTGTGACAGCAAGTAAGAAGGTCACCATTAGTTGCACGGCCAGTGAGAGCCTTTATTCAAGCAAACACAAGGTGCACTACTTGGCTTGGTACCAGAAGAAACCAGAGCAATCTCCTAAACTGCTGATATACGGGGCATCCAACCGATACATTGGGGTCCCTGATCGCTTCACAGGCAGTGGATCTGGGACAGATTTCACTCTGACCATCAGCAGTGTACAGGTTGAAGACCTCACACATTATTACTGTGCACAGTTTTACAGCTATCCGCT |
| IGKV8-18*01_129 | 129S1/SvImJ | Musmus IGKV8-18*01 ORF | 100 | 310 | GACATTGTGATGACCCAGTCTCCATCCTCCCTGGCTGTGACAGCAGGAGAGAAGGTCACTATGAGATGCAAGTCCAGTCAGAGTCTTTTGTGGAGTGTAAACCAAAATAACTACTTATCCTGGTACCAGCAGAAACAAGGGCAGCCTCCTAAACTGCTTATCTATGGGGCATCCATTAGAGAATCTTGGGTCCCTGATCGATTCACAGGAAGTGGATCTGGGACAGACTTCACTCTCACCATTAGCAATGTGCATGCTGAAGACCTAGCAGTTTATTACTGTCAGCACAATCATGGCAGCTTTCTCCCCC |
| IGKV4-58*01_129 | 129S1/SvImJ | Musmus IGKV4-58*01 F | 100 | 292 | GAAAATGTGCTCACCCAGTCTCCAGCAATAATGGCTGCCTCTCTGGGGCAGAAGGTCACCATGACCTGCAGTGCCAGCTCAAGTGTAAGTTCCAGTTACTTGCACTGGTACCAGCAGAAGTCAGGCGCTTCCCCCAAACCCTTGATTCATAGGACATCCAACCTGGCTTCTGGAGTCCCAGCTCGCTTCAGTGGCAGTGGGTCTGGGACCTCTTACTCTCTCACAATCAGCAGCGTGGAGGCTGAAGATGATGCAACTTATTACTGCCAGCAGTGGAGTGGTTACCCATTCA |
| IGKV1-132*01_129 | 129S1/SvImJ | Musmus IGKV1-132*01 F | 100 | 302 | GATGTTGTGATGACCCAGACTCCACTGTCTTTGTCGGTTACCATTGGACAACCAGCCTCTATCTCTTGCAAGTCAAGTCAGAGCCTCTTATATAGTAATGGAAAGACATATTTGAATTGGTTACAACAGAGGCCTGGCCAGGCTCCAAAGCACCTAATGTATCAGGTGTCCAAACTGGACCCTGGCATCCCTGACAGGTTCAGTGGCAGTGGATCAGAAACAGATTTTACACTTAAAATCAGCAGAGTGGAGGCTGAAGATTTGGGAGTTTATTACTGCTTGCAAGGTACATATTATCCTCA |
| IGKV1-99*01_129 | 129S1/SvImJ | Musmus IGKV1-99*01 F | 100 | 302 | GATGTTGTTCTGACCCAAACTCCACTCTCTCTGCCTGTCAATATTGGAGATCAAGCCTCTATCTCTTGCAAGTCTACTAAGAGTCTTCTGAATAGTGATGGATTCACTTATTTGGACTGGTACCTGCAGAAGCCAGGCCAGTCTCCACAGCTCCTAATATATTTGGTTTCTAATCGATTTTCTGGAGTTCCAGACAGGTTCAGTGGCAGTGGGTCAGGAACAGATTTCACACTCAAGATCAGCAGAGTGGAGGCTGAGGATTTGGGAGTTTATTATTGCTTCCAGAGTAACTATCTTCCTCT |
| IGKV4-57*01_S9219_129 | 129S1/SvImJ | Musmus IGKV4-57-1*01 F | 100 | 286 | CAAATTGTTCTCACCCAGTCTCCAGCAATCATGTCTGCATCTCCAGGGGAGAAGGTCACCATAACCTGCAGTGCCAGCTCAAGTGTAAGTTACATGCACTGGTTCCAGCAGAAGCCAGGCACTTCTCCCAAACTCTGGATTTATAGCACATCCAACCTGGCTTCTGGAGTCCCTGCTCGCTTCAGTGGCAGTGGATCTGGGACCTCTTACTCTCTCACAATCAGCCGAATGGAGGCTGAAGATGCTGCCACTTATTACTGCCAGCAAAGGAGTAGTTACCCACTCA |

| | | | | | |
|---|---|---|---|---|---|
| IGKV4-51*01_129 | 129S1/SvImJ | Musmus IGKV4-51*01 F | 100 | 292 | GAAAATGTGCTCACCCAGTCTCCAGCAATAATGGCTGCCTCTCTGGGGGAGAAGGTCACCATGACCTGCAGTGCCAGCTCAAGTGTAAGTTCCAGCTACTTGCACTGGTACCAGCAGAAGTCAGGCACTTCCCCCAAACTCTGGATTTATGGCACATCCAACCTGGCTTCTGGAGTCCCAGCTCGCTTCAGTGGCAGTGGGGCTGGGATCTCTTACTCTCTCACAATCAGCAGCATGGAGGCTGAAAATGATGCAACTTATTACTGCCAGCAGTGGAGTGGTTACCCATTCA |
| IGKV4-71*01_129 | 129S1/SvImJ | Musmus IGKV4-71*01 F | 100 | 286 | CAAATTGTTCTCACCCAGTCTCCAGCAATCATGTCTGCATCTCCAGGGGAGAAGGTCACCATGACCTGCAGTGCCAGCTCAAGTGTAAGTTACATGCACTGGTACCAGCAGAAGCCAGGATCCTCCCCCAGACTCTGGATTTATTTAACATTCAACTTGGCTTCTGGAGTCCCTGCTCGCTTCAGTGGCAGTGGGTCTGGGACCTCTTACTCTCTCTCAATCAGCAGCATGGAGGCTGAAGATGCTGCCACTTATTACTGCCAGCAGTGGAGTAGTAACCCACCCA |
| IGKV6-14*01_129 | 129S1/SvImJ | Musmus IGKV6-14*01 F | 100 | 287 | GACATTGTGATGACCCAGTCTCAAAAATTCATGTCCACATCAGTAGGAGACAGGGTCAGCATCACCTGCAAGGCCAGTCAGAATGTTCGTACTGCTGTAGCCTGGTATCAACAGAAACCAGGGCAGTCTCCTAAAGCACTGATTTACTTGGCATCCAACCGGCACACTGGAGTCCCTGATCGCTTCACAGGCAGTGGATCTGGGACAGATTTCACTCTCACCATTAGCAATGTGCAATCTGAAGACCTGGCAGATTATTTCTGTCTGCAACATTGGAATTATCCTCT |
| IGKV4-52*01_S5019_129 | 129S1/SvImJ | Musmus IGKV4-52*01 ORF, or Musmus IGKV4-54*01 ORF | 100 | 286 | CAAATTGTTCTCACCCAGTCTCCAGCAATCCTGTCTGCATCTCCAGGGGAGAAGGTCACCATGACCTGCAGTGCCAGCTCAAGTGTAAGTTACATGTACAGGTACCAGCAGAAGCCAGGATCCTCACCCAAACCCTGGATTTATGGCACATCCAACCTGGCTTCTGGAGTCCCTGCTCGCTTCAGTGGCAGTGGATCTGGGACCTCTTATTCTCTCACAATCAGCAGCATGGAGGCTGAAGATGCTGCCACTTATTACTGCCAGCAATATCATAGTTACCCACTCA |
| IGKV4-74*01_S2922_129 | 129S1/SvImJ | Musmus IGKV4-74*01 F | 100 | 292 | CAAATTGTTCTCACCCAGTCTCCAGCAATCATGTCTGCATCTCTAGGGGAACGGGTCACCATGACCTGCACTGCCAGCTCAAGTGTAAGTTCCAGTTACTTGCACTGGTACCAGCAGAAGCCAGGATCCTCCCCCAAACTCTGGATTTATAGCACATCCAACCTGGCTTCTGGAGTCCCAGCTCGCTTCAGTGGCAGTGGGTCTGGGACCTCTTACTCTCTCACAATCAGCAGCATGGAGGCTGAAGATGCTGCCACTTATTACTGCCACCAGTATCATCGTTCCCCACTCA |
| IGKV6-15*01_AJ | A/J | Musmus IGKV6-15*01 F | 100 | 287 | GACATTGTGATGACCCAGTCTCAAAAATTCATGTCCACATCAGTAGGAGACAGGGTCAGCGTCACCTGCAAGGCCAGTCAGAATGTGGGTACTAATGTAGCCTGGTATCAACAGAAACCAGGGCAATCTCCTAAAGCACTGATTTACTCGGCATCCTACCGGTACAGTGGAGTCCCTGATCGCTTCACAGGCAGTGGATCTGGGACAGATTTCACTCTCACCATCAGCAATGTGCAGTCTGAAGACTTGGCAGAGTATTTCTGTCAGCAATATAACAGCTATCCTCT |
| IGKV8-30*01_AJ | A/J | Musmus IGKV8-30*01 F | 100 | 305 | GACATTGTGATGTCACAGTCTCCATCCTCCCTAGCTGTGTCAGTTGGAGAGAAGGTTACTATGAGCTGCAAGTCCAGTCAGAGCCTTTTATATAGTAGCAATCAAAAGAACTACTTGGCCTGGTACCAGCAGAAACCAGGGCAGTCTCCTAAACTGCTGATTTACTGGGCATCCACTAGGGAATCTGGGGTCCCTGATCGCTTCACAGGCAGTGGATCTGGGACAGATTTCACTCTCACCATCAGCAGTGTGAAGGCTGAAGACCTGGCAGTTTATTACTGTCAGCAATATTATAGCTATCCTCC |
| IGKV12-44*01_AJ | A/J | Musmus IGKV12-44*01 F | 100 | 287 | GACATCCAGATGACTCAGTCTCCAGCCTCCCTATCTGCATCTGTGGGAGAAACTGTCACCATCACATGTCGAGCAAGTGAGAATATTTACAGTTATTTAGCATGGTATCAGCAGAAACAGGGAAAATCTCCTCAGCTCCTGGTCTATAATGCAAAAACCTTAGCAGAAGGTGTGCCATCAAGGTTCAGTGGCAGTGGATCAGGCACACAGTTTTCTCTGAAGATCAACAGCCTGCAGCCTGAAGATTTTGGGAGTTATTACTGTCAACATCATTATGGTACTCCTCC |

| | | | | | |
|---|---|---|---|---|---|
| IGKV8-27*01_AJ | A/J | Musmus IGKV8-27*01 F | 100 | 300 | AACATTATGATGACACAGTCGCCATCATCTCTGGCTGTGTCTGCAGGAGAAAAGGTCACTATGAGCTGTAAGTCCAGTCAAAGTGTTTTATACAGTTCAAATCAGAAGAACTACTTGGCCTGGTACCAGCAGAAACCAGGGCAGTCTCCTAAACTGCTGATCTACTGGGCATCCACTAGGGAATCTGGTGTCCCTGATCGCTTCACAGGCAGTGGATCTGGGACAGATTTTACTCTTACCATCAGCAGTGTACAAGCTGAAGACCTGGCAGTTTATTACTGTCATCAATACCTCTCCTCG |
| IGKV6-20*01_AJ | A/J | Musmus IGKV6-20*01 F | 100 | 287 | AACATTGTAATGACCCAATCTCCCAAATCCATGTCCATGTCAGTAGGAGAGAGGGTCACCTTGAGCTGCAAGGCCAGTGAGAATGTGGGTACTTATGTATCCTGGTATCAACAGAAACCAGAGCAGTCTCCTAAACTGCTGATATACGGGGCATCCAACCGGTACACTGGGGTCCCCGATCGCTTCACAGGCAGTGGATCTGCAACAGATTTCACTCTGACCATCAGCAGTGTGCAGGCTGAAGACCTTGCAGATTATCACTGTGGACAGAGTTACAGCTATCCTCC |
| IGKV6-23*01_AJ | A/J | Musmus IGKV6-23*01 F | 100 | 287 | GACATTGTGATGACCCAGTCTCACAAATTCATGTCCACATCAGTAGGAGACAGGGTCAGCATCACCTGCAAGGCCAGTCAGGATGTGGGTACTGCTGTAGCCTGGTATCAACAGAAACCAGGGCAATCTCCTAAACTACTGATTTACTGGGCATCCACCCGGCACACTGGAGTCCCTGATCGCTTCACAGGCAGTGGATCTGGGACAGATTTCACTCTCACCATTAGCAATGTGCAGTCTGAAGACTTGGCAGATTATTTCTGTCAGCAATATAGCAGCTATCCTCT |
| IGKV8-21*01_AJ | A/J | Musmus IGKV8-21*01 F | 100 | 302 | GACATTGTGATGTCACAGTCTCCATCCTCCCTGGCTGTGTCAGCAGGAGAGAAGGTCACTATGAGCTGCAAATCCAGTCAGAGTCTGCTCAACAGTAGAACCCGAAAGAACTACTTGGCTTGGTACCAGCAGAAACCAGGGCAGTCTCCTAAACTGCTGATCTACTGGGCATCCACTAGGGAATCTGGGGTCCCTGATCGCTTCACAGGCAGTGGATCTGGGACAGATTTCACTCTCACCATCAGCAGTGTGCAGGCTGAAGACCTGGCAGTTTATTACTGCAAGCAATCTTATAATCTTCC |
| IGKV1-135*01_AJ | A/J | Musmus IGKV1-135*01 F | 100 | 302 | GATGTTGTGATGACCCAGACTCCACTCACTTTGTCGGTTACCATTGGACAACCAGCCTCCATCTCTTGCAAGTCAAGTCAGAGCCTCTTAGATAGTGATGGAAAGACATATTTGAATTGGTTGTTACAGAGGCCAGGCCAGTCTCCAAAGCGCCTAATCTATCTGGTGTCTAAACTGGACTCTGGAGTCCCTGACAGGTTCACTGGCAGTGGATCAGGGACAGATTTCACACTGAAAATCAGCAGAGTGGAGGCTGAGGATTTGGGAGTTTATTATTGCTGGCAAGGTACACATTTTCCTCA |
| IGKV4-59*01_AJ | A/J | Musmus IGKV4-59*01 F | 100 | 286 | CAAATTGTTCTCACCCAGTCTCCAGCAATCATGTCTGCATCTCCAGGGGAGAAGGTCACCATGACCTGCAGTGCCAGCTCAAGTGTAAGTTACATGCACTGGTACCAGCAGAAGTCAGGCACCTCCCCCAAAAGATGGATTTATGACACATCCAAACTGGCTTCTGGAGTCCCTGCTCGCTTCAGTGGCAGTGGGTCTGGGACCTCTTACTCTCTCACAATCAGCAGCATGGAGGCTGAAGATGCTGCCACTTATTACTGCCAGCAGTGGAGTAGTAACCCACCCA |
| IGKV1-117*01_AJ | A/J | Musmus IGKV1-117*01 F | 100 | 302 | GATGTTTTGATGACCCAAACTCCACTCTCCCTGCCTGTCAGTCTTGGAGATCAAGCCTCCATCTCTTGCAGATCTAGTCAGAGCATTGTACATAGTAATGGAAACACCTATTTAGAATGGTACCTGCAGAAACCAGGCCAGTCTCCAAAGCTCCTGATCTACAAAGTTTCCAACCGATTTTCTGGGGTCCCAGACAGGTTCAGTGGCAGTGGATCAGGGACAGATTTCACACTCAAGATCAGCAGAGTGGAGGCTGAGGATCTGGGAGTTTATTACTGCTTTCAAGGTTCACATGTTCCTCC |

| | | | | | |
|---|---|---|---|---|---|
| IGKV8-19*01_AJ | A/J | Musmus IGKV8-19*01 F | 100 | 305 | GACATTGTGATGACACAGTCTCCATCCTCCCTGACTGTGACAGCAGGAGAGAAGGTCACTATGAGCTGCAAGTCCAGTCAGAGTCTGTTAAACAGTGGAAATCAAAAGAACTACTTGACCTGGTACCAGCAGAAACCAGGGCAGCCTCCTAAACTGTTGATCTACTGGGCATCCACTAGGGAATCTGGGGTCCCTGATCGCTTCACAGGCAGTGGATCTGGAACAGATTTCACTCTCACCATCAGCAGTGTGCAGGCTGAAGACCTGGCAGTTTATTACTGTCAGAATGATTATAGTTATCCTCC |
| IGKV6-32*01_AJ | A/J | Musmus IGKV6-32*01 F | 100 | 287 | AGTATTGTGATGACCCAGACTCCCAAATTCCTGCTTGTATCAGCAGGAGACAGGGTTACCATAACCTGCAAGGCCAGTCAGAGTGTGAGTAATGATGTAGCTTGGTACCAACAGAAGCCAGGGCAGTCTCCTAAACTGCTGATATACTATGCATCCAATCGCTACACTGGAGTCCCTGATCGCTTCACTGGCAGTGGATATGGGACGGATTTCACTTTCACCATCAGCACTGTGCAGGCTGAAGACCTGGCAGTTTATTTCTGTCAGCAGGATTATAGCTCTCCTCC |
| IGKV8-24*01_AJ | A/J | Musmus IGKV8-24*01 F | 100 | 305 | GACATTGTGATGACACAGTCTCCATCCTCCCTGGCTATGTCAGTAGGACAGAAGGTCACTATGAGCTGCAAGTCCAGTCAGAGCCTTTTAAATAGTAGCAATCAAAAGAACTATTTGGCCTGGTACCAGCAGAAACCAGGACAGTCTCCTAAACTTCTGGTATACTTTGCATCCACTAGGGAATCTGGGGTCCCTGATCGCTTCATAGGCAGTGGATCTGGGACAGATTTCACTCTTACCATCAGCAGTGTGCAGGCTGAAGACCTGGCAGATTACTTCTGTCAGCAACATTATAGCACTCCTCC |
| IGKV2-137*01_AJ | A/J | Musmus IGKV2-137*01 F | 100 | 302 | GATATTGTGATGACTCAGGCTGCACCCTCTGTACCTGTCACTCCTGGAGAGTCAGTATCCATCTCCTGCAGGTCTAGTAAGAGTCTCCTGCATAGTAATGGCAACACTTACTTGTATTGGTTCCTGCAGAGGCCAGGCCAGTCTCCTCAGCTCCTGATATATCGGATGTCCAACCTTGCCTCAGGAGTCCCAGACAGGTTCAGTGGCAGTGGGTCAGGAACTGCTTTCACACTGAGAATCAGTAGAGTGGAGGCTGAGGATGTGGGTGTTTATTACTGTATGCAACATCTAGAATATCCTTT |
| IGKV1-110*01_AJ | A/J | Musmus IGKV1-110*01 F | 100 | 302 | GATGTTGTGATGACCCAAACTCCACTCTCCCTGCCTGTCAGTCTTGGAGATCAAGCCTCCATCTCTTGCAGATCTAGTCAGAGCCTTGTACACAGTAATGGAAACACCTATTTACATTGGTACCTGCAGAAGCCAGGCCAGTCTCCAAAGCTCCTGATCTACAAAGTTTCCAACCGATTTTCTGGGGTCCCAGACAGGTTCAGTGGCAGTGGATCAGGGACAGATTTCACACTCAAGATCAGCAGAGTGGAGGCTGAGGATCTGGGAGTTTATTTCTGCTCTCAAAGTACACATGTTCCTCC |
| IGKV4-57*01_S6684_AJ | A/J | Musmus IGKV4-57-1*01 F | 100 | 286 | CAAATTGTTCTCACCCAGTCTCCAGCAATCATGTCTGCATCTCCAGGGGAGAAGGTCACCATAACCTGCAGTGCCAGCTCAAGTGTAAGTTACATGCACTGGTTCCAGCAGAAGCCAGGCACTTCTCCCAAACTCTGGATTTATAGCACATCCAACCTGGCTTCTGGAGTCCCTGCTCGCTTCAGTGGCAGTGGATCTGGGACCTCTTACTCTCTCACAATCAGCCGAATGGAGGCTGAAGATGCTGCCACTTATTACTGCCAGCAAAGGAGTAGTTACCCGCTCA |
| IGKV8-28*01_AJ | A/J | Musmus IGKV8-28*01 F | 100 | 305 | GACATTGTGATGACACAGTCTCCATCCTCCCTGAGTGTGTCAGCAGGAGAGAAGGTCACTATGAGCTGCAAGTCCAGTCAGAGTCTGTTAAACAGTGGAAATCAAAAGAACTACTTGGCCTGGTACCAGCAGAAACCAGGGCAGCCTCCTAAACTGTTGATCTACGGGGCATCCACTAGGGAATCTGGGGTCCCTGATCGCTTCACAGGCAGTGGATCTGGAACCGATTTCACTCTTACCATCAGCAGTGTGCAGGCTGAAGACCTGGCAGTTTATTACTGTCAGAATGATCATAGTTATCCTCC |

| | | | | | |
|---|---|---|---|---|---|
| IGKV6-17*01_AJ | A/J | Musmus IGKV6-17*01 F | 100 | 287 | GACATTGTGATGACCCAGTCTCACAAATTCATGTCCACATCAGTAGGAGACAGGGTCAGCATCACCTGCAAGGCCAGTCAGGATGTGAGTACTGCTGTAGCCTGGTATCAACAGAAACCAGGACAATCTCCTAAACTACTGATTTACTCGGCATCCTACCGGTACACTGGAGTCCCTGATCGCTTCACTGGCAGTGGATCTGGGACGGATTTCACTTTCACCATCAGCAGTGTGCAGGCTGAAGACCTGGCAGTTTATTACTGTCAGCAACATTATAGTACTCCTCC |
| IGKV4-57-1*01_AJ | A/J | Musmus IGKV4-57*01 F | 100 | 292 | GAAAATGTGCTCACCCAGTCTCCAGCAATCATGTCTGCATCTCCAGGGGAAAAGGTCACCATGACCTGCAGGGCCAGCTCAAGTGTAAGTTCCAGTTACTTGCACTGGTACCAGCAGAAGTCAGGTGCCTCCCCCAAACTCTGGATTTATAGCACATCCAACTTGGCTTCTGGAGTCCCTGCTCGCTTCAGTGGCAGTGGGTCTGGGACCTCTTACTCTCTCACAATCAGCAGTGTGGAGGCTGAAGATGCTGCCACTTATTACTGCCAGCAGTACAGTGGTTACCCACTCA |
| IGKV10-96*01_AJ | A/J | Musmus IGKV10-96*01 F | 100 | 286 | GATATCCAGATGACACAGACTACATCCTCCCTGTCTGCCTCTCTGGGAGACAGAGTCACCATCAGTTGCAGGGCAAGTCAGGACATTAGCAATTATTTAAACTGGTATCAGCAGAAACCAGATGGAACTGTTAAACTCCTGATCTACTACACATCAAGATTACACTCAGGAGTCCCATCAAGGTTCAGTGGCAGTGGGTCTGGAACAGATTATTCTCTCACCATTAGCAACCTGGAGCAAGAAGATATTGCCACTTACTTTTGCCAACAGGGTAATACGCTTCCTC |
| IGKV12-46*01_AJ | A/J | Musmus IGKV12-46*01 F | 100 | 287 | GACATCCAGATGACTCAGTCTCCAGCCTCCCTATCTGTATCTGTGGGAGAAACTGTCACCATCACATGTCGAGCAAGTGAGAATATTTACAGTAATTTAGCATGGTATCAGCAGAAACAGGGAAAATCTCCTCAGCTCCTGGTCTATGCTGCAACAAACTTAGCAGATGGTGTGCCATCAAGGTTCAGTGGCAGTGGATCAGGCACACAGTATTCCCTCAAGATCAACAGCCTGCAGTCTGAAGATTTTGGGAGTTATTACTGTCAACATTTTTGGGGTACTCCTCC |
| IGKV4-70*01_AJ | A/J | Musmus IGKV4-70*01 F | 100 | 282 | CAAATTGTTCTCACCCAGTCTCCAGCAATCATGTCTGCATCTCCAGGGGAGAAGGTCACCATGACCTGCAGTGCCAGCTCAAGTATAAGTTACATGCACTGGTACCAGCAGAAGCCAGGCACCTCCCCCAAAAGATGGATTTATGACACATCCAAACTGGCTTCTGGAGTCCCTGCTCGCTTCAGTGGCAGTGGGTCTGGGACCTCTTATTCTCTCACAATCAGCAGCATGGAGGCTGAAGATGCTGCCACTTATTACTGCCATCAGCGGAGTAGTTACCCA |
| IGKV12-89*01_AJ | A/J | Musmus IGKV12-89*01 F | 100 | 287 | GACATCCAGATGACTCAGTCTCCAGCTTCACTGTCTGCATCTGTGGGAGAAACTGTCACCATCACATGTGGAGCAAGTGAGAATATTTACGGTGCTTTAAATTGGTATCAGCGGAAACAGGGAAAATCTCCTCAGCTCCTGATCTATGGTGCAACCAACTTGGCAGATGGCATGTCATCGAGGTTCAGTGGCAGTGGATCTGGTAGACAGTATTCTCTCAAGATCAGTAGCCTGCATCCTGACGATGTTGCAACGTATTACTGTCAAAATGTGTTAAGTACTCCTCC |
| IGKV12-41*01_AJ | A/J | Musmus IGKV12-41*01 F | 100 | 287 | GACATCCAGATGACTCAGTCTCCAGCCTCCCTATCTGCATCTGTGGGAGAAACTGTCACCATCACATGTCGAGCAAGTGGGAATATTCACAATTATTTAGCATGGTATCAGCAGAAACAGGGAAAATCTCCTCAGCTCCTGGTCTATAATGCAAAAACCTTAGCAGATGGTGTGCCATCAAGGTTCAGTGGCAGTGGATCAGGAACACAATATTCTCTCAAGATCAACAGCCTGCAGCCTGAAGATTTTGGGAGTTATTACTGTCAACATTTTTGGAGTACTCCTCC |
| IGKV6-13*01_AJ | A/J | Musmus IGKV6-13*01 F | 100 | 285 | GACATTGTGATGACCCAGTCTCAAAAATTCATGTCCACATCAGTAGGAGACAGGGTCAGCATCACCTGCAAGGCCAGTCAGAATGTGGGTACTGCTGTAGCCTGGTATCAACAGAAACCAGGACAATCTCCTAAACTACTGATTTACTCGGCATCCAATCGGTACACTGGAGTCCCTGATCGCTTCACAGGCAGTGGATCTGGGACAGATTTCACTCTCACCATCAGCAATATGCAGTCTGAAGACCTGGCAGATTATTTCTGCCAGCAATATAGCAGCTATCCT |

| | | | | | |
|---|---|---|---|---|---|
| IGKV5-43*01_AJ | A/J | Musmus IGKV5-43*01 F | 100 | 287 | GATATTGTGCTAACTCAGTCTCCAGCCACCCTGTCTGTGACTCCAGGAGATAGCGTCAGTCTTTCCTGCAGGGCCAGCCAAAGTATTAGCAACAACCTACACTGGTATCAACAAAAATCACATGAGTCTCCAAGGCTTCTCATCAAGTATGCTTCCCAGTCCATCTCTGGGATCCCCTCCAGGTTCAGTGGCAGTGGATCAGGGACAGATTTCACTCTCAGTATCAACAGTGTGGAGACTGAAGATTTTGGAATGTATTTCTGTCAACAGAGTAACAGCTGGCCTCA |
| IGKV4-55*01_AJ | A/J | Musmus IGKV4-55*01 F | 100 | 286 | CAAATTGTTCTCACCCAGTCTCCAGCAATCATGTCTGCATCTCCAGGGGAGAAGGTCACCATGACCTGCAGTGCCAGCTCAAGTGTAAGTTACATGTACTGGTACCAGCAGAAGCCAGGATCCTCCCCCAGACTCCTGATTTATGACACATCCAACCTGGCTTCTGGAGTCCCTGTTCGCTTCAGTGGCAGTGGGTCTGGGACCTCTTACTCTCTCACAATCAGCCGAATGGAGGCTGAAGATGCTGCCACTTATTACTGCCAGCAGTGGAGTAGTTACCCACCCA |
| IGKV15-103*01_AJ | A/J | Musmus IGKV15-103*01 ORF | 100 | 287 | GACATCCAGATGAACCAGTCTCCATCCAGTCTGTCTGCATCCCTTGGAGACACAATTACCATCACTTGCCATGCCAGTCAGAACATTAATGTTTGGTTAAGCTGGTACCAGCAGAAACCAGGAAATATTCCTAAACTATTGATCTATAAGGCTTCCAACTTGCACACAGGCGTCCCATCAAGGTTTAGTGGCAGTGGATCTGGAACAGGTTTCACATTAACCATCAGCAGCCTGCAGCCTGAAGACATTGCCACTTACTACTGTCAACAGGGTCAAAGTTATCCTCT |
| IGKV9-120*01_S2628_AJ | A/J | Musmus IGKV9-120*02 F | 100 | 285 | GACATCCAGATGACCCAGTCTCCATCCTCCTTATCTGCCTCTCTGGGAGAAAGAGTCAGTCTCACTTGTCGGGCAAGTCAGGACATTGGTAGTAGCTTAAACTGGCTTCAGCAGGAACCAGATGGAACTATTAAACGCCTGATCTACGCCACATCCAGTTTAGATTCTGGTGTCCCCAAAAGGTTCAGTGGCAGTAGGTCTGGGTCAGATTATTCTCTCACCATCAGCAGCCTTGAGTCTGAAGATTTTGTAGACTATTACTGTCTACAATATGCTAGTTCTCCT |
| IGKV19-93*01_AJ | A/J | Musmus IGKV19-93*01 F | 100 | 287 | GACATCCAGATGACACAGTCTCCATCCTCACTGTCTGCATCTCTGGGAGGCAAAGTCACCATCACTTGCAAGGCAAGCCAAGACATTAACAAGTATATAGCTTGGTACCAACACAAGCCTGGAAAAGGTCCTAGGCTGCTCATACATTACACATCTACATTACAGCCAGGCATCCCATCAAGGTTCAGTGGAAGTGGGTCTGGGAGAGATTATTCCTTCAGCATCAGCAACCTGGAGCCTGAAGATATTGCAACTTATTATTGTCTACAGTATGATAATCTTCTACC |
| IGKV4-91*01_S5997_AJ | A/J | Musmus IGKV4-91*01 F | 100 | 292 | GAAATTGTGCTCACCCAGTCTCCAACCACCATGGCTGCATCTCCCGGGGAGAAGATCACTATCACCTGCAGTGCCAGCTCAAGTATAAGTTCCAATTACTTGCATTGGTATCAGCAGAAGCCAGGATTCTCCCCTAAACTCTTGATTTATAGGACATCCAATCTGGCTTCTGGAGTCCCAGCTCGCTTCAGTGGCAGTGGGTCTGGGACCTCTTACTCTCTCACAATTGGCACCATGGAGGCTGAAGATGTTGCCACTTACTACTGCCAGCAGGGTAGTAGTATACCACTCA |
| IGKV5-48*01_AJ | A/J | Musmus IGKV5-48*01 F | 100 | 287 | GACATCTTGCTGACTCAGTCTCCAGCCATCCTGTCTGTGAGTCCAGGAGAAAGAGTCAGTTTCTCCTGCAGGGCCAGTCAGAGCATTGGCACAAGCATACACTGGTATCAGCAAAGAACAAATGGTTCTCCAAGGCTTCTCATAAAGTATGCTTCTGAGTCTATCTCTGGGATCCCTTCCAGGTTTAGTGGCAGTGGATCAGGGACAGATTTTACTCTTAGCATCAACAGTGTGGAGTCTGAAGATATTGCAGATTATTACTGTCAACAAAGTAATAGCTGGCCAAC |
| IGKV4-68*01_AJ | A/J | Musmus IGKV4-68*01 F | 100 | 286 | CAAATTGTTCTCACCCAGTCTCCAGCACTCATGTCTGCATCTCCAGGGGAGAAGGTCACCATGACCTGCAGTGCCAGCTCAAGTGTAAGTTACATGTACTGGTACCAGCAGAAGCCAAGATCCTCCCCCAAACCCTGGATTTATCTCACATCCAACCTGGCTTCTGGAGTCCCTGCTCGCTTCAGTGGCAGTGGGTCTGGGACCTCTTACTCTCTCACAATCAGCAGCATGGAGGCTGAAGATGCTGCCACTTATTACTGCCAGCAGTGGAGTAGTAACCCACCCA |

| | | | | | |
|---|---|---|---|---|---|
| IGKV3-4*01_AJ | A/J | Musmus IGKV3-4*01 F | 100 | 299 | GACATTGTGCTGACCCAATCTCCAGCTTCTTTGGCTGTGTCTCTAGGGCAGAGGGCCACCATCTCCTGCAAGGCCAGCCAAAGTGTTGATTATGATGGTGATAGTTATATGAACTGGTACCAACAGAAACCAGGACAGCCACCCAAACTCCTCATCTATGCTGCATCCAATCTAGAATCTGGGATCCCAGCCAGGTTTAGTGGCAGTGGGTCTGGGACAGACTTCACCCTCAACATCCATCCTGTGGAGGAGGAGGATGCTGCAACCTATTACTGTCAGCAAAGTAATGAGGATCCTCC |
| IGKV3-12*01_AJ | A/J | Musmus IGKV3-12*01 F | 100 | 299 | GACATTGTGCTGACACAGTCTCCTGCTTCCTTAGCTGTATCTCTGGGGCAGAGGGCCACCATCTCATGCAGGGCCAGCAAAAGTGTCAGTACATCTGGCTATAGTTATATGCACTGGTACCAACAGAAACCAGGACAGCCACCCAAACTCCTCATCTATCTTGCATCCAACCTAGAATCTGGGGTCCCTGCCAGGTTCAGTGGCAGTGGGTCTGGGACAGACTTCACCCTCAACATCCATCCTGTGGAGGAGGAGGATGCTGCAACCTATTACTGTCAGCACAGTAGGGAGCTTCCTCC |
| IGKV4-80*01_AJ | A/J | Musmus IGKV4-80*01 F | 100 | 282 | CAAATTGTTCTCACCCAGTCTCCAGCAATCATGTCTGCATCTCTAGGGGAGGAGATCACCCTAACCTGCAGTGCCAGCTCGAGTGTAAGTTACATGCACTGGTACCAGCAGAAGTCAGGCACTTCTCCCAAACTCTTGATTTATAGCACATCCAACCTGGCTTCTGGAGTCCCTTCTCGCTTCAGTGGCAGTGGGTCTGGGACCTTTTATTCTCTCACAATCAGCAGTGTGGAGGCTGAAGATGCTGCCGATTATTACTGCCATCAGTGGAGTAGTTATCCA |
| IGKV8-16*01_AJ | A/J | Musmus IGKV8-16*01 F | 100 | 305 | GAAATTGTGTTGACCCAGTCTATACCATCCCTGACTGTGTCAGCAGGAGAGAGGGTCACTATCAGCTGCAAATCCAATCAGAATCTTTTATGGAGTGGAAACCAAAGGTACTGTTTGGTCTGGCACCAGTGGAAACCAGGGCAAACTCCTACACCGTTGATCACCTGGACATCTGATAGGTACTCTGGAGTCCCTGATCGTTTCATAGGCAGTGGATCTGTGACAGATTTCACTCTGACCATCAGCAGTGTGCAGGCTGAAGATGTGGCAGTTTATTTCTGTCAGCAGCATTTACACATTCCTCC |
| IGKV14-100*01_AJ | A/J | Musmus IGKV14-100*01 F | 100 | 287 | GACATCCTGATGACCCAATCTCCATCCTCCATGTCTGTATCTCTGGGAGACACAGTCAGCATCACTTGCCATGCAAGTCAGGGCATTAGCAGTAATATAGGGTGGTTGCAGCAGAAACCAGGGAAATCATTTAAGGGCCTGATCTATCATGGAACCAACTTGGAAGATGGAGTTCCATCAAGGTTCAGTGGCAGTGGATCTGGAGCAGATTATTCTCTCACCATCAGCAGCCTGGAATCTGAAGATTTTGCAGACTATTACTGTGTACAGTATGCTCAGTTTCCTCC |
| IGKV3-5*01_AJ | A/J | Musmus IGKV3-5*01 F | 100 | 299 | GACATTGTGCTGACCCAATCTCCAGCTTCTTTGGCTGTGTCTCTAGGGCAGAGGGCCACCATATCCTGCAGAGCCAGTGAAAGTGTTGATAGTTATGGCAATAGTTTTATGCACTGGTACCAGCAGAAACCAGGACAGCCACCCAAACTCCTCATCTATCGTGCATCCAACCTAGAATCTGGGATCCCTGCCAGGTTCAGTGGCAGTGGGTCTAGGACAGACTTCACCCTCACCATTAATCCTGTGGAGGCTGATGATGTTGCAACCTATTACTGTCAGCAAAGTAATGAGGATCCTCC |
| IGKV3-2*01_AJ | A/J | Musmus IGKV3-2*01 F | 100 | 299 | GACATTGTGCTGACCCAATCTCCAGCTTCTTTGGCTGTGTCTCTAGGGCAGAGGGCCACCATCTCCTGCAGAGCCAGCGAAAGTGTTGATAATTATGGCATTAGTTTTATGAACTGGTTCCAACAGAAACCAGGACAGCCACCCAAACTCCTCATCTATGCTGCATCCAACCAAGGATCCGGGGTCCCTGCCAGGTTTAGTGGCAGTGGGTCTGGGACAGACTTCAGCCTCAACATCCATCCTATGGAGGAGGATGATACTGCAATGTATTTCTGTCAGCAAAGTAAGGAGGTTCCTCC |

| | | | | | |
|---|---|---|---|---|---|
| IGKV10-94*01_AJ | A/J | Musmus IGKV10-94*01 F | 100 | 287 | GATATCCAGATGACACAGACTACATCCTCCCTGTCTGCCTCTCTGGGAGACAGAGTCACCATCAGTTGCAGTGCAAGTCAGGGCATTAGCAATTATTTAAACTGGTATCAGCAGAAACCAGATGGAACTGTTAAACTCCTGATCTATTACACATCAAGTTTACACTCAGGAGTCCCATCAAGGTTCAGTGGCAGTGGGTCTGGGACAGATTATTCTCTCACCATCAGCAACCTGGAACCTGAAGATATTGCCACTTACTATTGTCAGCAGTATAGTAAGCTTCCTCC |
| IGKV16-104*01_AJ | A/J | Musmus IGKV16-104*01 F | 100 | 288 | GATGTCCAGATAACCCAGTCTCCATCTTATCTTGCTGCATCTCCTGGAGAAACCATTACTATTAATTGCAGGGCAAGTAAGAGCATTAGCAAATATTTAGCCTGGTATCAAGAGAAACCTGGGAAAACTAATAAGCTTCTTATCTACTCTGGATCCACTTTGCAATCTGGAATTCCATCAAGGTTCAGTGGCAGTGGATCTGGTACAGATTTCACTCTCACCATCAGTAGCCTGGAGCCTGAAGATTTTGCAATGTATTACTGTCAACAGCATAATGAATACCCGTAC |
| IGKV6-29*01_AJ | A/J | Musmus IGKV6-29*01 F | 100 | 287 | AACATTGTAATGACCCAATCTCCCAAATCCATGTCCATGTCAGTAGGAGAGAGGGTCACCTTGAGCTGCAAGGCCAGTGAGAATGTGGGTACTTATGTATCCTGGTATCAACAGAAACCAGAGCAGTCTCCTAAACTGCTGATATACGGGGCATCCAACCGGTACCCTGGGGTCCCTGATCGCTTCACAGGCAGTGGATCTGCAACAGATTTCACTCTGACCATCAGCAGTCTGCAGGCTGAAGACCTTGCAGATTATCACTGTGGACAGGGTTACAGCTATCTTCC |
| IGKV3-7*01_S5296_AJ | A/J | Musmus IGKV3-7*02 F | 100 | 299 | GACATTGTGCTGACACAGTCTCCTGCTTCCTTAGCTGTATCTCTGGGGCAGAGGGCCACCATCTCATGCAGGGCCAGCCAAAGTGTCAGTACATCTAGCTATAGTTATATGCACTGGTACCAACAGAAACCAGGACAGCCACCCAAACTCCTCATCAAGTATGCATCCAACCTAGAATCTGGGGTCCCTGCCAGGTTCAGTGGCAGTGGGTCTGGGACAGACTTCACCCTCAACATCCATCCTGTGGAGGAGGAGGATACTGCAACATATTACTGTCAGCACAGTTGGGAGATTCCTCC |
| IGKV5-39*01_AJ | A/J | Musmus IGKV5-39*01 F | 100 | 287 | GACATTGTGATGACTCAGTCTCCAGCCACCCTGTCTGTGACTCCAGGAGATAGAGTCTCTCTTTCCTGCAGGGCCAGCCAGAGTATTAGCGACTACTTACACTGGTATCAACAAAAATCACATGAGTCTCCAAGGCTTCTCATCAAATATGCTTCCCAATCCATCTCTGGGATCCCCTCCAGGTTCAGTGGCAGTGGATCAGGGTCAGATTTCACTCTCAGTATCAACAGTGTGGAACCTGAAGATGTTGGAGTGTATTACTGTCAAAATGGTCACAGCTTTCCTCC |
| IGKV6-25*01_AJ | A/J | Musmus IGKV6-25*01 F | 100 | 287 | GACATTGTGATGACCCAGTCTCACAAATTCATGTCCACATCAGTAGGAGACAGGGTCAGCATCACCTGCAAGGCCAGTCAGGATGTGAGTACTGCTGTAGCCTGGTATCAACAAAAACCAGGGCAATCTCCTAAACTACTGATTTACTGGGCATCCACCCGGCACACTGGAGTCCCTGATCGCTTCACAGGCAGTGGATCTGGGACAGATTATACTCTCACCATCAGCAGTGTGCAGGCTGAAGACCTGGCACTTTATTACTGTCAGCAACATTATAGCACTCCTCC |
| IGKV17-127*01_AJ | A/J | Musmus IGKV17-127*01 F | 100 | 287 | GAAACAACTGTGACCCAGTCTCCAGCATCCCTGTCCGTGGCTACAGGAGAAAAAGTCACTATCAGATGCATAACCAGCACTGATATTGATGATGATATGAACTGGTACCAGCAGAAGCCAGGGGAACCTCCTAAGCTCCTTATTTCAGAAGGCAATACTCTTCGTCCTGGAGTCCCATCCCGATTCTCCAGCAGTGGCTATGGCACAGATTTTGTTTTTACAATTGAAAACACGCTCTCAGAAGATGTTGCAGATTACTACTGTTTGCAAAGTGATAACATGCCTCT |
| IGKV14-126*01_AJ | A/J | Musmus IGKV14-126*01 F | 100 | 287 | GACATCAAGATGACCCAGTCTCCATCCTCCATGTATGCATCGCTGGGAGAGAGAGTCACTATCACTTGCAAGGCGAGTCAGGACATTAAAAGCTATTTAAGCTGGTACCAGCAGAAACCATGGAAATCTCCTAAGACCCTGATCTATTATGCAACAAGCTTGGCAGATGGGGTCCCATCAAGATTCAGTGGCAGTGGATCTGGGCAAGATTATTCTCTAACCATCAGCAGCCTGGAGTCTGACGATACAGCAACTTATTACTGTCTACAGCATGGTGAGAGCCCTCC |

| | | | | | |
|---|---|---|---|---|---|
| IGKV4-74*01_S2922_AJ | A/J | Musmus IGKV4-74*01 F | 100 | 292 | CAAATTGTTCTCACCCAGTCTCCAGCAATCATGTCTGCATCTCTAGGGGAACGGGTCACCATGACCTGCACTGCCAGCTCAAGTGTAAGTTCCAGTTACTTGCACTGGTACCAGCAGAAGCCAGGATCCTCCCCCAAACTCTGGATTTATAGCACATCCAACCTGGCTTCTGGAGTCCCAGCTCGCTTCAGTGGCAGTGGGTCTGGGACCTCTTACTCTCTCACAATCAGCAGCATGGAGGCTGAAGATGCTGCCACTTATTACTGCCACCAGTATCATCGTTCCCCACTCA |
| IGKV13-84*01_AJ | A/J | Musmus IGKV13-84*01 F | 100 | 287 | GACATCCAGATGACACAATCTTCATCCTCCTTTTCTGTATCTCTAGGAGACAGAGTCACCATTACTTGCAAGGCAAGTGAGGACATATATAATCGGTTAGCCTGGTATCAGCAGAAACCAGGAAATGCTCCTAGGCTCTTAATATCTGGTGCAACCAGTTTGGAAACTGGGGTTCCTTCAAGATTCAGTGGCAGTGGATCTGGAAAGGATTACACTCTCAGCATTACCAGTCTTCAGACTGAAGATGTTGCTACTTATTACTGTCAACAGTATTGGAGTACTCCTCC |
| IGKV12-98*01_AJ | A/J | Musmus IGKV12-98*01 F | 100 | 287 | GACATTCAGATGACCCAGTCTCCTGCCTCCCAGTCTGCATCTCTGGGAGAAAGTGTCACCATCACATGCCTGGCAAGTCAGACCATTGGTACATGGTTAGCATGGTATCAGCAGAAACCAGGGAAATCTCCTCAGCTCCTGATTTATGCTGCAACCAGCTTGGCAGATGGGGTCCCATCAAGGTTCAGTGGTAGTGGATCTGGCACAAAATTTTCTTTCAAGATCAGCAGCCTACAGGCTGAAGATTTTGTAAGTTATTACTGTCAACAACTTTACAGTACTCCTCT |
| IGKV4-63*01_AJ | A/J | Musmus IGKV4-63*01 F | 100 | 286 | GAAAATGTTCTCACCCAGTCTCCAGCAATCATGTCTGCATCTCCAGGGGAAAAGGTCACCATGACCTGCAGTGCCAGCTCAAGTGTAAGTTACATGCACTGGTACCAGCAGAAGTCAAGCACCTCCCCCAAACTCTGGATTTATGACACATCCAAACTGGCTTCTGGAGTCCCAGGTCGCTTCAGTGGCAGTGGGTCTGGAAACTCTTACTCTCTCACGATCAGCAGCATGGAGGCTGAAGATGTTGCCACTTATTACTGTTTTCAGGGGAGTGGGTACCCACTCA |
| IGKV4-72*01_S3540_AJ | A/J | Musmus IGKV4-72*01 F | 100 | 286 | CAAATTGTTCTCTCCCAGTCTCCAGCAATCCTGTCTGCATCTCCAGGGGAGAAGGTCACAATGACTTGCAGGGCCAGCTCAAGTGTAAGTTACATGCACTGGTACCAGCAGAAGCCAGGATCCTCCCCCAAACCCTGGATTTATGCCACATCCAACCTGGCTTCTGGAGTCCCTGCTCGCTTCAGTGGCAGTGGGTCTGGGACCTCTTACTCTCTCACAATCAGCAGAGTGGAGGCTGAAGATGCTGCCACTTATTACTGCCAGCAGTGGAGTAGTAACCCACTCA |
| IGKV2-109*01_AJ | A/J | Musmus IGKV2-109*01 F | 100 | 302 | GATATTGTGATGACGCAGGCTGCATTCTCCAATCCAGTCACTCTTGGAACATCAGCTTCCATCTCCTGCAGGTCTAGTAAGAGTCTCCTACATAGTAATGGCATCACTTATTTGTATTGGTATCTGCAGAAGCCAGGCCAGTCTCCTCAGCTCCTGATTTATCAGATGTCCAACCTTGCCTCAGGAGTCCCAGACAGGTTCAGTAGCAGTGGGTCAGGAACTGATTTCACACTGAGAATCAGCAGAGTGGAGGCTGAGGATGTGGGTGTTTATTACTGTGCTCAAAATCTAGAACTTCCTCC |
| IGKV4-50*01_AJ | A/J | Musmus IGKV4-50*01 F | 100 | 286 | GAAAATGTGCTCACCCAGTCTCCAGCAATCATGTCTGCATCTCTAGGGGAGAAGGTCACCATGAGCTGCAGGGCCAGCTCAAGTGTAAATTACATGTACTGGTACCAGCAGAAGTCAGATGCCTCCCCCAAACTATGGATTTATTACACATCCAACCTGGCTCCTGGAGTCCCAGCTCGCTTCAGTGGCAGTGGGTCTGGGAACTCTTATTCTCTCACAATCAGCAGCATGGAGGGTGAAGATGCTGCCACTTATTACTGCCAGCAGTTTACTAGTTCCCCATCCA |
| IGKV4-53*01_AJ | A/J | Musmus IGKV4-53*01 F | 100 | 292 | GAAATTGTGCTCACCCAGTCTCCAGCACTCATGGCTGCATCTCCAGGGGAGAAGGTCACCATCACCTGCAGTGTCAGCTCAAGTATAAGTTCCAGCAACTTGCACTGGTACCAGCAGAAGTCAGAAACCTCCCCCAAACCCTGGATTTATGGCACATCCAACCTGGCTTCTGGAGTCCCTGTTCGCTTCAGTGGCAGTGGATCTGGGACCTCTTATTCTCTCACAATCAGCAGCATGGAGGCTGAAGATGCTGCCACTTATTACTGTCAACAGTGGAGTAGTTACCCACTCA |

| | | | | | |
|---|---|---|---|---|---|
| IGKV13-85*01_AJ | A/J | Musmus IGKV13-85*01 F | 100 | 287 | GACATCCAGATGACACAATCTTCATCCTACTTGTCTGTATCTCTAGGAGGCAGAGTCA<br>CCATTACTTGCAAGGCAAGTGACCACATTAATAATTGGTTAGCCTGGTATCAGCAGAA<br>ACCAGGAAATGCTCCTAGGCTCTTAATATCTGGTGCAACCAGTTTGGAAACTGGGGTT<br>CCTTCAAGATTCAGTGGCAGTGGATCTGGAAAGGATTACACTCTCAGCATTACCAGTC<br>TTCAGACTGAAGATGTTGCTACTTATTACTGTCAACAGTATTGGAGTACTCCTCC |
| IGKV8-34*01_AJ | A/J | Musmus IGKV8-34*01 F | 100 | 305 | GACATTTTGATGACCCAGTCTCCATCCTCCCTGACTGTGTCAGCAGGAGAGAAGGTCA<br>CTATGAGCTGCAAGTCCAGTCAGAGTCTTTTAGCTAGTGGCAACCAAAATAACTACTT<br>GGCCTGGCACCAGCAGAAACCAGGACGATCTCCTAAAATGCTGATAATTTGGGCATCC<br>ACTAGGGTATCTGGAGTCCCTGATCGCTTCATAGGCAGTGGATCTGGGACGGATTTCA<br>CTCTGACCATCAACAGTGTGCAGGCTGAAGATCTGGCTGTTTATTACTGTCAGCAGTC<br>CTACAGCGCTCCTAC |
| IGKV1-122*01_AJ | A/J | Musmus IGKV1-122*01 F | 100 | 302 | GATGCTGTGATGACCCAAACTCCACTCTCCCTGCCTGTCAGTCTTGGAGATCAAGCCT<br>CCATCTCTTGCAGGTCTAGTCAGAGCCTTGAAAACAGTAATGGAAACACCTATTTGAA<br>CTGGTACCTCCAGAAACCAGGCCAGTCTCCACAGCTCCTGATCTACAGGGTTTCCAAC<br>CGATTTTCTGGGGTCCTAGACAGGTTCAGTGGTAGTGGATCAGGGACAGATTTCACAC<br>TGAAAATCAGCAGAGTGGAGGCTGAGGATTTGGGAGTTTATTTCTGCCTCCAAGTTAC<br>ACATGTCCCTCC |
| IGKV14-111*01_AJ | A/J | Musmus IGKV14-111*01 F | 100 | 287 | GACATCAAGATGACCCAGTCTCCATCTTCCATGTATGCATCTCTAGGAGAGAGAGTCA<br>CTATCACTTGCAAGGCGAGTCAGGACATTAATAGCTATTTAAGCTGGTTCCAGCAGAA<br>ACCAGGGAAATCTCCTAAGACCCTGATCTATCGTGCAAACAGATTGGTAGATGGGGTC<br>CCATCAAGGTTCAGTGGCAGTGGATCTGGGCAAGATTATTCTCTCACCATCAGCAGCC<br>TGGAGTATGAAGATATGGGAATTTATTATTGTCTACAGTATGATGAGTTTCCTCC |
| IGKV4-79*01_AJ | A/J | Musmus IGKV4-79*01 F | 100 | 292 | CAAATTGTTCTCACCCAGTCTCCAGCAATCATGTCTGCATCTCCTGGGGAGAAGGTCA<br>CCTTGACCTGCAGTGCCAGCTCAAGTGTAAGTTCCAGCTACTTGTACTGGTACCAGCA<br>GAAGCCAGGATCCTCCCCCAAACTCTGGATTTATAGCACATCCAACCTGGCTTCTGGA<br>GTCCCTGCTCGCTTCAGTGGCAGTGGGTCTGGGACCTCTTACTCTCTCACAATCAGCAG<br>CATGGAGGCTGAAGATGCTGCCTCTTATTTCTGCCATCAGTGGAGTAGTTACCCACCC<br>C |
| IGKV17-121*01_AJ | A/J | Musmus IGKV17-121*01 F | 100 | 287 | GAAACAACTGTGACCCAGTCTCCAGCATCCCTGTCCATGGCTATAGGAGAAAAAGTCA<br>CCATCAGATGCATAACCAGCACTGATATTGATGATGATATGAACTGGTACCAGCAGAA<br>GCCAGGGGAACCTCCTAAGCTCCTTATTTCAGAAGGCAATACTCTTCGTCCTGGAGTC<br>CCATCCCGATTCTCCAGCAGTGGCTATGGTACAGATTTTGTTTTTACAATTGAAAACAT<br>GCTCTCAGAAGATGTTGCAGATTACTACTGTTTGCAAAGTGATAACTTGCCTCT |
| IGKV4-86*01_AJ | A/J | Musmus IGKV4-86*01 F | 100 | 283 | GAAATTGTGCTCACTCAGTCTCCAGCCATCACAGCTGCATCTCTGGGGCAAAAGGTCA<br>CCATCACCTGCAGTGCCAGCTCAAGTGTAAGTTACATGCACTGGTACCAGCAGAAGTC<br>AGGCACCTCCCCCAAACCATGGATTTATGAAATATCCAAACTGGCTTCTGGAGTCCCA<br>GCTCGCTTCAGTGGCAGTGGGTCTGGGACCTCTTACTCTCTCACAATCAGCAGCATGG<br>AGGCTGAAGATGCTGCCATTTATTACTGCCAGCAGTGGAATTATCCTCTTA |
| IGKV4-61*01_S2955_AJ | A/J | Musmus IGKV4-61*01 F | 100 | 286 | CAAATTGTTCTCACCCAGTCTCCAGCAATCATGTCTGCATCTCCAGGGGAGAAGGTCA<br>CCATATCCTGCAGTGCCAGCTCAAGTGTAAGTTACATGTACTGGTACCAGCAGAAGCC<br>AGGATCCTCCCCCAAACCCTGGATTTATCGCACATCCAACCTGGCTTCTGGAGTCCCT<br>GCTCGCTTCAGTGGCAGTGGGTCTGGGACCTCTTACTCTCTCACAATCAGCAGCATGG<br>AGGCTGAAGATGCTGCCACTTATTACTGCCAGCAGTATCATAGTTACCCACTCA |

| | | | | | |
|---|---|---|---|---|---|
| IGKV2-112*01_AJ | A/J | Musmus IGKV2-112*01 F | 100 | 300 | GATATTGTGATAACCCAGGATGAACTCTCCAATCCTGTCACTTCTGGAGAATCAGTTTCCATCTCCTGCAGGTCTAGTAAGAGTCTCCTATATAAGGATGGGAAGACATACTTGAATTGGTTTCTGCAGAGACCAGGACAATCTCCTCAGCTCCTGATCTATTTGATGTCCACCCGTGCATCAGGAGTCTCAGACCGGTTTAGTGGCAGTGGGTCAGGAACAGATTTCACCCTGGAAATCAGTAGAGTGAAGGCTGAGGATGTGGGTGTGTATTACTGTCAACAACTTGTAGAGTATCCT |
| IGKV4-58*01_AJ | A/J | Musmus IGKV4-58*01 F | 100 | 292 | GAAAATGTGCTCACCCAGTCTCCAGCAATAATGGCTGCCTCTCTGGGGCAGAAGGTCACCATGACCTGCAGTGCCAGCTCAAGTGTAAGTTCCAGTTACTTGCACTGGTACCAGCAGAAGTCAGGCGCTTCCCCCAAACCCTTGATTCATAGGACATCCAACCTGGCTTCTGGAGTCCCAGCTCGCTTCAGTGGCAGTGGGTCTGGGACCTCTTACTCTCTCACAATCAGCAGCGTGGAGGCTGAAGATGATGCAACTTATTACTGCCAGCAGTGGAGTGGTTACCCATTCA |
| IGKV1-133*01_AJ | A/J | Musmus IGKV1-133*01 F | 100 | 302 | GATGTTGTGATGACCCAGACTCCACTCACTTTGTCGGTTACCATTGGACAACCAGCCTCTATCTCTTGCAAGTCAAGTCAGAGCCTCTTATATAGTAATGGAAAAACCTATTTGAATTGGTTATTACAGAGGCCAGGCCAGTCTCCAAAGCGCCTAATCTATCTGGTGTCTAAACTGGACTCTGGAGTCCCTGACAGGTTCACTGGCAGTGGATCAGGAACAGATTTTACACTGAAAATCAGCAGAGTGGAGGCTGAGGATTTGGGAGTTTATTACTGCGTGCAAGGTACACATTTTCCTCA |
| IGKV3-10*01_AJ | A/J | Musmus IGKV3-10*01 F | 100 | 299 | AACATTGTGCTGACCCAATCTCCAGCTTCTTTGGCTGTGTCTCTAGGGCAGAGGGCCACCATATCCTGCAGAGCCAGTGAAAGTGTTGATAGTTATGGCAATAGTTTTATGCACTGGTACCAGCAGAAACCAGGACAGCCACCCAAACTCCTCATCTATCTTGCATCCAACCTAGAATCTGGGGTCCCTGCCAGGTTCAGTGGCAGTGGGTCTAGGACAGACTTCACCCTCACCATTGATCCTGTGGAGGCTGATGATGCTGCAACCTATTACTGTCAGCAAAATAATGAGGATCCTCC |
| IGKV4-68*01_S3092_AJ | A/J | Musmus IGKV4-68*01 F | 100 | 286 | CAAATTGTTCTCACCCAGTCTCCAGCACTCATGTCTGCATCTCCAGGGGAGAAGGTCACCATGACCTGCAGTGCCAGCTCAAGTGTAAGTTACATGTACTGGTACCAGCAGAAGCCAAGATCCTCCCCCAAACCCTGGATTTATCTCACATCCAACCTGGCTTCTGGAGTCCCTGCTCGCTTCAGTGGCAGTGGGTCTGGGACCTCTTACTCTCTCACAATCAGCAGCATGGAGGCTGAAGATGCTGCCACTTATTACTGCCAGCAGTGGAGTAGTAACCCACTCA |
| IGKV1-99*01_AJ | A/J | Musmus IGKV1-99*01 F | 100 | 302 | GATGTTGTTCTGACCCAAACTCCACTCTCTCTGCCTGTCAATATTGGAGATCAAGCCTCTATCTCTTGCAAGTCTACTAAGAGTCTTCTGAATAGTGATGGATTCACTTATTTGGACTGGTACCTGCAGAAGCCAGGCCAGTCTCCACAGCTCCTAATATATTTGGTTTCTAATCGATTTTCTGGAGTTCCAGACAGGTTCAGTGGCAGTGGGTCAGGAACAGATTTCACACTCAAGATCAGCAGAGTGGAGGCTGAGGATTTGGGAGTTTATTATTGCTTCCAGAGTAACTATCTTCCTCT |
| IGKV4-69*01_AJ | A/J | Musmus IGKV4-69*01 F | 100 | 282 | CAAATTCTTCTCACCCAGTCTCCAGCAATCATGTCTGCATCTCCAGGGGAGAAGGTCACCATGACCTGCAGTGCCAGCTCAAGTGTAAGTTACATGCACTGGTACCAGCAGAAGCCAGGATCCTCGCCCAAACCCTGGATTTATGACACATCCAACCTGGCTTCTGGATTCCCTGCTCGCTTCAGTGGCAGTGGGTCTGGGACCTCTTACTCTCTCATAATCAGCAGCATGGAGGCTGAAGATGCTGCCACTTATTACTGCCATCAGCGGAGTAGTTACCCA |

| | | | | | |
|---|---|---|---|---|---|
| IGKV8-18*01_AJ | A/J | Musmus IGKV8-18*01 ORF | 100 | 310 | GACATTGTGATGACCCAGTCTCCATCCTCCCTGGCTGTGACAGCAGGAGAGAAGGTCACTATGAGATGCAAGTCCAGTCAGAGTCTTTTGTGGAGTGTAAACCAAAATAACTACTTATCCTGGTACCAGCAGAAACAAGGGCAGCCTCCTAAACTGCTTATCTATGGGGCATCCATTAGAGAATCTTGGGTCCCTGATCGATTCACAGGAAGTGGATCTGGGACAGACTTCACTCTCACCATTAGCAATGTGCATGCTGAAGACCTAGCAGTTTATTACTGTCAGCACAATCATGGCAGCTTTCTCCCCC |
| IGKV9-124*01_AJ | A/J | Musmus IGKV9-124*01 F | 100 | 285 | GACATCCAGATGACCCAGTCTCCATCCTCCTTATCTGCCTCTCTGGGAGAAAGAGTCAGTCTCACTTGTCGGGCAAGTCAGGAAATTAGTGGTTACTTAAGCTGGCTTCAGCAGAAACCAGATGGAACTATTAAACGCCTGATCTACGCCGCATCCACTTTAGATTCTGGTGTCCCAAAAAGGTTCAGTGGCAGTAGGTCTGGGTCAGATTATTCTCTCACCATCAGCAGCCTTGAGTCTGAAGATTTTGCAGACTATTACTGTCTACAATATGCTAGTTATCCT |
| IGKV4-92*01_AJ | A/J | Musmus IGKV4-92*01 F | 100 | 291 | GAAATGGTTCTCACCCAGTCTCCAGTATCCATAACTGCATCTCGAGGGGAGAAGGTCACCATCACCTGCCGTGCCAGCTCAAGTATAAGTTCCAATTACTTACACTGGTACCAGCAGAAGCCAGGATCCTCCCCTAAACTTTTGATTTATAGGACATCCATCCTGGCATCTGGAGTCCTAGACAGCTTCAGTGGCAGTGGGTCTGAGAGCTCTTACACTCTGACAATCAGCTGCATGCAGGACGAAGTTGCTGCCACTTACTATTGTCAGCAGGGGAGTAGTAGCCCACCA |
| IGKV5-45*01_AJ | A/J | Musmus IGKV5-45*01 F | 100 | 287 | GATATTGTGCTAACTCAGTCTCCAGCCACCCTGTCTGTGACTCCAGGAGATAGAGTCAGTCTTTCCTGCAGGGCCAGTCAAAGTATTAGCAACTACCTACACTGGTATCAACAAAATCACATGAGTCTCCAAGGCTTCTCATCAAGTATGCTTCCCAGTCCATCTCTGGGATCCCCTCCAGGTTCAGTGGCAGTGGATCAGGGACAGATTTCACTCTCAGTATCAACAGTGTGGAGACTGAAGATTTTGGAATGTATTTCTGTCAACAGAGTAACAGCTGGCCTCA |
| IGKV1-88*01_AJ | A/J | Musmus IGKV1-88*01 F | 100 | 302 | GATGTTGTGGTGACTCAAACTCCACTCTCCCTGCCTGTCAGCTTTGGAGATCAAGTTTCTATCTCTTGCAGGTCTAGTCAGAGTCTTGCAAACAGTTATGGGAACACCTATTTGTCTTGGTACCTGCACAAGCCTGGCCAGTCTCCACAGCTCCTCATCTATGGGATTTCCAACAGATTTTCTGGGGTGCCAGACAGGTTCAGTGGCAGTGGTTCAGGGACAGATTTCACACTCAAGATCAGCACAATAAAGCCTGAGGACTTGGGAATGTATTACTGCTTACAAGGTACACATCAGCCTCC |
| IGKV7-33*01_S6039_AJ | A/J | Musmus IGKV7-33*01 F | 100 | 305 | GACATTGTGATGACTCAGTCTCCAACTTTCCTTGCTGTGACAGCAAGTAAGAAGGTCACCATTAGTTGCACGGCCAGTGAGAGCCTTTATTCAAGCAAACACAAGGTGCACTACTTGGCTTGGTACCAGAAGAAACCAGAGCAATCTCCTAAACTGCTGATATACGGGGCATCCAACCGATACATTGGGGTCCCTGATCGCTTCACAGGCAGTGGATCTGGGACAGATTTCACTCTGACCATCAGCAGTGTACAGGTTGAAGACCTCACACATTATTACTGTGCACAGTTTTACAGCTATCCGCT |
| IGKV8-23-1*01_AJ | A/J | Musmus IGKV8-16*01 F, or Musmus IGKV8-23-1*01 ORF | 100 | 284 | GAAATTGTGCTGACTGTGTCAGCAGGAGAGAGGGTCACTATCAGCTGCAAATCCAATCAGAATCTTTTATGGAGTGGAAACCAAAGGTACTGTTTGGTCTGGCACCAGTGGAAACCAGGGCAAACTCCTACACCGTTGATCACCTGGACATCTGATAGGTACTCTGGAGTCCCTGATCGTTTCATAGGCAGTGGATCTGTGACAGATTTCACTCTGACCATCAGCAGTGTGCAGGCTGAAGATGTGGCAGTTTATTTCTGTCAGCAGCATTTACACATTCCTCC |

| | | | | | |
|---|---|---|---|---|---|
| IGKV4-51*01_S0136_AJ | A/J | Musmus IGKV4-51*01 F | 100 | 292 | GAAAATGTGCTCACCCAGTCTCCAGCAATAATGGCTGCCTCTCTGGGGGAGAAGGTCACCATGACCTGCAGTGCCAGCTCAAGTGTAAGTTCCAGCTACTTGCACTGGTACCAGCAGAAGTCAGGCACTTCCCCCAAACTCTGGATTTATGGCACATCCAACCTGGCTTCTGGAGTCCCAGCTCGCTTCAGTGGCAGTGGGGCTGGGATCTCTTACTCTCTCACAATCAGCAGCATGGAGGCTGAAAATGATGCAACTTATTACTGCCAGCAGTGGAGTGGTTACCCGTACA |
| IGKV4-62*01_S5834_AJ | A/J | Musmus IGKV4-62*01 ORF | 100 | 286 | CAAATTGTTCTCACCCAGTCTCCAGCAATCATGTCTGCATCTCCAGGGGAAAAGGTCACCATGACCTGCAGTGCCAGCTCAAGTGTAAGTTACATGTACTGGTACCAGCAGAAGCCAGGCTCCTCCCCCAGACTCTGGATTTATGACACATCCAACCTGGTTTCTGGAGTCCCTGCTCGCTTCAGTGGCAGTAGGTCTGGGACCTCTTATTCTCTCACAATCAGCAGCATGGAGGCTGAAGATGCTGCCACTTATTACTGCCAGCAGTACAGTGGTTACCCATACA |
| IGKV14-130*01_AJ | A/J | Musmus IGKV14-130*01 F | 100 | 287 | GAAATCCAGATGACCCAGTCTCCATCCTCTATGTCTGCATCTCTGGGAGACAGAATAACCATCACTTGCCAGGCAACTCAAGACATTGTTAAGAATTTAAACTGGTATCAGCAGAAACCAGGGAAACCCCCTTCATTCCTGATCTATTATGCAACTGAACTGGCAGAAGGGGTCCCATCAAGGTTCAGTGGCAGTGGGTCTGGGTCAGACTATTCTCTGACAATCAGCAACCTGGAGTCTGAAGATTTTGCAGACTATTACTGTCTACAGTTTTATGAGTTTCCTCC |
| IGKV9-123*01_AJ | A/J | Musmus IGKV9-123*01 F | 100 | 285 | GACATCCAGATGATTCAGTCTCCATCGTCCATGTTTGCCTCTCTGGGAGACAGAGTCAGTCTCTCTTGTCGGGCTAGTCAGGGCATTAGAGGTAATTTAGACTGGTATCAGCAGAAACCAGGTGGAACTATTAAACTCCTGATCTACTCCACATCCAATTTAAATTCTGGTGTCCCATCAAGGTTCAGTGGCAGTGGGTCTGGGTCAGATTATTCTCTCACCATCAGCAGCCTAGAGTCTGAAGATTTTGCAGACTATTACTGTCTACAGCGTAATGCGTATCCT |
| IGKV10-95*01_AJ | A/J | Musmus IGKV10-95*01 F | 100 | 287 | GATATCCAGATGACACAGACTACTTCCTCCCTGTCTGCCTCTCTGGGAGACAGAGTCACCATCAGTTGCAGGGCAAGTGAGGACATTAGCACTTATTTAAACTGGTATCAGCAGAAACCAGATGGAACTGTTAAACTCCTGATCTATTACACATCAGGATTACACTCAGGAGTCCCATCAAGGTTCAGTGGCAGTGGGTCTGGGGCAGATTATTCTCTCACCATCAGCAACCTGGAACCTGAAGATATTGCCACTTACTATTGTCAGCAGTATAGTAAGCTTCCTCC |
| IGKV11-125*01_AJ | A/J | Musmus IGKV11-125*01 F | 100 | 287 | GATGTCCAGATGATTCAGTCTCCATCCTCCCTGTCTGCATCTTTGGGAGACATAGTCACCATGACTTGCCAGGCAAGTCAGGGCACTAGCATTAATTTAAACTGGTTTCAGCAAAAACCAGGGAAAGCTCCTAAGCTCCTGATCTATGGTGCAAGCAACTTGGAAGATGGGGTCCCATCAAGGTTCAGTGGCAGTAGATATGGGACAGATTTCACTCTCACCATCAGCAGCCTGGAGGATGAAGATATGGCAACTTATTTCTGTCTACAGCATAGTTATCTCCCTCC |
| IGKV3-1*01_AJ | A/J | Musmus IGKV3-1*01 F | 100 | 299 | GACATTGTGCTCACCCAATCTCCAGCTTCTTTGGCTGTGTCTCTAGGGCAGAGAGCCACCATCTCCTGCAGAGCCAGTGAAAGTGTTGAATATTATGGCACAAGTTTAATGCAGTGGTACCAACAGAAACCAGGACAGCCACCCAAACTCCTCATCTATGCTGCATCCAACGTAGAATCTGGGGTCCCTGCCAGGTTTAGTGGCAGTGGGTCTGGGACAGACTTCAGCCTCAACATCCATCCTGTGGAGGAGGATGATATTGCAATGTATTTCTGTCAGCAAAGTAGGAAGGTTCCTTC |
| IGKV1-132*01_AJ | A/J | Musmus IGKV1-132*01 F | 100 | 302 | GATGTTGTGATGACCCAGACTCCACTGTCTTTGTCGGTTACCATTGGACAACCAGCCTCTATCTCTTGCAAGTCAAGTCAGAGCCTCTTATATAGTAATGGAAAGACATATTTGAATTGGTTACAACAGAGGCCTGGCCAGGCTCCAAAGCACCTAATGTATCAGGTGTCCAAACTGGACCCTGGCATCCCTGACAGGTTCAGTGGCAGTGGATCAGAAACAGATTTTACACTTAAAATCAGCAGAGTGGAGGCTGAAGATTTGGGAGTTTATTACTGCTTGCAAGGTACATATTATCCTCA |

| | | | | | |
|---|---|---|---|---|---|
| IGKV2-109*03_S1521_AJ | A/J | Musmus IGKV2-116*01 F | 100 | 301 | GATATTGTGATGACGCAGGCTGCCTTCTCCAATCCAGTCACTCTTGGAACATCAGCTTCCATCTCCTGCAGGTCTAGTAAGAATCTCCTACATAGTAATGGCATCACTTATTTGTATTGGTATCTGCAGAGGCCAGGCCAGTCTCCTCAGCTCCTGATATATCGGGTGTCCAATCTGGCCTCAGGAGTCCCAAACAGGTTCAGTGGCAGTGAGTCAGGAACTGATTTCACACTGAGAATCAGCAGAGTGGAGGCTGAGGATGTGGGTGTTTATTACTGTGCTCAACTGCTAGAACTCCCTC |
| IGKV9-129*01_AJ | A/J | Musmus IGKV9-129*01 ORF | 100 | 287 | GACATCCAGATGACCCAGTCTCCATCCTCCTTATCTGCCTCTCTGGGAGAAAGAGTCAGTCTCACTTGCCGGGCAAGTCAGGACATTCATGGTTATTTAAACTTGTTTCAGCAGAAACCAGGTGAAACTATTAAACACCTGATCTATGAAACATCCAATTTAGATTCTGGTGTCCCAAAAAGGTTCAGTGGCAGTAGGTCTGGGTCAGATTATTCTCTCATTATCGGCAGCCTTGAGTCTGAAGATTTTGCAGACTATTACTGTCTACAATATGCTAGTTCTCCTCC |
| IGKV12-38*01_AJ | A/J | Musmus IGKV12-38*01 F | 100 | 287 | GACATCCAGATGACTCAGTCTCCAGCCTCCCTGGCTGCATCTGTGGGAGAAACTGTCACCATCACATGTCGAGCAAGTGAGAACATTTACTACAGTTTAGCATGGTATCAGCAGAAGCAAGGGAAATCTCCTCAGCTCCTGATCTATAATGCAAACAGCTTGGAAGATGGTGTCCCATCGAGGTTCAGTGGCAGTGGATCTGGGACACAGTATTCTATGAAGATCAACAGCATGCAGCCTGAAGATACCGCAACTTATTTCTGTAAACAGGCTTATGACGTTCCTCC |
| IGKV4-52*01_AJ | A/J | Musmus IGKV4-52*01 ORF, or Musmus IGKV4-54*01 ORF | 100 | 286 | CAAATTGTTCTCACCCAGTCTCCAGCAATCCTGTCTGCATCTCCAGGGGAGAAGGTCACCATGACCTGCAGTGCCAGCTCAAGTGTAAGTTACATGTACAGGTACCAGCAGAAGCCAGGATCCTCACCCAAACCCTGGATTTATGGCACATCCAACCTGGCTTCTGGAGTCCCTGCTCGCTTCAGTGGCAGTGGATCTGGGACCTCTTATTCTCTCACAATCAGCAGCATGGAGGCTGAAGATGCTGCCACTTATTACTGCCAGCAATATCATAGTTACCCACCCA |
| IGKV4-56*01_AJ | A/J | Musmus IGKV4-56*01 P | 100 | 286 | CAAATTGTTCTCACCCAGTCTCCAGCAATCATGTCTGCATCTCCAGGGCAGAAAGTCACCATAACCTGCAGTGCCATCTCAAGTGTAAATTACATGCACTGGTACCAGCAGAAGCCAGGATCCTCCCCCAAACTCTGGATTTATGCAACATCCAAACTGGCTCTTGGAGTCCCTGCTTGCTTCAGTGGCAGTGGGTCTGGGACCTCTTACTCTCTCACAATCAGCAGCATGGTGGCTGAAGATGCCACCTCTTATTTCTGTCATCAGTGGAGTAGTTACCCACCCA |
| IGKV4-81*01_AJ | A/J | Musmus IGKV4-81*01 F | 100 | 292 | GAAAATGTGCTGACCCAGTCTCCAGCAATCATGGCTGCATCTCCAGGGGAGAAGGTCACCATGACCTGCAGTGCCAGCTCAAGTGTAAGTTCTAGTAACTTGCACTGGTACCAGCAGAAGTCAGGCACTTCTACCAAATTCTGGATTTATAGGACATCCAACCTGGCTTCAGAAGTCCCAGCTCCCTTCAGTGGCAGTGGGTCTGGGACCTCTTACTCTCTTACAATCAGCAGCGTGGAGGCCGAAGATGCTGCCACTTATTACTGCCAGCAGTGGAGTGGTTACCCACCCA |
| IGKV4-60*01_S1112_AJ | A/J | Musmus IGKV4-60*01 P | 100 | 292 | GAAATTGTGCTCACCCAGTCTCCAGCACTCATGGCTGCATCTCCAGGGGAGAAGGTCACCATCACCTGCAGTGTCAGCTCAAGTATAAGTTCCAGCTACTTGCACTGGTACCAGCAGAAGTCAGGAATCTCCCCCAAACCCTGGATTTATGGCACATCCAACCTGGCTTCTGGAGTCCCTGCTCGCTTCAGTGGCAGTGGATCTGGGACCTCTTACTCTCTCACAATCAGCAGCATGGAGGCTGAAGATGCTGCCACTTATTACTGTCAACAGTGGAGTAGTTCCCCACTCA |
| IGKV4-78*01_AJ | A/J | Musmus IGKV4-78*01 F | 100 | 292 | CAAATTGTTCTCACCCAGTCTCCAGCAATCATGTCTGCATCTCCTGGGGAGAAGGTCACCATGACCTGCAGTGCCAGATCAAGTGTAAGTTCCAGCTACTTGTACTGGTACCAGCAGAAGCCAGGATCCTCCCCCAAACTCTGGATTTATAGCACATCCAACCTGGCTTCTGGAGTCCCTGCTCGCTTCAGTGGCAGTGGGTCTGGGACCTCTTATTCTCTCACAATCAGCAGCATGGAGGCTGAAGATGCTGCCACTTTTTACTGCCAGCAGTACAGTGGTTACCCATCCA |

| | | | | | |
|---|---|---|---|---|---|
| IGKV4-90*01_AJ | A/J | Musmus IGKV4-90*01 F | 100 | 285 | GAAATTTTGCTCACCCAGTCTCCAGCAATCATAGCTGCATCTCCTGGGGAGAAGGTCACCATCACCTGCAGTGCCAGCTCAAGTGTAAGTTACATGAACTGGTACCAGCAGAAACCAGGATCCTCCCCCAAAATATGGATTTATGGTATATCCAACCTGGCTTCTGGAGTTCCTGCTCGCTTCAGTGGCAGTGGGTCTGGGACATCTTTCTCTTTCACAATCAACAGCATGGAGGCTGAAGATGTTGCCACTTATTACTGTCAGCAAAGGAGTAGTTACCCACCC |
| IGKV5-37*01_AJ | A/J | Musmus IGKV5-37*01 F | 100 | 287 | GACATCCTGCTGACCCAGTCTCCAGCCACCCTGTCTGTGACTCCAGGAGAAACAGTCAGTCTTTCCTGTAGGGCCAGCCAGAGTATTTACAAGAACCTACACTGGTATCAACAGAAATCACATCGGTCTCCAAGGCTTCTCATCAAGTATGCTTCTGATTCCATCTCTGGGATCCCCTCCAGGTTCACTGGCAGTGGATCAGGGACAGATTACACTCTCAGTATCAACAGTGTGAAGCCCGAAGATGAAGGAATATATTACTGTCTTCAAGGTTACAGCACACCTTC |
| IGKV6-14*01_AJ | A/J | Musmus IGKV6-14*01 F | 100 | 287 | GACATTGTGATGACCCAGTCTCAAAAATTCATGTCCACATCAGTAGGAGACAGGGTCAGCATCACCTGCAAGGCCAGTCAGAATGTTCGTACTGCTGTAGCCTGGTATCAACAGAAACCAGGGCAGTCTCCTAAAGCACTGATTTACTTGGCATCCAACCGGCACACTGGAGTCCCTGATCGCTTCACAGGCAGTGGATCTGGGACAGATTTCACTCTCACCATTAGCAATGTGCAATCTGAAGACCTGGCAGATTATTTCTGTCTGCAACATTGGAATTATCCTCT |
| IGKV8-26*01_AJ | A/J | Musmus IGKV8-26*01 ORF | 100 | 311 | GACATTGTGATGACCCAGTCTCCATCCTCCCTGGCTGTGACAGCAGGAGAGAAGGTCACTATGAGCTGCAAGTCCAGTCAGAGTCTTTTGTGGAGTGTAAACCAAAAGAACTACTTGTCCTGGTACCAGCAGAAACAAAGGCAGCCTCCTAAACTGCTTATCTATGGGGCATCCATTAGAGAATCTTGGGTCCCTGATCGGTTCACAGGAAGTGGATCTGGGACAGACTTCACTCTCACCATTAGCAATGTGCATGCTGAAGACCTAGCAGTTTATTACTGTCAACACAATCATGGCAGCTTTCTCCCCCC |
| IGKV18-36*01_AJ | A/J | Musmus IGKV18-36*01 F | 100 | 287 | ACTGGAGAAACAACACAGGCTCCAGCTTCTCTGAGTTTTTCTCTTGGTGAAACAGCAACACTGTCATGCAGGTCCAGTGAGAGTGTTGGCAGCTACTTAGCCTGGTACCAGCAGAAAGCAGAGCAAGTTCCCCGGCTCCTTATCCATAGTGCCTCCACTAGGGCCGGTGGTGTCCCAGTCCGGTTCAGTGGCACTGGGTCTGGGACAGACTTCACTCTCACCATCAGCAGTCTAGAACCTGAAGATGCTGCAGTTTACTACTGTCAACCTTTCAAAAGTTGGTCATA |
| IGKV3-3*01_AJ | A/J | Musmus IGKV3-3*01 F | 100 | 299 | GACATTGTGCTGACCCAATCTCCAGCTTCTTTGGCTGTGTCTCTAGGACAGAGAGCCACTATCTTCTGCAGAGCCAGCCAGAGTGTCGATTATAATGGAATTAGTTATATGCACTGGTTCCAACAGAAACCAGGACAGCCACCCAAACTCCTCATCTATGCTGCATCCAACCTAGAATCTGGGATCCCTGCCAGGTTCAGTGGCAGTGGGTCTGGGACAGACTTCACCCTCAACATCCATCCTGTGGAGGAGGAAGATGCTGCAACCTATTACTGTCAGCAAAGTATTGAGGATCCTCC |
| IGKV4-52*01_S5019_AJ | A/J | Musmus IGKV4-52*01 ORF, or Musmus IGKV4-54*01 ORF | 100 | 286 | CAAATTGTTCTCACCCAGTCTCCAGCAATCCTGTCTGCATCTCCAGGGGAGAAGGTCACCATGACCTGCAGTGCCAGCTCAAGTGTAAGTTACATGTACAGGTACCAGCAGAAGCCAGGATCCTCACCCAAACCCTGGATTTATGGCACATCCAACCTGGCTTCTGGAGTCCCTGCTCGCTTCAGTGGCAGTGGATCTGGGACCTCTTATTCTCTCACAATCAGCAGCATGGAGGCTGAAGATGCTGCCACTTATTACTGCCAGCAATATCATAGTTACCCACTCA |
| IGKV4-81*01_S0130_AJ | A/J | Musmus IGKV4-81*01 F | 100 | 292 | GAAAATGTGCTGACCCAGTCTCCAGCAATCATGGCTGCATCTCCAGGGGAGAAGGTCACCATGACCTGCAGTGCCAGCTCAAGTGTAAGTTCTAGTAACTTGCACTGGTACCAGCAGAAGTCAGGCACTTCTACCAAATTCTGGATTTATAGGACATCCAACCTGGCTTCAGAAGTCCCAGCTCCCTTCAGTGGCAGTGGGTCTGGGACCTCTTACTCTCTTACAATCAGCAGCGTGGAGGCCGAAGATGCTGCCACTTATTACTGCCAGCAGTGGAGTGGTTACCCACTCA |

| | | | | | |
|---|---|---|---|---|---|
| IGKV6-15*01_S9105_AKR | AKR/J | Musmus IGKV6-15*01 F | 97.13 | 287 | GACATTGTGATGACCCAGTCTCAAAAATTCATGTCCACATCAGTAGGAGACAGGGTCAGCATCAGCTGCAAGGCCAGTCAGAATGTGGGTAATATTATAGCCTGGTATCAACAGAAACCAGGGCAATCTCCTAAAGCACTGATTTACTTGGCATCCTACCGGTACAGTGGAGTCCCTGATCGCTTCACAGGCAGTGGATCTGGGACAGATTTCACTCTCACCATTAGCAATGTGCAGTCTGAAGACTTGGCAGAGTATTTCTGTCAGCAATATAGCAGCTCTCCTCT |
| IGKV6-20*01_S2082_AKR | AKR/J | Musmus IGKV6-20*01 F | 99.64 | 287 | AACATTGTAATGACCCAATCTCCCAAATCCATGTCCATGTCAGTAGGAGAGAGGGTCACCTTGAGCTGCAAGGCCAGTGAGAATGTGGGTACTTATGTATCCTGGTATCAACAGAAACCAGAGCAGTCTCCTAAACTGCTGATATACGGGGCATCCAACCGGTACACTGGGGTCCCCGATCGCTTCACAGGCAGTGGATCTGCAACAGATTTCACTCTGACCATCAGCAGTGTGCAGGCTGAAGACCTTGCAGATTATTACTGTGGACAGAGTTACAGCTATCCTCC |
| IGKV8-19*01_AKR | AKR/J | Musmus IGKV8-19*01 F | 100 | 305 | GACATTGTGATGACACAGTCTCCATCCTCCCTGACTGTGACAGCAGGAGAGAAGGTCACTATGAGCTGCAAGTCCAGTCAGAGTCTGTTAAACAGTGGAAATCAAAAGAACTACTTGACCTGGTACCAGCAGAAACCAGGGCAGCCTCCTAAACTGTTGATCTACTGGGCATCCACTAGGGAATCTGGGGTCCCTGATCGCTTCACAGGCAGTGGATCTGGAACAGATTTCACTCTCACCATCAGCAGTGTGCAGGCTGAAGACCTGGCAGTTTATTACTGTCAGAATGATTATAGTTATCCTCC |
| IGKV1-135*01_AKR | AKR/J | Musmus IGKV1-135*01 F | 100 | 302 | GATGTTGTGATGACCCAGACTCCACTCACTTTGTCGGTTACCATTGGACAACCAGCCTCCATCTCTTGCAAGTCAAGTCAGAGCCTCTTAGATAGTGATGGAAAGACATATTTGAATTGGTTGTTACAGAGGCCAGGCCAGTCTCCAAAGCGCCTAATCTATCTGGTGTCTAAACTGGACTCTGGAGTCCCTGACAGGTTCACTGGCAGTGGATCAGGGACAGATTTCACACTGAAAATCAGCAGAGTGGAGGCTGAGGATTTGGGAGTTTATTATTGCTGGCAAGGTACACATTTTCCTCA |
| IGKV2-137*01_AKR | AKR/J | Musmus IGKV2-137*01 F | 100 | 302 | GATATTGTGATGACTCAGGCTGCACCCTCTGTACCTGTCACTCCTGGAGAGTCAGTATCCATCTCCTGCAGGTCTAGTAAGAGTCTCCTGCATAGTAATGGCAACACTTACTTGTATTGGTTCCTGCAGAGGCCAGGCCAGTCTCCTCAGCTCCTGATATATCGGATGTCCAACCTTGCCTCAGGAGTCCCAGACAGGTTCAGTGGCAGTGGGTCAGGAACTGCTTTCACACTGAGAATCAGTAGAGTGGAGGCTGAGGATGTGGGTGTTTATTACTGTATGCAACATCTAGAATATCCTTT |
| IGKV12-46*01_S1254_AKR | AKR/J | Musmus IGKV12-46*01 F | 94.62 | 287 | GACATCCAGATGACTCAGTCTCCAGCTTCCCTGTCTGCATCTGTGGGAGAAACTGTCACCATCACATGTCGAGCAAGTGAGAATATTGACAGTTATTTAGCATGGTATCAGCAGAAACAGGGAAAATCTCCTCAGCTCCTGGTCTATGCTGCAACACTCTTAGCAGATGGTGTGCCATCAAGGTTCAGTGGCAGTGGATCAGGCACACAGTATTCTCTCAAGATCAACAGCCTGCAGTCTGAAGATGTTGCGAGATATTACTGTCAACATTATTATAGTACTCCTCC |
| IGKV8-21*01_S1419_AKR | AKR/J | Musmus IGKV8-21*01 F | 99.33 | 302 | GACATTGTGATGTCACAGTCTCCATCCTCCCTGGCTGTGTCAGCAGGAGAGAAGGTCACTATGAGCTGCAAATCCAGTCAGAGTCTGTTCAACAGTAGAACCCGAAAGAACTACTTGGCTTGGTACCAGCAGAAACCAGGGCAGTCTCCTAAACTGCTGATCTACTGGGCATCCACTAGGGAATCTGGGGTCCCTGATCGCTTCACAGGCAGTGGATCTGGGACAGATTTCACTCTCACCATCAGCAGTGTGCAGGCTGAAGACCTGGCAGTTTATTACTGCAAGCAATCTTATTATCTTCC |
| IGKV12-46*01_S4541_AKR | AKR/J | Musmus IGKV12-46*01 F | 97.13 | 287 | GACATCCAGATGACTCAGTCTCCAGCCTCCCTATCTGCATCTGTGGGAGAAACTGTCACCATCACATGTCGAGCAAGTGAGAATATTTACAGTAATTTAGCATGGTATCAGCAGAAACAGGGAAAATCTCCTCAGCTCCTGGTCTATGCTGCAACAAATTTAGCAGATGGTGTGCCATCAAGGTTCAGTGGCAGTGGATCAGGCACACAGTTTTCTCTGAAGATCAACAGCCTGCAGCCTGAAGATTTTGGGAGTTATTACTGTCAACATTTTATGGTACTCCTCC |

| | | | | | |
|---|---|---|---|---|---|
| IGKV6-b*01_AKR | AKR/J | Musmus IGKV6-b*01 F | 100 | 287 | AGTATTGTGATGACCCAGACTCCCAAATTCCTGCCTGTATCAGCAGGAGACAGGGTTACCATGACCTGCAAGGCCAGTCAGAGTGTGGGTAATAATGTAGCCTGGTACCAACAGAAGCCAGGACAGTCTCCTAAACTGCTGATATACTATGCATCCAATCGCTACACTGGAGTCCCTGATCGCTTCACTGGCAGTGGATCTGGGACAGATTTCACTTTCACCATCAGCAGTGTGCAGGTTGAAGACCTGGCAGTTTATTTCTGTCAGCAGCATTATAGCTCTCCTCC |
| IGKV6-15*01_S9094_AKR | AKR/J | Musmus IGKV6-15*01 F | 96.06 | 287 | GACATTGTGATGACCCAGTCTCAAAAATTCATGTCCACATCAGTAGGAGACAGGGTCAGCGTCACCTGCAAGGCCAGTCAGTATGTGGGTACTTATGTAGCCTGGTATCAACAGAAACCAGGACAATCTCCTAAAGCACTGATTTACTCGGCATCCACCCGGCACACTGGAGTCCCTGATCGCTTCACAGGCAGTGGATCTGGGACAGATTTCACTCTCACCATTAGCAATGTGCAGTCTGAAGACTTGGCAGAGTATTTCTGTGAGCAATACAGCAGCTCTCCTCT |
| IGKV1-110*01_AKR | AKR/J | Musmus IGKV1-110*01 F | 100 | 302 | GATGTTGTGATGACCCAAACTCCACTCTCCCTGCCTGTCAGTCTTGGAGATCAAGCCTCCATCTCTTGCAGATCTAGTCAGAGCCTTGTACACAGTAATGGAAACACCTATTTACATTGGTACCTGCAGAAGCCAGGCCAGTCTCCAAAGCTCCTGATCTACAAAGTTTCCAACCGATTTTCTGGGGTCCCAGACAGGTTCAGTGGCAGTGGATCAGGGACAGATTTCACACTCAAGATCAGCAGAGTGGAGGCTGAGGATCTGGGAGTTTATTTCTGCTCTCAAAGTACACATGTTCCTCC |
| IGKV12-44*01_AKR | AKR/J | Musmus IGKV12-44*01 F | 100 | 287 | GACATCCAGATGACTCAGTCTCCAGCCTCCCTATCTGCATCTGTGGGAGAAACTGTCACCATCACATGTCGAGCAAGTGAGAATATTTACAGTTATTTAGCATGGTATCAGCAGAAACAGGGAAAATCTCCTCAGCTCCTGGTCTATAATGCAAAAACCTTAGCAGAAGGTGTGCCATCAAGGTTCAGTGGCAGTGGATCAGGCACACAGTTTTCTCTGAAGATCAACAGCCTGCAGCCTGAAGATTTTGGGAGTTATTACTGTCAACATCATTATGGTACTCCTCC |
| IGKV1-117*01_AKR | AKR/J | Musmus IGKV1-117*01 F | 100 | 302 | GATGTTTTGATGACCCAAACTCCACTCTCCCTGCCTGTCAGTCTTGGAGATCAAGCCTCCATCTCTTGCAGATCTAGTCAGAGCATTGTACATAGTAATGGAAACACCTATTTAGAATGGTACCTGCAGAAACCAGGCCAGTCTCCAAAGCTCCTGATCTACAAAGTTTCCAACCGATTTTCTGGGGTCCCAGACAGGTTCAGTGGCAGTGGATCAGGGACAGATTTCACACTCAAGATCAGCAGAGTGGAGGCTGAGGATCTGGGAGTTTATTACTGCTTTCAAGGTTCACATGTTCCTCC |
| IGKV3-1*01_S7345_AKR | AKR/J | Musmus IGKV3-1*01 F | 98.63 | 299 | GACATTGTGCTCACCCAATCTCCAGCTTCTTTGGCTGTGTCTCTAGGGCAGAGTGTCACCATCTCCTGCAGAGCCAGTGAAAGTGTTGAATATTATGGCACTAGTTTAATGCAGTGGTACCAACAGAAACCAGGACAGCCACCCAAACTCCTCATCTATGGTGCATCCAACGTAGAATCTGGGGTCCCTGCCAGGTTTAGTGGCAGTGGGTCTGGGACAGACTTCAGCCTCAACATCCATCCTGTGGAGGAGGATGATATTGCAATGTATTTCTGTCAGCAAAGTAGGAAGGTTCCTTC |
| IGKV6-23*01_S4960_AKR | AKR/J | Musmus IGKV6-23*01 F | 92.11 | 287 | GACATTGTGATGACCCAGTCTCAAAAATTCATGTCCACATCAGTAGGAGAGAGGGTCAGCATCACCTGCAAGGCCAGTCAGAATGTAGGTACTAATGTAGCCTGGTATCAGCAGAAAGCAGGGCAGTCTCTTGAACTGCTGATCTATGGGGCATCCAACCGGCACACTGGAGTCCCTGATCGCTTCACAGGCAGTGGATCTGGGACAGATTTCACTCTCACCATCACCAATGTGCAGTCTGAAGACATGACAAATTATTTCTGTGAGCAATATAGCAGCTATCCTCT |
| IGKV5-45*01_S0430_AKR | AKR/J | Musmus IGKV5-45*01 F | 99.64 | 287 | GATATTGTGCTAACTCAGTCTCCAGCCACCCTGTCTGTGACTCCAGGAGATAGAGTCAGTCTTTCCTGCAGGGCCAGCCAAAGTATTAGCAACTACCTACACTGGTATCAACAAAAATCACATGAGTCTCCAAGGCTTCTCATCAAGTATGCTTCCCAGTCCATCTCTGGGATCCCCTCCAGGTTCAGTGGCAGTGGATCAGGGACAGATTTCACTCTCAGTATCAACAGTGTGGAGACTGAAGATTTTGGAATGTATTTCTGTCAACAGAGTAACAGCTGGCCTCA |

| | | | | | |
|---|---|---|---|---|---|
| IGKV10-94*03_AKR | AKR/J | Musmus IGKV10-94*03 F, or Musmus IGKV10-94*08 F | 100 | 287 | GATATCCAGATGACACAGACTACATCCTCCCTGTCTGCCTCTCTGGGAGACAGAGTCACCATCAGTTGCAGGGCAAGTCAGGATATTAGCAATTATTTAAACTGGTATCAGCAGAAACCAGATGGAACTGTTAAACTCCTGATCTACTACACATCAAGATTACACTCAGGAGTCCCATCAAGGTTCAGTGGCAGTGGGTCTGGGACAGATTATTCTCTCACCATCAGCAACCTGGAACCTGAAGATATTGCCACTTACTATTGTCAGCAGTATAGTAAGCTTCCTCC |
| IGKV4-74*01_S3301_AKR | AKR/J | Musmus IGKV4-74*01 F | 94.68 | 292 | CAGATTGTTCTCACCCAGTCTCCAGCAATCATGTCTGCATCTCCAGGGGAGAAGGTCACCATGACCTGCAGGGCCAGCTCAAGTGTAAGTTCCAGTTACTTGCACTGGTACCAGCAGAAGCCAGGATCTTCCCCCAAACTCTGGATTTATAGCACATCCAACCTGGCTTCAGGAGTCCCAGCTCGCTTCAGTGGCAGTGGGTCTGGGACCTCTTACTCTCTCACAATCAGCAGTGTGGAGGCTGAGGATGCTGCCACTTATTACTGCCAGCAGTATGATAGTTCCCCACCCA |
| IGKV4-56*01_S5696_AKR | AKR/J | Musmus IGKV4-56*01 P | 96.38 | 286 | CAAATTGTTCTCACCCAGTCTCCAGCAATCATGTCTGCATCTCCAGGGCAGAAAGTCACCATAACCTGCAGTGCCAGCTCAAGTGTAAATTACATGCACTGGTACCAGCAGAAGCTAGGATCCTCCCCCAAACTCTGGATTTATGACACATCCAAACTGGCTCCTGGAGTCCCTGCTCGCTTCAGTGGCAGTGGGTCTGGGACCTCTTACTCTCTCACAATCAGCAGCATGGAGGCTGAAGATGCTGCCTCTTATTTCTGCCATCAGTGGAGTAGTTACCCACCCA |
| IGKV12-46*01_S4026_AKR | AKR/J | Musmus IGKV12-46*01 F | 99.64 | 287 | GACATCCAGATGACTCAGTCTCCAGCCTCCCTATCTGTATCTGTGGGAGAAACTGTCACCATCACATGTCGAGCAAGTGAGAATATTTACAGTAATTTAGCATGGTATCAGCAGAAACAGGGAAAATCTCCTCAGCTCCTGGTCTATGCTGCAACAAACTTAGCAGATGGTGTTCCATCAAGGTTCAGTGGCAGTGGATCAGGCACACAGTATTCCCTCAAGATCAACAGCCTGCAGTCTGAAGATTTTGGGAGTTATTACTGTCAACATTTTTGGGGTACTCCTCC |
| IGKV3-4*01_S8857_AKR | AKR/J | Musmus IGKV3-4*01 F | 97.25 | 299 | GACATTGTGCTGACCCAATCTCCAGCTTCTTTGGCTGTGTCTCTAGGGCAGAGGGCCACCATCTCCTGCAGAGCCAGCGAAAGTGTTGATTATGATGGCGATAGTTATATGAACTGGTACCAACAGAAACCAGGACAGCCACCGAAACTCCTCATCTATGCTGCATCCAATCTAGAATCTGGGATCCCTGCCAGGTTTAGTGGCAGTGGGTCTGGGACAGACTTCACCCTCAACATTCATCCTGTGGAGGAGGAGGATGCTGCAACCTATTACTGTCTGCAAAGTAATGAGGCTCCTCC |
| IGKV5-43*01_S9511_AKR | AKR/J | Musmus IGKV5-43*01 F | 99.64 | 287 | GATATTGTGCTAACTCAGTCTCCAGCCACCCTGTCTGTGACTCCAGGAGATAGCGTCAGTCTTTCCTGCAGGGCCAGCCAAAGTATTAGCAACAACCTACACTGGTATCAACAAAAATCACATGAGTCTCCAAGGCTTCTCATCAAGTATGCTTCCCAGTCCATCTCTGGGATCCCCTCCAAGTTCAGTGGCAGTGGATCAGGGACAGATTTCACTCTCAGTATCAACAGTGTGGAGACTGAAGATTTTGGAATGTATTTCTGTCAACAGAGTAACAGCTGGCCTCA |
| IGKV14-126*01_AKR | AKR/J | Musmus IGKV14-126*01 F | 100 | 287 | GACATCAAGATGACCCAGTCTCCATCCTCCATGTATGCATCGCTGGGAGAGAGAGTCACTATCACTTGCAAGGCGAGTCAGGACATTAAAAGCTATTTAAGCTGGTACCAGCAGAAACCATGGAAATCTCCTAAGACCCTGATCTATTATGCAACAAGCTTGGCAGATGGGGTCCCATCAAGATTCAGTGGCAGTGGATCTGGGCAAGATTATTCTCTAACCATCAGCAGCCTGGAGTCTGACGATACAGCAACTTATTACTGTCTACAGCATGGTGAGAGCCCTCC |
| IGKV4-59*01_S1025_AKR | AKR/J | Musmus IGKV4-59*01 F | 98.19 | 286 | CAAATTGTTCTCACCCAGTCTCCAGCAATCATGTCTGCATCTCCAGGGGAGAAGGTCACCATGACATGCAGTGCCAGCTCAAGTGTAAGTTACATGCACTGGTACCAGCAGAAGTCAGGCACCTCCCCCAAAAGATGGATTTATGACACATCCAAACTGGATTCTGGAGTCCCTGCTCGCTTCAGTGGCAGTGGGTCTGGGACCTCTTATTCTCTCACAATCAGCAGCATGGAGGCTGAAGATGCTGCCACTTATTACTGCTATCAGTGGAGTAGTTACCCACCCA |

| | | | | | |
|---|---|---|---|---|---|
| IGKV7-33*01_S2856_AKR | AKR/J | Musmus IGKV7-33*01 F | 97.31 | 305 | GACATTGTGATGACTCAGTCTCCAACTTTCCTTGCTGTGACAGCAAGTAAGAAGGTCACCATTAGTTGCACGGCCAGTGAGAGCCTTTATTCAAGCAAACACAAGGTGAACTACTTAGCTTGGTACCAGAAGAAACCAGAGCAATCTCCTAAACTGCTGATTTATGGGGCATCCAACCGATTCACTGGGGTCCCTGATCGCTTCACAGGCAGTGGATCTGGGACAGATTTCACTCTGACCATCAGCAGTGTGCAGGCTGAAGACCTCACACATTATTACTGTGCACAGTTTTACAGCTATCCTCT |
| IGKV8-24*01_S8843_AKR | AKR/J | Musmus IGKV8-24*01 F | 98.32 | 305 | GACATTGTGATGACACAGTCTCCATCCTCCCTGGCTATGTCAGTAGGACAGAAGGTCACTATGAGCTGCAAGTCCAGTCAGAGCCTTTTAAGTAGTAGCAATCAAAAGAACTATTTGGCCTGGTACCAGCAGAAACCAGGACAGTCTCCTAAACTTCTGGTATACTTTGCATCCACTAGGGAATCTGGGGTCCCTGATCGCTTCATAGGCAGTGGATCTGGGACAGATTTCACTCTTACCATCAGCAGTGTGCAGGCTGAAGACCTGGCACTTTATTACTGTCAGCAACATTATAGCACTCCTCC |
| IGKV4-53*01_S3575_AKR | AKR/J | Musmus IGKV4-53*01 F | 98.94 | 292 | GAAATTGTGCTCACCCAGTCTCCAGCACTCATGGCTGCATCTCCAGGGGAGAAGGTCACCATCACCTGCAGTGTCAGCTCAAGTATAAGTTCCAGCAACTTACACTGGTACCAGCAGAAGTCAGGAACCTCCCCCAAACCCTGGATTTATGGCACATCCAACCTTGCTTCTGGAGTCCCTGTTCGCTTCAGTGGCAGTGGATCTGGGACCTCTTATTCTCTCACAATCAGCAGCATGGAGGCTGAAGATGCTGCCACTTATTACTGTCAACAGTGGAGTAGTTACCCACTCA |
| IGKV8-30*01_S0907_AKR | AKR/J | Musmus IGKV8-30*01 F | 95.62 | 304 | GACATTGTGATGTCACAGTCTCCATCCGCCCTAGCTGTGTCAGTTGGAGAGAAGGTCACTATGAGCTGCAAGTCCAGTCAGAGCCTTTTATATAGTAGCAATCAAAAGAACTACTTGGCCTGGTACCAGCAGAAACCAGGGCAGTCTCCTAAACTGTTAATCTACTGGGCATCCACTAGGGAATCTGGGGTCCCTGACCGCTTCACAGGCAGTAGATCAGGGACAGATTTCACTCTCACCATCAGCAGTGTGCAGGCTGAAGACCTGGCCGTTTATTACTGCAAGCAATCTTATAGCTATCCTC |
| IGKV19-93*01_S8964_AKR | AKR/J | Musmus IGKV19-93*01 F | 98.21 | 287 | GACATCCAGATGACACAGTCTCCATCCTCACTGTCTGCATCTCTGGGAGGCAAAGTCACCATCACTTGCAAGGCAAGCCAAGACATTAACAAGAATATAGCTTGGTACCAACACAAGCCTGGAAAAGGTCCTAGGCTGCTCATATGGTACACATCTACATTACAGCCAGGCATCCCATCAAGGTTCAGTGGAAGTGGGTCTGGGAGAGATTATTCCTTCAGCATCAGCAACCTGGAGCCTGAAGATATTGCAACTTATTACTGTCTACAGTATGATAATCTTCCACC |
| IGKV9-120*01_S2628_AKR | AKR/J | Musmus IGKV9-120*02 F | 100 | 285 | GACATCCAGATGACCCAGTCTCCATCCTCCTTATCTGCCTCTCTGGGAGAAAGAGTCAGTCTCACTTGTCGGGCAAGTCAGGACATTGGTAGTAGCTTAAACTGGCTTCAGCAGGAACCAGATGGAACTATTAAACGCCTGATCTACGCCACATCCAGTTTAGATTCTGGTGTCCCCAAAAGGTTCAGTGGCAGTAGGTCTGGGTCAGATTATTCTCTCACCATCAGCAGCCTTGAGTCTGAAGATTTTGTAGACTATTACTGTCTACAATATGCTAGTTCTCCT |
| IGKV8-34*01_S3615_AKR | AKR/J | Musmus IGKV8-34*01 F | 99.33 | 305 | GACATTTTGATGACTCAGTCTCCATCCTCCCTGACTGTGTCAGCAGGAGAGAAGGTCACTATGAGCTGCAAGTCCAGTCAGAGTCTTTTAGCTAGTGCCAACCAAAATAACTACTTGGCCTGGCACCAGCAGAAACCAGGACGATCTCCTAAAATGCTGATAATTTGGGCATCCACTAGGGTATCTGGAGTCCCTGATCGCTTCATAGGCAGTGGATCTGGGACGGATTTCACTCTGACCATCAACAGTGTGCAGGCTGAAGATCTGGCTGTTTATTACTGTCAGCAGTCCTACAGCGCTCCTAC |

| | | | | | |
|---|---|---|---|---|---|
| IGKV3-10*01_S5437_AKR | AKR/J | Musmus IGKV3-10*01 F | 98.97 | 299 | AAAATTGTGCTGACCCAATCTCCAGCTTCTTTGGCTGTGTCTCTAAGGCAGAGGGCCACCATATCCTGCAGAGCCAGTGAAAGTGTTGATAGTTATGGCAATAGTTTTATGCACTGGTACCAGCAGAAACCAGGACAGCCACCCAAACTCCTCATCTATCGTGCATCCAACCTAGAATCTGGGGTCCCTGCCAGGTTCAGTGGCAGTGGGTCTAGGACAGACTTCACCCTCACCATTGATCCTGTGGAGGCTGATGATGCTGCAACCTATTACTGTCAGCAAAATAATGAGGATCCTCC |
| IGKV4-50*01_S2188_AKR | AKR/J | Musmus IGKV4-50*01 F | 97.1 | 286 | GAAAATGTGCTCACCCAGTCTCCAGCAATCATGTCTGCAACTCTAGGGGAGAAGGTCACCATGAGCTGCAGGGCCAGCTCAAATGTAAAGTACATGTACTGGTACCAGCAGAAGTCAGGTGCCTCCCCCAAACTATGGATTTATTACACATCCAACCTGGCTTCTGGAGTCCCAGCTCGCTTCAGTGGCAGTGGGTCTGGGACCTCTTATTCTCTCACAATCAGCAGCGTGGAGGCTGAAGATGCTGCCACTTATTACTGCCAGCAGTTTACTAGTTCCCCATCCA |
| IGKV8-16*01_S7495_AKR | AKR/J | Musmus IGKV8-16*01 F | 99.33 | 305 | GAAATTGTGTTGACGCAGTCTATACCATCCCTGACTGTGTCAGCAGGAGAGAGGGTCACTATCAGCTGCAAATCCAATCAGAATCTTTTATGGAGTGGAAACCAAAGGTACTGTTTGGTCTGGCACCAGTGGAAACCAGGGCAAACTCCTACACCGTTGATCACCTGGACATCTGATAGATACTCTGGAGTCCCTGATCGTTTCATAGGCAGTGGATCTGTGACAGATTTCACTCTGACCATCAGCAGTGTGCAGGCTGAAGATGTGGCAGTTTATTTCTGTCAGCAGCATTTACACATTCCTCC |
| IGKV3-7*01_S6705_AKR | AKR/J | Musmus IGKV3-7*02 F | 97.59 | 299 | GACATTGTGCTGACACAGTCTCCTGCTTCCTTAGCTGTATCTCTGGGGCAGAGGGCCACCATCTCCTGCAGGGCCAGCAAAAGTGTCAGTACATCTAGCTATAGTTACATGCACTGGTACCAACAGAAACCAGGACAGCCACCCAAACTCCTCATCAAGTATGCATCCTACCTAGAATCTGGGGTTCCTGCCAGGTTCAGTGGCAGTGGGTCTGGGACAGACTTCACCCTCAACATCCATCCTGTGGAGGAGGAGGATGCTGCAACATATTACTGTCAGCACAGTAGGGAGTTTCCTCC |
| IGKV4-91*01_S4811_AKR | AKR/J | Musmus IGKV4-91*01 F | 95.74 | 292 | GAAATTGTGCTTACCCAGTCTCCAACCACCATGGCTGCATCTCCTGGGGAGAAGGTCACCATCACCTGCAGTGCCAGCTCAAGTATAAGTTCCAATTACTTGCACTGGTATCAGCAGAAGCCAGGATTCCCTCCTAAACTCTTGATATATAGGACATCCAATCTGGCTTCTGGAGTCCCAGCTCGCTTCAGTGGCAGTGGGTCTGGGACCTCTTACTCTCTCACAATTGGCACCATGGAGGCTGAAGGTGCTGCCACTTATTACTGCCAGCATGGTAGTAGTTTACTACGCA |
| IGKV4-57-1*01_S8603_AKR | AKR/J | Musmus IGKV4-57-1*01 F | 92.91 | 292 | CAAATTGTTCTCACCCAGTCTCCAGCAATCATGTCTGCCTCTCCAGGGGAGAAGGTCACCATGACCTGCAGTGCCAGCTCAAGTGTAAGTTCCAGGTACTTGCACTGGTACCAGCAGAAGTCAGGAGCCTCCCCCAAACTCTGGATTTATGGCACATCCAACCTGGCTTCTGGAGTCCCTGCTCGCTTCAGTGGCAGTGGGTCTGGGACCTCTTACTCTCTCACAATCAGCAGCGTGGAGGCTGAAGATGCTGCCACTTATTACTGCCAGCAGTATCATAGTGACCCACTCA |
| IGKV4-62*01_S2199_AKR | AKR/J | Musmus IGKV4-62*01 ORF | 96.74 | 286 | CAAATTGTTCTCACCCAGTCTCCAGCAATCATGTCTGCATCTCCAGGGGAGAAGGTCACCATGACCTGCAGTGCCAGCTCAAGTGTAAGTTACATGTACTGGTACCAGCAGAAGCCAGGATCCTCCCCCAGACTCTGGATTTATGACACATCCAACCTGGCTTCTGGAGTCCCCGCTCGCTTCAGTGGCAGTAGGTCTGGGACCTCTTATTCTCTCACAATCAGCAGCATGGAGGCTGAAGATGCTGCCACTTATTACTGCCATCAGCGGAGTAGTTACCCATTCA |

| | | | | | |
|---|---|---|---|---|---|
| IGKV12-41*01_S6800_AKR | AKR/J | Musmus IGKV12-41*01 F | 97.13 | 287 | GATATCCAGATGACTCAGTCTCCAGCCTCCCTATCTGCATCTGTGGGAGAAACTGTCACCATCACATGTCGAGCAAGTGGGAATATTCACAATTATTTAGCATGGTATCAGCAGAAACAGGGAAAATCTCCTCAGCTCCTGGTCTATAATGCAAAAACCTTAGCGGAAGGTGTGCCATCAAGGTTCAGTGGCAGTGGATCAGGAACACAATATTCTCTCAAGATCAACAGCCTGCAGCCTGAGGATTTTGGGAGTTATTACTGTCATCATTATTATAGTACTCCTCC |
| IGKV3-4*01_S3473_AKR | AKR/J | Musmus IGKV3-4*01 F | 92.44 | 299 | GACATTGTGCTGACCCAATCTCCAGCTTCTTTGGCTGTGTCTCTAGGGCAGAGGGCCACCATATCCTGCAAGGCCAGCCAGAGTATCAATTATTACGGTGATAATTATATGCACTGGTTCCAACAGAAACCAGGACAGCCAACCAAACTCCTTATCTATGATGCATCCAACCTAGAATCTGGAGTTCCTGACAGGTTCAGTGGCAGTGGGTCTGGGACAGACTTCACCCTCAACATCCATCCTGTGGAGGAGGAGGATGCTGCAACCTATTACTGTCAGCAAAGCAAGGAGTTTCCTCC |
| IGKV4-80*01_S4247_AKR | AKR/J | Musmus IGKV4-80*01 F | 99.64 | 282 | CAAATTGTTCTCACCCAGTCTCCAGCAATCATGTCTGCATCTCTAGGGGAGGAGATCACCCTAACCTGCAGTGCCAGCTCGAGTGTAAGTTACATGCACTGGTTCCAGCAGAAGTCAGGCACTTCTCCCAAACTCTTGATTTATAGCACATCCAACCTGGCTTCTGGAGTCCCTTCTCGCTTCAGTGGCAGTGGGTCTGGGACCTTTTATTCTCTCACAATCAGCAGTGTGGAGGCTGAAGATGCTGCCGATTATTACTGCCATCAGTGGAGTAGTTATCCA |
| IGKV15-103*01_AKR | AKR/J | Musmus IGKV15-103*01 ORF | 100 | 287 | GACATCCAGATGAACCAGTCTCCATCCAGTCTGTCTGCATCCCTTGGAGACACAATTACCATCACTTGCCATGCCAGTCAGAACATTAATGTTTGGTTAAGCTGGTACCAGCAGAAACCAGGAAATATTCCTAAACTATTGATCTATAAGGCTTCCAACTTGCACACAGGCGTCCCATCAAGGTTTAGTGGCAGTGGATCTGGAACAGGTTTCACATTAACCATCAGCAGCCTGCAGCCTGAAGACATTGCCACTTACTACTGTCAACAGGGTCAAAGTTATCCTCT |
| IGKV4-57-1*01_S1729_AKR | AKR/J | Musmus IGKV4-57*01 F | 100 | 292 | GAAATTGTGCTCACCCAGTCTCCAGCACTCATGGCTGCATCTCCAGGGGAGAAGGTCACCATGACCTGCAGTGCCAGCTCAAGTGTAGGTTCCAGTTACTTGCACTGGTACCAGCAGAAGTCAGGAGCCTCCCCCAAACTCTGGATTTACGGCACATCCAACCTGGCTTCTGGAGTCCCTGCTTGCTTCAGTGGCAGTGGGTCTGGGACCTCTTACTCTCTCACAATCAGCAGCGTGGAGGCTGAAGATGATGCAACTTATTACTGCCAGCAGGGGTGGGATTACCCACTCA |
| IGKV16-104*01_AKR | AKR/J | Musmus IGKV16-104*01 F | 100 | 288 | GATGTCCAGATAACCCAGTCTCCATCTTATCTTGCTGCATCTCCTGGAGAAACCATTACTATTAATTGCAGGGCAAGTAAGAGCATTAGCAAATATTTAGCCTGGTATCAAGAGAAACCTGGGAAAACTAATAAGCTTCTTATCTACTCTGGATCCACTTTGCAATCTGGAATTCCATCAAGGTTCAGTGGCAGTGGATCTGGTACAGATTTCACTCTCACCATCAGTAGCCTGGAGCCTGAAGATTTTGCAATGTATTACTGTCAACAGCATAATGAATACCCGTAC |
| IGKV2-109*01_AKR | AKR/J | Musmus IGKV2-109*01 F | 100 | 302 | GATATTGTGATGACGCAGGCTGCATTCTCCAATCCAGTCACTCTTGGAACATCAGCTTCCATCTCCTGCAGGTCTAGTAAGAGTCTCCTACATAGTAATGGCATCACTTATTTGTATTGGTATCTGCAGAAGCCAGGCCAGTCTCCTCAGCTCCTGATTTATCAGATGTCCAACCTTGCCTCAGGAGTCCCAGACAGGTTCAGTAGCAGTGGGTCAGGAACTGATTTCACACTGAGAATCAGCAGAGTGGAGGCTGAGGATGTGGGTGTTTATTACTGTGCTCAAAATCTAGAACTTCCTCC |
| IGKV10-96*03_S5544_AKR | AKR/J | Musmus IGKV10-96*03 F | 99.64 | 287 | GATATCCAGATGACACAGACTACATCCTCCCTGTCTGCCTCTCTGGGAGACAGAGTCACCATCAGTTGCAGGGCAAGTCAGGACATTAGCAATTATTTAAACTGGTATCAGCAGAAACCAGATGGAACTGTTAAACTCCTGATCTACTACACATCAAGATTACACTCAGGAGTCCCATCAAGGTTCAGTGGCAGTGGGTCTGGGACAGATTATTCTCTCACTATCAGCAACCTGGAACAAGAAGATATTGCCACTTACTTTTGCCAACAGGATAGTAAGCATCCTCC |

| | | | | | |
|---|---|---|---|---|---|
| IGKV2-112*01_AKR | AKR/J | Musmus IGKV2-112*01 F | 100 | 300 | GATATTGTGATAACCCAGGATGAACTCTCCAATCCTGTCACTTCTGGAGAATCAGTTTCCATCTCCTGCAGGTCTAGTAAGAGTCTCCTATATAAGGATGGGAAGACATACTTGAATTGGTTTCTGCAGAGACCAGGACAATCTCCTCAGCTCCTGATCTATTTGATGTCCACCCGTGCATCAGGAGTCTCAGACCGGTTTAGTGGCAGTGGGTCAGGAACAGATTTCACCCTGGAAATCAGTAGAGTGAAGGCTGAGGATGTGGGTGTGTATTACTGTCAACAACTTGTAGAGTATCCT |
| IGKV5-37*01_S9729_AKR | AKR/J | Musmus IGKV5-37*01 F | 97.13 | 285 | GACATCCTGATGACCCAGTCTCCAGCCACCCTGTCTGTGACTCCAGGAGAAACAGTCAGTCTTTCCTGTAGGGCCAGCCAGAATACTTACAAGAACCTACACTGGTATCAACAGAAATCACATGGGACTCCAAAGCTTCTCATCAAGTATGCATCTGATCCCATCTCTGGGATCCCCTCCAGGTTCACTGGCAGTGGATCAGGGACAGATTACACTCTCAGTATCAACAGTGTGAAGCCCGAAGATGAAGGAATATATTACTGTCTTCAAGGTTACAGCATGCCT |
| IGKV14-111*01_AKR | AKR/J | Musmus IGKV14-111*01 F | 100 | 287 | GACATCAAGATGACCCAGTCTCCATCTTCCATGTATGCATCTCTAGGAGAGAGAGTCACTATCACTTGCAAGGCGAGTCAGGACATTAATAGCTATTTAAGCTGGTTCCAGCAGAAACCAGGGAAATCTCCTAAGACCCTGATCTATCGTGCAAACAGATTGGTAGATGGGGTCCCATCAAGGTTCAGTGGCAGTGGATCTGGGCAAGATTATTCTCTCACCATCAGCAGCCTGGAGTATGAAGATATGGGAATTTATTATTGTCTACAGTATGATGAGTTTCCTCC |
| IGKV14-100*01_S4794_AKR | AKR/J | Musmus IGKV14-100*01 F | 99.64 | 287 | GACATCCTGATGACCCAATCTCCATCCTCCATGTCTGTATCTCTGGGAGACACAGTCAGCATCACTTGCCATGCAAGTCAGGGCATTAGCAGTAATATAGGGTGGTTGCAGCAGAAACCAGGGAAATCATTTAAGGGCCTGATCTATCATGCAACCAACTTGGAAGATGGAGTTCCATCAAGGTTCAGTGGCAGTGGATCTGGAGCAGATTATTCTCTCACCATCAGCAGCCTGGAATCTGAAGATTTTGCAGACTATTACTGTGTACAGTATGCTCAGTTTCCTCC |
| IGKV4-74*01_S3912_AKR | AKR/J | Musmus IGKV4-74*01 F | 95.04 | 292 | CAAATTGTTCTCACCCAGTCTCCAGCAATCATGTCTGCCTCTCCAGGGGAGAAGGTCACCATGACCTGCAGTGCCAGCTCAAGTGTAAGTTCCAGGTACTTGCACTGGTACCAGCAGAAGTCAGGAGCCTCCCCCAAACTCTGGATTTATGGCACATCCAACCTGGCTTCTGGAGTCCCTGCTCGCTTCAGTGGCAGTGGGTCTGGGACCTCTTACTCTCTCACAATCAGCAGCGTGGAGGCTGAAGATGCTGCCACTTATTACTGCCAGCAGTATCATAGTGACCCACCCA |
| IGKV4-78*01_S3978_AKR | AKR/J | Musmus IGKV4-78*01 F | 97.87 | 292 | CAAATTGTTCTCACCCAGTCTCCAGCAATCATGTCTGCATCTCCTGGGGAACGGGTCACCATGACCTGCAGTGCCAGCTCAAGTGTAAGTTCCAGCTACTTGTACTGGTACCAGCAGAAGCCAGGATCCTCCCCCAAACTATGGATTTATAGCACATCCAACCTGGCTTCTGGAGTCCCTGCTCGCTTCAGTGGCAGTGGGTCTGGGACCTCTTATTCTCTCACAATCAGCAGCATGGAGGCTGAAGATGCTGCCACTTATTACTGCCAGCAGTACAGTGGTTACCCATCCA |
| IGKV8-28*02_AKR | AKR/J | Musmus IGKV8-28*02 F | 100 | 302 | GACATTGTGATGACACAGTCTCCATCCTCCCTGAGTGTGTCAGCAGGAGATAAGGTCACTATGAGCTGCAAGTCCAGTCAGAGTCTGTTAAACAGTAGAAACCAAAAGAACTACTTGGCCTGGTACCAGCAGAAACCATGGCAGCCTCCTAAACTGCTGATCTACGGGGCATCCACTAGGGAATCTGGGGTCCCTGATCGCTTCACAGGCAGTGGATCTGGAACAGATTTCACTCTCACCATCAGCAGTGTGCAGGCTGAAGACCTGGCAGTTTATTACTGTCAGAATGATTATAGTTATCC |
| IGKV9-124*01_AKR | AKR/J | Musmus IGKV9-124*01 F | 100 | 285 | GACATCCAGATGACCCAGTCTCCATCCTCCTTATCTGCCTCTCTGGGAGAAAGAGTCAGTCTCACTTGTCGGGCAAGTCAGGAAATTAGTGGTTACTTAAGCTGGCTTCAGCAGAAACCAGATGGAACTATTAAACGCCTGATCTACGCCGCATCCACTTTAGATTCTGGTGTCCCAAAAAGGTTCAGTGGCAGTAGGTCTGGGTCAGATTATTCTCTCACCATCAGCAGCCTTGAGTCTGAAGATTTTGCAGACTATTACTGTCTACAATATGCTAGTTATCCT |

| | | | | | |
|---|---|---|---|---|---|
| IGKV17-127*01_AKR | AKR/J | Musmus IGKV17-127*01 F | 100 | 287 | GAAACAACTGTGACCCAGTCTCCAGCATCCCTGTCCGTGGCTACAGGAGAAAAAGTCACTATCAGATGCATAACCAGCACTGATATTGATGATGATATGAACTGGTACCAGCAGAAGCCAGGGGAACCTCCTAAGCTCCTTATTTCAGAAGGCAATACTCTTCGTCCTGGAGTCCCATCCCGATTCTCCAGCAGTGGCTATGGCACAGATTTTGTTTTTACAATTGAAAACACGCTCTCAGAAGATGTTGCAGATTACTACTGTTTGCAAAGTGATAACATGCCTCT |
| IGKV8-16*01_S1056_AKR | AKR/J | Musmus IGKV8-16*01 F | 98.99 | 305 | GAAATTGTGTTGACACAGTCTATACCATCCCTGACTGTGTCAGCAGGAGAGAGGGTCACTATCAGCTGCAAATCCAATCAGAATCTTTTATGGAGTGGAAACCAACGGTACTGTTTGGTCTGGCACCAGTGGAAACCAGGGCAAACTCCTACACCGTTGATCACCTGGACATCTGATAGGTACTCTGGAGTCCCTGATCGTTTCATAGGCAGTGGATCTGTGACAGATTTCACTCTGACCATCAGCAGTGTGCAGGATGAAGATGTGGCAGTTTATTTCTGTCAGCAGCATTTACACATTCCTCC |
| IGKV17-121*01_AKR | AKR/J | Musmus IGKV17-121*01 F | 100 | 287 | GAAACAACTGTGACCCAGTCTCCAGCATCCCTGTCCATGGCTATAGGAGAAAAAGTCACCATCAGATGCATAACCAGCACTGATATTGATGATGATATGAACTGGTACCAGCAGAAGCCAGGGGAACCTCCTAAGCTCCTTATTTCAGAAGGCAATACTCTTCGTCCTGGAGTCCCATCCCGATTCTCCAGCAGTGGCTATGGTACAGATTTTGTTTTTACAATTGAAAACATGCTCTCAGAAGATGTTGCAGATTACTACTGTTTGCAAAGTGATAACTTGCCTCT |
| IGKV12-38*01_S2611_AKR | AKR/J | Musmus IGKV12-38*01 F | 98.21 | 287 | GACATCCAGATGACTCAGTCTCCAGCCTCCCTGGCTGCATCTGTGGGAGAAACCATCACCATCACATGTCAAGCAAGTGAGAACATTTACTTCAGTTTAGCATGGTATCAGCAGAAGCAAGGGAAATCTCCTCAGCTCCTGATCTATAATGCAAACAGCTTGGAAGATGGTGTCCCATCGAGGTTCAGTGGCAGTGGATCTGGGACACAGTATTCTATGAAGATCAACAGCATGCAGCCTGAAGATACTGCAACTTATTTCTGTAAACAGGCTTATGACTTTCCTCC |
| IGKV4-81*01_S2422_AKR | AKR/J | Musmus IGKV4-81*01 F | 96.45 | 292 | GAAAATGTGCTGACCCAGTCTCCAGCAATCATGGCTGCATCTCCAGGGGAGAAGGTCACCATGACCTGCAGTGCCAGCTCAAGTGTAAGTTCTGGTAACTTTCACTGGTACCAGCAGAAGCCAGGCACTTCTCCCAAACTCTGGATTTATAGGACATCCAACCTGGCTTCTGGAGTCCCCGCTCGCTTCAGTGGCAGTGGGTCTGGGACCTCTTACTCTCTTACAATCAGCAGCATGGAGGCCGAAGATGCTGCCACTTATTACTGCCAGCAGTGGAGTGGTTACCCACCCA |
| IGKV1-133*01_AKR | AKR/J | Musmus IGKV1-133*01 F | 100 | 302 | GATGTTGTGATGACCCAGACTCCACTCACTTTGTCGGTTACCATTGGACAACCAGCCTCTATCTCTTGCAAGTCAAGTCAGAGCCTCTTATATAGTAATGGAAAAACCTATTTGAATTGGTTATTACAGAGGCCAGGCCAGTCTCCAAAGCGCCTAATCTATCTGGTGTCTAAACTGGACTCTGGAGTCCCTGACAGGTTCACTGGCAGTGGATCAGGAACAGATTTTACACTGAAAATCAGCAGAGTGGAGGCTGAGGATTTGGGAGTTTATTACTGCGTGCAAGGTACACATTTTCCTCA |
| IGKV8-18*01_S2898_AKR | AKR/J | Musmus IGKV8-18*01 ORF | 99.66 | 310 | GACATTGTGATGACCCAGTCTCCATCCTCCCTGGCTGTGACAGCAGGAGAGAAGGTCACTATGAGATGCAAGTCCAGTCAGAGTCTTTTGTGGAGTGTAAACCAAAAGAACTACTTATCCTGGTACCAGCAGAAACAAGGGCAGCCTCCTAAACTGCTTATCTATGGGGCATCCATTAGAGAATCTTGGGTCCCTGATCGATTCACAGGAAGTGGATCTGGGACAGACTTCACTCTCACCATTAGCAATGTGCATGCTGAAGACCTAGCAGTTTATTACTGTCAGCACAATCATGGCAGCTTTCTCCCCC |

| | | | | | |
|---|---|---|---|---|---|
| IGKV1-122*01_AKR | AKR/J | Musmus IGKV1-122*01 F | 100 | 302 | GATGCTGTGATGACCCAAACTCCACTCTCCCTGCCTGTCAGTCTTGGAGATCAAGCCTCCATCTCTTGCAGGTCTAGTCAGAGCCTTGAAAACAGTAATGGAAACACCTATTTGAACTGGTACCTCCAGAAACCAGGCCAGTCTCCACAGCTCCTGATCTACAGGGTTTCCAACCGATTTTCTGGGGTCCTAGACAGGTTCAGTGGTAGTGGATCAGGGACAGATTTCACACTGAAAATCAGCAGAGTGGAGGCTGAGGATTTGGGAGTTTATTTCTGCCTCCAAGTTACACATGTCCCTCC |
| IGKV4-86*01_S4316_AKR | AKR/J | Musmus IGKV4-86*01 F | 100 | 283 | GAAATTGTGCTCACTCAGTCTCCAGCCATCACAGCTGCATCTCTGGGGCAAAAGGTCACCATCACCTGCAGTGCCAGCTCAAGTGTAAGTTACATGCACTGGTACCAGCAGAAGTCAGGCACCTCCCCCAAACCATGGATTTATGAAATATCCAAACTGGCTTCTGGAGTCCCAGCTCGCTTCAGTGGCAGTGGGTCTGGGACCTCTTACTCTCTCACAATCAGCAGCATGGAGGCTGAAGATGCTGCCATTTATTACTGCCAGCAGTGGAATTATCCACTTA |
| IGKV11-125*01_AKR | AKR/J | Musmus IGKV11-125*01 F | 100 | 287 | GATGTCCAGATGATTCAGTCTCCATCCTCCCTGTCTGCATCTTTGGGAGACATAGTCACCATGACTTGCCAGGCAAGTCAGGGCACTAGCATTAATTTAAACTGGTTTCAGCAAAAACCAGGGAAAGCTCCTAAGCTCCTGATCTATGGTGCAAGCAACTTGGAAGATGGGGTCCCATCAAGGTTCAGTGGCAGTAGATATGGGACAGATTTCACTCTCACCATCAGCAGCCTGGAGGATGAAGATATGGCAACTTATTTCTGTCTACAGCATAGTTATCTCCCTCC |
| IGKV1-99*01_S3206_AKR | AKR/J | Musmus IGKV1-99*01 F | 98.3 | 302 | GATGTTGTTCTGACCCAAACTCCACTCTCTCTGCCTGTCAATATTGGAGATCAAGCCTCTATCTCTTGCAAGTCTACTAAGAGCCTTCTGAATAGTGATGGATTCACTTATTTGGGCTGGTACCTGCAGAAGCCAGGCCAGTCTCCACAGCTCCTAATATATTTGGTTTCTAATCGATTTTCTGGAGTTCCAGACAGGTTCAGTGGTAGTGGGTCAGGGACAGATTTCACCCTCAAGATCAGCAGAGTGGAGGCTGAGGATTTGGGAGTTTATTATTGCTTCCAGAGTAACTATCTTCCTCT |
| IGKV5-39*01_S3142_AKR | AKR/J | Musmus IGKV5-39*01 F | 98.92 | 287 | GACATTGTGATGACTCAGTCTCCAGCCACCCTGTCTGTGACTCCAGGAGATAGGGTCTCTCTTTCCTGCAGGGCCAGTCAGAGTATTAGCGACTACTTACACTGGTATCAACAAAAATCACATGAGTCTCCAAGGCTTCTCATCAAGTATGCTTCCCAATCCATCTCTGGGATCCCCTCCAGGTTCAGTGGCAGTGGATCAGGGTCAGATTTCACTCTCAGTATCAACAGTGTGGAACCTGAAGATGTTGGAGTGTATTACTGTCAAAATGGTCACAGCTTTCCTCC |
| IGKV9-123*01_AKR | AKR/J | Musmus IGKV9-123*01 F | 100 | 285 | GACATCCAGATGATTCAGTCTCCATCGTCCATGTTTGCCTCTCTGGGAGACAGAGTCAGTCTCTCTTGTCGGGCTAGTCAGGGCATTAGAGGTAATTTAGACTGGTATCAGCAGAAACCAGGTGGAACTATTAAACTCCTGATCTACTCCACATCCAATTTAAATTCTGGTGTCCCATCAAGGTTCAGTGGCAGTGGGTCTGGGTCAGATTATTCTCTCACCATCAGCAGCCTAGAGTCTGAAGATTTTGCAGACTATTACTGTCTACAGCGTAATGCGTATCCT |
| IGKV16-104*01_S9985_AKR | AKR/J | Musmus IGKV14-87-2*01 F | 99.64 | 287 | GACATCCAAATGACCCAGTCCTCATCATTTCTCTCTGCATCTCTAGGAGATCATCTTACAATCAACTGCAGGGCCAGTAAGGATATTAACAAGTATTTTGCTTGGGTTCAACAGAAGCCAGGGAAGGCTCCAAGGATGTTGATTCATTTTGCTTCCACCTTGCTACCTGGGGTTCCAGAAAAGTTCAGTGGGAGTGGATCTGGGACAGATTTTTCTCTCACTATCAGAAACATAGAGTCTGAAGATATTGCAATGTATTACTGTCTACAGTATTCTGAGCATCCGTA |
| IGKV1-132*01_AKR | AKR/J | Musmus IGKV1-132*01 F | 100 | 302 | GATGTTGTGATGACCCAGACTCCACTGTCTTTGTCGGTTACCATTGGACAACCAGCCTCTATCTCTTGCAAGTCAAGTCAGAGCCTCTTATATAGTAATGGAAAGACATATTTGAATTGGTTACAACAGAGGCCTGGCCAGGCTCCAAAGCACCTAATGTATCAGGTGTCCAAACTGGACCCTGGCATCCCTGACAGGTTCAGTGGCAGTGGATCAGAAACAGATTTTACACTTAAAATCAGCAGAGTGGAGGCTGAAGATTTGGGAGTTTATTACTGCTTGCAAGGTACATATTATCCTCA |

| | | | | | |
|---|---|---|---|---|---|
| IGKV6-25*01_S3519_AKR | AKR/J | Musmus IGKV6-25*01 F | 98.21 | 287 | TGCATTGTGATGACCCAGTCTCACAAATTCATGTCCACATCAGTAGGAGACAGGGTCAGCATCACCTGCAAGGCCAGTCAGGATGTGAGTACTGCTGTAGCCTGGTATCAACAAAAACCAGGGCAATCTCCTAAACTACTGATTTACTGGGCATCCACCCGGCACACTGGAGTCCCTGATCGCTTCACAGGCAGTGGATCTGGGACAGATTTCACTCTTACCATCAGCAGTGTGCAGGCTGAAGACCTGGCACTTTATTACTGTCAGCAACATTATAGCACTCCTCC |
| IGKV13-85*01_S1266_AKR | AKR/J | Musmus IGKV13-85*01 F | 97.13 | 287 | GACATCCAGATGACACAATCTTCATCCTCCTTGTCTGTATCTCTAGGAGACAGAGTCACCATTACTTGCAAGGCAAGTGAGCACATTAATAGTTGGTTAGCCTGGTATCAGCAAAAACCAGGAAATGCTCCTAGGCTCTTAATATCTGGTGCAACCAGTTTGGAAACTGGGGTTCCTTCAAGATTCAGTGGCAGTGCATCTGGAAAGGATTACACTCTCAGCATTACTAGTCTTCAGACTGAAGATGTTGCTACTTATTACTGTCAACAGTATTGGGGTACTCCTCC |
| IGKV1-88*01_S0716_AKR | AKR/J | Musmus IGKV1-88*01 F | 99.66 | 302 | GATGTTGTGGTGACTCAAACTCCACTCTCCCTGCCTGTCAGCTTTGGAGATCAAGTTTCTATCTCTTGCAGGTCTAGTCAGAGTCTTGCAAACAGTTATGGGAACACCTATTTGTCTTGGTACCTGCACAAGCCTGGCCAGTCTCCACAGCTCCTCATCTATGGGATTTCCAACAGATTTTCCGGGGTGCCAGACAGGTTCAGTGGCAGTGGTTCAGGGACAGATTTCACACTCAAGATCAGCACAATAAAGCCTGAGGACTTGGGAATGTATTACTGCTTACAAGGTACACATCAGCCTCC |
| IGKV3-9*01_S5964_AKR | AKR/J | Musmus IGKV3-9*01 F | 90.38 | 299 | GACATTGTGCTGACCCAATCTCCAGCTTCTTTGGCTGTGTCTCTAGGGCAGAGGGCCACCATCTCCTGCAGAGCCAGCGAAAGTGTCAGTATTCGTGGTACTAGTTTAATGCACTGGTACCAACAGAAACCAGGATATCCACCCAAACTCCTCATCTATGCTGCATCCAACCTAGAATCTGGGGTTCCTGCCAGGTTTAGTGGCAGAAGGTCTGGGACAGACTTCACCCTCAACATTCATCCTGTGGAGGAAGATGATGCTGCAACCTATTACTGTCAGCAAAGTAGGGAATATCCTCC |
| IGKV6-25*01_S9703_AKR | AKR/J | Musmus IGKV6-25*01 F | 97.49 | 287 | GACATTGTGATGACCCAGTCTCACAAATTCATGTCCACATCAGTAGGAGACAGGGTCAGCATCACCTGCAAGGCCAGTCAGGATGTGGGTACTGCTGTAGCCTGGTATCAACAGAAACCAGGGCAATCTCCTAAACTACTGATTTACTGGGCATCCACCCGGCACACTGGAGTCCCTGATCCCTTCACAGGCAGTGGATCTGGGACAGATTTCACTCTCACCATCAGCAGTGTGCAGGCTGAAGACCTGGCAGTTTATTACTGTCAGCAAGATTATAGCACTCCTCC |
| IGKV9-129*01_AKR | AKR/J | Musmus IGKV9-129*01 ORF | 100 | 287 | GACATCCAGATGACCCAGTCTCCATCCTCCTTATCTGCCTCTCTGGGAGAAAGAGTCAGTCTCACTTGCCGGGCAAGTCAGGACATTCATGGTTATTTAAACTTGTTTCAGCAGAAACCAGGTGAAACTATTAAACACCTGATCTATGAAACATCCAATTTAGATTCTGGTGTCCCAAAAAGGTTCAGTGGCAGTAGGTCTGGGTCAGATTATTCTCTCATTATCGGCAGCCTTGAGTCTGAAGATTTTGCAGACTATTACTGTCTACAATATGCTAGTTCTCCTCC |
| IGKV14-130*01_AKR | AKR/J | Musmus IGKV14-130*01 F | 100 | 287 | GAAATCCAGATGACCCAGTCTCCATCCTCTATGTCTGCATCTCTGGGAGACAGAATAACCATCACTTGCCAGGCAACTCAAGACATTGTTAAGAATTTAAACTGGTATCAGCAGAAACCAGGGAAACCCCCTTCATTCCTGATCTATTATGCAACTGAACTGGCAGAAGGGGTCCCATCAAGGTTCAGTGGCAGTGGGTCTGGGTCAGACTATTCTCTGACAATCAGCAACCTGGAGTCTGAAGATTTTGCAGACTATTACTGTCTACAGTTTTATGAGTTTCCTCC |
| IGKV18-36*01_S0889_AKR | AKR/J | Musmus IGKV18-36*01 F | 99.28 | 287 | ACTGGAGAAACAACACAGTCTCCAGCTTCTCTGAGTTTTTCTCTTGGTGAAACAGCAACACTGTCATGCAGGTCCAGTGAGAGTGTTGGCAGCTACTTAGCCTGGTACCAGCAGAAAGCAGAGCAAGTTCCCCGGCTCCTTATCCATAGTGCCTCCACTAGGGCCGGTGGTGTCCCAGTCCGATTCAGTGGCACTGGGTCTGGGACAGACTTCACTCTCACCATCAGCAGTCTAGAACCTGAAGATGCTGCAGTTTACTACTGTCAACCTTTCAAAAGTTGGTCATA |

| | | | | | |
|---|---|---|---|---|---|
| IGKV1-35*01_S0416_AKR | AKR/J | Musmus IGKV1-35*01 ORF | 97.28 | 301 | GACATTGTGATGACCCAGACTCCACTCACTTTATCAGCTACCATTGGACAATCAGCCTCCATCTCTTGCAGGTCAAGTCAGAGTCTCTTACATAGTAATGGAAACACATACTTGAATTGGTTTCTACAGAGGCCAGGCCAATCTCCACAGCTTCTGATTTATGGGGTGTTTGAACGGGAATCTGGGGTTCCTGACAGGTTCAGTGGCAGTGGGTCAGGAACAGATTTCACACTCAAGATCAGCAGAGTGGAGGCTGAGGATTTGGGAGTTTATTACTGCATGCAAGCTACCTATGAACCTC |
| IGKV4-92*01_S4902_AKR | AKR/J | Musmus IGKV4-92*01 F | 98.58 | 291 | GAAATGGTTCTCACCCAGTCTCCAGTATCCATAACTGCATCTCGAGGGGAGAAGATCACCATCACCTGCCGTGCCAGCTCAAGTATAAGTTCCAATTACTTACACTGGTACCAGCAGAAGCCAGGATCCTCCCCTAAACTTTTGATTTATAGGACATCCATCCTGGCATCTGGAGTCCTAGACAACTTCAGTGGCAGTGGGTCTGAGAGCTCTTACACTCTGACAATCAGCTGCATGCAGGACGATGTTGCTGCCACTTACTACTGTCAGCAGGGGAGTAGTAGCCCACCA |
| IGKV8-27*01_AKR | AKR/J | Musmus IGKV8-27*01 F | 100 | 300 | AACATTATGATGACACAGTCGCCATCATCTCTGGCTGTGTCTGCAGGAGAAAAGGTCACTATGAGCTGTAAGTCCAGTCAAAGTGTTTTATACAGTTCAAATCAGAAGAACTACTTGGCCTGGTACCAGCAGAAACCAGGGCAGTCTCCTAAACTGCTGATCTACTGGGCATCCACTAGGGAATCTGGTGTCCCTGATCGCTTCACAGGCAGTGGATCTGGGACAGATTTTACTCTTACCATCAGCAGTGTACAAGCTGAAGACCTGGCAGTTTATTACTGTCATCAATACCTCTCCTCG |
| IGKV3-5*01_AKR | AKR/J | Musmus IGKV3-5*01 F | 100 | 299 | GACATTGTGCTGACCCAATCTCCAGCTTCTTTGGCTGTGTCTCTAGGGCAGAGGGCCACCATATCCTGCAGAGCCAGTGAAAGTGTTGATAGTTATGGCAATAGTTTTATGCACTGGTACCAGCAGAAACCAGGACAGCCACCCAAACTCCTCATCTATCGTGCATCCAACCTAGAATCTGGGATCCCTGCCAGGTTCAGTGGCAGTGGGTCTAGGACAGACTTCACCCTCACCATTAATCCTGTGGAGGCTGATGATGTTGCAACCTATTACTGTCAGCAAAGTAATGAGGATCCTCC |
| IGKV4-55*01_AKR | AKR/J | Musmus IGKV4-55*01 F | 100 | 286 | CAAATTGTTCTCACCCAGTCTCCAGCAATCATGTCTGCATCTCCAGGGGAGAAGGTCACCATGACCTGCAGTGCCAGCTCAAGTGTAAGTTACATGTACTGGTACCAGCAGAAGCCAGGATCCTCCCCCAGACTCCTGATTTATGACACATCCAACCTGGCTTCTGGAGTCCCTGTTCGCTTCAGTGGCAGTGGGTCTGGGACCTCTTACTCTCTCACAATCAGCCGAATGGAGGCTGAAGATGCTGCCACTTATTACTGCCAGCAGTGGAGTAGTTACCCACCCA |
| IGKV4-72*01_AKR | AKR/J | Musmus IGKV4-72*01 F | 100 | 286 | CAAATTGTTCTCTCCCAGTCTCCAGCAATCCTGTCTGCATCTCCAGGGGAGAAGGTCACAATGACTTGCAGGGCCAGCTCAAGTGTAAGTTACATGCACTGGTACCAGCAGAAGCCAGGATCCTCCCCCAAACCCTGGATTTATGCCACATCCAACCTGGCTTCTGGAGTCCCTGCTCGCTTCAGTGGCAGTGGGTCTGGGACCTCTTACTCTCTCACAATCAGCAGAGTGGAGGCTGAAGATGCTGCCACTTATTACTGCCAGCAGTGGAGTAGTAACCCACCCA |
| IGKV6-32*01_AKR | AKR/J | Musmus IGKV6-32*01 F | 100 | 287 | AGTATTGTGATGACCCAGACTCCCAAATTCCTGCTTGTATCAGCAGGAGACAGGGTTACCATAACCTGCAAGGCCAGTCAGAGTGTGAGTAATGATGTAGCTTGGTACCAACAGAAGCCAGGGCAGTCTCCTAAACTGCTGATATACTATGCATCCAATCGCTACACTGGAGTCCCTGATCGCTTCACTGGCAGTGGATATGGGACGGATTTCACTTTCACCATCAGCACTGTGCAGGCTGAAGACCTGGCAGTTTATTTCTGTCAGCAGGATTATAGCTCTCCTCC |
| IGKV4-57*01_AKR | AKR/J | Musmus IGKV4-74*01 F | 95.04 | 286 | CAAATTGTTCTCACCCAGTCTCCAGCAATCATGTCTGCATCTCCAGGGGAGAAGGTCACCATAACCTGCAGTGCCAGCTCAAGTGTAAGTTACATGCACTGGTTCCAGCAGAAGCCAGGCACTTCTCCCAAACTCTGGATTTATAGCACATCCAACCTGGCTTCTGGAGTCCCTGCTCGCTTCAGTGGCAGTGGATCTGGGACCTCTTACTCTCTCACAATCAGCCGAATGGAGGCTGAAGATGCTGCCACTTATTACTGCCAGCAAAGGAGTAGTTACCCACCCA |

| | | | | | |
|---|---|---|---|---|---|
| IGKV6-15*01_BALB | BALB/cByJ | Musmus IGKV6-15*01 F | 100 | 287 | GACATTGTGATGACCCAGTCTCAAAAATTCATGTCCACATCAGTAGGAGACAGGGTCAGCGTCACCTGCAAGGCCAGTCAGAATGTGGGTACTAATGTAGCCTGGTATCAACAGAAACCAGGGCAATCTCCTAAAGCACTGATTTACTCGGCATCCTACCGGTACAGTGGAGTCCCTGATCGCTTCACAGGCAGTGGATCTGGGACAGATTTCACTCTCACCATCAGCAATGTGCAGTCTGAAGACTTGGCAGAGTATTTCTGTCAGCAATATAACAGCTATCCTCT |
| IGKV8-30*01_BALB | BALB/cByJ | Musmus IGKV8-30*01 F | 100 | 305 | GACATTGTGATGTCACAGTCTCCATCCTCCCTAGCTGTGTCAGTTGGAGAGAAGGTTACTATGAGCTGCAAGTCCAGTCAGAGCCTTTTATATAGTAGCAATCAAAAGAACTACTTGGCCTGGTACCAGCAGAAACCAGGGCAGTCTCCTAAACTGCTGATTTACTGGGCATCCACTAGGGAATCTGGGGTCCCTGATCGCTTCACAGGCAGTGGATCTGGGACAGATTTCACTCTCACCATCAGCAGTGTGAAGGCTGAAGACCTGGCAGTTTATTACTGTCAGCAATATTATAGCTATCCTCC |
| IGKV12-44*01_BALB | BALB/cByJ | Musmus IGKV12-44*01 F | 100 | 287 | GACATCCAGATGACTCAGTCTCCAGCCTCCCTATCTGCATCTGTGGGAGAAACTGTCACCATCACATGTCGAGCAAGTGAGAATATTTACAGTTATTTAGCATGGTATCAGCAGAAACAGGGAAAATCTCCTCAGCTCCTGGTCTATAATGCAAAAACCTTAGCAGAAGGTGTGCCATCAAGGTTCAGTGGCAGTGGATCAGGCACACAGTTTTCTCTGAAGATCAACAGCCTGCAGCCTGAAGATTTTGGGAGTTATTACTGTCAACATCATTATGGTACTCCTCC |
| IGKV8-27*01_BALB | BALB/cByJ | Musmus IGKV8-27*01 F | 100 | 300 | AACATTATGATGACACAGTCGCCATCATCTCTGGCTGTGTCTGCAGGAGAAAAGGTCACTATGAGCTGTAAGTCCAGTCAAAGTGTTTTATACAGTTCAAATCAGAAGAACTACTTGGCCTGGTACCAGCAGAAACCAGGGCAGTCTCCTAAACTGCTGATCTACTGGGCATCCACTAGGGAATCTGGTGTCCCTGATCGCTTCACAGGCAGTGGATCTGGGACAGATTTTACTCTTACCATCAGCAGTGTACAAGCTGAAGACCTGGCAGTTTATTACTGTCATCAATACCTCTCCTCG |
| IGKV6-23*01_BALB | BALB/cByJ | Musmus IGKV6-23*01 F | 100 | 287 | GACATTGTGATGACCCAGTCTCACAAATTCATGTCCACATCAGTAGGAGACAGGGTCAGCATCACCTGCAAGGCCAGTCAGGATGTGGGTACTGCTGTAGCCTGGTATCAACAGAAACCAGGGCAATCTCCTAAACTACTGATTTACTGGGCATCCACCCGGCACACTGGAGTCCCTGATCGCTTCACAGGCAGTGGATCTGGGACAGATTTCACTCTCACCATTAGCAATGTGCAGTCTGAAGACTTGGCAGATTATTTCTGTCAGCAATATAGCAGCTATCCTCT |
| IGKV4-59*01_BALB | BALB/cByJ | Musmus IGKV4-59*01 F | 100 | 286 | CAAATTGTTCTCACCCAGTCTCCAGCAATCATGTCTGCATCTCCAGGGGAGAAGGTCACCATGACCTGCAGTGCCAGCTCAAGTGTAAGTTACATGCACTGGTACCAGCAGAAGTCAGGCACCTCCCCCAAAAGATGGATTTATGACACATCCAAACTGGCTTCTGGAGTCCCTGCTCGCTTCAGTGGCAGTGGGTCTGGGACCTCTTACTCTCTCACAATCAGCAGCATGGAGGCTGAAGATGCTGCCACTTATTACTGCCAGCAGTGGAGTAGTAACCCACCCA |
| IGKV14-126*01_BALB | BALB/cByJ | Musmus IGKV14-126*01 F | 100 | 287 | GACATCAAGATGACCCAGTCTCCATCCTCCATGTATGCATCGCTGGGAGAGAGAGTCACTATCACTTGCAAGGCGAGTCAGGACATTAAAAGCTATTTAAGCTGGTACCAGCAGAAACCATGGAAATCTCCTAAGACCCTGATCTATTATGCAACAAGCTTGGCAGATGGGGTCCCATCAAGATTCAGTGGCAGTGGATCTGGGCAAGATTATTCTCTAACCATCAGCAGCCTGGAGTCTGACGATACAGCAACTTATTACTGTCTACAGCATGGTGAGAGCCCTCC |
| IGKV8-21*01_BALB | BALB/cByJ | Musmus IGKV8-21*01 F | 100 | 302 | GACATTGTGATGTCACAGTCTCCATCCTCCCTGGCTGTGTCAGCAGGAGAGAAGGTCACTATGAGCTGCAAATCCAGTCAGAGTCTGCTCAACAGTAGAACCCGAAAGAACTACTTGGCTTGGTACCAGCAGAAACCAGGGCAGTCTCCTAAACTGCTGATCTACTGGGCATCCACTAGGGAATCTGGGGTCCCTGATCGCTTCACAGGCAGTGGATCTGGGACAGATTTCACTCTCACCATCAGCAGTGTGCAGGCTGAAGACCTGGCAGTTTATTACTGCAAGCAATCTTATAATCTTCC |

| | | | | | |
|---|---|---|---|---|---|
| IGKV6-13*01_BALB | BALB/cByJ | Musmus IGKV6-13*01 F | 100 | 285 | GACATTGTGATGACCCAGTCTCAAAAATTCATGTCCACATCAGTAGGAGACAGGGTCAGCATCACCTGCAAGGCCAGTCAGAATGTGGGTACTGCTGTAGCCTGGTATCAACAGAAACCAGGACAATCTCCTAAACTACTGATTTACTCGGCATCCAATCGGTACACTGGAGTCCCTGATCGCTTCACAGGCAGTGGATCTGGGACAGATTTCACTCTCACCATCAGCAATATGCAGTCTGAAGACCTGGCAGATTATTTCTGCCAGCAATATAGCAGCTATCCT |
| IGKV1-135*01_BALB | BALB/cByJ | Musmus IGKV1-135*01 F | 100 | 302 | GATGTTGTGATGACCCAGACTCCACTCACTTTGTCGGTTACCATTGGACAACCAGCCTCCATCTCTTGCAAGTCAAGTCAGAGCCTCTTAGATAGTGATGGAAAGACATATTTGAATTGGTTGTTACAGAGGCCAGGCCAGTCTCCAAAGCGCCTAATCTATCTGGTGTCTAAACTGGACTCTGGAGTCCCTGACAGGTTCACTGGCAGTGGATCAGGGACAGATTTCACACTGAAAATCAGCAGAGTGGAGGCTGAGGATTTGGGAGTTTATTATTGCTGGCAAGGTACACATTTTCCTCA |
| IGKV4-70*01_BALB | BALB/cByJ | Musmus IGKV4-70*01 F | 100 | 282 | CAAATTGTTCTCACCCAGTCTCCAGCAATCATGTCTGCATCTCCAGGGGAGAAGGTCACCATGACCTGCAGTGCCAGCTCAAGTATAAGTTACATGCACTGGTACCAGCAGAAGCCAGGCACCTCCCCCAAAAGATGGATTTATGACACATCCAAACTGGCTTCTGGAGTCCCTGCTCGCTTCAGTGGCAGTGGGTCTGGGACCTCTTATTCTCTCACAATCAGCAGCATGGAGGCTGAAGATGCTGCCACTTATTACTGCCATCAGCGGAGTAGTTACCCA |
| IGKV15-103*01_BALB | BALB/cByJ | Musmus IGKV15-103*01 ORF | 100 | 287 | GACATCCAGATGAACCAGTCTCCATCCAGTCTGTCTGCATCCCTTGGAGACACAATTACCATCACTTGCCATGCCAGTCAGAACATTAATGTTTGGTTAAGCTGGTACCAGCAGAAACCAGGAAATATTCCTAAACTATTGATCTATAAGGCTTCCAACTTGCACACAGGCGTCCCATCAAGGTTTAGTGGCAGTGGATCTGGAACAGGTTTCACATTAACCATCAGCAGCCTGCAGCCTGAAGACATTGCCACTTACTACTGTCAACAGGGTCAAAGTTATCCTCT |
| IGKV8-19*01_BALB | BALB/cByJ | Musmus IGKV8-19*01 F | 100 | 305 | GACATTGTGATGACACAGTCTCCATCCTCCCTGACTGTGACAGCAGGAGAGAAGGTCACTATGAGCTGCAAGTCCAGTCAGAGTCTGTTAAACAGTGGAAATCAAAAGAACTACTTGACCTGGTACCAGCAGAAACCAGGGCAGCCTCCTAAACTGTTGATCTACTGGGCATCCACTAGGGAATCTGGGGTCCCTGATCGCTTCACAGGCAGTGGATCTGGAACAGATTTCACTCTCACCATCAGCAGTGTGCAGGCTGAAGACCTGGCAGTTTATTACTGTCAGAATGATTATAGTTATCCTCC |
| IGKV10-96*01_BALB | BALB/cByJ | Musmus IGKV10-96*01 F | 100 | 286 | GATATCCAGATGACACAGACTACATCCTCCCTGTCTGCCTCTCTGGGAGACAGAGTCACCATCAGTTGCAGGGCAAGTCAGGACATTAGCAATTATTTAAACTGGTATCAGCAGAAACCAGATGGAACTGTTAAACTCCTGATCTACTACACATCAAGATTACACTCAGGAGTCCCATCAAGGTTCAGTGGCAGTGGGTCTGGAACAGATTATTCTCTCACCATTAGCAACCTGGAGCAAGAAGATATTGCCACTTACTTTTGCCAACAGGGTAATACGCTTCCTC |
| IGKV12-41*01_BALB | BALB/cByJ | Musmus IGKV12-41*01 F | 100 | 287 | GACATCCAGATGACTCAGTCTCCAGCCTCCCTATCTGCATCTGTGGGAGAAACTGTCACCATCACATGTCGAGCAAGTGGGAATATTCACAATTATTTAGCATGGTATCAGCAGAAACAGGGAAAATCTCCTCAGCTCCTGGTCTATAATGCAAAAACCTTAGCAGATGGTGTGCCATCAAGGTTCAGTGGCAGTGGATCAGGAACACAATATTCTCTCAAGATCAACAGCCTGCAGCCTGAAGATTTTGGGAGTTATTACTGTCAACATTTTTGGAGTACTCCTCC |
| IGKV5-43*01_BALB | BALB/cByJ | Musmus IGKV5-43*01 F | 100 | 287 | GATATTGTGCTAACTCAGTCTCCAGCCACCCTGTCTGTGACTCCAGGAGATAGCGTCAGTCTTTCCTGCAGGGCCAGCCAAAGTATTAGCAACAACCTACACTGGTATCAACAAAAATCACATGAGTCTCCAAGGCTTCTCATCAAGTATGCTTCCCAGTCCATCTCTGGGATCCCCTCCAGGTTCAGTGGCAGTGGATCAGGGACAGATTTCACTCTCAGTATCAACAGTGTGGAGACTGAAGATTTTGGAATGTATTTCTGTCAACAGAGTAACAGCTGGCCTCA |

| | | | | | |
|---|---|---|---|---|---|
| IGKV1-117*01_BALB | BALB/cByJ | Musmus IGKV1-117*01 F | 100 | 302 | GATGTTTTGATGACCCAAACTCCACTCTCCCTGCCTGTCAGTCTTGGAGATCAAGCCTCCATCTCTTGCAGATCTAGTCAGAGCATTGTACATAGTAATGGAAACACCTATTTAGAATGGTACCTGCAGAAACCAGGCCAGTCTCCAAAGCTCCTGATCTACAAAGTTTCCAACCGATTTTCTGGGGTCCCAGACAGGTTCAGTGGCAGTGGATCAGGGACAGATTTCACACTCAAGATCAGCAGAGTGGAGGCTGAGGATCTGGGAGTTTATTACTGCTTTCAAGGTTCACATGTTCCTCC |
| IGKV1-110*01_BALB | BALB/cByJ | Musmus IGKV1-110*01 F | 100 | 302 | GATGTTGTGATGACCCAAACTCCACTCTCCCTGCCTGTCAGTCTTGGAGATCAAGCCTCCATCTCTTGCAGATCTAGTCAGAGCCTTGTACACAGTAATGGAAACACCTATTTACATTGGTACCTGCAGAAGCCAGGCCAGTCTCCAAAGCTCCTGATCTACAAAGTTTCCAACCGATTTTCTGGGGTCCCAGACAGGTTCAGTGGCAGTGGATCAGGGACAGATTTCACACTCAAGATCAGCAGAGTGGAGGCTGAGGATCTGGGAGTTTATTTCTGCTCTCAAAGTACACATGTTCCTCC |
| IGKV8-16*01_BALB | BALB/cByJ | Musmus IGKV8-16*01 F | 100 | 305 | GAAATTGTGTTGACCCAGTCTATACCATCCCTGACTGTGTCAGCAGGAGAGAGGGTCACTATCAGCTGCAAATCCAATCAGAATCTTTTATGGAGTGGAAACCAAAGGTACTGTTTGGTCTGGCACCAGTGGAAACCAGGGCAAACTCCTACACCGTTGATCACCTGGACATCTGATAGGTACTCTGGAGTCCCTGATCGTTTCATAGGCAGTGGATCTGTGACAGATTTCACTCTGACCATCAGCAGTGTGCAGGCTGAAGATGTGGCAGTTTATTTCTGTCAGCAGCATTTACACATTCCTCC |
| IGKV4-57-1*01_BALB | BALB/cByJ | Musmus IGKV4-57*01 F | 100 | 292 | GAAAATGTGCTCACCCAGTCTCCAGCAATCATGTCTGCATCTCCAGGGGAAAAGGTCACCATGACCTGCAGGGCCAGCTCAAGTGTAAGTTCCAGTTACTTGCACTGGTACCAGCAGAAGTCAGGTGCCTCCCCCAAACTCTGGATTTATAGCACATCCAACTTGGCTTCTGGAGTCCCTGCTCGCTTCAGTGGCAGTGGGTCTGGGACCTCTTACTCTCTCACAATCAGCAGTGTGGAGGCTGAAGATGCTGCCACTTATTACTGCCAGCAGTACAGTGGTTACCCACTCA |
| IGKV4-55*01_BALB | BALB/cByJ | Musmus IGKV4-55*01 F | 100 | 286 | CAAATTGTTCTCACCCAGTCTCCAGCAATCATGTCTGCATCTCCAGGGGAGAAGGTCACCATGACCTGCAGTGCCAGCTCAAGTGTAAGTTACATGTACTGGTACCAGCAGAAGCCAGGATCCTCCCCCAGACTCCTGATTTATGACACATCCAACCTGGCTTCTGGAGTCCCTGTTCGCTTCAGTGGCAGTGGGTCTGGGACCTCTTACTCTCTCACAATCAGCCGAATGGAGGCTGAAGATGCTGCCACTTATTACTGCCAGCAGTGGAGTAGTTACCCACCCA |
| IGKV6-32*01_BALB | BALB/cByJ | Musmus IGKV6-32*01 F | 100 | 287 | AGTATTGTGATGACCCAGACTCCCAAATTCCTGCTTGTATCAGCAGGAGACAGGGTTACCATAACCTGCAAGGCCAGTCAGAGTGTGAGTAATGATGTAGCTTGGTACCAACAGAAGCCAGGGCAGTCTCCTAAACTGCTGATATACTATGCATCCAATCGCTACACTGGAGTCCCTGATCGCTTCACTGGCAGTGGATATGGGACGGATTTCACTTTCACCATCAGCACTGTGCAGGCTGAAGACCTGGCAGTTTATTTCTGTCAGCAGGATTATAGCTCTCCTCC |
| IGKV4-57*01_S9219_BALB | BALB/cByJ | Musmus IGKV4-57-1*01 F | 100 | 286 | CAAATTGTTCTCACCCAGTCTCCAGCAATCATGTCTGCATCTCCAGGGGAGAAGGTCACCATAACCTGCAGTGCCAGCTCAAGTGTAAGTTACATGCACTGGTTCCAGCAGAAGCCAGGCACTTCTCCCAAACTCTGGATTTATAGCACATCCAACCTGGCTTCTGGAGTCCCTGCTCGCTTCAGTGGCAGTGGATCTGGGACCTCTTACTCTCTCACAATCAGCCGAATGGAGGCTGAAGATGCTGCCACTTATTACTGCCAGCAAAGGAGTAGTTACCCACTCA |
| IGKV12-46*01_BALB | BALB/cByJ | Musmus IGKV12-46*01 F | 100 | 287 | GACATCCAGATGACTCAGTCTCCAGCCTCCCTATCTGTATCTGTGGGAGAAACTGTCACCATCACATGTCGAGCAAGTGAGAATATTTACAGTAATTTAGCATGGTATCAGCAGAAACAGGGAAAATCTCCTCAGCTCCTGGTCTATGCTGCAACAAACTTAGCAGATGGTGTGCCATCAAGGTTCAGTGGCAGTGGATCAGGCACACAGTATTCCCTCAAGATCAACAGCCTGCAGTCTGAAGATTTTGGGAGTTATTACTGTCAACATTTTGGGGTACTCCTCC |

| | | | | | |
|---|---|---|---|---|---|
| IGKV2-137*01_BALB | BALB/cByJ | Musmus IGKV2-137*01 F | 100 | 302 | GATATTGTGATGACTCAGGCTGCACCCTCTGTACCTGTCACTCCTGGAGAGTCAGTATCCATCTCCTGCAGGTCTAGTAAGAGTCTCCTGCATAGTAATGGCAACACTTACTTGTATTGGTTCCTGCAGAGGCCAGGCCAGTCTCCTCAGCTCCTGATATATCGGATGTCCAACCTTGCCTCAGGAGTCCCAGACAGGTTCAGTGGCAGTGGGTCAGGAACTGCTTTCACACTGAGAATCAGTAGAGTGGAGGCTGAGGATGTGGGTGTTTATTACTGTATGCAACATCTAGAATATCCTTT |
| IGKV8-28*01_BALB | BALB/cByJ | Musmus IGKV8-28*01 F | 100 | 305 | GACATTGTGATGACACAGTCTCCATCCTCCCTGAGTGTGTCAGCAGGAGAGAAGGTCACTATGAGCTGCAAGTCCAGTCAGAGTCTGTTAAACAGTGGAAATCAAAAGAACTACTTGGCCTGGTACCAGCAGAAACCAGGGCAGCCTCCTAAACTGTTGATCTACGGGGCATCCACTAGGGAATCTGGGGTCCCTGATCGCTTCACAGGCAGTGGATCTGGAACCGATTTCACTCTTACCATCAGCAGTGTGCAGGCTGAAGACCTGGCAGTTTATTACTGTCAGAATGATCATAGTTATCCTCC |
| IGKV4-74*01_BALB | BALB/cByJ | Musmus IGKV4-74*01 F | 100 | 292 | CAAATTGTTCTCACCCAGTCTCCAGCAATCATGTCTGCATCTCTAGGGGAACGGGTCACCATGACCTGCACTGCCAGCTCAAGTGTAAGTTCCAGTTACTTGCACTGGTACCAGCAGAAGCCAGGATCCTCCCCCAAACTCTGGATTTATAGCACATCCAACCTGGCTTCTGGAGTCCCAGCTCGCTTCAGTGGCAGTGGGTCTGGGACCTCTTACTCTCTCACAATCAGCAGCATGGAGGCTGAAGATGCTGCCACTTATTACTGCCACCAGTATCATCGTTCCCCACCCA |
| IGKV6-20*01_BALB | BALB/cByJ | Musmus IGKV6-20*01 F | 100 | 287 | AACATTGTAATGACCCAATCTCCCAAATCCATGTCCATGTCAGTAGGAGAGAGGGTCACCTTGAGCTGCAAGGCCAGTGAGAATGTGGGTACTTATGTATCCTGGTATCAACAGAAACCAGAGCAGTCTCCTAAACTGCTGATATACGGGGCATCCAACCGGTACACTGGGGTCCCCGATCGCTTCACAGGCAGTGGATCTGCAACAGATTTCACTCTGACCATCAGCAGTGTGCAGGCTGAAGACCTTGCAGATTATCACTGTGGACAGAGTTACAGCTATCCTCC |
| IGKV8-24*01_BALB | BALB/cByJ | Musmus IGKV8-24*01 F | 100 | 305 | GACATTGTGATGACACAGTCTCCATCCTCCCTGGCTATGTCAGTAGGACAGAAGGTCACTATGAGCTGCAAGTCCAGTCAGAGCCTTTTAAATAGTAGCAATCAAAAGAACTATTTGGCCTGGTACCAGCAGAAACCAGGACAGTCTCCTAAACTTCTGGTATACTTTGCATCCACTAGGGAATCTGGGGTCCCTGATCGCTTCATAGGCAGTGGATCTGGGACAGATTTCACTCTTACCATCAGCAGTGTGCAGGCTGAAGACCTGGCAGATTACTTCTGTCAGCAACATTATAGCACTCCTCC |
| IGKV12-89*01_BALB | BALB/cByJ | Musmus IGKV12-89*01 F | 100 | 287 | GACATCCAGATGACTCAGTCTCCAGCTTCACTGTCTGCATCTGTGGGAGAAACTGTCACCATCACATGTGGAGCAAGTGAGAATATTTACGGTGCTTTAAATTGGTATCAGCGGAAACAGGGAAAATCTCCTCAGCTCCTGATCTATGGTGCAACCAACTTGGCAGATGGCATGTCATCGAGGTTCAGTGGCAGTGGATCTGGTAGACAGTATTCTCTCAAGATCAGTAGCCTGCATCCTGACGATGTTGCAACGTATTACTGTCAAAATGTGTTAAGTACTCCTCC |
| IGKV4-53*01_BALB | BALB/cByJ | Musmus IGKV4-53*01 F | 100 | 292 | GAAATTGTGCTCACCCAGTCTCCAGCACTCATGGCTGCATCTCCAGGGGAGAAGGTCACCATCACCTGCAGTGTCAGCTCAAGTATAAGTTCCAGCAACTTGCACTGGTACCAGCAGAAGTCAGAAACCTCCCCCAAACCCTGGATTTATGGCACATCCAACCTGGCTTCTGGAGTCCCTGTTCGCTTCAGTGGCAGTGGATCTGGGACCTCTTATTCTCTCACAATCAGCAGCATGGAGGCTGAAGATGCTGCCACTTATTACTGTCAACAGTGGAGTAGTTACCCACTCA |

| | | | | | |
|---|---|---|---|---|---|
| IGKV4-68*01_BALB | BALB/cByJ | Musmus IGKV4-68*01 F | 100 | 286 | CAAATTGTTCTCACCCAGTCTCCAGCACTCATGTCTGCATCTCCAGGGGAGAAGGTCACCATGACCTGCAGTGCCAGCTCAAGTGTAAGTTACATGTACTGGTACCAGCAGAAGCCAAGATCCTCCCCCAAACCCTGGATTTATCTCACATCCAACCTGGCTTCTGGAGTCCCTGCTCGCTTCAGTGGCAGTGGGTCTGGGACCTCTTACTCTCTCACAATCAGCAGCATGGAGGCTGAAGATGCTGCCACTTATTACTGCCAGCAGTGGAGTAGTAACCCACCCA |
| IGKV19-93*01_BALB | BALB/cByJ | Musmus IGKV19-93*01 F | 100 | 287 | GACATCCAGATGACACAGTCTCCATCCTCACTGTCTGCATCTCTGGGAGGCAAAGTCACCATCACTTGCAAGGCAAGCCAAGACATTAACAAGTATATAGCTTGGTACCAACACAAGCCTGGAAAAGGTCCTAGGCTGCTCATACATTACACATCTACATTACAGCCAGGCATCCCATCAAGGTTCAGTGGAAGTGGGTCTGGGAGAGATTATTCCTTCAGCATCAGCAACCTGGAGCCTGAAGATATTGCAACTTATTATTGTCTACAGTATGATAATCTTCTACC |
| IGKV6-17*01_BALB | BALB/cByJ | Musmus IGKV6-17*01 F | 100 | 287 | GACATTGTGATGACCCAGTCTCACAAATTCATGTCCACATCAGTAGGAGACAGGGTCAGCATCACCTGCAAGGCCAGTCAGGATGTGAGTACTGCTGTAGCCTGGTATCAACAGAAACCAGGACAATCTCCTAAACTACTGATTTACTCGGCATCCTACCGGTACACTGGAGTCCCTGATCGCTTCACTGGCAGTGGATCTGGGACGGATTTCACTTTCACCATCAGCAGTGTGCAGGCTGAAGACCTGGCAGTTTATTACTGTCAGCAACATTATAGTACTCCTCC |
| IGKV4-80*01_BALB | BALB/cByJ | Musmus IGKV4-80*01 F | 100 | 282 | CAAATTGTTCTCACCCAGTCTCCAGCAATCATGTCTGCATCTCTAGGGGAGGAGATCACCCTAACCTGCAGTGCCAGCTCGAGTGTAAGTTACATGCACTGGTACCAGCAGAAGTCAGGCACTTCTCCCAAACTCTTGATTTATAGCACATCCAACCTGGCTTCTGGAGTCCCTTCTCGCTTCAGTGGCAGTGGGTCTGGGACCTTTTATTCTCTCACAATCAGCAGTGTGGAGGCTGAAGATGCTGCCGATTATTACTGCCATCAGTGGAGTAGTTATCCA |
| IGKV3-2*01_BALB | BALB/cByJ | Musmus IGKV3-2*01 F | 100 | 299 | GACATTGTGCTGACCCAATCTCCAGCTTCTTTGGCTGTGTCTCTAGGGCAGAGGGCCACCATCTCCTGCAGAGCCAGCGAAAGTGTTGATAATTATGGCATTAGTTTTATGAACTGGTTCCAACAGAAACCAGGACAGCCACCCAAACTCCTCATCTATGCTGCATCCAACCAAGGATCCGGGGTCCCTGCCAGGTTTAGTGGCAGTGGGTCTGGGACAGACTTCAGCCTCAACATCCATCCTATGGAGGAGGATGATACTGCAATGTATTTCTGTCAGCAAAGTAAGGAGGTTCCTCC |
| IGKV13-84*01_BALB | BALB/cByJ | Musmus IGKV13-84*01 F | 100 | 287 | GACATCCAGATGACACAATCTTCATCCTCCTTTTCTGTATCTCTAGGAGACAGAGTCACCATTACTTGCAAGGCAAGTGAGGACATATATAATCGGTTAGCCTGGTATCAGCAGAAACCAGGAAATGCTCCTAGGCTCTTAATATCTGGTGCAACCAGTTTGGAAACTGGGGTTCCTTCAAGATTCAGTGGCAGTGGATCTGGAAAGGATTACACTCTCAGCATTACCAGTCTTCAGACTGAAGATGTTGCTACTTATTACTGTCAACAGTATTGGAGTACTCCTCC |
| IGKV4-72*01_BALB | BALB/cByJ | Musmus IGKV4-72*01 F | 100 | 286 | CAAATTGTTCTCTCCCAGTCTCCAGCAATCCTGTCTGCATCTCCAGGGGAGAAGGTCACAATGACTTGCAGGGCCAGCTCAAGTGTAAGTTACATGCACTGGTACCAGCAGAAGCCAGGATCCTCCCCCAAACCCTGGATTTATGCCACATCCAACCTGGCTTCTGGAGTCCCTGCTCGCTTCAGTGGCAGTGGGTCTGGGACCTCTTACTCTCTCACAATCAGCAGAGTGGAGGCTGAAGATGCTGCCACTTATTACTGCCAGCAGTGGAGTAGTAACCCACCCA |
| IGKV9-120*01_S2628_BALB | BALB/cByJ | Musmus IGKV9-120*02 F | 100 | 285 | GACATCCAGATGACCCAGTCTCCATCCTCCTTATCTGCCTCTCTGGGAGAAAGAGTCAGTCTCACTTGTCGGGCAAGTCAGGACATTGGTAGTAGCTTAAACTGGCTTCAGCAGGAACCAGATGGAACTATTAAACGCCTGATCTACGCCACATCCAGTTTAGATTCTGGTGTCCCCAAAAGGTTCAGTGGCAGTAGGTCTGGGTCAGATTATTCTCTCACCATCAGCAGCCTTGAGTCTGAAGATTTTGTAGACTATTACTGTCTACAATATGCTAGTTCTCCT |

| | | | | | |
|---|---|---|---|---|---|
| IGKV4-50*01_BALB | BALB/cByJ | Musmus IGKV4-50*01 F | 100 | 286 | GAAAATGTGCTCACCCAGTCTCCAGCAATCATGTCTGCATCTCTAGGGGAGAAGGTCACCATGAGCTGCAGGGCCAGCTCAAGTGTAAATTACATGTACTGGTACCAGCAGAAGTCAGATGCCTCCCCCAAACTATGGATTTATTACACATCCAACCTGGCTCCTGGAGTCCCAGCTCGCTTCAGTGGCAGTGGGTCTGGGAACTCTTATTCTCTCACAATCAGCAGCATGGAGGGTGAAGATGCTGCCACTTATTACTGCCAGCAGTTTACTAGTTCCCCATCCA |
| IGKV6-25*01_BALB | BALB/cByJ | Musmus IGKV6-25*01 F | 100 | 287 | GACATTGTGATGACCCAGTCTCACAAATTCATGTCCACATCAGTAGGAGACAGGGTCAGCATCACCTGCAAGGCCAGTCAGGATGTGAGTACTGCTGTAGCCTGGTATCAACAAAAACCAGGGCAATCTCCTAAACTACTGATTTACTGGGCATCCACCCGGCACACTGGAGTCCCTGATCGCTTCACAGGCAGTGGATCTGGGACAGATTATACTCTCACCATCAGCAGTGTGCAGGCTGAAGACCTGGCACTTTATTACTGTCAGCAACATTATAGCACTCCTCC |
| IGKV4-79*01_BALB | BALB/cByJ | Musmus IGKV4-79*01 F | 100 | 292 | CAAATTGTTCTCACCCAGTCTCCAGCAATCATGTCTGCATCTCCTGGGGAGAAGGTCACCTTGACCTGCAGTGCCAGCTCAAGTGTAAGTTCCAGCTACTTGTACTGGTACCAGCAGAAGCCAGGATCCTCCCCCAAACTCTGGATTTATAGCACATCCAACCTGGCTTCTGGAGTCCCTGCTCGCTTCAGTGGCAGTGGGTCTGGGACCTCTTACTCTCTCACAATCAGCAGCATGGAGGCTGAAGATGCTGCCTCTTATTTCTGCCATCAGTGGAGTAGTTACCCACCCC |
| IGKV3-12*01_BALB | BALB/cByJ | Musmus IGKV3-12*01 F | 100 | 299 | GACATTGTGCTGACACAGTCTCCTGCTTCCTTAGCTGTATCTCTGGGGCAGAGGGCCACCATCTCATGCAGGGCCAGCAAAAGTGTCAGTACATCTGGCTATAGTTATATGCACTGGTACCAACAGAAACCAGGACAGCCACCCAAACTCCTCATCTATCTTGCATCCAACCTAGAATCTGGGGTCCCTGCCAGGTTCAGTGGCAGTGGGTCTGGGACAGACTTCACCCTCAACATCCATCCTGTGGAGGAGGAGGATGCTGCAACCTATTACTGTCAGCACAGTAGGGAGCTTCCTCC |
| IGKV16-104*01_BALB | BALB/cByJ | Musmus IGKV16-104*01 F | 100 | 288 | GATGTCCAGATAACCCAGTCTCCATCTTATCTTGCTGCATCTCCTGGAGAAACCATTACTATTAATTGCAGGGCAAGTAAGAGCATTAGCAAATATTTAGCCTGGTATCAAGAGAAACCTGGGAAAACTAATAAGCTTCTTATCTACTCTGGATCCACTTTGCAATCTGGAATTCCATCAAGGTTCAGTGGCAGTGGATCTGGTACAGATTTCACTCTCACCATCAGTAGCCTGGAGCCTGAAGATTTTGCAATGTATTACTGTCAACAGCATAATGAATACCCGTAC |
| IGKV1-88*01_BALB | BALB/cByJ | Musmus IGKV1-88*01 F | 100 | 302 | GATGTTGTGGTGACTCAAACTCCACTCTCCCTGCCTGTCAGCTTTGGAGATCAAGTTTCTATCTCTTGCAGGTCTAGTCAGAGTCTTGCAAACAGTTATGGGAACACCTATTTGTCTTGGTACCTGCACAAGCCTGGCCAGTCTCCACAGCTCCTCATCTATGGGATTTCCAACAGATTTTCTGGGGTGCCAGACAGGTTCAGTGGCAGTGGTTCAGGGACAGATTTCACACTCAAGATCAGCACAATAAAGCCTGAGGACTTGGGAATGTATTACTGCTTACAAGGTACACATCAGCCTCC |
| IGKV3-7*01_S5296_BALB | BALB/cByJ | Musmus IGKV3-7*02 F | 100 | 299 | GACATTGTGCTGACACAGTCTCCTGCTTCCTTAGCTGTATCTCTGGGGCAGAGGGCCACCATCTCATGCAGGGCCAGCCAAAGTGTCAGTACATCTAGCTATAGTTATATGCACTGGTACCAACAGAAACCAGGACAGCCACCCAAACTCCTCATCAAGTATGCATCCAACCTAGAATCTGGGGTCCCTGCCAGGTTCAGTGGCAGTGGGTCTGGGACAGACTTCACCCTCAACATCCATCCTGTGGAGGAGGAGGATACTGCAACATATTACTGTCAGCACAGTTGGGAGATTCCTCC |
| IGKV14-100*01_BALB | BALB/cByJ | Musmus IGKV14-100*01 F | 100 | 287 | GACATCCTGATGACCCAATCTCCATCCTCCATGTCTGTATCTCTGGGAGACACAGTCAGCATCACTTGCCATGCAAGTCAGGGCATTAGCAGTAATATAGGGTGGTTGCAGCAGAAACCAGGGAAATCATTTAAGGGCCTGATCTATCATGGAACCAACTTGGAAGATGGAGTTCCATCAAGGTTCAGTGGCAGTGGATCTGGAGCAGATTATTCTCTCACCATCAGCAGCCTGGAATCTGAAGATTTTGCAGACTATTACTGTGTACAGTATGCTCAGTTTCCTCC |

| | | | | | |
|---|---|---|---|---|---|
| IGKV10-94*01_BALB | BALB/cByJ | Musmus IGKV10-94*01 F | 100 | 287 | GATATCCAGATGACACAGACTACATCCTCCCTGTCTGCCTCTCTGGGAGACAGAGTCACCATCAGTTGCAGTGCAAGTCAGGGCATTAGCAATTATTTAAACTGGTATCAGCAGAAACCAGATGGAACTGTTAAACTCCTGATCTATTACACATCAAGTTTACACTCAGGAGTCCCATCAAGGTTCAGTGGCAGTGGGTCTGGGACAGATTATTCTCTCACCATCAGCAACCTGGAACCTGAAGATATTGCCACTTACTATTGTCAGCAGTATAGTAAGCTTCCTCC |
| IGKV4-91*01_S5997_BALB | BALB/cByJ | Musmus IGKV4-91*01 F | 100 | 292 | GAAATTGTGCTCACCCAGTCTCCAACCACCATGGCTGCATCTCCCGGGGAGAAGATCACTATCACCTGCAGTGCCAGCTCAAGTATAAGTTCCAATTACTTGCATTGGTATCAGCAGAAGCCAGGATTCTCCCCTAAACTCTTGATTTATAGGACATCCAATCTGGCTTCTGGAGTCCCAGCTCGCTTCAGTGGCAGTGGGTCTGGGACCTCTTACTCTCTCACAATTGGCACCATGGAGGCTGAAGATGTTGCCACTTACTACTGCCAGCAGGGTAGTAGTATACCACTCA |
| IGKV3-4*01_BALB | BALB/cByJ | Musmus IGKV3-4*01 F | 100 | 299 | GACATTGTGCTGACCCAATCTCCAGCTTCTTTGGCTGTGTCTCTAGGGCAGAGGGCCACCATCTCCTGCAAGGCCAGCCAAAGTGTTGATTATGATGGTGATAGTTATATGAACTGGTACCAACAGAAACCAGGACAGCCACCCAAACTCCTCATCTATGCTGCATCCAATCTAGAATCTGGGATCCCAGCCAGGTTTAGTGGCAGTGGGTCTGGGACAGACTTCACCCTCAACATCCATCCTGTGGAGGAGGAGGATGCTGCAACCTATTACTGTCAGCAAAGTAATGAGGATCCTCC |
| IGKV4-61*01_BALB | BALB/cByJ | Musmus IGKV4-61*01 F | 100 | 286 | CAAATTGTTCTCACCCAGTCTCCAGCAATCATGTCTGCATCTCCAGGGGAGAAGGTCACCATATCCTGCAGTGCCAGCTCAAGTGTAAGTTACATGTACTGGTACCAGCAGAAGCCAGGATCCTCCCCCAAACCCTGGATTTATCGCACATCCAACCTGGCTTCTGGAGTCCCTGCTCGCTTCAGTGGCAGTGGGTCTGGGACCTCTTACTCTCTCACAATCAGCAGCATGGAGGCTGAAGATGCTGCCACTTATTACTGCCAGCAGTATCATAGTTACCCACCCA |
| IGKV5-48*01_BALB | BALB/cByJ | Musmus IGKV5-48*01 F | 100 | 287 | GACATCTTGCTGACTCAGTCTCCAGCCATCCTGTCTGTGAGTCCAGGAGAAAGAGTCAGTTTCTCCTGCAGGGCCAGTCAGAGCATTGGCACAAGCATACACTGGTATCAGCAAAGAACAAATGGTTCTCCAAGGCTTCTCATAAAGTATGCTTCTGAGTCTATCTCTGGGATCCCTTCCAGGTTTAGTGGCAGTGGATCAGGGACAGATTTTACTCTTAGCATCAACAGTGTGGAGTCTGAAGATATTGCAGATTATTACTGTCAACAAAGTAATAGCTGGCCAAC |
| IGKV17-127*01_BALB | BALB/cByJ | Musmus IGKV17-127*01 F | 100 | 287 | GAAACAACTGTGACCCAGTCTCCAGCATCCCTGTCCGTGGCTACAGGAGAAAAAGTCACTATCAGATGCATAACCAGCACTGATATTGATGATGATATGAACTGGTACCAGCAGAAGCCAGGGGAACCTCCTAAGCTCCTTATTTCAGAAGGCAATACTCTTCGTCCTGGAGTCCCATCCCGATTCTCCAGCAGTGGCTATGGCACAGATTTTGTTTTTACAATTGAAAACACGCTCTCAGAAGATGTTGCAGATTACTACTGTTTGCAAAGTGATAACATGCCTCT |
| IGKV1-122*01_BALB | BALB/cByJ | Musmus IGKV1-122*01 F | 100 | 302 | GATGCTGTGATGACCCAAACTCCACTCTCCCTGCCTGTCAGTCTTGGAGATCAAGCCTCCATCTCTTGCAGGTCTAGTCAGAGCCTTGAAAACAGTAATGGAAACACCTATTTGAACTGGTACCTCCAGAAACCAGGCCAGTCTCCACAGCTCCTGATCTACAGGGTTTCCAACCGATTTTCTGGGGTCCTAGACAGGTTCAGTGGTAGTGGATCAGGGACAGATTTCACACTGAAAATCAGCAGAGTGGAGGCTGAGGATTTGGGAGTTTATTTCTGCCTCCAAGTTACACATGTCCCTCC |
| IGKV5-39*01_BALB | BALB/cByJ | Musmus IGKV5-39*01 F | 100 | 287 | GACATTGTGATGACTCAGTCTCCAGCCACCCTGTCTGTGACTCCAGGAGATAGAGTCTCTCTTTCCTGCAGGGCCAGCCAGAGTATTAGCGACTACTTACACTGGTATCAACAAAAATCACATGAGTCTCCAAGGCTTCTCATCAAATATGCTTCCCAATCCATCTCTGGGATCCCCTCCAGGTTCAGTGGCAGTGGATCAGGGTCAGATTTCACTCTCAGTATCAACAGTGTGGAACCTGAAGATGTTGGAGTGTATTACTGTCAAAATGGTCACAGCTTTCCTCC |

| | | | | | |
|---|---|---|---|---|---|
| IGKV2-109*01_BALB | BALB/cByJ | Musmus IGKV2-109*01 F | 100 | 302 | GATATTGTGATGACGCAGGCTGCATTCTCCAATCCAGTCACTCTTGGAACATCAGCTTCCATCTCCTGCAGGTCTAGTAAGAGTCTCCTACATAGTAATGGCATCACTTATTTGTATTGGTATCTGCAGAAGCCAGGCCAGTCTCCTCAGCTCCTGATTTATCAGATGTCCAACCTTGCCTCAGGAGTCCCAGACAGGTTCAGTAGCAGTGGGTCAGGAACTGATTTCACACTGAGAATCAGCAGAGTGGAGGCTGAGGATGTGGGTGTTTATTACTGTGCTCAAAATCTAGAACTTCCTCC |
| IGKV3-5*01_BALB | BALB/cByJ | Musmus IGKV3-5*01 F | 100 | 299 | GACATTGTGCTGACCCAATCTCCAGCTTCTTTGGCTGTGTCTCTAGGGCAGAGGGCCACCATATCCTGCAGAGCCAGTGAAAGTGTTGATAGTTATGGCAATAGTTTTATGCACTGGTACCAGCAGAAACCAGGACAGCCACCCAAACTCCTCATCTATCGTGCATCCAACCTAGAATCTGGGATCCCTGCCAGGTTCAGTGGCAGTGGGTCTAGGACAGACTTCACCCTCACCATTAATCCTGTGGAGGCTGATGATGTTGCAACCTATTACTGTCAGCAAAGTAATGAGGATCCTCC |
| IGKV2-112*01_BALB | BALB/cByJ | Musmus IGKV2-112*01 F | 100 | 300 | GATATTGTGATAACCCAGGATGAACTCTCCAATCCTGTCACTTCTGGAGAATCAGTTTCCATCTCCTGCAGGTCTAGTAAGAGTCTCCTATATAAGGATGGGAAGACATACTTGAATTGGTTTCTGCAGAGACCAGGACAATCTCCTCAGCTCCTGATCTATTTGATGTCCACCCGTGCATCAGGAGTCTCAGACCGGTTTAGTGGCAGTGGGTCAGGAACAGATTTCACCCTGGAAATCAGTAGAGTGAAGGCTGAGGATGTGGGTGTGTATTACTGTCAACAACTTGTAGAGTATCCT |
| IGKV7-33*01_BALB | BALB/cByJ | Musmus IGKV7-33*01 F | 100 | 305 | GACATTGTGATGACTCAGTCTCCAACTTTCCTTGCTGTGACAGCAAGTAAGAAGGTCACCATTAGTTGCACGGCCAGTGAGAGCCTTTATTCAAGCAAACACAAGGTGCACTACTTGGCTTGGTACCAGAAGAAACCAGAGCAATCTCCTAAACTGCTGATATACGGGGCATCCAACCGATACATTGGGGTCCCTGATCGCTTCACAGGCAGTGGATCTGGGACAGATTTCACTCTGACCATCAGCAGTGTACAGGTTGAAGACCTCACACATTATTACTGTGCACAGTTTTACAGCTATCCTCT |
| IGKV14-111*01_BALB | BALB/cByJ | Musmus IGKV14-111*01 F | 100 | 287 | GACATCAAGATGACCCAGTCTCCATCTTCCATGTATGCATCTCTAGGAGAGAGAGTCACTATCACTTGCAAGGCGAGTCAGGACATTAATAGCTATTTAAGCTGGTTCCAGCAGAAACCAGGGAAATCTCCTAAGACCCTGATCTATCGTGCAAACAGATTGGTAGATGGGGTCCCATCAAGGTTCAGTGGCAGTGGATCTGGGCAAGATTATTCTCTCACCATCAGCAGCCTGGAGTATGAAGATATGGGAATTTATTATTGTCTACAGTATGATGAGTTTCCTCC |
| IGKV4-63*01_BALB | BALB/cByJ | Musmus IGKV4-63*01 F | 100 | 286 | GAAAATGTTCTCACCCAGTCTCCAGCAATCATGTCTGCATCTCCAGGGGAAAAGGTCACCATGACCTGCAGTGCCAGCTCAAGTGTAAGTTACATGCACTGGTACCAGCAGAAGTCAAGCACCTCCCCCAAACTCTGGATTTATGACACATCCAAACTGGCTTCTGGAGTCCCAGGTCGCTTCAGTGGCAGTGGGTCTGGAAACTCTTACTCTCTCACGATCAGCAGCATGGAGGCTGAAGATGTTGCCACTTATTACTGTTTTCAGGGGAGTGGGTACCCACTCA |
| IGKV5-45*01_BALB | BALB/cByJ | Musmus IGKV5-45*01 F | 100 | 287 | GATATTGTGCTAACTCAGTCTCCAGCCACCCTGTCTGTGACTCCAGGAGATAGAGTCAGTCTTTCCTGCAGGGCCAGTCAAAGTATTAGCAACTACCTACACTGGTATCAACAAAAATCACATGAGTCTCCAAGGCTTCTCATCAAGTATGCTTCCCAGTCCATCTCTGGGATCCCCTCCAGGTTCAGTGGCAGTGGATCAGGGACAGATTTCACTCTCAGTATCAACAGTGTGGAGACTGAAGATTTTGGAATGTATTTCTGTCAACAGAGTAACAGCTGGCCTCA |

| | | | | | |
|---|---|---|---|---|---|
| IGKV8-34*01_BALB | BALB/cByJ | Musmus IGKV8-34*01 F | 100 | 305 | GACATTTTGATGACCCAGTCTCCATCCTCCCTGACTGTGTCAGCAGGAGAGAAGGTCA CTATGAGCTGCAAGTCCAGTCAGAGTCTTTTAGCTAGTGGCAACCAAAATAACTACTT GGCCTGGCACCAGCAGAAACCAGGACGATCTCCTAAAATGCTGATAATTTGGGCATCC ACTAGGGTATCTGGAGTCCCTGATCGCTTCATAGGCAGTGGATCTGGGACGGATTTCA CTCTGACCATCAACAGTGTGCAGGCTGAAGATCTGGCTGTTTATTACTGTCAGCAGTC CTACAGCGCTCCTAC |
| IGKV4-86*01_BALB | BALB/cByJ | Musmus IGKV4-86*01 F | 100 | 283 | GAAATTGTGCTCACTCAGTCTCCAGCCATCACAGCTGCATCTCTGGGGCAAAAGGTCA CCATCACCTGCAGTGCCAGCTCAAGTGTAAGTTACATGCACTGGTACCAGCAGAAGTC AGGCACCTCCCCCAAACCATGGATTTATGAAATATCCAAACTGGCTTCTGGAGTCCCA GCTCGCTTCAGTGGCAGTGGGTCTGGGACCTCTTACTCTCTCACAATCAGCAGCATGG AGGCTGAAGATGCTGCCATTTATTACTGCCAGCAGTGGAATTATCCTCTTA |
| IGKV4-69*01_BALB | BALB/cByJ | Musmus IGKV4-69*01 F | 100 | 282 | CAAATTCTTCTCACCCAGTCTCCAGCAATCATGTCTGCATCTCCAGGGGAGAAGGTCA CCATGACCTGCAGTGCCAGCTCAAGTGTAAGTTACATGCACTGGTACCAGCAGAAGCC AGGATCCTCGCCCAAACCCTGGATTTATGACACATCCAACCTGGCTTCTGGATTCCCT GCTCGCTTCAGTGGCAGTGGGTCTGGGACCTCTTACTCTCTCATAATCAGCAGCATGG AGGCTGAAGATGCTGCCACTTATTACTGCCATCAGCGGAGTAGTTACCCA |
| IGKV12-98*01_BALB | BALB/cByJ | Musmus IGKV12-98*01 F | 100 | 287 | GACATTCAGATGACCCAGTCTCCTGCCTCCCAGTCTGCATCTCTGGGAGAAAGTGTCA CCATCACATGCCTGGCAAGTCAGACCATTGGTACATGGTTAGCATGGTATCAGCAGAA ACCAGGGAAATCTCCTCAGCTCCTGATTTATGCTGCAACCAGCTTGGCAGATGGGGTC CCATCAAGGTTCAGTGGTAGTGGATCTGGCACAAAATTTTCTTTCAAGATCAGCAGCC TACAGGCTGAAGATTTTGTAAGTTATTACTGTCAACAACTTTACAGTACTCCTCT |
| IGKV17-121*01_BALB | BALB/cByJ | Musmus IGKV17-121*01 F | 100 | 287 | GAAACAACTGTGACCCAGTCTCCAGCATCCCTGTCCATGGCTATAGGAGAAAAAGTCA CCATCAGATGCATAACCAGCACTGATATTGATGATGATATGAACTGGTACCAGCAGAA GCCAGGGGAACCTCCTAAGCTCCTTATTTCAGAAGGCAATACTCTTCGTCCTGGAGTC CCATCCCGATTCTCCAGCAGTGGCTATGGTACAGATTTTGTTTTTACAATTGAAAACAT GCTCTCAGAAGATGTTGCAGATTACTACTGTTTGCAAAGTGATAACTTGCCTCT |
| IGKV13-85*01_BALB | BALB/cByJ | Musmus IGKV13-85*01 F | 100 | 287 | GACATCCAGATGACACAATCTTCATCCTACTTGTCTGTATCTCTAGGAGGCAGAGTCA CCATTACTTGCAAGGCAAGTGACCACATTAATAATTGGTTAGCCTGGTATCAGCAGAA ACCAGGAAATGCTCCTAGGCTCTTAATATCTGGTGCAACCAGTTTGGAAACTGGGGTT CCTTCAAGATTCAGTGGCAGTGGATCTGGAAAGGATTACACTCTCAGCATTACCAGTC TTCAGACTGAAGATGTTGCTACTTATTACTGTCAACAGTATTGGAGTACTCCTCC |
| IGKV6-29*01_BALB | BALB/cByJ | Musmus IGKV6-29*01 F | 100 | 287 | AACATTGTAATGACCCAATCTCCCAAATCCATGTCCATGTCAGTAGGAGAGAGGGTCA CCTTGAGCTGCAAGGCCAGTGAGAATGTGGGTACTTATGTATCCTGGTATCAACAGAA ACCAGAGCAGTCTCCTAAACTGCTGATATACGGGGCATCCAACCGGTACCCTGGGGTC CCTGATCGCTTCACAGGCAGTGGATCTGCAACAGATTTCACTCTGACCATCAGCAGTC TGCAGGCTGAAGACCTTGCAGATTATCACTGTGGACAGGGTTACAGCTATCTTCC |
| IGKV3-10*01_BALB | BALB/cByJ | Musmus IGKV3-10*01 F | 100 | 299 | AACATTGTGCTGACCCAATCTCCAGCTTCTTTGGCTGTGTCTCTAGGGCAGAGGGCCA CCATATCCTGCAGAGCCAGTGAAAGTGTTGATAGTTATGGCAATAGTTTTATGCACTG GTACCAGCAGAAACCAGGACAGCCACCCAAACTCCTCATCTATCTTGCATCCAACCTA GAATCTGGGGTCCCTGCCAGGTTCAGTGGCAGTGGGTCTAGGACAGACTTCACCCTCA CCATTGATCCTGTGGAGGCTGATGATGCTGCAACCTATTACTGTCAGCAAAATAATGA GGATCCTCC |

| | | | | | |
|---|---|---|---|---|---|
| IGKV11-125*01_BALB | BALB/cByJ | Musmus IGKV11-125*01 F | 100 | 287 | GATGTCCAGATGATTCAGTCTCCATCCTCCCTGTCTGCATCTTTGGGAGACATAGTCACCATGACTTGCCAGGCAAGTCAGGGCACTAGCATTAATTTAAACTGGTTTCAGCAAAAACCAGGGAAAGCTCCTAAGCTCCTGATCTATGGTGCAAGCAACTTGGAAGATGGGGTCCCATCAAGGTTCAGTGGCAGTAGATATGGGACAGATTTCACTCTCACCATCAGCAGCCTGGAGGATGAAGATATGGCAACTTATTTCTGTCTACAGCATAGTTATCTCCCTCC |
| IGKV4-58*01_BALB | BALB/cByJ | Musmus IGKV4-58*01 F | 100 | 292 | GAAAATGTGCTCACCCAGTCTCCAGCAATAATGGCTGCCTCTCTGGGGCAGAAGGTCACCATGACCTGCAGTGCCAGCTCAAGTGTAAGTTCCAGTTACTTGCACTGGTACCAGCAGAAGTCAGGCGCTTCCCCCAAACCCTTGATTCATAGGACATCCAACCTGGCTTCTGGAGTCCCAGCTCGCTTCAGTGGCAGTGGGTCTGGGACCTCTTACTCTCTCACAATCAGCAGCGTGGAGGCTGAAGATGATGCAACTTATTACTGCCAGCAGTGGAGTGGTTACCCATTCA |
| IGKV4-52*01_BALB | BALB/cByJ | Musmus IGKV4-52*01 ORF, or Musmus IGKV4-54*01 ORF | 100 | 286 | CAAATTGTTCTCACCCAGTCTCCAGCAATCCTGTCTGCATCTCCAGGGGAGAAGGTCACCATGACCTGCAGTGCCAGCTCAAGTGTAAGTTACATGTACAGGTACCAGCAGAAGCCAGGATCCTCACCCAAACCCTGGATTTATGGCACATCCAACCTGGCTTCTGGAGTCCCTGCTCGCTTCAGTGGCAGTGGATCTGGGACCTCTTATTCTCTCACAATCAGCAGCATGGAGGCTGAAGATGCTGCCACTTATTACTGCCAGCAATATCATAGTTACCCACCCA |
| IGKV1-133*01_BALB | BALB/cByJ | Musmus IGKV1-133*01 F | 100 | 302 | GATGTTGTGATGACCCAGACTCCACTCACTTTGTCGGTTACCATTGGACAACCAGCCTCTATCTCTTGCAAGTCAAGTCAGAGCCTCTTATATAGTAATGGAAAAACCTATTTGAATTGGTTATTACAGAGGCCAGGCCAGTCTCCAAAGCGCCTAATCTATCTGGTGTCTAAACTGGACTCTGGAGTCCCTGACAGGTTCACTGGCAGTGGATCAGGAACAGATTTTACACTGAAAATCAGCAGAGTGGAGGCTGAGGATTTGGGAGTTTATTACTGCGTGCAAGGTACACATTTTCCTCA |
| IGKV5-37*01_BALB | BALB/cByJ | Musmus IGKV5-37*01 F | 100 | 287 | GACATCCTGCTGACCCAGTCTCCAGCCACCCTGTCTGTGACTCCAGGAGAAACAGTCAGTCTTTCCTGTAGGGCCAGCCAGAGTATTTACAAGAACCTACACTGGTATCAACAGAAATCACATCGGTCTCCAAGGCTTCTCATCAAGTATGCTTCTGATTCCATCTCTGGGATCCCCTCCAGGTTCACTGGCAGTGGATCAGGGACAGATTACACTCTCAGTATCAACAGTGTGAAGCCCGAAGATGAAGGAATATATTACTGTCTTCAAGGTTACAGCACACCTTC |
| IGKV8-23-1*01_BALB | BALB/cByJ | Musmus IGKV8-16*01 F, or Musmus IGKV8-23-1*01 ORF | 100 | 284 | GAAATTGTGCTGACTGTGTCAGCAGGAGAGAGGGTCACTATCAGCTGCAAATCCAATCAGAATCTTTTATGGAGTGGAAACCAAAGGTACTGTTTGGTCTGGCACCAGTGGAAACCAGGGCAAACTCCTACACCGTTGATCACCTGGACATCTGATAGGTACTCTGGAGTCCCTGATCGTTTCATAGGCAGTGGATCTGTGACAGATTTCACTCTGACCATCAGCAGTGTGCAGGCTGAAGATGTGGCAGTTTATTTCTGTCAGCAGCATTTACACATTCCTCC |
| IGKV9-124*01_BALB | BALB/cByJ | Musmus IGKV9-124*01 F | 100 | 285 | GACATCCAGATGACCCAGTCTCCATCCTCCTTATCTGCCTCTCTGGGAGAAAGAGTCAGTCTCACTTGTCGGGCAAGTCAGGAAATTAGTGGTTACTTAAGCTGGCTTCAGCAGAAACCAGATGGAACTATTAAACGCCTGATCTACGCCGCATCCACTTTAGATTCTGGTGTCCCAAAAAGGTTCAGTGGCAGTAGGTCTGGGTCAGATTATTCTCTCACCATCAGCAGCCTTGAGTCTGAAGATTTTGCAGACTATTACTGTCTACAATATGCTAGTTATCCT |
| IGKV3-1*01_BALB | BALB/cByJ | Musmus IGKV3-1*01 F | 100 | 299 | GACATTGTGCTCACCCAATCTCCAGCTTCTTTGGCTGTGTCTCTAGGGCAGAGAGCCACCATCTCCTGCAGAGCCAGTGAAAGTGTTGAATATTATGGCACAAGTTTAATGCAGTGGTACCAACAGAAACCAGGACAGCCACCCAAACTCCTCATCTATGCTGCATCCAACGTAGAATCTGGGGTCCCTGCCAGGTTTAGTGGCAGTGGGTCTGGGACAGACTTCAGCCTCAACATCCATCCTGTGGAGGAGGATGATATTGCAATGTATTTCTGTCAGCAAAGTAGGAAGGTTCCTTC |

| | | | | | |
|---|---|---|---|---|---|
| IGKV9-123*01_BALB | BALB/cByJ | Musmus IGKV9-123*01 F | 100 | 285 | GACATCCAGATGATTCAGTCTCCATCGTCCATGTTTGCCTCTCTGGGAGACAGAGTCAGTCTCTCTTGTCGGGCTAGTCAGGGCATTAGAGGTAATTTAGACTGGTATCAGCAGAAACCAGGTGGAACTATTAAACTCCTGATCTACTCCACATCCAATTTAAATTCTGGTGTCCCATCAAGGTTCAGTGGCAGTGGGTCTGGGTCAGATTATTCTCTCACCATCAGCAGCCTAGAGTCTGAAGATTTTGCAGACTATTACTGTCTACAGCGTAATGCGTATCCT |
| IGKV12-38*01_BALB | BALB/cByJ | Musmus IGKV12-38*01 F | 100 | 287 | GACATCCAGATGACTCAGTCTCCAGCCTCCCTGGCTGCATCTGTGGGAGAAACTGTCACCATCACATGTCGAGCAAGTGAGAACATTTACTACAGTTTAGCATGGTATCAGCAGAAGCAAGGGAAATCTCCTCAGCTCCTGATCTATAATGCAAACAGCTTGGAAGATGGTGTCCCATCGAGGTTCAGTGGCAGTGGATCTGGGACACAGTATTCTATGAAGATCAACAGCATGCAGCCTGAAGATACCGCAACTTATTTCTGTAAACAGGCTTATGACGTTCCTCC |
| IGKV4-51*01_BALB | BALB/cByJ | Musmus IGKV4-51*01 F | 100 | 292 | GAAAATGTGCTCACCCAGTCTCCAGCAATAATGGCTGCCTCTCTGGGGGAGAAGGTCACCATGACCTGCAGTGCCAGCTCAAGTGTAAGTTCCAGCTACTTGCACTGGTACCAGCAGAAGTCAGGCACTTCCCCCAAACTCTGGATTTATGGCACATCCAACCTGGCTTCTGGAGTCCCAGCTCGCTTCAGTGGCAGTGGGGCTGGGATCTCTTACTCTCTCACAATCAGCAGCATGGAGGCTGAAAATGATGCAACTTATTACTGCCAGCAGTGGAGTGGTTACCCATTCA |
| IGKV4-92*01_BALB | BALB/cByJ | Musmus IGKV4-92*01 F | 100 | 291 | GAAATGGTTCTCACCCAGTCTCCAGTATCCATAACTGCATCTCGAGGGGAGAAGGTCACCATCACCTGCCGTGCCAGCTCAAGTATAAGTTCCAATTACTTACACTGGTACCAGCAGAAGCCAGGATCCTCCCCTAAACTTTTGATTTATAGGACATCCATCCTGGCATCTGGAGTCCTAGACAGCTTCAGTGGCAGTGGGTCTGAGAGCTCTTACACTCTGACAATCAGCTGCATGCAGGACGAAGTTGCTGCCACTTACTATTGTCAGCAGGGGAGTAGTAGCCCACCA |
| IGKV4-56*01_BALB | BALB/cByJ | Musmus IGKV4-56*01 P | 100 | 286 | CAAATTGTTCTCACCCAGTCTCCAGCAATCATGTCTGCATCTCCAGGGCAGAAAGTCACCATAACCTGCAGTGCCATCTCAAGTGTAAATTACATGCACTGGTACCAGCAGAAGCCAGGATCCTCCCCCAAACTCTGGATTTATGCAACATCCAAACTGGCTCTTGGAGTCCCTGCTTGCTTCAGTGGCAGTGGGTCTGGGACCTCTTACTCTCTCACAATCAGCAGCATGGTGGCTGAAGATGCCACCTCTTATTTCTGTCATCAGTGGAGTAGTTACCCACCCA |
| IGKV1-99*01_BALB | BALB/cByJ | Musmus IGKV1-99*01 F | 100 | 302 | GATGTTGTTCTGACCCAAACTCCACTCTCTCTGCCTGTCAATATTGGAGATCAAGCCTCTATCTCTTGCAAGTCTACTAAGAGTCTTCTGAATAGTGATGGATTCACTTATTTGGACTGGTACCTGCAGAAGCCAGGCCAGTCTCCACAGCTCCTAATATATTTGGTTTCTAATCGATTTTCTGGAGTTCCAGACAGGTTCAGTGGCAGTGGGTCAGGAACAGATTTCACACTCAAGATCAGCAGAGTGGAGGCTGAGGATTTGGGAGTTTATTATTGCTTCCAGAGTAACTATCTTCCTCT |
| IGKV10-95*01_BALB | BALB/cByJ | Musmus IGKV10-95*01 F | 100 | 287 | GATATCCAGATGACACAGACTACTTCCTCCCTGTCTGCCTCTCTGGGAGACAGAGTCACCATCAGTTGCAGGGCAAGTGAGGACATTAGCACTTATTTAAACTGGTATCAGCAGAAACCAGATGGAACTGTTAAACTCCTGATCTATTACACATCAGGATTACACTCAGGAGTCCCATCAAGGTTCAGTGGCAGTGGGTCTGGGGCAGATTATTCTCTCACCATCAGCAACCTGGAACCTGAAGATATTGCCACTTACTATTGTCAGCAGTATAGTAAGCTTCCTCC |
| IGKV8-18*01_BALB | BALB/cByJ | Musmus IGKV8-18*01 ORF | 100 | 310 | GACATTGTGATGACCCAGTCTCCATCCTCCCTGGCTGTGACAGCAGGAGAGAAGGTCACTATGAGATGCAAGTCCAGTCAGAGTCTTTTGTGGAGTGTAAACCAAAATAACTACTTATCCTGGTACCAGCAGAAACAAGGGCAGCCTCCTAAACTGCTTATCTATGGGGCATCCATTAGAGAATCTTGGGTCCCTGATCGATTCACAGGAAGTGGATCTGGGACAGACTTCACTCTCACCATTAGCAATGTGCATGCTGAAGACCTAGCAGTTTATTACTGTCAGCACAATCATGGCAGCTTTCTCCCCC |

| | | | | | |
|---|---|---|---|---|---|
| IGKV9-129*01_BALB | BALB/cByJ | Musmus IGKV9-129*01 ORF | 100 | 287 | GACATCCAGATGACCCAGTCTCCATCCTCCTTATCTGCCTCTCTGGGAGAAAGAGTCAGTCTCACTTGCCGGGCAAGTCAGGACATTCATGGTTATTTAAACTTGTTTCAGCAGAAACCAGGTGAAACTATTAAACACCTGATCTATGAAACATCCAATTTAGATTCTGGTGTCCCAAAAAGGTTCAGTGGCAGTAGGTCTGGGTCAGATTATTCTCTCATTATCGGCAGCCTTGAGTCTGAAGATTTTGCAGACTATTACTGTCTACAATATGCTAGTTCTCCTCC |
| IGKV1-132*01_BALB | BALB/cByJ | Musmus IGKV1-132*01 F | 100 | 302 | GATGTTGTGATGACCCAGACTCCACTGTCTTTGTCGGTTACCATTGGACAACCAGCCTCTATCTCTTGCAAGTCAAGTCAGAGCCTCTTATATAGTAATGGAAAGACATATTTGAATTGGTTACAACAGAGGCCTGGCCAGGCTCCAAAGCACCTAATGTATCAGGTGTCCAAACTGGACCCTGGCATCCCTGACAGGTTCAGTGGCAGTGGATCAGAAACAGATTTTACACTTAAAATCAGCAGAGTGGAGGCTGAAGATTTGGGAGTTTATTACTGCTTGCAAGGTACATATTATCCTCA |
| IGKV14-130*01_BALB | BALB/cByJ | Musmus IGKV14-130*01 F | 100 | 287 | GAAATCCAGATGACCCAGTCTCCATCCTCTATGTCTGCATCTCTGGGAGACAGAATAACCATCACTTGCCAGGCAACTCAAGACATTGTTAAGAATTTAAACTGGTATCAGCAGAAACCAGGGAAACCCCCTTCATTCCTGATCTATTATGCAACTGAACTGGCAGAAGGGGTCCCATCAAGGTTCAGTGGCAGTGGGTCTGGGTCAGACTATTCTCTGACAATCAGCAACCTGGAGTCTGAAGATTTTGCAGACTATTACTGTCTACAGTTTTATGAGTTTCCTCC |
| IGKV4-81*01_S0130_BALB | BALB/cByJ | Musmus IGKV4-81*01 F | 100 | 292 | GAAAATGTGCTGACCCAGTCTCCAGCAATCATGGCTGCATCTCCAGGGGAGAAGGTCACCATGACCTGCAGTGCCAGCTCAAGTGTAAGTTCTAGTAACTTGCACTGGTACCAGCAGAAGTCAGGCACTTCTACCAAATTCTGGATTTATAGGACATCCAACCTGGCTTCAGAAGTCCCAGCTCCCTTCAGTGGCAGTGGGTCTGGGACCTCTTACTCTCTTACAATCAGCAGCGTGGAGGCCGAAGATGCTGCCACTTATTACTGCCAGCAGTGGAGTGGTTACCCACTCA |
| IGKV4-73*01_BALB | BALB/cByJ | Musmus IGKV4-73*01 ORF | 100 | 292 | CAAATTGTTCTCACCCAGTCTCCAGCAATCATGTCTGCATCTCCTGGGGAACGGGTCACCATGACCTGCAGTGCCAGCTCAAGTGTAAGTTCCAGCTACTTGTACTGGTACCAGCAGAAGTCAGGATCCTCCCCAAAACTCTGGATTTATAGCATATCCAACCTGGCTTCTGGAGTCCCAGCTCGCTTCAGTGGCAGTGGGTCTGGGACCTCTTACTCTCTCACAATCAACAGCATGGAGGCTGAAGATGCTGCCACTTATTACTGCCAGCAGTGGAGTAGTAACCCACCCA |
| IGKV3-9*01_BALB | BALB/cByJ | Musmus IGKV3-9*01 F | 100 | 299 | GACATTGTGCTGACCCAATCTCCAGCTTCTTTGGCTGTGTCTCTAGGACAGAGGGCCACCATATCCTGCCAAGCCAGCGAAAGTGTCAGTTTTGCTGGTACAAGTTTAATGCACTGGTACCAACAGAAACCAGGACAGCCACCCAAACTCCTCATCTATCGTGCATCCAACCTAGAATCTGGAGTCCCTGCCAGGTTCAGTGGCAGTGGGTCTGAGTCAGACTTCACTCTCACCATCGATCCTGTGGAGGAAGATGATGCTGCAATGTATTACTGTATGCAAAGTATGGAAGATCCTCC |
| IGKV4-60*01_BALB | BALB/cByJ | Musmus IGKV4-60*01 P | 100 | 292 | GAAATTGTGCTCACCCAGTCTCCAGCACTCATGGCTGCATCTCCAGGGGAGAAGGTCACCATCACCTGCAGTGTCAGCTCAAGTATAAGTTCCAGCTACTTGCACTGGTACCAGCAGAAGTCAGGAATCTCCCCCAAACCCTGGATTTATGGCACATCCAACCTGGCTTCTGGAGTCCCTGCTCGCTTCAGTGGCAGTGGATCTGGGACCTCTTACTCTCTCACAATCAGCAGCATGGAGGCTGAAGATGCTGCCACTTATTACTGTCAACAGTGGAGTAGTTCCCCACCCA |

| | | | | | |
|---|---|---|---|---|---|
| IGKV4-62*01_BALB | BALB/cByJ | Musmus IGKV4-62*01 ORF | 100 | 286 | CAAATTGTTCTCACCCAGTCTCCAGCAATCATGTCTGCATCTCCAGGGGAAAAGGTCACCATGACCTGCAGTGCCAGCTCAAGTGTAAGTTACATGTACTGGTACCAGCAGAAGCCAGGCTCCTCCCCCAGACTCTGGATTTATGACACATCCAACCTGGTTTCTGGAGTCCCTGCTCGCTTCAGTGGCAGTAGGTCTGGGACCTCTTATTCTCTCACAATCAGCAGCATGGAGGCTGAAGATGCTGCCACTTATTACTGCCAGCAGTACAGTGGTTACCCATCCA |
| IGKV4-78*01_BALB | BALB/cByJ | Musmus IGKV4-78*01 F | 100 | 292 | CAAATTGTTCTCACCCAGTCTCCAGCAATCATGTCTGCATCTCCTGGGGAGAAGGTCACCATGACCTGCAGTGCCAGATCAAGTGTAAGTTCCAGCTACTTGTACTGGTACCAGCAGAAGCCAGGATCCTCCCCCAAACTCTGGATTTATAGCACATCCAACCTGGCTTCTGGAGTCCCTGCTCGCTTCAGTGGCAGTGGGTCTGGGACCTCTTATTCTCTCACAATCAGCAGCATGGAGGCTGAAGATGCTGCCACTTTTTACTGCCAGCAGTACAGTGGTTACCCATCCA |
| IGKV4-90*01_BALB | BALB/cByJ | Musmus IGKV4-90*01 F | 100 | 285 | GAAATTTTGCTCACCCAGTCTCCAGCAATCATAGCTGCATCTCCTGGGGAGAAGGTCACCATCACCTGCAGTGCCAGCTCAAGTGTAAGTTACATGAACTGGTACCAGCAGAAACCAGGATCCTCCCCCAAAATATGGATTTATGGTATATCCAACCTGGCTTCTGGAGTTCCTGCTCGCTTCAGTGGCAGTGGGTCTGGGACATCTTTCTCTTTCACAATCAACAGCATGGAGGCTGAAGATGTTGCCACTTATTACTGTCAGCAAAGGAGTAGTTACCCACCC |
| IGKV6-14*01_BALB | BALB/cByJ | Musmus IGKV6-14*01 F | 100 | 287 | GACATTGTGATGACCCAGTCTCAAAAATTCATGTCCACATCAGTAGGAGACAGGGTCAGCATCACCTGCAAGGCCAGTCAGAATGTTCGTACTGCTGTAGCCTGGTATCAACAGAAACCAGGGCAGTCTCCTAAAGCACTGATTTACTTGGCATCCAACCGGCACACTGGAGTCCCTGATCGCTTCACAGGCAGTGGATCTGGGACAGATTTCACTCTCACCATTAGCAATGTGCAATCTGAAGACCTGGCAGATTATTTCTGTCTGCAACATTGGAATTATCCTCT |
| IGKV8-26*01_BALB | BALB/cByJ | Musmus IGKV8-26*01 ORF | 100 | 311 | GACATTGTGATGACCCAGTCTCCATCCTCCCTGGCTGTGACAGCAGGAGAGAAGGTCACTATGAGCTGCAAGTCCAGTCAGAGTCTTTTGTGGAGTGTAAACCAAAAGAACTACTTGTCCTGGTACCAGCAGAAACAAAGGCAGCCTCCTAAACTGCTTATCTATGGGGCATCCATTAGAGAATCTTGGGTCCCTGATCGGTTCACAGGAAGTGGATCTGGGACAGACTTCACTCTCACCATTAGCAATGTGCATGCTGAAGACCTAGCAGTTTATTACTGTCAACACAATCATGGCAGCTTTCTCCCCCC |
| IGKV8-30*01_C3H | C3H/HeJ | Musmus IGKV8-30*01 F | 100 | 305 | GACATTGTGATGTCACAGTCTCCATCCTCCCTAGCTGTGTCAGTTGGAGAGAAGGTTACTATGAGCTGCAAGTCCAGTCAGAGCCTTTTATATAGTAGCAATCAAAAGAACTACTTGGCCTGGTACCAGCAGAAACCAGGGCAGTCTCCTAAACTGCTGATTTACTGGGCATCCACTAGGGAATCTGGGGTCCCTGATCGCTTCACAGGCAGTGGATCTGGGACAGATTTCACTCTCACCATCAGCAGTGTGAAGGCTGAAGACCTGGCAGTTTATTACTGTCAGCAATATTATAGCTATCCTCC |
| IGKV6-15*01_C3H | C3H/HeJ | Musmus IGKV6-15*01 F | 100 | 287 | GACATTGTGATGACCCAGTCTCAAAAATTCATGTCCACATCAGTAGGAGACAGGGTCAGCGTCACCTGCAAGGCCAGTCAGAATGTGGGTACTAATGTAGCCTGGTATCAACAGAAACCAGGGCAATCTCCTAAAGCACTGATTTACTCGGCATCCTACCGGTACAGTGGAGTCCCTGATCGCTTCACAGGCAGTGGATCTGGGACAGATTTCACTCTCACCATCAGCAATGTGCAGTCTGAAGACTTGGCAGAGTATTTCTGTCAGCAATATAACAGCTATCCTCT |
| IGKV14-126*01_C3H | C3H/HeJ | Musmus IGKV14-126*01 F | 100 | 287 | GACATCAAGATGACCCAGTCTCCATCCTCCATGTATGCATCGCTGGGAGAGAGAGTCACTATCACTTGCAAGGCGAGTCAGGACATTAAAAGCTATTTAAGCTGGTACCAGCAGAAACCATGGAAATCTCCTAAGACCCTGATCTATTATGCAACAAGCTTGGCAGATGGGGTCCCATCAAGATTCAGTGGCAGTGGATCTGGGCAAGATTATTCTCTAACCATCAGCAGCCTGGAGTCTGACGATACAGCAACTTATTACTGTCTACAGCATGGTGAGAGCCCTCC |

| | | | | | |
|---|---|---|---|---|---|
| IGKV8-27*01_C3H | C3H/HeJ | Musmus IGKV8-27*01 F | 100 | 300 | AACATTATGATGACACAGTCGCCATCATCTCTGGCTGTGTCTGCAGGAGAAAAGGTCACTATGAGCTGTAAGTCCAGTCAAAGTGTTTTATACAGTTCAAATCAGAAGAACTACTTGGCCTGGTACCAGCAGAAACCAGGGCAGTCTCCTAAACTGCTGATCTACTGGGCATCCACTAGGGAATCTGGTGTCCCTGATCGCTTCACAGGCAGTGGATCTGGGACAGATTTTACTCTTACCATCAGCAGTGTACAAGCTGAAGACCTGGCAGTTTATTACTGTCATCAATACCTCTCCTCG |
| IGKV4-57*01_S9174_C3H | C3H/HeJ | Musmus IGKV4-57-1*01 F | 100 | 286 | CAAATTGTTCTCACCCAGTCTCCAGCAATCATGTCTGCATCTCCAGGGGAGAAGGTCACCATAACCTGCAGTGCCAGCTCAAGTGTAAGTTACATGCACTGGTTCCAGCAGAAGCCAGGCACTTCTCCCAAACTCTGGATTTATAGCACATCCAACCTGGCTTCTGGAGTCCCTGCTCGCTTCAGTGGCAGTGGATCTGGGACCTCTTACTCTCTCACAATCAGCCGAATGGAGGCTGAAGATGCTGCCACTTATTACTGCCAGCAAAGGAGTAGTTACCCACACA |
| IGKV1-135*01_C3H | C3H/HeJ | Musmus IGKV1-135*01 F | 100 | 302 | GATGTTGTGATGACCCAGACTCCACTCACTTTGTCGGTTACCATTGGACAACCAGCCTCCATCTCTTGCAAGTCAAGTCAGAGCCTCTTAGATAGTGATGGAAAGACATATTTGAATTGGTTGTTACAGAGGCCAGGCCAGTCTCCAAAGCGCCTAATCTATCTGGTGTCTAAACTGGACTCTGGAGTCCCTGACAGGTTCACTGGCAGTGGATCAGGGACAGATTTCACACTGAAAATCAGCAGAGTGGAGGCTGAGGATTTGGGAGTTTATTATTGCTGGCAAGGTACACATTTTCCTCA |
| IGKV12-44*01_C3H | C3H/HeJ | Musmus IGKV12-44*01 F | 100 | 287 | GACATCCAGATGACTCAGTCTCCAGCCTCCCTATCTGCATCTGTGGGAGAAACTGTCACCATCACATGTCGAGCAAGTGAGAATATTTACAGTTATTTAGCATGGTATCAGCAGAAACAGGGAAAATCTCCTCAGCTCCTGGTCTATAATGCAAAAACCTTAGCAGAAGGTGTGCCATCAAGGTTCAGTGGCAGTGGATCAGGCACACAGTTTTCTCTGAAGATCAACAGCCTGCAGCCTGAAGATTTTGGGAGTTATTACTGTCAACATCATTATGGTACTCCTCC |
| IGKV12-46*01_C3H | C3H/HeJ | Musmus IGKV12-46*01 F | 100 | 287 | GACATCCAGATGACTCAGTCTCCAGCCTCCCTATCTGTATCTGTGGGAGAAACTGTCACCATCACATGTCGAGCAAGTGAGAATATTTACAGTAATTTAGCATGGTATCAGCAGAAACAGGGAAAATCTCCTCAGCTCCTGGTCTATGCTGCAACAAACTTAGCAGATGGTGTGCCATCAAGGTTCAGTGGCAGTGGATCAGGCACACAGTATTCCCTCAAGATCAACAGCCTGCAGTCTGAAGATTTTGGGAGTTATTACTGTCAACATTTTTGGGGTACTCCTCC |
| IGKV6-23*01_C3H | C3H/HeJ | Musmus IGKV6-23*01 F | 100 | 287 | GACATTGTGATGACCCAGTCTCACAAATTCATGTCCACATCAGTAGGAGACAGGGTCAGCATCACCTGCAAGGCCAGTCAGGATGTGGGTACTGCTGTAGCCTGGTATCAACAGAAACCAGGGCAATCTCCTAAACTACTGATTTACTGGGCATCCACCCGGCACACTGGAGTCCCTGATCGCTTCACAGGCAGTGGATCTGGGACAGATTTCACTCTCACCATTAGCAATGTGCAGTCTGAAGACTTGGCAGATTATTTCTGTCAGCAATATAGCAGCTATCCTCT |
| IGKV2-137*01_C3H | C3H/HeJ | Musmus IGKV2-137*01 F | 100 | 302 | GATATTGTGATGACTCAGGCTGCACCCTCTGTACCTGTCACTCCTGGAGAGTCAGTATCCATCTCCTGCAGGTCTAGTAAGAGTCTCCTGCATAGTAATGGCAACACTTACTTGTATTGGTTCCTGCAGAGGCCAGGCCAGTCTCCTCAGCTCCTGATATATCGGATGTCCAACCTTGCCTCAGGAGTCCCAGACAGGTTCAGTGGCAGTGGGTCAGGAACTGCTTTCACACTGAGAATCAGTAGAGTGGAGGCTGAGGATGTGGGTGTTTATTACTGTATGCAACATCTAGAATATCCTTT |
| IGKV8-19*01_C3H | C3H/HeJ | Musmus IGKV8-19*01 F | 100 | 305 | GACATTGTGATGACACAGTCTCCATCCTCCCTGACTGTGACAGCAGGAGAGAAGGTCACTATGAGCTGCAAGTCCAGTCAGAGTCTGTTAAACAGTGGAAATCAAAAGAACTACTTGACCTGGTACCAGCAGAAACCAGGGCAGCCTCCTAAACTGTTGATCTACTGGGCATCCACTAGGGAATCTGGGGTCCCTGATCGCTTCACAGGCAGTGGATCTGGAACAGATTTCACTCTCACCATCAGCAGTGTGCAGGCTGAAGACCTGGCAGTTTATTACTGTCAGAATGATTATAGTTATCCTCC |

| | | | | | |
|---|---|---|---|---|---|
| IGKV1-117*01_C3H | C3H/HeJ | Musmus IGKV1-117*01 F | 100 | 302 | GATGTTTTGATGACCCAAACTCCACTCTCCCTGCCTGTCAGTCTTGGAGATCAAGCCTCCATCTCTTGCAGATCTAGTCAGAGCATTGTACATAGTAATGGAAACACCTATTTAGAATGGTACCTGCAGAAACCAGGCCAGTCTCCAAAGCTCCTGATCTACAAAGTTTCCAACCGATTTTCTGGGGTCCCAGACAGGTTCAGTGGCAGTGGATCAGGGACAGATTTCACACTCAAGATCAGCAGAGTGGAGGCTGAGGATCTGGGAGTTTATTACTGCTTTCAAGGTTCACATGTTCCTCC |
| IGKV1-110*01_C3H | C3H/HeJ | Musmus IGKV1-110*01 F | 100 | 302 | GATGTTGTGATGACCCAAACTCCACTCTCCCTGCCTGTCAGTCTTGGAGATCAAGCCTCCATCTCTTGCAGATCTAGTCAGAGCCTTGTACACAGTAATGGAAACACCTATTTACATTGGTACCTGCAGAAGCCAGGCCAGTCTCCAAAGCTCCTGATCTACAAAGTTTCCAACCGATTTTCTGGGGTCCCAGACAGGTTCAGTGGCAGTGGATCAGGGACAGATTTCACACTCAAGATCAGCAGAGTGGAGGCTGAGGATCTGGGAGTTTATTTCTGCTCTCAAAGTACACATGTTCCTCC |
| IGKV8-21*01_C3H | C3H/HeJ | Musmus IGKV8-21*01 F | 100 | 302 | GACATTGTGATGTCACAGTCTCCATCCTCCCTGGCTGTGTCAGCAGGAGAGAAGGTCACTATGAGCTGCAAATCCAGTCAGAGTCTGCTCAACAGTAGAACCCGAAAGAACTACTTGGCTTGGTACCAGCAGAAACCAGGGCAGTCTCCTAAACTGCTGATCTACTGGGCATCCACTAGGGAATCTGGGGTCCCTGATCGCTTCACAGGCAGTGGATCTGGGACAGATTTCACTCTCACCATCAGCAGTGTGCAGGCTGAAGACCTGGCAGTTTATTACTGCAAGCAATCTTATAATCTTCC |
| IGKV4-59*01_C3H | C3H/HeJ | Musmus IGKV4-59*01 F | 100 | 286 | CAAATTGTTCTCACCCAGTCTCCAGCAATCATGTCTGCATCTCCAGGGGAGAAGGTCACCATGACCTGCAGTGCCAGCTCAAGTGTAAGTTACATGCACTGGTACCAGCAGAAGTCAGGCACCTCCCCCAAAAGATGGATTTATGACACATCCAAACTGGCTTCTGGAGTCCCTGCTCGCTTCAGTGGCAGTGGGTCTGGGACCTCTTACTCTCTCACAATCAGCAGCATGGAGGCTGAAGATGCTGCCACTTATTACTGCCAGCAGTGGAGTAGTAACCCACCCA |
| IGKV10-96*01_C3H | C3H/HeJ | Musmus IGKV10-96*01 F | 100 | 286 | GATATCCAGATGACACAGACTACATCCTCCCTGTCTGCCTCTCTGGGAGACAGAGTCACCATCAGTTGCAGGGCAAGTCAGGACATTAGCAATTATTTAAACTGGTATCAGCAGAAACCAGATGGAACTGTTAAACTCCTGATCTACTACACATCAAGATTACACTCAGGAGTCCCATCAAGGTTCAGTGGCAGTGGGTCTGGAACAGATTATTCTCTCACCATTAGCAACCTGGAGCAAGAAGATATTGCCACTTACTTTTGCCAACAGGGTAATACGCTTCCTC |
| IGKV5-43*01_C3H | C3H/HeJ | Musmus IGKV5-43*01 F | 100 | 287 | GATATTGTGCTAACTCAGTCTCCAGCCACCCTGTCTGTGACTCCAGGAGATAGCGTCAGTCTTTCCTGCAGGGCCAGCCAAAGTATTAGCAACAACCTACACTGGTATCAACAAAAATCACATGAGTCTCCAAGGCTTCTCATCAAGTATGCTTCCCAGTCCATCTCTGGGATCCCCTCCAGGTTCAGTGGCAGTGGATCAGGGACAGATTTCACTCTCAGTATCAACAGTGTGGAGACTGAAGATTTTGGAATGTATTTCTGTCAACAGAGTAACAGCTGGCCTCA |
| IGKV4-70*01_C3H | C3H/HeJ | Musmus IGKV4-70*01 F | 100 | 282 | CAAATTGTTCTCACCCAGTCTCCAGCAATCATGTCTGCATCTCCAGGGGAGAAGGTCACCATGACCTGCAGTGCCAGCTCAAGTATAAGTTACATGCACTGGTACCAGCAGAAGCCAGGCACCTCCCCCAAAAGATGGATTTATGACACATCCAAACTGGCTTCTGGAGTCCCTGCTCGCTTCAGTGGCAGTGGGTCTGGGACCTCTTATTCTCTCACAATCAGCAGCATGGAGGCTGAAGATGCTGCCACTTATTACTGCCATCAGCGGAGTAGTTACCCA |
| IGKV12-41*01_C3H | C3H/HeJ | Musmus IGKV12-41*01 F | 100 | 287 | GACATCCAGATGACTCAGTCTCCAGCCTCCCTATCTGCATCTGTGGGAGAAACTGTCACCATCACATGTCGAGCAAGTGGGAATATTCACAATTATTTAGCATGGTATCAGCAGAAACAGGGAAAATCTCCTCAGCTCCTGGTCTATAATGCAAAAACCTTAGCAGATGGTGTGCCATCAAGGTTCAGTGGCAGTGGATCAGGAACACAATATTCTCTCAAGATCAACAGCCTGCAGCCTGAAGATTTTGGGAGTTATTACTGTCAACATTTTTGGAGTACTCCTCC |

| | | | | | |
|---|---|---|---|---|---|
| IGKV6-20*01_C3H | C3H/HeJ | Musmus IGKV6-20*01 F | 100 | 287 | AACATTGTAATGACCCAATCTCCCAAATCCATGTCCATGTCAGTAGGAGAGAGGGTCACCTTGAGCTGCAAGGCCAGTGAGAATGTGGGTACTTATGTATCCTGGTATCAACAGAAACCAGAGCAGTCTCCTAAACTGCTGATATACGGGGCATCCAACCGGTACACTGGGGTCCCCGATCGCTTCACAGGCAGTGGATCTGCAACAGATTTCACTCTGACCATCAGCAGTGTGCAGGCTGAAGACCTTGCAGATTATCACTGTGGACAGAGTTACAGCTATCCTCC |
| IGKV8-28*01_C3H | C3H/HeJ | Musmus IGKV8-28*01 F | 100 | 305 | GACATTGTGATGACACAGTCTCCATCCTCCCTGAGTGTGTCAGCAGGAGAGAAGGTCACTATGAGCTGCAAGTCCAGTCAGAGTCTGTTAAACAGTGGAAATCAAAAGAACTACTTGGCCTGGTACCAGCAGAAACCAGGGCAGCCTCCTAAACTGTTGATCTACGGGGCATCCACTAGGGAATCTGGGGTCCCTGATCGCTTCACAGGCAGTGGATCTGGAACCGATTTCACTCTTACCATCAGCAGTGTGCAGGCTGAAGACCTGGCAGTTTATTACTGTCAGAATGATCATAGTTATCCTCC |
| IGKV6-32*01_C3H | C3H/HeJ | Musmus IGKV6-32*01 F | 100 | 287 | AGTATTGTGATGACCCAGACTCCCAAATTCCTGCTTGTATCAGCAGGAGACAGGGTTACCATAACCTGCAAGGCCAGTCAGAGTGTGAGTAATGATGTAGCTTGGTACCAACAGAAGCCAGGGCAGTCTCCTAAACTGCTGATATACTATGCATCCAATCGCTACACTGGAGTCCCTGATCGCTTCACTGGCAGTGGATATGGGACGGATTTCACTTTCACCATCAGCACTGTGCAGGCTGAAGACCTGGCAGTTTATTTCTGTCAGCAGGATTATAGCTCTCCTCC |
| IGKV8-24*01_C3H | C3H/HeJ | Musmus IGKV8-24*01 F | 100 | 305 | GACATTGTGATGACACAGTCTCCATCCTCCCTGGCTATGTCAGTAGGACAGAAGGTCACTATGAGCTGCAAGTCCAGTCAGAGCCTTTTAAATAGTAGCAATCAAAAGAACTATTTGGCCTGGTACCAGCAGAAACCAGGACAGTCTCCTAAACTTCTGGTATACTTTGCATCCACTAGGGAATCTGGGGTCCCTGATCGCTTCATAGGCAGTGGATCTGGGACAGATTTCACTCTTACCATCAGCAGTGTGCAGGCTGAAGACCTGGCAGATTACTTCTGTCAGCAACATTATAGCACTCCTCC |
| IGKV6-13*01_C3H | C3H/HeJ | Musmus IGKV6-13*01 F | 100 | 285 | GACATTGTGATGACCCAGTCTCAAAAATTCATGTCCACATCAGTAGGAGACAGGGTCAGCATCACCTGCAAGGCCAGTCAGAATGTGGGTACTGCTGTAGCCTGGTATCAACAGAAACCAGGACAATCTCCTAAACTACTGATTTACTCGGCATCCAATCGGTACACTGGAGTCCCTGATCGCTTCACAGGCAGTGGATCTGGGACAGATTTCACTCTCACCATCAGCAATATGCAGTCTGAAGACCTGGCAGATTATTTCTGCCAGCAATATAGCAGCTATCCT |
| IGKV8-18*01_C3H | C3H/HeJ | Musmus IGKV8-18*01 ORF | 100 | 310 | GACATTGTGATGACCCAGTCTCCATCCTCCCTGGCTGTGACAGCAGGAGAGAAGGTCACTATGAGATGCAAGTCCAGTCAGAGTCTTTTGTGGAGTGTAAACCAAAATAACTACTTATCCTGGTACCAGCAGAAACAAGGGCAGCCTCCTAAACTGCTTATCTATGGGGCATCCATTAGAGAATCTTGGGTCCCTGATCGATTCACAGGAAGTGGATCTGGGACAGACTTCACTCTCACCATTAGCAATGTGCATGCTGAAGACCTAGCAGTTTATTACTGTCAGCACAATCATGGCAGCTTTCTCCCCC |
| IGKV15-103*01_C3H | C3H/HeJ | Musmus IGKV15-103*01 ORF | 100 | 287 | GACATCCAGATGAACCAGTCTCCATCCAGTCTGTCTGCATCCCTTGGAGACACAATTACCATCACTTGCCATGCCAGTCAGAACATTAATGTTTGGTTAAGCTGGTACCAGCAGAAACCAGGAAATATTCCTAAACTATTGATCTATAAGGCTTCCAACTTGCACACAGGCGTCCCATCAAGGTTTAGTGGCAGTGGATCTGGAACAGGTTTCACATTAACCATCAGCAGCCTGCAGCCTGAAGACATTGCCACTTACTACTGTCAACAGGGTCAAAGTTATCCTCT |
| IGKV4-57-1*01_C3H | C3H/HeJ | Musmus IGKV4-57*01 F | 100 | 292 | GAAAATGTGCTCACCCAGTCTCCAGCAATCATGTCTGCATCTCCAGGGGAAAAGGTCACCATGACCTGCAGGGCCAGCTCAAGTGTAAGTTCCAGTTACTTGCACTGGTACCAGCAGAAGTCAGGTGCCTCCCCCAAACTCTGGATTTATAGCACATCCAACTTGGCTTCTGGAGTCCCTGCTCGCTTCAGTGGCAGTGGGTCTGGGACCTCTTACTCTCTCACAATCAGCAGTGTGGAGGCTGAAGATGCTGCCACTTATTACTGCCAGCAGTACAGTGGTTACCCACTCA |

| | | | | | |
|---|---|---|---|---|---|
| IGKV4-80*01_C3H | C3H/HeJ | Musmus IGKV4-80*01 F | 100 | 282 | CAAATTGTTCTCACCCAGTCTCCAGCAATCATGTCTGCATCTCTAGGGGAGGAGATCACCCTAACCTGCAGTGCCAGCTCGAGTGTAAGTTACATGCACTGGTACCAGCAGAAGTCAGGCACTTCTCCCAAACTCTTGATTTATAGCACATCCAACCTGGCTTCTGGAGTCCCTTCTCGCTTCAGTGGCAGTGGGTCTGGGACCTTTTATTCTCTCACAATCAGCAGTGTGGAGGCTGAAGATGCTGCCGATTATTACTGCCATCAGTGGAGTAGTTATCCA |
| IGKV3-2*01_C3H | C3H/HeJ | Musmus IGKV3-2*01 F | 100 | 299 | GACATTGTGCTGACCCAATCTCCAGCTTCTTTGGCTGTGTCTCTAGGGCAGAGGGCCACCATCTCCTGCAGAGCCAGCGAAAGTGTTGATAATTATGGCATTAGTTTTATGAACTGGTTCCAACAGAAACCAGGACAGCCACCCAAACTCCTCATCTATGCTGCATCCAACCAAGGATCCGGGGTCCCTGCCAGGTTTAGTGGCAGTGGGTCTGGGACAGACTTCAGCCTCAACATCCATCCTATGGAGGAGGATGATACTGCAATGTATTTCTGTCAGCAAAGTAAGGAGGTTCCTCC |
| IGKV19-93*01_C3H | C3H/HeJ | Musmus IGKV19-93*01 F | 100 | 287 | GACATCCAGATGACACAGTCTCCATCCTCACTGTCTGCATCTCTGGGAGGCAAAGTCACCATCACTTGCAAGGCAAGCCAAGACATTAACAAGTATATAGCTTGGTACCAACACAAGCCTGGAAAAGGTCCTAGGCTGCTCATACATTACACATCTACATTACAGCCAGGCATCCCATCAAGGTTCAGTGGAAGTGGGTCTGGGAGAGATTATTCCTTCAGCATCAGCAACCTGGAGCCTGAAGATATTGCAACTTATTATTGTCTACAGTATGATAATCTTCTACC |
| IGKV3-7*01_S5296_C3H | C3H/HeJ | Musmus IGKV3-7*02 F | 100 | 299 | GACATTGTGCTGACACAGTCTCCTGCTTCCTTAGCTGTATCTCTGGGGCAGAGGGCCACCATCTCATGCAGGGCCAGCCAAAGTGTCAGTACATCTAGCTATAGTTATATGCACTGGTACCAACAGAAACCAGGACAGCCACCCAAACTCCTCATCAAGTATGCATCCAACCTAGAATCTGGGGTCCCTGCCAGGTTCAGTGGCAGTGGGTCTGGGACAGACTTCACCCTCAACATCCATCCTGTGGAGGAGGAGGATACTGCAACATATTACTGTCAGCACAGTTGGGAGATTCCTCC |
| IGKV3-5*01_C3H | C3H/HeJ | Musmus IGKV3-5*01 F | 100 | 299 | GACATTGTGCTGACCCAATCTCCAGCTTCTTTGGCTGTGTCTCTAGGGCAGAGGGCCACCATATCCTGCAGAGCCAGTGAAAGTGTTGATAGTTATGGCAATAGTTTTATGCACTGGTACCAGCAGAAACCAGGACAGCCACCCAAACTCCTCATCTATCGTGCATCCAACCTAGAATCTGGGATCCCTGCCAGGTTCAGTGGCAGTGGGTCTAGGACAGACTTCACCCTCACCATTAATCCTGTGGAGGCTGATGATGTTGCAACCTATTACTGTCAGCAAAGTAATGAGGATCCTCC |
| IGKV4-53*01_C3H | C3H/HeJ | Musmus IGKV4-53*01 F | 100 | 292 | GAAATTGTGCTCACCCAGTCTCCAGCACTCATGGCTGCATCTCCAGGGGAGAAGGTCACCATCACCTGCAGTGTCAGCTCAAGTATAAGTTCCAGCAACTTGCACTGGTACCAGCAGAAGTCAGAAACCTCCCCCAAACCCTGGATTTATGGCACATCCAACCTGGCTTCTGGAGTCCCTGTTCGCTTCAGTGGCAGTGGATCTGGGACCTCTTATTCTCTCACAATCAGCAGCATGGAGGCTGAAGATGCTGCCACTTATTACTGTCAACAGTGGAGTAGTTACCCACTCA |
| IGKV6-17*01_C3H | C3H/HeJ | Musmus IGKV6-17*01 F | 100 | 287 | GACATTGTGATGACCCAGTCTCACAAATTCATGTCCACATCAGTAGGAGACAGGGTCAGCATCACCTGCAAGGCCAGTCAGGATGTGAGTACTGCTGTAGCCTGGTATCAACAGAAACCAGGACAATCTCCTAAACTACTGATTTACTCGGCATCCTACCGGTACACTGGAGTCCCTGATCGCTTCACTGGCAGTGGATCTGGGACGGATTTCACTTTCACCATCAGCAGTGTGCAGGCTGAAGACCTGGCAGTTTATTACTGTCAGCAACATTATAGTACTCCTCC |
| IGKV9-120*01_S2628_C3H | C3H/HeJ | Musmus IGKV9-120*02 F | 100 | 285 | GACATCCAGATGACCCAGTCTCCATCCTCCTTATCTGCCTCTCTGGGAGAAAGAGTCAGTCTCACTTGTCGGGCAAGTCAGGACATTGGTAGTAGCTTAAACTGGCTTCAGCAGGAACCAGATGGAACTATTAAACGCCTGATCTACGCCACATCCAGTTTAGATTCTGGTGTCCCCAAAAGGTTCAGTGGCAGTAGGTCTGGGTCAGATTATTCTCTCACCATCAGCAGCCTTGAGTCTGAAGATTTTGTAGACTATTACTGTCTACAATATGCTAGTTCTCCT |

| | | | | | |
|---|---|---|---|---|---|
| IGKV3-4*01_C3H | C3H/HeJ | Musmus IGKV3-4*01 F | 100 | 299 | GACATTGTGCTGACCCAATCTCCAGCTTCTTTGGCTGTGTCTCTAGGGCAGAGGGCCACCATCTCCTGCAAGGCCAGCCAAAGTGTTGATTATGATGGTGATAGTTATATGAACTGGTACCAACAGAAACCAGGACAGCCACCCAAACTCCTCATCTATGCTGCATCCAATCTAGAATCTGGGATCCCAGCCAGGTTTAGTGGCAGTGGGTCTGGGACAGACTTCACCCTCAACATCCATCCTGTGGAGGAGGAGGATGCTGCAACCTATTACTGTCAGCAAAGTAATGAGGATCCTCC |
| IGKV8-16*01_C3H | C3H/HeJ | Musmus IGKV8-16*01 F | 100 | 305 | GAAATTGTGTTGACCCAGTCTATACCATCCCTGACTGTGTCAGCAGGAGAGAGGGTCACTATCAGCTGCAAATCCAATCAGAATCTTTTATGGAGTGGAAACCAAAGGTACTGTTTGGTCTGGCACCAGTGGAAACCAGGGCAAACTCCTACACCGTTGATCACCTGGACATCTGATAGGTACTCTGGAGTCCCTGATCGTTTCATAGGCAGTGGATCTGTGACAGATTTCACTCTGACCATCAGCAGTGTGCAGGCTGAAGATGTGGCAGTTTATTTCTGTCAGCAGCATTTACACATTCCTCC |
| IGKV10-94*01_C3H | C3H/HeJ | Musmus IGKV10-94*01 F | 100 | 287 | GATATCCAGATGACACAGACTACATCCTCCCTGTCTGCCTCTCTGGGAGACAGAGTCACCATCAGTTGCAGTGCAAGTCAGGGCATTAGCAATTATTTAAACTGGTATCAGCAGAAACCAGATGGAACTGTTAAACTCCTGATCTATTACACATCAAGTTTACACTCAGGAGTCCCATCAAGGTTCAGTGGCAGTGGGTCTGGGACAGATTATTCTCTCACCATCAGCAACCTGGAACCTGAAGATATTGCCACTTACTATTGTCAGCAGTATAGTAAGCTTCCTCC |
| IGKV4-91*01_S5997_C3H | C3H/HeJ | Musmus IGKV4-91*01 F | 100 | 292 | GAAATTGTGCTCACCCAGTCTCCAACCACCATGGCTGCATCTCCCGGGGAGAAGATCACTATCACCTGCAGTGCCAGCTCAAGTATAAGTTCCAATTACTTGCATTGGTATCAGCAGAAGCCAGGATTCTCCCCTAAACTCTTGATTTATAGGACATCCAATCTGGCTTCTGGAGTCCCAGCTCGCTTCAGTGGCAGTGGGTCTGGGACCTCTTACTCTCTCACAATTGGCACCATGGAGGCTGAAGATGTTGCCACTTACTACTGCCAGCAGGGTAGTAGTATACCACTCA |
| IGKV3-12*01_C3H | C3H/HeJ | Musmus IGKV3-12*01 F | 100 | 299 | GACATTGTGCTGACACAGTCTCCTGCTTCCTTAGCTGTATCTCTGGGGCAGAGGGCCACCATCTCATGCAGGGCCAGCAAAAGTGTCAGTACATCTGGCTATAGTTATATGCACTGGTACCAACAGAAACCAGGACAGCCACCCAAACTCCTCATCTATCTTGCATCCAACCTAGAATCTGGGGTCCCTGCCAGGTTCAGTGGCAGTGGGTCTGGGACAGACTTCACCCTCAACATCCATCCTGTGGAGGAGGAGGATGCTGCAACCTATTACTGTCAGCACAGTAGGGAGCTTCCTCC |
| IGKV5-48*01_C3H | C3H/HeJ | Musmus IGKV5-48*01 F | 100 | 287 | GACATCTTGCTGACTCAGTCTCCAGCCATCCTGTCTGTGAGTCCAGGAGAAAGAGTCAGTTTCTCCTGCAGGGCCAGTCAGAGCATTGGCACAAGCATACACTGGTATCAGCAAAGAACAAATGGTTCTCCAAGGCTTCTCATAAAGTATGCTTCTGAGTCTATCTCTGGGATCCCTTCCAGGTTTAGTGGCAGTGGATCAGGGACAGATTTTACTCTTAGCATCAACAGTGTGGAGTCTGAAGATATTGCAGATTATTACTGTCAACAAAGTAATAGCTGGCCAAC |
| IGKV5-39*01_C3H | C3H/HeJ | Musmus IGKV5-39*01 F | 100 | 287 | GACATTGTGATGACTCAGTCTCCAGCCACCCTGTCTGTGACTCCAGGAGATAGAGTCTCTCTTTCCTGCAGGGCCAGCCAGAGTATTAGCGACTACTTACACTGGTATCAACAAAAATCACATGAGTCTCCAAGGCTTCTCATCAAATATGCTTCCCAATCCATCTCTGGGATCCCCTCCAGGTTCAGTGGCAGTGGATCAGGGTCAGATTTCACTCTCAGTATCAACAGTGTGGAACCTGAAGATGTTGGAGTGTATTACTGTCAAAATGGTCACAGCTTTCCTCC |
| IGKV6-25*01_C3H | C3H/HeJ | Musmus IGKV6-25*01 F | 100 | 287 | GACATTGTGATGACCCAGTCTCACAAATTCATGTCCACATCAGTAGGAGACAGGGTCAGCATCACCTGCAAGGCCAGTCAGGATGTGAGTACTGCTGTAGCCTGGTATCAACAAAAACCAGGGCAATCTCCTAAACTACTGATTTACTGGGCATCCACCCGGCACACTGGAGTCCCTGATCGCTTCACAGGCAGTGGATCTGGGACAGATTATACTCTCACCATCAGCAGTGTGCAGGCTGAAGACCTGGCACTTTATTACTGTCAGCAACATTATAGCACTCCTCC |

| | | | | | |
|---|---|---|---|---|---|
| IGKV12-89*01_C3H | C3H/HeJ | Musmus IGKV12-89*01 F | 100 | 287 | GACATCCAGATGACTCAGTCTCCAGCTTCACTGTCTGCATCTGTGGGAGAAACTGTCACCATCACATGTGGAGCAAGTGAGAATATTTACGGTGCTTTAAATTGGTATCAGCGGAAACAGGGAAAATCTCCTCAGCTCCTGATCTATGGTGCAACCAACTTGGCAGATGGCATGTCATCGAGGTTCAGTGGCAGTGGATCTGGTAGACAGTATTCTCTCAAGATCAGTAGCCTGCATCCTGACGATGTTGCAACGTATTACTGTCAAAATGTGTTAAGTACTCCTCC |
| IGKV4-55*01_S6279_C3H | C3H/HeJ | Musmus IGKV4-55*01 F | 100 | 286 | CAAATTGTTCTCACCCAGTCTCCAGCAATCATGTCTGCATCTCCAGGGGAGAAGGTCACCATGACCTGCAGTGCCAGCTCAAGTGTAAGTTACATGTACTGGTACCAGCAGAAGCCAGGATCCTCCCCCAGACTCCTGATTTATGACACATCCAACCTGGCTTCTGGAGTCCCTGTTCGCTTCAGTGGCAGTGGGTCTGGGACCTCTTACTCTCTCACAATCAGCCGAATGGAGGCTGAAGATGCTGCCACTTATTACTGCCAGCAGTGGAGTAGTTACCCACTCA |
| IGKV13-84*01_C3H | C3H/HeJ | Musmus IGKV13-84*01 F | 100 | 287 | GACATCCAGATGACACAATCTTCATCCTCCTTTTCTGTATCTCTAGGAGACAGAGTCACCATTACTTGCAAGGCAAGTGAGGACATATATAATCGGTTAGCCTGGTATCAGCAGAAACCAGGAAATGCTCCTAGGCTCTTAATATCTGGTGCAACCAGTTTGGAAACTGGGGTTCCTTCAAGATTCAGTGGCAGTGGATCTGGAAAGGATTACACTCTCAGCATTACCAGTCTTCAGACTGAAGATGTTGCTACTTATTACTGTCAACAGTATTGGAGTACTCCTCC |
| IGKV4-68*01_C3H | C3H/HeJ | Musmus IGKV4-68*01 F | 100 | 286 | CAAATTGTTCTCACCCAGTCTCCAGCACTCATGTCTGCATCTCCAGGGGAGAAGGTCACCATGACCTGCAGTGCCAGCTCAAGTGTAAGTTACATGTACTGGTACCAGCAGAAGCCAAGATCCTCCCCCAAACCCTGGATTTATCTCACATCCAACCTGGCTTCTGGAGTCCCTGCTCGCTTCAGTGGCAGTGGGTCTGGGACCTCTTACTCTCTCACAATCAGCAGCATGGAGGCTGAAGATGCTGCCACTTATTACTGCCAGCAGTGGAGTAGTAACCCACCCA |
| IGKV17-121*01_C3H | C3H/HeJ | Musmus IGKV17-121*01 F | 100 | 287 | GAAACAACTGTGACCCAGTCTCCAGCATCCCTGTCCATGGCTATAGGAGAAAAAGTCACCATCAGATGCATAACCAGCACTGATATTGATGATGATATGAACTGGTACCAGCAGAAGCCAGGGGAACCTCCTAAGCTCCTTATTTCAGAAGGCAATACTCTTCGTCCTGGAGTCCCATCCCGATTCTCCAGCAGTGGCTATGGTACAGATTTTGTTTTTACAATTGAAAACATGCTCTCAGAAGATGTTGCAGATTACTACTGTTTGCAAAGTGATAACTTGCCTCT |
| IGKV4-72*01_C3H | C3H/HeJ | Musmus IGKV4-72*01 F | 100 | 286 | CAAATTGTTCTCTCCCAGTCTCCAGCAATCCTGTCTGCATCTCCAGGGGAGAAGGTCACAATGACTTGCAGGGCCAGCTCAAGTGTAAGTTACATGCACTGGTACCAGCAGAAGCCAGGATCCTCCCCCAAACCCTGGATTTATGCCACATCCAACCTGGCTTCTGGAGTCCCTGCTCGCTTCAGTGGCAGTGGGTCTGGGACCTCTTACTCTCTCACAATCAGCAGAGTGGAGGCTGAAGATGCTGCCACTTATTACTGCCAGCAGTGGAGTAGTAACCCACCCA |
| IGKV13-85*01_C3H | C3H/HeJ | Musmus IGKV13-85*01 F | 100 | 287 | GACATCCAGATGACACAATCTTCATCCTACTTGTCTGTATCTCTAGGAGGCAGAGTCACCATTACTTGCAAGGCAAGTGACCACATTAATAATTGGTTAGCCTGGTATCAGCAGAAACCAGGAAATGCTCCTAGGCTCTTAATATCTGGTGCAACCAGTTTGGAAACTGGGGTTCCTTCAAGATTCAGTGGCAGTGGATCTGGAAAGGATTACACTCTCAGCATTACCAGTCTTCAGACTGAAGATGTTGCTACTTATTACTGTCAACAGTATTGGAGTACTCCTCC |
| IGKV14-100*01_C3H | C3H/HeJ | Musmus IGKV14-100*01 F | 100 | 287 | GACATCCTGATGACCCAATCTCCATCCTCCATGTCTGTATCTCTGGGAGACACAGTCAGCATCACTTGCCATGCAAGTCAGGGCATTAGCAGTAATATAGGGTGGTTGCAGCAGAAACCAGGGAAATCATTTAAGGGCCTGATCTATCATGGAACCAACTTGGAAGATGGAGTTCCATCAAGGTTCAGTGGCAGTGGATCTGGAGCAGATTATTCTCTCACCATCAGCAGCCTGGAATCTGAAGATTTTGCAGACTATTACTGTGTACAGTATGCTCAGTTTCCTCC |

| | | | | | |
|---|---|---|---|---|---|
| IGKV17-127*01_C3H | C3H/HeJ | Musmus IGKV17-127*01 F | 100 | 287 | GAAACAACTGTGACCCAGTCTCCAGCATCCCTGTCCGTGGCTACAGGAGAAAAAGTCACTATCAGATGCATAACCAGCACTGATATTGATGATGATATGAACTGGTACCAGCAGAAGCCAGGGGAACCTCCTAAGCTCCTTATTTCAGAAGGCAATACTCTTCGTCCTGGAGTCCCATCCCGATTCTCCAGCAGTGGCTATGGCACAGATTTTGTTTTTACAATTGAAAACACGCTCTCAGAAGATGTTGCAGATTACTACTGTTTGCAAAGTGATAACATGCCTCT |
| IGKV14-111*01_C3H | C3H/HeJ | Musmus IGKV14-111*01 F | 100 | 287 | GACATCAAGATGACCCAGTCTCCATCTTCCATGTATGCATCTCTAGGAGAGAGAGTCACTATCACTTGCAAGGCGAGTCAGGACATTAATAGCTATTTAAGCTGGTTCCAGCAGAAACCAGGGAAATCTCCTAAGACCCTGATCTATCGTGCAAACAGATTGGTAGATGGGGTCCCATCAAGGTTCAGTGGCAGTGGATCTGGGCAAGATTATTCTCTCACCATCAGCAGCCTGGAGTATGAAGATATGGGAATTTATTATTGTCTACAGTATGATGAGTTTCCTCC |
| IGKV12-98*01_C3H | C3H/HeJ | Musmus IGKV12-98*01 F | 100 | 287 | GACATTCAGATGACCCAGTCTCCTGCCTCCCAGTCTGCATCTCTGGGAGAAAGTGTCACCATCACATGCCTGGCAAGTCAGACCATTGGTACATGGTTAGCATGGTATCAGCAGAAACCAGGGAAATCTCCTCAGCTCCTGATTTATGCTGCAACCAGCTTGGCAGATGGGGTCCCATCAAGGTTCAGTGGTAGTGGATCTGGCACAAAATTTTCTTTCAAGATCAGCAGCCTACAGGCTGAAGATTTTGTAAGTTATTACTGTCAACAACTTTACAGTACTCCTCT |
| IGKV2-109*01_C3H | C3H/HeJ | Musmus IGKV2-109*01 F | 100 | 302 | GATATTGTGATGACGCAGGCTGCATTCTCCAATCCAGTCACTCTTGGAACATCAGCTTCCATCTCCTGCAGGTCTAGTAAGAGTCTCCTACATAGTAATGGCATCACTTATTTGTATTGGTATCTGCAGAAGCCAGGCCAGTCTCCTCAGCTCCTGATTTATCAGATGTCCAACCTTGCCTCAGGAGTCCCAGACAGGTTCAGTAGCAGTGGGTCAGGAACTGATTTCACACTGAGAATCAGCAGAGTGGAGGCTGAGGATGTGGGTGTTTATTACTGTGCTCAAAATCTAGAACTTCCTCC |
| IGKV8-34*01_C3H | C3H/HeJ | Musmus IGKV8-34*01 F | 100 | 305 | GACATTTTGATGACCCAGTCTCCATCCTCCCTGACTGTGTCAGCAGGAGAGAAGGTCACTATGAGCTGCAAGTCCAGTCAGAGTCTTTTAGCTAGTGGCAACCAAAATAACTACTTGGCCTGGCACCAGCAGAAACCAGGACGATCTCCTAAAATGCTGATAATTTGGGCATCCACTAGGGTATCTGGAGTCCCTGATCGCTTCATAGGCAGTGGATCTGGGACGGATTTCACTCTGACCATCAACAGTGTGCAGGCTGAAGATCTGGCTGTTTATTACTGTCAGCAGTCCTACAGCGCTCCTAC |
| IGKV16-104*01_C3H | C3H/HeJ | Musmus IGKV16-104*01 F | 100 | 288 | GATGTCCAGATAACCCAGTCTCCATCTTATCTTGCTGCATCTCCTGGAGAAACCATTACTATTAATTGCAGGGCAAGTAAGAGCATTAGCAAATATTTAGCCTGGTATCAAGAGAAACCTGGGAAAACTAATAAGCTTCTTATCTACTCTGGATCCACTTTGCAATCTGGAATTCCATCAAGGTTCAGTGGCAGTGGATCTGGTACAGATTTCACTCTCACCATCAGTAGCCTGGAGCCTGAAGATTTTGCAATGTATTACTGTCAACAGCATAATGAATACCCGTAC |
| IGKV1-122*01_C3H | C3H/HeJ | Musmus IGKV1-122*01 F | 100 | 302 | GATGCTGTGATGACCCAAACTCCACTCTCCCTGCCTGTCAGTCTTGGAGATCAAGCCTCCATCTCTTGCAGGTCTAGTCAGAGCCTTGAAAACAGTAATGGAAACACCTATTTGAACTGGTACCTCCAGAAACCAGGCCAGTCTCCACAGCTCCTGATCTACAGGGTTTCCAACCGATTTTCTGGGGTCCTAGACAGGTTCAGTGGTAGTGGATCAGGGACAGATTTCACACTGAAAATCAGCAGAGTGGAGGCTGAGGATTTGGGAGTTTATTTCTGCCTCCAAGTTACACATGTCCCTCC |
| IGKV4-63*01_C3H | C3H/HeJ | Musmus IGKV4-63*01 F | 100 | 286 | GAAAATGTTCTCACCCAGTCTCCAGCAATCATGTCTGCATCTCCAGGGGAAAAGGTCACCATGACCTGCAGTGCCAGCTCAAGTGTAAGTTACATGCACTGGTACCAGCAGAAGTCAAGCACCTCCCCCAAACTCTGGATTTATGACACATCCAAACTGGCTTCTGGAGTCCCAGGTCGCTTCAGTGGCAGTGGGTCTGGAAACTCTTACTCTCTCACGATCAGCAGCATGGAGGCTGAAGATGTTGCCACTTATTACTGTTTTCAGGGGAGTGGGTACCCACTCA |

| | | | | | |
|---|---|---|---|---|---|
| IGKV4-74*01_S2922_C3H | C3H/HeJ | Musmus IGKV4-74*01 F | 100 | 292 | CAAATTGTTCTCACCCAGTCTCCAGCAATCATGTCTGCATCTCTAGGGGAACGGGTCACCATGACCTGCACTGCCAGCTCAAGTGTAAGTTCCAGTTACTTGCACTGGTACCAGCAGAAGCCAGGATCCTCCCCCAAACTCTGGATTTATAGCACATCCAACCTGGCTTCTGGAGTCCCAGCTCGCTTCAGTGGCAGTGGGTCTGGGACCTCTTACTCTCTCACAATCAGCAGCATGGAGGCTGAAGATGCTGCCACTTATTACTGCCACCAGTATCATCGTTCCCCACTCA |
| IGKV9-124*01_C3H | C3H/HeJ | Musmus IGKV9-124*01 F | 100 | 285 | GACATCCAGATGACCCAGTCTCCATCCTCCTTATCTGCCTCTCTGGGAGAAAGAGTCAGTCTCACTTGTCGGGCAAGTCAGGAAATTAGTGGTTACTTAAGCTGGCTTCAGCAGAAACCAGATGGAACTATTAAACGCCTGATCTACGCCGCATCCACTTTAGATTCTGGTGTCCCAAAAAGGTTCAGTGGCAGTAGGTCTGGGTCAGATTATTCTCTCACCATCAGCAGCCTTGAGTCTGAAGATTTTGCAGACTATTACTGTCTACAATATGCTAGTTATCCT |
| IGKV3-10*01_C3H | C3H/HeJ | Musmus IGKV3-10*01 F | 100 | 299 | AACATTGTGCTGACCCAATCTCCAGCTTCTTTGGCTGTGTCTCTAGGGCAGAGGGCCACCATATCCTGCAGAGCCAGTGAAAGTGTTGATAGTTATGGCAATAGTTTTATGCACTGGTACCAGCAGAAACCAGGACAGCCACCCAAACTCCTCATCTATCTTGCATCCAACCTAGAATCTGGGGTCCCTGCCAGGTTCAGTGGCAGTGGGTCTAGGACAGACTTCACCCTCACCATTGATCCTGTGGAGGCTGATGATGCTGCAACCTATTACTGTCAGCAAAATAATGAGGATCCTCC |
| IGKV4-61*01_S2955_C3H | C3H/HeJ | Musmus IGKV4-61*01 F | 100 | 286 | CAAATTGTTCTCACCCAGTCTCCAGCAATCATGTCTGCATCTCCAGGGGAGAAGGTCACCATATCCTGCAGTGCCAGCTCAAGTGTAAGTTACATGTACTGGTACCAGCAGAAGCCAGGATCCTCCCCCAAACCCTGGATTTATCGCACATCCAACCTGGCTTCTGGAGTCCCTGCTCGCTTCAGTGGCAGTGGGTCTGGGACCTCTTACTCTCTCACAATCAGCAGCATGGAGGCTGAAGATGCTGCCACTTATTACTGCCAGCAGTATCATAGTTACCCACTCA |
| IGKV4-79*01_C3H | C3H/HeJ | Musmus IGKV4-79*01 F | 100 | 292 | CAAATTGTTCTCACCCAGTCTCCAGCAATCATGTCTGCATCTCCTGGGGAGAAGGTCACCTTGACCTGCAGTGCCAGCTCAAGTGTAAGTTCCAGCTACTTGTACTGGTACCAGCAGAAGCCAGGATCCTCCCCCAAACTCTGGATTTATAGCACATCCAACCTGGCTTCTGGAGTCCCTGCTCGCTTCAGTGGCAGTGGGTCTGGGACCTCTTACTCTCTCACAATCAGCAGCATGGAGGCTGAAGATGCTGCCTCTTATTTCTGCCATCAGTGGAGTAGTTACCCACCCC |
| IGKV1-133*01_C3H | C3H/HeJ | Musmus IGKV1-133*01 F | 100 | 302 | GATGTTGTGATGACCCAGACTCCACTCACTTTGTCGGTTACCATTGGACAACCAGCCTCTATCTCTTGCAAGTCAAGTCAGAGCCTCTTATATAGTAATGGAAAAACCTATTTGAATTGGTTATTACAGAGGCCAGGCCAGTCTCCAAAGCGCCTAATCTATCTGGTGTCTAAACTGGACTCTGGAGTCCCTGACAGGTTCACTGGCAGTGGATCAGGAACAGATTTTACACTGAAAATCAGCAGAGTGGAGGCTGAGGATTTGGGAGTTTATTACTGCGTGCAAGGTACACATTTTCCTCA |
| IGKV4-50*01_C3H | C3H/HeJ | Musmus IGKV4-50*01 F | 100 | 286 | GAAAATGTGCTCACCCAGTCTCCAGCAATCATGTCTGCATCTCTAGGGGAGAAGGTCACCATGAGCTGCAGGGCCAGCTCAAGTGTAAATTACATGTACTGGTACCAGCAGAAGTCAGATGCCTCCCCCAAACTATGGATTTATTACACATCCAACCTGGCTCCTGGAGTCCCAGCTCGCTTCAGTGGCAGTGGGTCTGGGAACTCTTATTCTCTCACAATCAGCAGCATGGAGGGTGAAGATGCTGCCACTTATTACTGCCAGCAGTTTACTAGTTCCCCATCCA |
| IGKV5-45*01_C3H | C3H/HeJ | Musmus IGKV5-45*01 F | 100 | 287 | GATATTGTGCTAACTCAGTCTCCAGCCACCCTGTCTGTGACTCCAGGAGATAGAGTCAGTCTTTCCTGCAGGGCCAGTCAAAGTATTAGCAACTACCTACACTGGTATCAACAAAATCACATGAGTCTCCAAGGCTTCTCATCAAGTATGCTTCCCAGTCCATCTCTGGGATCCCCTCCAGGTTCAGTGGCAGTGGATCAGGGACAGATTTCACTCTCAGTATCAACAGTGTGGAGACTGAAGATTTTGGAATGTATTTCTGTCAACAGAGTAACAGCTGGCCTCA |

| | | | | | |
|---|---|---|---|---|---|
| IGKV2-112*01_C3H | C3H/HeJ | Musmus IGKV2-112*01 F | 100 | 300 | GATATTGTGATAACCCAGGATGAACTCTCCAATCCTGTCACTTCTGGAGAATCAGTTTCCATCTCCTGCAGGTCTAGTAAGAGTCTCCTATATAAGGATGGGAAGACATACTTGAATTGGTTTCTGCAGAGACCAGGACAATCTCCTCAGCTCCTGATCTATTTGATGTCCACCCGTGCATCAGGAGTCTCAGACCGGTTTAGTGGCAGTGGGTCAGGAACAGATTTCACCCTGGAAATCAGTAGAGTGAAGGCTGAGGATGTGGGTGTGTATTACTGTCAACAACTTGTAGAGTATCCT |
| IGKV9-123*01_C3H | C3H/HeJ | Musmus IGKV9-123*01 F | 100 | 285 | GACATCCAGATGATTCAGTCTCCATCGTCCATGTTTGCCTCTCTGGGAGACAGAGTCAGTCTCTCTTGTCGGGCTAGTCAGGGCATTAGAGGTAATTTAGACTGGTATCAGCAGAAACCAGGTGGAACTATTAAACTCCTGATCTACTCCACATCCAATTTAAATTCTGGTGTCCCATCAAGGTTCAGTGGCAGTGGGTCTGGGTCAGATTATTCTCTCACCATCAGCAGCCTAGAGTCTGAAGATTTTGCAGACTATTACTGTCTACAGCGTAATGCGTATCCT |
| IGKV11-125*01_C3H | C3H/HeJ | Musmus IGKV11-125*01 F | 100 | 287 | GATGTCCAGATGATTCAGTCTCCATCCTCCCTGTCTGCATCTTTGGGAGACATAGTCACCATGACTTGCCAGGCAAGTCAGGGCACTAGCATTAATTTAAACTGGTTTCAGCAAAAACCAGGGAAAGCTCCTAAGCTCCTGATCTATGGTGCAAGCAACTTGGAAGATGGGGTCCCATCAAGGTTCAGTGGCAGTAGATATGGGACAGATTTCACTCTCACCATCAGCAGCCTGGAGGATGAAGATATGGCAACTTATTTCTGTCTACAGCATAGTTATCTCCCTCC |
| IGKV4-58*01_C3H | C3H/HeJ | Musmus IGKV4-58*01 F | 100 | 292 | GAAAATGTGCTCACCCAGTCTCCAGCAATAATGGCTGCCTCTCTGGGGCAGAAGGTCACCATGACCTGCAGTGCCAGCTCAAGTGTAAGTTCCAGTTACTTGCACTGGTACCAGCAGAAGTCAGGCGCTTCCCCCAAACCCTTGATTCATAGGACATCCAACCTGGCTTCTGGAGTCCCAGCTCGCTTCAGTGGCAGTGGGTCTGGGACCTCTTACTCTCTCACAATCAGCAGCGTGGAGGCTGAAGATGATGCAACTTATTACTGCCAGCAGTGGAGTGGTTACCCATTCA |
| IGKV4-86*01_C3H | C3H/HeJ | Musmus IGKV4-86*01 F | 100 | 283 | GAAATTGTGCTCACTCAGTCTCCAGCCATCACAGCTGCATCTCTGGGGCAAAAGGTCACCATCACCTGCAGTGCCAGCTCAAGTGTAAGTTACATGCACTGGTACCAGCAGAAGTCAGGCACCTCCCCCAAACCATGGATTTATGAAATATCCAAACTGGCTTCTGGAGTCCCAGCTCGCTTCAGTGGCAGTGGGTCTGGGACCTCTTACTCTCTCACAATCAGCAGCATGGAGGCTGAAGATGCTGCCATTTATTACTGCCAGCAGTGGAATTATCCTCTTA |
| IGKV8-26*01_C3H | C3H/HeJ | Musmus IGKV8-26*01 ORF | 100 | 311 | GACATTGTGATGACCCAGTCTCCATCCTCCCTGGCTGTGACAGCAGGAGAGAAGGTCACTATGAGCTGCAAGTCCAGTCAGAGTCTTTTGTGGAGTGTAAACCAAAAGAACTACTTGTCCTGGTACCAGCAGAAACAAAGGCAGCCTCCTAAACTGCTTATCTATGGGGCATCCATTAGAGAATCTTGGGTCCCTGATCGGTTCACAGGAAGTGGATCTGGGACAGACTTCACTCTCACCATTAGCAATGTGCATGCTGAAGACCTAGCAGTTTATTACTGTCAACACAATCATGGCAGCTTTCTCCCCCC |
| IGKV4-68*01_S3092_C3H | C3H/HeJ | Musmus IGKV4-68*01 F | 100 | 286 | CAAATTGTTCTCACCCAGTCTCCAGCACTCATGTCTGCATCTCCAGGGGAGAAGGTCACCATGACCTGCAGTGCCAGCTCAAGTGTAAGTTACATGTACTGGTACCAGCAGAAGCCAAGATCCTCCCCCAAACCCTGGATTTATCTCACATCCAACCTGGCTTCTGGAGTCCCTGCTCGCTTCAGTGGCAGTGGGTCTGGGACCTCTTACTCTCTCACAATCAGCAGCATGGAGGCTGAAGATGCTGCCACTTATTACTGCCAGCAGTGGAGTAGTAACCCACTCA |
| IGKV4-69*01_C3H | C3H/HeJ | Musmus IGKV4-69*01 F | 100 | 282 | CAAATTCTTCTCACCCAGTCTCCAGCAATCATGTCTGCATCTCCAGGGGAGAAGGTCACCATGACCTGCAGTGCCAGCTCAAGTGTAAGTTACATGCACTGGTACCAGCAGAAGCCAGGATCCTCGCCCAAACCCTGGATTTATGACACATCCAACCTGGCTTCTGGATTCCCTGCTCGCTTCAGTGGCAGTGGGTCTGGGACCTCTTACTCTCTCATAATCAGCAGCATGGAGGCTGAAGATGCTGCCACTTATTACTGCCATCAGCGGAGTAGTTACCCA |

| | | | | | |
|---|---|---|---|---|---|
| IGKV14-130*01_C3H | C3H/HeJ | Musmus IGKV14-130*01 F | 100 | 287 | GAAATCCAGATGACCCAGTCTCCATCCTCTATGTCTGCATCTCTGGGAGACAGAATAACCATCACTTGCCAGGCAACTCAAGACATTGTTAAGAATTTAAACTGGTATCAGCAGAAACCAGGGAAACCCCCTTCATTCCTGATCTATTATGCAACTGAACTGGCAGAAGGGGTCCCATCAAGGTTCAGTGGCAGTGGGTCTGGGTCAGACTATTCTCTGACAATCAGCAACCTGGAGTCTGAAGATTTTGCAGACTATTACTGTCTACAGTTTTATGAGTTTCCTCC |
| IGKV3-1*01_C3H | C3H/HeJ | Musmus IGKV3-1*01 F | 100 | 299 | GACATTGTGCTCACCCAATCTCCAGCTTCTTTGGCTGTGTCTCTAGGGCAGAGAGCCACCATCTCCTGCAGAGCCAGTGAAAGTGTTGAATATTATGGCACAAGTTTAATGCAGTGGTACCAACAGAAACCAGGACAGCCACCCAAACTCCTCATCTATGCTGCATCCAACGTAGAATCTGGGGTCCCTGCCAGGTTTAGTGGCAGTGGGTCTGGGACAGACTTCAGCCTCAACATCCATCCTGTGGAGGAGGATGATATTGCAATGTATTTCTGTCAGCAAAGTAGGAAGGTTCCTTC |
| IGKV4-92*01_C3H | C3H/HeJ | Musmus IGKV4-92*01 F | 100 | 291 | GAAATGGTTCTCACCCAGTCTCCAGTATCCATAACTGCATCTCGAGGGGAGAAGGTCACCATCACCTGCCGTGCCAGCTCAAGTATAAGTTCCAATTACTTACACTGGTACCAGCAGAAGCCAGGATCCTCCCCTAAACTTTTGATTTATAGGACATCCATCCTGGCATCTGGAGTCCTAGACAGCTTCAGTGGCAGTGGGTCTGAGAGCTCTTACACTCTGACAATCAGCTGCATGCAGGACGAAGTTGCTGCCACTTACTATTGTCAGCAGGGGAGTAGTAGCCCACCA |
| IGKV10-95*01_C3H | C3H/HeJ | Musmus IGKV10-95*01 F | 100 | 287 | GATATCCAGATGACACAGACTACTTCCTCCCTGTCTGCCTCTCTGGGAGACAGAGTCACCATCAGTTGCAGGGCAAGTGAGGACATTAGCACTTATTTAAACTGGTATCAGCAGAAACCAGATGGAACTGTTAAACTCCTGATCTATTACACATCAGGATTACACTCAGGAGTCCCATCAAGGTTCAGTGGCAGTGGGTCTGGGGCAGATTATTCTCTCACCATCAGCAACCTGGAACCTGAAGATATTGCCACTTACTATTGTCAGCAGTATAGTAAGCTTCCTCC |
| IGKV5-37*01_C3H | C3H/HeJ | Musmus IGKV5-37*01 F | 100 | 287 | GACATCCTGCTGACCCAGTCTCCAGCCACCCTGTCTGTGACTCCAGGAGAAACAGTCAGTCTTTCCTGTAGGGCCAGCCAGAGTATTTACAAGAACCTACACTGGTATCAACAGAAATCACATCGGTCTCCAAGGCTTCTCATCAAGTATGCTTCTGATTCCATCTCTGGGATCCCCTCCAGGTTCACTGGCAGTGGATCAGGGACAGATTACACTCTCAGTATCAACAGTGTGAAGCCCGAAGATGAAGGAATATATTACTGTCTTCAAGGTTACAGCACACCTTC |
| IGKV8-23-1*01_C3H | C3H/HeJ | Musmus IGKV8-16*01 F, or Musmus IGKV8-23-1*01 ORF | 100 | 284 | GAAATTGTGCTGACTGTGTCAGCAGGAGAGAGGGTCACTATCAGCTGCAAATCCAATCAGAATCTTTTATGGAGTGGAAACCAAAGGTACTGTTTGGTCTGGCACCAGTGGAAACCAGGGCAAACTCCTACACCGTTGATCACCTGGACATCTGATAGGTACTCTGGAGTCCCTGATCGTTTCATAGGCAGTGGATCTGTGACAGATTTCACTCTGACCATCAGCAGTGTGCAGGCTGAAGATGTGGCAGTTTATTTCTGTCAGCAGCATTTACACATTCCTCC |
| IGKV1-88*01_C3H | C3H/HeJ | Musmus IGKV1-88*01 F | 100 | 302 | GATGTTGTGGTGACTCAAACTCCACTCTCCCTGCCTGTCAGCTTTGGAGATCAAGTTTCTATCTCTTGCAGGTCTAGTCAGAGTCTTGCAAACAGTTATGGGAACACCTATTTGTCTTGGTACCTGCACAAGCCTGGCCAGTCTCCACAGCTCCTCATCTATGGGATTTCCAACAGATTTTCTGGGGTGCCAGACAGGTTCAGTGGCAGTGGTTCAGGGACAGATTTCACACTCAAGATCAGCACAATAAAGCCTGAGGACTTGGGAATGTATTACTGCTTACAAGGTACACATCAGCCTCC |
| IGKV4-50*01_S0901_C3H | C3H/HeJ | Musmus IGKV4-50*01 F | 100 | 286 | GAAAATGTGCTCACCCAGTCTCCAGCAATCATGTCTGCATCTCTAGGGGAGAAGGTCACCATGAGCTGCAGGGCCAGCTCAAGTGTAAATTACATGTACTGGTACCAGCAGAAGTCAGATGCCTCCCCCAAACTATGGATTTATTACACATCCAACCTGGCTCCTGGAGTCCCAGCTCGCTTCAGTGGCAGTGGGTCTGGGAACTCTTATTCTCTCACAATCAGCAGCATGGAGGGTGAAGATGCTGCCACTTATTACTGCCAGCAGTTTACTAGTTCCCCATACA |

| | | | | | |
|---|---|---|---|---|---|
| IGKV4-72*01_S3540_C3H | C3H/HeJ | Musmus IGKV4-72*01 F | 100 | 286 | CAAATTGTTCTCTCCCAGTCTCCAGCAATCCTGTCTGCATCTCCAGGGGAGAAGGTCACAATGACTTGCAGGGCCAGCTCAAGTGTAAGTTACATGCACTGGTACCAGCAGAAGCCAGGATCCTCCCCCAAACCCTGGATTTATGCCACATCCAACCTGGCTTCTGGAGTCCCTGCTCGCTTCAGTGGCAGTGGGTCTGGGACCTCTTACTCTCTCACAATCAGCAGAGTGGAGGCTGAAGATGCTGCCACTTATTACTGCCAGCAGTGGAGTAGTAACCCACTCA |
| IGKV6-14*01_C3H | C3H/HeJ | Musmus IGKV6-14*01 F | 100 | 287 | GACATTGTGATGACCCAGTCTCAAAAATTCATGTCCACATCAGTAGGAGACAGGGTCAGCATCACCTGCAAGGCCAGTCAGAATGTTCGTACTGCTGTAGCCTGGTATCAACAGAAACCAGGGCAGTCTCCTAAAGCACTGATTTACTTGGCATCCAACCGGCACACTGGAGTCCCTGATCGCTTCACAGGCAGTGGATCTGGGACAGATTTCACTCTCACCATTAGCAATGTGCAATCTGAAGACCTGGCAGATTATTTCTGTCTGCAACATTGGAATTATCCTCT |
| IGKV1-132*01_C3H | C3H/HeJ | Musmus IGKV1-132*01 F | 100 | 302 | GATGTTGTGATGACCCAGACTCCACTGTCTTTGTCGGTTACCATTGGACAACCAGCCTCTATCTCTTGCAAGTCAAGTCAGAGCCTCTTATATAGTAATGGAAAGACATATTTGAATTGGTTACAACAGAGGCCTGGCCAGGCTCCAAAGCACCTAATGTATCAGGTGTCCAAACTGGACCCTGGCATCCCTGACAGGTTCAGTGGCAGTGGATCAGAAACAGATTTTACACTTAAAATCAGCAGAGTGGAGGCTGAAGATTTGGGAGTTTATTACTGCTTGCAAGGTACATATTATCCTCA |
| IGKV12-38*01_C3H | C3H/HeJ | Musmus IGKV12-38*01 F | 100 | 287 | GACATCCAGATGACTCAGTCTCCAGCCTCCCTGGCTGCATCTGTGGGAGAAACTGTCACCATCACATGTCGAGCAAGTGAGAACATTTACTACAGTTTAGCATGGTATCAGCAGAAGCAAGGGAAATCTCCTCAGCTCCTGATCTATAATGCAAACAGCTTGGAAGATGGTGTCCCATCGAGGTTCAGTGGCAGTGGATCTGGGACACAGTATTCTATGAAGATCAACAGCATGCAGCCTGAAGATACCGCAACTTATTTCTGTAAACAGGCTTATGACGTTCCTCC |
| IGKV1-99*01_C3H | C3H/HeJ | Musmus IGKV1-99*01 F | 100 | 302 | GATGTTGTTCTGACCCAAACTCCACTCTCTCTGCCTGTCAATATTGGAGATCAAGCCTCTATCTCTTGCAAGTCTACTAAGAGTCTTCTGAATAGTGATGGATTCACTTATTTGGACTGGTACCTGCAGAAGCCAGGCCAGTCTCCACAGCTCCTAATATATTTGGTTTCTAATCGATTTTCTGGAGTTCCAGACAGGTTCAGTGGCAGTGGGTCAGGAACAGATTTCACACTCAAGATCAGCAGAGTGGAGGCTGAGGATTTGGGAGTTTATTATTGCTTCCAGAGTAACTATCTTCCTCT |
| IGKV4-51*01_C3H | C3H/HeJ | Musmus IGKV4-51*01 F | 100 | 292 | GAAAATGTGCTCACCCAGTCTCCAGCAATAATGGCTGCCTCTCTGGGGGAGAAGGTCACCATGACCTGCAGTGCCAGCTCAAGTGTAAGTTCCAGCTACTTGCACTGGTACCAGCAGAAGTCAGGCACTTCCCCCAAACTCTGGATTTATGGCACATCCAACCTGGCTTCTGGAGTCCCAGCTCGCTTCAGTGGCAGTGGGGCTGGGATCTCTTACTCTCTCACAATCAGCAGCATGGAGGCTGAAAATGATGCAACTTATTACTGCCAGCAGTGGAGTGGTTACCCATTCA |
| IGKV4-52*01_S5019_C3H | C3H/HeJ | Musmus IGKV4-52*01 ORF, or Musmus IGKV4-54*01 ORF | 100 | 286 | CAAATTGTTCTCACCCAGTCTCCAGCAATCCTGTCTGCATCTCCAGGGGAGAAGGTCACCATGACCTGCAGTGCCAGCTCAAGTGTAAGTTACATGTACAGGTACCAGCAGAAGCCAGGATCCTCACCCAAACCCTGGATTTATGGCACATCCAACCTGGCTTCTGGAGTCCCTGCTCGCTTCAGTGGCAGTGGATCTGGGACCTCTTATTCTCTCACAATCAGCAGCATGGAGGCTGAAGATGCTGCCACTTATTACTGCCAGCAATATCATAGTTACCCACTCA |
| IGKV4-60*01_S1112_C3H | C3H/HeJ | Musmus IGKV4-60*01 P | 100 | 292 | GAAATTGTGCTCACCCAGTCTCCAGCACTCATGGCTGCATCTCCAGGGGAGAAGGTCACCATCACCTGCAGTGTCAGCTCAAGTATAAGTTCCAGCTACTTGCACTGGTACCAGCAGAAGTCAGGAATCTCCCCCAAACCCTGGATTTATGGCACATCCAACCTGGCTTCTGGAGTCCCTGCTCGCTTCAGTGGCAGTGGATCTGGGACCTCTTACTCTCTCACAATCAGCAGCATGGAGGCTGAAGATGCTGCCACTTATTACTGTCAACAGTGGAGTAGTTCCCCACTCA |

| | | | | | |
|---|---|---|---|---|---|
| IGKV4-81*01_C3H | C3H/HeJ | Musmus IGKV4-81*01 F | 100 | 292 | GAAAATGTGCTGACCCAGTCTCCAGCAATCATGGCTGCATCTCCAGGGGAGAAGGTCACCATGACCTGCAGTGCCAGCTCAAGTGTAAGTTCTAGTAACTTGCACTGGTACCAGCAGAAGTCAGGCACTTCTACCAAATTCTGGATTTATAGGACATCCAACCTGGCTTCAGAAGTCCCAGCTCCCTTCAGTGGCAGTGGGTCTGGGACCTCTTACTCTCTTACAATCAGCAGCGTGGAGGCCGAAGATGCTGCCACTTATTACTGCCAGCAGTGGAGTGGTTACCCACCCA |
| IGKV4-90*01_C3H | C3H/HeJ | Musmus IGKV4-90*01 F | 100 | 285 | GAAATTTTGCTCACCCAGTCTCCAGCAATCATAGCTGCATCTCCTGGGGAGAAGGTCACCATCACCTGCAGTGCCAGCTCAAGTGTAAGTTACATGAACTGGTACCAGCAGAAACCAGGATCCTCCCCCAAAATATGGATTTATGGTATATCCAACCTGGCTTCTGGAGTTCCTGCTCGCTTCAGTGGCAGTGGGTCTGGGACATCTTTCTCTTTCACAATCAACAGCATGGAGGCTGAAGATGTTGCCACTTATTACTGTCAGCAAAGGAGTAGTTACCCACCC |
| IGKV6-29*01_C3H | C3H/HeJ | Musmus IGKV6-29*01 F | 100 | 287 | AACATTGTAATGACCCAATCTCCCAAATCCATGTCCATGTCAGTAGGAGAGAGGGTCACCTTGAGCTGCAAGGCCAGTGAGAATGTGGGTACTTATGTATCCTGGTATCAACAGAAACCAGAGCAGTCTCCTAAACTGCTGATATACGGGGCATCCAACCGGTACCCTGGGGTCCCTGATCGCTTCACAGGCAGTGGATCTGCAACAGATTTCACTCTGACCATCAGCAGTCTGCAGGCTGAAGACCTTGCAGATTATCACTGTGGACAGGGTTACAGCTATCTTCC |
| IGKV9-129*01_C3H | C3H/HeJ | Musmus IGKV9-129*01 ORF | 100 | 287 | GACATCCAGATGACCCAGTCTCCATCCTCCTTATCTGCCTCTCTGGGAGAAAGAGTCAGTCTCACTTGCCGGGCAAGTCAGGACATTCATGGTTATTTAAACTTGTTTCAGCAGAAACCAGGTGAAACTATTAAACACCTGATCTATGAAACATCCAATTTAGATTCTGGTGTCCCAAAAAGGTTCAGTGGCAGTAGGTCTGGGTCAGATTATTCTCTCATTATCGGCAGCCTTGAGTCTGAAGATTTTGCAGACTATTACTGTCTACAATATGCTAGTTCTCCTCC |
| IGKV4-78*01_C3H | C3H/HeJ | Musmus IGKV4-78*01 F | 100 | 292 | CAAATTGTTCTCACCCAGTCTCCAGCAATCATGTCTGCATCTCCTGGGGAGAAGGTCACCATGACCTGCAGTGCCAGATCAAGTGTAAGTTCCAGCTACTTGTACTGGTACCAGCAGAAGCCAGGATCCTCCCCCAAACTCTGGATTTATAGCACATCCAACCTGGCTTCTGGAGTCCCTGCTCGCTTCAGTGGCAGTGGGTCTGGGACCTCTTATTCTCTCACAATCAGCAGCATGGAGGCTGAAGATGCTGCCACTTTTTACTGCCAGCAGTACAGTGGTTACCCATCCA |
| IGKV6-15*01_B6 | C57BL/6J | Musmus IGKV6-15*01 F | 100 | 287 | GACATTGTGATGACCCAGTCTCAAAAATTCATGTCCACATCAGTAGGAGACAGGGTCAGCGTCACCTGCAAGGCCAGTCAGAATGTGGGTACTAATGTAGCCTGGTATCAACAGAAACCAGGGCAATCTCCTAAAGCACTGATTTACTCGGCATCCTACCGGTACAGTGGAGTCCCTGATCGCTTCACAGGCAGTGGATCTGGGACAGATTTCACTCTCACCATCAGCAATGTGCAGTCTGAAGACTTGGCAGAGTATTTCTGTCAGCAATATAACAGCTATCCTCT |
| IGKV8-30*01_B6 | C57BL/6J | Musmus IGKV8-30*01 F | 100 | 305 | GACATTGTGATGTCACAGTCTCCATCCTCCCTAGCTGTGTCAGTTGGAGAGAAGGTTACTATGAGCTGCAAGTCCAGTCAGAGCCTTTTATATAGTAGCAATCAAAAGAACTACTTGGCCTGGTACCAGCAGAAACCAGGGCAGTCTCCTAAACTGCTGATTTACTGGGCATCCACTAGGGAATCTGGGGTCCCTGATCGCTTCACAGGCAGTGGATCTGGGACAGATTTCACTCTCACCATCAGCAGTGTGAAGGCTGAAGACCTGGCAGTTTATTACTGTCAGCAATATTATAGCTATCCTCC |
| IGKV1-135*01_B6 | C57BL/6J | Musmus IGKV1-135*01 F | 100 | 302 | GATGTTGTGATGACCCAGACTCCACTCACTTTGTCGGTTACCATTGGACAACCAGCCTCCATCTCTTGCAAGTCAAGTCAGAGCCTCTTAGATAGTGATGGAAAGACATATTTGAATTGGTTGTTACAGAGGCCAGGCCAGTCTCCAAAGCGCCTAATCTATCTGGTGTCTAAACTGGACTCTGGAGTCCCTGACAGGTTCACTGGCAGTGGATCAGGGACAGATTTCACACTGAAAATCAGCAGAGTGGAGGCTGAGGATTTGGGAGTTTATTATTGCTGGCAAGGTACACATTTTCCTCA |

| | | | | | |
|---|---|---|---|---|---|
| IGKV8-27*01_B6 | C57BL/6J | Musmus IGKV8-27*01 F | 100 | 300 | AACATTATGATGACACAGTCGCCATCATCTCTGGCTGTGTCTGCAGGAGAAAAGGTCACTATGAGCTGTAAGTCCAGTCAAAGTGTTTTATACAGTTCAAATCAGAAGAACTACTTGGCCTGGTACCAGCAGAAACCAGGGCAGTCTCCTAAACTGCTGATCTACTGGGCATCCACTAGGGAATCTGGTGTCCCTGATCGCTTCACAGGCAGTGGATCTGGGACAGATTTTACTCTTACCATCAGCAGTGTACAAGCTGAAGACCTGGCAGTTTATTACTGTCATCAATACCTCTCCTCG |
| IGKV12-44*01_B6 | C57BL/6J | Musmus IGKV12-44*01 F | 100 | 287 | GACATCCAGATGACTCAGTCTCCAGCCTCCCTATCTGCATCTGTGGGAGAAACTGTCACCATCACATGTCGAGCAAGTGAGAATATTTACAGTTATTTAGCATGGTATCAGCAGAAACAGGGAAAATCTCCTCAGCTCCTGGTCTATAATGCAAAAACCTTAGCAGAAGGTGTGCCATCAAGGTTCAGTGGCAGTGGATCAGGCACACAGTTTTCTCTGAAGATCAACAGCCTGCAGCCTGAAGATTTTGGGAGTTATTACTGTCAACATCATTATGGTACTCCTCC |
| IGKV6-23*01_B6 | C57BL/6J | Musmus IGKV6-23*01 F | 100 | 287 | GACATTGTGATGACCCAGTCTCACAAATTCATGTCCACATCAGTAGGAGACAGGGTCAGCATCACCTGCAAGGCCAGTCAGGATGTGGGTACTGCTGTAGCCTGGTATCAACAGAAACCAGGGCAATCTCCTAAACTACTGATTTACTGGGCATCCACCCGGCACACTGGAGTCCCTGATCGCTTCACAGGCAGTGGATCTGGGACAGATTTCACTCTCACCATTAGCAATGTGCAGTCTGAAGACTTGGCAGATTATTTCTGTCAGCAATATAGCAGCTATCCTCT |
| IGKV10-96*01_B6 | C57BL/6J | Musmus IGKV10-96*01 F | 100 | 286 | GATATCCAGATGACACAGACTACATCCTCCCTGTCTGCCTCTCTGGGAGACAGAGTCACCATCAGTTGCAGGGCAAGTCAGGACATTAGCAATTATTTAAACTGGTATCAGCAGAAACCAGATGGAACTGTTAAACTCCTGATCTACTACACATCAAGATTACACTCAGGAGTCCCATCAAGGTTCAGTGGCAGTGGGTCTGGAACAGATTATTCTCTCACCATTAGCAACCTGGAGCAAGAAGATATTGCCACTTACTTTTGCCAACAGGGTAATACGCTTCCTC |
| IGKV12-46*01_B6 | C57BL/6J | Musmus IGKV12-46*01 F | 100 | 287 | GACATCCAGATGACTCAGTCTCCAGCCTCCCTATCTGTATCTGTGGGAGAAACTGTCACCATCACATGTCGAGCAAGTGAGAATATTTACAGTAATTTAGCATGGTATCAGCAGAAACAGGGAAAATCTCCTCAGCTCCTGGTCTATGCTGCAACAAACTTAGCAGATGGTGTGCCATCAAGGTTCAGTGGCAGTGGATCAGGCACACAGTATTCCCTCAAGATCAACAGCCTGCAGTCTGAAGATTTTGGGAGTTATTACTGTCAACATTTTTGGGGTACTCCTCC |
| IGKV2-137*01_B6 | C57BL/6J | Musmus IGKV2-137*01 F | 100 | 302 | GATATTGTGATGACTCAGGCTGCACCCTCTGTACCTGTCACTCCTGGAGAGTCAGTATCCATCTCCTGCAGGTCTAGTAAGAGTCTCCTGCATAGTAATGGCAACACTTACTTGTATTGGTTCCTGCAGAGGCCAGGCCAGTCTCCTCAGCTCCTGATATATCGGATGTCCAACCTTGCCTCAGGAGTCCCAGACAGGTTCAGTGGCAGTGGGTCAGGAACTGCTTTCACACTGAGAATCAGTAGAGTGGAGGCTGAGGATGTGGGTGTTTATTACTGTATGCAACATCTAGAATATCCTTT |
| IGKV6-20*01_B6 | C57BL/6J | Musmus IGKV6-20*01 F | 100 | 287 | AACATTGTAATGACCCAATCTCCCAAATCCATGTCCATGTCAGTAGGAGAGAGGGTCACCTTGAGCTGCAAGGCCAGTGAGAATGTGGGTACTTATGTATCCTGGTATCAACAGAAACCAGAGCAGTCTCCTAAACTGCTGATATACGGGGCATCCAACCGGTACACTGGGGTCCCCGATCGCTTCACAGGCAGTGGATCTGCAACAGATTTCACTCTGACCATCAGCAGTGTGCAGGCTGAAGACCTTGCAGATTATCACTGTGGACAGAGTTACAGCTATCCTCC |
| IGKV4-57*01_S6684_B6 | C57BL/6J | Musmus IGKV4-57-1*01 F | 100 | 286 | CAAATTGTTCTCACCCAGTCTCCAGCAATCATGTCTGCATCTCCAGGGGAGAAGGTCACCATAACCTGCAGTGCCAGCTCAAGTGTAAGTTACATGCACTGGTTCCAGCAGAAGCCAGGCACTTCTCCCAAACTCTGGATTTATAGCACATCCAACCTGGCTTCTGGAGTCCCTGCTCGCTTCAGTGGCAGTGGATCTGGGACCTCTTACTCTCTCACAATCAGCCGAATGGAGGCTGAAGATGCTGCCACTTATTACTGCCAGCAAAGGAGTAGTTACCCGCTCA |

| | | | | | |
|---|---|---|---|---|---|
| IGKV12-89*01_B6 | C57BL/6J | Musmus IGKV12-89*01 F | 100 | 287 | GACATCCAGATGACTCAGTCTCCAGCTTCACTGTCTGCATCTGTGGGAGAAACTGTCACCATCACATGTGGAGCAAGTGAGAATATTTACGGTGCTTTAAATTGGTATCAGCGGAAACAGGGAAAATCTCCTCAGCTCCTGATCTATGGTGCAACCAACTTGGCAGATGGCATGTCATCGAGGTTCAGTGGCAGTGGATCTGGTAGACAGTATTCTCTCAAGATCAGTAGCCTGCATCCTGACGATGTTGCAACGTATTACTGTCAAAATGTGTTAAGTACTCCTCC |
| IGKV8-21*01_B6 | C57BL/6J | Musmus IGKV8-21*01 F | 100 | 302 | GACATTGTGATGTCACAGTCTCCATCCTCCCTGGCTGTGTCAGCAGGAGAGAAGGTCACTATGAGCTGCAAATCCAGTCAGAGTCTGCTCAACAGTAGAACCCGAAAGAACTACTTGGCTTGGTACCAGCAGAAACCAGGGCAGTCTCCTAAACTGCTGATCTACTGGGCATCCACTAGGGAATCTGGGGTCCCTGATCGCTTCACAGGCAGTGGATCTGGGACAGATTTCACTCTCACCATCAGCAGTGTGCAGGCTGAAGACCTGGCAGTTTATTACTGCAAGCAATCTTATAATCTTCC |
| IGKV1-117*01_B6 | C57BL/6J | Musmus IGKV1-117*01 F | 100 | 302 | GATGTTTTGATGACCCAAACTCCACTCTCCCTGCCTGTCAGTCTTGGAGATCAAGCCTCCATCTCTTGCAGATCTAGTCAGAGCATTGTACATAGTAATGGAAACACCTATTTAGAATGGTACCTGCAGAAACCAGGCCAGTCTCCAAAGCTCCTGATCTACAAAGTTTCCAACCGATTTTCTGGGGTCCCAGACAGGTTCAGTGGCAGTGGATCAGGGACAGATTTCACACTCAAGATCAGCAGAGTGGAGGCTGAGGATCTGGGAGTTTATTACTGCTTTCAAGGTTCACATGTTCCTCC |
| IGKV8-19*01_B6 | C57BL/6J | Musmus IGKV8-19*01 F | 100 | 305 | GACATTGTGATGACACAGTCTCCATCCTCCCTGACTGTGACAGCAGGAGAGAAGGTCACTATGAGCTGCAAGTCCAGTCAGAGTCTGTTAAACAGTGGAAATCAAAAGAACTACTTGACCTGGTACCAGCAGAAACCAGGGCAGCCTCCTAAACTGTTGATCTACTGGGCATCCACTAGGGAATCTGGGGTCCCTGATCGCTTCACAGGCAGTGGATCTGGAACAGATTTCACTCTCACCATCAGCAGTGTGCAGGCTGAAGACCTGGCAGTTTATTACTGTCAGAATGATTATAGTTATCCTCC |
| IGKV4-57-1*01_B6 | C57BL/6J | Musmus IGKV4-57*01 F | 100 | 292 | GAAAATGTGCTCACCCAGTCTCCAGCAATCATGTCTGCATCTCCAGGGGAAAAGGTCACCATGACCTGCAGGGCCAGCTCAAGTGTAAGTTCCAGTTACTTGCACTGGTACCAGCAGAAGTCAGGTGCCTCCCCCAAACTCTGGATTTATAGCACATCCAACTTGGCTTCTGGAGTCCCTGCTCGCTTCAGTGGCAGTGGGTCTGGGACCTCTTACTCTCTCACAATCAGCAGTGTGGAGGCTGAAGATGCTGCCACTTATTACTGCCAGCAGTACAGTGGTTACCCACTCA |
| IGKV4-68*01_S3092_B6 | C57BL/6J | Musmus IGKV4-68*01 F | 100 | 286 | CAAATTGTTCTCACCCAGTCTCCAGCACTCATGTCTGCATCTCCAGGGGAGAAGGTCACCATGACCTGCAGTGCCAGCTCAAGTGTAAGTTACATGTACTGGTACCAGCAGAAGCCAAGATCCTCCCCCAAACCCTGGATTTATCTCACATCCAACCTGGCTTCTGGAGTCCCTGCTCGCTTCAGTGGCAGTGGGTCTGGGACCTCTTACTCTCTCACAATCAGCAGCATGGAGGCTGAAGATGCTGCCACTTATTACTGCCAGCAGTGGAGTAGTAACCCACTCA |
| IGKV8-28*01_B6 | C57BL/6J | Musmus IGKV8-28*01 F | 100 | 305 | GACATTGTGATGACACAGTCTCCATCCTCCCTGAGTGTGTCAGCAGGAGAGAAGGTCACTATGAGCTGCAAGTCCAGTCAGAGTCTGTTAAACAGTGGAAATCAAAAGAACTACTTGGCCTGGTACCAGCAGAAACCAGGGCAGCCTCCTAAACTGTTGATCTACGGGGCATCCACTAGGGAATCTGGGGTCCCTGATCGCTTCACAGGCAGTGGATCTGGAACCGATTTCACTCTTACCATCAGCAGTGTGCAGGCTGAAGACCTGGCAGTTTATTACTGTCAGAATGATCATAGTTATCCTCC |

| | | | | | |
|---|---|---|---|---|---|
| IGKV4-59*01_B6 | C57BL/6J | Musmus IGKV4-59*01 F | 100 | 286 | CAAATTGTTCTCACCCAGTCTCCAGCAATCATGTCTGCATCTCCAGGGGAGAAGGTCACCATGACCTGCAGTGCCAGCTCAAGTGTAAGTTACATGCACTGGTACCAGCAGAAGTCAGGCACCTCCCCCAAAAGATGGATTTATGACACATCCAAACTGGCTTCTGGAGTCCCTGCTCGCTTCAGTGGCAGTGGGTCTGGGACCTCTTACTCTCTCACAATCAGCAGCATGGAGGCTGAAGATGCTGCCACTTATTACTGCCAGCAGTGGAGTAGTAACCCACCCA |
| IGKV6-32*01_B6 | C57BL/6J | Musmus IGKV6-32*01 F | 100 | 287 | AGTATTGTGATGACCCAGACTCCCAAATTCCTGCTTGTATCAGCAGGAGACAGGGTTACCATAACCTGCAAGGCCAGTCAGAGTGTGAGTAATGATGTAGCTTGGTACCAACAGAAGCCAGGGCAGTCTCCTAAACTGCTGATATACTATGCATCCAATCGCTACACTGGAGTCCCTGATCGCTTCACTGGCAGTGGATATGGGACGGATTTCACTTTCACCATCAGCACTGTGCAGGCTGAAGACCTGGCAGTTTATTTCTGTCAGCAGGATTATAGCTCTCCTCC |
| IGKV1-110*01_B6 | C57BL/6J | Musmus IGKV1-110*01 F | 100 | 302 | GATGTTGTGATGACCCAAACTCCACTCTCCCTGCCTGTCAGTCTTGGAGATCAAGCCTCCATCTCTTGCAGATCTAGTCAGAGCCTTGTACACAGTAATGGAAACACCTATTTACATTGGTACCTGCAGAAGCCAGGCCAGTCTCCAAAGCTCCTGATCTACAAAGTTTCCAACCGATTTTCTGGGGTCCCAGACAGGTTCAGTGGCAGTGGATCAGGGACAGATTTCACACTCAAGATCAGCAGAGTGGAGGCTGAGGATCTGGGAGTTTATTTCTGCTCTCAAAGTACACATGTTCCTCC |
| IGKV6-13*01_B6 | C57BL/6J | Musmus IGKV6-13*01 F | 100 | 285 | GACATTGTGATGACCCAGTCTCAAAAATTCATGTCCACATCAGTAGGAGACAGGGTCAGCATCACCTGCAAGGCCAGTCAGAATGTGGGTACTGCTGTAGCCTGGTATCAACAGAAACCAGGACAATCTCCTAAACTACTGATTTACTCGGCATCCAATCGGTACACTGGAGTCCCTGATCGCTTCACAGGCAGTGGATCTGGGACAGATTTCACTCTCACCATCAGCAATATGCAGTCTGAAGACCTGGCAGATTATTTCTGCCAGCAATATAGCAGCTATCCT |
| IGKV4-80*01_B6 | C57BL/6J | Musmus IGKV4-80*01 F | 100 | 282 | CAAATTGTTCTCACCCAGTCTCCAGCAATCATGTCTGCATCTCTAGGGGAGGAGATCACCCTAACCTGCAGTGCCAGCTCGAGTGTAAGTTACATGCACTGGTACCAGCAGAAGTCAGGCACTTCTCCCAAACTCTTGATTTATAGCACATCCAACCTGGCTTCTGGAGTCCCTTCTCGCTTCAGTGGCAGTGGGTCTGGGACCTTTTATTCTCTCACAATCAGCAGTGTGGAGGCTGAAGATGCTGCCGATTATTACTGCCATCAGTGGAGTAGTTATCCA |
| IGKV12-41*01_B6 | C57BL/6J | Musmus IGKV12-41*01 F | 100 | 287 | GACATCCAGATGACTCAGTCTCCAGCCTCCCTATCTGCATCTGTGGGAGAAACTGTCACCATCACATGTCGAGCAAGTGGGAATATTCACAATTATTTAGCATGGTATCAGCAGAAACAGGGAAAATCTCCTCAGCTCCTGGTCTATAATGCAAAAACCTTAGCAGATGGTGTGCCATCAAGGTTCAGTGGCAGTGGATCAGGAACACAATATTCTCTCAAGATCAACAGCCTGCAGCCTGAAGATTTTGGGAGTTATTACTGTCAACATTTTTGGAGTACTCCTCC |
| IGKV14-111*01_B6 | C57BL/6J | Musmus IGKV14-111*01 F | 100 | 287 | GACATCAAGATGACCCAGTCTCCATCTTCCATGTATGCATCTCTAGGAGAGAGAGTCACTATCACTTGCAAGGCGAGTCAGGACATTAATAGCTATTTAAGCTGGTTCCAGCAGAAACCAGGGAAATCTCCTAAGACCCTGATCTATCGTGCAAACAGATTGGTAGATGGGGTCCCATCAAGGTTCAGTGGCAGTGGATCTGGGCAAGATTATTCTCTCACCATCAGCAGCCTGGAGTATGAAGATATGGGAATTTATTATTGTCTACAGTATGATGAGTTTCCTCC |
| IGKV5-43*01_B6 | C57BL/6J | Musmus IGKV5-43*01 F | 100 | 287 | GATATTGTGCTAACTCAGTCTCCAGCCACCCTGTCTGTGACTCCAGGAGATAGCGTCAGTCTTTCCTGCAGGGCCAGCCAAAGTATTAGCAACAACCTACACTGGTATCAACAAAAATCACATGAGTCTCCAAGGCTTCTCATCAAGTATGCTTCCCAGTCCATCTCTGGGATCCCCTCCAGGTTCAGTGGCAGTGGATCAGGGACAGATTTCACTCTCAGTATCAACAGTGTGGAGACTGAAGATTTTGGAATGTATTTCTGTCAACAGAGTAACAGCTGGCCTCA |

| | | | | | |
|---|---|---|---|---|---|
| IGKV15-103*01_B6 | C57BL/6J | Musmus IGKV15-103*01 ORF | 100 | 287 | GACATCCAGATGAACCAGTCTCCATCCAGTCTGTCTGCATCCCTTGGAGACACAATTA CCATCACTTGCCATGCCAGTCAGAACATTAATGTTTGGTTAAGCTGGTACCAGCAGAA ACCAGGAAATATTCCTAAACTATTGATCTATAAGGCTTCCAACTTGCACACAGGCGTC CCATCAAGGTTTAGTGGCAGTGGATCTGGAACAGGTTTCACATTAACCATCAGCAGCC TGCAGCCTGAAGACATTGCCACTTACTACTGTCAACAGGGTCAAAGTTATCCTCT |
| IGKV8-24*01_B6 | C57BL/6J | Musmus IGKV8-24*01 F | 100 | 305 | GACATTGTGATGACACAGTCTCCATCCTCCCTGGCTATGTCAGTAGGACAGAAGGTCA CTATGAGCTGCAAGTCCAGTCAGAGCCTTTTAAATAGTAGCAATCAAAAGAACTATTT GGCCTGGTACCAGCAGAAACCAGGACAGTCTCCTAAACTTCTGGTATACTTTGCATCC ACTAGGGAATCTGGGGTCCCTGATCGCTTCATAGGCAGTGGATCTGGGACAGATTTCA CTCTTACCATCAGCAGTGTGCAGGCTGAAGACCTGGCAGATTACTTCTGTCAGCAACA TTATAGCACTCCTCC |
| IGKV17-127*01_B6 | C57BL/6J | Musmus IGKV17-127*01 F | 100 | 287 | GAAACAACTGTGACCCAGTCTCCAGCATCCCTGTCCGTGGCTACAGGAGAAAAAGTC ACTATCAGATGCATAACCAGCACTGATATTGATGATGATATGAACTGGTACCAGCAGA AGCCAGGGGAACCTCCTAAGCTCCTTATTTCAGAAGGCAATACTCTTCGTCCTGGAGT CCCATCCCGATTCTCCAGCAGTGGCTATGGCACAGATTTTGTTTTTACAATTGAAAACA CGCTCTCAGAAGATGTTGCAGATTACTACTGTTTGCAAAGTGATAACATGCCTCT |
| IGKV8-16*01_B6 | C57BL/6J | Musmus IGKV8-16*01 F | 100 | 305 | GAAATTGTGTTGACCCAGTCTATACCATCCCTGACTGTGTCAGCAGGAGAGAGGGTCA CTATCAGCTGCAAATCCAATCAGAATCTTTTATGGAGTGGAAACCAAAGGTACTGTTT GGTCTGGCACCAGTGGAAACCAGGGCAAACTCCTACACCGTTGATCACCTGGACATCT GATAGGTACTCTGGAGTCCCTGATCGTTTCATAGGCAGTGGATCTGTGACAGATTTCA CTCTGACCATCAGCAGTGTGCAGGCTGAAGATGTGGCAGTTTATTTCTGTCAGCAGCA TTTACACATTCCTCC |
| IGKV6-17*01_B6 | C57BL/6J | Musmus IGKV6-17*01 F | 100 | 287 | GACATTGTGATGACCCAGTCTCACAAATTCATGTCCACATCAGTAGGAGACAGGGTCA GCATCACCTGCAAGGCCAGTCAGGATGTGAGTACTGCTGTAGCCTGGTATCAACAGAA ACCAGGACAATCTCCTAAACTACTGATTTACTCGGCATCCTACCGGTACACTGGAGTC CCTGATCGCTTCACTGGCAGTGGATCTGGGACGGATTTCACTTTCACCATCAGCAGTG TGCAGGCTGAAGACCTGGCAGTTTATTACTGTCAGCAACATTATAGTACTCCTCC |
| IGKV19-93*01_B6 | C57BL/6J | Musmus IGKV19-93*01 F | 100 | 287 | GACATCCAGATGACACAGTCTCCATCCTCACTGTCTGCATCTCTGGGAGGCAAAGTCA CCATCACTTGCAAGGCAAGCCAAGACATTAACAAGTATATAGCTTGGTACCAACACA AGCCTGGAAAAGGTCCTAGGCTGCTCATACATTACACATCTACATTACAGCCAGGCAT CCCATCAAGGTTCAGTGGAAGTGGGTCTGGGAGAGATTATTCCTTCAGCATCAGCAAC CTGGAGCCTGAAGATATTGCAACTTATTATTGTCTACAGTATGATAATCTTCTACC |
| IGKV9-120*01_S2628_B6 | C57BL/6J | Musmus IGKV9-120*02 F | 100 | 285 | GACATCCAGATGACCCAGTCTCCATCCTCCTTATCTGCCTCTCTGGGAGAAAGAGTCA GTCTCACTTGTCGGGCAAGTCAGGACATTGGTAGTAGCTTAAACTGGCTTCAGCAGGA ACCAGATGGAACTATTAAACGCCTGATCTACGCCACATCCAGTTTAGATTCTGGTGTC CCCAAAAGGTTCAGTGGCAGTAGGTCTGGGTCAGATTATTCTCTCACCATCAGCAGCC TTGAGTCTGAAGATTTTGTAGACTATTACTGTCTACAATATGCTAGTTCTCCT |
| IGKV4-70*01_B6 | C57BL/6J | Musmus IGKV4-70*01 F | 100 | 282 | CAAATTGTTCTCACCCAGTCTCCAGCAATCATGTCTGCATCTCCAGGGGAGAAGGTCA CCATGACCTGCAGTGCCAGCTCAAGTATAAGTTACATGCACTGGTACCAGCAGAAGCC AGGCACCTCCCCCAAAAGATGGATTTATGACACATCCAAACTGGCTTCTGGAGTCCCT GCTCGCTTCAGTGGCAGTGGGTCTGGGACCTCTTATTCTCTCACAATCAGCAGCATGG AGGCTGAAGATGCTGCCACTTATTACTGCCATCAGCGGAGTAGTTACCCA |

| | | | | | |
|---|---|---|---|---|---|
| IGKV13-84*01_B6 | C57BL/6J | Musmus IGKV13-84*01 F | 100 | 287 | GACATCCAGATGACACAATCTTCATCCTCCTTTTCTGTATCTCTAGGAGACAGAGTCACCATTACTTGCAAGGCAAGTGAGGACATATATAATCGGTTAGCCTGGTATCAGCAGAAACCAGGAAATGCTCCTAGGCTCTTAATATCTGGTGCAACCAGTTTGGAAACTGGGGTTCCTTCAAGATTCAGTGGCAGTGGATCTGGAAAGGATTACACTCTCAGCATTACCAGTCTTCAGACTGAAGATGTTGCTACTTATTACTGTCAACAGTATTGGAGTACTCCTCC |
| IGKV4-72*01_B6 | C57BL/6J | Musmus IGKV4-72*01 F | 100 | 286 | CAAATTGTTCTCTCCCAGTCTCCAGCAATCCTGTCTGCATCTCCAGGGGAGAAGGTCACAATGACTTGCAGGGCCAGCTCAAGTGTAAGTTACATGCACTGGTACCAGCAGAAGCCAGGATCCTCCCCCAAACCCTGGATTTATGCCACATCCAACCTGGCTTCTGGAGTCCCTGCTCGCTTCAGTGGCAGTGGGTCTGGGACCTCTTACTCTCTCACAATCAGCAGAGTGGAGGCTGAAGATGCTGCCACTTATTACTGCCAGCAGTGGAGTAGTAACCCACCCA |
| IGKV10-94*01_B6 | C57BL/6J | Musmus IGKV10-94*01 F | 100 | 287 | GATATCCAGATGACACAGACTACATCCTCCCTGTCTGCCTCTCTGGGAGACAGAGTCACCATCAGTTGCAGTGCAAGTCAGGGCATTAGCAATTATTTAAACTGGTATCAGCAGAAACCAGATGGAACTGTTAAACTCCTGATCTATTACACATCAAGTTTACACTCAGGAGTCCCATCAAGGTTCAGTGGCAGTGGGTCTGGGACAGATTATTCTCTCACCATCAGCAACCTGGAACCTGAAGATATTGCCACTTACTATTGTCAGCAGTATAGTAAGCTTCCTCC |
| IGKV17-121*01_B6 | C57BL/6J | Musmus IGKV17-121*01 F | 100 | 287 | GAAACAACTGTGACCCAGTCTCCAGCATCCCTGTCCATGGCTATAGGAGAAAAAGTCACCATCAGATGCATAACCAGCACTGATATTGATGATGATATGAACTGGTACCAGCAGAAGCCAGGGGAACCTCCTAAGCTCCTTATTTCAGAAGGCAATACTCTTCGTCCTGGAGTCCCATCCCGATTCTCCAGCAGTGGCTATGGTACAGATTTTGTTTTTACAATTGAAAACATGCTCTCAGAAGATGTTGCAGATTACTACTGTTTGCAAAGTGATAACTTGCCTCT |
| IGKV14-126*01_B6 | C57BL/6J | Musmus IGKV14-126*01 F | 100 | 287 | GACATCAAGATGACCCAGTCTCCATCCTCCATGTATGCATCGCTGGGAGAGAGAGTCACTATCACTTGCAAGGCGAGTCAGGACATTAAAAGCTATTTAAGCTGGTACCAGCAGAAACCATGGAAATCTCCTAAGACCCTGATCTATTATGCAACAAGCTTGGCAGATGGGGTCCCATCAAGATTCAGTGGCAGTGGATCTGGGCAAGATTATTCTCTAACCATCAGCAGCCTGGAGTCTGACGATACAGCAACTTATTACTGTCTACAGCATGGTGAGAGCCCTCC |
| IGKV5-39*01_B6 | C57BL/6J | Musmus IGKV5-39*01 F | 100 | 287 | GACATTGTGATGACTCAGTCTCCAGCCACCCTGTCTGTGACTCCAGGAGATAGAGTCTCTCTTTCCTGCAGGGCCAGCCAGAGTATTAGCGACTACTTACACTGGTATCAACAAAAATCACATGAGTCTCCAAGGCTTCTCATCAAATATGCTTCCCAATCCATCTCTGGGATCCCCTCCAGGTTCAGTGGCAGTGGATCAGGGTCAGATTTCACTCTCAGTATCAACAGTGTGGAACCTGAAGATGTTGGAGTGTATTACTGTCAAAATGGTCACAGCTTTCCTCC |
| IGKV5-48*01_B6 | C57BL/6J | Musmus IGKV5-48*01 F | 100 | 287 | GACATCTTGCTGACTCAGTCTCCAGCCATCCTGTCTGTGAGTCCAGGAGAAAGAGTCAGTTTCTCCTGCAGGGCCAGTCAGAGCATTGGCACAAGCATACACTGGTATCAGCAAAGAACAAATGGTTCTCCAAGGCTTCTCATAAAGTATGCTTCTGAGTCTATCTCTGGGATCCCTTCCAGGTTTAGTGGCAGTGGATCAGGGACAGATTTTACTCTTAGCATCAACAGTGTGGAGTCTGAAGATATTGCAGATTATTACTGTCAACAAAGTAATAGCTGGCCAAC |
| IGKV13-85*01_B6 | C57BL/6J | Musmus IGKV13-85*01 F | 100 | 287 | GACATCCAGATGACACAATCTTCATCCTACTTGTCTGTATCTCTAGGAGGCAGAGTCACCATTACTTGCAAGGCAAGTGACCACATTAATAATTGGTTAGCCTGGTATCAGCAGAAACCAGGAAATGCTCCTAGGCTCTTAATATCTGGTGCAACCAGTTTGGAAACTGGGGTTCCTTCAAGATTCAGTGGCAGTGGATCTGGAAAGGATTACACTCTCAGCATTACCAGTCTTCAGACTGAAGATGTTGCTACTTATTACTGTCAACAGTATTGGAGTACTCCTCC |

| | | | | | |
|---|---|---|---|---|---|
| IGKV3-4*01_B6 | C57BL/6J | Musmus IGKV3-4*01 F | 100 | 299 | GACATTGTGCTGACCCAATCTCCAGCTTCTTTGGCTGTGTCTCTAGGGCAGAGGGCCACCATCTCCTGCAAGGCCAGCCAAAGTGTTGATTATGATGGTGATAGTTATATGAACTGGTACCAACAGAAACCAGGACAGCCACCCAAACTCCTCATCTATGCTGCATCCAATCTAGAATCTGGGATCCCAGCCAGGTTTAGTGGCAGTGGGTCTGGGACAGACTTCACCCTCAACATCCATCCTGTGGAGGAGGAGGATGCTGCAACCTATTACTGTCAGCAAAGTAATGAGGATCCTCC |
| IGKV4-91*01_S5997_B6 | C57BL/6J | Musmus IGKV4-91*01 F | 100 | 292 | GAAATTGTGCTCACCCAGTCTCCAACCACCATGGCTGCATCTCCCGGGGAGAAGATCACTATCACCTGCAGTGCCAGCTCAAGTATAAGTTCCAATTACTTGCATTGGTATCAGCAGAAGCCAGGATTCTCCCCTAAACTCTTGATTTATAGGACATCCAATCTGGCTTCTGGAGTCCCAGCTCGCTTCAGTGGCAGTGGGTCTGGGACCTCTTACTCTCTCACAATTGGCACCATGGAGGCTGAAGATGTTGCCACTTACTACTGCCAGCAGGGTAGTAGTATACCACTCA |
| IGKV8-34*01_B6 | C57BL/6J | Musmus IGKV8-34*01 F | 100 | 305 | GACATTTTGATGACCCAGTCTCCATCCTCCCTGACTGTGTCAGCAGGAGAGAAGGTCACTATGAGCTGCAAGTCCAGTCAGAGTCTTTTAGCTAGTGGCAACCAAAATAACTACTTGGCCTGGCACCAGCAGAAACCAGGACGATCTCCTAAAATGCTGATAATTTGGGCATCCACTAGGGTATCTGGAGTCCCTGATCGCTTCATAGGCAGTGGATCTGGGACGGATTTCACTCTGACCATCAACAGTGTGCAGGCTGAAGATCTGGCTGTTTATTACTGTCAGCAGTCCTACAGCGCTCCTAC |
| IGKV3-12*01_B6 | C57BL/6J | Musmus IGKV3-12*01 F | 100 | 299 | GACATTGTGCTGACACAGTCTCCTGCTTCCTTAGCTGTATCTCTGGGGCAGAGGGCCACCATCTCATGCAGGGCCAGCAAAAGTGTCAGTACATCTGGCTATAGTTATATGCACTGGTACCAACAGAAACCAGGACAGCCACCCAAACTCCTCATCTATCTTGCATCCAACCTAGAATCTGGGGTCCCTGCCAGGTTCAGTGGCAGTGGGTCTGGGACAGACTTCACCCTCAACATCCATCCTGTGGAGGAGGAGGATGCTGCAACCTATTACTGTCAGCACAGTAGGGAGCTTCCTCC |
| IGKV3-2*01_B6 | C57BL/6J | Musmus IGKV3-2*01 F | 100 | 299 | GACATTGTGCTGACCCAATCTCCAGCTTCTTTGGCTGTGTCTCTAGGGCAGAGGGCCACCATCTCCTGCAGAGCCAGCGAAAGTGTTGATAATTATGGCATTAGTTTTATGAACTGGTTCCAACAGAAACCAGGACAGCCACCCAAACTCCTCATCTATGCTGCATCCAACCAAGGATCCGGGGTCCCTGCCAGGTTTAGTGGCAGTGGGTCTGGGACAGACTTCAGCCTCAACATCCATCCTATGGAGGAGGATGATACTGCAATGTATTTCTGTCAGCAAAGTAAGGAGGTTCCTCC |
| IGKV14-100*01_B6 | C57BL/6J | Musmus IGKV14-100*01 F | 100 | 287 | GACATCCTGATGACCCAATCTCCATCCTCCATGTCTGTATCTCTGGGAGACACAGTCAGCATCACTTGCCATGCAAGTCAGGGCATTAGCAGTAATATAGGGTGGTTGCAGCAGAAACCAGGGAAATCATTTAAGGGCCTGATCTATCATGGAACCAACTTGGAAGATGGAGTTCCATCAAGGTTCAGTGGCAGTGGATCTGGAGCAGATTATTCTCTCACCATCAGCAGCCTGGAATCTGAAGATTTTGCAGACTATTACTGTGTACAGTATGCTCAGTTTCCTCC |
| IGKV4-55*01_B6 | C57BL/6J | Musmus IGKV4-55*01 F | 100 | 286 | CAAATTGTTCTCACCCAGTCTCCAGCAATCATGTCTGCATCTCCAGGGGAGAAGGTCACCATGACCTGCAGTGCCAGCTCAAGTGTAAGTTACATGTACTGGTACCAGCAGAAGCCAGGATCCTCCCCCAGACTCCTGATTTATGACACATCCAACCTGGCTTCTGGAGTCCCTGTTCGCTTCAGTGGCAGTGGGTCTGGGACCTCTTACTCTCTCACAATCAGCCGAATGGAGGCTGAAGATGCTGCCACTTATTACTGCCAGCAGTGGAGTAGTTACCCACCCA |

| | | | | | |
|---|---|---|---|---|---|
| IGKV12-98*01_B6 | C57BL/6J | Musmus IGKV12-98*01 F | 100 | 287 | GACATTCAGATGACCCAGTCTCCTGCCTCCCAGTCTGCATCTCTGGGAGAAAGTGTCACCATCACATGCCTGGCAAGTCAGACCATTGGTACATGGTTAGCATGGTATCAGCAGAAACCAGGGAAATCTCCTCAGCTCCTGATTTATGCTGCAACCAGCTTGGCAGATGGGGTCCCATCAAGGTTCAGTGGTAGTGGATCTGGCACAAAATTTTCTTTCAAGATCAGCAGCCTACAGGCTGAAGATTTTGTAAGTTATTACTGTCAACAACTTTACAGTACTCCTCT |
| IGKV4-63*01_B6 | C57BL/6J | Musmus IGKV4-63*01 F | 100 | 286 | GAAAATGTTCTCACCCAGTCTCCAGCAATCATGTCTGCATCTCCAGGGGAAAAGGTCACCATGACCTGCAGTGCCAGCTCAAGTGTAAGTTACATGCACTGGTACCAGCAGAAGTCAAGCACCTCCCCCAAACTCTGGATTTATGACACATCCAAACTGGCTTCTGGAGTCCCAGGTCGCTTCAGTGGCAGTGGGTCTGGAAACTCTTACTCTCTCACGATCAGCAGCATGGAGGCTGAAGATGTTGCCACTTATTACTGTTTTCAGGGGAGTGGGTACCCACTCA |
| IGKV6-25*01_B6 | C57BL/6J | Musmus IGKV6-25*01 F | 100 | 287 | GACATTGTGATGACCCAGTCTCACAAATTCATGTCCACATCAGTAGGAGACAGGGTCAGCATCACCTGCAAGGCCAGTCAGGATGTGAGTACTGCTGTAGCCTGGTATCAACAAAAACCAGGGCAATCTCCTAAACTACTGATTTACTGGGCATCCACCCGGCACACTGGAGTCCCTGATCGCTTCACAGGCAGTGGATCTGGGACAGATTATACTCTCACCATCAGCAGTGTGCAGGCTGAAGACCTGGCACTTTATTACTGTCAGCAACATTATAGCACTCCTCC |
| IGKV16-104*01_B6 | C57BL/6J | Musmus IGKV16-104*01 F | 100 | 288 | GATGTCCAGATAACCCAGTCTCCATCTTATCTTGCTGCATCTCCTGGAGAAACCATTACTATTAATTGCAGGGCAAGTAAGAGCATTAGCAAATATTTAGCCTGGTATCAAGAGAAACCTGGGAAAACTAATAAGCTTCTTATCTACTCTGGATCCACTTTGCAATCTGGAATTCCATCAAGGTTCAGTGGCAGTGGATCTGGTACAGATTTCACTCTCACCATCAGTAGCCTGGAGCCTGAAGATTTTGCAATGTATTACTGTCAACAGCATAATGAATACCCGTAC |
| IGKV2-109*01_B6 | C57BL/6J | Musmus IGKV2-109*01 F | 100 | 302 | GATATTGTGATGACGCAGGCTGCATTCTCCAATCCAGTCACTCTTGGAACATCAGCTTCCATCTCCTGCAGGTCTAGTAAGAGTCTCCTACATAGTAATGGCATCACTTATTTGTATTGGTATCTGCAGAAGCCAGGCCAGTCTCCTCAGCTCCTGATTTATCAGATGTCCAACCTTGCCTCAGGAGTCCCAGACAGGTTCAGTAGCAGTGGGTCAGGAACTGATTTCACACTGAGAATCAGCAGAGTGGAGGCTGAGGATGTGGGTGTTTATTACTGTGCTCAAAATCTAGAACTTCCTCC |
| IGKV4-74*01_B6 | C57BL/6J | Musmus IGKV4-74*01 F | 100 | 292 | CAAATTGTTCTCACCCAGTCTCCAGCAATCATGTCTGCATCTCTAGGGGAACGGGTCACCATGACCTGCACTGCCAGCTCAAGTGTAAGTTCCAGTTACTTGCACTGGTACCAGCAGAAGCCAGGATCCTCCCCCAAACTCTGGATTTATAGCACATCCAACCTGGCTTCTGGAGTCCCAGCTCGCTTCAGTGGCAGTGGGTCTGGGACCTCTTACTCTCTCACAATCAGCAGCATGGAGGCTGAAGATGCTGCCACTTATTACTGCCACCAGTATCATCGTTCCCCACCCA |
| IGKV3-5*01_B6 | C57BL/6J | Musmus IGKV3-5*01 F | 100 | 299 | GACATTGTGCTGACCCAATCTCCAGCTTCTTTGGCTGTGTCTCTAGGGCAGAGGGCCACCATATCCTGCAGAGCCAGTGAAAGTGTTGATAGTTATGGCAATAGTTTTATGCACTGGTACCAGCAGAAACCAGGACAGCCACCCAAACTCCTCATCTATCGTGCATCCAACCTAGAATCTGGGATCCCTGCCAGGTTCAGTGGCAGTGGGTCTAGGACAGACTTCACCCTCACCATTAATCCTGTGGAGGCTGATGATGTTGCAACCTATTACTGTCAGCAAAGTAATGAGGATCCTCC |
| IGKV4-86*01_B6 | C57BL/6J | Musmus IGKV4-86*01 F | 100 | 283 | GAAATTGTGCTCACTCAGTCTCCAGCCATCACAGCTGCATCTCTGGGGCAAAAGGTCACCATCACCTGCAGTGCCAGCTCAAGTGTAAGTTACATGCACTGGTACCAGCAGAAGTCAGGCACCTCCCCCAAACCATGGATTTATGAAATATCCAAACTGGCTTCTGGAGTCCCAGCTCGCTTCAGTGGCAGTGGGTCTGGGACCTCTTACTCTCTCACAATCAGCAGCATGGAGGCTGAAGATGCTGCCATTTATTACTGCCAGCAGTGGAATTATCCTCTTA |

| | | | | | |
|---|---|---|---|---|---|
| IGKV3-7*01_S5296_B6 | C57BL/6J | Musmus IGKV3-7*02 F | 100 | 299 | GACATTGTGCTGACACAGTCTCCTGCTTCCTTAGCTGTATCTCTGGGGCAGAGGGCCACCATCTCATGCAGGGCCAGCCAAAGTGTCAGTACATCTAGCTATAGTTATATGCACTGGTACCAACAGAAACCAGGACAGCCACCCAAACTCCTCATCAAGTATGCATCCAACCTAGAATCTGGGGTCCCTGCCAGGTTCAGTGGCAGTGGGTCTGGGACAGACTTCACCCTCAACATCCATCCTGTGGAGGAGGAGGATACTGCAACATATTACTGTCAGCACAGTTGGGAGATTCCTCC |
| IGKV4-53*01_B6 | C57BL/6J | Musmus IGKV4-53*01 F | 100 | 292 | GAAATTGTGCTCACCCAGTCTCCAGCACTCATGGCTGCATCTCCAGGGGAGAAGGTCACCATCACCTGCAGTGTCAGCTCAAGTATAAGTTCCAGCAACTTGCACTGGTACCAGCAGAAGTCAGAAACCTCCCCCAAACCCTGGATTTATGGCACATCCAACCTGGCTTCTGGAGTCCCTGTTCGCTTCAGTGGCAGTGGATCTGGGACCTCTTATTCTCTCACAATCAGCAGCATGGAGGCTGAAGATGCTGCCACTTATTACTGTCAACAGTGGAGTAGTTACCCACTCA |
| IGKV5-45*01_B6 | C57BL/6J | Musmus IGKV5-45*01 F | 100 | 287 | GATATTGTGCTAACTCAGTCTCCAGCCACCCTGTCTGTGACTCCAGGAGATAGAGTCAGTCTTTCCTGCAGGGCCAGTCAAAGTATTAGCAACTACCTACACTGGTATCAACAAAAATCACATGAGTCTCCAAGGCTTCTCATCAAGTATGCTTCCCAGTCCATCTCTGGGATCCCCTCCAGGTTCAGTGGCAGTGGATCAGGGACAGATTTCACTCTCAGTATCAACAGTGTGGAGACTGAAGATTTTGGAATGTATTTCTGTCAACAGAGTAACAGCTGGCCTCA |
| IGKV4-69*01_B6 | C57BL/6J | Musmus IGKV4-69*01 F | 100 | 282 | CAAATTCTTCTCACCCAGTCTCCAGCAATCATGTCTGCATCTCCAGGGGAGAAGGTCACCATGACCTGCAGTGCCAGCTCAAGTGTAAGTTACATGCACTGGTACCAGCAGAAGCCAGGATCCTCGCCCAAACCCTGGATTTATGACACATCCAACCTGGCTTCTGGATTCCCTGCTCGCTTCAGTGGCAGTGGGTCTGGGACCTCTTACTCTCTCATAATCAGCAGCATGGAGGCTGAAGATGCTGCCACTTATTACTGCCATCAGCGGAGTAGTTACCCA |
| IGKV9-124*01_B6 | C57BL/6J | Musmus IGKV9-124*01 F | 100 | 285 | GACATCCAGATGACCCAGTCTCCATCCTCCTTATCTGCCTCTCTGGGAGAAAGAGTCAGTCTCACTTGTCGGGCAAGTCAGGAAATTAGTGGTTACTTAAGCTGGCTTCAGCAGAAACCAGATGGAACTATTAAACGCCTGATCTACGCCGCATCCACTTTAGATTCTGGTGTCCCAAAAAGGTTCAGTGGCAGTAGGTCTGGGTCAGATTATTCTCTCACCATCAGCAGCCTTGAGTCTGAAGATTTTGCAGACTATTACTGTCTACAATATGCTAGTTATCCT |
| IGKV1-133*01_B6 | C57BL/6J | Musmus IGKV1-133*01 F | 100 | 302 | GATGTTGTGATGACCCAGACTCCACTCACTTTGTCGGTTACCATTGGACAACCAGCCTCTATCTCTTGCAAGTCAAGTCAGAGCCTCTTATATAGTAATGGAAAAACCTATTTGAATTGGTTATTACAGAGGCCAGGCCAGTCTCCAAAGCGCCTAATCTATCTGGTGTCTAAACTGGACTCTGGAGTCCCTGACAGGTTCACTGGCAGTGGATCAGGAACAGATTTTACACTGAAAATCAGCAGAGTGGAGGCTGAGGATTTGGGAGTTTATTACTGCGTGCAAGGTACACATTTTCCTCA |
| IGKV4-50*01_S0901_B6 | C57BL/6J | Musmus IGKV4-50*01 F | 100 | 286 | GAAAATGTGCTCACCCAGTCTCCAGCAATCATGTCTGCATCTCTAGGGGAGAAGGTCACCATGAGCTGCAGGGCCAGCTCAAGTGTAAATTACATGTACTGGTACCAGCAGAAGTCAGATGCCTCCCCCAAACTATGGATTTATTACACATCCAACCTGGCTCCTGGAGTCCCAGCTCGCTTCAGTGGCAGTGGGTCTGGGAACTCTTATTCTCTCACAATCAGCAGCATGGAGGGTGAAGATGCTGCCACTTATTACTGCCAGCAGTTTACTAGTTCCCCATACA |
| IGKV4-61*01_B6 | C57BL/6J | Musmus IGKV4-61*01 F | 100 | 286 | CAAATTGTTCTCACCCAGTCTCCAGCAATCATGTCTGCATCTCCAGGGGAGAAGGTCACCATATCCTGCAGTGCCAGCTCAAGTGTAAGTTACATGTACTGGTACCAGCAGAAGCCAGGATCCTCCCCCAAACCCTGGATTTATCGCACATCCAACCTGGCTTCTGGAGTCCCTGCTCGCTTCAGTGGCAGTGGGTCTGGGACCTCTTACTCTCTCACAATCAGCAGCATGGAGGCTGAAGATGCTGCCACTTATTACTGCCAGCAGTATCATAGTTACCCACCCA |

| | | | | | |
|---|---|---|---|---|---|
| IGKV1-122*01_B6 | C57BL/6J | Musmus IGKV1-122*01 F | 100 | 302 | GATGCTGTGATGACCCAAACTCCACTCTCCCTGCCTGTCAGTCTTGGAGATCAAGCCTCCATCTCTTGCAGGTCTAGTCAGAGCCTTGAAAACAGTAATGGAAACACCTATTTGAACTGGTACCTCCAGAAACCAGGCCAGTCTCCACAGCTCCTGATCTACAGGGTTTCCAACCGATTTTCTGGGGTCCTAGACAGGTTCAGTGGTAGTGGATCAGGGACAGATTTCACACTGAAAATCAGCAGAGTGGAGGCTGAGGATTTGGGAGTTTATTTCTGCCTCCAAGTTACACATGTCCCTCC |
| IGKV4-79*01_B6 | C57BL/6J | Musmus IGKV4-79*01 F | 100 | 292 | CAAATTGTTCTCACCCAGTCTCCAGCAATCATGTCTGCATCTCCTGGGGAGAAGGTCACCTTGACCTGCAGTGCCAGCTCAAGTGTAAGTTCCAGCTACTTGTACTGGTACCAGCAGAAGCCAGGATCCTCCCCCAAACTCTGGATTTATAGCACATCCAACCTGGCTTCTGGAGTCCCTGCTCGCTTCAGTGGCAGTGGGTCTGGGACCTCTTACTCTCTCACAATCAGCAGCATGGAGGCTGAAGATGCTGCCTCTTATTTCTGCCATCAGTGGAGTAGTTACCCACCCC |
| IGKV4-92*01_B6 | C57BL/6J | Musmus IGKV4-92*01 F | 100 | 291 | GAAATGGTTCTCACCCAGTCTCCAGTATCCATAACTGCATCTCGAGGGGAGAAGGTCACCATCACCTGCCGTGCCAGCTCAAGTATAAGTTCCAATTACTTACACTGGTACCAGCAGAAGCCAGGATCCTCCCCTAAACTTTTGATTTATAGGACATCCATCCTGGCATCTGGAGTCCTAGACAGCTTCAGTGGCAGTGGGTCTGAGAGCTCTTACACTCTGACAATCAGCTGCATGCAGGACGAAGTTGCTGCCACTTACTATTGTCAGCAGGGGAGTAGTAGCCCACCA |
| IGKV2-112*01_B6 | C57BL/6J | Musmus IGKV2-112*01 F | 100 | 300 | GATATTGTGATAACCCAGGATGAACTCTCCAATCCTGTCACTTCTGGAGAATCAGTTTCCATCTCCTGCAGGTCTAGTAAGAGTCTCCTATATAAGGATGGGAAGACATACTTGAATTGGTTTCTGCAGAGACCAGGACAATCTCCTCAGCTCCTGATCTATTTGATGTCCACCCGTGCATCAGGAGTCTCAGACCGGTTTAGTGGCAGTGGGTCAGGAACAGATTTCACCCTGGAAATCAGTAGAGTGAAGGCTGAGGATGTGGGTGTGTATTACTGTCAACAACTTGTAGAGTATCCT |
| IGKV8-23-1*01_B6 | C57BL/6J | Musmus IGKV8-16*01 F, or Musmus IGKV8-23-1*01 ORF | 100 | 284 | GAAATTGTGCTGACTGTGTCAGCAGGAGAGAGGGTCACTATCAGCTGCAAATCCAATCAGAATCTTTTATGGAGTGGAAACCAAAGGTACTGTTTGGTCTGGCACCAGTGGAAACCAGGGCAAACTCCTACACCGTTGATCACCTGGACATCTGATAGGTACTCTGGAGTCCCTGATCGTTTCATAGGCAGTGGATCTGTGACAGATTTCACTCTGACCATCAGCAGTGTGCAGGCTGAAGATGTGGCAGTTTATTTCTGTCAGCAGCATTTACACATTCCTCC |
| IGKV3-10*01_B6 | C57BL/6J | Musmus IGKV3-10*01 F | 100 | 299 | AACATTGTGCTGACCCAATCTCCAGCTTCTTTGGCTGTGTCTCTAGGGCAGAGGGCCACCATATCCTGCAGAGCCAGTGAAAGTGTTGATAGTTATGGCAATAGTTTTATGCACTGGTACCAGCAGAAACCAGGACAGCCACCCAAACTCCTCATCTATCTTGCATCCAACCTAGAATCTGGGGTCCCTGCCAGGTTCAGTGGCAGTGGGTCTAGGACAGACTTCACCCTCACCATTGATCCTGTGGAGGCTGATGATGCTGCAACCTATTACTGTCAGCAAAATAATGAGGATCCTCC |
| IGKV8-18*01_B6 | C57BL/6J | Musmus IGKV8-18*01 ORF | 100 | 310 | GACATTGTGATGACCCAGTCTCCATCCTCCCTGGCTGTGACAGCAGGAGAGAAGGTCACTATGAGATGCAAGTCCAGTCAGAGTCTTTTGTGGAGTGTAAACCAAAATAACTACTTATCCTGGTACCAGCAGAAACAAGGGCAGCCTCCTAAACTGCTTATCTATGGGGCATCCATTAGAGAATCTTGGGTCCCTGATCGATTCACAGGAAGTGGATCTGGGACAGACTTCACTCTCACCATTAGCAATGTGCATGCTGAAGACCTAGCAGTTTATTACTGTCAGCACAATCATGGCAGCTTTCTCCCCC |

| | | | | | |
|---|---|---|---|---|---|
| IGKV1-99*01_B6 | C57BL/6J | Musmus IGKV1-99*01 F | 100 | 302 | GATGTTGTTCTGACCCAAACTCCACTCTCTCTGCCTGTCAATATTGGAGATCAAGCCTCTATCTCTTGCAAGTCTACTAAGAGTCTTCTGAATAGTGATGGATTCACTTATTTGGACTGGTACCTGCAGAAGCCAGGCCAGTCTCCACAGCTCCTAATATATTTGGTTTCTAATCGATTTTCTGGAGTTCCAGACAGGTTCAGTGGCAGTGGGTCAGGAACAGATTTCACACTCAAGATCAGCAGAGTGGAGGCTGAGGATTTGGGAGTTTATTATTGCTTCCAGAGTAACTATCTTCCTCT |
| IGKV12-38*01_B6 | C57BL/6J | Musmus IGKV12-38*01 F | 100 | 287 | GACATCCAGATGACTCAGTCTCCAGCCTCCCTGGCTGCATCTGTGGGAGAAACTGTCACCATCACATGTCGAGCAAGTGAGAACATTTACTACAGTTTAGCATGGTATCAGCAGAAGCAAGGGAAATCTCCTCAGCTCCTGATCTATAATGCAAACAGCTTGGAAGATGGTGTCCCATCGAGGTTCAGTGGCAGTGGATCTGGGACACAGTATTCTATGAAGATCAACAGCATGCAGCCTGAAGATACCGCAACTTATTTCTGTAAACAGGCTTATGACGTTCCTCC |
| IGKV14-130*01_B6 | C57BL/6J | Musmus IGKV14-130*01 F | 100 | 287 | GAAATCCAGATGACCCAGTCTCCATCCTCTATGTCTGCATCTCTGGGAGACAGAATAACCATCACTTGCCAGGCAACTCAAGACATTGTTAAGAATTTAAACTGGTATCAGCAGAAACCAGGGAAACCCCCTTCATTCCTGATCTATTATGCAACTGAACTGGCAGAAGGGGTCCCATCAAGGTTCAGTGGCAGTGGGTCTGGGTCAGACTATTCTCTGACAATCAGCAACCTGGAGTCTGAAGATTTTGCAGACTATTACTGTCTACAGTTTTATGAGTTTCCTCC |
| IGKV9-123*01_B6 | C57BL/6J | Musmus IGKV9-123*01 F | 100 | 285 | GACATCCAGATGATTCAGTCTCCATCGTCCATGTTTGCCTCTCTGGGAGACAGAGTCAGTCTCTCTTGTCGGGCTAGTCAGGGCATTAGAGGTAATTTAGACTGGTATCAGCAGAAACCAGGTGGAACTATTAAACTCCTGATCTACTCCACATCCAATTTAAATTCTGGTGTCCCATCAAGGTTCAGTGGCAGTGGGTCTGGGTCAGATTATTCTCTCACCATCAGCAGCCTAGAGTCTGAAGATTTTGCAGACTATTACTGTCTACAGCGTAATGCGTATCCT |
| IGKV4-58*01_B6 | C57BL/6J | Musmus IGKV4-58*01 F | 100 | 292 | GAAAATGTGCTCACCCAGTCTCCAGCAATAATGGCTGCCTCTCTGGGGCAGAAGGTCACCATGACCTGCAGTGCCAGCTCAAGTGTAAGTTCCAGTTACTTGCACTGGTACCAGCAGAAGTCAGGCGCTTCCCCCAAACCCTTGATTCATAGGACATCCAACCTGGCTTCTGGAGTCCCAGCTCGCTTCAGTGGCAGTGGGTCTGGGACCTCTTACTCTCTCACAATCAGCAGCGTGGAGGCTGAAGATGATGCAACTTATTACTGCCAGCAGTGGAGTGGTTACCCATTCA |
| IGKV7-33*01_B6 | C57BL/6J | Musmus IGKV7-33*01 F | 100 | 305 | GACATTGTGATGACTCAGTCTCCAACTTTCCTTGCTGTGACAGCAAGTAAGAAGGTCACCATTAGTTGCACGGCCAGTGAGAGCCTTTATTCAAGCAAACACAAGGTGCACTACTTGGCTTGGTACCAGAAGAAACCAGAGCAATCTCCTAAACTGCTGATATACGGGGCATCCAACCGATACATTGGGGTCCCTGATCGCTTCACAGGCAGTGGATCTGGGACAGATTTCACTCTGACCATCAGCAGTGTACAGGTTGAAGACCTCACACATTATTACTGTGCACAGTTTTACAGCTATCCTCT |
| IGKV11-125*01_B6 | C57BL/6J | Musmus IGKV11-125*01 F | 100 | 287 | GATGTCCAGATGATTCAGTCTCCATCCTCCCTGTCTGCATCTTTGGGAGACATAGTCACCATGACTTGCCAGGCAAGTCAGGGCACTAGCATTAATTTAAACTGGTTTCAGCAAAAACCAGGGAAAGCTCCTAAGCTCCTGATCTATGGTGCAAGCAACTTGGAAGATGGGGTCCCATCAAGGTTCAGTGGCAGTAGATATGGGACAGATTTCACTCTCACCATCAGCAGCCTGGAGGATGAAGATATGGCAACTTATTTCTGTCTACAGCATAGTTATCTCCCTCC |
| IGKV5-37*01_B6 | C57BL/6J | Musmus IGKV5-37*01 F | 100 | 287 | GACATCCTGCTGACCCAGTCTCCAGCCACCCTGTCTGTGACTCCAGGAGAAACAGTCAGTCTTTCCTGTAGGGCCAGCCAGAGTATTTACAAGAACCTACACTGGTATCAACAGAAATCACATCGGTCTCCAAGGCTTCTCATCAAGTATGCTTCTGATTCCATCTCTGGGATCCCCTCCAGGTTCACTGGCAGTGGATCAGGGACAGATTACACTCTCAGTATCAACAGTGTGAAGCCCGAAGATGAAGGAATATATTACTGTCTTCAAGGTTACAGCACACCTTC |

| | | | | | |
|---|---|---|---|---|---|
| IGKV1-88*01_B6 | C57BL/6J | Musmus IGKV1-88*01 F | 100 | 302 | GATGTTGTGGTGACTCAAACTCCACTCTCCCTGCCTGTCAGCTTTGGAGATCAAGTTTCTATCTCTTGCAGGTCTAGTCAGAGTCTTGCAAACAGTTATGGGAACACCTATTTGTCTTGGTACCTGCACAAGCCTGGCCAGTCTCCACAGCTCCTCATCTATGGGATTTCCAACAGATTTTCTGGGGTGCCAGACAGGTTCAGTGGCAGTGGTTCAGGGACAGATTTCACACTCAAGATCAGCACAATAAAGCCTGAGGACTTGGGAATGTATTACTGCTTACAAGGTACACATCAGCCTCC |
| IGKV4-73*01_S8836_B6 | C57BL/6J | Musmus IGKV4-73*01 ORF | 100 | 292 | CAAATTGTTCTCACCCAGTCTCCAGCAATCATGTCTGCATCTCCTGGGGAACGGGTCACCATGACCTGCAGTGCCAGCTCAAGTGTAAGTTCCAGCTACTTGTACTGGTACCAGCAGAAGTCAGGATCCTCCCCAAAACTCTGGATTTATAGCATATCCAACCTGGCTTCTGGAGTCCCAGCTCGCTTCAGTGGCAGTGGGTCTGGGACCTCTTACTCTCTCACAATCAACAGCATGGAGGCTGAAGATGCTGCCACTTATTACTGCCAGCAGTGGAGTAGTAACCCACTCA |
| IGKV4-52*01_S2975_B6 | C57BL/6J | Musmus IGKV4-52*01 ORF, or Musmus IGKV4-54*01 ORF | 100 | 286 | CAAATTGTTCTCACCCAGTCTCCAGCAATCCTGTCTGCATCTCCAGGGGAGAAGGTCACCATGACCTGCAGTGCCAGCTCAAGTGTAAGTTACATGTACAGGTACCAGCAGAAGCCAGGATCCTCACCCAAACCCTGGATTTATGGCACATCCAACCTGGCTTCTGGAGTCCCTGCTCGCTTCAGTGGCAGTGGATCTGGGACCTCTTATTCTCTCACAATCAGCAGCATGGAGGCTGAAGATGCTGCCACTTATTACTGCCAGCAATATCATAGTTACCCGCTCA |
| IGKV9-129*01_B6 | C57BL/6J | Musmus IGKV9-129*01 ORF | 100 | 287 | GACATCCAGATGACCCAGTCTCCATCCTCCTTATCTGCCTCTCTGGGAGAAAGAGTCAGTCTCACTTGCCGGGCAAGTCAGGACATTCATGGTTATTTAAACTTGTTTCAGCAGAAACCAGGTGAAACTATTAAACACCTGATCTATGAAACATCCAATTTAGATTCTGGTGTCCCAAAAAGGTTCAGTGGCAGTAGGTCTGGGTCAGATTATTCTCTCATTATCGGCAGCCTTGAGTCTGAAGATTTTGCAGACTATTACTGTCTACAATATGCTAGTTCTCCTCC |
| IGKV4-61*01_S2955_B6 | C57BL/6J | Musmus IGKV4-61*01 F | 100 | 286 | CAAATTGTTCTCACCCAGTCTCCAGCAATCATGTCTGCATCTCCAGGGGAGAAGGTCACCATATCCTGCAGTGCCAGCTCAAGTGTAAGTTACATGTACTGGTACCAGCAGAAGCCAGGATCCTCCCCCAAACCCTGGATTTATCGCACATCCAACCTGGCTTCTGGAGTCCCTGCTCGCTTCAGTGGCAGTGGGTCTGGGACCTCTTACTCTCTCACAATCAGCAGCATGGAGGCTGAAGATGCTGCCACTTATTACTGCCAGCAGTATCATAGTTACCCACTCA |
| IGKV4-71*01_B6 | C57BL/6J | Musmus IGKV4-71*01 F | 100 | 286 | CAAATTGTTCTCACCCAGTCTCCAGCAATCATGTCTGCATCTCCAGGGGAGAAGGTCACCATGACCTGCAGTGCCAGCTCAAGTGTAAGTTACATGCACTGGTACCAGCAGAAGCCAGGATCCTCCCCCAGACTCTGGATTTATTTAACATTCAACTTGGCTTCTGGAGTCCCTGCTCGCTTCAGTGGCAGTGGGTCTGGGACCTCTTACTCTCTCTCAATCAGCAGCATGGAGGCTGAAGATGCTGCCACTTATTACTGCCAGCAGTGGAGTAGTAACCCACCCA |
| IGKV4-51*01_B6 | C57BL/6J | Musmus IGKV4-51*01 F | 100 | 292 | GAAAATGTGCTCACCCAGTCTCCAGCAATAATGGCTGCCTCTCTGGGGGAGAAGGTCACCATGACCTGCAGTGCCAGCTCAAGTGTAAGTTCCAGCTACTTGCACTGGTACCAGCAGAAGTCAGGCACTTCCCCCAAACTCTGGATTTATGGCACATCCAACCTGGCTTCTGGAGTCCCAGCTCGCTTCAGTGGCAGTGGGGCTGGGATCTCTTACTCTCTCACAATCAGCAGCATGGAGGCTGAAAATGATGCAACTTATTACTGCCAGCAGTGGAGTGGTTACCCATTCA |
| IGKV4-78*01_B6 | C57BL/6J | Musmus IGKV4-78*01 F | 100 | 292 | CAAATTGTTCTCACCCAGTCTCCAGCAATCATGTCTGCATCTCCTGGGGAGAAGGTCACCATGACCTGCAGTGCCAGATCAAGTGTAAGTTCCAGCTACTTGTACTGGTACCAGCAGAAGCCAGGATCCTCCCCCAAACTCTGGATTTATAGCACATCCAACCTGGCTTCTGGAGTCCCTGCTCGCTTCAGTGGCAGTGGGTCTGGGACCTCTTATTCTCTCACAATCAGCAGCATGGAGGCTGAAGATGCTGCCACTTTTTACTGCCAGCAGTACAGTGGTTACCCATCCA |

| | | | | | |
|---|---|---|---|---|---|
| IGKV4-60*01_S1112_B6 | C57BL/6J | Musmus IGKV4-60*01 P | 100 | 292 | GAAATTGTGCTCACCCAGTCTCCAGCACTCATGGCTGCATCTCCAGGGGAGAAGGTCA<br>CCATCACCTGCAGTGTCAGCTCAAGTATAAGTTCCAGCTACTTGCACTGGTACCAGCA<br>GAAGTCAGGAATCTCCCCCAAACCCTGGATTTATGGCACATCCAACCTGGCTTCTGGA<br>GTCCCTGCTCGCTTCAGTGGCAGTGGATCTGGGACCTCTTACTCTCTCACAATCAGCAG<br>CATGGAGGCTGAAGATGCTGCCACTTATTACTGTCAACAGTGGAGTAGTTCCCCACTC<br>A |
| IGKV4-81*01_B6 | C57BL/6J | Musmus IGKV4-81*01 F | 100 | 292 | GAAAATGTGCTGACCCAGTCTCCAGCAATCATGGCTGCATCTCCAGGGGAGAAGGTC<br>ACCATGACCTGCAGTGCCAGCTCAAGTGTAAGTTCTAGTAACTTGCACTGGTACCAGC<br>AGAAGTCAGGCACTTCTACCAAATTCTGGATTTATAGGACATCCAACCTGGCTTCAGA<br>AGTCCCAGCTCCCTTCAGTGGCAGTGGGTCTGGGACCTCTTACTCTCTTACAATCAGCA<br>GCGTGGAGGCCGAAGATGCTGCCACTTATTACTGCCAGCAGTGGAGTGGTTACCCACC<br>CA |
| IGKV4-81*01_S0130_B6 | C57BL/6J | Musmus IGKV4-81*01 F | 100 | 292 | GAAAATGTGCTGACCCAGTCTCCAGCAATCATGGCTGCATCTCCAGGGGAGAAGGTC<br>ACCATGACCTGCAGTGCCAGCTCAAGTGTAAGTTCTAGTAACTTGCACTGGTACCAGC<br>AGAAGTCAGGCACTTCTACCAAATTCTGGATTTATAGGACATCCAACCTGGCTTCAGA<br>AGTCCCAGCTCCCTTCAGTGGCAGTGGGTCTGGGACCTCTTACTCTCTTACAATCAGCA<br>GCGTGGAGGCCGAAGATGCTGCCACTTATTACTGCCAGCAGTGGAGTGGTTACCCACT<br>CA |
| IGKV4-80*01_S4247_CAST | CAST/EiJ | Musmus IGKV4-80*01 F | 99.64 | 282 | CAAATTGTTCTCACCCAGTCTCCAGCAATCATGTCTGCATCTCTAGGGGAGGAGATCA<br>CCCTAACCTGCAGTGCCAGCTCGAGTGTAAGTTACATGCACTGGTTCCAGCAGAAGTC<br>AGGCACTTCTCCCAAACTCTTGATTTATAGCACATCCAACCTGGCTTCTGGAGTCCCTT<br>CTCGCTTCAGTGGCAGTGGGTCTGGGACCTTTTATTCTCTCACAATCAGCAGTGTGGA<br>GGCTGAAGATGCTGCCGATTATTACTGCCATCAGTGGAGTAGTTATCCA |
| IGKV4-86*01_S4316_CAST | CAST/EiJ | Musmus IGKV4-86*01 F | 100 | 283 | GAAATTGTGCTCACTCAGTCTCCAGCCATCACAGCTGCATCTCTGGGGCAAAAGGTCA<br>CCATCACCTGCAGTGCCAGCTCAAGTGTAAGTTACATGCACTGGTACCAGCAGAAGTC<br>AGGCACCTCCCCCAAACCATGGATTTATGAAATATCCAAACTGGCTTCTGGAGTCCCA<br>GCTCGCTTCAGTGGCAGTGGGTCTGGGACCTCTTACTCTCTCACAATCAGCAGCATGG<br>AGGCTGAAGATGCTGCCATTTATTACTGCCAGCAGTGGAATTATCCACTTA |
| IGKV8-16*01_S7495_CAST | CAST/EiJ | Musmus IGKV8-16*01 F | 99.33 | 305 | GAAATTGTGTTGACGCAGTCTATACCATCCCTGACTGTGTCAGCAGGAGAGAGGGTCA<br>CTATCAGCTGCAAATCCAATCAGAATCTTTTATGGAGTGGAAACCAAAGGTACTGTTT<br>GGTCTGGCACCAGTGGAAACCAGGGCAAACTCCTACACCGTTGATCACCTGGACATCT<br>GATAGATACTCTGGAGTCCCTGATCGTTTCATAGGCAGTGGATCTGTGACAGATTTCA<br>CTCTGACCATCAGCAGTGTGCAGGCTGAAGATGTGGCAGTTTATTTCTGTCAGCAGCA<br>TTTACACATTCCTCC |
| IGKV4-59*01_S3253_CAST | CAST/EiJ | Musmus IGKV4-59*01 F | 100 | 286 | CAAATTGTTCTCACCCAGTCTCCAGCAATCATGTCTGCATCTCCAGGGGAGAAGGTCA<br>CCATGACCTGCAGTGCCAGCTCAAGTGTAAGTTACATGCACTGGTACCAGCAGAAGTC<br>AGGCACCTCCCCCAAAAGATGGATTTATGACACATCCAAACTGGCTTCTGGAGTCCCT<br>GCTCGCTTCAGTGGCAGTGGGTCTGGGACCTCTTACTCTCTCACAATCAGCAGCATGG<br>AGGCTGAAGATGCTGCCACTTATTACTGCCAGCAGTGGAGTAGTTACCCACTCA |
| IGKV6-20*01_S2082_CAST | CAST/EiJ | Musmus IGKV6-20*01 F | 99.64 | 287 | AACATTGTAATGACCCAATCTCCCAAATCCATGTCCATGTCAGTAGGAGAGAGGGTCA<br>CCTTGAGCTGCAAGGCCAGTGAGAATGTGGGTACTTATGTATCCTGGTATCAACAGAA<br>ACCAGAGCAGTCTCCTAAACTGCTGATATACGGGGCATCCAACCGGTACACTGGGGTC<br>CCCGATCGCTTCACAGGCAGTGGATCTGCAACAGATTTCACTCTGACCATCAGCAGTG<br>TGCAGGCTGAAGACCTTGCAGATTATTACTGTGGACAGAGTTACAGCTATCCTCC |

| | | | | | |
|---|---|---|---|---|---|
| IGKV12-46*01_S2465_CAST | CAST/EiJ | Musmus IGKV12-46*01 F | 97.13 | 287 | GACATCCAGATGACTCAGTCTCCAGCCTCCCTATCTGCATCTGTAGGAGAAACTGTCACCATCACATGTCGAGCAAGTGAGAATATTTACAGTTATTTAGCCTGGTATCAGCAGAAACAGGGAAAATCTCCTCAGCTCCTGGTCTATGCTGCAACAAACTTAGCAGATGGTGTGCCATCAAGGTTCAGTGGCAGTGGATCAGGCACACAGTATTCCCTCAAGATCAACAGCCTGCAGCCTGAAGATTTTGGGAGTTATTACTGTCAACATTATTATGGTACTCCTCC |
| IGKV2-137*01_S5808_CAST | CAST/EiJ | Musmus IGKV2-137*01 F | 94.56 | 302 | GATATTGTGATGACTCAGGCTGCACCCTCTGTACCTGTCACTACTGGAGAGTCAGTATCCATTTCTTGCAAGTCTAGTAAGAGTCTTCTGCATAGTAATGGCATCACTTATTTGTATTGGTTCCTACAGAGGCCAGGCCAGTCTCCTCAGCGCCTGATATATTATATGTCCAACCTTGCCTCAGGAGTCCCAGACAGGTTCAGTGGCAGAGGGTCAGGAACTGATTTCACACTGAGAATCAGTAGAGTGGAGGCTGAGGATGTGGGTGTTTATTACTGTATGCAAAGTCTAGAATATCCTTT |
| IGKV6-b*01_CAST | CAST/EiJ | Musmus IGKV6-b*01 F | 100 | 287 | AGTATTGTGATGACCCAGACTCCCAAATTCCTGCCTGTATCAGCAGGAGACAGGGTTACCATGACCTGCAAGGCCAGTCAGAGTGTGGGTAATAATGTAGCCTGGTACCAACAGAAGCCAGGACAGTCTCCTAAACTGCTGATATACTATGCATCCAATCGCTACACTGGAGTCCCTGATCGCTTCACTGGCAGTGGATCTGGGACAGATTTCACTTTCACCATCAGCAGTGTGCAGGTTGAAGACCTGGCAGTTTATTTCTGTCAGCAGCATTATAGCTCTCCTCC |
| IGKV10-94*03_CAST | CAST/EiJ | Musmus IGKV10-94*03 F, or Musmus IGKV10-94*08 F | 100 | 287 | GATATCCAGATGACACAGACTACATCCTCCCTGTCTGCCTCTCTGGGAGACAGAGTCACCATCAGTTGCAGGGCAAGTCAGGATATTAGCAATTATTTAAACTGGTATCAGCAGAAACCAGATGGAACTGTTAAACTCCTGATCTACTACACATCAAGATTACACTCAGGAGTCCCATCAAGGTTCAGTGGCAGTGGGTCTGGGACAGATTATTCTCTCACCATCAGCAACCTGGAACCTGAAGATATTGCCACTTACTATTGTCAGCAGTATAGTAAGCTTCCTCC |
| IGKV6-15*01_S4042_CAST | CAST/EiJ | Musmus IGKV6-15*01 F | 97.13 | 287 | GACATTGTGATGACCCAGTCTCAAAAATTCATGTCCACATCAGTAGGAGACAGGGTCAGCGTCACCTGCAAGGCCAGTCAGTATGTGGGTACTTATGTAGCCTGGTATCAACAGAAACCAGGGCAATCTCCTAAAGCACTGATTTACTCGGCATCCACCCGGCACACTGGAGTCCCTGATCGCTTCACAGGCAGTGGATCTGGGACAGATTTCACTCTCACCATTAGCAATGTGCAGTCTGAAGACTTGGCAGAGTATTTCTGTCAGCAATATAGCAGCTCTCCTCT |
| IGKV8-21*01_S1419_CAST | CAST/EiJ | Musmus IGKV8-21*01 F | 99.33 | 302 | GACATTGTGATGTCACAGTCTCCATCCTCCCTGGCTGTGTCAGCAGGAGAGAAGGTCACTATGAGCTGCAAATCCAGTCAGAGTCTGTTCAACAGTAGAACCCGAAAGAACTACTTGGCTTGGTACCAGCAGAAACCAGGGCAGTCTCCTAAACTGCTGATCTACTGGGCATCCACTAGGGAATCTGGGGTCCCTGATCGCTTCACAGGCAGTGGATCTGGGACAGATTTCACTCTCACCATCAGCAGTGTGCAGGCTGAAGACCTGGCAGTTTATTACTGCAAGCAATCTTATTATCTTCC |
| IGKV15-103*01_S2403_CAST | CAST/EiJ | Musmus IGKV15-103*01 ORF | 98.21 | 287 | GACATCCAGATGAACCAGTCTCCATCCAGTCTGTCTGCATCCCTCGGAGACACAATTACCATCACTTGCCGTGCCAGTCAGAACATTAATATTTGGTTAAGCTGGTACCAGCAGAAACCAGGAAATATTCCTAAACTATTGATCTATAAGGCTTCCAACTTGCACACAGGCGTCCCATCAAGGTTTAGTGGCAGTGGATCTGGAACAGATTTCACATTAACCATCAGCAGCCTGCAGCCTGAAGACATTGCCACTTACTACTGTCTACAGGGTCAAAGTTATCCTCT |
| IGKV1-110*02_S9753_CAST | CAST/EiJ | Musmus IGKV1-110*02 F | 97.28 | 302 | GATGTTGTGATGACCCAAACTCCACTCTCCCTGCCTGTCAGTCTTGGAGATCAAGCCTCCATCTCTTGCAGATCTAGTCAGAGCATTGTAGCCAGTAATGGAAACACCTATTTATATTGGTACCTGCAGAAGCCAGGCCAGTCTCCAAAGCTCCTGATCTACAGGGTTTCCAACAGATTTTCTGGGGTCCCAGACAGGTTCAGTGGCAGTGGATCAGGGACAGATTTCACTCTCAAGATCAGCAGAGTGGAGGCTGAGGATCTGGGAGTTTATTACTGCTTTCAAGGTACACATGTTCCTCA |

| | | | | | |
|---|---|---|---|---|---|
| IGKV12-44*01_S1769_CAST | CAST/EiJ | Musmus IGKV12-44*01 F | 97.13 | 287 | GACATCCAGATGACTCAGTCTCCAGCTTCCCTATCTGCATCTGTGGGAGAAACTGTCACCATCACATGTCGAGCAAGTGAGAATATTTACAGTAATTTAGCCTGGTATCAGCAGAAACAGGGAAAATCTCCTCAGCTCCTGGTCTATAATGCAAAAACCTTAGCAGAAGGTGTGCCATCAAGGTTCAGTGGCAGTGGATCAGGAACACAATATTCTCTCAAGATCAACAGCCTGCAGCCTGAAGATTTTGGGAGTTATTACTGTCAACATTATTATGGTACTCCTCC |
| IGKV2-137*01_S2608_CAST | CAST/EiJ | Musmus IGKV2-137*01 F | 94.22 | 302 | GATATTGTGATGACTCAGGCTGCACCCTCTGTACCTGTCACTCCTGGAGAGTCAGTATCCATCTCCTGCAGGTCTAGTAAGAGTCTCCTGCACAGTAGTGGCAAGCATAGGTTGTATTGGTTCCTACAGAGGCCAGGCCAATCTCCTCAGCGCCTGATATATTATATGTCCAACCTTGCCTCAGGAGTCCCAGACAGGTTCAGTGGCAGTGGGTCAGGAACTGATTTCACACTGAGAATCAGTAGAGTGGAGGCTGAGGATGTGGGTGTTTATTACTGTATGCAAAGTCTAGAATATCCTTT |
| IGKV4-69*01_S2827_CAST | CAST/EiJ | Musmus IGKV4-69*01 F | 97.1 | 282 | CAAATTCTTCTCACCCAGTCTCCAGCAATCATGTCTGCATCTCCAGGAGAGAAGGTCACCATGACCTGCAGTGCCAGCTCAAGTGTAAGTTATATGCACTGGTACCAGCAGAAGCCAGGATCCTCGCCCAAACCCTGGATTTATGGCACATCCAACCTGGCTTCTGGAGTCCCTGTTCGCTTCAGTGGCAGTGGGTCTGGGACCTCTTACTCTCTCACAATCAGCAGCATGGAGGCTGAAGATGCTGCCACTTATTACTGCCAGCAGTGGAGTAGTTACCCA |
| IGKV12-40*01_S2386_CAST | CAST/EiJ | Musmus IGKV12-40*01 P | 97.49 | 287 | GACATCCAGATGACTCAGTCTCCAGCCTCCCTGTCTGCATCTGTGGGAGAAACTGTCACCATCACATGTCGAGCAAGTGAGAATATTGCCAGTAATTTAGCATGGTATCAGCAGAAACAGGGAAAATCTCCTCAGCTCCTGGTCTATGATGCGAGAAACTTAGCAGATGGTGTGCCATCAAGGTTCAGTGGCAGTGGATCAGGCACACAGTATTCTCTCAATATCAACAGCCTGCAGTCTGAAGATGTTGCGAGATATTACTGTCAACATTATTATGGTACTCCTCC |
| IGKV8-19*01_CAST | CAST/EiJ | Musmus IGKV8-19*01 F | 100 | 305 | GACATTGTGATGACACAGTCTCCATCCTCCCTGACTGTGACAGCAGGAGAGAAGGTCACTATGAGCTGCAAGTCCAGTCAGAGTCTGTTAAACAGTGGAAATCAAAAGAACTACTTGACCTGGTACCAGCAGAAACCAGGGCAGCCTCCTAAACTGTTGATCTACTGGGCATCCACTAGGGAATCTGGGGTCCCTGATCGCTTCACAGGCAGTGGATCTGGAACAGATTTCACTCTCACCATCAGCAGTGTGCAGGCTGAAGACCTGGCAGTTTATTACTGTCAGAATGATTATAGTTATCCTCC |
| IGKV3-12*01_S3040_CAST | CAST/EiJ | Musmus IGKV3-12*01 F | 99.31 | 299 | GACATTGTGCTGACACAGTCTCCTGCTTCCTTAGCTGTATCTCTGGGGCAGAGGGCCACCATCTCATGCAGGGCCAGCAAAAGTGTCAGTACATCTAGCTATAGTTATATGCACTGGTACCAACAGAAACCAGGACAGCCACCCAAACTCCTCATCTATCGTGCATCCAACCTAGAATCTGGGGTCCCTGCCAGGTTCAGTGGCAGTGGGTCTGGGACAGACTTCACCCTCAACATCCATCCTGTGGAGGAGGAGGATGCTGCAACCTATTACTGTCAGCACAGTAGGGAGCTTCCTCC |
| IGKV4-77*01_S4503_CAST | CAST/EiJ | Musmus IGKV4-77*01 P | 97.83 | 286 | CAAATTGTTCTCACCCAGTCTCCAGCAATCCTGACTGCATCTCCAGGGGAGAAGGTCACCATGACCTGCAGTGCCAGCTCAAGTGTAAGTTACATGTACTGGTTCCAGCAGAAGCCAGGATCCTCCCCCAAACTCTGGATTTATAGCATATCCAACCTGGCTTCTGGAGTCCCTGCTCGCTTCAGTGGCAGTGGGTCTGGGACCTCTTACTCTCTTACAATCAGCAGTGTGAAGGCTGAAGATGCTGCCACTTATTACTGCCAGCAATGGAGTAGTTCCCCACCCA |
| IGKV1-110*01_S6010_CAST | CAST/EiJ | Musmus IGKV1-110*01 F | 98.3 | 301 | GATGTTGTGATGACCCAAACTCCACTCTCCCTGCCTGTCAGTCTTGGAGATCAAGCCTCCATCTCTTGCAGATCTAGTCAGAGCCTTGTACACAGTAATGGAAACACCTATTTACATTGGTACCTGCAGAAACCAGGCCAGTCTCCAAAGCTCCTGATCTACAGGGTTTCCAACCGATTTTCTGGGGTCCCAGACAGGTTCAGTGGCAGTGGATCAGGGACAGATTTCACACTCAAGATCAGCAGAGTGGAGGCTGAGGATCTGGGAGTTTATTACTGCTTTCAAAGTACACATGTTCCTC |

| | | | | | |
|---|---|---|---|---|---|
| IGKV1-117*01_S1348_CAST | CAST/EiJ | Musmus IGKV1-117*01 F | 99.32 | 301 | GATGTTGTGATGACCCAAACTCCACTCTCCCTGCCTGTCAGTCTTGGAGATCAAGCCTCCATCTCTTGCAGATCTAGTCAGAGCATTGTACATAGTAATGGAAACACCTATTTAGAATGGTACCTGCAGAAGCCAGGCCAGTCTCCAAAGCTCCTGATCTACAAAGTTTCCAACCGATTTTCTGGGGTCCCAGACAGGTTCAGTGGCAGTGGATCAGGGACAGATTTCACACTCAAGATCAGCAGAGTGGAGGCTGAGGATCTGGGAGTTTATTACTGCTTTCAAGGTTCACATGTTCCTC |
| IGKV4-58*01_S3374_CAST | CAST/EiJ | Musmus IGKV4-58*01 F | 99.29 | 292 | GAAAATGTGCTCACCCAGTCTCCAGCAATAATGGCTGCCTCTCTGGGGGAGAAGGTCACCATGACCTGCAGTGCCAGCTCAAGTGTAAGTTCCAGTTACTTGCACTGGTACCAGCAGAAGTCAGGCGCTTCCCCCAAACCCTTGATTCATAGGACATCCAACCTGGCTTCTGGTGTCCCAGCTCGCTTCAGTGGCAGTGGGTCTGGGACCTCTTACTCTCTCACAATCAGCAGCGTGGAGGCTGAAGATGATGCAACTTATTACTGCCAGCAGTGGAGTGGTTACCCATTCA |
| IGKV4-68*01_S2180_CAST | CAST/EiJ | Musmus IGKV4-68*01 F | 97.83 | 286 | CAAATTGTTCTCACCCAGTCTCCAGCAATCATGTCTGCATCTCCAGGGGAGAAGGTCACCATGACCTGCAGTGCCAGCTCAAGTGTAAGTTACATGTACTGGTACCAGCAGAAGCCAGGATCCTCACCCAAACCCTGGATTTATCGCACATCCAACCTGGCTTCTGGAGTCCCAACTCGCTTCAGTGGCAGTGGGTCTGGGACCTCTTACTCTCTCACAATCAGCAGCATGGAGGCTGAAGATGCTGCCACTTATTACTGCCAGCAGTGGAGTAGTAACCCACTCA |
| IGKV5-43*01_S4957_CAST | CAST/EiJ | Musmus IGKV5-43*01 F | 98.57 | 287 | GATATTGTGCTAACTCAGTCTCCAGCCACCCTGTCTGTGATTCCAGGAGATAGCGTCAGTCTTTCCTGCAGGGCGAGCCAAAGTATTAGCAACAACCTACACTGGTATCAACAAAAACACATGAGTCTCCAAGGCTTCTCATCAAGTATGCTTCCCAATCCATCTCTGGGATCCCCTCCAGGTTCAGTGGCAGTGGATCAGGGACAGATTTCACTCTCAGTATCAACAGTGTGGAGACTGAAGATTTTGGAATGTATTTCTGTCAACAGAGTAACAGCTGGCCTCA |
| IGKV1-135*01_S9254_CAST | CAST/EiJ | Musmus IGKV1-135*01 F | 95.58 | 302 | GATGTTGTGATGACTCAGACCCCAATCACTATGTCGGTTACCATTGGACAACCAGCCTCCATCTCTTGCAAGTCAAGTCAGAGCCTCTTACATAGTAATGGAAAGACATATTTGAATTGGTTATTACAGAGGCCAGGCCAGTCTCCAAAGCTCCTAATCTATCTGGTGTCTAAACTGGAATCTGGAGTCCCTGACAGGTTCAGTGGCAGTGGATCAGGGACAGATTTCACACTGAAAATCAGCAGAGTGGAGGCTGAGGATTTGGGAGTTTATTACTGCTTGCAAGGTACATATTATCCTCA |
| IGKV11-125*01_S0249_CAST | CAST/EiJ | Musmus IGKV11-106*02 F | 96.77 | 287 | GATGTTCAAATGACCCAGTCTCCATCCTCCCTGTCTGCATCTTTGGGAGAGAGAGTCTCCCTGACTTGCCAGGCAAGTCAGGGCATTAGCAATTATTTAAACTGGTATCAGCAAACACCAGGGAAAGCTCCTAGGCTCTTGATCTATGGTGCAAGCAACTTGGAAGATGGGGTCCCTTCAAGGTTCAGTGGCACTGGATATGGGACAGATTTCACTTTCACCATCAGCAGCCTGGAGGAAGAAGATGTGGCAACTTATTTCTGTCTACAGCATGGTTATCTCCCTCC |
| IGKV19-93*01_S8964_CAST | CAST/EiJ | Musmus IGKV19-93*01 F | 98.21 | 287 | GACATCCAGATGACACAGTCTCCATCCTCACTGTCTGCATCTCTGGGAGGCAAAGTCACCATCACTTGCAAGGCAAGCCAAGACATTAACAAGAATATAGCTTGGTACCAACACAAGCCTGGAAAAGGTCCTAGGCTGCTCATATGGTACACATCTACATTACAGCCAGGCATCCCATCAAGGTTCAGTGGAAGTGGGTCTGGGAGAGATTATTCCTTCAGCATCAGCAACCTGGAGCCTGAAGATATTGCAACTTATTACTGTCTACAGTATGATAATCTTCCACC |
| IGKV3-1*01_S6564_CAST | CAST/EiJ | Musmus IGKV3-1*01 F | 95.88 | 299 | GACATTGTGCTGACCCAATCTCCAGCTTCTTTGGCTGTGTCTCTAGGGCAGAGAGCCACCATCTCCTGCAGAACCAGTGAAAGTGTTGATTATGATGGCGATAGTTATATGAACTGGTACCAACAGAAACCAGGACAGCCACCCAAACTCCTCATCTATGGTGCATCCAACGTAGAATCTGGGGTCCCTGCCAGGTTTAGTGGCAGTGGGTCTGGGACAGACTTCAGCCTCAACATCCATCCTGTGGAGGAGGATGATATTGCAATGTATTTCTGTCAGCAAAGTAGGAAGGTTCCTTC |

| | | | | | |
|---|---|---|---|---|---|
| IGKV4-74*01_S3773_CAST | CAST/EiJ | Musmus IGKV4-74*01 F | 95.04 | 292 | CAAATTGTTCTCACCCAGTCTCCAGCAATCATGTCTGCATCTCCAGGGGAGAAGGTCACCATGACCTGCAGGGCCAGCTCAAGTGTAAGTTCCAGTTACTTGCACTGGTACCAGCAGAAGCCAGGATCTTCCCCCAAACTCTGGATTTATAGCACATCCAACCTGGCTTCAGGAGTCCCAGCTCGCTTCAGTGGCAGTGGGTCTGGGACCTCTTACTCTCTCACAATCAGCAGTGTGGAGGCTGAGGATGCTGCCACTTATTACTGCCAGCAGTATGATAGTTCCCCATCCA |
| IGKV8-30*01_S1609_CAST | CAST/EiJ | Musmus IGKV8-30*01 F | 95.61 | 296 | GACATTGTGATGTCACAGTCTCCATCCGCCCTAGCTGTGTCAGTTGGAGAGAAGGTCACTATGAGCTGCAAGTCCAGTCAGAGCCTTTTATATAGTAGCAATCAAAAGAACTACTTGGCCTGGTACCAGCAGAAACCAGGGCAGTCTCCTAAACTGTTAATCTACTGGGCATCCACTAGGGAATCTGGGGTCCCTGACCGCTTCACAGGCAGTAGATCAGGGACAGATTTCACTCTCACCATCAGCAGTGTGCAGGCTGAAGACCTGGCCGTTTATTACTGCAAGCAATCTTATAG |
| IGKV4-50*01_S6254_CAST | CAST/EiJ | Musmus IGKV4-50*01 F | 97.83 | 286 | GAAAATGTGCTCACCCAGTCTCCAGCAATCATGTCTGCATCTCTAGGGGAGAAGGTCACCATGAGCTGCAGGGCCAGCTCAAGAGTAAATTACATGTACTGGTACCAGCAGAAGTCAGGTGCCTCCCCCAAACTATGGATTTATTACACATCCAACCTGGCTCCTGGAGTCCCAGCTCGCTTCAGTGGCAGTGGGTCTGGGACCTCTTATTCTCTCACAATCAGCAGCGTGGAGGCTGAAGATGCTGCCACTTATTACTGCCAGCAGTTAACTAGTTCCCCATCCA |
| IGKV8-24*01_S8843_CAST | CAST/EiJ | Musmus IGKV8-24*01 F | 98.32 | 305 | GACATTGTGATGACACAGTCTCCATCCTCCCTGGCTATGTCAGTAGGACAGAAGGTCACTATGAGCTGCAAGTCCAGTCAGAGCCTTTTAAGTAGTAGCAATCAAAAGAACTATTTGGCCTGGTACCAGCAGAAACCAGGACAGTCTCCTAAACTTCTGGTATACTTTGCATCCACTAGGGAATCTGGGGTCCCTGATCGCTTCATAGGCAGTGGATCTGGGACAGATTTCACTCTTACCATCAGCAGTGTGCAGGCTGAAGACCTGGCACTTTATTACTGTCAGCAACATTATAGCACTCCTCC |
| IGKV4-53*01_S1762_CAST | CAST/EiJ | Musmus IGKV4-53*01 F | 99.65 | 292 | GAAATTGTGCTCACCCAGTCTCCAGCACTCATGGCTGCATCTCCAGGGGAGAAGGTCACCATCACCTGCAGTGTCAGCTCAAGTATAAGTTCCAGCAACTTGCACTGGTACCAGCAGAAGTCAGGAACCTCCCCCAAACCCTGGATTTATGGCACATCCAACCTGGCTTCTGGAGTCCCTGTTCGCTTCAGTGGCAGTGGATCTGGGACCTCTTATTCTCTCACAATCAGCAGCATGGAGGCTGAAGATGCTGCCACTTATTACTGTCAACAGTGGAGTAGTTACCCACTCA |
| IGKV8-34*01_S3615_CAST | CAST/EiJ | Musmus IGKV8-34*01 F | 99.33 | 305 | GACATTTTGATGACTCAGTCTCCATCCTCCCTGACTGTGTCAGCAGGAGAGAAGGTCACTATGAGCTGCAAGTCCAGTCAGAGTCTTTTAGCTAGTGCCAACCAAAATAACTACTTGGCCTGGCACCAGCAGAAACCAGGACGATCTCCTAAAATGCTGATAATTTGGGCATCCACTAGGGTATCTGGAGTCCCTGATCGCTTCATAGGCAGTGGATCTGGGACGGATTTCACTCTGACCATCAACAGTGTGCAGGCTGAAGATCTGGCTGTTTATTACTGTCAGCAGTCCTACAGCGCTCCTAC |
| IGKV4-68*01_S6732_CAST | CAST/EiJ | Musmus IGKV4-68*01 F | 97.83 | 286 | CAAATTGTTCTCACCCAGTCTCCAGCAATCATGTCTGCATCTCCAGGGGAGAAGGTCACCATGACCTGCAGTGCCAGCTCAAGTGTAAGTTACATGTACTGGTACCAGCAGAAGCCAGGATCCTCACCCAAACCCTGGATTTATCGCACATCCAACCTGGCTTCTGGAGTCCCAACTCGCTTCAGTGGCAGTGGGTCTGGGACCTCTTACTCTCTCACAATCAGCAGCATGGAGGCTGAAGATGCTGCCACTTATTACTGCCAGCAGTGGAGTAGTAACCCACCCA |

| | | | | | |
|---|---|---|---|---|---|
| IGKV3-7*01_S8881_CAST | CAST/EiJ | Musmus IGKV3-7*02 F | 99.66 | 299 | GACATTGTGCTGACACAGTCTCCTGCTTCCTTAGCTGTATCTCTGGGGCAGAGGGCCACCATCTCATGCAGGGCCAGCCAAAGTGTCAGTACATCTAGCTATAGTTATATGCACTGGTACCAACAGAAACCAGGACAGCCACCCAAACTCCTCATCAGGTATGCATCCAACCTAGAATCTGGGGTCCCTGCCAGGTTCAGTGGCAGTGGGTCTGGGACAGACTTCACCCTCAACATCCATCCTGTGGAGGAGGAGGATACTGCAACATATTACTGTCAGCACAGTTGGGAGATTCCTCC |
| IGKV4-55*01_S5216_CAST | CAST/EiJ | Musmus IGKV4-55*01 F | 97.46 | 286 | CAAATTGTTCTCACCCAGTCTCCAGCAATCATGTCTGCATCTCCAGGGGAGAAGGTCACCATGACCTGCAGTGCCAGCTCAAGTGTAAGTTACATGTACTGGTACCAGCAGAAGCCAGGATCCTCCCCCAGACTCTGGATTTATGACACATCCAACCTGGCTTCTGGAGTCCCCGCTCGCTTCAGTGGCAGTGGGTCTGGGACCTCTTATTCTCTCACAATCAGCAGCATGGAGGCTGAAGATGCTGCCACTTATTACTGCCAGCAGTGGAGTAGTTACCCACTCA |
| IGKV17-127*01_S4378_CAST | CAST/EiJ | Musmus IGKV17-127*01 F | 99.64 | 287 | GAAACAACTGTGACCCAGTCTCCAGCATCCCTGTCCGTGGCTACAGGAGAAAAAGTCACTATCAGATGCATAACCAACACTGATATTGATGATGATATGAACTGGTACCAGCAGAAGCCAGGGGAACCTCCTAAGCTCCTTATTTCAGAAGGCAATACTCTTCGTCCTGGAGTCCCATCCCGATTCTCCAGCAGTGGCTATGGCACAGATTTTGTTTTTACAATTGAAAACACGCTCTCAGAAGATGTTGCAGATTACTACTGTTTGCAAAGTGATAACATGCCTCT |
| IGKV14-100*01_S6933_CAST | CAST/EiJ | Musmus IGKV14-100*01 F | 99.64 | 287 | GACATCCTGATGACCCAATCTCCATCCTCCATGTCTGTATCTCTGGGAGACACAGTCAGCATCACTTGCCATGCAAGTCAGGGCATTAGCAGTAATATAGGGTGGTTGCAGCAGAAACCAGGGAAATCATTTAAGGGCCTGATCTATCATGGAACCAACTTGGAAGATGGAGTTCCATCAAGGTTCAGTGGCAGTGGATCTGGAGCAGATTATTCTCTCACCATCAGCAGCCTGGAATCTGAGGATTTTGCAGACTATTACTGTGTACAGTATGCTCAGTTTCCTCC |
| IGKV3-4*01_S3446_CAST | CAST/EiJ | Musmus IGKV3-4*01 F | 92.44 | 299 | GACATTGTGCTGACCCAATCTCCAGCTTCTTTGGCTGTGTCTATAGGGCAGAGGGCCACCATATCCTGCAAGGCCAGCCAGAGTATCAATTATTACGGTGATAATTATATGCACTGGTTCCAACAGAAACCAGGACAGCCAACCAAACTCCTTATCTATGATGCATCCAACCTAGAATCTGGAGTTCCTGACAGGTTCAGTGGCAGTGGGTCTGGGACAGACTTCACCCTCAACATCCATCCTGTGGAGGAGGAGGATGCTGCAACCTATTACTGTCAGCAAAGTAAGGAGTTTCCTCC |
| IGKV4-90*01_S5868_CAST | CAST/EiJ | Musmus IGKV4-90*01 F | 99.28 | 285 | GAAATTTTGCTCACCCAGTCTCCAGCAATCATAGCTGCATCTCCTGGGGAGAAGGTCACCATCACCTGCAGTGCCAGCTCAAGTGTAAGTTACATGCACTGGTACCAGCAGAAACCAGGATCCTCTCCCAAAATATGGATTTATGGTATATCCAACCTGGCTTCTGGAGTTCCTGCTCGCTTCAGTGGCAGTGGGTCTGGGACATCTTTCTCTTTCACAATCAACAGCATGGAGGCTGAAGATGTTGCCACTTATTACTGTCAGCAAAGGAGTAGTTACCCACCC |
| IGKV13-85*01_S1266_CAST | CAST/EiJ | Musmus IGKV13-85*01 F | 97.13 | 287 | GACATCCAGATGACACAATCTTCATCCTCCTTGTCTGTATCTCTAGGAGACAGAGTCACCATTACTTGCAAGGCAAGTGAGCACATTAATAGTTGGTTAGCCTGGTATCAGCAAAAACCAGGAAATGCTCCTAGGCTCTTAATATCTGGTGCAACCAGTTTGGAAACTGGGGTTCCTTCAAGATTCAGTGGCAGTGCATCTGGAAAGGATTACACTCTCAGCATTACTAGTCTTCAGACTGAAGATGTTGCTACTTATTACTGTCAACAGTATTGGGGTACTCCTCC |
| IGKV4-56*01_S2201_CAST | CAST/EiJ | Musmus IGKV4-56*01 P | 96.38 | 286 | CAAATTGTTCTCACCCAGTCTCCAGCAATCATGTCTGCATCTCCGGGGCAGAAAGTCACCATAACCTGCAGTGCCAGCTCAAGTGTAAATTACATGCACTGGTACCAGCAGAAGCCAGGATCCTCCCCCAAACTCTGGATTTATGACACATCCAAACTGGCTCCTGGAGTCCCTGCTCGCTTCAGTGGCAGTGGGTCTGGGACCTCTTACTCTCTCACAATCAGCAGCATGGAGGCTGAAGATGCTGCCTCTTATTTCTGCCATCAGTGGAGTAGTTACCCACCCA |

| | | | | | |
|---|---|---|---|---|---|
| IGKV4-78*01_S3637_CAST | CAST/EiJ | Musmus IGKV4-78*01 F | 98.23 | 292 | CAAATTGTTCTCACCCAGTCTCCAGCAATCATGTCTGCATCTCCTGGGGAACGGGTCACCATGACCTGCAGTGCCAGCTCAAGTGTAAGTTCCAGCTACTTGTACTGGTACCAGCAGAAGCCAGGATCCTCCCCCAAACTCTGGATTTATAGCACATCCAACCTGGCTTCTGGAGTCCCTGCTCGCTTCAGTGGCAGTGGGTCTGGGACCTCTTATTCTCTCACAATCAGCAGCATGGAGGCTGAAGATGCTGCCACTTATTACTGCCAGCAGTACAGTGGTTACCCATTCA |
| IGKV5-48*01_S4608_CAST | CAST/EiJ | Musmus IGKV5-48*01 F | 97.13 | 287 | GATATCTTGCTGACTCAGTCTCCAGCCATCCTGTCTGTGACTCTAGGAGAAAGAGTCAGTTTCTCCTGCAGGGCCAGTCAGAGCATTGGCACAAGCATACACTGGTATCAGCAAAGAACAAATGGTTCTCCAAGGCTTCTCATAAAGTATGCTTCTGAGTCTATCTCTGGGATCCCATCCAGGTTTAGTGGCAGTGGATCAGGGACAGATTTCACTCTTAGCATCAACAGTGTGGAGTCTGAAGATGTTGCAGATTATTACTGTCAACAGAGTTATAGCTGGCCAAC |
| IGKV2-109*03_S0318_CAST | CAST/EiJ | Musmus IGKV2-109*03 F | 99.32 | 302 | GATATTGTGATGACACAGGCTGCATTCTCCAATCCAGTCACTCTTGGAACATCAGCTTCCATCTCTTGCAGTTCTAGTAAGAGTCTCCTACATAGTAATGGCATCACTTATTTGTATTGGTATCTGCAGAGGCCAGGCCAGTCTCCTCAGCTCCTGATATATCGGATGTCCAACCTTGCCTCAGGAGTCCCAGACAGGTTCAGTGGCAGAGGGTCAGGAACTGATTTCACACTGAGAATCAGCAGAGTGGAGGCTGAGGATGTGGGTGTTTATTACTGTGCTCAAATGCTAGAACGCCCTCC |
| IGKV9-124*01_S2023_CAST | CAST/EiJ | Musmus IGKV9-124*01 F | 98.92 | 285 | GACATCCAGATGACCCAGTCTCCATCCTCCTTATCTGCCTCTCTGGGAGAAAGAGTCAGTCTCACTTGTCGGGCAAGTCAGGAAATTAGTGGTTACTTAAGCTGGCTTCAGCAGAAACCAGATGGAACTATTAAACGCCTGATCTACAGCACATCCACTTTAGATTCTGGTGTCCCAAAAAGGTTCAGTGGCAGTAGGTCTGGGTCAGATTATTCTCTCACCATCAGCAGCCTTGAGTCTGAAGATTTTGCAGACTATTACTGTCTACAATATGCTAGTTCTCCT |
| IGKV11-125*01_S0413_CAST | CAST/EiJ | Musmus IGKV11-106*02 F | 97.13 | 287 | GATGTTCAAATGACCCAGTCTCCATCCTCCCTGTCTGCATCTTTGGGAGAGAGAGTCTCCCTGACTTGCCAGGCAAGTCAGAGCGTTAGCAATAATTTAAACTGGTATCAGCAAACACCAGGGAAAGCTCCTAGGCTCTTGATCTATGGTGCAAGCAAATTGGAAGATGGGGTCCCTTCAAGGTTCAGTGGCACTGGATATGGGACAGATTTCACTTTCACCATCAGCAGCCTGGAGGAAGAAGATGTGGCAACTTATTTCTGCCTACAGCATAGGTATCTCCCTCC |
| IGKV1-115*01_S3088_CAST | CAST/EiJ | Musmus IGKV1-115*01 P | 95.24 | 302 | GATGTTCTGTTGACCCAAACTCCACTTTTCCTGCATGTCAGCCTTGGAGATCAAGCCTCTATCTCTTGCACATCTAGTCAGAGTCTTGTACACAGTAATGGAAATTCCTATTTGGATTGGCACCTGCAGAAGCCAGACCAGTCTCTACAACTCCTGATCTATGAGGTTTCTAAACGAAATTCTGGGGTTCCAGACAGGTTCAGTGGCAGTGGATCAGGAACAGATTTCACACTTAAGATCAGCAGAGTAGAGCCTGAGGATTTGGGAGTTTATTACTGCTTCCAACGTACACATTTACCTCC |
| IGKV5-37*01_S6352_CAST | CAST/EiJ | Musmus IGKV5-37*01 F | 97.85 | 285 | GACATCCTGATGACCCAGTCTCCAGCCACCCTGTCTGTGACTCCAGGAGAAACAGTCAGTCTTTCCTGTAGGGCCAGCCAGAATATTTACAAGAACCTACACTGGTATCAACAGAAATCACATGGGACTCCAAGGCTTCTCATCAAGTATGCATCTGATCCCATCTCTGGGATCCCCTCCAGGTTCACTGGCAGTGGATCAGGGACAGATTACACTCTCAGTATCAACAGTGTGAAGCCCGAAGATGAAGGAATATATTACTGTCTTCAAGGTTACAGCATGCCT |
| IGKV12-89*01_S9010_CAST | CAST/EiJ | Musmus IGKV12-89*01 F | 99.28 | 287 | GACATCCAGATGACTCAGTCTCCAGCTTCACTGTCTGCATCTGTGGGAGAAACTGTCACCATCACATGTGGAGCAAGTGAGAATATTTACGGTGCTTTAAATTGGTATCAGCAGAAACAGGGAAAATCTCCTCAGCTCCTGATCTATGGTGCAACCAACTTGGCAGATGGAATGTCATCGAGGTTCAGTGGCAGTGGATCTGGTAGACAGTATTCTCTCAAGATCAGTAGCCTGCATCCTGACGATGTTGCAACGTATTACTGTCAAAATGTGTTAAGTACCCCTCC |

| | | | | | |
|---|---|---|---|---|---|
| IGKV9-120*01_S7607_CAST | CAST/EiJ | Musmus IGKV9-120*02 F | 96.42 | 285 | GACATCCAGATGACCCAGTCTCCATCCTCCTTATCTGCCTCTCTGGGAGAAAGAATCAGTCTCACTTGCCGGGCAAGTCAGGACATTTATGGTAGCTTAAACTGGTTTCAGCAGAAACCAGATGGAACTATTAAACTCCTGATCTACGGCACATCCAGTTTAGATTCTGGTGTCCCCAAAAGGTTCAGTGGCAGTAGGTCTGGGTCAGATTATTCTCTCACCATCAGCAGCCTTGAGTCTGAAGATTTTGCAGACTATTACTGTCTACAATATGCTAGTTCTCCT |
| IGKV12-38*01_S2611_CAST | CAST/EiJ | Musmus IGKV12-38*01 F | 98.21 | 287 | GACATCCAGATGACTCAGTCTCCAGCCTCCCTGGCTGCATCTGTGGGAGAAACCATCACCATCACATGTCAAGCAAGTGAGAACATTTACTTCAGTTTAGCATGGTATCAGCAGAAGCAAGGGAAATCTCCTCAGCTCCTGATCTATAATGCAAACAGCTTGGAAGATGGTGTCCCATCGAGGTTCAGTGGCAGTGGATCTGGGACACAGTATTCTATGAAGATCAACAGCATGCAGCCTGAAGATACTGCAACTTATTTCTGTAAACAGGCTTATGACTTTCCTCC |
| IGKV6-23*01_S5820_CAST | CAST/EiJ | Musmus IGKV6-23*01 F | 92.11 | 287 | GACATTGTGATGACCCAGTCTCAAAAATTCATGTCCACATCAGTAGGAGAGAGGGTCAGCATCACCTGCAAGGCCAGTCAGAATGTAGGTACTAATGTAGCCTGGTATCAGCAGAAAGCAGGGCAGTCTCTTGAACTGCTGATCTATGGGGCATCCAACCGGCACACTGGAGTCCCTGATCGCTTCACAGGCAGTGGATCTGGGACAGATTTCACTCTCACCATCACCAATGTGCAGTCTGAAGACATGACAAATTATTTCTGTGAGCAATATAGCAGCTATCCGCT |
| IGKV6-23*01_S4960_CAST | CAST/EiJ | Musmus IGKV6-23*01 F | 92.11 | 287 | GACATTGTGATGACCCAGTCTCAAAAATTCATGTCCACATCAGTAGGAGAGAGGGTCAGCATCACCTGCAAGGCCAGTCAGAATGTAGGTACTAATGTAGCCTGGTATCAGCAGAAAGCAGGGCAGTCTCTTGAACTGCTGATCTATGGGGCATCCAACCGGCACACTGGAGTCCCTGATCGCTTCACAGGCAGTGGATCTGGGACAGATTTCACTCTCACCATCACCAATGTGCAGTCTGAAGACATGACAAATTATTTCTGTGAGCAATATAGCAGCTATCCTCT |
| IGKV4-91*01_S3435_CAST | CAST/EiJ | Musmus IGKV4-91*01 F | 96.45 | 292 | GAAATTGTGCTCACCCAGTCTCCAACCATCATGGCTGCATCTCCCGGGGAGAAGGTCACCATCACCTGCAGTGCCAGCTCAAGTATAAGTTCCAATTACTTGCACTGGTATCAGCAGAAGCCAGGATTCCCTCCTAAACTCTTGATATATAGGACATCCAATCTGGCTTCTGGAGTCCCAGCTCGCTTCAGTGGCAGTGGGTCTGGGACCTCTTACTCTCTCACAATTGGCACCATGGAGGCTGAAGGTGCTGCCACTTATTACTGCCAGCAGGGTAGTAGTTTACCACTCA |
| IGKV7-33*01_S2856_CAST | CAST/EiJ | Musmus IGKV7-33*01 F | 97.31 | 305 | GACATTGTGATGACTCAGTCTCCAACTTTCCTTGCTGTGACAGCAAGTAAGAAGGTCACCATTAGTTGCACGGCCAGTGAGAGCCTTTATTCAAGCAAACACAAGGTGAACTACTTAGCTTGGTACCAGAAGAAACCAGAGCAATCTCCTAAACTGCTGATTTATGGGGCATCCAACCGATTCACTGGGGTCCCTGATCGCTTCACAGGCAGTGGATCTGGGACAGATTTCACTCTGACCATCAGCAGTGTGCAGGCTGAAGACCTCACACATTATTACTGTGCACAGTTTTACAGCTATCCTCT |
| IGKV1-99*01_S3206_CAST | CAST/EiJ | Musmus IGKV1-99*01 F | 98.3 | 302 | GATGTTGTTCTGACCCAAACTCCACTCTCTCTGCCTGTCAATATTGGAGATCAAGCCTCTATCTCTTGCAAGTCTACTAAGAGCCTTCTGAATAGTGATGGATTCACTTATTTGGGCTGGTACCTGCAGAAGCCAGGCCAGTCTCCACAGCTCCTAATATATTTGGTTTCTAATCGATTTTCTGGAGTTCCAGACAGGTTCAGTGGTAGTGGGTCAGGGACAGATTTCACCCTCAAGATCAGCAGAGTGGAGGCTGAGGATTTGGGAGTTTATTATTGCTTCCAGAGTAACTATCTTCCTCT |
| IGKV16-104*01_S6011_CAST | CAST/EiJ | Musmus IGKV16-104*01 F | 100 | 288 | GATGTCCAGATAACCCAGTCTCCATCTTATCTTGCTGCATCTCCTGGAGAAACCATTACTATTAATTGCAGGGCAAGTAAGAGCATTAGCAAATATTTAGCCTGGTATCAAGAGAAACCTGGGAAAACTAATAAGCTTCTTATCTACTCTGGATCCACTTTGCAATCTGGAATTCCATCAAGGTTCAGTGGCAGTGGATCTGGTACAGATTTCACTCTCACCATCAGTAGCCTGGAGCCTGAAGATTTTGCAATGTATTACTGTCAACAGCATAATGAATCCCCGTAC |

| | | | | | |
|---|---|---|---|---|---|
| IGKV14-111*01_S5416_CAST | CAST/EiJ | Musmus IGKV14-111*01 F | 95.7 | 287 | GACATCAAGATGACCCAGTCTGCATCCTCCATGTATGCATCGCTGGGAGAGAGAGTCACTATCACTTGCAAGGCGAGTCAGGACATTAAAAGCTATTTAAGCTGGTACCAGCAGAAACCAGGGAAATCTCCTAAGACCCTGATCTATCGTGCAAACAGCTTGGTAGATGGGGTCCCATCAAGGTTCAGTGGCAGTGGATCTGGGCAAGATTATTCTCTCACCATCAGCAGCCTGGAGTATGAAGATCCGGCAATTTATTTTTGTCTACAGTATAATGAGTTTCCTCC |
| IGKV11-125*01_S4527_CAST | CAST/EiJ | Musmus IGKV11-106*02 F | 96.06 | 286 | AATGTTCAAATGACCCAGTCTCCATCCTCCCTGTCTGCATCTTTGGGAGAGAGAGTCTCCCTGACTTGCCAGGCAAGTCAGAGCGTTAGCAATAATTTAAACTGGTATCAGCAAACACCAGGGAAAGCTCCTAGGCTCTTGATCTATGGTGCAAGCAAATTGGAAGATGGGGTCCCTTCAAGGTTCAGTGGCACTGGATATGGGACAGATTTCACTTTCACCATCAGCAGCCTGGAGGAAGAAGATGTGGCAATTTATTTCTGTCTACAGGATAGTTATCTCCCTC |
| IGKV12-98*01_S2656_CAST | CAST/EiJ | Musmus IGKV12-98*01 F | 98.92 | 287 | GACATTCAGATGACCCAGTCTCCTGCCTCCCAGTCTGCATCTCTGGGAGAAAGTGTCACCATCACATGCCTGGCAAGTCAGACCATTGGTACATGGTTAGCATGGTATCAGCAGAAACCAGGGAAATCTCCTCAGCTCCTGATTTATGCTGCAACCAGCTTGGCAGATGGGGTCCCATCAAGGTTCAGTGGTAGTGGATCTGGCACAAAGTTTTCTTTCAAGATCAGCAGCCTACAGGCTGAAGATTTTGCAAGTTATTACTGTCAACAACATTACAGTACTCCTCT |
| IGKV3-4*01_S0127_CAST | CAST/EiJ | Musmus IGKV3-4*01 F | 96.91 | 299 | GACATTGTGCTGACCCAATCTCCAGCTTCTTTGGCTGTGTCTCTAGGGCAGAGGGCCACCATCTCCTGCAGAGCCAGCGAAAGTGTTGATTATGATGGCGATAGTTATATGAACTGGTACCAACAGAAACCAGGACAGCCACCGAAACTCCTCATCTATGCTGCATCCAATCTAGAATCTGGGATCCCTGCCAGGTTTAGTGGCAGTGGGTCTGGGACAGACTTTACCCTCAACATTCATCCTGTGGAGGAGGAGGATGCTGCAACCTATTACTGTCTGCAAAGTAATGAGGCTCCTCC |
| IGKV8-28*02_CAST | CAST/EiJ | Musmus IGKV8-28*02 F | 100 | 302 | GACATTGTGATGACACAGTCTCCATCCTCCCTGAGTGTGTCAGCAGGAGATAAGGTCACTATGAGCTGCAAGTCCAGTCAGAGTCTGTTAAACAGTAGAAACCAAAAGAACTACTTGGCCTGGTACCAGCAGAAACCATGGCAGCCTCCTAAACTGCTGATCTACGGGGCATCCACTAGGGAATCTGGGGTCCCTGATCGCTTCACAGGCAGTGGATCTGGAACAGATTTCACTCTCACCATCAGCAGTGTGCAGGCTGAAGACCTGGCAGTTTATTACTGTCAGAATGATTATAGTTATCC |
| IGKV1-122*01_S7970_CAST | CAST/EiJ | Musmus IGKV1-122*01 F | 98.64 | 302 | GATGCTGTGATGACCCAAACTCCACTCTCCCTGCCTGTCAGTCTTGGAGATCAAGCCTCCATCTCTTGCAGGTCTAGTCAGAGCCTTGAAAACAGTAATGGAAACACCTATTTGAACTGGTACCTCCAGAAACCAGGCCAGTCTCCACAGCTCCTGATCTACAGGGTTTCCAACCGATTTTCTGGGGTCCTAGACAGGTTCAGTGGCAGTGGTTCAGGGACAGATTTCACACTCAAGATCAGCAGAGTGGAGGCTGAGGATTTGGGAGTTTATTTCTGCCTCCAAGTTACACATGTCCCTCC |
| IGKV1-133*01_S0950_CAST | CAST/EiJ | Musmus IGKV1-133*01 F | 99.32 | 302 | GATGTTGTGATGACCCAGACTCCACTCACTTTGTTGGTTACCATTGGACAACCAGCCTCTATCTCTTGCAAGTCAAGTCAGAGCCTCTTATATAGTAATGGAAAAACCTATTTGAATTGGTTATTACAGAGGCCAGGCCAGTCTCCAAAGCGCCTAATCTATCTGGTGTCTAAACTGGACTCTGGAGTCCCTGACAGGTTCACTGGCAGTGGATCAGGAACAGATTTTACACTGAAAATCAGCAGAGTGGAGGCTGAGGATTTGGGAGTTTATTACTGCTTGCAAGGTACACATTTTCCTCA |
| IGKV4-55*01_S8231_CAST | CAST/EiJ | Musmus IGKV4-55*01 F | 95.65 | 286 | CAAATTGTTCTCACCCAGTCTCCAGCAATCATGTCTGCATCTCCAGGGGAGAAGGTCACCATGACCTGCAGTGCCAGCTCAAGTGTAAGTTACATGTACTGGTACCAGCAGAAGCCAGGATCCTCCCCCAGACTCTGGATTTATGACACATCCAACCTGGCTTCTGGAGTTTCCGCTCGCTTCCGTGGCAGTGGGTCTGGGACCTCTTATTCTCTCACAATCAGCAGCATGGAGGCTGAAGATGCTGCCACTTATTACTGCTATCAGTGGAGTAGTTACCCACCCA |

| | | | | | |
|---|---|---|---|---|---|
| IGKV5-39*01_S7300_CAST | CAST/EiJ | Musmus IGKV5-39*01 F | 99.28 | 287 | GACATTGTGATGACTCAGTCTCCAGCCACCCTGTCTGTGACTCCAGGAGATAGAGTCTCTCTTTCCTGCAGGGCCAGCCAGAGTATTAGCAACTACTTACATTGGTATCAACAAAAATCACATGAGTCTCCAAGGCTTCTCATCAAATATGCTTCCCAATCCATCTCTGGGATCCCCTCCAGGTTCAGTGGCAGTGGATCAGGGTCAGATTTCACTCTCAGTATCAACAGTGTGGAACCTGAAGATGTTGGAGTGTATTACTGTCAAAATGGTCACAGCTTTCCTCC |
| IGKV11-125*01_CAST | CAST/EiJ | Musmus IGKV11-125*01 F | 100 | 287 | GATGTCCAGATGATTCAGTCTCCATCCTCCCTGTCTGCATCTTTGGGAGACATAGTCACCATGACTTGCCAGGCAAGTCAGGGCACTAGCATTAATTTAAACTGGTTTCAGCAAAAACCAGGGAAAGCTCCTAAGCTCCTGATCTATGGTGCAAGCAACTTGGAAGATGGGGTCCCATCAAGGTTCAGTGGCAGTAGATATGGGACAGATTTCACTCTCACCATCAGCAGCCTGGAGGATGAAGATATGGCAACTTATTTCTGTCTACAGCATAGTTATCTCCCTCC |
| IGKV14-126*01_CAST | CAST/EiJ | Musmus IGKV14-126*01 F | 100 | 287 | GACATCAAGATGACCCAGTCTCCATCCTCCATGTATGCATCGCTGGGAGAGAGAGTCACTATCACTTGCAAGGCGAGTCAGGACATTAAAAGCTATTTAAGCTGGTACCAGCAGAAACCATGGAAATCTCCTAAGACCCTGATCTATTATGCAACAAGCTTGGCAGATGGGGTCCCATCAAGATTCAGTGGCAGTGGATCTGGGCAAGATTATTCTCTAACCATCAGCAGCCTGGAGTCTGACGATACAGCAACTTATTACTGTCTACAGCATGGTGAGAGCCCTCC |
| IGKV9-123*01_S4220_CAST | CAST/EiJ | Musmus IGKV9-123*01 F | 99.28 | 285 | GACATCCAGATGATTCAGTCTCCATCGTCCATGTTTGGCTCTCTGGGAGACAGAGTCAGTCTCTCTTGCCGGGCTAGTCAGGGCATTAGAGGTAATTTAGACTGGTATCAGCAGAAACCAGGTGGAACTATTAAACTCCTGATCTACTCCACATCCAATTTAAATTCTGGTGTCCCATCAAGGTTCAGTGGCAGTGGGTCTGGGTCAGATTATTCTCTCACCATCAGCAGCCTAGAGTCTGAAGATTTTGCAGACTATTACTGTCTACAGCGTAATGCGTATCCT |
| IGKV20-101-2*01_S0678_CAST | CAST/EiJ | Musmus IGKV20-101-2*01 F | 98.19 | 281 | AATATCCAGGTGATCCAGTCACCATTTCTGTCTGCATCTGTGGGAGAGAGGGTCACAATCAGCTGCAAGACACATCAGCATATTAACAGTTCCATAGCCTGGTACCAGCAAAAAGTTGGAAAAGCTCCCAAACTCCTGATAAGAGATGCAAGTTTTCTCTAACAGACACCCCATCAAGGTTCACTGGGAATGGATTTGGCACAGATTTCACACTCAGCATCAGCAGTATGCAGCCTGAAGATGGTGCCACATACTTCTGCCAGCAGCATTTTAACTATTA |
| IGKV2-112*02_S2310_CAST | CAST/EiJ | Musmus IGKV2-112*02 F | 99.66 | 301 | GATATTGTGATAACCCAGGATGAACTCTCCAATCCTGTCACTTCTGGAGAATCAGTTTCCATCTCCTGCAGGTCTAGTAAGAGTCTCCTATATAAGGATGGGAAGACATACTTGAATTGGTTTCTGCAGAGGCCAGGACAGTCTCCTCAGCTCCTGGTCTATTGGATGTCCACCCGTGCATCAGGAGTCTCAGACCGGTTTAGTGGCAGTGGGTCAGGAACAGTTTTCACACTGGAAATCAGTAGAGTGAAGGCTGAGGATGTCGGTGTGTATTACTGTCAACAAGTTGTAGAGTATCCTC |
| IGKV4-74*01_S4432_CAST | CAST/EiJ | Musmus IGKV4-74*01 F | 95.04 | 292 | CAAATTGTTCTCACCCAGTCTCCAGCAATCATGTCTGCATCTCCAGGGGAGAAGGTCACCATGACCTGCAGGGCCAGCTCAAGTGTAAGTTCCAGTTACTTGCACTGGTACCAGCAGAAGCCAGGATCTTCCCCCAAACTCTGGATTTATAGCACATCCAACCTGGCTTCAGGAGTCCCAGCTCGCTTCAGTGGCAGTGGGTCTGGGACCTCTTACTCTCTCACAATCAGCAGTGTGGAGGCTGAGGATGCTGCCACTTATTACTGCCAGCAGTATGATAGTTCCCCACTCA |
| IGKV4-78*01_S9469_CAST | CAST/EiJ | Musmus IGKV4-78*01 F | 98.23 | 292 | CAAATTGTTCTCACCCAGTCTCCAGCAATCATGTCTGCATCTCCTGGGGAACGGGTCACCATGACCTGCAGTGCCAGCTCAAGTGTAAGTTCCAGCTACTTGTACTGGTACCAGCAGAAGCCAGGATCCTCCCCCAAACTCTGGATTTATAGCACATCCAACCTGGCTTCTGGAGTCCCTGCTCGCTTCAGTGGCAGTGGGTCTGGGACCTCTTATTCTCTCACAATCAGCAGCATGGAGGCTGAAGATGCTGCCACTTATTACTGCCAGCAGTACAGTGGTTACCCATCCA |

| | | | | | |
|---|---|---|---|---|---|
| IGKV10-96*06_CAST | CAST/EiJ | Musmus IGKV10-96*06 ORF | 100 | 287 | GATATCCAGATGACACAGACTACATCCTCCCTGTCTGCCTCTCTGGGAGACAGAGTCACCATCAGTTGCAGGGCAAGTCAGGACATTAGCAATTATTTAAACGGGTATCAGCAGAAACCAGATGGAACTGTTAAACTCCTGATCTACTACACATCAAGATTACACTCAGGAGTCCCATCAAGGTTCAGTGGCAGTGGGTCTGGGACAGATTATTCTCTCACTATTAGCAACCTGGAACAAGAAGATATTGCCACTTACTTTTGCCAACAGGATAGTAAGCATCCTCC |
| IGKV2-109*03_S4111_CAST | CAST/EiJ | Musmus IGKV2-109*03 F, or Musmus IGKV2-109*04 F | 97.96 | 301 | GATATTGTGATGACACAGGCTGCATTCTCCAATCCAGTCACTCTTGGAACATCAGCTTCCATCTCCTGCAGGTCTAGTAAGAGTCTCCTACATAGTAATGGCATCACTTATTTGTATTGGTATCTGCAGAGGCCAGGCCAGTCTCCTCAGCTCCTGATACATCGGATGTCCAACCTTGCCTCAGGAGTCCCAGACAGGTTCAGTAGCAGTGGGTCAGGAACTGATTTCACACTGAGAATCAGCAGAGTGGAGGCTGAGGATGTGGGTGTTTATTACTGTGCTCAAAGGCTAGAACTCCCTC |
| IGKV6-25*01_S3519_CAST | CAST/EiJ | Musmus IGKV6-25*01 F | 98.21 | 287 | TGCATTGTGATGACCCAGTCTCACAAATTCATGTCCACATCAGTAGGAGACAGGGTCAGCATCACCTGCAAGGCCAGTCAGGATGTGAGTACTGCTGTAGCCTGGTATCAACAAAAACCAGGGCAATCTCCTAAACTACTGATTTACTGGGCATCCACCCGGCACACTGGAGTCCCTGATCGCTTCACAGGCAGTGGATCTGGGACAGATTTCACTCTTACCATCAGCAGTGTGCAGGCTGAAGACCTGGCACTTTATTACTGTCAGCAACATTATAGCACTCCTCC |
| IGKV14-130*01_CAST | CAST/EiJ | Musmus IGKV14-130*01 F | 100 | 287 | GAAATCCAGATGACCCAGTCTCCATCCTCTATGTCTGCATCTCTGGGAGACAGAATAACCATCACTTGCCAGGCAACTCAAGACATTGTTAAGAATTTAAACTGGTATCAGCAGAAACCAGGGAAACCCCCTTCATTCCTGATCTATTATGCAACTGAACTGGCAGAAGGGGTCCCATCAAGGTTCAGTGGCAGTGGGTCTGGGTCAGACTATTCTCTGACAATCAGCAACCTGGAGTCTGAAGATTTTGCAGACTATTACTGTCTACAGTTTTATGAGTTTCCTCC |
| IGKV1-88*01_S1851_CAST | CAST/EiJ | Musmus IGKV1-88*01 F | 98.3 | 299 | GATGTTGTGGTGACCCAAACTCCACTCTCCCTGCCTGTCAGCTTTGGAGATCAGGTTTCTATCTCTTGCAGGTCTAGTCAGAGTCTTGCAAACAGTTATGGGAACACCTATTTGTCTTGGTACCTGCACAAGCCTGGCCAGTCTCCTCAGCTCCTCATCTATGGGATTTTCAACAGATTTTCCGGGGTGCCAGACAGGTTCAGTGGCAGTGGTTCAGGGACAGATTTCACACTCAAGATCAGCACAATAAAGCCTGAGGACTTGGGAATGTATTACTGCTTACAAGGTACACATCAGCC |
| IGKV4-92*01_S4169_CAST | CAST/EiJ | Musmus IGKV4-92*01 F | 98.94 | 291 | GAAATTGTTCTCACCCAGTCTCCAGTATCCATAACTGCATCTCGAGGGGAGAAGGTCACCATCACCTGCCGTGCCAGCTCAAGTATAAGTTCCAAATACTTACACTGGTACCAGCAGAAGCCAGAATCCTCCCCTAAACTTTTGATTTATAGGACATCCATCCTGGCATCTGGAGTCCTAGACAGCTTCAGTGGCAGTGGGTCTGAGAGCTCTTACACTCTGACAATCAGCTGCATGCAGGACGAAGTTGCTGCCACTTACTATTGTCAGCAGGGGAGTAGTAGCCCACCA |
| IGKV18-36*01_S3639_CAST | CAST/EiJ | Musmus IGKV18-36*01 F | 99.64 | 287 | ACTGGAGAAACAACACAGGCTCCAGCTTCTCTGAGTTTTTCTCTTGGTGAAACAGCAACACTGTCATGCAGGTCCAGTGAGAGTGTTGGCAGCTACTTAGCCTGGTACCAGCAGAAAGCAGAGCAAGTTCCCCGGCTCCTTATCCATAGTGCCTCCACTAGGACCGGTGGTGTCCCAGTCCGGTTCAGTGGCACTGGGTCTGGGACAGACTTCACTCTCACCATCAGCAGTCTAGAACCTGAAGATGCTGCAGTTTACTACTGTCAACCTTTCAAAAGTTGGTCATA |
| IGKV1-35*01_S2930_CAST | CAST/EiJ | Musmus IGKV1-35*01 ORF | 96.94 | 302 | GACATTGTGATGACCCAGACTCCACTCACTTTATCAGCTACCATTGGACAATCAGCCTCCATCTCTTGCAGGTCAAGTCAGAGTCTCTTACATAGTAATGGAAACACATACTTGAATTGGTTTCTACAGAGGCCAGGCCAATCTCCACAGCTTCTGATTTATGGGGTGTTTGAACGGGAATCTGGGGTTCCTGACAGGTTCAGTGGCAGTGGGTCAGGAACAGATTTCACACTCAAGATCAGTAGAGTGGAGGCTGAGGATTTGGGAGTTTATTACTGCATGCAAGCTACCTATGAACCTCC |

| | | | | | |
|---|---|---|---|---|---|
| IGKV4-61*01_S4622_CAST | CAST/EiJ | Musmus IGKV4-61*01 F | 96.01 | 286 | CAAATTGTTCTCACCCAGTCCCCAGCAATCCTGTCTGCATCTCCAGGGGAGAAGGTCACCATATCCTGCAGTGCCAGCTCAAGTGTAAATTACATGTACTGGTACCAGCAGAAGCCAGGATCCTCACCCAAACCCTGGATTTATCCCACATCCAACCTGGCTTCTGGAGTCCCTGCTCGCTTCAATGGCAGTGGGTCTGCGACCTCTTATTCTCTCACAATCAGCAGCATGGAGGCTGAAGATGCTGCCACTTATTACTGCCAGCAGTTTACTAGTTCCCCATCCA |
| IGKV4-62*01_S7049_CAST | CAST/EiJ | Musmus IGKV4-55*01 F | 95.65 | 286 | CAAATTGTTCTCACCCAGTCTCCAGCAATCATGTCTGCATCTCCAGGGGAGAAGGTCACCATGACCTGCAGTGCCAGCTCAAGTGTAAGTTACATGTACTGGTACCAGCAGAAGCCAGGATCCTCCCCCAGACTCTGGATTTATGACACATCCAACCTGGCTTCTGGAGTTTCCGCTCGCTTCCGTGGCAGTGGGTCTGGGACCTCTTATTCTCTCACAATCAGCAGCATGGAGGCTGAAGATGCTGCCACTTATTACTGCTATCAGTGGAGTAGTTACCCATTCA |
| IGKV4-63*01_S6121_CAST | CAST/EiJ | Musmus IGKV4-63*01 F | 97.1 | 286 | GAAAATGTTCTCACCCAGTCTCCAGCAATCATGTCTGCATCTCCAGGGGAGAAGGTCACCATGACCTGCAGTGCCAGCTCAAGTGTAAGTTACATGCACTGGTACCAGCAGAAGTCAGGCACCTCTCCCAAACTCTGGATTTATGACACATCCAAACTGGCTTCTGGAGTCCCAGGTCGCTTCAGTGGCAATGGGTCTGGAAACTCTTACTCTCTCACGATCAGCAGCATGGAGGCTGAAGATGCTGCCACTTATTACTGCTTTCAGGGGAGTCTGTACCCATTCA |
| IGKV4-81*01_S8073_CAST | CAST/EiJ | Musmus IGKV4-81*01 F | 95.39 | 290 | GAAAATGTGCTGACCCAGTCTCCAGCAATCATGGCTGCATCTCCAGGGGAGAAGGTCACCATGACCTGCAGTGCCAGCTCAAGTGTAAGTTCTGGTAACTTTCACTGGTACCAGCAGAAGCCAGGCACTTCTCACAAACTCTGGATTTATAGGACATCCAACCTGGCTTCTGGAGTCCCAGCTCGCTTCAGTGGCAGTGGGTCTGGGACCTCTTATTCTCTTACAATCAACAGCATGGAAGCCGAAGATGCTGCCACTTATTACTGCCAGCAGTGGAGTGGTTACCCACC |
| IGKV6-15*01_CBA | CBA/J | Musmus IGKV6-15*01 F | 100 | 287 | GACATTGTGATGACCCAGTCTCAAAAATTCATGTCCACATCAGTAGGAGACAGGGTCAGCGTCACCTGCAAGGCCAGTCAGAATGTGGGTACTAATGTAGCCTGGTATCAACAGAAACCAGGGCAATCTCCTAAAGCACTGATTTACTCGGCATCCTACCGGTACAGTGGAGTCCCTGATCGCTTCACAGGCAGTGGATCTGGGACAGATTTCACTCTCACCATCAGCAATGTGCAGTCTGAAGACTTGGCAGAGTATTTCTGTCAGCAATATAACAGCTATCCTCT |
| IGKV8-30*01_CBA | CBA/J | Musmus IGKV8-30*01 F | 100 | 305 | GACATTGTGATGTCACAGTCTCCATCCTCCCTAGCTGTGTCAGTTGGAGAGAAGGTTACTATGAGCTGCAAGTCCAGTCAGAGCCTTTTATATAGTAGCAATCAAAAGAACTACTTGGCCTGGTACCAGCAGAAACCAGGGCAGTCTCCTAAACTGCTGATTTACTGGGCATCCACTAGGGAATCTGGGGTCCCTGATCGCTTCACAGGCAGTGGATCTGGGACAGATTTCACTCTCACCATCAGCAGTGTGAAGGCTGAAGACCTGGCAGTTTATTACTGTCAGCAATATTATAGCTATCCTCC |
| IGKV8-27*01_CBA | CBA/J | Musmus IGKV8-27*01 F | 100 | 300 | AACATTATGATGACACAGTCGCCATCATCTCTGGCTGTGTCTGCAGGAGAAAAGGTCACTATGAGCTGTAAGTCCAGTCAAAGTGTTTTATACAGTTCAAATCAGAAGAACTACTTGGCCTGGTACCAGCAGAAACCAGGGCAGTCTCCTAAACTGCTGATCTACTGGGCATCCACTAGGGAATCTGGTGTCCCTGATCGCTTCACAGGCAGTGGATCTGGGACAGATTTTACTCTTACCATCAGCAGTGTACAAGCTGAAGACCTGGCAGTTTATTACTGTCATCAATACCTCTCCTCG |
| IGKV8-19*01_CBA | CBA/J | Musmus IGKV8-19*01 F | 100 | 305 | GACATTGTGATGACACAGTCTCCATCCTCCCTGACTGTGACAGCAGGAGAGAAGGTCACTATGAGCTGCAAGTCCAGTCAGAGTCTGTTAAACAGTGGAAATCAAAAGAACTACTTGACCTGGTACCAGCAGAAACCAGGGCAGCCTCCTAAACTGTTGATCTACTGGGCATCCACTAGGGAATCTGGGGTCCCTGATCGCTTCACAGGCAGTGGATCTGGAACAGATTTCACTCTCACCATCAGCAGTGTGCAGGCTGAAGACCTGGCAGTTTATTACTGTCAGAATGATTATAGTTATCCTCC |

| | | | | | |
|---|---|---|---|---|---|
| IGKV14-126*01_CBA | CBA/J | Musmus IGKV14-126*01 F | 100 | 287 | GACATCAAGATGACCCAGTCTCCATCCTCCATGTATGCATCGCTGGGAGAGAGAGTCACTATCACTTGCAAGGCGAGTCAGGACATTAAAAGCTATTTAAGCTGGTACCAGCAGAAACCATGGAAATCTCCTAAGACCCTGATCTATTATGCAACAAGCTTGGCAGATGGGGTCCCATCAAGATTCAGTGGCAGTGGATCTGGGCAAGATTATTCTCTAACCATCAGCAGCCTGGAGTCTGACGATACAGCAACTTATTACTGTCTACAGCATGGTGAGAGCCCTCC |
| IGKV12-44*01_CBA | CBA/J | Musmus IGKV12-44*01 F | 100 | 287 | GACATCCAGATGACTCAGTCTCCAGCCTCCCTATCTGCATCTGTGGGAGAAACTGTCACCATCACATGTCGAGCAAGTGAGAATATTTACAGTTATTTAGCATGGTATCAGCAGAAACAGGGAAAATCTCCTCAGCTCCTGGTCTATAATGCAAAAACCTTAGCAGAAGGTGTGCCATCAAGGTTCAGTGGCAGTGGATCAGGCACACAGTTTTCTCTGAAGATCAACAGCCTGCAGCCTGAAGATTTTGGGAGTTATTACTGTCAACATCATTATGGTACTCCTCC |
| IGKV6-23*01_CBA | CBA/J | Musmus IGKV6-23*01 F | 100 | 287 | GACATTGTGATGACCCAGTCTCACAAATTCATGTCCACATCAGTAGGAGACAGGGTCAGCATCACCTGCAAGGCCAGTCAGGATGTGGGTACTGCTGTAGCCTGGTATCAACAGAAACCAGGGCAATCTCCTAAACTACTGATTTACTGGGCATCCACCCGGCACACTGGAGTCCCTGATCGCTTCACAGGCAGTGGATCTGGGACAGATTTCACTCTCACCATTAGCAATGTGCAGTCTGAAGACTTGGCAGATTATTTCTGTCAGCAATATAGCAGCTATCCTCT |
| IGKV1-135*01_CBA | CBA/J | Musmus IGKV1-135*01 F | 100 | 302 | GATGTTGTGATGACCCAGACTCCACTCACTTTGTCGGTTACCATTGGACAACCAGCCTCCATCTCTTGCAAGTCAAGTCAGAGCCTCTTAGATAGTGATGGAAAGACATATTTGAATTGGTTGTTACAGAGGCCAGGCCAGTCTCCAAAGCGCCTAATCTATCTGGTGTCTAAACTGGACTCTGGAGTCCCTGACAGGTTCACTGGCAGTGGATCAGGGACAGATTTCACACTGAAAATCAGCAGAGTGGAGGCTGAGGATTTGGGAGTTTATTATTGCTGGCAAGGTACACATTTTCCTCA |
| IGKV4-57*01_S6684_CBA | CBA/J | Musmus IGKV4-57-1*01 F | 100 | 286 | CAAATTGTTCTCACCCAGTCTCCAGCAATCATGTCTGCATCTCCAGGGGAGAAGGTCACCATAACCTGCAGTGCCAGCTCAAGTGTAAGTTACATGCACTGGTTCCAGCAGAAGCCAGGCACTTCTCCCAAACTCTGGATTTATAGCACATCCAACCTGGCTTCTGGAGTCCCTGCTCGCTTCAGTGGCAGTGGATCTGGGACCTCTTACTCTCTCACAATCAGCCGAATGGAGGCTGAAGATGCTGCCACTTATTACTGCCAGCAAAGGAGTAGTTACCCGCTCA |
| IGKV15-103*01_CBA | CBA/J | Musmus IGKV15-103*01 ORF | 100 | 287 | GACATCCAGATGAACCAGTCTCCATCCAGTCTGTCTGCATCCCTTGGAGACACAATTACCATCACTTGCCATGCCAGTCAGAACATTAATGTTTGGTTAAGCTGGTACCAGCAGAAACCAGGAAATATTCCTAAACTATTGATCTATAAGGCTTCCAACTTGCACACAGGCGTCCCATCAAGGTTTAGTGGCAGTGGATCTGGAACAGGTTTCACATTAACCATCAGCAGCCTGCAGCCTGAAGACATTGCCACTTACTACTGTCAACAGGGTCAAAGTTATCCTCT |
| IGKV2-137*01_CBA | CBA/J | Musmus IGKV2-137*01 F | 100 | 302 | GATATTGTGATGACTCAGGCTGCACCCTCTGTACCTGTCACTCCTGGAGAGTCAGTATCCATCTCCTGCAGGTCTAGTAAGAGTCTCCTGCATAGTAATGGCAACACTTACTTGTATTGGTTCCTGCAGAGGCCAGGCCAGTCTCCTCAGCTCCTGATATATCGGATGTCCAACCTTGCCTCAGGAGTCCCAGACAGGTTCAGTGGCAGTGGGTCAGGAACTGCTTTCACACTGAGAATCAGTAGAGTGGAGGCTGAGGATGTGGGTGTTTATTACTGTATGCAACATCTAGAATATCCTTT |
| IGKV4-59*01_CBA | CBA/J | Musmus IGKV4-59*01 F | 100 | 286 | CAAATTGTTCTCACCCAGTCTCCAGCAATCATGTCTGCATCTCCAGGGGAGAAGGTCACCATGACCTGCAGTGCCAGCTCAAGTGTAAGTTACATGCACTGGTACCAGCAGAAGTCAGGCACCTCCCCCAAAAGATGGATTTATGACACATCCAAACTGGCTTCTGGAGTCCCTGCTCGCTTCAGTGGCAGTGGGTCTGGGACCTCTTACTCTCTCACAATCAGCAGCATGGAGGCTGAAGATGCTGCCACTTATTACTGCCAGCAGTGGAGTAGTAACCCACCCA |

| | | | | | |
|---|---|---|---|---|---|
| IGKV8-21*01_CBA | CBA/J | Musmus IGKV8-21*01 F | 100 | 302 | GACATTGTGATGTCACAGTCTCCATCCTCCCTGGCTGTGTCAGCAGGAGAGAAGGTCACTATGAGCTGCAAATCCAGTCAGAGTCTGCTCAACAGTAGAACCCGAAAGAACTACTTGGCTTGGTACCAGCAGAAACCAGGGCAGTCTCCTAAACTGCTGATCTACTGGGCATCCACTAGGGAATCTGGGGTCCCTGATCGCTTCACAGGCAGTGGATCTGGGACAGATTTCACTCTCACCATCAGCAGTGTGCAGGCTGAAGACCTGGCAGTTTATTACTGCAAGCAATCTTATAATCTTCC |
| IGKV12-41*01_CBA | CBA/J | Musmus IGKV12-41*01 F | 100 | 287 | GACATCCAGATGACTCAGTCTCCAGCCTCCCTATCTGCATCTGTGGGAGAAACTGTCACCATCACATGTCGAGCAAGTGGGAATATTCACAATTATTTAGCATGGTATCAGCAGAAACAGGGAAAATCTCCTCAGCTCCTGGTCTATAATGCAAAAACCTTAGCAGATGGTGTGCCATCAAGGTTCAGTGGCAGTGGATCAGGAACACAATATTCTCTCAAGATCAACAGCCTGCAGCCTGAAGATTTTGGGAGTTATTACTGTCAACATTTTTGGAGTACTCCTCC |
| IGKV6-13*01_CBA | CBA/J | Musmus IGKV6-13*01 F | 100 | 285 | GACATTGTGATGACCCAGTCTCAAAAATTCATGTCCACATCAGTAGGAGACAGGGTCAGCATCACCTGCAAGGCCAGTCAGAATGTGGGTACTGCTGTAGCCTGGTATCAACAGAAACCAGGACAATCTCCTAAACTACTGATTTACTCGGCATCCAATCGGTACACTGGAGTCCCTGATCGCTTCACAGGCAGTGGATCTGGGACAGATTTCACTCTCACCATCAGCAATATGCAGTCTGAAGACCTGGCAGATTATTTCTGCCAGCAATATAGCAGCTATCCT |
| IGKV8-28*01_CBA | CBA/J | Musmus IGKV8-28*01 F | 100 | 305 | GACATTGTGATGACACAGTCTCCATCCTCCCTGAGTGTGTCAGCAGGAGAGAAGGTCACTATGAGCTGCAAGTCCAGTCAGAGTCTGTTAAACAGTGGAAATCAAAAGAACTACTTGGCCTGGTACCAGCAGAAACCAGGGCAGCCTCCTAAACTGTTGATCTACGGGGCATCCACTAGGGAATCTGGGGTCCCTGATCGCTTCACAGGCAGTGGATCTGGAACCGATTTCACTCTTACCATCAGCAGTGTGCAGGCTGAAGACCTGGCAGTTTATTACTGTCAGAATGATCATAGTTATCCTCC |
| IGKV12-46*01_CBA | CBA/J | Musmus IGKV12-46*01 F | 100 | 287 | GACATCCAGATGACTCAGTCTCCAGCCTCCCTATCTGTATCTGTGGGAGAAACTGTCACCATCACATGTCGAGCAAGTGAGAATATTTACAGTAATTTAGCATGGTATCAGCAGAAACAGGGAAAATCTCCTCAGCTCCTGGTCTATGCTGCAACAAACTTAGCAGATGGTGTGCCATCAAGGTTCAGTGGCAGTGGATCAGGCACACAGTATTCCCTCAAGATCAACAGCCTGCAGTCTGAAGATTTTGGGAGTTATTACTGTCAACATTTTTGGGGTACTCCTCC |
| IGKV10-96*01_CBA | CBA/J | Musmus IGKV10-96*01 F | 100 | 286 | GATATCCAGATGACACAGACTACATCCTCCCTGTCTGCCTCTCTGGGAGACAGAGTCACCATCAGTTGCAGGGCAAGTCAGGACATTAGCAATTATTTAAACTGGTATCAGCAGAAACCAGATGGAACTGTTAAACTCCTGATCTACTACACATCAAGATTACACTCAGGAGTCCCATCAAGGTTCAGTGGCAGTGGGTCTGGAACAGATTATTCTCTCACCATTAGCAACCTGGAGCAAGAAGATATTGCCACTTACTTTTGCCAACAGGGTAATACGCTTCCTC |
| IGKV4-57-1*01_CBA | CBA/J | Musmus IGKV4-57*01 F | 100 | 292 | GAAAATGTGCTCACCCAGTCTCCAGCAATCATGTCTGCATCTCCAGGGGAAAAGGTCACCATGACCTGCAGGGCCAGCTCAAGTGTAAGTTCCAGTTACTTGCACTGGTACCAGCAGAAGTCAGGTGCCTCCCCCAAACTCTGGATTTATAGCACATCCAACTTGGCTTCTGGAGTCCCTGCTCGCTTCAGTGGCAGTGGGTCTGGGACCTCTTACTCTCTCACAATCAGCAGTGTGGAGGCTGAAGATGCTGCCACTTATTACTGCCAGCAGTACAGTGGTTACCCACTCA |
| IGKV3-5*01_CBA | CBA/J | Musmus IGKV3-5*01 F | 100 | 299 | GACATTGTGCTGACCCAATCTCCAGCTTCTTTGGCTGTGTCTCTAGGGCAGAGGGCCACCATATCCTGCAGAGCCAGTGAAAGTGTTGATAGTTATGGCAATAGTTTTATGCACTGGTACCAGCAGAAACCAGGACAGCCACCCAAACTCCTCATCTATCGTGCATCCAACCTAGAATCTGGGATCCCTGCCAGGTTCAGTGGCAGTGGGTCTAGGACAGACTTCACCCTCACCATTAATCCTGTGGAGGCTGATGATGTTGCAACCTATTACTGTCAGCAAAGTAATGAGGATCCTCC |

| | | | | | |
|---|---|---|---|---|---|
| IGKV1-110*01_CBA | CBA/J | Musmus IGKV1-110*01 F | 100 | 302 | GATGTTGTGATGACCCAAACTCCACTCTCCCTGCCTGTCAGTCTTGGAGATCAAGCCTCCATCTCTTGCAGATCTAGTCAGAGCCTTGTACACAGTAATGGAAACACCTATTTACATTGGTACCTGCAGAAGCCAGGCCAGTCTCCAAAGCTCCTGATCTACAAAGTTTCCAACCGATTTTCTGGGGTCCCAGACAGGTTCAGTGGCAGTGGATCAGGGACAGATTTCACACTCAAGATCAGCAGAGTGGAGGCTGAGGATCTGGGAGTTTATTTCTGCTCTCAAAGTACACATGTTCCTCC |
| IGKV1-117*01_CBA | CBA/J | Musmus IGKV1-117*01 F | 100 | 302 | GATGTTTTGATGACCCAAACTCCACTCTCCCTGCCTGTCAGTCTTGGAGATCAAGCCTCCATCTCTTGCAGATCTAGTCAGAGCATTGTACATAGTAATGGAAACACCTATTTAGAATGGTACCTGCAGAAACCAGGCCAGTCTCCAAAGCTCCTGATCTACAAAGTTTCCAACCGATTTTCTGGGGTCCCAGACAGGTTCAGTGGCAGTGGATCAGGGACAGATTTCACACTCAAGATCAGCAGAGTGGAGGCTGAGGATCTGGGAGTTTATTACTGCTTTCAAGGTTCACATGTTCCTCC |
| IGKV4-70*01_CBA | CBA/J | Musmus IGKV4-70*01 F | 100 | 282 | CAAATTGTTCTCACCCAGTCTCCAGCAATCATGTCTGCATCTCCAGGGGAGAAGGTCACCATGACCTGCAGTGCCAGCTCAAGTATAAGTTACATGCACTGGTACCAGCAGAAGCCAGGCACCTCCCCCAAAAGATGGATTTATGACACATCCAAACTGGCTTCTGGAGTCCCTGCTCGCTTCAGTGGCAGTGGGTCTGGGACCTCTTATTCTCTCACAATCAGCAGCATGGAGGCTGAAGATGCTGCCACTTATTACTGCCATCAGCGGAGTAGTTACCCA |
| IGKV8-24*01_CBA | CBA/J | Musmus IGKV8-24*01 F | 100 | 305 | GACATTGTGATGACACAGTCTCCATCCTCCCTGGCTATGTCAGTAGGACAGAAGGTCACTATGAGCTGCAAGTCCAGTCAGAGCCTTTTAAATAGTAGCAATCAAAAGAACTATTTGGCCTGGTACCAGCAGAAACCAGGACAGTCTCCTAAACTTCTGGTATACTTTGCATCCACTAGGGAATCTGGGGTCCCTGATCGCTTCATAGGCAGTGGATCTGGGACAGATTTCACTCTTACCATCAGCAGTGTGCAGGCTGAAGACCTGGCAGATTACTTCTGTCAGCAACATTATAGCACTCCTCC |
| IGKV6-20*01_CBA | CBA/J | Musmus IGKV6-20*01 F | 100 | 287 | AACATTGTAATGACCCAATCTCCCAAATCCATGTCCATGTCAGTAGGAGAGAGGGTCACCTTGAGCTGCAAGGCCAGTGAGAATGTGGGTACTTATGTATCCTGGTATCAACAGAAACCAGAGCAGTCTCCTAAACTGCTGATATACGGGGCATCCAACCGGTACACTGGGGTCCCCGATCGCTTCACAGGCAGTGGATCTGCAACAGATTTCACTCTGACCATCAGCAGTGTGCAGGCTGAAGACCTTGCAGATTATCACTGTGGACAGAGTTACAGCTATCCTCC |
| IGKV4-80*01_CBA | CBA/J | Musmus IGKV4-80*01 F | 100 | 282 | CAAATTGTTCTCACCCAGTCTCCAGCAATCATGTCTGCATCTCTAGGGGAGGAGATCACCCTAACCTGCAGTGCCAGCTCGAGTGTAAGTTACATGCACTGGTACCAGCAGAAGTCAGGCACTTCTCCCAAACTCTTGATTTATAGCACATCCAACCTGGCTTCTGGAGTCCCTTCTCGCTTCAGTGGCAGTGGGTCTGGGACCTTTTATTCTCTCACAATCAGCAGTGTGGAGGCTGAAGATGCTGCCGATTATTACTGCCATCAGTGGAGTAGTTATCCA |
| IGKV3-7*01_S5296_CBA | CBA/J | Musmus IGKV3-7*02 F | 100 | 299 | GACATTGTGCTGACACAGTCTCCTGCTTCCTTAGCTGTATCTCTGGGGCAGAGGGCCACCATCTCATGCAGGGCCAGCCAAAGTGTCAGTACATCTAGCTATAGTTATATGCACTGGTACCAACAGAAACCAGGACAGCCACCCAAACTCCTCATCAAGTATGCATCCAACCTAGAATCTGGGGTCCCTGCCAGGTTCAGTGGCAGTGGGTCTGGGACAGACTTCACCCTCAACATCCATCCTGTGGAGGAGGAGGATACTGCAACATATTACTGTCAGCACAGTTGGGAGATTCCTCC |
| IGKV5-43*01_CBA | CBA/J | Musmus IGKV5-43*01 F | 100 | 287 | GATATTGTGCTAACTCAGTCTCCAGCCACCCTGTCTGTGACTCCAGGAGATAGCGTCAGTCTTTCCTGCAGGGCCAGCCAAAGTATTAGCAACAACCTACACTGGTATCAACAAAAATCACATGAGTCTCCAAGGCTTCTCATCAAGTATGCTTCCCAGTCCATCTCTGGGATCCCCTCCAGGTTCAGTGGCAGTGGATCAGGGACAGATTTCACTCTCAGTATCAACAGTGTGGAGACTGAAGATTTTGGAATGTATTTCTGTCAACAGAGTAACAGCTGGCCTCA |

| | | | | | |
|---|---|---|---|---|---|
| IGKV4-53*01_CBA | CBA/J | Musmus IGKV4-53*01 F | 100 | 292 | GAAATTGTGCTCACCCAGTCTCCAGCACTCATGGCTGCATCTCCAGGGGAGAAGGTCACCATCACCTGCAGTGTCAGCTCAAGTATAAGTTCCAGCAACTTGCACTGGTACCAGCAGAAGTCAGAAACCTCCCCCAAACCCTGGATTTATGGCACATCCAACCTGGCTTCTGGAGTCCCTGTTCGCTTCAGTGGCAGTGGATCTGGGACCTCTTATTCTCTCACAATCAGCAGCATGGAGGCTGAAGATGCTGCCACTTATTACTGTCAACAGTGGAGTAGTTACCCACTCA |
| IGKV8-16*01_CBA | CBA/J | Musmus IGKV8-16*01 F | 100 | 305 | GAAATTGTGTTGACCCAGTCTATACCATCCCTGACTGTGTCAGCAGGAGAGAGGGTCACTATCAGCTGCAAATCCAATCAGAATCTTTTATGGAGTGGAAACCAAAGGTACTGTTTGGTCTGGCACCAGTGGAAACCAGGGCAAACTCCTACACCGTTGATCACCTGGACATCTGATAGGTACTCTGGAGTCCCTGATCGTTTCATAGGCAGTGGATCTGTGACAGATTTCACTCTGACCATCAGCAGTGTGCAGGCTGAAGATGTGGCAGTTTATTTCTGTCAGCAGCATTTACACATTCCTCC |
| IGKV4-72*01_CBA | CBA/J | Musmus IGKV4-72*01 F | 100 | 286 | CAAATTGTTCTCTCCCAGTCTCCAGCAATCCTGTCTGCATCTCCAGGGGAGAAGGTCACAATGACTTGCAGGGCCAGCTCAAGTGTAAGTTACATGCACTGGTACCAGCAGAAGCCAGGATCCTCCCCCAAACCCTGGATTTATGCCACATCCAACCTGGCTTCTGGAGTCCCTGCTCGCTTCAGTGGCAGTGGGTCTGGGACCTCTTACTCTCTCACAATCAGCAGAGTGGAGGCTGAAGATGCTGCCACTTATTACTGCCAGCAGTGGAGTAGTAACCCACCCA |
| IGKV12-89*01_CBA | CBA/J | Musmus IGKV12-89*01 F | 100 | 287 | GACATCCAGATGACTCAGTCTCCAGCTTCACTGTCTGCATCTGTGGGAGAAACTGTCACCATCACATGTGGAGCAAGTGAGAATATTTACGGTGCTTTAAATTGGTATCAGCGGAAACAGGGAAAATCTCCTCAGCTCCTGATCTATGGTGCAACCAACTTGGCAGATGGCATGTCATCGAGGTTCAGTGGCAGTGGATCTGGTAGACAGTATTCTCTCAAGATCAGTAGCCTGCATCCTGACGATGTTGCAACGTATTACTGTCAAAATGTGTTAAGTACTCCTCC |
| IGKV6-32*01_CBA | CBA/J | Musmus IGKV6-32*01 F | 100 | 287 | AGTATTGTGATGACCCAGACTCCCAAATTCCTGCTTGTATCAGCAGGAGACAGGGTTACCATAACCTGCAAGGCCAGTCAGAGTGTGAGTAATGATGTAGCTTGGTACCAACAGAAGCCAGGGCAGTCTCCTAAACTGCTGATATACTATGCATCCAATCGCTACACTGGAGTCCCTGATCGCTTCACTGGCAGTGGATATGGGACGGATTTCACTTTCACCATCAGCACTGTGCAGGCTGAAGACCTGGCAGTTTATTTCTGTCAGCAGGATTATAGCTCTCCTCC |
| IGKV5-45*01_CBA | CBA/J | Musmus IGKV5-45*01 F | 100 | 287 | GATATTGTGCTAACTCAGTCTCCAGCCACCCTGTCTGTGACTCCAGGAGATAGAGTCAGTCTTTCCTGCAGGGCCAGTCAAAGTATTAGCAACTACCTACACTGGTATCAACAAAAATCACATGAGTCTCCAAGGCTTCTCATCAAGTATGCTTCCCAGTCCATCTCTGGGATCCCCTCCAGGTTCAGTGGCAGTGGATCAGGGACAGATTTCACTCTCAGTATCAACAGTGTGGAGACTGAAGATTTTGGAATGTATTTCTGTCAACAGAGTAACAGCTGGCCTCA |
| IGKV3-4*01_CBA | CBA/J | Musmus IGKV3-4*01 F | 100 | 299 | GACATTGTGCTGACCCAATCTCCAGCTTCTTTGGCTGTGTCTCTAGGGCAGAGGGCCACCATCTCCTGCAAGGCCAGCCAAAGTGTTGATTATGATGGTGATAGTTATATGAACTGGTACCAACAGAAACCAGGACAGCCACCCAAACTCCTCATCTATGCTGCATCCAATCTAGAATCTGGGATCCCAGCCAGGTTTAGTGGCAGTGGGTCTGGGACAGACTTCACCCTCAACATCCATCCTGTGGAGGAGGAGGATGCTGCAACCTATTACTGTCAGCAAAGTAATGAGGATCCTCC |
| IGKV8-18*01_CBA | CBA/J | Musmus IGKV8-18*01 ORF | 100 | 310 | GACATTGTGATGACCCAGTCTCCATCCTCCCTGGCTGTGACAGCAGGAGAGAAGGTCACTATGAGATGCAAGTCCAGTCAGAGTCTTTTGTGGAGTGTAAACCAAAATAACTACTTATCCTGGTACCAGCAGAAACAAGGGCAGCCTCCTAAACTGCTTATCTATGGGGCATCCATTAGAGAATCTTGGGTCCCTGATCGATTCACAGGAAGTGGATCTGGGACAGACTTCACTCTCACCATTAGCAATGTGCATGCTGAAGACCTAGCAGTTTATTACTGTCAGCACAATCATGGCAGCTTTCTCCCCC |

| | | | | | |
|---|---|---|---|---|---|
| IGKV10-94*01_CBA | CBA/J | Musmus IGKV10-94*01 F | 100 | 287 | GATATCCAGATGACACAGACTACATCCTCCCTGTCTGCCTCTCTGGGAGACAGAGTCACCATCAGTTGCAGTGCAAGTCAGGGCATTAGCAATTATTTAAACTGGTATCAGCAGAAACCAGATGGAACTGTTAAACTCCTGATCTATTACACATCAAGTTTACACTCAGGAGTCCCATCAAGGTTCAGTGGCAGTGGGTCTGGGACAGATTATTCTCTCACCATCAGCAACCTGGAACCTGAAGATATTGCCACTTACTATTGTCAGCAGTATAGTAAGCTTCCTCC |
| IGKV17-127*01_CBA | CBA/J | Musmus IGKV17-127*01 F | 100 | 287 | GAAACAACTGTGACCCAGTCTCCAGCATCCCTGTCCGTGGCTACAGGAGAAAAAGTCACTATCAGATGCATAACCAGCACTGATATTGATGATGATATGAACTGGTACCAGCAGAAGCCAGGGGAACCTCCTAAGCTCCTTATTTCAGAAGGCAATACTCTTCGTCCTGGAGTCCCATCCCGATTCTCCAGCAGTGGCTATGGCACAGATTTTGTTTTTACAATTGAAAACACGCTCTCAGAAGATGTTGCAGATTACTACTGTTTGCAAAGTGATAACATGCCTCT |
| IGKV4-91*01_S5997_CBA | CBA/J | Musmus IGKV4-91*01 F | 100 | 292 | GAAATTGTGCTCACCCAGTCTCCAACCACCATGGCTGCATCTCCCGGGGAGAAGATCACTATCACCTGCAGTGCCAGCTCAAGTATAAGTTCCAATTACTTGCATTGGTATCAGCAGAAGCCAGGATTCTCCCCTAAACTCTTGATTTATAGGACATCCAATCTGGCTTCTGGAGTCCCAGCTCGCTTCAGTGGCAGTGGGTCTGGGACCTCTTACTCTCTCACAATTGGCACCATGGAGGCTGAAGATGTTGCCACTTACTACTGCCAGCAGGGTAGTAGTATACCACTCA |
| IGKV6-17*01_CBA | CBA/J | Musmus IGKV6-17*01 F | 100 | 287 | GACATTGTGATGACCCAGTCTCACAAATTCATGTCCACATCAGTAGGAGACAGGGTCAGCATCACCTGCAAGGCCAGTCAGGATGTGAGTACTGCTGTAGCCTGGTATCAACAGAAACCAGGACAATCTCCTAAACTACTGATTTACTCGGCATCCTACCGGTACACTGGAGTCCCTGATCGCTTCACTGGCAGTGGATCTGGGACGGATTTCACTTTCACCATCAGCAGTGTGCAGGCTGAAGACCTGGCAGTTTATTACTGTCAGCAACATTATAGTACTCCTCC |
| IGKV3-12*01_CBA | CBA/J | Musmus IGKV3-12*01 F | 100 | 299 | GACATTGTGCTGACACAGTCTCCTGCTTCCTTAGCTGTATCTCTGGGGCAGAGGGCCACCATCTCATGCAGGGCCAGCAAAAGTGTCAGTACATCTGGCTATAGTTATATGCACTGGTACCAACAGAAACCAGGACAGCCACCCAAACTCCTCATCTATCTTGCATCCAACCTAGAATCTGGGGTCCCTGCCAGGTTCAGTGGCAGTGGGTCTGGGACAGACTTCACCCTCAACATCCATCCTGTGGAGGAGGAGGATGCTGCAACCTATTACTGTCAGCACAGTAGGGAGCTTCCTCC |
| IGKV4-55*01_S6279_CBA | CBA/J | Musmus IGKV4-55*01 F | 100 | 286 | CAAATTGTTCTCACCCAGTCTCCAGCAATCATGTCTGCATCTCCAGGGGAGAAGGTCACCATGACCTGCAGTGCCAGCTCAAGTGTAAGTTACATGTACTGGTACCAGCAGAAGCCAGGATCCTCCCCCAGACTCCTGATTTATGACACATCCAACCTGGCTTCTGGAGTCCCTGTTCGCTTCAGTGGCAGTGGGTCTGGGACCTCTTACTCTCTCACAATCAGCCGAATGGAGGCTGAAGATGCTGCCACTTATTACTGCCAGCAGTGGAGTAGTTACCCACTCA |
| IGKV5-48*01_CBA | CBA/J | Musmus IGKV5-48*01 F | 100 | 287 | GACATCTTGCTGACTCAGTCTCCAGCCATCCTGTCTGTGAGTCCAGGAGAAAGAGTCAGTTTCTCCTGCAGGGCCAGTCAGAGCATTGGCACAAGCATACACTGGTATCAGCAAAGAACAAATGGTTCTCCAAGGCTTCTCATAAAGTATGCTTCTGAGTCTATCTCTGGGATCCCTTCCAGGTTTAGTGGCAGTGGATCAGGGACAGATTTTACTCTTAGCATCAACAGTGTGGAGTCTGAAGATATTGCAGATTATTACTGTCAACAAAGTAATAGCTGGCCAAC |
| IGKV13-85*01_CBA | CBA/J | Musmus IGKV13-85*01 F | 100 | 287 | GACATCCAGATGACACAATCTTCATCCTACTTGTCTGTATCTCTAGGAGGCAGAGTCACCATTACTTGCAAGGCAAGTGACCACATTAATAATTGGTTAGCCTGGTATCAGCAGAAACCAGGAAATGCTCCTAGGCTCTTAATATCTGGTGCAACCAGTTTGGAAACTGGGGTTCCTTCAAGATTCAGTGGCAGTGGATCTGGAAAGGATTACACTCTCAGCATTACCAGTCTTCAGACTGAAGATGTTGCTACTTATTACTGTCAACAGTATTGGAGTACTCCTCC |

| | | | | | |
|---|---|---|---|---|---|
| IGKV19-93*01_CBA | CBA/J | Musmus IGKV19-93*01 F | 100 | 287 | GACATCCAGATGACACAGTCTCCATCCTCACTGTCTGCATCTCTGGGAGGCAAAGTCACCATCACTTGCAAGGCAAGCCAAGACATTAACAAGTATATAGCTTGGTACCAACACAAGCCTGGAAAAGGTCCTAGGCTGCTCATACATTACACATCTACATTACAGCCAGGCATCCCATCAAGGTTCAGTGGAAGTGGGTCTGGGAGAGATTATTCCTTCAGCATCAGCAACCTGGAGCCTGAAGATATTGCAACTTATTATTGTCTACAGTATGATAATCTTCTACC |
| IGKV6-25*01_CBA | CBA/J | Musmus IGKV6-25*01 F | 100 | 287 | GACATTGTGATGACCCAGTCTCACAAATTCATGTCCACATCAGTAGGAGACAGGGTCAGCATCACCTGCAAGGCCAGTCAGGATGTGAGTACTGCTGTAGCCTGGTATCAACAAAAACCAGGGCAATCTCCTAAACTACTGATTTACTGGGCATCCACCCGGCACACTGGAGTCCCTGATCGCTTCACAGGCAGTGGATCTGGGACAGATTATACTCTCACCATCAGCAGTGTGCAGGCTGAAGACCTGGCACTTTATTACTGTCAGCAACATTATAGCACTCCTCC |
| IGKV9-120*01_S2628_CBA | CBA/J | Musmus IGKV9-120*02 F | 100 | 285 | GACATCCAGATGACCCAGTCTCCATCCTCCTTATCTGCCTCTCTGGGAGAAAGAGTCAGTCTCACTTGTCGGGCAAGTCAGGACATTGGTAGTAGCTTAAACTGGCTTCAGCAGGAACCAGATGGAACTATTAAACGCCTGATCTACGCCACATCCAGTTTAGATTCTGGTGTCCCCAAAAGGTTCAGTGGCAGTAGGTCTGGGTCAGATTATTCTCTCACCATCAGCAGCCTTGAGTCTGAAGATTTTGTAGACTATTACTGTCTACAATATGCTAGTTCTCCT |
| IGKV4-59*01_S4622_CBA | CBA/J | Musmus IGKV4-59*01 F | 100 | 286 | CAAATTGTTCTCACCCAGTCTCCAGCAATCATGTCTGCATCTCCAGGGGAGAAGGTCACCATGACCTGCAGTGCCAGCTCAAGTGTAAGTTACATGCACTGGTACCAGCAGAAGTCAGGCACCTCCCCCAAAAGATGGATTTATGACACATCCAAACTGGCTTCTGGAGTCCCTGCTCGCTTCAGTGGCAGTGGGTCTGGGACCTCTTACTCTCTCACAATCAGCAGCATGGAGGCTGAAGATGCTGCCACTTATTACTGCCAGCAGTGGAGTAGTAACCCACTCA |
| IGKV17-121*01_CBA | CBA/J | Musmus IGKV17-121*01 F | 100 | 287 | GAAACAACTGTGACCCAGTCTCCAGCATCCCTGTCCATGGCTATAGGAGAAAAAGTCACCATCAGATGCATAACCAGCACTGATATTGATGATGATATGAACTGGTACCAGCAGAAGCCAGGGGAACCTCCTAAGCTCCTTATTTCAGAAGGCAATACTCTTCGTCCTGGAGTCCCATCCCGATTCTCCAGCAGTGGCTATGGTACAGATTTTGTTTTTACAATTGAAAACATGCTCTCAGAAGATGTTGCAGATTACTACTGTTTGCAAAGTGATAACTTGCCTCT |
| IGKV4-50*01_CBA | CBA/J | Musmus IGKV4-50*01 F | 100 | 286 | GAAAATGTGCTCACCCAGTCTCCAGCAATCATGTCTGCATCTCTAGGGGAGAAGGTCACCATGAGCTGCAGGGCCAGCTCAAGTGTAAATTACATGTACTGGTACCAGCAGAAGTCAGATGCCTCCCCCAAACTATGGATTTATTACACATCCAACCTGGCTCCTGGAGTCCCAGCTCGCTTCAGTGGCAGTGGGTCTGGGAACTCTTATTCTCTCACAATCAGCAGCATGGAGGGTGAAGATGCTGCCACTTATTACTGCCAGCAGTTTACTAGTTCCCCATCCA |
| IGKV4-68*01_S3092_CBA | CBA/J | Musmus IGKV4-68*01 F | 100 | 286 | CAAATTGTTCTCACCCAGTCTCCAGCACTCATGTCTGCATCTCCAGGGGAGAAGGTCACCATGACCTGCAGTGCCAGCTCAAGTGTAAGTTACATGTACTGGTACCAGCAGAAGCCAAGATCCTCCCCCAAACCCTGGATTTATCTCACATCCAACCTGGCTTCTGGAGTCCCTGCTCGCTTCAGTGGCAGTGGGTCTGGGACCTCTTACTCTCTCACAATCAGCAGCATGGAGGCTGAAGATGCTGCCACTTATTACTGCCAGCAGTGGAGTAGTAACCCACTCA |
| IGKV16-104*01_CBA | CBA/J | Musmus IGKV16-104*01 F | 100 | 288 | GATGTCCAGATAACCCAGTCTCCATCTTATCTTGCTGCATCTCCTGGAGAAACCATTACTATTAATTGCAGGGCAAGTAAGAGCATTAGCAAATATTTAGCCTGGTATCAAGAGAAACCTGGGAAAACTAATAAGCTTCTTATCTACTCTGGATCCACTTTGCAATCTGGAATTCCATCAAGGTTCAGTGGCAGTGGATCTGGTACAGATTTCACTCTCACCATCAGTAGCCTGGAGCCTGAAGATTTTGCAATGTATTACTGTCAACAGCATAATGAATACCCGTAC |

| | | | | | |
|---|---|---|---|---|---|
| IGKV2-109*01_CBA | CBA/J | Musmus IGKV2-109*01 F | 100 | 302 | GATATTGTGATGACGCAGGCTGCATTCTCCAATCCAGTCACTCTTGGAACATCAGCTTCCATCTCCTGCAGGTCTAGTAAGAGTCTCCTACATAGTAATGGCATCACTTATTTGTATTGGTATCTGCAGAAGCCAGGCCAGTCTCCTCAGCTCCTGATTTATCAGATGTCCAACCTTGCCTCAGGAGTCCCAGACAGGTTCAGTAGCAGTGGGTCAGGAACTGATTTCACACTGAGAATCAGCAGAGTGGAGGCTGAGGATGTGGGTGTTTATTACTGTGCTCAAAATCTAGAACTTCCTCC |
| IGKV4-79*01_CBA | CBA/J | Musmus IGKV4-79*01 F | 100 | 292 | CAAATTGTTCTCACCCAGTCTCCAGCAATCATGTCTGCATCTCCTGGGGAGAAGGTCACCTTGACCTGCAGTGCCAGCTCAAGTGTAAGTTCCAGCTACTTGTACTGGTACCAGCAGAAGCCAGGATCCTCCCCCAAACTCTGGATTTATAGCACATCCAACCTGGCTTCTGGAGTCCCTGCTCGCTTCAGTGGCAGTGGGTCTGGGACCTCTTACTCTCTCACAATCAGCAGCATGGAGGCTGAAGATGCTGCCTCTTATTTCTGCCATCAGTGGAGTAGTTACCCACCCC |
| IGKV13-84*01_CBA | CBA/J | Musmus IGKV13-84*01 F | 100 | 287 | GACATCCAGATGACACAATCTTCATCCTCCTTTTCTGTATCTCTAGGAGACAGAGTCACCATTACTTGCAAGGCAAGTGAGGACATATATAATCGGTTAGCCTGGTATCAGCAGAAACCAGGAAATGCTCCTAGGCTCTTAATATCTGGTGCAACCAGTTTGGAAACTGGGGTTCCTTCAAGATTCAGTGGCAGTGGATCTGGAAAGGATTACACTCTCAGCATTACCAGTCTTCAGACTGAAGATGTTGCTACTTATTACTGTCAACAGTATTGGAGTACTCCTCC |
| IGKV3-2*01_CBA | CBA/J | Musmus IGKV3-2*01 F | 100 | 299 | GACATTGTGCTGACCCAATCTCCAGCTTCTTTGGCTGTGTCTCTAGGGCAGAGGGCCACCATCTCCTGCAGAGCCAGCGAAAGTGTTGATAATTATGGCATTAGTTTTATGAACTGGTTCCAACAGAAACCAGGACAGCCACCCAAACTCCTCATCTATGCTGCATCCAACCAAGGATCCGGGGTCCCTGCCAGGTTTAGTGGCAGTGGGTCTGGGACAGACTTCAGCCTCAACATCCATCCTATGGAGGAGGATGATACTGCAATGTATTTCTGTCAGCAAAGTAAGGAGGTTCCTCC |
| IGKV4-74*01_CBA | CBA/J | Musmus IGKV4-74*01 F | 100 | 292 | CAAATTGTTCTCACCCAGTCTCCAGCAATCATGTCTGCATCTCTAGGGGAACGGGTCACCATGACCTGCACTGCCAGCTCAAGTGTAAGTTCCAGTTACTTGCACTGGTACCAGCAGAAGCCAGGATCCTCCCCCAAACTCTGGATTTATAGCACATCCAACCTGGCTTCTGGAGTCCCAGCTCGCTTCAGTGGCAGTGGGTCTGGGACCTCTTACTCTCTCACAATCAGCAGCATGGAGGCTGAAGATGCTGCCACTTATTACTGCCACCAGTATCATCGTTCCCCACCCA |
| IGKV4-63*01_S2117_CBA | CBA/J | Musmus IGKV4-63*01 F | 100 | 286 | GAAAATGTTCTCACCCAGTCTCCAGCAATCATGTCTGCATCTCCAGGGGAAAAGGTCACCATGACCTGCAGTGCCAGCTCAAGTGTAAGTTACATGCACTGGTACCAGCAGAAGTCAAGCACCTCCCCCAAACTCTGGATTTATGACACATCCAAACTGGCTTCTGGAGTCCCAGGTCGCTTCAGTGGCAGTGGGTCTGGAAACTCTTACTCTCTCACGATCAGCAGCATGGAGGCTGAAGATGTTGCCACTTATTACTGTTTTCAGGGGAGTGGGTACCCATTCA |
| IGKV3-10*01_CBA | CBA/J | Musmus IGKV3-10*01 F | 100 | 299 | AACATTGTGCTGACCCAATCTCCAGCTTCTTTGGCTGTGTCTCTAGGGCAGAGGGCCACCATATCCTGCAGAGCCAGTGAAAGTGTTGATAGTTATGGCAATAGTTTTATGCACTGGTACCAGCAGAAACCAGGACAGCCACCCAAACTCCTCATCTATCTTGCATCCAACCTAGAATCTGGGGTCCCTGCCAGGTTCAGTGGCAGTGGGTCTAGGACAGACTTCACCCTCACCATTGATCCTGTGGAGGCTGATGATGCTGCAACCTATTACTGTCAGCAAAATAATGAGGATCCTCC |

| | | | | | |
|---|---|---|---|---|---|
| IGKV4-61*01_S2955_CBA | CBA/J | Musmus IGKV4-61*01 F | 100 | 286 | CAAATTGTTCTCACCCAGTCTCCAGCAATCATGTCTGCATCTCCAGGGGAGAAGGTCACCATATCCTGCAGTGCCAGCTCAAGTGTAAGTTACATGTACTGGTACCAGCAGAAGCCAGGATCCTCCCCCAAACCCTGGATTTATCGCACATCCAACCTGGCTTCTGGAGTCCCTGCTCGCTTCAGTGGCAGTGGGTCTGGGACCTCTTACTCTCTCACAATCAGCAGCATGGAGGCTGAAGATGCTGCCACTTATTACTGCCAGCAGTATCATAGTTACCCACTCA |
| IGKV4-69*01_CBA | CBA/J | Musmus IGKV4-69*01 F | 100 | 282 | CAAATTCTTCTCACCCAGTCTCCAGCAATCATGTCTGCATCTCCAGGGGAGAAGGTCACCATGACCTGCAGTGCCAGCTCAAGTGTAAGTTACATGCACTGGTACCAGCAGAAGCCAGGATCCTCGCCCAAACCCTGGATTTATGACACATCCAACCTGGCTTCTGGATTCCCTGCTCGCTTCAGTGGCAGTGGGTCTGGGACCTCTTACTCTCTCATAATCAGCAGCATGGAGGCTGAAGATGCTGCCACTTATTACTGCCATCAGCGGAGTAGTTACCCA |
| IGKV1-122*01_CBA | CBA/J | Musmus IGKV1-122*01 F | 100 | 302 | GATGCTGTGATGACCCAAACTCCACTCTCCCTGCCTGTCAGTCTTGGAGATCAAGCCTCCATCTCTTGCAGGTCTAGTCAGAGCCTTGAAAACAGTAATGGAAACACCTATTTGAACTGGTACCTCCAGAAACCAGGCCAGTCTCCACAGCTCCTGATCTACAGGGTTTCCAACCGATTTTCTGGGGTCCTAGACAGGTTCAGTGGTAGTGGATCAGGGACAGATTTCACACTGAAAATCAGCAGAGTGGAGGCTGAGGATTTGGGAGTTTATTTCTGCCTCCAAGTTACACATGTCCCTCC |
| IGKV1-133*01_CBA | CBA/J | Musmus IGKV1-133*01 F | 100 | 302 | GATGTTGTGATGACCCAGACTCCACTCACTTTGTCGGTTACCATTGGACAACCAGCCTCTATCTCTTGCAAGTCAAGTCAGAGCCTCTTATATAGTAATGGAAAAACCTATTTGAATTGGTTATTACAGAGGCCAGGCCAGTCTCCAAAGCGCCTAATCTATCTGGTGTCTAAACTGGACTCTGGAGTCCCTGACAGGTTCACTGGCAGTGGATCAGGAACAGATTTTACACTGAAAATCAGCAGAGTGGAGGCTGAGGATTTGGGAGTTTATTACTGCGTGCAAGGTACACATTTTCCTCA |
| IGKV14-100*01_CBA | CBA/J | Musmus IGKV14-100*01 F | 100 | 287 | GACATCCTGATGACCCAATCTCCATCCTCCATGTCTGTATCTCTGGGAGACACAGTCAGCATCACTTGCCATGCAAGTCAGGGCATTAGCAGTAATATAGGGTGGTTGCAGCAGAAACCAGGGAAATCATTTAAGGGCCTGATCTATCATGGAACCAACTTGGAAGATGGAGTTCCATCAAGGTTCAGTGGCAGTGGATCTGGAGCAGATTATTCTCTCACCATCAGCAGCCTGGAATCTGAAGATTTTGCAGACTATTACTGTGTACAGTATGCTCAGTTTCCTCC |
| IGKV4-92*01_CBA | CBA/J | Musmus IGKV4-92*01 F | 100 | 291 | GAAATGGTTCTCACCCAGTCTCCAGTATCCATAACTGCATCTCGAGGGGAGAAGGTCACCATCACCTGCCGTGCCAGCTCAAGTATAAGTTCCAATTACTTACACTGGTACCAGCAGAAGCCAGGATCCTCCCCTAAACTTTTGATTTATAGGACATCCATCCTGGCATCTGGAGTCCTAGACAGCTTCAGTGGCAGTGGGTCTGAGAGCTCTTACACTCTGACAATCAGCTGCATGCAGGACGAAGTTGCTGCCACTTACTATTGTCAGCAGGGGAGTAGTAGCCCACCA |
| IGKV8-34*01_CBA | CBA/J | Musmus IGKV8-34*01 F | 100 | 305 | GACATTTTGATGACCCAGTCTCCATCCTCCCTGACTGTGTCAGCAGGAGAGAAGGTCACTATGAGCTGCAAGTCCAGTCAGAGTCTTTTAGCTAGTGGCAACCAAAATAACTACTTGGCCTGGCACCAGCAGAAACCAGGACGATCTCCTAAAATGCTGATAATTTGGGCATCCACTAGGGTATCTGGAGTCCCTGATCGCTTCATAGGCAGTGGATCTGGGACGGATTTCACTCTGACCATCAACAGTGTGCAGGCTGAAGATCTGGCTGTTTATTACTGTCAGCAGTCCTACAGCGCTCCTAC |
| IGKV9-124*01_CBA | CBA/J | Musmus IGKV9-124*01 F | 100 | 285 | GACATCCAGATGACCCAGTCTCCATCCTCCTTATCTGCCTCTCTGGGAGAAAGAGTCAGTCTCACTTGTCGGGCAAGTCAGGAAATTAGTGGTTACTTAAGCTGGCTTCAGCAGAAACCAGATGGAACTATTAAACGCCTGATCTACGCCGCATCCACTTTAGATTCTGGTGTCCCAAAAAGGTTCAGTGGCAGTAGGTCTGGGTCAGATTATTCTCTCACCATCAGCAGCCTTGAGTCTGAAGATTTTGCAGACTATTACTGTCTACAATATGCTAGTTATCCT |

| | | | | | |
|---|---|---|---|---|---|
| IGKV5-39*01_CBA | CBA/J | Musmus IGKV5-39*01 F | 100 | 287 | GACATTGTGATGACTCAGTCTCCAGCCACCCTGTCTGTGACTCCAGGAGATAGAGTCTCTCTTTCCTGCAGGGCCAGCCAGAGTATTAGCGACTACTTACACTGGTATCAACAAAAATCACATGAGTCTCCAAGGCTTCTCATCAAATATGCTTCCCAATCCATCTCTGGGATCCCCTCCAGGTTCAGTGGCAGTGGATCAGGGTCAGATTTCACTCTCAGTATCAACAGTGTGGAACCTGAAGATGTTGGAGTGTATTACTGTCAAAATGGTCACAGCTTTCCTCC |
| IGKV4-86*01_CBA | CBA/J | Musmus IGKV4-86*01 F | 100 | 283 | GAAATTGTGCTCACTCAGTCTCCAGCCATCACAGCTGCATCTCTGGGGCAAAAGGTCACCATCACCTGCAGTGCCAGCTCAAGTGTAAGTTACATGCACTGGTACCAGCAGAAGTCAGGCACCTCCCCCAAACCATGGATTTATGAAATATCCAAACTGGCTTCTGGAGTCCCAGCTCGCTTCAGTGGCAGTGGGTCTGGGACCTCTTACTCTCTCACAATCAGCAGCATGGAGGCTGAAGATGCTGCCATTTATTACTGCCAGCAGTGGAATTATCCTCTTA |
| IGKV11-125*01_CBA | CBA/J | Musmus IGKV11-125*01 F | 100 | 287 | GATGTCCAGATGATTCAGTCTCCATCCTCCCTGTCTGCATCTTTGGGAGACATAGTCACCATGACTTGCCAGGCAAGTCAGGGCACTAGCATTAATTTAAACTGGTTTCAGCAAAAACCAGGGAAAGCTCCTAAGCTCCTGATCTATGGTGCAAGCAACTTGGAAGATGGGGTCCCATCAAGGTTCAGTGGCAGTAGATATGGGACAGATTTCACTCTCACCATCAGCAGCCTGGAGGATGAAGATATGGCAACTTATTTCTGTCTACAGCATAGTTATCTCCCTCC |
| IGKV4-58*01_CBA | CBA/J | Musmus IGKV4-58*01 F | 100 | 292 | GAAAATGTGCTCACCCAGTCTCCAGCAATAATGGCTGCCTCTCTGGGGCAGAAGGTCACCATGACCTGCAGTGCCAGCTCAAGTGTAAGTTCCAGTTACTTGCACTGGTACCAGCAGAAGTCAGGCGCTTCCCCCAAACCCTTGATTCATAGGACATCCAACCTGGCTTCTGGAGTCCCAGCTCGCTTCAGTGGCAGTGGGTCTGGGACCTCTTACTCTCTCACAATCAGCAGCGTGGAGGCTGAAGATGATGCAACTTATTACTGCCAGCAGTGGAGTGGTTACCCATTCA |
| IGKV12-38*01_CBA | CBA/J | Musmus IGKV12-38*01 F | 100 | 287 | GACATCCAGATGACTCAGTCTCCAGCCTCCCTGGCTGCATCTGTGGGAGAAACTGTCACCATCACATGTCGAGCAAGTGAGAACATTTACTACAGTTTAGCATGGTATCAGCAGAAGCAAGGGAAATCTCCTCAGCTCCTGATCTATAATGCAAACAGCTTGGAAGATGGTGTCCCATCGAGGTTCAGTGGCAGTGGATCTGGGACACAGTATTCTATGAAGATCAACAGCATGCAGCCTGAAGATACCGCAACTTATTTCTGTAAACAGGCTTATGACGTTCCTCC |
| IGKV8-23-1*01_CBA | CBA/J | Musmus IGKV8-16*01 F, or Musmus IGKV8-23-1*01 ORF | 100 | 284 | GAAATTGTGCTGACTGTGTCAGCAGGAGAGAGGGTCACTATCAGCTGCAAATCCAATCAGAATCTTTTATGGAGTGGAAACCAAAGGTACTGTTTGGTCTGGCACCAGTGGAAACCAGGGCAAACTCCTACACCGTTGATCACCTGGACATCTGATAGGTACTCTGGAGTCCCTGATCGTTTCATAGGCAGTGGATCTGTGACAGATTTCACTCTGACCATCAGCAGTGTGCAGGCTGAAGATGTGGCAGTTTATTTCTGTCAGCAGCATTTACACATTCCTCC |
| IGKV14-111*01_CBA | CBA/J | Musmus IGKV14-111*01 F | 100 | 287 | GACATCAAGATGACCCAGTCTCCATCTTCCATGTATGCATCTCTAGGAGAGAGAGTCACTATCACTTGCAAGGCGAGTCAGGACATTAATAGCTATTTAAGCTGGTTCCAGCAGAAACCAGGGAAATCTCCTAAGACCCTGATCTATCGTGCAAACAGATTGGTAGATGGGGTCCCATCAAGGTTCAGTGGCAGTGGATCTGGGCAAGATTATTCTCTCACCATCAGCAGCCTGGAGTATGAAGATATGGGAATTTATTATTGTCTACAGTATGATGAGTTTCCTCC |
| IGKV9-123*01_CBA | CBA/J | Musmus IGKV9-123*01 F | 100 | 285 | GACATCCAGATGATTCAGTCTCCATCGTCCATGTTTGCCTCTCTGGGAGACAGAGTCAGTCTCTCTTGTCGGGCTAGTCAGGGCATTAGAGGTAATTTAGACTGGTATCAGCAGAAACCAGGTGGAACTATTAAACTCCTGATCTACTCCACATCCAATTTAAATTCTGGTGTCCCATCAAGGTTCAGTGGCAGTGGGTCTGGGTCAGATTATTCTCTCACCATCAGCAGCCTAGAGTCTGAAGATTTTGCAGACTATTACTGTCTACAGCGTAATGCGTATCCT |

| | | | | | |
|---|---|---|---|---|---|
| IGKV14-130*01_CBA | CBA/J | Musmus IGKV14-130*01 F | 100 | 287 | GAAATCCAGATGACCCAGTCTCCATCCTCTATGTCTGCATCTCTGGGAGACAGAATAACCATCACTTGCCAGGCAACTCAAGACATTGTTAAGAATTTAAACTGGTATCAGCAGAAACCAGGGAAACCCCCTTCATTCCTGATCTATTATGCAACTGAACTGGCAGAAGGGGTCCCATCAAGGTTCAGTGGCAGTGGGTCTGGGTCAGACTATTCTCTGACAATCAGCAACCTGGAGTCTGAAGATTTTGCAGACTATTACTGTCTACAGTTTTATGAGTTTCCTCC |
| IGKV10-95*01_CBA | CBA/J | Musmus IGKV10-95*01 F | 100 | 287 | GATATCCAGATGACACAGACTACTTCCTCCCTGTCTGCCTCTCTGGGAGACAGAGTCACCATCAGTTGCAGGGCAAGTGAGGACATTAGCACTTATTTAAACTGGTATCAGCAGAAACCAGATGGAACTGTTAAACTCCTGATCTATTACACATCAGGATTACACTCAGGAGTCCCATCAAGGTTCAGTGGCAGTGGGTCTGGGGCAGATTATTCTCTCACCATCAGCAACCTGGAACCTGAAGATATTGCCACTTACTATTGTCAGCAGTATAGTAAGCTTCCTCC |
| IGKV2-112*01_CBA | CBA/J | Musmus IGKV2-112*01 F | 100 | 300 | GATATTGTGATAACCCAGGATGAACTCTCCAATCCTGTCACTTCTGGAGAATCAGTTTCCATCTCCTGCAGGTCTAGTAAGAGTCTCCTATATAAGGATGGGAAGACATACTTGAATTGGTTTCTGCAGAGACCAGGACAATCTCCTCAGCTCCTGATCTATTTGATGTCCACCCGTGCATCAGGAGTCTCAGACCGGTTTAGTGGCAGTGGGTCAGGAACAGATTTCACCCTGGAAATCAGTAGAGTGAAGGCTGAGGATGTGGGTGTGTATTACTGTCAACAACTTGTAGAGTATCCT |
| IGKV4-56*01_CBA | CBA/J | Musmus IGKV4-56*01 P | 100 | 286 | CAAATTGTTCTCACCCAGTCTCCAGCAATCATGTCTGCATCTCCAGGGCAGAAAGTCACCATAACCTGCAGTGCCATCTCAAGTGTAAATTACATGCACTGGTACCAGCAGAAGCCAGGATCCTCCCCCAAACTCTGGATTTATGCAACATCCAAACTGGCTCTTGGAGTCCCTGCTTGCTTCAGTGGCAGTGGGTCTGGGACCTCTTACTCTCTCACAATCAGCAGCATGGTGGCTGAAGATGCCACCTCTTATTTCTGTCATCAGTGGAGTAGTTACCCACCCA |
| IGKV4-74*01_S2922_CBA | CBA/J | Musmus IGKV4-74*01 F | 100 | 292 | CAAATTGTTCTCACCCAGTCTCCAGCAATCATGTCTGCATCTCTAGGGGAACGGGTCACCATGACCTGCACTGCCAGCTCAAGTGTAAGTTCCAGTTACTTGCACTGGTACCAGCAGAAGCCAGGATCCTCCCCCAAACTCTGGATTTATAGCACATCCAACCTGGCTTCTGGAGTCCCAGCTCGCTTCAGTGGCAGTGGGTCTGGGACCTCTTACTCTCTCACAATCAGCAGCATGGAGGCTGAAGATGCTGCCACTTATTACTGCCACCAGTATCATCGTTCCCCACTCA |
| IGKV4-78*01_S5980_CBA | CBA/J | Musmus IGKV4-78*01 F | 100 | 292 | CAAATTGTTCTCACCCAGTCTCCAGCAATCATGTCTGCATCTCCTGGGGAGAAGGTCACCATGACCTGCAGTGCCAGATCAAGTGTAAGTTCCAGCTACTTGTACTGGTACCAGCAGAAGCCAGGATCCTCCCCCAAACTCTGGATTTATAGCACATCCAACCTGGCTTCTGGAGTCCCTGCTCGCTTCAGTGGCAGTGGGTCTGGGACCTCTTATTCTCTCACAATCAGCAGCATGGAGGCTGAAGATGCTGCCACTTTTTACTGCCAGCAGTACAGTGGTTACCCATTCA |
| IGKV4-51*01_CBA | CBA/J | Musmus IGKV4-51*01 F | 100 | 292 | GAAAATGTGCTCACCCAGTCTCCAGCAATAATGGCTGCCTCTCTGGGGGAGAAGGTCACCATGACCTGCAGTGCCAGCTCAAGTGTAAGTTCCAGCTACTTGCACTGGTACCAGCAGAAGTCAGGCACTTCCCCCAAACTCTGGATTTATGGCACATCCAACCTGGCTTCTGGAGTCCCAGCTCGCTTCAGTGGCAGTGGGGCTGGGATCTCTTACTCTCTCACAATCAGCAGCATGGAGGCTGAAAATGATGCAACTTATTACTGCCAGCAGTGGAGTGGTTACCCATTCA |

| | | | | | |
|---|---|---|---|---|---|
| IGKV8-30*01_DBA1 | DBA/1J | Musmus IGKV8-30*01 F | 100 | 305 | GACATTGTGATGTCACAGTCTCCATCCTCCCTAGCTGTGTCAGTTGGAGAGAAGGTTACTATGAGCTGCAAGTCCAGTCAGAGCCTTTTATATAGTAGCAATCAAAAGAACTACTTGGCCTGGTACCAGCAGAAACCAGGGCAGTCTCCTAAACTGCTGATTTACTGGGCATCCACTAGGGAATCTGGGGTCCCTGATCGCTTCACAGGCAGTGGATCTGGGACAGATTTCACTCTCACCATCAGCAGTGTGAAGGCTGAAGACCTGGCAGTTTATTACTGTCAGCAATATTATAGCTATCCTCC |
| IGKV6-15*01_DBA1 | DBA/1J | Musmus IGKV6-15*01 F | 100 | 287 | GACATTGTGATGACCCAGTCTCAAAAATTCATGTCCACATCAGTAGGAGACAGGGTCAGCGTCACCTGCAAGGCCAGTCAGAATGTGGGTACTAATGTAGCCTGGTATCAACAGAAACCAGGGCAATCTCCTAAAGCACTGATTTACTCGGCATCCTACCGGTACAGTGGAGTCCCTGATCGCTTCACAGGCAGTGGATCTGGGACAGATTTCACTCTCACCATCAGCAATGTGCAGTCTGAAGACTTGGCAGAGTATTTCTGTCAGCAATATAACAGCTATCCTCT |
| IGKV8-27*01_DBA1 | DBA/1J | Musmus IGKV8-27*01 F | 100 | 300 | AACATTATGATGACACAGTCGCCATCATCTCTGGCTGTGTCTGCAGGAGAAAAGGTCACTATGAGCTGTAAGTCCAGTCAAAGTGTTTTATACAGTTCAAATCAGAAGAACTACTTGGCCTGGTACCAGCAGAAACCAGGGCAGTCTCCTAAACTGCTGATCTACTGGGCATCCACTAGGGAATCTGGTGTCCCTGATCGCTTCACAGGCAGTGGATCTGGGACAGATTTTACTCTTACCATCAGCAGTGTACAAGCTGAAGACCTGGCAGTTTATTACTGTCATCAATACCTCTCCTCG |
| IGKV12-44*01_DBA1 | DBA/1J | Musmus IGKV12-44*01 F | 100 | 287 | GACATCCAGATGACTCAGTCTCCAGCCTCCCTATCTGCATCTGTGGGAGAAACTGTCACCATCACATGTCGAGCAAGTGAGAATATTTACAGTTATTTAGCATGGTATCAGCAGAAACAGGGAAAATCTCCTCAGCTCCTGGTCTATAATGCAAAAACCTTAGCAGAAGGTGTGCCATCAAGGTTCAGTGGCAGTGGATCAGGCACACAGTTTTCTCTGAAGATCAACAGCCTGCAGCCTGAAGATTTTGGGAGTTATTACTGTCAACATCATTATGGTACTCCTCC |
| IGKV1-135*01_DBA1 | DBA/1J | Musmus IGKV1-135*01 F | 100 | 302 | GATGTTGTGATGACCCAGACTCCACTCACTTTGTCGGTTACCATTGGACAACCAGCCTCCATCTCTTGCAAGTCAAGTCAGAGCCTCTTAGATAGTGATGGAAAGACATATTTGAATTGGTTGTTACAGAGGCCAGGCCAGTCTCCAAAGCGCCTAATCTATCTGGTGTCTAAACTGGACTCTGGAGTCCCTGACAGGTTCACTGGCAGTGGATCAGGGACAGATTTCACACTGAAAATCAGCAGAGTGGAGGCTGAGGATTTGGGAGTTTATTATTGCTGGCAAGGTACACATTTTCCTCA |
| IGKV10-96*01_DBA1 | DBA/1J | Musmus IGKV10-96*01 F | 100 | 286 | GATATCCAGATGACACAGACTACATCCTCCCTGTCTGCCTCTCTGGGAGACAGAGTCACCATCAGTTGCAGGGCAAGTCAGGACATTAGCAATTATTTAAACTGGTATCAGCAGAAACCAGATGGAACTGTTAAACTCCTGATCTACTACACATCAAGATTACACTCAGGAGTCCCATCAAGGTTCAGTGGCAGTGGGTCTGGAACAGATTATTCTCTCACCATTAGCAACCTGGAGCAAGAAGATATTGCCACTTACTTTTGCCAACAGGGTAATACGCTTCCTC |
| IGKV4-59*01_DBA1 | DBA/1J | Musmus IGKV4-59*01 F | 100 | 286 | CAAATTGTTCTCACCCAGTCTCCAGCAATCATGTCTGCATCTCCAGGGGAGAAGGTCACCATGACCTGCAGTGCCAGCTCAAGTGTAAGTTACATGCACTGGTACCAGCAGAAGTCAGGCACCTCCCCCAAAAGATGGATTTATGACACATCCAAACTGGCTTCTGGAGTCCCTGCTCGCTTCAGTGGCAGTGGGTCTGGGACCTCTTACTCTCTCACAATCAGCAGCATGGAGGCTGAAGATGCTGCCACTTATTACTGCCAGCAGTGGAGTAGTAACCCACCCA |
| IGKV2-137*01_DBA1 | DBA/1J | Musmus IGKV2-137*01 F | 100 | 302 | GATATTGTGATGACTCAGGCTGCACCCTCTGTACCTGTCACTCCTGGAGAGTCAGTATCCATCTCCTGCAGGTCTAGTAAGAGTCTCCTGCATAGTAATGGCAACACTTACTTGTATTGGTTCCTGCAGAGGCCAGGCCAGTCTCCTCAGCTCCTGATATATCGGATGTCCAACCTTGCCTCAGGAGTCCCAGACAGGTTCAGTGGCAGTGGGTCAGGAACTGCTTTCACACTGAGAATCAGTAGAGTGGAGGCTGAGGATGTGGGTGTTTATTACTGTATGCAACATCTAGAATATCCTTT |

| | | | | | |
|---|---|---|---|---|---|
| IGKV8-19*01_DBA1 | DBA/1J | Musmus IGKV8-19*01 F | 100 | 305 | GACATTGTGATGACACAGTCTCCATCCTCCCTGACTGTGACAGCAGGAGAGAAGGTCACTATGAGCTGCAAGTCCAGTCAGAGTCTGTTAAACAGTGGAAATCAAAAGAACTACTTGACCTGGTACCAGCAGAAACCAGGGCAGCCTCCTAAACTGTTGATCTACTGGGCATCCACTAGGGAATCTGGGGTCCCTGATCGCTTCACAGGCAGTGGATCTGGAACAGATTTCACTCTCACCATCAGCAGTGTGCAGGCTGAAGACCTGGCAGTTTATTACTGTCAGAATGATTATAGTTATCCTCC |
| IGKV1-110*01_DBA1 | DBA/1J | Musmus IGKV1-110*01 F | 100 | 302 | GATGTTGTGATGACCCAAACTCCACTCTCCCTGCCTGTCAGTCTTGGAGATCAAGCCTCCATCTCTTGCAGATCTAGTCAGAGCCTTGTACACAGTAATGGAAACACCTATTTACATTGGTACCTGCAGAAGCCAGGCCAGTCTCCAAAGCTCCTGATCTACAAAGTTTCCAACCGATTTTCTGGGGTCCCAGACAGGTTCAGTGGCAGTGGATCAGGGACAGATTTCACACTCAAGATCAGCAGAGTGGAGGCTGAGGATCTGGGAGTTTATTTCTGCTCTCAAAGTACACATGTTCCTCC |
| IGKV6-23*01_DBA1 | DBA/1J | Musmus IGKV6-23*01 F | 100 | 287 | GACATTGTGATGACCCAGTCTCACAAATTCATGTCCACATCAGTAGGAGACAGGGTCAGCATCACCTGCAAGGCCAGTCAGGATGTGGGTACTGCTGTAGCCTGGTATCAACAGAAACCAGGGCAATCTCCTAAACTACTGATTTACTGGGCATCCACCCGGCACACTGGAGTCCCTGATCGCTTCACAGGCAGTGGATCTGGGACAGATTTCACTCTCACCATTAGCAATGTGCAGTCTGAAGACTTGGCAGATTATTTCTGTCAGCAATATAGCAGCTATCCTCT |
| IGKV1-117*01_DBA1 | DBA/1J | Musmus IGKV1-117*01 F | 100 | 302 | GATGTTTTGATGACCCAAACTCCACTCTCCCTGCCTGTCAGTCTTGGAGATCAAGCCTCCATCTCTTGCAGATCTAGTCAGAGCATTGTACATAGTAATGGAAACACCTATTTAGAATGGTACCTGCAGAAACCAGGCCAGTCTCCAAAGCTCCTGATCTACAAAGTTTCCAACCGATTTTCTGGGGTCCCAGACAGGTTCAGTGGCAGTGGATCAGGGACAGATTTCACACTCAAGATCAGCAGAGTGGAGGCTGAGGATCTGGGAGTTTATTACTGCTTTCAAGGTTCACATGTTCCTCC |
| IGKV12-41*01_DBA1 | DBA/1J | Musmus IGKV12-41*01 F | 100 | 287 | GACATCCAGATGACTCAGTCTCCAGCCTCCCTATCTGCATCTGTGGGAGAAACTGTCACCATCACATGTCGAGCAAGTGGGAATATTCACAATTATTTAGCATGGTATCAGCAGAAACAGGGAAAATCTCCTCAGCTCCTGGTCTATAATGCAAAAACCTTAGCAGATGGTGTGCCATCAAGGTTCAGTGGCAGTGGATCAGGAACACAATATTCTCTCAAGATCAACAGCCTGCAGCCTGAAGATTTTGGGAGTTATTACTGTCAACATTTTTGGAGTACTCCTCC |
| IGKV8-21*01_DBA1 | DBA/1J | Musmus IGKV8-21*01 F | 100 | 302 | GACATTGTGATGTCACAGTCTCCATCCTCCCTGGCTGTGTCAGCAGGAGAGAAGGTCACTATGAGCTGCAAATCCAGTCAGAGTCTGCTCAACAGTAGAACCCGAAAGAACTACTTGGCTTGGTACCAGCAGAAACCAGGGCAGTCTCCTAAACTGCTGATCTACTGGGCATCCACTAGGGAATCTGGGGTCCCTGATCGCTTCACAGGCAGTGGATCTGGGACAGATTTCACTCTCACCATCAGCAGTGTGCAGGCTGAAGACCTGGCAGTTTATTACTGCAAGCAATCTTATAATCTTCC |
| IGKV8-24*01_DBA1 | DBA/1J | Musmus IGKV8-24*01 F | 100 | 305 | GACATTGTGATGACACAGTCTCCATCCTCCCTGGCTATGTCAGTAGGACAGAAGGTCACTATGAGCTGCAAGTCCAGTCAGAGCCTTTTAAATAGTAGCAATCAAAAGAACTATTTGGCCTGGTACCAGCAGAAACCAGGACAGTCTCCTAAACTTCTGGTATACTTTGCATCCACTAGGGAATCTGGGGTCCCTGATCGCTTCATAGGCAGTGGATCTGGGACAGATTTCACTCTTACCATCAGCAGTGTGCAGGCTGAAGACCTGGCAGATTACTTCTGTCAGCAACATTATAGCACTCCTCC |

| | | | | | |
|---|---|---|---|---|---|
| IGKV7-33*01_DBA1 | DBA/1J | Musmus IGKV7-33*01 F | 100 | 305 | GACATTGTGATGACTCAGTCTCCAACTTTCCTTGCTGTGACAGCAAGTAAGAAGGTCACCATTAGTTGCACGGCCAGTGAGAGCCTTTATTCAAGCAAACACAAGGTGCACTACTTGGCTTGGTACCAGAAGAAACCAGAGCAATCTCCTAAACTGCTGATATACGGGGCATCCAACCGATACATTGGGGTCCCTGATCGCTTCACAGGCAGTGGATCTGGGACAGATTTCACTCTGACCATCAGCAGTGTACAGGTTGAAGACCTCACACATTATTACTGTGCACAGTTTTACAGCTATCCTCT |
| IGKV8-28*01_DBA1 | DBA/1J | Musmus IGKV8-28*01 F | 100 | 305 | GACATTGTGATGACACAGTCTCCATCCTCCCTGAGTGTGTCAGCAGGAGAGAAGGTCACTATGAGCTGCAAGTCCAGTCAGAGTCTGTTAAACAGTGGAAATCAAAAGAACTACTTGGCCTGGTACCAGCAGAAACCAGGGCAGCCTCCTAAACTGTTGATCTACGGGGCATCCACTAGGGAATCTGGGGTCCCTGATCGCTTCACAGGCAGTGGATCTGGAACCGATTTCACTCTTACCATCAGCAGTGTGCAGGCTGAAGACCTGGCAGTTTATTACTGTCAGAATGATCATAGTTATCCTCC |
| IGKV6-20*01_DBA1 | DBA/1J | Musmus IGKV6-20*01 F | 100 | 287 | AACATTGTAATGACCCAATCTCCCAAATCCATGTCCATGTCAGTAGGAGAGAGGGTCACCTTGAGCTGCAAGGCCAGTGAGAATGTGGGTACTTATGTATCCTGGTATCAACAGAAACCAGAGCAGTCTCCTAAACTGCTGATATACGGGGCATCCAACCGGTACACTGGGGTCCCCGATCGCTTCACAGGCAGTGGATCTGCAACAGATTTCACTCTGACCATCAGCAGTGTGCAGGCTGAAGACCTTGCAGATTATCACTGTGGACAGAGTTACAGCTATCCTCC |
| IGKV12-46*01_DBA1 | DBA/1J | Musmus IGKV12-46*01 F | 100 | 287 | GACATCCAGATGACTCAGTCTCCAGCCTCCCTATCTGTATCTGTGGGAGAAACTGTCACCATCACATGTCGAGCAAGTGAGAATATTTACAGTAATTTAGCATGGTATCAGCAGAAACAGGGAAAATCTCCTCAGCTCCTGGTCTATGCTGCAACAAACTTAGCAGATGGTGTGCCATCAAGGTTCAGTGGCAGTGGATCAGGCACACAGTATTCCCTCAAGATCAACAGCCTGCAGTCTGAAGATTTTGGGAGTTATTACTGTCAACATTTTTGGGGTACTCCTCC |
| IGKV4-72*01_DBA1 | DBA/1J | Musmus IGKV4-72*01 F | 100 | 286 | CAAATTGTTCTCTCCCAGTCTCCAGCAATCCTGTCTGCATCTCCAGGGGAGAAGGTCACAATGACTTGCAGGGCCAGCTCAAGTGTAAGTTACATGCACTGGTACCAGCAGAAGCCAGGATCCTCCCCCAAACCCTGGATTTATGCCACATCCAACCTGGCTTCTGGAGTCCCTGCTCGCTTCAGTGGCAGTGGGTCTGGGACCTCTTACTCTCTCACAATCAGCAGAGTGGAGGCTGAAGATGCTGCCACTTATTACTGCCAGCAGTGGAGTAGTAACCCACCCA |
| IGKV5-43*01_DBA1 | DBA/1J | Musmus IGKV5-43*01 F | 100 | 287 | GATATTGTGCTAACTCAGTCTCCAGCCACCCTGTCTGTGACTCCAGGAGATAGCGTCAGTCTTTCCTGCAGGGCCAGCCAAAGTATTAGCAACAACCTACACTGGTATCAACAAAAATCACATGAGTCTCCAAGGCTTCTCATCAAGTATGCTTCCCAGTCCATCTCTGGGATCCCCTCCAGGTTCAGTGGCAGTGGATCAGGGACAGATTTCACTCTCAGTATCAACAGTGTGGAGACTGAAGATTTTGGAATGTATTTCTGTCAACAGAGTAACAGCTGGCCTCA |
| IGKV6-13*01_DBA1 | DBA/1J | Musmus IGKV6-13*01 F | 100 | 285 | GACATTGTGATGACCCAGTCTCAAAAATTCATGTCCACATCAGTAGGAGACAGGGTCAGCATCACCTGCAAGGCCAGTCAGAATGTGGGTACTGCTGTAGCCTGGTATCAACAGAAACCAGGACAATCTCCTAAACTACTGATTTACTCGGCATCCAATCGGTACACTGGAGTCCCTGATCGCTTCACAGGCAGTGGATCTGGGACAGATTTCACTCTCACCATCAGCAATATGCAGTCTGAAGACCTGGCAGATTATTTCTGCCAGCAATATAGCAGCTATCCT |
| IGKV9-120*01_S2628_DBA1 | DBA/1J | Musmus IGKV9-120*02 F | 100 | 285 | GACATCCAGATGACCCAGTCTCCATCCTCCTTATCTGCCTCTCTGGGAGAAAGAGTCAGTCTCACTTGTCGGGCAAGTCAGGACATTGGTAGTAGCTTAAACTGGCTTCAGCAGGAACCAGATGGAACTATTAAACGCCTGATCTACGCCACATCCAGTTTAGATTCTGGTGTCCCCAAAAGGTTCAGTGGCAGTAGGTCTGGGTCAGATTATTCTCTCACCATCAGCAGCCTTGAGTCTGAAGATTTTGTAGACTATTACTGTCTACAATATGCTAGTTCTCCT |

| | | | | | |
|---|---|---|---|---|---|
| IGKV3-4*01_DBA1 | DBA/1J | Musmus IGKV3-4*01 F | 100 | 299 | GACATTGTGCTGACCCAATCTCCAGCTTCTTTGGCTGTGTCTCTAGGGCAGAGGGCCACCATCTCCTGCAAGGCCAGCCAAAGTGTTGATTATGATGGTGATAGTTATATGAACTGGTACCAACAGAAACCAGGACAGCCACCCAAACTCCTCATCTATGCTGCATCCAATCTAGAATCTGGGATCCCAGCCAGGTTTAGTGGCAGTGGGTCTGGGACAGACTTCACCCTCAACATCCATCCTGTGGAGGAGGAGGATGCTGCAACCTATTACTGTCAGCAAAGTAATGAGGATCCTCC |
| IGKV4-61*01_DBA1 | DBA/1J | Musmus IGKV4-61*01 F | 100 | 286 | CAAATTGTTCTCACCCAGTCTCCAGCAATCATGTCTGCATCTCCAGGGGAGAAGGTCACCATATCCTGCAGTGCCAGCTCAAGTGTAAGTTACATGTACTGGTACCAGCAGAAGCCAGGATCCTCCCCCAAACCCTGGATTTATCGCACATCCAACCTGGCTTCTGGAGTCCCTGCTCGCTTCAGTGGCAGTGGGTCTGGGACCTCTTACTCTCTCACAATCAGCAGCATGGAGGCTGAAGATGCTGCCACTTATTACTGCCAGCAGTATCATAGTTACCCACCCA |
| IGKV1-88*01_DBA1 | DBA/1J | Musmus IGKV1-88*01 F | 100 | 302 | GATGTTGTGGTGACTCAAACTCCACTCTCCCTGCCTGTCAGCTTTGGAGATCAAGTTTCTATCTCTTGCAGGTCTAGTCAGAGTCTTGCAAACAGTTATGGGAACACCTATTTGTCTTGGTACCTGCACAAGCCTGGCCAGTCTCCACAGCTCCTCATCTATGGGATTTCCAACAGATTTTCTGGGGTGCCAGACAGGTTCAGTGGCAGTGGTTCAGGGACAGATTTCACACTCAAGATCAGCACAATAAAGCCTGAGGACTTGGGAATGTATTACTGCTTACAAGGTACACATCAGCCTCC |
| IGKV6-32*01_DBA1 | DBA/1J | Musmus IGKV6-32*01 F | 100 | 287 | AGTATTGTGATGACCCAGACTCCCAAATTCCTGCTTGTATCAGCAGGAGACAGGGTTACCATAACCTGCAAGGCCAGTCAGAGTGTGAGTAATGATGTAGCTTGGTACCAACAGAAGCCAGGGCAGTCTCCTAAACTGCTGATATACTATGCATCCAATCGCTACACTGGAGTCCCTGATCGCTTCACTGGCAGTGGATATGGGACGGATTTCACTTTCACCATCAGCACTGTGCAGGCTGAAGACCTGGCAGTTTATTTCTGTCAGCAGGATTATAGCTCTCCTCC |
| IGKV4-70*01_DBA1 | DBA/1J | Musmus IGKV4-70*01 F | 100 | 282 | CAAATTGTTCTCACCCAGTCTCCAGCAATCATGTCTGCATCTCCAGGGGAGAAGGTCACCATGACCTGCAGTGCCAGCTCAAGTATAAGTTACATGCACTGGTACCAGCAGAAGCCAGGCACCTCCCCCAAAAGATGGATTTATGACACATCCAAACTGGCTTCTGGAGTCCCTGCTCGCTTCAGTGGCAGTGGGTCTGGGACCTCTTATTCTCTCACAATCAGCAGCATGGAGGCTGAAGATGCTGCCACTTATTACTGCCATCAGCGGAGTAGTTACCCA |
| IGKV4-57*01_S9219_DBA1 | DBA/1J | Musmus IGKV4-57-1*01 F | 100 | 286 | CAAATTGTTCTCACCCAGTCTCCAGCAATCATGTCTGCATCTCCAGGGGAGAAGGTCACCATAACCTGCAGTGCCAGCTCAAGTGTAAGTTACATGCACTGGTTCCAGCAGAAGCCAGGCACTTCTCCCAAACTCTGGATTTATAGCACATCCAACCTGGCTTCTGGAGTCCCTGCTCGCTTCAGTGGCAGTGGATCTGGGACCTCTTACTCTCTCACAATCAGCCGAATGGAGGCTGAAGATGCTGCCACTTATTACTGCCAGCAAAGGAGTAGTTACCCACTCA |
| IGKV10-94*01_DBA1 | DBA/1J | Musmus IGKV10-94*01 F | 100 | 287 | GATATCCAGATGACACAGACTACATCCTCCCTGTCTGCCTCTCTGGGAGACAGAGTCACCATCAGTTGCAGTGCAAGTCAGGGCATTAGCAATTATTTAAACTGGTATCAGCAGAAACCAGATGGAACTGTTAAACTCCTGATCTATTACACATCAAGTTTACACTCAGGAGTCCCATCAAGGTTCAGTGGCAGTGGGTCTGGGACAGATTATTCTCTCACCATCAGCAACCTGGAACCTGAAGATATTGCCACTTACTATTGTCAGCAGTATAGTAAGCTTCCTCC |
| IGKV15-103*01_DBA1 | DBA/1J | Musmus IGKV15-103*01 ORF | 100 | 287 | GACATCCAGATGAACCAGTCTCCATCCAGTCTGTCTGCATCCCTTGGAGACACAATTACCATCACTTGCCATGCCAGTCAGAACATTAATGTTTGGTTAAGCTGGTACCAGCAGAAACCAGGAAATATTCCTAAACTATTGATCTATAAGGCTTCCAACTTGCACACAGGCGTCCCATCAAGGTTTAGTGGCAGTGGATCTGGAACAGGTTTCACATTAACCATCAGCAGCCTGCAGCCTGAAGACATTGCCACTTACTACTGTCAACAGGGTCAAAGTTATCCTCT |

| | | | | | |
|---|---|---|---|---|---|
| IGKV12-89*01_DBA1 | DBA/1J | Musmus IGKV12-89*01 F | 100 | 287 | GACATCCAGATGACTCAGTCTCCAGCTTCACTGTCTGCATCTGTGGGAGAAACTGTCACCATCACATGTGGAGCAAGTGAGAATATTTACGGTGCTTTAAATTGGTATCAGCGGAAACAGGGAAAATCTCCTCAGCTCCTGATCTATGGTGCAACCAACTTGGCAGATGGCATGTCATCGAGGTTCAGTGGCAGTGGATCTGGTAGACAGTATTCTCTCAAGATCAGTAGCCTGCATCCTGACGATGTTGCAACGTATTACTGTCAAAATGTGTTAAGTACTCCTCC |
| IGKV4-91*01_DBA1 | DBA/1J | Musmus IGKV4-91*01 F | 100 | 292 | GAAATTGTGCTCACCCAGTCTCCAACCACCATGGCTGCATCTCCCGGGGAGAAGATCACTATCACCTGCAGTGCCAGCTCAAGTATAAGTTCCAATTACTTGCATTGGTATCAGCAGAAGCCAGGATTCTCCCCTAAACTCTTGATTTATAGGACATCCAATCTGGCTTCTGGAGTCCCAGCTCGCTTCAGTGGCAGTGGGTCTGGGACCTCTTACTCTCTCACAATTGGCACCATGGAGGCTGAAGATGTTGCCACTTACTACTGCCAGCAGGGTAGTAGTATACCACGCA |
| IGKV4-57-1*01_DBA1 | DBA/1J | Musmus IGKV4-57*01 F | 100 | 292 | GAAAATGTGCTCACCCAGTCTCCAGCAATCATGTCTGCATCTCCAGGGGAAAAGGTCACCATGACCTGCAGGGCCAGCTCAAGTGTAAGTTCCAGTTACTTGCACTGGTACCAGCAGAAGTCAGGTGCCTCCCCCAAACTCTGGATTTATAGCACATCCAACTTGGCTTCTGGAGTCCCTGCTCGCTTCAGTGGCAGTGGGTCTGGGACCTCTTACTCTCTCACAATCAGCAGTGTGGAGGCTGAAGATGCTGCCACTTATTACTGCCAGCAGTACAGTGGTTACCCACTCA |
| IGKV4-55*01_DBA1 | DBA/1J | Musmus IGKV4-55*01 F | 100 | 286 | CAAATTGTTCTCACCCAGTCTCCAGCAATCATGTCTGCATCTCCAGGGGAGAAGGTCACCATGACCTGCAGTGCCAGCTCAAGTGTAAGTTACATGTACTGGTACCAGCAGAAGCCAGGATCCTCCCCCAGACTCCTGATTTATGACACATCCAACCTGGCTTCTGGAGTCCCTGTTCGCTTCAGTGGCAGTGGGTCTGGGACCTCTTACTCTCTCACAATCAGCCGAATGGAGGCTGAAGATGCTGCCACTTATTACTGCCAGCAGTGGAGTAGTTACCCACCCA |
| IGKV4-80*01_DBA1 | DBA/1J | Musmus IGKV4-80*01 F | 100 | 282 | CAAATTGTTCTCACCCAGTCTCCAGCAATCATGTCTGCATCTCTAGGGGAGGAGATCACCCTAACCTGCAGTGCCAGCTCGAGTGTAAGTTACATGCACTGGTACCAGCAGAAGTCAGGCACTTCTCCCAAACTCTTGATTTATAGCACATCCAACCTGGCTTCTGGAGTCCCTTCTCGCTTCAGTGGCAGTGGGTCTGGGACCTTTTATTCTCTCACAATCAGCAGTGTGGAGGCTGAAGATGCTGCCGATTATTACTGCCATCAGTGGAGTAGTTATCCA |
| IGKV19-93*01_DBA1 | DBA/1J | Musmus IGKV19-93*01 F | 100 | 287 | GACATCCAGATGACACAGTCTCCATCCTCACTGTCTGCATCTCTGGGAGGCAAAGTCACCATCACTTGCAAGGCAAGCCAAGACATTAACAAGTATATAGCTTGGTACCAACACAAGCCTGGAAAAGGTCCTAGGCTGCTCATACATTACACATCTACATTACAGCCAGGCATCCCATCAAGGTTCAGTGGAAGTGGGTCTGGGAGAGATTATTCCTTCAGCATCAGCAACCTGGAGCCTGAAGATATTGCAACTTATTATTGTCTACAGTATGATAATCTTCTACC |
| IGKV6-17*01_DBA1 | DBA/1J | Musmus IGKV6-17*01 F | 100 | 287 | GACATTGTGATGACCCAGTCTCACAAATTCATGTCCACATCAGTAGGAGACAGGGTCAGCATCACCTGCAAGGCCAGTCAGGATGTGAGTACTGCTGTAGCCTGGTATCAACAGAAACCAGGACAATCTCCTAAACTACTGATTTACTCGGCATCCTACCGGTACACTGGAGTCCCTGATCGCTTCACTGGCAGTGGATCTGGGACGGATTTCACTTTCACCATCAGCAGTGTGCAGGCTGAAGACCTGGCAGTTTATTACTGTCAGCAACATTATAGTACTCCTCC |
| IGKV3-2*01_DBA1 | DBA/1J | Musmus IGKV3-2*01 F | 100 | 299 | GACATTGTGCTGACCCAATCTCCAGCTTCTTTGGCTGTGTCTCTAGGGCAGAGGGCCACCATCTCCTGCAGAGCCAGCGAAAGTGTTGATAATTATGGCATTAGTTTTATGAACTGGTTCCAACAGAAACCAGGACAGCCACCCAAACTCCTCATCTATGCTGCATCCAACCAAGGATCCGGGGTCCCTGCCAGGTTTAGTGGCAGTGGGTCTGGGACAGACTTCAGCCTCAACATCCATCCTATGGAGGAGGATGATACTGCAATGTATTTCTGTCAGCAAAGTAAGGAGGTTCCTCC |

| | | | | | |
|---|---|---|---|---|---|
| IGKV3-5*01_DBA1 | DBA/1J | Musmus IGKV3-5*01 F | 100 | 299 | GACATTGTGCTGACCCAATCTCCAGCTTCTTTGGCTGTGTCTCTAGGGCAGAGGGCCACCATATCCTGCAGAGCCAGTGAAAGTGTTGATAGTTATGGCAATAGTTTTATGCACTGGTACCAGCAGAAACCAGGACAGCCACCCAAACTCCTCATCTATCGTGCATCCAACCTAGAATCTGGGATCCCTGCCAGGTTCAGTGGCAGTGGGTCTAGGACAGACTTCACCCTCACCATTAATCCTGTGGAGGCTGATGATGTTGCAACCTATTACTGTCAGCAAAGTAATGAGGATCCTCC |
| IGKV3-7*01_S5296_DBA1 | DBA/1J | Musmus IGKV3-7*02 F | 100 | 299 | GACATTGTGCTGACACAGTCTCCTGCTTCCTTAGCTGTATCTCTGGGGCAGAGGGCCACCATCTCATGCAGGGCCAGCCAAAGTGTCAGTACATCTAGCTATAGTTATATGCACTGGTACCAACAGAAACCAGGACAGCCACCCAAACTCCTCATCAAGTATGCATCCAACCTAGAATCTGGGGTCCCTGCCAGGTTCAGTGGCAGTGGGTCTGGGACAGACTTCACCCTCAACATCCATCCTGTGGAGGAGGAGGATACTGCAACATATTACTGTCAGCACAGTTGGGAGATTCCTCC |
| IGKV8-16*01_DBA1 | DBA/1J | Musmus IGKV8-16*01 F | 100 | 305 | GAAATTGTGTTGACCCAGTCTATACCATCCCTGACTGTGTCAGCAGGAGAGAGGGTCACTATCAGCTGCAAATCCAATCAGAATCTTTTATGGAGTGGAAACCAAAGGTACTGTTTGGTCTGGCACCAGTGGAAACCAGGGCAAACTCCTACACCGTTGATCACCTGGACATCTGATAGGTACTCTGGAGTCCCTGATCGTTTCATAGGCAGTGGATCTGTGACAGATTTCACTCTGACCATCAGCAGTGTGCAGGCTGAAGATGTGGCAGTTTATTTCTGTCAGCAGCATTTACACATTCCTCC |
| IGKV3-12*01_DBA1 | DBA/1J | Musmus IGKV3-12*01 F | 100 | 299 | GACATTGTGCTGACACAGTCTCCTGCTTCCTTAGCTGTATCTCTGGGGCAGAGGGCCACCATCTCATGCAGGGCCAGCAAAAGTGTCAGTACATCTGGCTATAGTTATATGCACTGGTACCAACAGAAACCAGGACAGCCACCCAAACTCCTCATCTATCTTGCATCCAACCTAGAATCTGGGGTCCCTGCCAGGTTCAGTGGCAGTGGGTCTGGGACAGACTTCACCCTCAACATCCATCCTGTGGAGGAGGAGGATGCTGCAACCTATTACTGTCAGCACAGTAGGGAGCTTCCTCC |
| IGKV5-48*01_DBA1 | DBA/1J | Musmus IGKV5-48*01 F | 100 | 287 | GACATCTTGCTGACTCAGTCTCCAGCCATCCTGTCTGTGAGTCCAGGAGAAAGAGTCAGTTTCTCCTGCAGGGCCAGTCAGAGCATTGGCACAAGCATACACTGGTATCAGCAAAGAACAAATGGTTCTCCAAGGCTTCTCATAAAGTATGCTTCTGAGTCTATCTCTGGGATCCCTTCCAGGTTTAGTGGCAGTGGATCAGGGACAGATTTTACTCTTAGCATCAACAGTGTGGAGTCTGAAGATATTGCAGATTATTACTGTCAACAAAGTAATAGCTGGCCAAC |
| IGKV6-25*01_DBA1 | DBA/1J | Musmus IGKV6-25*01 F | 100 | 287 | GACATTGTGATGACCCAGTCTCACAAATTCATGTCCACATCAGTAGGAGACAGGGTCAGCATCACCTGCAAGGCCAGTCAGGATGTGAGTACTGCTGTAGCCTGGTATCAACAAAAACCAGGGCAATCTCCTAAACTACTGATTTACTGGGCATCCACCCGGCACACTGGAGTCCCTGATCGCTTCACAGGCAGTGGATCTGGGACAGATTATACTCTCACCATCAGCAGTGTGCAGGCTGAAGACCTGGCACTTTATTACTGTCAGCAACATTATAGCACTCCTCC |
| IGKV4-53*01_DBA1 | DBA/1J | Musmus IGKV4-53*01 F | 100 | 292 | GAAATTGTGCTCACCCAGTCTCCAGCACTCATGGCTGCATCTCCAGGGGAGAAGGTCACCATCACCTGCAGTGTCAGCTCAAGTATAAGTTCCAGCAACTTGCACTGGTACCAGCAGAAGTCAGAAACCTCCCCCAAACCCTGGATTTATGGCACATCCAACCTGGCTTCTGGAGTCCCTGTTCGCTTCAGTGGCAGTGGATCTGGGACCTCTTATTCTCTCACAATCAGCAGCATGGAGGCTGAAGATGCTGCCACTTATTACTGTCAACAGTGGAGTAGTTACCCACTCA |

| | | | | | |
|---|---|---|---|---|---|
| IGKV14-126*01_DBA1 | DBA/1J | Musmus IGKV14-126*01 F | 100 | 287 | GACATCAAGATGACCCAGTCTCCATCCTCCATGTATGCATCGCTGGGAGAGAGAGTCACTATCACTTGCAAGGCGAGTCAGGACATTAAAAGCTATTTAAGCTGGTACCAGCAGAAACCATGGAAATCTCCTAAGACCCTGATCTATTATGCAACAAGCTTGGCAGATGGGGTCCCATCAAGATTCAGTGGCAGTGGATCTGGGCAAGATTATTCTCTAACCATCAGCAGCCTGGAGTCTGACGATACAGCAACTTATTACTGTCTACAGCATGGTGAGAGCCCTCC |
| IGKV13-85*01_DBA1 | DBA/1J | Musmus IGKV13-85*01 F | 100 | 287 | GACATCCAGATGACACAATCTTCATCCTACTTGTCTGTATCTCTAGGAGGCAGAGTCACCATTACTTGCAAGGCAAGTGACCACATTAATAATTGGTTAGCCTGGTATCAGCAGAAACCAGGAAATGCTCCTAGGCTCTTAATATCTGGTGCAACCAGTTTGGAAACTGGGGTTCCTTCAAGATTCAGTGGCAGTGGATCTGGAAAGGATTACACTCTCAGCATTACCAGTCTTCAGACTGAAGATGTTGCTACTTATTACTGTCAACAGTATTGGAGTACTCCTCC |
| IGKV1-122*01_DBA1 | DBA/1J | Musmus IGKV1-122*01 F | 100 | 302 | GATGCTGTGATGACCCAAACTCCACTCTCCCTGCCTGTCAGTCTTGGAGATCAAGCCTCCATCTCTTGCAGGTCTAGTCAGAGCCTTGAAAACAGTAATGGAAACACCTATTTGAACTGGTACCTCCAGAAACCAGGCCAGTCTCCACAGCTCCTGATCTACAGGGTTTCCAACCGATTTTCTGGGGTCCTAGACAGGTTCAGTGGTAGTGGATCAGGGACAGATTTCACACTGAAAATCAGCAGAGTGGAGGCTGAGGATTTGGGAGTTTATTTCTGCCTCCAAGTTACACATGTCCCTCC |
| IGKV13-84*01_DBA1 | DBA/1J | Musmus IGKV13-84*01 F | 100 | 287 | GACATCCAGATGACACAATCTTCATCCTCCTTTTCTGTATCTCTAGGAGACAGAGTCACCATTACTTGCAAGGCAAGTGAGGACATATATAATCGGTTAGCCTGGTATCAGCAGAAACCAGGAAATGCTCCTAGGCTCTTAATATCTGGTGCAACCAGTTTGGAAACTGGGGTTCCTTCAAGATTCAGTGGCAGTGGATCTGGAAAGGATTACACTCTCAGCATTACCAGTCTTCAGACTGAAGATGTTGCTACTTATTACTGTCAACAGTATTGGAGTACTCCTCC |
| IGKV4-68*01_DBA1 | DBA/1J | Musmus IGKV4-68*01 F | 100 | 286 | CAAATTGTTCTCACCCAGTCTCCAGCACTCATGTCTGCATCTCCAGGGGAGAAGGTCACCATGACCTGCAGTGCCAGCTCAAGTGTAAGTTACATGTACTGGTACCAGCAGAAGCCAAGATCCTCCCCCAAACCCTGGATTTATCTCACATCCAACCTGGCTTCTGGAGTCCCTGCTCGCTTCAGTGGCAGTGGGTCTGGGACCTCTTACTCTCTCACAATCAGCAGCATGGAGGCTGAAGATGCTGCCACTTATTACTGCCAGCAGTGGAGTAGTAACCCACCCA |
| IGKV4-86*01_DBA1 | DBA/1J | Musmus IGKV4-86*01 F | 100 | 283 | GAAATTGTGCTCACTCAGTCTCCAGCCATCACAGCTGCATCTCTGGGGCAAAAGGTCACCATCACCTGCAGTGCCAGCTCAAGTGTAAGTTACATGCACTGGTACCAGCAGAAGTCAGGCACCTCCCCCAAACCATGGATTTATGAAATATCCAAACTGGCTTCTGGAGTCCCAGCTCGCTTCAGTGGCAGTGGGTCTGGGACCTCTTACTCTCTCACAATCAGCAGCATGGAGGCTGAAGATGCTGCCATTTATTACTGCCAGCAGTGGAATTATCCTCTTA |
| IGKV17-121*01_DBA1 | DBA/1J | Musmus IGKV17-121*01 F | 100 | 287 | GAAACAACTGTGACCCAGTCTCCAGCATCCCTGTCCATGGCTATAGGAGAAAAAGTCACCATCAGATGCATAACCAGCACTGATATTGATGATGATATGAACTGGTACCAGCAGAAGCCAGGGGAACCTCCTAAGCTCCTTATTTCAGAAGGCAATACTCTTCGTCCTGGAGTCCCATCCCGATTCTCCAGCAGTGGCTATGGTACAGATTTTGTTTTTACAATTGAAAACATGCTCTCAGAAGATGTTGCAGATTACTACTGTTTGCAAAGTGATAACTTGCCTCT |
| IGKV17-127*01_DBA1 | DBA/1J | Musmus IGKV17-127*01 F | 100 | 287 | GAAACAACTGTGACCCAGTCTCCAGCATCCCTGTCCGTGGCTACAGGAGAAAAAGTCACTATCAGATGCATAACCAGCACTGATATTGATGATGATATGAACTGGTACCAGCAGAAGCCAGGGGAACCTCCTAAGCTCCTTATTTCAGAAGGCAATACTCTTCGTCCTGGAGTCCCATCCCGATTCTCCAGCAGTGGCTATGGCACAGATTTTGTTTTTACAATTGAAAACACGCTCTCAGAAGATGTTGCAGATTACTACTGTTTGCAAAGTGATAACATGCCTCT |

| | | | | | |
|---|---|---|---|---|---|
| IGKV16-104*01_DBA1 | DBA/1J | Musmus IGKV16-104*01 F | 100 | 288 | GATGTCCAGATAACCCAGTCTCCATCTTATCTTGCTGCATCTCCTGGAGAAACCATTACTATTAATTGCAGGGCAAGTAAGAGCATTAGCAAATATTTAGCCTGGTATCAAGAGAAACCTGGGAAAACTAATAAGCTTCTTATCTACTCTGGATCCACTTTGCAATCTGGAATTCCATCAAGGTTCAGTGGCAGTGGATCTGGTACAGATTTCACTCTCACCATCAGTAGCCTGGAGCCTGAAGATTTTGCAATGTATTACTGTCAACAGCATAATGAATACCCGTAC |
| IGKV4-74*01_S2922_DBA1 | DBA/1J | Musmus IGKV4-74*01 F | 100 | 292 | CAAATTGTTCTCACCCAGTCTCCAGCAATCATGTCTGCATCTCTAGGGGAACGGGTCACCATGACCTGCACTGCCAGCTCAAGTGTAAGTTCCAGTTACTTGCACTGGTACCAGCAGAAGCCAGGATCCTCCCCCAAACTCTGGATTTATAGCACATCCAACCTGGCTTCTGGAGTCCCAGCTCGCTTCAGTGGCAGTGGGTCTGGGACCTCTTACTCTCTCACAATCAGCAGCATGGAGGCTGAAGATGCTGCCACTTATTACTGCCACCAGTATCATCGTTCCCCACTCA |
| IGKV5-45*01_DBA1 | DBA/1J | Musmus IGKV5-45*01 F | 100 | 287 | GATATTGTGCTAACTCAGTCTCCAGCCACCCTGTCTGTGACTCCAGGAGATAGAGTCAGTCTTTCCTGCAGGGCCAGTCAAAGTATTAGCAACTACCTACACTGGTATCAACAAAAATCACATGAGTCTCCAAGGCTTCTCATCAAGTATGCTTCCCAGTCCATCTCTGGGATCCCCTCCAGGTTCAGTGGCAGTGGATCAGGGACAGATTTCACTCTCAGTATCAACAGTGTGGAGACTGAAGATTTTGGAATGTATTTCTGTCAACAGAGTAACAGCTGGCCTCA |
| IGKV14-111*01_DBA1 | DBA/1J | Musmus IGKV14-111*01 F | 100 | 287 | GACATCAAGATGACCCAGTCTCCATCTTCCATGTATGCATCTCTAGGAGAGAGAGTCACTATCACTTGCAAGGCGAGTCAGGACATTAATAGCTATTTAAGCTGGTTCCAGCAGAAACCAGGGAAATCTCCTAAGACCCTGATCTATCGTGCAAACAGATTGGTAGATGGGGTCCCATCAAGGTTCAGTGGCAGTGGATCTGGGCAAGATTATTCTCTCACCATCAGCAGCCTGGAGTATGAAGATATGGGAATTTATTATTGTCTACAGTATGATGAGTTTCCTCC |
| IGKV8-34*01_DBA1 | DBA/1J | Musmus IGKV8-34*01 F | 100 | 305 | GACATTTTGATGACCCAGTCTCCATCCTCCCTGACTGTGTCAGCAGGAGAGAAGGTCACTATGAGCTGCAAGTCCAGTCAGAGTCTTTTAGCTAGTGGCAACCAAAATAACTACTTGGCCTGGCACCAGCAGAAACCAGGACGATCTCCTAAAATGCTGATAATTTGGGCATCCACTAGGGTATCTGGAGTCCCTGATCGCTTCATAGGCAGTGGATCTGGGACGGATTTCACTCTGACCATCAACAGTGTGCAGGCTGAAGATCTGGCTGTTTATTACTGTCAGCAGTCCTACAGCGCTCCTAC |
| IGKV2-112*01_DBA1 | DBA/1J | Musmus IGKV2-112*01 F | 100 | 300 | GATATTGTGATAACCCAGGATGAACTCTCCAATCCTGTCACTTCTGGAGAATCAGTTTCCATCTCCTGCAGGTCTAGTAAGAGTCTCCTATATAAGGATGGGAAGACATACTTGAATTGGTTTCTGCAGAGACCAGGACAATCTCCTCAGCTCCTGATCTATTTGATGTCCACCCGTGCATCAGGAGTCTCAGACCGGTTTAGTGGCAGTGGGTCAGGAACAGATTTCACCCTGGAAATCAGTAGAGTGAAGGCTGAGGATGTGGGTGTGTATTACTGTCAACAACTTGTAGAGTATCCT |
| IGKV3-10*01_DBA1 | DBA/1J | Musmus IGKV3-10*01 F | 100 | 299 | AACATTGTGCTGACCCAATCTCCAGCTTCTTTGGCTGTGTCTCTAGGGCAGAGGGCCACCATATCCTGCAGAGCCAGTGAAAGTGTTGATAGTTATGGCAATAGTTTTATGCACTGGTACCAGCAGAAACCAGGACAGCCACCCAAACTCCTCATCTATCTTGCATCCAACCTAGAATCTGGGGTCCCTGCCAGGTTCAGTGGCAGTGGGTCTAGGACAGACTTCACCCTCACCATTGATCCTGTGGAGGCTGATGATGCTGCAACCTATTACTGTCAGCAAAATAATGAGGATCCTCC |

| | | | | | |
|---|---|---|---|---|---|
| IGKV1-133*01_DBA1 | DBA/1J | Musmus IGKV1-133*01 F | 100 | 302 | GATGTTGTGATGACCCAGACTCCACTCACTTTGTCGGTTACCATTGGACAACCAGCCTCTATCTCTTGCAAGTCAAGTCAGAGCCTCTTATATAGTAATGGAAAAACCTATTTGAATTGGTTATTACAGAGGCCAGGCCAGTCTCCAAAGCGCCTAATCTATCTGGTGTCTAAACTGGACTCTGGAGTCCCTGACAGGTTCACTGGCAGTGGATCAGGAACAGATTTTACACTGAAAATCAGCAGAGTGGAGGCTGAGGATTTGGGAGTTTATTACTGCGTGCAAGGTACACATTTTCCTCA |
| IGKV2-109*01_DBA1 | DBA/1J | Musmus IGKV2-109*01 F | 100 | 302 | GATATTGTGATGACGCAGGCTGCATTCTCCAATCCAGTCACTCTTGGAACATCAGCTTCCATCTCCTGCAGGTCTAGTAAGAGTCTCCTACATAGTAATGGCATCACTTATTTGTATTGGTATCTGCAGAAGCCAGGCCAGTCTCCTCAGCTCCTGATTTATCAGATGTCCAACCTTGCCTCAGGAGTCCCAGACAGGTTCAGTAGCAGTGGGTCAGGAACTGATTTCACACTGAGAATCAGCAGAGTGGAGGCTGAGGATGTGGGTGTTTATTACTGTGCTCAAAATCTAGAACTTCCTCC |
| IGKV11-125*01_DBA1 | DBA/1J | Musmus IGKV11-125*01 F | 100 | 287 | GATGTCCAGATGATTCAGTCTCCATCCTCCCTGTCTGCATCTTTGGGAGACATAGTCACCATGACTTGCCAGGCAAGTCAGGGCACTAGCATTAATTTAAACTGGTTTCAGCAAAAACCAGGGAAAGCTCCTAAGCTCCTGATCTATGGTGCAAGCAACTTGGAAGATGGGGTCCCATCAAGGTTCAGTGGCAGTAGATATGGGACAGATTTCACTCTCACCATCAGCAGCCTGGAGGATGAAGATATGGCAACTTATTTCTGTCTACAGCATAGTTATCTCCCTCC |
| IGKV14-100*01_DBA1 | DBA/1J | Musmus IGKV14-100*01 F | 100 | 287 | GACATCCTGATGACCCAATCTCCATCCTCCATGTCTGTATCTCTGGGAGACACAGTCAGCATCACTTGCCATGCAAGTCAGGGCATTAGCAGTAATATAGGGTGGTTGCAGCAGAAACCAGGGAAATCATTTAAGGGCCTGATCTATCATGGAACCAACTTGGAAGATGGAGTTCCATCAAGGTTCAGTGGCAGTGGATCTGGAGCAGATTATTCTCTCACCATCAGCAGCCTGGAATCTGAAGATTTTGCAGACTATTACTGTGTACAGTATGCTCAGTTTCCTCC |
| IGKV4-63*01_DBA1 | DBA/1J | Musmus IGKV4-63*01 F | 100 | 286 | GAAAATGTTCTCACCCAGTCTCCAGCAATCATGTCTGCATCTCCAGGGGAAAAGGTCACCATGACCTGCAGTGCCAGCTCAAGTGTAAGTTACATGCACTGGTACCAGCAGAAGTCAAGCACCTCCCCCAAACTCTGGATTTATGACACATCCAAACTGGCTTCTGGAGTCCCAGGTCGCTTCAGTGGCAGTGGGTCTGGAAACTCTTACTCTCTCACGATCAGCAGCATGGAGGCTGAAGATGTTGCCACTTATTACTGTTTTCAGGGGAGTGGGTACCCACTCA |
| IGKV4-79*01_DBA1 | DBA/1J | Musmus IGKV4-79*01 F | 100 | 292 | CAAATTGTTCTCACCCAGTCTCCAGCAATCATGTCTGCATCTCCTGGGGAGAAGGTCACCTTGACCTGCAGTGCCAGCTCAAGTGTAAGTTCCAGCTACTTGTACTGGTACCAGCAGAAGCCAGGATCCTCCCCCAAACTCTGGATTTATAGCACATCCAACCTGGCTTCTGGAGTCCCTGCTCGCTTCAGTGGCAGTGGGTCTGGGACCTCTTACTCTCTCACAATCAGCAGCATGGAGGCTGAAGATGCTGCCTCTTATTTCTGCCATCAGTGGAGTAGTTACCCACCCC |
| IGKV4-51*01_DBA1 | DBA/1J | Musmus IGKV4-51*01 F | 100 | 292 | GAAAATGTGCTCACCCAGTCTCCAGCAATAATGGCTGCCTCTCTGGGGGAGAAGGTCACCATGACCTGCAGTGCCAGCTCAAGTGTAAGTTCCAGCTACTTGCACTGGTACCAGCAGAAGTCAGGCACTTCCCCCAAACTCTGGATTTATGGCACATCCAACCTGGCTTCTGGAGTCCCAGCTCGCTTCAGTGGCAGTGGGGCTGGGATCTCTTACTCTCTCACAATCAGCAGCATGGAGGCTGAAAATGATGCAACTTATTACTGCCAGCAGTGGAGTGGTTACCCATTCA |

| | | | | | |
|---|---|---|---|---|---|
| IGKV4-91*01_S5997_DBA1 | DBA/1J | Musmus IGKV4-91*01 F | 100 | 292 | GAAATTGTGCTCACCCAGTCTCCAACCACCATGGCTGCATCTCCCGGGGAGAAGATCACTATCACCTGCAGTGCCAGCTCAAGTATAAGTTCCAATTACTTGCATTGGTATCAGCAGAAGCCAGGATTCTCCCCTAAACTCTTGATTTATAGGACATCCAATCTGGCTTCTGGAGTCCCAGCTCGCTTCAGTGGCAGTGGGTCTGGGACCTCTTACTCTCTCACAATTGGCACCATGGAGGCTGAAGATGTTGCCACTTACTACTGCCAGCAGGGTAGTAGTATACCACTCA |
| IGKV5-39*01_DBA1 | DBA/1J | Musmus IGKV5-39*01 F | 100 | 287 | GACATTGTGATGACTCAGTCTCCAGCCACCCTGTCTGTGACTCCAGGAGATAGAGTCTCTCTTTCCTGCAGGGCCAGCCAGAGTATTAGCGACTACTTACACTGGTATCAACAAAAATCACATGAGTCTCCAAGGCTTCTCATCAAATATGCTTCCCAATCCATCTCTGGGATCCCCTCCAGGTTCAGTGGCAGTGGATCAGGGTCAGATTTCACTCTCAGTATCAACAGTGTGGAACCTGAAGATGTTGGAGTGTATTACTGTCAAAATGGTCACAGCTTTCCTCC |
| IGKV9-123*01_DBA1 | DBA/1J | Musmus IGKV9-123*01 F | 100 | 285 | GACATCCAGATGATTCAGTCTCCATCGTCCATGTTTGCCTCTCTGGGAGACAGAGTCAGTCTCTCTTGTCGGGCTAGTCAGGGCATTAGAGGTAATTTAGACTGGTATCAGCAGAAACCAGGTGGAACTATTAAACTCCTGATCTACTCCACATCCAATTTAAATTCTGGTGTCCCATCAAGGTTCAGTGGCAGTGGGTCTGGGTCAGATTATTCTCTCACCATCAGCAGCCTAGAGTCTGAAGATTTTGCAGACTATTACTGTCTACAGCGTAATGCGTATCCT |
| IGKV9-124*01_DBA1 | DBA/1J | Musmus IGKV9-124*01 F | 100 | 285 | GACATCCAGATGACCCAGTCTCCATCCTCCTTATCTGCCTCTCTGGGAGAAAGAGTCAGTCTCACTTGTCGGGCAAGTCAGGAAATTAGTGGTTACTTAAGCTGGCTTCAGCAGAAACCAGATGGAACTATTAAACGCCTGATCTACGCCGCATCCACTTTAGATTCTGGTGTCCCAAAAAGGTTCAGTGGCAGTAGGTCTGGGTCAGATTATTCTCTCACCATCAGCAGCCTTGAGTCTGAAGATTTTGCAGACTATTACTGTCTACAATATGCTAGTTATCCT |
| IGKV10-95*01_DBA1 | DBA/1J | Musmus IGKV10-95*01 F | 100 | 287 | GATATCCAGATGACACAGACTACTTCCTCCCTGTCTGCCTCTCTGGGAGACAGAGTCACCATCAGTTGCAGGGCAAGTGAGGACATTAGCACTTATTTAAACTGGTATCAGCAGAAACCAGATGGAACTGTTAAACTCCTGATCTATTACACATCAGGATTACACTCAGGAGTCCCATCAAGGTTCAGTGGCAGTGGGTCTGGGGCAGATTATTCTCTCACCATCAGCAACCTGGAACCTGAAGATATTGCCACTTACTATTGTCAGCAGTATAGTAAGCTTCCTCC |
| IGKV4-50*01_S0901_DBA1 | DBA/1J | Musmus IGKV4-50*01 F | 100 | 286 | GAAAATGTGCTCACCCAGTCTCCAGCAATCATGTCTGCATCTCTAGGGGAGAAGGTCACCATGAGCTGCAGGGCCAGCTCAAGTGTAAATTACATGTACTGGTACCAGCAGAAGTCAGATGCCTCCCCCAAACTATGGATTTATTACACATCCAACCTGGCTCCTGGAGTCCCAGCTCGCTTCAGTGGCAGTGGGTCTGGGAACTCTTATTCTCTCACAATCAGCAGCATGGAGGGTGAAGATGCTGCCACTTATTACTGCCAGCAGTTTACTAGTTCCCCATACA |
| IGKV12-98*01_DBA1 | DBA/1J | Musmus IGKV12-98*01 F | 100 | 287 | GACATTCAGATGACCCAGTCTCCTGCCTCCCAGTCTGCATCTCTGGGAGAAAGTGTCACCATCACATGCCTGGCAAGTCAGACCATTGGTACATGGTTAGCATGGTATCAGCAGAAACCAGGGAAATCTCCTCAGCTCCTGATTTATGCTGCAACCAGCTTGGCAGATGGGGTCCCATCAAGGTTCAGTGGTAGTGGATCTGGCACAAAATTTTCTTTCAAGATCAGCAGCCTACAGGCTGAAGATTTTGTAAGTTATTACTGTCAACAACTTTACAGTACTCCTCT |
| IGKV4-92*01_DBA1 | DBA/1J | Musmus IGKV4-92*01 F | 100 | 291 | GAAATGGTTCTCACCCAGTCTCCAGTATCCATAACTGCATCTCGAGGGGAGAAGGTCACCATCACCTGCCGTGCCAGCTCAAGTATAAGTTCCAATTACTTACACTGGTACCAGCAGAAGCCAGGATCCTCCCCTAAACTTTTGATTTATAGGACATCCATCCTGGCATCTGGAGTCCTAGACAGCTTCAGTGGCAGTGGGTCTGAGAGCTCTTACACTCTGACAATCAGCTGCATGCAGGACGAAGTTGCTGCCACTTACTATTGTCAGCAGGGGAGTAGTAGCCCACCA |

| | | | | | |
|---|---|---|---|---|---|
| IGKV8-23-1*01_DBA1 | DBA/1J | Musmus IGKV8-16*01 F, or Musmus IGKV8-23-1*01 ORF | 100 | 284 | GAAATTGTGCTGACTGTGTCAGCAGGAGAGAGGGTCACTATCAGCTGCAAATCCAATCAGAATCTTTTATGGAGTGGAAACCAAAGGTACTGTTTGGTCTGGCACCAGTGGAAACCAGGGCAAACTCCTACACCGTTGATCACCTGGACATCTGATAGGTACTCTGGAGTCCCTGATCGTTTCATAGGCAGTGGATCTGTGACAGATTTCACTCTGACCATCAGCAGTGTGCAGGCTGAAGATGTGGCAGTTTATTTCTGTCAGCAGCATTTACACATTCCTCC |
| IGKV4-69*01_DBA1 | DBA/1J | Musmus IGKV4-69*01 F | 100 | 282 | CAAATTCTTCTCACCCAGTCTCCAGCAATCATGTCTGCATCTCCAGGGGAGAAGGTCACCATGACCTGCAGTGCCAGCTCAAGTGTAAGTTACATGCACTGGTACCAGCAGAAGCCAGGATCCTCGCCCAAACCCTGGATTTATGACACATCCAACCTGGCTTCTGGATTCCCTGCTCGCTTCAGTGGCAGTGGGTCTGGGACCTCTTACTCTCTCATAATCAGCAGCATGGAGGCTGAAGATGCTGCCACTTATTACTGCCATCAGCGGAGTAGTTACCCA |
| IGKV8-18*01_DBA1 | DBA/1J | Musmus IGKV8-18*01 ORF | 100 | 310 | GACATTGTGATGACCCAGTCTCCATCCTCCCTGGCTGTGACAGCAGGAGAGAAGGTCACTATGAGATGCAAGTCCAGTCAGAGTCTTTTGTGGAGTGTAAACCAAAATAACTACTTATCCTGGTACCAGCAGAAACAAGGGCAGCCTCCTAAACTGCTTATCTATGGGGCATCCATTAGAGAATCTTGGGTCCCTGATCGATTCACAGGAAGTGGATCTGGGACAGACTTCACTCTCACCATTAGCAATGTGCATGCTGAAGACCTAGCAGTTTATTACTGTCAGCACAATCATGGCAGCTTTCTCCCCC |
| IGKV1-99*01_DBA1 | DBA/1J | Musmus IGKV1-99*01 F | 100 | 302 | GATGTTGTTCTGACCCAAACTCCACTCTCTCTGCCTGTCAATATTGGAGATCAAGCCTCTATCTCTTGCAAGTCTACTAAGAGTCTTCTGAATAGTGATGGATTCACTTATTTGGACTGGTACCTGCAGAAGCCAGGCCAGTCTCCACAGCTCCTAATATATTTGGTTTCTAATCGATTTTCTGGAGTTCCAGACAGGTTCAGTGGCAGTGGGTCAGGAACAGATTTCACACTCAAGATCAGCAGAGTGGAGGCTGAGGATTTGGGAGTTTATTATTGCTTCCAGAGTAACTATCTTCCTCT |
| IGKV14-130*01_DBA1 | DBA/1J | Musmus IGKV14-130*01 F | 100 | 287 | GAAATCCAGATGACCCAGTCTCCATCCTCTATGTCTGCATCTCTGGGAGACAGAATAACCATCACTTGCCAGGCAACTCAAGACATTGTTAAGAATTTAAACTGGTATCAGCAGAAACCAGGGAAACCCCCTTCATTCCTGATCTATTATGCAACTGAACTGGCAGAAGGGGTCCCATCAAGGTTCAGTGGCAGTGGGTCTGGGTCAGACTATTCTCTGACAATCAGCAACCTGGAGTCTGAAGATTTTGCAGACTATTACTGTCTACAGTTTTATGAGTTTCCTCC |
| IGKV3-1*01_DBA1 | DBA/1J | Musmus IGKV3-1*01 F | 100 | 299 | GACATTGTGCTCACCCAATCTCCAGCTTCTTTGGCTGTGTCTCTAGGGCAGAGAGCCACCATCTCCTGCAGAGCCAGTGAAAGTGTTGAATATTATGGCACAAGTTTAATGCAGTGGTACCAACAGAAACCAGGACAGCCACCCAAACTCCTCATCTATGCTGCATCCAACGTAGAATCTGGGGTCCCTGCCAGGTTTAGTGGCAGTGGGTCTGGGACAGACTTCAGCCTCAACATCCATCCTGTGGAGGAGGATGATATTGCAATGTATTTCTGTCAGCAAAGTAGGAAGGTTCCTTC |
| IGKV4-55*01_S6279_DBA1 | DBA/1J | Musmus IGKV4-55*01 F | 100 | 286 | CAAATTGTTCTCACCCAGTCTCCAGCAATCATGTCTGCATCTCCAGGGGAGAAGGTCACCATGACCTGCAGTGCCAGCTCAAGTGTAAGTTACATGTACTGGTACCAGCAGAAGCCAGGATCCTCCCCCAGACTCCTGATTTATGACACATCCAACCTGGCTTCTGGAGTCCCTGTTCGCTTCAGTGGCAGTGGGTCTGGGACCTCTTACTCTCTCACAATCAGCCGAATGGAGGCTGAAGATGCTGCCACTTATTACTGCCAGCAGTGGAGTAGTTACCCACTCA |
| IGKV4-73*01_S8836_DBA1 | DBA/1J | Musmus IGKV4-73*01 ORF | 100 | 292 | CAAATTGTTCTCACCCAGTCTCCAGCAATCATGTCTGCATCTCCTGGGGAACGGGTCACCATGACCTGCAGTGCCAGCTCAAGTGTAAGTTCCAGCTACTTGTACTGGTACCAGCAGAAGTCAGGATCCTCCCCAAAACTCTGGATTTATAGCATATCCAACCTGGCTTCTGGAGTCCCAGCTCGCTTCAGTGGCAGTGGGTCTGGGACCTCTTACTCTCTCACAATCAACAGCATGGAGGCTGAAGATGCTGCCACTTATTACTGCCAGCAGTGGAGTAGTAACCCACTCA |

| | | | | | |
|---|---|---|---|---|---|
| IGKV4-50*01_S2743_DBA1 | DBA/1J | Musmus IGKV4-50*01 F | 100 | 286 | GAAAATGTGCTCACCCAGTCTCCAGCAATCATGTCTGCATCTCTAGGGGAGAAGGTCACCATGAGCTGCAGGGCCAGCTCAAGTGTAAATTACATGTACTGGTACCAGCAGAAGTCAGATGCCTCCCCCAAACTATGGATTTATTACACATCCAACCTGGCTCCTGGAGTCCCAGCTCGCTTCAGTGGCAGTGGGTCTGGGAACTCTTATTCTCTCACAATCAGCAGCATGGAGGGTGAAGATGCTGCCACTTATTACTGCCAGCAGTTTACTAGTTCCCCATTCA |
| IGKV4-58*01_DBA1 | DBA/1J | Musmus IGKV4-58*01 F | 100 | 292 | GAAAATGTGCTCACCCAGTCTCCAGCAATAATGGCTGCCTCTCTGGGGCAGAAGGTCACCATGACCTGCAGTGCCAGCTCAAGTGTAAGTTCCAGTTACTTGCACTGGTACCAGCAGAAGTCAGGCGCTTCCCCCAAACCCTTGATTCATAGGACATCCAACCTGGCTTCTGGAGTCCCAGCTCGCTTCAGTGGCAGTGGGTCTGGGACCTCTTACTCTCTCACAATCAGCAGCGTGGAGGCTGAAGATGATGCAACTTATTACTGCCAGCAGTGGAGTGGTTACCCATTCA |
| IGKV4-61*01_S2955_DBA1 | DBA/1J | Musmus IGKV4-61*01 F | 100 | 286 | CAAATTGTTCTCACCCAGTCTCCAGCAATCATGTCTGCATCTCCAGGGGAGAAGGTCACCATATCCTGCAGTGCCAGCTCAAGTGTAAGTTACATGTACTGGTACCAGCAGAAGCCAGGATCCTCCCCCAAACCCTGGATTTATCGCACATCCAACCTGGCTTCTGGAGTCCCTGCTCGCTTCAGTGGCAGTGGGTCTGGGACCTCTTACTCTCTCACAATCAGCAGCATGGAGGCTGAAGATGCTGCCACTTATTACTGCCAGCAGTATCATAGTTACCCACTCA |
| IGKV4-68*01_S3092_DBA1 | DBA/1J | Musmus IGKV4-68*01 F | 100 | 286 | CAAATTGTTCTCACCCAGTCTCCAGCACTCATGTCTGCATCTCCAGGGGAGAAGGTCACCATGACCTGCAGTGCCAGCTCAAGTGTAAGTTACATGTACTGGTACCAGCAGAAGCCAAGATCCTCCCCCAAACCCTGGATTTATCTCACATCCAACCTGGCTTCTGGAGTCCCTGCTCGCTTCAGTGGCAGTGGGTCTGGGACCTCTTACTCTCTCACAATCAGCAGCATGGAGGCTGAAGATGCTGCCACTTATTACTGCCAGCAGTGGAGTAGTAACCCACTCA |
| IGKV12-38*01_DBA1 | DBA/1J | Musmus IGKV12-38*01 F | 100 | 287 | GACATCCAGATGACTCAGTCTCCAGCCTCCCTGGCTGCATCTGTGGGAGAAACTGTCACCATCACATGTCGAGCAAGTGAGAACATTTACTACAGTTTAGCATGGTATCAGCAGAAGCAAGGGAAATCTCCTCAGCTCCTGATCTATAATGCAAACAGCTTGGAAGATGGTGTCCCATCGAGGTTCAGTGGCAGTGGATCTGGGACACAGTATTCTATGAAGATCAACAGCATGCAGCCTGAAGATACCGCAACTTATTTCTGTAAACAGGCTTATGACGTTCCTCC |
| IGKV4-52*01_S2975_DBA1 | DBA/1J | Musmus IGKV4-52*01 ORF, or Musmus IGKV4-54*01 ORF | 100 | 286 | CAAATTGTTCTCACCCAGTCTCCAGCAATCCTGTCTGCATCTCCAGGGGAGAAGGTCACCATGACCTGCAGTGCCAGCTCAAGTGTAAGTTACATGTACAGGTACCAGCAGAAGCCAGGATCCTCACCCAAACCCTGGATTTATGGCACATCCAACCTGGCTTCTGGAGTCCCTGCTCGCTTCAGTGGCAGTGGATCTGGGACCTCTTATTCTCTCACAATCAGCAGCATGGAGGCTGAAGATGCTGCCACTTATTACTGCCAGCAATATCATAGTTACCCGCTCA |
| IGKV4-62*01_S5834_DBA1 | DBA/1J | Musmus IGKV4-62*01 ORF | 100 | 286 | CAAATTGTTCTCACCCAGTCTCCAGCAATCATGTCTGCATCTCCAGGGGAAAAGGTCACCATGACCTGCAGTGCCAGCTCAAGTGTAAGTTACATGTACTGGTACCAGCAGAAGCCAGGCTCCTCCCCCAGACTCTGGATTTATGACACATCCAACCTGGTTTCTGGAGTCCCTGCTCGCTTCAGTGGCAGTAGGTCTGGGACCTCTTATTCTCTCACAATCAGCAGCATGGAGGCTGAAGATGCTGCCACTTATTACTGCCAGCAGTACAGTGGTTACCCATACA |
| IGKV5-37*01_DBA1 | DBA/1J | Musmus IGKV5-37*01 F | 100 | 287 | GACATCCTGCTGACCCAGTCTCCAGCCACCCTGTCTGTGACTCCAGGAGAAACAGTCAGTCTTTCCTGTAGGGCCAGCCAGAGTATTTACAAGAACCTACACTGGTATCAACAGAAATCACATCGGTCTCCAAGGCTTCTCATCAAGTATGCTTCTGATTCCATCTCTGGGATCCCCTCCAGGTTCACTGGCAGTGGATCAGGGACAGATTACACTCTCAGTATCAACAGTGTGAAGCCCGAAGATGAAGGAATATATTACTGTCTTCAAGGTTACAGCACACCTTC |

| | | | | | |
|---|---|---|---|---|---|
| IGKV9-123*01_S4103_DBA1 | DBA/1J | Musmus IGKV9-123*01 F | 100 | 285 | GACATCCAGATGATTCAGTCTCCATCGTCCATGTTTGCCTCTCTGGGAGACAGAGTCAGTCTCTCTTGTCGGGCTAGTCAGGGCATTAGAGGTAATTTAGACTGGTATCAGCAGAAACCAGGTGGAACTATTAAACTCCTGATCTACTCCACATCCAATTTAAATTCTGGTGTCCCATCAAGGTTCAGTGGCAGTGGGTCTGGGTCAGATTATTCTCTCACCATCAGCAGCCTAGAGTCTGAAGATTTTGCAGACTATTACTGTCTACAGCGTAATGCGTTTCCT |
| IGKV9-129*01_DBA1 | DBA/1J | Musmus IGKV9-129*01 ORF | 100 | 287 | GACATCCAGATGACCCAGTCTCCATCCTCCTTATCTGCCTCTCTGGGAGAAAGAGTCAGTCTCACTTGCCGGGCAAGTCAGGACATTCATGGTTATTTAAACTTGTTTCAGCAGAAACCAGGTGAAACTATTAAACACCTGATCTATGAAACATCCAATTTAGATTCTGGTGTCCCAAAAAGGTTCAGTGGCAGTAGGTCTGGGTCAGATTATTCTCTCATTATCGGCAGCCTTGAGTCTGAAGATTTTGCAGACTATTACTGTCTACAATATGCTAGTTCTCCTCC |
| IGKV1-132*01_DBA1 | DBA/1J | Musmus IGKV1-132*01 F | 100 | 302 | GATGTTGTGATGACCCAGACTCCACTGTCTTTGTCGGTTACCATTGGACAACCAGCCTCTATCTCTTGCAAGTCAAGTCAGAGCCTCTTATATAGTAATGGAAAGACATATTTGAATTGGTTACAACAGAGGCCTGGCCAGGCTCCAAAGCACCTAATGTATCAGGTGTCCAAACTGGACCCTGGCATCCCTGACAGGTTCAGTGGCAGTGGATCAGAAACAGATTTTACACTTAAAATCAGCAGAGTGGAGGCTGAAGATTTGGGAGTTTATTACTGCTTGCAAGGTACATATTATCCTCA |
| IGKV4-52*01_S5019_DBA1 | DBA/1J | Musmus IGKV4-52*01 ORF, or Musmus IGKV4-54*01 ORF | 100 | 286 | CAAATTGTTCTCACCCAGTCTCCAGCAATCCTGTCTGCATCTCCAGGGGAGAAGGTCACCATGACCTGCAGTGCCAGCTCAAGTGTAAGTTACATGTACAGGTACCAGCAGAAGCCAGGATCCTCACCCAAACCCTGGATTTATGGCACATCCAACCTGGCTTCTGGAGTCCCTGCTCGCTTCAGTGGCAGTGGATCTGGGACCTCTTATTCTCTCACAATCAGCAGCATGGAGGCTGAAGATGCTGCCACTTATTACTGCCAGCAATATCATAGTTACCCACTCA |
| IGKV6-14*01_DBA1 | DBA/1J | Musmus IGKV6-14*01 F | 100 | 287 | GACATTGTGATGACCCAGTCTCAAAAATTCATGTCCACATCAGTAGGAGACAGGGTCAGCATCACCTGCAAGGCCAGTCAGAATGTTCGTACTGCTGTAGCCTGGTATCAACAGAAACCAGGGCAGTCTCCTAAAGCACTGATTTACTTGGCATCCAACCGGCACACTGGAGTCCCTGATCGCTTCACAGGCAGTGGATCTGGGACAGATTTCACTCTCACCATTAGCAATGTGCAATCTGAAGACCTGGCAGATTATTTCTGTCTGCAACATTGGAATTATCCTCT |
| IGKV4-56*01_DBA1 | DBA/1J | Musmus IGKV4-56*01 P | 100 | 286 | CAAATTGTTCTCACCCAGTCTCCAGCAATCATGTCTGCATCTCCAGGGCAGAAAGTCACCATAACCTGCAGTGCCATCTCAAGTGTAAATTACATGCACTGGTACCAGCAGAAGCCAGGATCCTCCCCCAAACTCTGGATTTATGCAACATCCAAACTGGCTCTTGGAGTCCCTGCTTGCTTCAGTGGCAGTGGGTCTGGGACCTCTTACTCTCTCACAATCAGCAGCATGGTGGCTGAAGATGCCACCTCTTATTTCTGTCATCAGTGGAGTAGTTACCCACCCA |
| IGKV4-90*01_DBA1 | DBA/1J | Musmus IGKV4-90*01 F | 100 | 285 | GAAATTTTGCTCACCCAGTCTCCAGCAATCATAGCTGCATCTCCTGGGGAGAAGGTCACCATCACCTGCAGTGCCAGCTCAAGTGTAAGTTACATGAACTGGTACCAGCAGAAACCAGGATCCTCCCCCAAAATATGGATTTATGGTATATCCAACCTGGCTTCTGGAGTTCCTGCTCGCTTCAGTGGCAGTGGGTCTGGGACATCTTTCTCTTTCACAATCAACAGCATGGAGGCTGAAGATGTTGCCACTTATTACTGTCAGCAAAGGAGTAGTTACCCACCC |
| IGKV6-29*01_DBA1 | DBA/1J | Musmus IGKV6-29*01 F | 100 | 287 | AACATTGTAATGACCCAATCTCCCAAATCCATGTCCATGTCAGTAGGAGAGAGGGTCACCTTGAGCTGCAAGGCCAGTGAGAATGTGGGTACTTATGTATCCTGGTATCAACAGAAACCAGAGCAGTCTCCTAAACTGCTGATATACGGGGCATCCAACCGGTACCCTGGGGTCCCTGATCGCTTCACAGGCAGTGGATCTGCAACAGATTTCACTCTGACCATCAGCAGTCTGCAGGCTGAAGACCTTGCAGATTATCACTGTGGACAGGGTTACAGCTATCTTCC |

| | | | | | |
|---|---|---|---|---|---|
| IGKV3-3*01_DBA1 | DBA/1J | Musmus IGKV3-3*01 F | 100 | 299 | GACATTGTGCTGACCCAATCTCCAGCTTCTTTGGCTGTGTCTCTAGGACAGAGAGCCACTATCTTCTGCAGAGCCAGCCAGAGTGTCGATTATAATGGAATTAGTTATATGCACTGGTTCCAACAGAAACCAGGACAGCCACCCAAACTCCTCATCTATGCTGCATCCAACCTAGAATCTGGGATCCCTGCCAGGTTCAGTGGCAGTGGGTCTGGGACAGACTTCACCCTCAACATCCATCCTGTGGAGGAGGAAGATGCTGCAACCTATTACTGTCAGCAAAGTATTGAGGATCCTCC |
| IGKV3-9*01_S2885_DBA1 | DBA/1J | Musmus IGKV3-9*01 F | 99.66 | 296 | GACATTGTGCTGACCCAATCTCCAGCTTCTTTGGCTGTGTCTCTAGGACAGAGGGCCACCATATCCTGCCAAGCCAGCGAAAGTGTCAGTTTTGCTGGTACAAGTTTAATGCACTGGTACCAACAGAAACCAGGACAGCCACCCAAACTCCTCATCTATCGTGCATCCAACCTGGAATCTGGAGTCCCTGCCAGGTTCAGTGGCAGTGGGTCTGAGTCAGACTTCACTCTCACCATCGATCCTGTGGAGGAAGATGATGCTGCAATGTATTACTGTATGCAAAGTATGGAAGATCC |
| IGKV4-78*01_S6302_DBA1 | DBA/1J | Musmus IGKV4-78*01 F | 100 | 292 | CAAATTGTTCTCACCCAGTCTCCAGCAATCATGTCTGCATCTCCTGGGGAGAAGGTCACCATGACCTGCAGTGCCAGATCAAGTGTAAGTTCCAGCTACTTGTACTGGTACCAGCAGAAGCCAGGATCCTCCCCCAAACTCTGGATTTATAGCACATCCAACCTGGCTTCTGGAGTCCCTGCTCGCTTCAGTGGCAGTGGGTCTGGGACCTCTTATTCTCTCACAATCAGCAGCATGGAGGCTGAAGATGCTGCCACTTTTTACTGCCAGCAGTACAGTGGTTACCCATACA |
| IGKV6-b*01_DBA1 | DBA/1J | Musmus IGKV6-b*01 F | 100 | 287 | AGTATTGTGATGACCCAGACTCCCAAATTCCTGCCTGTATCAGCAGGAGACAGGGTTACCATGACCTGCAAGGCCAGTCAGAGTGTGGGTAATAATGTAGCCTGGTACCAACAGAAGCCAGGACAGTCTCCTAAACTGCTGATATACTATGCATCCAATCGCTACACTGGAGTCCCTGATCGCTTCACTGGCAGTGGATCTGGGACAGATTTCACTTTCACCATCAGCAGTGTGCAGGTTGAAGACCTGGCAGTTTATTTCTGTCAGCAGCATTATAGCTCTCCTCC |
| IGKV6-15*01_DBA2 | DBA/2J | Musmus IGKV6-15*01 F | 100 | 287 | GACATTGTGATGACCCAGTCTCAAAAATTCATGTCCACATCAGTAGGAGACAGGGTCAGCGTCACCTGCAAGGCCAGTCAGAATGTGGGTACTAATGTAGCCTGGTATCAACAGAAACCAGGGCAATCTCCTAAAGCACTGATTTACTCGGCATCCTACCGGTACAGTGGAGTCCCTGATCGCTTCACAGGCAGTGGATCTGGGACAGATTTCACTCTCACCATCAGCAATGTGCAGTCTGAAGACTTGGCAGAGTATTTCTGTCAGCAATATAACAGCTATCCTCT |
| IGKV8-30*01_DBA2 | DBA/2J | Musmus IGKV8-30*01 F | 100 | 305 | GACATTGTGATGTCACAGTCTCCATCCTCCCTAGCTGTGTCAGTTGGAGAGAAGGTTACTATGAGCTGCAAGTCCAGTCAGAGCCTTTTATATAGTAGCAATCAAAAGAACTACTTGGCCTGGTACCAGCAGAAACCAGGGCAGTCTCCTAAACTGCTGATTTACTGGGCATCCACTAGGGAATCTGGGGTCCCTGATCGCTTCACAGGCAGTGGATCTGGGACAGATTTCACTCTCACCATCAGCAGTGTGAAGGCTGAAGACCTGGCAGTTTATTACTGTCAGCAATATTATAGCTATCCTCC |
| IGKV8-27*01_DBA2 | DBA/2J | Musmus IGKV8-27*01 F | 100 | 300 | AACATTATGATGACACAGTCGCCATCATCTCTGGCTGTGTCTGCAGGAGAAAAGGTCACTATGAGCTGTAAGTCCAGTCAAAGTGTTTTATACAGTTCAAATCAGAAGAACTACTTGGCCTGGTACCAGCAGAAACCAGGGCAGTCTCCTAAACTGCTGATCTACTGGGCATCCACTAGGGAATCTGGTGTCCCTGATCGCTTCACAGGCAGTGGATCTGGGACAGATTTTACTCTTACCATCAGCAGTGTACAAGCTGAAGACCTGGCAGTTTATTACTGTCATCAATACCTCTCCTCG |

| | | | | | |
|---|---|---|---|---|---|
| IGKV12-44*01_DBA2 | DBA/2J | Musmus IGKV12-44*01 F | 100 | 287 | GACATCCAGATGACTCAGTCTCCAGCCTCCCTATCTGCATCTGTGGGAGAAACTGTCACCATCACATGTCGAGCAAGTGAGAATATTTACAGTTATTTAGCATGGTATCAGCAGAAACAGGGAAAATCTCCTCAGCTCCTGGTCTATAATGCAAAAACCTTAGCAGAAGGTGTGCCATCAAGGTTCAGTGGCAGTGGATCAGGCACACAGTTTTCTCTGAAGATCAACAGCCTGCAGCCTGAAGATTTTGGGAGTTATTACTGTCAACATCATTATGGTACTCCTCC |
| IGKV1-110*01_DBA2 | DBA/2J | Musmus IGKV1-110*01 F | 100 | 302 | GATGTTGTGATGACCCAAACTCCACTCTCCCTGCCTGTCAGTCTTGGAGATCAAGCCTCCATCTCTTGCAGATCTAGTCAGAGCCTTGTACACAGTAATGGAAACACCTATTTACATTGGTACCTGCAGAAGCCAGGCCAGTCTCCAAAGCTCCTGATCTACAAAGTTTCCAACCGATTTTCTGGGGTCCCAGACAGGTTCAGTGGCAGTGGATCAGGGACAGATTTCACACTCAAGATCAGCAGAGTGGAGGCTGAGGATCTGGGAGTTTATTTCTGCTCTCAAAGTACACATGTTCCTCC |
| IGKV4-59*01_DBA2 | DBA/2J | Musmus IGKV4-59*01 F | 100 | 286 | CAAATTGTTCTCACCCAGTCTCCAGCAATCATGTCTGCATCTCCAGGGGAGAAGGTCACCATGACCTGCAGTGCCAGCTCAAGTGTAAGTTACATGCACTGGTACCAGCAGAAGTCAGGCACCTCCCCCAAAAGATGGATTTATGACACATCCAAACTGGCTTCTGGAGTCCCTGCTCGCTTCAGTGGCAGTGGGTCTGGGACCTCTTACTCTCTCACAATCAGCAGCATGGAGGCTGAAGATGCTGCCACTTATTACTGCCAGCAGTGGAGTAGTAACCCACCCA |
| IGKV6-23*01_DBA2 | DBA/2J | Musmus IGKV6-23*01 F | 100 | 287 | GACATTGTGATGACCCAGTCTCACAAATTCATGTCCACATCAGTAGGAGACAGGGTCAGCATCACCTGCAAGGCCAGTCAGGATGTGGGTACTGCTGTAGCCTGGTATCAACAGAAACCAGGGCAATCTCCTAAACTACTGATTTACTGGGCATCCACCCGGCACACTGGAGTCCCTGATCGCTTCACAGGCAGTGGATCTGGGACAGATTTCACTCTCACCATTAGCAATGTGCAGTCTGAAGACTTGGCAGATTATTTCTGTCAGCAATATAGCAGCTATCCTCT |
| IGKV8-19*01_DBA2 | DBA/2J | Musmus IGKV8-19*01 F | 100 | 305 | GACATTGTGATGACACAGTCTCCATCCTCCCTGACTGTGACAGCAGGAGAGAAGGTCACTATGAGCTGCAAGTCCAGTCAGAGTCTGTTAAACAGTGGAAATCAAAAGAACTACTTGACCTGGTACCAGCAGAAACCAGGGCAGCCTCCTAAACTGTTGATCTACTGGGCATCCACTAGGGAATCTGGGGTCCCTGATCGCTTCACAGGCAGTGGATCTGGAACAGATTTCACTCTCACCATCAGCAGTGTGCAGGCTGAAGACCTGGCAGTTTATTACTGTCAGAATGATTATAGTTATCCTCC |
| IGKV1-135*01_DBA2 | DBA/2J | Musmus IGKV1-135*01 F | 100 | 302 | GATGTTGTGATGACCCAGACTCCACTCACTTTGTCGGTTACCATTGGACAACCAGCCTCCATCTCTTGCAAGTCAAGTCAGAGCCTCTTAGATAGTGATGGAAAGACATATTTGAATTGGTTGTTACAGAGGCCAGGCCAGTCTCCAAAGCGCCTAATCTATCTGGTGTCTAAACTGGACTCTGGAGTCCCTGACAGGTTCACTGGCAGTGGATCAGGGACAGATTTCACACTGAAAATCAGCAGAGTGGAGGCTGAGGATTTGGGAGTTTATTATTGCTGGCAAGGTACACATTTTCCTCA |
| IGKV2-137*01_DBA2 | DBA/2J | Musmus IGKV2-137*01 F | 100 | 302 | GATATTGTGATGACTCAGGCTGCACCCTCTGTACCTGTCACTCCTGGAGAGTCAGTATCCATCTCCTGCAGGTCTAGTAAGAGTCTCCTGCATAGTAATGGCAACACTTACTTGTATTGGTTCCTGCAGAGGCCAGGCCAGTCTCCTCAGCTCCTGATATATCGGATGTCCAACCTTGCCTCAGGAGTCCCAGACAGGTTCAGTGGCAGTGGGTCAGGAACTGCTTTCACACTGAGAATCAGTAGAGTGGAGGCTGAGGATGTGGGTGTTTATTACTGTATGCAACATCTAGAATATCCTTT |
| IGKV14-126*01_DBA2 | DBA/2J | Musmus IGKV14-126*01 F | 100 | 287 | GACATCAAGATGACCCAGTCTCCATCCTCCATGTATGCATCGCTGGGAGAGAGAGTCACTATCACTTGCAAGGCGAGTCAGGACATTAAAAGCTATTTAAGCTGGTACCAGCAGAAACCATGGAAATCTCCTAAGACCCTGATCTATTATGCAACAAGCTTGGCAGATGGGGTCCCATCAAGATTCAGTGGCAGTGGATCTGGGCAAGATTATTCTCTAACCATCAGCAGCCTGGAGTCTGACGATACAGCAACTTATTACTGTCTACAGCATGGTGAGAGCCCTCC |

| | | | | | |
|---|---|---|---|---|---|
| IGKV8-21*01_DBA2 | DBA/2J | Musmus IGKV8-21*01 F | 100 | 302 | GACATTGTGATGTCACAGTCTCCATCCTCCCTGGCTGTGTCAGCAGGAGAGAAGGTCACTATGAGCTGCAAATCCAGTCAGAGTCTGCTCAACAGTAGAACCCGAAAGAACTACTTGGCTTGGTACCAGCAGAAACCAGGGCAGTCTCCTAAACTGCTGATCTACTGGGCATCCACTAGGGAATCTGGGGTCCCTGATCGCTTCACAGGCAGTGGATCTGGGACAGATTTCACTCTCACCATCAGCAGTGTGCAGGCTGAAGACCTGGCAGTTTATTACTGCAAGCAATCTTATAATCTTCC |
| IGKV12-89*01_DBA2 | DBA/2J | Musmus IGKV12-89*01 F | 100 | 287 | GACATCCAGATGACTCAGTCTCCAGCTTCACTGTCTGCATCTGTGGGAGAAACTGTCACCATCACATGTGGAGCAAGTGAGAATATTTACGGTGCTTTAAATTGGTATCAGCGGAAACAGGGAAAATCTCCTCAGCTCCTGATCTATGGTGCAACCAACTTGGCAGATGGCATGTCATCGAGGTTCAGTGGCAGTGGATCTGGTAGACAGTATTCTCTCAAGATCAGTAGCCTGCATCCTGACGATGTTGCAACGTATTACTGTCAAAATGTGTTAAGTACTCCTCC |
| IGKV10-96*01_DBA2 | DBA/2J | Musmus IGKV10-96*01 F | 100 | 286 | GATATCCAGATGACACAGACTACATCCTCCCTGTCTGCCTCTCTGGGAGACAGAGTCACCATCAGTTGCAGGGCAAGTCAGGACATTAGCAATTATTTAAACTGGTATCAGCAGAAACCAGATGGAACTGTTAAACTCCTGATCTACTACACATCAAGATTACACTCAGGAGTCCCATCAAGGTTCAGTGGCAGTGGGTCTGGAACAGATTATTCTCTCACCATTAGCAACCTGGAGCAAGAAGATATTGCCACTTACTTTTGCCAACAGGGTAATACGCTTCCTC |
| IGKV8-24*01_DBA2 | DBA/2J | Musmus IGKV8-24*01 F | 100 | 305 | GACATTGTGATGACACAGTCTCCATCCTCCCTGGCTATGTCAGTAGGACAGAAGGTCACTATGAGCTGCAAGTCCAGTCAGAGCCTTTTAAATAGTAGCAATCAAAAGAACTATTTGGCCTGGTACCAGCAGAAACCAGGACAGTCTCCTAAACTTCTGGTATACTTTGCATCCACTAGGGAATCTGGGGTCCCTGATCGCTTCATAGGCAGTGGATCTGGGACAGATTTCACTCTTACCATCAGCAGTGTGCAGGCTGAAGACCTGGCAGATTACTTCTGTCAGCAACATTATAGCACTCCTCC |
| IGKV10-94*01_DBA2 | DBA/2J | Musmus IGKV10-94*01 F | 100 | 287 | GATATCCAGATGACACAGACTACATCCTCCCTGTCTGCCTCTCTGGGAGACAGAGTCACCATCAGTTGCAGTGCAAGTCAGGGCATTAGCAATTATTTAAACTGGTATCAGCAGAAACCAGATGGAACTGTTAAACTCCTGATCTATTACACATCAAGTTTACACTCAGGAGTCCCATCAAGGTTCAGTGGCAGTGGGTCTGGGACAGATTATTCTCTCACCATCAGCAACCTGGAACCTGAAGATATTGCCACTTACTATTGTCAGCAGTATAGTAAGCTTCCTCC |
| IGKV1-88*01_DBA2 | DBA/2J | Musmus IGKV1-88*01 F | 100 | 302 | GATGTTGTGGTGACTCAAACTCCACTCTCCCTGCCTGTCAGCTTTGGAGATCAAGTTTCTATCTCTTGCAGGTCTAGTCAGAGTCTTGCAAACAGTTATGGGAACACCTATTTGTCTTGGTACCTGCACAAGCCTGGCCAGTCTCCACAGCTCCTCATCTATGGGATTTCCAACAGATTTTCTGGGGTGCCAGACAGGTTCAGTGGCAGTGGTTCAGGGACAGATTTCACACTCAAGATCAGCACAATAAAGCCTGAGGACTTGGGAATGTATTACTGCTTACAAGGTACACATCAGCCTCC |
| IGKV1-117*01_DBA2 | DBA/2J | Musmus IGKV1-117*01 F | 100 | 302 | GATGTTTTGATGACCCAAACTCCACTCTCCCTGCCTGTCAGTCTTGGAGATCAAGCCTCCATCTCTTGCAGATCTAGTCAGAGCATTGTACATAGTAATGGAAACACCTATTTAGAATGGTACCTGCAGAAACCAGGCCAGTCTCCAAAGCTCCTGATCTACAAAGTTTCCAACCGATTTTCTGGGGTCCCAGACAGGTTCAGTGGCAGTGGATCAGGGACAGATTTCACACTCAAGATCAGCAGAGTGGAGGCTGAGGATCTGGGAGTTTATTACTGCTTTCAAGGTTCACATGTTCCTCC |
| IGKV6-32*01_DBA2 | DBA/2J | Musmus IGKV6-32*01 F | 100 | 287 | AGTATTGTGATGACCCAGACTCCCAAATTCCTGCTTGTATCAGCAGGAGACAGGGTTACCATAACCTGCAAGGCCAGTCAGAGTGTGAGTAATGATGTAGCTTGGTACCAACAGAAGCCAGGGCAGTCTCCTAAACTGCTGATATACTATGCATCCAATCGCTACACTGGAGTCCCTGATCGCTTCACTGGCAGTGGATATGGGACGGATTTCACTTTCACCATCAGCACTGTGCAGGCTGAAGACCTGGCAGTTTATTTCTGTCAGCAGGATTATAGCTCTCCTCC |

| | | | | | |
|---|---|---|---|---|---|
| IGKV12-46*01_DBA2 | DBA/2J | Musmus IGKV12-46*01 F | 100 | 287 | GACATCCAGATGACTCAGTCTCCAGCCTCCCTATCTGTATCTGTGGGAGAAACTGTCACCATCACATGTCGAGCAAGTGAGAATATTTACAGTAATTTAGCATGGTATCAGCAGAAACAGGGAAAATCTCCTCAGCTCCTGGTCTATGCTGCAACAAACTTAGCAGATGGTGTGCCATCAAGGTTCAGTGGCAGTGGATCAGGCACACAGTATTCCCTCAAGATCAACAGCCTGCAGTCTGAAGATTTTGGGAGTTATTACTGTCAACATTTTTGGGGTACTCCTCC |
| IGKV6-20*01_DBA2 | DBA/2J | Musmus IGKV6-20*01 F | 100 | 287 | AACATTGTAATGACCCAATCTCCCAAATCCATGTCCATGTCAGTAGGAGAGAGGGTCACCTTGAGCTGCAAGGCCAGTGAGAATGTGGGTACTTATGTATCCTGGTATCAACAGAAACCAGAGCAGTCTCCTAAACTGCTGATATACGGGGCATCCAACCGGTACACTGGGGTCCCCGATCGCTTCACAGGCAGTGGATCTGCAACAGATTTCACTCTGACCATCAGCAGTGTGCAGGCTGAAGACCTTGCAGATTATCACTGTGGACAGAGTTACAGCTATCCTCC |
| IGKV12-41*01_DBA2 | DBA/2J | Musmus IGKV12-41*01 F | 100 | 287 | GACATCCAGATGACTCAGTCTCCAGCCTCCCTATCTGCATCTGTGGGAGAAACTGTCACCATCACATGTCGAGCAAGTGGGAATATTCACAATTATTTAGCATGGTATCAGCAGAAACAGGGAAAATCTCCTCAGCTCCTGGTCTATAATGCAAAAACCTTAGCAGATGGTGTGCCATCAAGGTTCAGTGGCAGTGGATCAGGAACACAATATTCTCTCAAGATCAACAGCCTGCAGCCTGAAGATTTTGGGAGTTATTACTGTCAACATTTTTGGAGTACTCCTCC |
| IGKV4-70*01_DBA2 | DBA/2J | Musmus IGKV4-70*01 F | 100 | 282 | CAAATTGTTCTCACCCAGTCTCCAGCAATCATGTCTGCATCTCCAGGGGAGAAGGTCACCATGACCTGCAGTGCCAGCTCAAGTATAAGTTACATGCACTGGTACCAGCAGAAGCCAGGCACCTCCCCCAAAAGATGGATTTATGACACATCCAAACTGGCTTCTGGAGTCCCTGCTCGCTTCAGTGGCAGTGGGTCTGGGACCTCTTATTCTCTCACAATCAGCAGCATGGAGGCTGAAGATGCTGCCACTTATTACTGCCATCAGCGGAGTAGTTACCCA |
| IGKV8-28*01_DBA2 | DBA/2J | Musmus IGKV8-28*01 F | 100 | 305 | GACATTGTGATGACACAGTCTCCATCCTCCCTGAGTGTGTCAGCAGGAGAGAAGGTCACTATGAGCTGCAAGTCCAGTCAGAGTCTGTTAAACAGTGGAAATCAAAAGAACTACTTGGCCTGGTACCAGCAGAAACCAGGGCAGCCTCCTAAACTGTTGATCTACGGGGCATCCACTAGGGAATCTGGGGTCCCTGATCGCTTCACAGGCAGTGGATCTGGAACCGATTTCACTCTTACCATCAGCAGTGTGCAGGCTGAAGACCTGGCAGTTTATTACTGTCAGAATGATCATAGTTATCCTCC |
| IGKV6-13*01_DBA2 | DBA/2J | Musmus IGKV6-13*01 F | 100 | 285 | GACATTGTGATGACCCAGTCTCAAAAATTCATGTCCACATCAGTAGGAGACAGGGTCAGCATCACCTGCAAGGCCAGTCAGAATGTGGGTACTGCTGTAGCCTGGTATCAACAGAAACCAGGACAATCTCCTAAACTACTGATTTACTCGGCATCCAATCGGTACACTGGAGTCCCTGATCGCTTCACAGGCAGTGGATCTGGGACAGATTTCACTCTCACCATCAGCAATATGCAGTCTGAAGACCTGGCAGATTATTTCTGCCAGCAATATAGCAGCTATCCT |
| IGKV7-33*01_DBA2 | DBA/2J | Musmus IGKV7-33*01 F | 100 | 305 | GACATTGTGATGACTCAGTCTCCAACTTTCCTTGCTGTGACAGCAAGTAAGAAGGTCACCATTAGTTGCACGGCCAGTGAGAGCCTTTATTCAAGCAAACACAAGGTGCACTACTTGGCTTGGTACCAGAAGAAACCAGAGCAATCTCCTAAACTGCTGATATACGGGGCATCCAACCGATACATTGGGGTCCCTGATCGCTTCACAGGCAGTGGATCTGGGACAGATTTCACTCTGACCATCAGCAGTGTACAGGTTGAAGACCTCACACATTATTACTGTGCACAGTTTTACAGCTATCCTCT |
| IGKV5-43*01_DBA2 | DBA/2J | Musmus IGKV5-43*01 F | 100 | 287 | GATATTGTGCTAACTCAGTCTCCAGCCACCCTGTCTGTGACTCCAGGAGATAGCGTCAGTCTTTCCTGCAGGGCCAGCCAAAGTATTAGCAACAACCTACACTGGTATCAACAAAAATCACATGAGTCTCCAAGGCTTCTCATCAAGTATGCTTCCCAGTCCATCTCTGGGATCCCCTCCAGGTTCAGTGGCAGTGGATCAGGGACAGATTTCACTCTCAGTATCAACAGTGTGGAGACTGAAGATTTTGGAATGTATTTCTGTCAACAGAGTAACAGCTGGCCTCA |

| | | | | | |
|---|---|---|---|---|---|
| IGKV6-17*01_DBA2 | DBA/2J | Musmus IGKV6-17*01 F | 100 | 287 | GACATTGTGATGACCCAGTCTCACAAATTCATGTCCACATCAGTAGGAGACAGGGTCAGCATCACCTGCAAGGCCAGTCAGGATGTGAGTACTGCTGTAGCCTGGTATCAACAGAAACCAGGACAATCTCCTAAACTACTGATTTACTCGGCATCCTACCGGTACACTGGAGTCCCTGATCGCTTCACTGGCAGTGGATCTGGGACGGATTTCACTTTCACCATCAGCAGTGTGCAGGCTGAAGACCTGGCAGTTTATTACTGTCAGCAACATTATAGTACTCCTCC |
| IGKV3-7*01_S5296_DBA2 | DBA/2J | Musmus IGKV3-7*02 F | 100 | 299 | GACATTGTGCTGACACAGTCTCCTGCTTCCTTAGCTGTATCTCTGGGGCAGAGGGCCACCATCTCATGCAGGGCCAGCCAAAGTGTCAGTACATCTAGCTATAGTTATATGCACTGGTACCAACAGAAACCAGGACAGCCACCCAAACTCCTCATCAAGTATGCATCCAACCTAGAATCTGGGGTCCCTGCCAGGTTCAGTGGCAGTGGGTCTGGGACAGACTTCACCCTCAACATCCATCCTGTGGAGGAGGAGGATACTGCAACATATTACTGTCAGCACAGTTGGGAGATTCCTCC |
| IGKV8-16*01_DBA2 | DBA/2J | Musmus IGKV8-16*01 F | 100 | 305 | GAAATTGTGTTGACCCAGTCTATACCATCCCTGACTGTGTCAGCAGGAGAGAGGGTCACTATCAGCTGCAAATCCAATCAGAATCTTTTATGGAGTGGAAACCAAAGGTACTGTTTGGTCTGGCACCAGTGGAAACCAGGGCAAACTCCTACACCGTTGATCACCTGGACATCTGATAGGTACTCTGGAGTCCCTGATCGTTTCATAGGCAGTGGATCTGTGACAGATTTCACTCTGACCATCAGCAGTGTGCAGGCTGAAGATGTGGCAGTTTATTTCTGTCAGCAGCATTTACACATTCCTCC |
| IGKV3-4*01_DBA2 | DBA/2J | Musmus IGKV3-4*01 F | 100 | 299 | GACATTGTGCTGACCCAATCTCCAGCTTCTTTGGCTGTGTCTCTAGGGCAGAGGGCCACCATCTCCTGCAAGGCCAGCCAAAGTGTTGATTATGATGGTGATAGTTATATGAACTGGTACCAACAGAAACCAGGACAGCCACCCAAACTCCTCATCTATGCTGCATCCAATCTAGAATCTGGGATCCCAGCCAGGTTTAGTGGCAGTGGGTCTGGGACAGACTTCACCCTCAACATCCATCCTGTGGAGGAGGAGGATGCTGCAACCTATTACTGTCAGCAAAGTAATGAGGATCCTCC |
| IGKV19-93*01_DBA2 | DBA/2J | Musmus IGKV19-93*01 F | 100 | 287 | GACATCCAGATGACACAGTCTCCATCCTCACTGTCTGCATCTCTGGGAGGCAAAGTCACCATCACTTGCAAGGCAAGCCAAGACATTAACAAGTATATAGCTTGGTACCAACACAAGCCTGGAAAAGGTCCTAGGCTGCTCATACATTACACATCTACATTACAGCCAGGCATCCCATCAAGGTTCAGTGGAAGTGGGTCTGGGAGAGATTATTCCTTCAGCATCAGCAACCTGGAGCCTGAAGATATTGCAACTTATTATTGTCTACAGTATGATAATCTTCTACC |
| IGKV3-2*01_DBA2 | DBA/2J | Musmus IGKV3-2*01 F | 100 | 299 | GACATTGTGCTGACCCAATCTCCAGCTTCTTTGGCTGTGTCTCTAGGGCAGAGGGCCACCATCTCCTGCAGAGCCAGCGAAAGTGTTGATAATTATGGCATTAGTTTTATGAACTGGTTCCAACAGAAACCAGGACAGCCACCCAAACTCCTCATCTATGCTGCATCCAACCAAGGATCCGGGGTCCCTGCCAGGTTTAGTGGCAGTGGGTCTGGGACAGACTTCAGCCTCAACATCCATCCTATGGAGGAGGATGATACTGCAATGTATTTCTGTCAGCAAAGTAAGGAGGTTCCTCC |
| IGKV3-12*01_DBA2 | DBA/2J | Musmus IGKV3-12*01 F | 100 | 299 | GACATTGTGCTGACACAGTCTCCTGCTTCCTTAGCTGTATCTCTGGGGCAGAGGGCCACCATCTCATGCAGGGCCAGCAAAAGTGTCAGTACATCTGGCTATAGTTATATGCACTGGTACCAACAGAAACCAGGACAGCCACCCAAACTCCTCATCTATCTTGCATCCAACCTAGAATCTGGGGTCCCTGCCAGGTTCAGTGGCAGTGGGTCTGGGACAGACTTCACCCTCAACATCCATCCTGTGGAGGAGGAGGATGCTGCAACCTATTACTGTCAGCACAGTAGGGAGCTTCCTCC |

| | | | | | |
|---|---|---|---|---|---|
| IGKV3-5*01_DBA2 | DBA/2J | Musmus IGKV3-5*01 F | 100 | 299 | GACATTGTGCTGACCCAATCTCCAGCTTCTTTGGCTGTGTCTCTAGGGCAGAGGGCCACCATATCCTGCAGAGCCAGTGAAAGTGTTGATAGTTATGGCAATAGTTTTATGCACTGGTACCAGCAGAAACCAGGACAGCCACCCAAACTCCTCATCTATCGTGCATCCAACCTAGAATCTGGGATCCCTGCCAGGTTCAGTGGCAGTGGGTCTAGGACAGACTTCACCCTCACCATTAATCCTGTGGAGGCTGATGATGTTGCAACCTATTACTGTCAGCAAAGTAATGAGGATCCTCC |
| IGKV4-74*01_DBA2 | DBA/2J | Musmus IGKV4-74*01 F | 100 | 292 | CAAATTGTTCTCACCCAGTCTCCAGCAATCATGTCTGCATCTCTAGGGGAACGGGTCACCATGACCTGCACTGCCAGCTCAAGTGTAAGTTCCAGTTACTTGCACTGGTACCAGCAGAAGCCAGGATCCTCCCCCAAACTCTGGATTTATAGCACATCCAACCTGGCTTCTGGAGTCCCAGCTCGCTTCAGTGGCAGTGGGTCTGGGACCTCTTACTCTCTCACAATCAGCAGCATGGAGGCTGAAGATGCTGCCACTTATTACTGCCACCAGTATCATCGTTCCCCACCCA |
| IGKV4-57*01_S9219_DBA2 | DBA/2J | Musmus IGKV4-57-1*01 F | 100 | 286 | CAAATTGTTCTCACCCAGTCTCCAGCAATCATGTCTGCATCTCCAGGGGAGAAGGTCACCATAACCTGCAGTGCCAGCTCAAGTGTAAGTTACATGCACTGGTTCCAGCAGAAGCCAGGCACTTCTCCCAAACTCTGGATTTATAGCACATCCAACCTGGCTTCTGGAGTCCCTGCTCGCTTCAGTGGCAGTGGATCTGGGACCTCTTACTCTCTCACAATCAGCCGAATGGAGGCTGAAGATGCTGCCACTTATTACTGCCAGCAAAGGAGTAGTTACCCACTCA |
| IGKV4-80*01_DBA2 | DBA/2J | Musmus IGKV4-80*01 F | 100 | 282 | CAAATTGTTCTCACCCAGTCTCCAGCAATCATGTCTGCATCTCTAGGGGAGGAGATCACCCTAACCTGCAGTGCCAGCTCGAGTGTAAGTTACATGCACTGGTACCAGCAGAAGTCAGGCACTTCTCCCAAACTCTTGATTTATAGCACATCCAACCTGGCTTCTGGAGTCCCTTCTCGCTTCAGTGGCAGTGGGTCTGGGACCTTTTATTCTCTCACAATCAGCAGTGTGGAGGCTGAAGATGCTGCCGATTATTACTGCCATCAGTGGAGTAGTTATCCA |
| IGKV4-57-1*01_DBA2 | DBA/2J | Musmus IGKV4-57*01 F | 100 | 292 | GAAAATGTGCTCACCCAGTCTCCAGCAATCATGTCTGCATCTCCAGGGGAAAAGGTCACCATGACCTGCAGGGCCAGCTCAAGTGTAAGTTCCAGTTACTTGCACTGGTACCAGCAGAAGTCAGGTGCCTCCCCCAAACTCTGGATTTATAGCACATCCAACTTGGCTTCTGGAGTCCCTGCTCGCTTCAGTGGCAGTGGGTCTGGGACCTCTTACTCTCTCACAATCAGCAGTGTGGAGGCTGAAGATGCTGCCACTTATTACTGCCAGCAGTACAGTGGTTACCCACTCA |
| IGKV9-120*01_S2628_DBA2 | DBA/2J | Musmus IGKV9-120*02 F | 100 | 285 | GACATCCAGATGACCCAGTCTCCATCCTCCTTATCTGCCTCTCTGGGAGAAAGAGTCAGTCTCACTTGTCGGGCAAGTCAGGACATTGGTAGTAGCTTAAACTGGCTTCAGCAGGAACCAGATGGAACTATTAAACGCCTGATCTACGCCACATCCAGTTTAGATTCTGGTGTCCCCAAAAGGTTCAGTGGCAGTAGGTCTGGGTCAGATTATTCTCTCACCATCAGCAGCCTTGAGTCTGAAGATTTTGTAGACTATTACTGTCTACAATATGCTAGTTCTCCT |
| IGKV15-103*01_DBA2 | DBA/2J | Musmus IGKV15-103*01 ORF | 100 | 287 | GACATCCAGATGAACCAGTCTCCATCCAGTCTGTCTGCATCCCTTGGAGACACAATTACCATCACTTGCCATGCCAGTCAGAACATTAATGTTTGGTTAAGCTGGTACCAGCAGAAACCAGGAAATATTCCTAAACTATTGATCTATAAGGCTTCCAACTTGCACACAGGCGTCCCATCAAGGTTTAGTGGCAGTGGATCTGGAACAGGTTTCACATTAACCATCAGCAGCCTGCAGCCTGAAGACATTGCCACTTACTACTGTCAACAGGGTCAAAGTTATCCTCT |
| IGKV16-104*01_DBA2 | DBA/2J | Musmus IGKV16-104*01 F | 100 | 288 | GATGTCCAGATAACCCAGTCTCCATCTTATCTTGCTGCATCTCCTGGAGAAACCATTACTATTAATTGCAGGGCAAGTAAGAGCATTAGCAAATATTTAGCCTGGTATCAAGAGAAACCTGGGAAAACTAATAAGCTTCTTATCTACTCTGGATCCACTTTGCAATCTGGAATTCCATCAAGGTTCAGTGGCAGTGGATCTGGTACAGATTTCACTCTCACCATCAGTAGCCTGGAGCCTGAAGATTTTGCAATGTATTACTGTCAACAGCATAATGAATACCCGTAC |

| | | | | | |
|---|---|---|---|---|---|
| IGKV4-68*01_DBA2 | DBA/2J | Musmus IGKV4-68*01 F | 100 | 286 | CAAATTGTTCTCACCCAGTCTCCAGCACTCATGTCTGCATCTCCAGGGGAGAAGGTCACCATGACCTGCAGTGCCAGCTCAAGTGTAAGTTACATGTACTGGTACCAGCAGAAGCCAAGATCCTCCCCCAAACCCTGGATTTATCTCACATCCAACCTGGCTTCTGGAGTCCCTGCTCGCTTCAGTGGCAGTGGGTCTGGGACCTCTTACTCTCTCACAATCAGCAGCATGGAGGCTGAAGATGCTGCCACTTATTACTGCCAGCAGTGGAGTAGTAACCCACCCA |
| IGKV17-121*01_DBA2 | DBA/2J | Musmus IGKV17-121*01 F | 100 | 287 | GAAACAACTGTGACCCAGTCTCCAGCATCCCTGTCCATGGCTATAGGAGAAAAAGTCACCATCAGATGCATAACCAGCACTGATATTGATGATGATATGAACTGGTACCAGCAGAAGCCAGGGGAACCTCCTAAGCTCCTTATTTCAGAAGGCAATACTCTTCGTCCTGGAGTCCCATCCCGATTCTCCAGCAGTGGCTATGGTACAGATTTTGTTTTTACAATTGAAAACATGCTCTCAGAAGATGTTGCAGATTACTACTGTTTGCAAAGTGATAACTTGCCTCT |
| IGKV4-55*01_S6279_DBA2 | DBA/2J | Musmus IGKV4-55*01 F | 100 | 286 | CAAATTGTTCTCACCCAGTCTCCAGCAATCATGTCTGCATCTCCAGGGGAGAAGGTCACCATGACCTGCAGTGCCAGCTCAAGTGTAAGTTACATGTACTGGTACCAGCAGAAGCCAGGATCCTCCCCCAGACTCCTGATTTATGACACATCCAACCTGGCTTCTGGAGTCCCTGTTCGCTTCAGTGGCAGTGGGTCTGGGACCTCTTACTCTCTCACAATCAGCCGAATGGAGGCTGAAGATGCTGCCACTTATTACTGCCAGCAGTGGAGTAGTTACCCACTCA |
| IGKV13-85*01_DBA2 | DBA/2J | Musmus IGKV13-85*01 F | 100 | 287 | GACATCCAGATGACACAATCTTCATCCTACTTGTCTGTATCTCTAGGAGGCAGAGTCACCATTACTTGCAAGGCAAGTGACCACATTAATAATTGGTTAGCCTGGTATCAGCAGAAACCAGGAAATGCTCCTAGGCTCTTAATATCTGGTGCAACCAGTTTGGAAACTGGGGTTCCTTCAAGATTCAGTGGCAGTGGATCTGGAAAGGATTACACTCTCAGCATTACCAGTCTTCAGACTGAAGATGTTGCTACTTATTACTGTCAACAGTATTGGAGTACTCCTCC |
| IGKV17-127*01_DBA2 | DBA/2J | Musmus IGKV17-127*01 F | 100 | 287 | GAAACAACTGTGACCCAGTCTCCAGCATCCCTGTCCGTGGCTACAGGAGAAAAAGTCACTATCAGATGCATAACCAGCACTGATATTGATGATGATATGAACTGGTACCAGCAGAAGCCAGGGGAACCTCCTAAGCTCCTTATTTCAGAAGGCAATACTCTTCGTCCTGGAGTCCCATCCCGATTCTCCAGCAGTGGCTATGGCACAGATTTTGTTTTTACAATTGAAAACACGCTCTCAGAAGATGTTGCAGATTACTACTGTTTGCAAAGTGATAACATGCCTCT |
| IGKV3-10*01_DBA2 | DBA/2J | Musmus IGKV3-10*01 F | 100 | 299 | AACATTGTGCTGACCCAATCTCCAGCTTCTTTGGCTGTGTCTCTAGGGCAGAGGGCCACCATATCCTGCAGAGCCAGTGAAAGTGTTGATAGTTATGGCAATAGTTTTATGCACTGGTACCAGCAGAAACCAGGACAGCCACCCAAACTCCTCATCTATCTTGCATCCAACCTAGAATCTGGGGTCCCTGCCAGGTTCAGTGGCAGTGGGTCTAGGACAGACTTCACCCTCACCATTGATCCTGTGGAGGCTGATGATGCTGCAACCTATTACTGTCAGCAAAATAATGAGGATCCTCC |
| IGKV6-25*01_DBA2 | DBA/2J | Musmus IGKV6-25*01 F | 100 | 287 | GACATTGTGATGACCCAGTCTCACAAATTCATGTCCACATCAGTAGGAGACAGGGTCAGCATCACCTGCAAGGCCAGTCAGGATGTGAGTACTGCTGTAGCCTGGTATCAACAAAAACCAGGGCAATCTCCTAAACTACTGATTTACTGGGCATCCACCCGGCACACTGGAGTCCCTGATCGCTTCACAGGCAGTGGATCTGGGACAGATTATACTCTCACCATCAGCAGTGTGCAGGCTGAAGACCTGGCACTTTATTACTGTCAGCAACATTATAGCACTCCTCC |
| IGKV1-122*01_DBA2 | DBA/2J | Musmus IGKV1-122*01 F | 100 | 302 | GATGCTGTGATGACCCAAACTCCACTCTCCCTGCCTGTCAGTCTTGGAGATCAAGCCTCCATCTCTTGCAGGTCTAGTCAGAGCCTTGAAAACAGTAATGGAAACACCTATTTGAACTGGTACCTCCAGAAACCAGGCCAGTCTCCACAGCTCCTGATCTACAGGGTTTCCAACCGATTTTCTGGGGTCCTAGACAGGTTCAGTGGTAGTGGATCAGGGACAGATTTCACACTGAAAATCAGCAGAGTGGAGGCTGAGGATTTGGGAGTTTATTTCTGCCTCCAAGTTACACATGTCCCTCC |

| | | | | | |
|---|---|---|---|---|---|
| IGKV4-72*01_DBA2 | DBA/2J | Musmus IGKV4-72*01 F | 100 | 286 | CAAATTGTTCTCTCCCAGTCTCCAGCAATCCTGTCTGCATCTCCAGGGGAGAAGGTCACAATGACTTGCAGGGCCAGCTCAAGTGTAAGTTACATGCACTGGTACCAGCAGAAGCCAGGATCCTCCCCCAAACCCTGGATTTATGCCACATCCAACCTGGCTTCTGGAGTCCCTGCTCGCTTCAGTGGCAGTGGGTCTGGGACCTCTTACTCTCTCACAATCAGCAGAGTGGAGGCTGAAGATGCTGCCACTTATTACTGCCAGCAGTGGAGTAGTAACCCACCCA |
| IGKV10-95*01_DBA2 | DBA/2J | Musmus IGKV10-95*01 F | 100 | 287 | GATATCCAGATGACACAGACTACTTCCTCCCTGTCTGCCTCTCTGGGAGACAGAGTCACCATCAGTTGCAGGGCAAGTGAGGACATTAGCACTTATTTAAACTGGTATCAGCAGAAACCAGATGGAACTGTTAAACTCCTGATCTATTACACATCAGGATTACACTCAGGAGTCCCATCAAGGTTCAGTGGCAGTGGGTCTGGGGCAGATTATTCTCTCACCATCAGCAACCTGGAACCTGAAGATATTGCCACTTACTATTGTCAGCAGTATAGTAAGCTTCCTCC |
| IGKV13-84*01_DBA2 | DBA/2J | Musmus IGKV13-84*01 F | 100 | 287 | GACATCCAGATGACACAATCTTCATCCTCCTTTTCTGTATCTCTAGGAGACAGAGTCACCATTACTTGCAAGGCAAGTGAGGACATATATAATCGGTTAGCCTGGTATCAGCAGAAACCAGGAAATGCTCCTAGGCTCTTAATATCTGGTGCAACCAGTTTGGAAACTGGGGTTCCTTCAAGATTCAGTGGCAGTGGATCTGGAAAGGATTACACTCTCAGCATTACCAGTCTTCAGACTGAAGATGTTGCTACTTATTACTGTCAACAGTATTGGAGTACTCCTCC |
| IGKV11-125*01_DBA2 | DBA/2J | Musmus IGKV11-125*01 F | 100 | 287 | GATGTCCAGATGATTCAGTCTCCATCCTCCCTGTCTGCATCTTTGGGAGACATAGTCACCATGACTTGCCAGGCAAGTCAGGGCACTAGCATTAATTTAAACTGGTTTCAGCAAAAACCAGGGAAAGCTCCTAAGCTCCTGATCTATGGTGCAAGCAACTTGGAAGATGGGGTCCCATCAAGGTTCAGTGGCAGTAGATATGGGACAGATTTCACTCTCACCATCAGCAGCCTGGAGGATGAAGATATGGCAACTTATTTCTGTCTACAGCATAGTTATCTCCCTCC |
| IGKV14-111*01_DBA2 | DBA/2J | Musmus IGKV14-111*01 F | 100 | 287 | GACATCAAGATGACCCAGTCTCCATCTTCCATGTATGCATCTCTAGGAGAGAGAGTCACTATCACTTGCAAGGCGAGTCAGGACATTAATAGCTATTTAAGCTGGTTCCAGCAGAAACCAGGGAAATCTCCTAAGACCCTGATCTATCGTGCAAACAGATTGGTAGATGGGGTCCCATCAAGGTTCAGTGGCAGTGGATCTGGGCAAGATTATTCTCTCACCATCAGCAGCCTGGAGTATGAAGATATGGGAATTTATTATTGTCTACAGTATGATGAGTTTCCTCC |
| IGKV4-53*01_DBA2 | DBA/2J | Musmus IGKV4-53*01 F | 100 | 292 | GAAATTGTGCTCACCCAGTCTCCAGCACTCATGGCTGCATCTCCAGGGGAGAAGGTCACCATCACCTGCAGTGTCAGCTCAAGTATAAGTTCCAGCAACTTGCACTGGTACCAGCAGAAGTCAGAAACCTCCCCCAAACCCTGGATTTATGGCACATCCAACCTGGCTTCTGGAGTCCCTGTTCGCTTCAGTGGCAGTGGATCTGGGACCTCTTATTCTCTCACAATCAGCAGCATGGAGGCTGAAGATGCTGCCACTTATTACTGTCAACAGTGGAGTAGTTACCCACTCA |
| IGKV5-48*01_DBA2 | DBA/2J | Musmus IGKV5-48*01 F | 100 | 287 | GACATCTTGCTGACTCAGTCTCCAGCCATCCTGTCTGTGAGTCCAGGAGAAAGAGTCAGTTTCTCCTGCAGGGCCAGTCAGAGCATTGGCACAAGCATACACTGGTATCAGCAAAGAACAAATGGTTCTCCAAGGCTTCTCATAAAGTATGCTTCTGAGTCTATCTCTGGGATCCCTTCCAGGTTTAGTGGCAGTGGATCAGGGACAGATTTTACTCTTAGCATCAACAGTGTGGAGTCTGAAGATATTGCAGATTATTACTGTCAACAAAGTAATAGCTGGCCAAC |
| IGKV2-109*01_DBA2 | DBA/2J | Musmus IGKV2-109*01 F | 100 | 302 | GATATTGTGATGACGCAGGCTGCATTCTCCAATCCAGTCACTCTTGGAACATCAGCTTCCATCTCCTGCAGGTCTAGTAAGAGTCTCCTACATAGTAATGGCATCACTTATTTGTATTGGTATCTGCAGAAGCCAGGCCAGTCTCCTCAGCTCCTGATTTATCAGATGTCCAACCTTGCCTCAGGAGTCCCAGACAGGTTCAGTAGCAGTGGGTCAGGAACTGATTTCACACTGAGAATCAGCAGAGTGGAGGCTGAGGATGTGGGTGTTTATTACTGTGCTCAAAATCTAGAACTTCCTCC |

| | | | | | |
|---|---|---|---|---|---|
| IGKV4-91*01_S5997_DBA2 | DBA/2J | Musmus IGKV4-91*01 F | 100 | 292 | GAAATTGTGCTCACCCAGTCTCCAACCACCATGGCTGCATCTCCCGGGGAGAAGATCACTATCACCTGCAGTGCCAGCTCAAGTATAAGTTCCAATTACTTGCATTGGTATCAGCAGAAGCCAGGATTCTCCCCTAAACTCTTGATTTATAGGACATCCAATCTGGCTTCTGGAGTCCCAGCTCGCTTCAGTGGCAGTGGGTCTGGGACCTCTTACTCTCTCACAATTGGCACCATGGAGGCTGAAGATGTTGCCACTTACTACTGCCAGCAGGGTAGTAGTATACCACTCA |
| IGKV14-100*01_DBA2 | DBA/2J | Musmus IGKV14-100*01 F | 100 | 287 | GACATCCTGATGACCCAATCTCCATCCTCCATGTCTGTATCTCTGGGAGACACAGTCAGCATCACTTGCCATGCAAGTCAGGGCATTAGCAGTAATATAGGGTGGTTGCAGCAGAAACCAGGGAAATCATTTAAGGGCCTGATCTATCATGGAACCAACTTGGAAGATGGAGTTCCATCAAGGTTCAGTGGCAGTGGATCTGGAGCAGATTATTCTCTCACCATCAGCAGCCTGGAATCTGAAGATTTTGCAGACTATTACTGTGTACAGTATGCTCAGTTTCCTCC |
| IGKV5-39*01_DBA2 | DBA/2J | Musmus IGKV5-39*01 F | 100 | 287 | GACATTGTGATGACTCAGTCTCCAGCCACCCTGTCTGTGACTCCAGGAGATAGAGTCTCTCTTTCCTGCAGGGCCAGCCAGAGTATTAGCGACTACTTACACTGGTATCAACAAAAATCACATGAGTCTCCAAGGCTTCTCATCAAATATGCTTCCCAATCCATCTCTGGGATCCCCTCCAGGTTCAGTGGCAGTGGATCAGGGTCAGATTTCACTCTCAGTATCAACAGTGTGGAACCTGAAGATGTTGGAGTGTATTACTGTCAAAATGGTCACAGCTTTCCTCC |
| IGKV5-45*01_DBA2 | DBA/2J | Musmus IGKV5-45*01 F | 100 | 287 | GATATTGTGCTAACTCAGTCTCCAGCCACCCTGTCTGTGACTCCAGGAGATAGAGTCAGTCTTTCCTGCAGGGCCAGTCAAAGTATTAGCAACTACCTACACTGGTATCAACAAAAATCACATGAGTCTCCAAGGCTTCTCATCAAGTATGCTTCCCAGTCCATCTCTGGGATCCCCTCCAGGTTCAGTGGCAGTGGATCAGGGACAGATTTCACTCTCAGTATCAACAGTGTGGAGACTGAAGATTTTGGAATGTATTTCTGTCAACAGAGTAACAGCTGGCCTCA |
| IGKV2-112*01_DBA2 | DBA/2J | Musmus IGKV2-112*01 F | 100 | 300 | GATATTGTGATAACCCAGGATGAACTCTCCAATCCTGTCACTTCTGGAGAATCAGTTTCCATCTCCTGCAGGTCTAGTAAGAGTCTCCTATATAAGGATGGGAAGACATACTTGAATTGGTTTCTGCAGAGACCAGGACAATCTCCTCAGCTCCTGATCTATTTGATGTCCACCCGTGCATCAGGAGTCTCAGACCGGTTTAGTGGCAGTGGGTCAGGAACAGATTTCACCCTGGAAATCAGTAGAGTGAAGGCTGAGGATGTGGGTGTGTATTACTGTCAACAACTTGTAGAGTATCCT |
| IGKV4-61*01_DBA2 | DBA/2J | Musmus IGKV4-61*01 F | 100 | 286 | CAAATTGTTCTCACCCAGTCTCCAGCAATCATGTCTGCATCTCCAGGGGAGAAGGTCACCATATCCTGCAGTGCCAGCTCAAGTGTAAGTTACATGTACTGGTACCAGCAGAAGCCAGGATCCTCCCCCAAACCCTGGATTTATCGCACATCCAACCTGGCTTCTGGAGTCCCTGCTCGCTTCAGTGGCAGTGGGTCTGGGACCTCTTACTCTCTCACAATCAGCAGCATGGAGGCTGAAGATGCTGCCACTTATTACTGCCAGCAGTATCATAGTTACCCACCCA |
| IGKV9-123*01_DBA2 | DBA/2J | Musmus IGKV9-123*01 F | 100 | 285 | GACATCCAGATGATTCAGTCTCCATCGTCCATGTTTGCCTCTCTGGGAGACAGAGTCAGTCTCTCTTGTCGGGCTAGTCAGGGCATTAGAGGTAATTTAGACTGGTATCAGCAGAAACCAGGTGGAACTATTAAACTCCTGATCTACTCCACATCCAATTTAAATTCTGGTGTCCCATCAAGGTTCAGTGGCAGTGGGTCTGGGTCAGATTATTCTCTCACCATCAGCAGCCTAGAGTCTGAAGATTTTGCAGACTATTACTGTCTACAGCGTAATGCGTATCCT |
| IGKV1-133*01_DBA2 | DBA/2J | Musmus IGKV1-133*01 F | 100 | 302 | GATGTTGTGATGACCCAGACTCCACTCACTTTGTCGGTTACCATTGGACAACCAGCCTCTATCTCTTGCAAGTCAAGTCAGAGCCTCTTATATAGTAATGGAAAAACCTATTTGAATTGGTTATTACAGAGGCCAGGCCAGTCTCCAAAGCGCCTAATCTATCTGGTGTCTAAACTGGACTCTGGAGTCCCTGACAGGTTCACTGGCAGTGGATCAGGAACAGATTTTACACTGAAAATCAGCAGAGTGGAGGCTGAGGATTTGGGAGTTTATTACTGCGTGCAAGGTACACATTTTCCTCA |

| | | | | | |
|---|---|---|---|---|---|
| IGKV4-86*01_DBA2 | DBA/2J | Musmus IGKV4-86*01 F | 100 | 283 | GAAATTGTGCTCACTCAGTCTCCAGCCATCACAGCTGCATCTCTGGGGCAAAAGGTCACCATCACCTGCAGTGCCAGCTCAAGTGTAAGTTACATGCACTGGTACCAGCAGAAGTCAGGCACCTCCCCCAAACCATGGATTTATGAAATATCCAAACTGGCTTCTGGAGTCCCAGCTCGCTTCAGTGGCAGTGGGTCTGGGACCTCTTACTCTCTCACAATCAGCAGCATGGAGGCTGAAGATGCTGCCATTTATTACTGCCAGCAGTGGAATTATCCTCTTA |
| IGKV9-124*01_DBA2 | DBA/2J | Musmus IGKV9-124*01 F | 100 | 285 | GACATCCAGATGACCCAGTCTCCATCCTCCTTATCTGCCTCTCTGGGAGAAAGAGTCAGTCTCACTTGTCGGGCAAGTCAGGAAATTAGTGGTTACTTAAGCTGGCTTCAGCAGAAACCAGATGGAACTATTAAACGCCTGATCTACGCCGCATCCACTTTAGATTCTGGTGTCCCAAAAAGGTTCAGTGGCAGTAGGTCTGGGTCAGATTATTCTCTCACCATCAGCAGCCTTGAGTCTGAAGATTTTGCAGACTATTACTGTCTACAATATGCTAGTTATCCT |
| IGKV14-130*01_DBA2 | DBA/2J | Musmus IGKV14-130*01 F | 100 | 287 | GAAATCCAGATGACCCAGTCTCCATCCTCTATGTCTGCATCTCTGGGAGACAGAATAACCATCACTTGCCAGGCAACTCAAGACATTGTTAAGAATTTAAACTGGTATCAGCAGAAACCAGGGAAACCCCCTTCATTCCTGATCTATTATGCAACTGAACTGGCAGAAGGGGTCCCATCAAGGTTCAGTGGCAGTGGGTCTGGGTCAGACTATTCTCTGACAATCAGCAACCTGGAGTCTGAAGATTTTGCAGACTATTACTGTCTACAGTTTTATGAGTTTCCTCC |
| IGKV4-63*01_DBA2 | DBA/2J | Musmus IGKV4-63*01 F | 100 | 286 | GAAAATGTTCTCACCCAGTCTCCAGCAATCATGTCTGCATCTCCAGGGGAAAAGGTCACCATGACCTGCAGTGCCAGCTCAAGTGTAAGTTACATGCACTGGTACCAGCAGAAGTCAAGCACCTCCCCCAAACTCTGGATTTATGACACATCCAAACTGGCTTCTGGAGTCCCAGGTCGCTTCAGTGGCAGTGGGTCTGGAAACTCTTACTCTCTCACGATCAGCAGCATGGAGGCTGAAGATGTTGCCACTTATTACTGTTTTCAGGGGAGTGGGTACCCACTCA |
| IGKV4-50*01_S0901_DBA2 | DBA/2J | Musmus IGKV4-50*01 F | 100 | 286 | GAAAATGTGCTCACCCAGTCTCCAGCAATCATGTCTGCATCTCTAGGGGAGAAGGTCACCATGAGCTGCAGGGCCAGCTCAAGTGTAAATTACATGTACTGGTACCAGCAGAAGTCAGATGCCTCCCCCAAACTATGGATTTATTACACATCCAACCTGGCTCCTGGAGTCCCAGCTCGCTTCAGTGGCAGTGGGTCTGGGAACTCTTATTCTCTCACAATCAGCAGCATGGAGGGTGAAGATGCTGCCACTTATTACTGCCAGCAGTTTACTAGTTCCCCATACA |
| IGKV8-18*01_DBA2 | DBA/2J | Musmus IGKV8-18*01 ORF | 100 | 310 | GACATTGTGATGACCCAGTCTCCATCCTCCCTGGCTGTGACAGCAGGAGAGAAGGTCACTATGAGATGCAAGTCCAGTCAGAGTCTTTTGTGGAGTGTAAACCAAAATAACTACTTATCCTGGTACCAGCAGAAACAAGGGCAGCCTCCTAAACTGCTTATCTATGGGGCATCCATTAGAGAATCTTGGGTCCCTGATCGATTCACAGGAAGTGGATCTGGGACAGACTTCACTCTCACCATTAGCAATGTGCATGCTGAAGACCTAGCAGTTTATTACTGTCAGCACAATCATGGCAGCTTTCTCCCCC |
| IGKV4-69*01_DBA2 | DBA/2J | Musmus IGKV4-69*01 F | 100 | 282 | CAAATTCTTCTCACCCAGTCTCCAGCAATCATGTCTGCATCTCCAGGGGAGAAGGTCACCATGACCTGCAGTGCCAGCTCAAGTGTAAGTTACATGCACTGGTACCAGCAGAAGCCAGGATCCTCGCCCAAACCCTGGATTTATGACACATCCAACCTGGCTTCTGGATTCCCTGCTCGCTTCAGTGGCAGTGGGTCTGGGACCTCTTACTCTCTCATAATCAGCAGCATGGAGGCTGAAGATGCTGCCACTTATTACTGCCATCAGCGGAGTAGTTACCCA |
| IGKV4-79*01_DBA2 | DBA/2J | Musmus IGKV4-79*01 F | 100 | 292 | CAAATTGTTCTCACCCAGTCTCCAGCAATCATGTCTGCATCTCCTGGGGAGAAGGTCACCTTGACCTGCAGTGCCAGCTCAAGTGTAAGTTCCAGCTACTTGTACTGGTACCAGCAGAAGCCAGGATCCTCCCCCAAACTCTGGATTTATAGCACATCCAACCTGGCTTCTGGAGTCCCTGCTCGCTTCAGTGGCAGTGGGTCTGGGACCTCTTACTCTCTCACAATCAGCAGCATGGAGGCTGAAGATGCTGCCTCTTATTTCTGCCATCAGTGGAGTAGTTACCCACCCC |

| | | | | | |
|---|---|---|---|---|---|
| IGKV8-34*01_DBA2 | DBA/2J | Musmus IGKV8-34*01 F | 100 | 305 | GACATTTTGATGACCCAGTCTCCATCCTCCCTGACTGTGTCAGCAGGAGAGAAGGTCACTATGAGCTGCAAGTCCAGTCAGAGTCTTTTAGCTAGTGGCAACCAAAATAACTACTTGGCCTGGCACCAGCAGAAACCAGGACGATCTCCTAAAATGCTGATAATTTGGGCATCCACTAGGGTATCTGGAGTCCCTGATCGCTTCATAGGCAGTGGATCTGGGACGGATTTCACTCTGACCATCAACAGTGTGCAGGCTGAAGATCTGGCTGTTTATTACTGTCAGCAGTCCTACAGCGCTCCTAC |
| IGKV4-92*01_DBA2 | DBA/2J | Musmus IGKV4-92*01 F | 100 | 291 | GAAATGGTTCTCACCCAGTCTCCAGTATCCATAACTGCATCTCGAGGGGAGAAGGTCACCATCACCTGCCGTGCCAGCTCAAGTATAAGTTCCAATTACTTACACTGGTACCAGCAGAAGCCAGGATCCTCCCCTAAACTTTTGATTTATAGGACATCCATCCTGGCATCTGGAGTCCTAGACAGCTTCAGTGGCAGTGGGTCTGAGAGCTCTTACACTCTGACAATCAGCTGCATGCAGGACGAAGTTGCTGCCACTTACTATTGTCAGCAGGGGAGTAGTAGCCCACCA |
| IGKV12-98*01_DBA2 | DBA/2J | Musmus IGKV12-98*01 F | 100 | 287 | GACATTCAGATGACCCAGTCTCCTGCCTCCCAGTCTGCATCTCTGGGAGAAAGTGTCACCATCACATGCCTGGCAAGTCAGACCATTGGTACATGGTTAGCATGGTATCAGCAGAAACCAGGGAAATCTCCTCAGCTCCTGATTTATGCTGCAACCAGCTTGGCAGATGGGGTCCCATCAAGGTTCAGTGGTAGTGGATCTGGCACAAAATTTTCTTTCAAGATCAGCAGCCTACAGGCTGAAGATTTTGTAAGTTATTACTGTCAACAACTTTACAGTACTCCTCT |
| IGKV3-1*01_DBA2 | DBA/2J | Musmus IGKV3-1*01 F | 100 | 299 | GACATTGTGCTCACCCAATCTCCAGCTTCTTTGGCTGTGTCTCTAGGGCAGAGAGCCACCATCTCCTGCAGAGCCAGTGAAAGTGTTGAATATTATGGCACAAGTTTAATGCAGTGGTACCAACAGAAACCAGGACAGCCACCCAAACTCCTCATCTATGCTGCATCCAACGTAGAATCTGGGGTCCCTGCCAGGTTTAGTGGCAGTGGGTCTGGGACAGACTTCAGCCTCAACATCCATCCTGTGGAGGAGGATGATATTGCAATGTATTTCTGTCAGCAAAGTAGGAAGGTTCCTTC |
| IGKV8-23-1*01_DBA2 | DBA/2J | Musmus IGKV8-16*01 F, or Musmus IGKV8-23-1*01 ORF | 100 | 284 | GAAATTGTGCTGACTGTGTCAGCAGGAGAGAGGGTCACTATCAGCTGCAAATCCAATCAGAATCTTTTATGGAGTGGAAACCAAAGGTACTGTTTGGTCTGGCACCAGTGGAAACCAGGGCAAACTCCTACACCGTTGATCACCTGGACATCTGATAGGTACTCTGGAGTCCCTGATCGTTTCATAGGCAGTGGATCTGTGACAGATTTCACTCTGACCATCAGCAGTGTGCAGGCTGAAGATGTGGCAGTTTATTTCTGTCAGCAGCATTTACACATTCCTCC |
| IGKV12-38*01_DBA2 | DBA/2J | Musmus IGKV12-38*01 F | 100 | 287 | GACATCCAGATGACTCAGTCTCCAGCCTCCCTGGCTGCATCTGTGGGAGAAACTGTCACCATCACATGTCGAGCAAGTGAGAACATTTACTACAGTTTAGCATGGTATCAGCAGAAGCAAGGGAAATCTCCTCAGCTCCTGATCTATAATGCAAACAGCTTGGAAGATGGTGTCCCATCGAGGTTCAGTGGCAGTGGATCTGGGACACAGTATTCTATGAAGATCAACAGCATGCAGCCTGAAGATACCGCAACTTATTTCTGTAAACAGGCTTATGACGTTCCTCC |
| IGKV1-132*01_DBA2 | DBA/2J | Musmus IGKV1-132*01 F | 100 | 302 | GATGTTGTGATGACCCAGACTCCACTGTCTTTGTCGGTTACCATTGGACAACCAGCCTCTATCTCTTGCAAGTCAAGTCAGAGCCTCTTATATAGTAATGGAAAGACATATTTGAATTGGTTACAACAGAGGCCTGGCCAGGCTCCAAAGCACCTAATGTATCAGGTGTCCAAACTGGACCCTGGCATCCCTGACAGGTTCAGTGGCAGTGGATCAGAAACAGATTTTACACTTAAAATCAGCAGAGTGGAGGCTGAAGATTTGGGAGTTTATTACTGCTTGCAAGGTACATATTATCCTCA |
| IGKV9-129*01_DBA2 | DBA/2J | Musmus IGKV9-129*01 ORF | 100 | 287 | GACATCCAGATGACCCAGTCTCCATCCTCCTTATCTGCCTCTCTGGGAGAAAGAGTCAGTCTCACTTGCCGGGCAAGTCAGGACATTCATGGTTATTTAAACTTGTTTCAGCAGAAACCAGGTGAAACTATTAAACACCTGATCTATGAAACATCCAATTTAGATTCTGGTGTCCCAAAAAGGTTCAGTGGCAGTAGGTCTGGGTCAGATTATTCTCTCATTATCGGCAGCCTTGAGTCTGAAGATTTTGCAGACTATTACTGTCTACAATATGCTAGTTCTCCTCC |

| | | | | | |
|---|---|---|---|---|---|
| IGKV1-99*01_S1751_DBA2 | DBA/2J | Musmus IGKV1-99*01 F | 100 | 302 | GATGTTGTTCTGACCCAAACTCCACTCTCTCTGCCTGTCAATATTGGAGATCAAGCCTCTATCTCTTGCAAGTCTACTAAGAGTCTTCTGAATAGTGATGGATTCACTTATTTGGACTGGTACCTGCAGAAGCCAGGCCAGTCTCCACAGCTCCTAATATATTTGGTTTCTAATCGATTTTCTGGAGTTCCAGACAGGTTCAGTGGCAGTGGGTCAGGAACAGATTTCACACTCAAGATCAGCAGAGTGGAGGCTGAGGATTTGGGAGTTTATTATTGCTTCCAGAGTAACTATCTTCCGCT |
| IGKV4-58*01_DBA2 | DBA/2J | Musmus IGKV4-58*01 F | 100 | 292 | GAAAATGTGCTCACCCAGTCTCCAGCAATAATGGCTGCCTCTCTGGGGCAGAAGGTCACCATGACCTGCAGTGCCAGCTCAAGTGTAAGTTCCAGTTACTTGCACTGGTACCAGCAGAAGTCAGGCGCTTCCCCCAAACCCTTGATTCATAGGACATCCAACCTGGCTTCTGGAGTCCCAGCTCGCTTCAGTGGCAGTGGGTCTGGGACCTCTTACTCTCTCACAATCAGCAGCGTGGAGGCTGAAGATGATGCAACTTATTACTGCCAGCAGTGGAGTGGTTACCCATTCA |
| IGKV5-37*01_DBA2 | DBA/2J | Musmus IGKV5-37*01 F | 100 | 287 | GACATCCTGCTGACCCAGTCTCCAGCCACCCTGTCTGTGACTCCAGGAGAAACAGTCAGTCTTTCCTGTAGGGCCAGCCAGAGTATTTACAAGAACCTACACTGGTATCAACAGAAATCACATCGGTCTCCAAGGCTTCTCATCAAGTATGCTTCTGATTCCATCTCTGGGATCCCCTCCAGGTTCACTGGCAGTGGATCAGGGACAGATTACACTCTCAGTATCAACAGTGTGAAGCCCGAAGATGAAGGAATATATTACTGTCTTCAAGGTTACAGCACACCTTC |
| IGKV3-3*01_DBA2 | DBA/2J | Musmus IGKV3-3*01 F | 100 | 299 | GACATTGTGCTGACCCAATCTCCAGCTTCTTTGGCTGTGTCTCTAGGACAGAGAGCCACTATCTTCTGCAGAGCCAGCCAGAGTGTCGATTATAATGGAATTAGTTATATGCACTGGTTCCAACAGAAACCAGGACAGCCACCCAAACTCCTCATCTATGCTGCATCCAACCTAGAATCTGGGATCCCTGCCAGGTTCAGTGGCAGTGGGTCTGGGACAGACTTCACCCTCAACATCCATCCTGTGGAGGAGGAAGATGCTGCAACCTATTACTGTCAGCAAAGTATTGAGGATCCTCC |
| IGKV4-51*01_S4777_DBA2 | DBA/2J | Musmus IGKV4-51*01 F | 100 | 292 | GAAAATGTGCTCACCCAGTCTCCAGCAATAATGGCTGCCTCTCTGGGGGAGAAGGTCACCATGACCTGCAGTGCCAGCTCAAGTGTAAGTTCCAGCTACTTGCACTGGTACCAGCAGAAGTCAGGCACTTCCCCCAAACTCTGGATTTATGGCACATCCAACCTGGCTTCTGGAGTCCCAGCTCGCTTCAGTGGCAGTGGGGCTGGGATCTCTTACTCTCTCACAATCAGCAGCATGGAGGCTGAAAATGATGCAACTTATTACTGCCAGCAGTGGAGTGGTTACCCATACA |
| IGKV4-52*01_DBA2 | DBA/2J | Musmus IGKV4-52*01 ORF, or Musmus IGKV4-54*01 ORF | 100 | 286 | CAAATTGTTCTCACCCAGTCTCCAGCAATCCTGTCTGCATCTCCAGGGGAGAAGGTCACCATGACCTGCAGTGCCAGCTCAAGTGTAAGTTACATGTACAGGTACCAGCAGAAGCCAGGATCCTCACCCAAACCCTGGATTTATGGCACATCCAACCTGGCTTCTGGAGTCCCTGCTCGCTTCAGTGGCAGTGGATCTGGGACCTCTTATTCTCTCACAATCAGCAGCATGGAGGCTGAAGATGCTGCCACTTATTACTGCCAGCAATATCATAGTTACCCACCCA |
| IGKV4-90*01_DBA2 | DBA/2J | Musmus IGKV4-90*01 F | 100 | 285 | GAAATTTTGCTCACCCAGTCTCCAGCAATCATAGCTGCATCTCCTGGGGAGAAGGTCACCATCACCTGCAGTGCCAGCTCAAGTGTAAGTTACATGAACTGGTACCAGCAGAAACCAGGATCCTCCCCCAAAATATGGATTTATGGTATATCCAACCTGGCTTCTGGAGTTCCTGCTCGCTTCAGTGGCAGTGGGTCTGGGACATCTTTCTCTTTCACAATCAACAGCATGGAGGCTGAAGATGTTGCCACTTATTACTGTCAGCAAAGGAGTAGTTACCCACCC |
| IGKV6-29*01_DBA2 | DBA/2J | Musmus IGKV6-29*01 F | 100 | 287 | AACATTGTAATGACCCAATCTCCCAAATCCATGTCCATGTCAGTAGGAGAGAGGGTCACCTTGAGCTGCAAGGCCAGTGAGAATGTGGGTACTTATGTATCCTGGTATCAACAGAAACCAGAGCAGTCTCCTAAACTGCTGATATACGGGGCATCCAACCGGTACCCTGGGGTCCCTGATCGCTTCACAGGCAGTGGATCTGCAACAGATTTCACTCTGACCATCAGCAGTCTGCAGGCTGAAGACCTTGCAGATTATCACTGTGGACAGGGTTACAGCTATCTTCC |

| | | | | | |
|---|---|---|---|---|---|
| IGKV3-9*01_DBA2 | DBA/2J | Musmus IGKV3-9*01 F | 100 | 299 | GACATTGTGCTGACCCAATCTCCAGCTTCTTTGGCTGTGTCTCTAGGACAGAGGGCCACCATATCCTGCCAAGCCAGCGAAAGTGTCAGTTTTGCTGGTACAAGTTTAATGCACTGGTACCAACAGAAACCAGGACAGCCACCCAAACTCCTCATCTATCGTGCATCCAACCTAGAATCTGGAGTCCCTGCCAGGTTCAGTGGCAGTGGGTCTGAGTCAGACTTCACTCTCACCATCGATCCTGTGGAGGAAGATGATGCTGCAATGTATTACTGTATGCAAAGTATGGAAGATCCTCC |
| IGKV4-56*01_DBA2 | DBA/2J | Musmus IGKV4-56*01 P | 100 | 286 | CAAATTGTTCTCACCCAGTCTCCAGCAATCATGTCTGCATCTCCAGGGCAGAAAGTCACCATAACCTGCAGTGCCATCTCAAGTGTAAATTACATGCACTGGTACCAGCAGAAGCCAGGATCCTCCCCCAAACTCTGGATTTATGCAACATCCAAACTGGCTCTTGGAGTCCCTGCTTGCTTCAGTGGCAGTGGGTCTGGGACCTCTTACTCTCTCACAATCAGCAGCATGGTGGCTGAAGATGCCACCTCTTATTTCTGTCATCAGTGGAGTAGTTACCCACCCA |
| IGKV4-60*01_DBA2 | DBA/2J | Musmus IGKV4-60*01 P | 100 | 292 | GAAATTGTGCTCACCCAGTCTCCAGCACTCATGGCTGCATCTCCAGGGGAGAAGGTCACCATCACCTGCAGTGTCAGCTCAAGTATAAGTTCCAGCTACTTGCACTGGTACCAGCAGAAGTCAGGAATCTCCCCCAAACCCTGGATTTATGGCACATCCAACCTGGCTTCTGGAGTCCCTGCTCGCTTCAGTGGCAGTGGATCTGGGACCTCTTACTCTCTCACAATCAGCAGCATGGAGGCTGAAGATGCTGCCACTTATTACTGTCAACAGTGGAGTAGTTCCCCACCCA |
| IGKV4-78*01_DBA2 | DBA/2J | Musmus IGKV4-78*01 F | 100 | 292 | CAAATTGTTCTCACCCAGTCTCCAGCAATCATGTCTGCATCTCCTGGGGAGAAGGTCACCATGACCTGCAGTGCCAGATCAAGTGTAAGTTCCAGCTACTTGTACTGGTACCAGCAGAAGCCAGGATCCTCCCCCAAACTCTGGATTTATAGCACATCCAACCTGGCTTCTGGAGTCCCTGCTCGCTTCAGTGGCAGTGGGTCTGGGACCTCTTATTCTCTCACAATCAGCAGCATGGAGGCTGAAGATGCTGCCACTTTTTACTGCCAGCAGTACAGTGGTTACCCATCCA |
| IGKV4-81*01_DBA2 | DBA/2J | Musmus IGKV4-81*01 F | 100 | 292 | GAAAATGTGCTGACCCAGTCTCCAGCAATCATGGCTGCATCTCCAGGGGAGAAGGTCACCATGACCTGCAGTGCCAGCTCAAGTGTAAGTTCTAGTAACTTGCACTGGTACCAGCAGAAGTCAGGCACTTCTACCAAATTCTGGATTTATAGGACATCCAACCTGGCTTCAGAAGTCCCAGCTCCCTTCAGTGGCAGTGGGTCTGGGACCTCTTACTCTCTTACAATCAGCAGCGTGGAGGCCGAAGATGCTGCCACTTATTACTGCCAGCAGTGGAGTGGTTACCCACCCA |
| IGKV6-14*01_DBA2 | DBA/2J | Musmus IGKV6-14*01 F | 100 | 287 | GACATTGTGATGACCCAGTCTCAAAAATTCATGTCCACATCAGTAGGAGACAGGGTCAGCATCACCTGCAAGGCCAGTCAGAATGTTCGTACTGCTGTAGCCTGGTATCAACAGAAACCAGGGCAGTCTCCTAAAGCACTGATTTACTTGGCATCCAACCGGCACACTGGAGTCCCTGATCGCTTCACAGGCAGTGGATCTGGGACAGATTTCACTCTCACCATTAGCAATGTGCAATCTGAAGACCTGGCAGATTATTTCTGTCTGCAACATTGGAATTATCCTCT |
| IGKV8-26*01_DBA2 | DBA/2J | Musmus IGKV8-26*01 ORF | 100 | 311 | GACATTGTGATGACCCAGTCTCCATCCTCCCTGGCTGTGACAGCAGGAGAGAAGGTCACTATGAGCTGCAAGTCCAGTCAGAGTCTTTTGTGGAGTGTAAACCAAAAGAACTACTTGTCCTGGTACCAGCAGAAACAAAGGCAGCCTCCTAAACTGCTTATCTATGGGGCATCCATTAGAGAATCTTGGGTCCCTGATCGGTTCACAGGAAGTGGATCTGGGACAGACTTCACTCTCACCATTAGCAATGTGCATGCTGAAGACCTAGCAGTTTATTACTGTCAACACAATCATGGCAGCTTTCTCCCCCC |

| | | | | | |
|---|---|---|---|---|---|
| IGKV18-36*01_DBA2 | DBA/2J | Musmus IGKV18-36*01 F | 100 | 287 | ACTGGAGAAACAACACAGGCTCCAGCTTCTCTGAGTTTTTCTCTTGGTGAAACAGCAACACTGTCATGCAGGTCCAGTGAGAGTGTTGGCAGCTACTTAGCCTGGTACCAGCAGAAAGCAGAGCAAGTTCCCCGGCTCCTTATCCATAGTGCCTCCACTAGGGCCGGTGGTGTCCCAGTCCGGTTCAGTGGCACTGGGTCTGGGACAGACTTCACTCTCACCATCAGCAGTCTAGAACCTGAAGATGCTGCAGTTTACTACTGTCAACCTTTCAAAAGTTGGTCATA |
| IGKV4-52*01_S5019_DBA2 | DBA/2J | Musmus IGKV4-52*01 ORF, or Musmus IGKV4-54*01 ORF | 100 | 286 | CAAATTGTTCTCACCCAGTCTCCAGCAATCCTGTCTGCATCTCCAGGGGAGAAGGTCACCATGACCTGCAGTGCCAGCTCAAGTGTAAGTTACATGTACAGGTACCAGCAGAAGCCAGGATCCTCACCCAAACCCTGGATTTATGGCACATCCAACCTGGCTTCTGGAGTCCCTGCTCGCTTCAGTGGCAGTGGATCTGGGACCTCTTATTCTCTCACAATCAGCAGCATGGAGGCTGAAGATGCTGCCACTTATTACTGCCAGCAATATCATAGTTACCCACTCA |
| IGKV4-56*01_S5954_DBA2 | DBA/2J | Musmus IGKV4-56*01 P | 100 | 286 | CAAATTGTTCTCACCCAGTCTCCAGCAATCATGTCTGCATCTCCAGGGCAGAAAGTCACCATAACCTGCAGTGCCATCTCAAGTGTAAATTACATGCACTGGTACCAGCAGAAGCCAGGATCCTCCCCCAAACTCTGGATTTATGCAACATCCAAACTGGCTCTTGGAGTCCCTGCTTGCTTCAGTGGCAGTGGGTCTGGGACCTCTTACTCTCTCACAATCAGCAGCATGGTGGCTGAAGATGCCACCTCTTATTTCTGTCATCAGTGGAGTAGTTACCCACTCA |
| IGKV8-27*01_LEWES | LEWES/EiJ | Musmus IGKV8-27*01 F | 100 | 300 | AACATTATGATGACACAGTCGCCATCATCTCTGGCTGTGTCTGCAGGAGAAAAGGTCACTATGAGCTGTAAGTCCAGTCAAAGTGTTTTATACAGTTCAAATCAGAAGAACTACTTGGCCTGGTACCAGCAGAAACCAGGGCAGTCTCCTAAACTGCTGATCTACTGGGCATCCACTAGGGAATCTGGTGTCCCTGATCGCTTCACAGGCAGTGGATCTGGGACAGATTTTACTCTTACCATCAGCAGTGTACAAGCTGAAGACCTGGCAGTTTATTACTGTCATCAATACCTCTCCTCG |
| IGKV8-30*01_LEWES | LEWES/EiJ | Musmus IGKV8-30*01 F | 100 | 305 | GACATTGTGATGTCACAGTCTCCATCCTCCCTAGCTGTGTCAGTTGGAGAGAAGGTTACTATGAGCTGCAAGTCCAGTCAGAGCCTTTTATATAGTAGCAATCAAAAGAACTACTTGGCCTGGTACCAGCAGAAACCAGGGCAGTCTCCTAAACTGCTGATTTACTGGGCATCCACTAGGGAATCTGGGGTCCCTGATCGCTTCACAGGCAGTGGATCTGGGACAGATTTCACTCTCACCATCAGCAGTGTGAAGGCTGAAGACCTGGCAGTTTATTACTGTCAGCAATATTATAGCTATCCTCC |
| IGKV6-15*01_LEWES | LEWES/EiJ | Musmus IGKV6-15*01 F | 100 | 287 | GACATTGTGATGACCCAGTCTCAAAAATTCATGTCCACATCAGTAGGAGACAGGGTCAGCGTCACCTGCAAGGCCAGTCAGAATGTGGGTACTAATGTAGCCTGGTATCAACAGAAACCAGGGCAATCTCCTAAAGCACTGATTTACTCGGCATCCTACCGGTACAGTGGAGTCCCTGATCGCTTCACAGGCAGTGGATCTGGGACAGATTTCACTCTCACCATCAGCAATGTGCAGTCTGAAGACTTGGCAGAGTATTTCTGTCAGCAATATAACAGCTATCCTCT |
| IGKV8-21*01_LEWES | LEWES/EiJ | Musmus IGKV8-21*01 F | 100 | 302 | GACATTGTGATGTCACAGTCTCCATCCTCCCTGGCTGTGTCAGCAGGAGAGAAGGTCACTATGAGCTGCAAATCCAGTCAGAGTCTGCTCAACAGTAGAACCCGAAAGAACTACTTGGCTTGGTACCAGCAGAAACCAGGGCAGTCTCCTAAACTGCTGATCTACTGGGCATCCACTAGGGAATCTGGGGTCCCTGATCGCTTCACAGGCAGTGGATCTGGGACAGATTTCACTCTCACCATCAGCAGTGTGCAGGCTGAAGACCTGGCAGTTTATTACTGCAAGCAATCTTATAATCTTCC |
| IGKV12-44*01_LEWES | LEWES/EiJ | Musmus IGKV12-44*01 F | 100 | 287 | GACATCCAGATGACTCAGTCTCCAGCCTCCCTATCTGCATCTGTGGGAGAAACTGTCACCATCACATGTCGAGCAAGTGAGAATATTTACAGTTATTTAGCATGGTATCAGCAGAAACAGGGAAAATCTCCTCAGCTCCTGGTCTATAATGCAAAAACCTTAGCAGAAGGTGTGCCATCAAGGTTCAGTGGCAGTGGATCAGGCACACAGTTTTCTCTGAAGATCAACAGCCTGCAGCCTGAAGATTTTGGGAGTTATTACTGTCAACATCATTATGGTACTCCTCC |

| | | | | | |
|---|---|---|---|---|---|
| IGKV2-137*01_S4012_LEWES | LEWES/EiJ | Musmus IGKV2-137*01 F | 96.94 | 302 | GATATTGTGATGACTCAGGCTGCACCCTCTGTACCTGTCACTCCTGGAGAGTCAGTATCCATCTCCTGCAGGTCTAGTAAGAGTCTTCTGCATAGTAATGGCAACACTTACTTGTATTGGTTCCTGCAGAGGCCAGGCCAGTCTCCTCAGCGCCTGATATATTATATGTCCAACCTTGCCTCAGGAGTCCCAGACAGGTTCAGTGGCAGAGGGTCAGGAACTGATTTCACACTGAGAATCAGTAGAGTGGAGGCTGAGGATGTGGGTGTTTATTACTGTATGCAAAGTCTAGAATATCCTTT |
| IGKV4-59*01_LEWES | LEWES/EiJ | Musmus IGKV4-59*01 F | 100 | 286 | CAAATTGTTCTCACCCAGTCTCCAGCAATCATGTCTGCATCTCCAGGGGAGAAGGTCACCATGACCTGCAGTGCCAGCTCAAGTGTAAGTTACATGCACTGGTACCAGCAGAAGTCAGGCACCTCCCCCAAAAGATGGATTTATGACACATCCAAACTGGCTTCTGGAGTCCCTGCTCGCTTCAGTGGCAGTGGGTCTGGGACCTCTTACTCTCTCACAATCAGCAGCATGGAGGCTGAAGATGCTGCCACTTATTACTGCCAGCAGTGGAGTAGTAACCCACCCA |
| IGKV12-46*01_LEWES | LEWES/EiJ | Musmus IGKV12-46*01 F | 100 | 287 | GACATCCAGATGACTCAGTCTCCAGCCTCCCTATCTGTATCTGTGGGAGAAACTGTCACCATCACATGTCGAGCAAGTGAGAATATTTACAGTAATTTAGCATGGTATCAGCAGAAACAGGGAAAATCTCCTCAGCTCCTGGTCTATGCTGCAACAAACTTAGCAGATGGTGTGCCATCAAGGTTCAGTGGCAGTGGATCAGGCACACAGTATTCCCTCAAGATCAACAGCCTGCAGTCTGAAGATTTTGGGAGTTATTACTGTCAACATTTTTGGGGTACTCCTCC |
| IGKV10-94*01_LEWES | LEWES/EiJ | Musmus IGKV10-94*01 F | 100 | 287 | GATATCCAGATGACACAGACTACATCCTCCCTGTCTGCCTCTCTGGGAGACAGAGTCACCATCAGTTGCAGTGCAAGTCAGGGCATTAGCAATTATTTAAACTGGTATCAGCAGAAACCAGATGGAACTGTTAAACTCCTGATCTATTACACATCAAGTTTACACTCAGGAGTCCCATCAAGGTTCAGTGGCAGTGGGTCTGGGACAGATTATTCTCTCACCATCAGCAACCTGGAACCTGAAGATATTGCCACTTACTATTGTCAGCAGTATAGTAAGCTTCCTCC |
| IGKV4-55*01_S8319_LEWES | LEWES/EiJ | Musmus IGKV4-55*01 F | 100 | 286 | CAAATTGTTCTCACCCAGTCTCCAGCAATCATGTCTGCATCTCCAGGGGAGAAGGTCACCATGACCTGCAGTGCCAGCTCAAGTGTAAGTTACATGTACTGGTACCAGCAGAAGCCAGGATCCTCCCCCAGACTCCTGATTTATGACACATCCAACCTGGCTTCTGGAGTCCCTGTTCGCTTCAGTGGCAGTGGGTCTGGGACCTCTTACTCTCTCACAATCAGCCGAATGGAGGCTGAAGATGCTGCCACTTATTACTGCCAGCAGTGGAGTAGTTACCCGCTCA |
| IGKV10-96*01_LEWES | LEWES/EiJ | Musmus IGKV10-96*01 F | 100 | 286 | GATATCCAGATGACACAGACTACATCCTCCCTGTCTGCCTCTCTGGGAGACAGAGTCACCATCAGTTGCAGGGCAAGTCAGGACATTAGCAATTATTTAAACTGGTATCAGCAGAAACCAGATGGAACTGTTAAACTCCTGATCTACTACACATCAAGATTACACTCAGGAGTCCCATCAAGGTTCAGTGGCAGTGGGTCTGGAACAGATTATTCTCTCACCATTAGCAACCTGGAGCAAGAAGATATTGCCACTTACTTTTGCCAACAGGGTAATACGCTTCCTC |
| IGKV4-57-1*01_LEWES | LEWES/EiJ | Musmus IGKV4-57*01 F | 100 | 292 | GAAAATGTGCTCACCCAGTCTCCAGCAATCATGTCTGCATCTCCAGGGGAAAAGGTCACCATGACCTGCAGGGCCAGCTCAAGTGTAAGTTCCAGTTACTTGCACTGGTACCAGCAGAAGTCAGGTGCCTCCCCCAAACTCTGGATTTATAGCACATCCAACTTGGCTTCTGGAGTCCCTGCTCGCTTCAGTGGCAGTGGGTCTGGGACCTCTTACTCTCTCACAATCAGCAGTGTGGAGGCTGAAGATGCTGCCACTTATTACTGCCAGCAGTACAGTGGTTACCCACTCA |
| IGKV8-19*01_LEWES | LEWES/EiJ | Musmus IGKV8-19*01 F | 100 | 305 | GACATTGTGATGACACAGTCTCCATCCTCCCTGACTGTGACAGCAGGAGAGAAGGTCACTATGAGCTGCAAGTCCAGTCAGAGTCTGTTAAACAGTGGAAATCAAAAGAACTACTTGACCTGGTACCAGCAGAAACCAGGGCAGCCTCCTAAACTGTTGATCTACTGGGCATCCACTAGGGAATCTGGGGTCCCTGATCGCTTCACAGGCAGTGGATCTGGAACAGATTTCACTCTCACCATCAGCAGTGTGCAGGCTGAAGACCTGGCAGTTTATTACTGTCAGAATGATTATAGTTATCCTCC |

| | | | | | |
|---|---|---|---|---|---|
| IGKV6-13*01_LEWES | LEWES/EiJ | Musmus IGKV6-13*01 F | 100 | 285 | GACATTGTGATGACCCAGTCTCAAAAATTCATGTCCACATCAGTAGGAGACAGGGTCAGCATCACCTGCAAGGCCAGTCAGAATGTGGGTACTGCTGTAGCCTGGTATCAACAGAAACCAGGACAATCTCCTAAACTACTGATTTACTCGGCATCCAATCGGTACACTGGAGTCCCTGATCGCTTCACAGGCAGTGGATCTGGGACAGATTTCACTCTCACCATCAGCAATATGCAGTCTGAAGACCTGGCAGATTATTTCTGCCAGCAATATAGCAGCTATCCT |
| IGKV6-23*01_LEWES | LEWES/EiJ | Musmus IGKV6-23*01 F | 100 | 287 | GACATTGTGATGACCCAGTCTCACAAATTCATGTCCACATCAGTAGGAGACAGGGTCAGCATCACCTGCAAGGCCAGTCAGGATGTGGGTACTGCTGTAGCCTGGTATCAACAGAAACCAGGGCAATCTCCTAAACTACTGATTTACTGGGCATCCACCCGGCACACTGGAGTCCCTGATCGCTTCACAGGCAGTGGATCTGGGACAGATTTCACTCTCACCATTAGCAATGTGCAGTCTGAAGACTTGGCAGATTATTTCTGTCAGCAATATAGCAGCTATCCTCT |
| IGKV9-120*01_S5626_LEWES | LEWES/EiJ | Musmus IGKV9-120*02 F | 97.49 | 285 | GACATCCAGATGACCCAGTCTCCATCCTCCTTATCTGCCTCTCTGGGAGAAAGAGTCAGTCTCACTTGCCGGGCAAGTCAGGACATTGGTGGTAGCTTAAACTGGTTTCAGCAGAAACCAGATGGAACTATTAAACGCCTGATCTACGGCACATCCAGTTTAGATTCTGGTGTCCCCAAAAGGTTCAGTGGCAGTAGGTCTGGGTCAGATTATTCTCTCACCATCAGCAGCCTAGAGTCTGAAGATTTTGCAGACTATTACTGTCTACAATATGCTAGTTCTCCT |
| IGKV4-80*01_LEWES | LEWES/EiJ | Musmus IGKV4-80*01 F | 100 | 282 | CAAATTGTTCTCACCCAGTCTCCAGCAATCATGTCTGCATCTCTAGGGGAGGAGATCACCCTAACCTGCAGTGCCAGCTCGAGTGTAAGTTACATGCACTGGTACCAGCAGAAGTCAGGCACTTCTCCCAAACTCTTGATTTATAGCACATCCAACCTGGCTTCTGGAGTCCCTTCTCGCTTCAGTGGCAGTGGGTCTGGGACCTTTTATTCTCTCACAATCAGCAGTGTGGAGGCTGAAGATGCTGCCGATTATTACTGCCATCAGTGGAGTAGTTATCCA |
| IGKV4-86*01_LEWES | LEWES/EiJ | Musmus IGKV4-86*01 F | 100 | 283 | GAAATTGTGCTCACTCAGTCTCCAGCCATCACAGCTGCATCTCTGGGGCAAAAGGTCACCATCACCTGCAGTGCCAGCTCAAGTGTAAGTTACATGCACTGGTACCAGCAGAAGTCAGGCACCTCCCCCAAACCATGGATTTATGAAATATCCAAACTGGCTTCTGGAGTCCCAGCTCGCTTCAGTGGCAGTGGGTCTGGGACCTCTTACTCTCTCACAATCAGCAGCATGGAGGCTGAAGATGCTGCCATTTATTACTGCCAGCAGTGGAATTATCCTCTTA |
| IGKV1-135*01_S1357_LEWES | LEWES/EiJ | Musmus IGKV1-135*01 F | 95.92 | 302 | GATGTTGTGATGACTCAGACCCCACTCACTTTGTCGGTTACCATTGGACAACCAGCCTCCATCTCTTGCAAGTCAAGTCAGAGCCTCTTATATAGTGATGGAAAGACATATTTGGATTGGTTATTACAGAGGCCAGGTCAGTCTCCAAAGAGCCTAATCTATCTGGTGTCTAAACTGGAATCTGGAGTCCCTGACAGGTTCAGTGGCAGTGGATCAGGGACAGATTTCACACTGAAAATCAGCAGAGTGGAGGCTGAGGATTTGGGAGTTTATTACTGCTTGCAAGCTACACATTTTCCTCA |
| IGKV12-41*01_LEWES | LEWES/EiJ | Musmus IGKV12-41*01 F | 100 | 287 | GACATCCAGATGACTCAGTCTCCAGCCTCCCTATCTGCATCTGTGGGAGAAACTGTCACCATCACATGTCGAGCAAGTGGGAATATTCACAATTATTTAGCATGGTATCAGCAGAAACAGGGAAAATCTCCTCAGCTCCTGGTCTATAATGCAAAAACCTTAGCAGATGGTGTGCCATCAAGGTTCAGTGGCAGTGGATCAGGAACACAATATTCTCTCAAGATCAACAGCCTGCAGCCTGAAGATTTTGGGAGTTATTACTGTCAACATTTTTGGAGTACTCCTCC |
| IGKV6-20*01_LEWES | LEWES/EiJ | Musmus IGKV6-20*01 F | 100 | 287 | AACATTGTAATGACCCAATCTCCCAAATCCATGTCCATGTCAGTAGGAGAGAGGGTCACCTTGAGCTGCAAGGCCAGTGAGAATGTGGGTACTTATGTATCCTGGTATCAACAGAAACCAGAGCAGTCTCCTAAACTGCTGATATACGGGGCATCCAACCGGTACACTGGGGTCCCCGATCGCTTCACAGGCAGTGGATCTGCAACAGATTTCACTCTGACCATCAGCAGTGTGCAGGCTGAAGACCTTGCAGATTATCACTGTGGACAGAGTTACAGCTATCCTCC |

| | | | | | |
|---|---|---|---|---|---|
| IGKV9-124*01_S5726_LEWES | LEWES/EiJ | Musmus IGKV9-124*01 F | 97.49 | 285 | GACATCCAAATGACCCAGTCTCCATCTTCCTTATCTGCCTCTCTGGGAGAAAGAGTCAGTCTCACTTGTCGGGCAAGTCAGGATATTAGTGGTTACTTAAGCTGGCTTCAGCAGAAACCAGATGGAACTATTAAACGCCTGATCTACAGCACATCCACTTTAGATTCTGGTGTCCCAAAAAGGTTCAGTGGCAGTAGGTCTGGGTCAGATTATTCTCTCACCATCAGCAGCCTAGAGTCTGAAGATTTTGCAGACTATTACTGTCTACAATATGCTAGTTCTCCT |
| IGKV5-43*01_LEWES | LEWES/EiJ | Musmus IGKV5-43*01 F | 100 | 287 | GATATTGTGCTAACTCAGTCTCCAGCCACCCTGTCTGTGACTCCAGGAGATAGCGTCAGTCTTTCCTGCAGGGCCAGCCAAAGTATTAGCAACAACCTACACTGGTATCAACAAAAATCACATGAGTCTCCAAGGCTTCTCATCAAGTATGCTTCCCAGTCCATCTCTGGGATCCCCTCCAGGTTCAGTGGCAGTGGATCAGGGACAGATTTCACTCTCAGTATCAACAGTGTGGAGACTGAAGATTTTGGAATGTATTTCTGTCAACAGAGTAACAGCTGGCCTCA |
| IGKV4-91*01_S5997_LEWES | LEWES/EiJ | Musmus IGKV4-91*01 F | 100 | 292 | GAAATTGTGCTCACCCAGTCTCCAACCACCATGGCTGCATCTCCCGGGGAGAAGATCACTATCACCTGCAGTGCCAGCTCAAGTATAAGTTCCAATTACTTGCATTGGTATCAGCAGAAGCCAGGATTCTCCCCTAAACTCTTGATTTATAGGACATCCAATCTGGCTTCTGGAGTCCCAGCTCGCTTCAGTGGCAGTGGGTCTGGGACCTCTTACTCTCTCACAATTGGCACCATGGAGGCTGAAGATGTTGCCACTTACTACTGCCAGCAGGGTAGTAGTATACCACTCA |
| IGKV8-28*01_LEWES | LEWES/EiJ | Musmus IGKV8-28*01 F | 100 | 305 | GACATTGTGATGACACAGTCTCCATCCTCCCTGAGTGTGTCAGCAGGAGAGAAGGTCACTATGAGCTGCAAGTCCAGTCAGAGTCTGTTAAACAGTGGAAATCAAAAGAACTACTTGGCCTGGTACCAGCAGAAACCAGGGCAGCCTCCTAAACTGTTGATCTACGGGGCATCCACTAGGGAATCTGGGGTCCCTGATCGCTTCACAGGCAGTGGATCTGGAACCGATTTCACTCTTACCATCAGCAGTGTGCAGGCTGAAGACCTGGCAGTTTATTACTGTCAGAATGATCATAGTTATCCTCC |
| IGKV1-135*01_LEWES | LEWES/EiJ | Musmus IGKV1-135*01 F | 100 | 302 | GATGTTGTGATGACCCAGACTCCACTCACTTTGTCGGTTACCATTGGACAACCAGCCTCCATCTCTTGCAAGTCAAGTCAGAGCCTCTTAGATAGTGATGGAAAGACATATTTGAATTGGTTGTTACAGAGGCCAGGCCAGTCTCCAAAGCGCCTAATCTATCTGGTGTCTAAACTGGACTCTGGAGTCCCTGACAGGTTCACTGGCAGTGGATCAGGGACAGATTTCACACTGAAAATCAGCAGAGTGGAGGCTGAGGATTTGGGAGTTTATTATTGCTGGCAAGGTACACATTTTCCTCA |
| IGKV8-24*01_LEWES | LEWES/EiJ | Musmus IGKV8-24*01 F | 100 | 305 | GACATTGTGATGACACAGTCTCCATCCTCCCTGGCTATGTCAGTAGGACAGAAGGTCACTATGAGCTGCAAGTCCAGTCAGAGCCTTTTAAATAGTAGCAATCAAAAGAACTATTTGGCCTGGTACCAGCAGAAACCAGGACAGTCTCCTAAACTTCTGGTATACTTTGCATCCACTAGGGAATCTGGGGTCCCTGATCGCTTCATAGGCAGTGGATCTGGGACAGATTTCACTCTTACCATCAGCAGTGTGCAGGCTGAAGACCTGGCAGATTACTTCTGTCAGCAACATTATAGCACTCCTCC |
| IGKV15-103*01_S4601_LEWES | LEWES/EiJ | Musmus IGKV15-103*01 ORF | 98.21 | 287 | GACATCCAGATGAACCAGTCTCCATCCAGTCTGTCTGCATCCCTCGGAGACACAATTACCATCACTTGCCGTGCCAGTCAGAACATTAATATTTGGTTAAGCTGGTACCAGCAGAAACCAGGAAATATTCCTAAACTATTGATCTATAAGGCTTCCAACTTGCACACAGGCGTCCCATCAAGGTTTAGTGGCAGTGGATCTGGAACAGATTTCACATTAACCATCAGCAGCCTGCAGCCTGAAGACATTGCCACTTACTACTGTCTACAGGGTCAAAGTCATCCTCT |
| IGKV6-17*01_LEWES | LEWES/EiJ | Musmus IGKV6-17*01 F | 100 | 287 | GACATTGTGATGACCCAGTCTCACAAATTCATGTCCACATCAGTAGGAGACAGGGTCAGCATCACCTGCAAGGCCAGTCAGGATGTGAGTACTGCTGTAGCCTGGTATCAACAGAAACCAGGACAATCTCCTAAACTACTGATTTACTCGGCATCCTACCGGTACACTGGAGTCCCTGATCGCTTCACTGGCAGTGGATCTGGGACGGATTTCACTTTCACCATCAGCAGTGTGCAGGCTGAAGACCTGGCAGTTTATTACTGTCAGCAACATTATAGTACTCCTCC |

| | | | | | |
|---|---|---|---|---|---|
| IGKV8-16*01_LEWES | LEWES/EiJ | Musmus IGKV8-16*01 F | 100 | 305 | GAAATTGTGTTGACCCAGTCTATACCATCCCTGACTGTGTCAGCAGGAGAGAGGGTCACTATCAGCTGCAAATCCAATCAGAATCTTTTATGGAGTGGAAACCAAAGGTACTGTTTGGTCTGGCACCAGTGGAAACCAGGGCAAACTCCTACACCGTTGATCACCTGGACATCTGATAGGTACTCTGGAGTCCCTGATCGTTTCATAGGCAGTGGATCTGTGACAGATTTCACTCTGACCATCAGCAGTGTGCAGGCTGAAGATGTGGCAGTTTATTTCTGTCAGCAGCATTTACACATTCCTCC |
| IGKV1-122*01_S5263_LEWES | LEWES/EiJ | Musmus IGKV1-122*01 F | 97.28 | 302 | GATGCTGTGATGACCCAAACTCCACTCTCCCTGCCTGTCAGTCTTGGAGATCAAGCCTCCATCTCTTGCAGATCTAGTCAGAGCCTTGTACACAGTAATGGAAACACCTATTTGAACTGGTACCTCCAGAAACCAGGCCAGTCTCCACAGCTCTTGATCTACAGGGTTTCCAACCGATTTTCTGGGGTCCTAGACAGGTTCAGTGGCAGTGGTTCAGGGACAGATTTCACACTCAAGATCAGCAGAGTGGAGGCTGAGGATTTGGGAGTTTATTTCTGCCTCCAAGTTACACATATCCCTCC |
| IGKV4-57*01_LEWES | LEWES/EiJ | Musmus IGKV4-57*01 F | 100 | 286 | CAAATTGTTCTCACCCAGTCTCCAGCAATCATGTCTGCATCTCCAGGGGAGAAGGTCACCATAACCTGCAGTGCCAGCTCAAGTGTAAGTTACATGCACTGGTTCCAGCAGAAGCCAGGCACTTCTCCCAAACTCTGGATTTATAGCACATCCAACCTGGCTTCTGGAGTCCCTGCTCGCTTCAGTGGCAGTGGATCTGGGACCTCTTACTCTCTCACAATCAGCCGAATGGAGGCTGAAGATGCTGCCACTTATTACTGCCAGCAAAGGAGTAGTTACCCACCCA |
| IGKV1-117*02_S5705_LEWES | LEWES/EiJ | Musmus IGKV1-117*02 F | 99.66 | 301 | GATGTTGTGATGACCCAAACTCCACTCTCCCTGCCTGTCAGTCTTGGAGATCAAGCCTCCATCTCTTGCAGATCTAGTCAGAGCATTGTACATAGTAATGGAAACACCTATTTAGAATGGTACCTGCAGAAACCAGGCCAGTCTCCAAAGCTCCTGATCTACAAAGTTTCCAACCGATTGTCTGGGGTCCCAGACAGGTTCAGTGGCAGTGGATCAGGGACAGACTTCACACTCAAGATCAGCAGAGTGGAGGCTGAGGATCTGGGAGTTTATTACTGCTTTCAAGGTTCACATGTTCCTC |
| IGKV19-93*01_LEWES | LEWES/EiJ | Musmus IGKV19-93*01 F | 100 | 287 | GACATCCAGATGACACAGTCTCCATCCTCACTGTCTGCATCTCTGGGAGGCAAAGTCACCATCACTTGCAAGGCAAGCCAAGACATTAACAAGTATATAGCTTGGTACCAACACAAGCCTGGAAAAGGTCCTAGGCTGCTCATACATTACACATCTACATTACAGCCAGGCATCCCATCAAGGTTCAGTGGAAGTGGGTCTGGGAGAGATTATTCCTTCAGCATCAGCAACCTGGAGCCTGAAGATATTGCAACTTATTATTGTCTACAGTATGATAATCTTCTACC |
| IGKV4-68*01_S3092_LEWES | LEWES/EiJ | Musmus IGKV4-68*01 F | 100 | 286 | CAAATTGTTCTCACCCAGTCTCCAGCACTCATGTCTGCATCTCCAGGGGAGAAGGTCACCATGACCTGCAGTGCCAGCTCAAGTGTAAGTTACATGTACTGGTACCAGCAGAAGCCAAGATCCTCCCCCAAACCCTGGATTTATCTCACATCCAACCTGGCTTCTGGAGTCCCTGCTCGCTTCAGTGGCAGTGGGTCTGGGACCTCTTACTCTCTCACAATCAGCAGCATGGAGGCTGAAGATGCTGCCACTTATTACTGCCAGCAGTGGAGTAGTAACCCACTCA |
| IGKV3-12*01_LEWES | LEWES/EiJ | Musmus IGKV3-12*01 F | 100 | 299 | GACATTGTGCTGACACAGTCTCCTGCTTCCTTAGCTGTATCTCTGGGGCAGAGGGCCACCATCTCATGCAGGGCCAGCAAAAGTGTCAGTACATCTGGCTATAGTTATATGCACTGGTACCAACAGAAACCAGGACAGCCACCCAAACTCCTCATCTATCTTGCATCCAACCTAGAATCTGGGGTCCCTGCCAGGTTCAGTGGCAGTGGGTCTGGGACAGACTTCACCCTCAACATCCATCCTGTGGAGGAGGAGGATGCTGCAACCTATTACTGTCAGCACAGTAGGGAGCTTCCTCC |
| IGKV5-48*01_LEWES | LEWES/EiJ | Musmus IGKV5-48*01 F | 100 | 287 | GACATCTTGCTGACTCAGTCTCCAGCCATCCTGTCTGTGAGTCCAGGAGAAAGAGTCAGTTTCTCCTGCAGGGCCAGTCAGAGCATTGGCACAAGCATACACTGGTATCAGCAAAGAACAAATGGTTCTCCAAGGCTTCTCATAAAGTATGCTTCTGAGTCTATCTCTGGGATCCCTTCCAGGTTTAGTGGCAGTGGATCAGGGACAGATTTTACTCTTAGCATCAACAGTGTGGAGTCTGAAGATATTGCAGATTATTACTGTCAACAAAGTAATAGCTGGCCAAC |

| | | | | | |
|---|---|---|---|---|---|
| IGKV3-7*01_S5296_LEWES | LEWES/EiJ | Musmus IGKV3-7*02 F | 100 | 299 | GACATTGTGCTGACACAGTCTCCTGCTTCCTTAGCTGTATCTCTGGGGCAGAGGGCCACCATCTCATGCAGGGCCAGCCAAAGTGTCAGTACATCTAGCTATAGTTATATGCACTGGTACCAACAGAAACCAGGACAGCCACCCAAACTCCTCATCAAGTATGCATCCAACCTAGAATCTGGGGTCCCTGCCAGGTTCAGTGGCAGTGGGTCTGGGACAGACTTCACCCTCAACATCCATCCTGTGGAGGAGGAGGATACTGCAACATATTACTGTCAGCACAGTTGGGAGATTCCTCC |
| IGKV4-53*01_LEWES | LEWES/EiJ | Musmus IGKV4-53*01 F | 100 | 292 | GAAATTGTGCTCACCCAGTCTCCAGCACTCATGGCTGCATCTCCAGGGGAGAAGGTCACCATCACCTGCAGTGTCAGCTCAAGTATAAGTTCCAGCAACTTGCACTGGTACCAGCAGAAGTCAGAAACCTCCCCCAAACCCTGGATTTATGGCACATCCAACCTGGCTTCTGGAGTCCCTGTTCGCTTCAGTGGCAGTGGATCTGGGACCTCTTATTCTCTCACAATCAGCAGCATGGAGGCTGAAGATGCTGCCACTTATTACTGTCAACAGTGGAGTAGTTACCCACTCA |
| IGKV3-4*01_LEWES | LEWES/EiJ | Musmus IGKV3-4*01 F | 100 | 299 | GACATTGTGCTGACCCAATCTCCAGCTTCTTTGGCTGTGTCTCTAGGGCAGAGGGCCACCATCTCCTGCAAGGCCAGCCAAAGTGTTGATTATGATGGTGATAGTTATATGAACTGGTACCAACAGAAACCAGGACAGCCACCCAAACTCCTCATCTATGCTGCATCCAATCTAGAATCTGGGATCCCAGCCAGGTTTAGTGGCAGTGGGTCTGGGACAGACTTCACCCTCAACATCCATCCTGTGGAGGAGGAGGATGCTGCAACCTATTACTGTCAGCAAAGTAATGAGGATCCTCC |
| IGKV13-85*01_LEWES | LEWES/EiJ | Musmus IGKV13-85*01 F | 100 | 287 | GACATCCAGATGACACAATCTTCATCCTACTTGTCTGTATCTCTAGGAGGCAGAGTCACCATTACTTGCAAGGCAAGTGACCACATTAATAATTGGTTAGCCTGGTATCAGCAGAAACCAGGAAATGCTCCTAGGCTCTTAATATCTGGTGCAACCAGTTTGGAAACTGGGGTTCCTTCAAGATTCAGTGGCAGTGGATCTGGAAAGGATTACACTCTCAGCATTACCAGTCTTCAGACTGAAGATGTTGCTACTTATTACTGTCAACAGTATTGGAGTACTCCTCC |
| IGKV4-50*01_LEWES | LEWES/EiJ | Musmus IGKV4-50*01 F | 100 | 286 | GAAAATGTGCTCACCCAGTCTCCAGCAATCATGTCTGCATCTCTAGGGGAGAAGGTCACCATGAGCTGCAGGGCCAGCTCAAGTGTAAATTACATGTACTGGTACCAGCAGAAGTCAGATGCCTCCCCCAAACTATGGATTTATTACACATCCAACCTGGCTCCTGGAGTCCCAGCTCGCTTCAGTGGCAGTGGGTCTGGGAACTCTTATTCTCTCACAATCAGCAGCATGGAGGGTGAAGATGCTGCCACTTATTACTGCCAGCAGTTTACTAGTTCCCCATCCA |
| IGKV1-110*01_S3077_LEWES | LEWES/EiJ | Musmus IGKV1-110*01 F, or Musmus IGKV1-117*01 F | 97.62 | 301 | GATGTTATGATGACCCAAACTCCACTCTCCCTGCCTGTCAGTCTTGGAGATCAAGCCTCCATCTCTTGCAGATCTAGTCAGAGCCTTGTACACAGTAATGGAAACACCTATTTATATTGGTTCCTGCAGAAACCAGGCCAGTCTCCAAAGCTCCTGATCTACAAAGTTTCCAACCGATTTTCTGGGGTCCCAGACAGGTTCAGTGGCAGTGGATCAGGGACAGATTTCACACTCAAGATCAGCAGAGTGGAGGCTGAGGATCTGGGAGTTTATTACTGCTTTCAAGGTACACATGTTCCTC |
| IGKV9-120*01_S2575_LEWES | LEWES/EiJ | Musmus IGKV9-120*02 F | 94.98 | 285 | GACATCCAGATGACACAGTCTCCATCCTCCTTATCTGCCTCTCTGGGAGACAGAATCAGTCTCACTTGCCGGGCAAGTCAGGACATTGGTGGGAGCTTAAACTGGTTTCAGCAGAAACCAGATGAAACTTTTAAACGCCTGATATACGCCACATCCAATTTAGATTCTGGTGTCCCAAAAAGGTTCAGTGGCAGTAGGTCTGGGTCAGATTATTCTCTCACCATCAGCAGCCTTGAGTCTGAAGATTTTGCAGACTATTACTGTCTACAATATGCTAGTTATCCT |
| IGKV13-84*01_LEWES | LEWES/EiJ | Musmus IGKV13-84*01 F | 100 | 287 | GACATCCAGATGACACAATCTTCATCCTCCTTTTCTGTATCTCTAGGAGACAGAGTCACCATTACTTGCAAGGCAAGTGAGGACATATATAATCGGTTAGCCTGGTATCAGCAGAAACCAGGAAATGCTCCTAGGCTCTTAATATCTGGTGCAACCAGTTTGGAAACTGGGGTTCCTTCAAGATTCAGTGGCAGTGGATCTGGAAAGGATTACACTCTCAGCATTACCAGTCTTCAGACTGAAGATGTTGCTACTTATTACTGTCAACAGTATTGGAGTACTCCTCC |

| | | | | | |
|---|---|---|---|---|---|
| IGKV8-34*01_LEWES | LEWES/EiJ | Musmus IGKV8-34*01 F | 100 | 305 | GACATTTTGATGACCCAGTCTCCATCCTCCCTGACTGTGTCAGCAGGAGAGAAGGTCACTATGAGCTGCAAGTCCAGTCAGAGTCTTTTAGCTAGTGGCAACCAAAATAACTACTTGGCCTGGCACCAGCAGAAACCAGGACGATCTCCTAAAATGCTGATAATTTGGGCATCCACTAGGGTATCTGGAGTCCCTGATCGCTTCATAGGCAGTGGATCTGGGACGGATTTCACTCTGACCATCAACAGTGTGCAGGCTGAAGATCTGGCTGTTTATTACTGTCAGCAGTCCTACAGCGCTCCTAC |
| IGKV6-32*01_LEWES | LEWES/EiJ | Musmus IGKV6-32*01 F | 100 | 287 | AGTATTGTGATGACCCAGACTCCCAAATTCCTGCTTGTATCAGCAGGAGACAGGGTTACCATAACCTGCAAGGCCAGTCAGAGTGTGAGTAATGATGTAGCTTGGTACCAACAGAAGCCAGGGCAGTCTCCTAAACTGCTGATATACTATGCATCCAATCGCTACACTGGAGTCCCTGATCGCTTCACTGGCAGTGGATATGGGACGGATTTCACTTTCACCATCAGCACTGTGCAGGCTGAAGACCTGGCAGTTTATTTCTGTCAGCAGGATTATAGCTCTCCTCC |
| IGKV3-2*01_LEWES | LEWES/EiJ | Musmus IGKV3-2*01 F | 100 | 299 | GACATTGTGCTGACCCAATCTCCAGCTTCTTTGGCTGTGTCTCTAGGGCAGAGGGCCACCATCTCCTGCAGAGCCAGCGAAAGTGTTGATAATTATGGCATTAGTTTTATGAACTGGTTCCAACAGAAACCAGGACAGCCACCCAAACTCCTCATCTATGCTGCATCCAACCAAGGATCCGGGGTCCCTGCCAGGTTTAGTGGCAGTGGGTCTGGGACAGACTTCAGCCTCAACATCCATCCTATGGAGGAGGATGATACTGCAATGTATTTCTGTCAGCAAAGTAAGGAGGTTCCTCC |
| IGKV3-5*01_LEWES | LEWES/EiJ | Musmus IGKV3-5*01 F | 100 | 299 | GACATTGTGCTGACCCAATCTCCAGCTTCTTTGGCTGTGTCTCTAGGGCAGAGGGCCACCATATCCTGCAGAGCCAGTGAAAGTGTTGATAGTTATGGCAATAGTTTTATGCACTGGTACCAGCAGAAACCAGGACAGCCACCCAAACTCCTCATCTATCGTGCATCCAACCTAGAATCTGGGATCCCTGCCAGGTTCAGTGGCAGTGGGTCTAGGACAGACTTCACCCTCACCATTAATCCTGTGGAGGCTGATGATGTTGCAACCTATTACTGTCAGCAAAGTAATGAGGATCCTCC |
| IGKV4-74*01_LEWES | LEWES/EiJ | Musmus IGKV4-74*01 F | 100 | 292 | CAAATTGTTCTCACCCAGTCTCCAGCAATCATGTCTGCATCTCTAGGGGAACGGGTCACCATGACCTGCACTGCCAGCTCAAGTGTAAGTTCCAGTTACTTGCACTGGTACCAGCAGAAGCCAGGATCCTCCCCCAAACTCTGGATTTATAGCACATCCAACCTGGCTTCTGGAGTCCCAGCTCGCTTCAGTGGCAGTGGGTCTGGGACCTCTTACTCTCTCACAATCAGCAGCATGGAGGCTGAAGATGCTGCCACTTATTACTGCCACCAGTATCATCGTTCCCCACCCA |
| IGKV4-70*01_LEWES | LEWES/EiJ | Musmus IGKV4-70*01 F | 100 | 282 | CAAATTGTTCTCACCCAGTCTCCAGCAATCATGTCTGCATCTCCAGGGGAGAAGGTCACCATGACCTGCAGTGCCAGCTCAAGTATAAGTTACATGCACTGGTACCAGCAGAAGCCAGGCACCTCCCCCAAAAGATGGATTTATGACACATCCAAACTGGCTTCTGGAGTCCCTGCTCGCTTCAGTGGCAGTGGGTCTGGGACCTCTTATTCTCTCACAATCAGCAGCATGGAGGCTGAAGATGCTGCCACTTATTACTGCCATCAGCGGAGTAGTTACCCA |
| IGKV4-72*01_S3540_LEWES | LEWES/EiJ | Musmus IGKV4-72*01 F | 100 | 286 | CAAATTGTTCTCTCCCAGTCTCCAGCAATCCTGTCTGCATCTCCAGGGGAGAAGGTCACAATGACTTGCAGGGCCAGCTCAAGTGTAAGTTACATGCACTGGTACCAGCAGAAGCCAGGATCCTCCCCCAAACCCTGGATTTATGCCACATCCAACCTGGCTTCTGGAGTCCCTGCTCGCTTCAGTGGCAGTGGGTCTGGGACCTCTTACTCTCTCACAATCAGCAGAGTGGAGGCTGAAGATGCTGCCACTTATTACTGCCAGCAGTGGAGTAGTAACCCACTCA |
| IGKV6-25*01_LEWES | LEWES/EiJ | Musmus IGKV6-25*01 F | 100 | 287 | GACATTGTGATGACCCAGTCTCACAAATTCATGTCCACATCAGTAGGAGACAGGGTCAGCATCACCTGCAAGGCCAGTCAGGATGTGAGTACTGCTGTAGCCTGGTATCAACAAAAACCAGGGCAATCTCCTAAACTACTGATTTACTGGGCATCCACCCGGCACACTGGAGTCCCTGATCGCTTCACAGGCAGTGGATCTGGGACAGATTATACTCTCACCATCAGCAGTGTGCAGGCTGAAGACCTGGCACTTTATTACTGTCAGCAACATTATAGCACTCCTCC |

| | | | | | |
|---|---|---|---|---|---|
| IGKV4-61*01_LEWES | LEWES/EiJ | Musmus IGKV4-61*01 F | 100 | 286 | CAAATTGTTCTCACCCAGTCTCCAGCAATCATGTCTGCATCTCCAGGGGAGAAGGTCACCATATCCTGCAGTGCCAGCTCAAGTGTAAGTTACATGTACTGGTACCAGCAGAAGCCAGGATCCTCCCCCAAACCCTGGATTTATCGCACATCCAACCTGGCTTCTGGAGTCCCTGCTCGCTTCAGTGGCAGTGGGTCTGGGACCTCTTACTCTCTCACAATCAGCAGCATGGAGGCTGAAGATGCTGCCACTTATTACTGCCAGCAGTATCATAGTTACCCACCCA |
| IGKV14-100*01_LEWES | LEWES/EiJ | Musmus IGKV14-100*01 F | 100 | 287 | GACATCCTGATGACCCAATCTCCATCCTCCATGTCTGTATCTCTGGGAGACACAGTCAGCATCACTTGCCATGCAAGTCAGGGCATTAGCAGTAATATAGGGTGGTTGCAGCAGAAACCAGGGAAATCATTTAAGGGCCTGATCTATCATGGAACCAACTTGGAAGATGGAGTTCCATCAAGGTTCAGTGGCAGTGGATCTGGAGCAGATTATTCTCTCACCATCAGCAGCCTGGAATCTGAAGATTTTGCAGACTATTACTGTGTACAGTATGCTCAGTTTCCTCC |
| IGKV11-125*01_S3701_LEWES | LEWES/EiJ | Musmus IGKV11-106*02 F | 96.06 | 287 | GATGTTCAAATGACCCAGTCTCCATCCTCCCCGTCTGCATCTTTGGGAGAGAGAGTCTCCCTGACTTGCCAGGCAAGTCAGGACATTAGCAATTATTTAAACTGGTATCAGCAAACACCAGGGAAAGCTCCTAGGCTCTTGATCTATGGTGCAAGCAACTTGGAAGATGGGGTCCCTTCAAGGTTCAGTGGCACTGGATATGGGACAGATTTCACTTTCACCATCAGCAGCCTGGAGGAAGAAGATGTGGCAACTTATTTCTGTCTACAGCATGGTTATCTCCCTCC |
| IGKV2-112*02_S7480_LEWES | LEWES/EiJ | Musmus IGKV2-112*02 F | 98.3 | 301 | GATATTGTGATAACCCAGGATGAACTCTCCAATCCTGTCACTTCTGGAGAATCGGTTTCCATCTCCTGCAGGTCTAGTAAGAGTCTCCTATATACAGATGGGAAGACATACTTGAATTGGTTTCTGCAGAGGCCAGGACAGTCTCCTCAGCTCCTGGTCTATTGGATGTCCACTCGTGCATCAGGAGTCTCAGACCGGTTTAGTGGCAGTGGGTCAGGAACAGATTTCACACTGGAAATCAGTAGAGTGAAGGCTGAGGATGTGGGTGTGTATTACTGTCAACAAGTTGTAGAGTATCCTC |
| IGKV1-133*01_S9974_LEWES | LEWES/EiJ | Musmus IGKV1-133*01 F | 96.26 | 302 | GATGTTGTGATGACCCAGACTCCACTCACTTTGTCTGTTACCATTGGACAGCCAGCTTCCATCTCTTGCAAGTCAAGTCAGAGCCTCTTATATAGTGATGGAAAAACCTATTTGAATTGGTTATTACAGAGTCCAGGCCAGTCTCCAAAGCTCCTAATCTATCTGGTGTCTAAACTGGAATCTGGAGTCCCTGACAGATTCAGTGGCAGTGGATCAGGGACAGATTTTACACTGAAAATCAGCAGAGTGGAGGCTGAGGATTTGGGAGTTTATTACTGCGTGCAAGGTACACATTTCCCTCA |
| IGKV4-92*01_LEWES | LEWES/EiJ | Musmus IGKV4-92*01 F | 100 | 291 | GAAATGGTTCTCACCCAGTCTCCAGTATCCATAACTGCATCTCGAGGGGAGAAGGTCACCATCACCTGCCGTGCCAGCTCAAGTATAAGTTCCAATTACTTACACTGGTACCAGCAGAAGCCAGGATCCTCCCCTAAACTTTTGATTTATAGGACATCCATCCTGGCATCTGGAGTCCTAGACAGCTTCAGTGGCAGTGGGTCTGAGAGCTCTTACACTCTGACAATCAGCTGCATGCAGGACGAAGTTGCTGCCACTTACTATTGTCAGCAGGGGAGTAGTAGCCCACCA |
| IGKV11-125*01_S3707_LEWES | LEWES/EiJ | Musmus IGKV11-106*02 F | 96.42 | 287 | AATGTTCAAATGACCCAGTCTCCATCCTCCCTGTCTGCATCTTTGGGAGAGAGAGTCTCCCTGACTTGCCAGGCAAGACAGAGCGTTAGCAATAATTTAAACTGGTATCAGCAAACACCAGGGAAAGCTCCTAGGCTCTTGATCTATGGTGCAAGCAAATTGGAAGATGGGGTCCCTTCAAGGTTCAGTGGCACTGGATATGGGACAGATTTCACTTTCACCATCAGCAGCCTGGAGGAAGAAGATGTGGCAACTTATTTCTGTCTACAGCATAGTTATCTCCCTCC |
| IGKV3-10*01_LEWES | LEWES/EiJ | Musmus IGKV3-10*01 F | 100 | 299 | AACATTGTGCTGACCCAATCTCCAGCTTCTTTGGCTGTGTCTCTAGGGCAGAGGGCCACCATATCCTGCAGAGCCAGTGAAAGTGTTGATAGTTATGGCAATAGTTTTATGCACTGGTACCAGCAGAAACCAGGACAGCCACCCAAACTCCTCATCTATCTTGCATCCAACCTAGAATCTGGGGTCCCTGCCAGGTTCAGTGGCAGTGGGTCTAGGACAGACTTCACCCTCACCATTGATCCTGTGGAGGCTGATGATGCTGCAACCTATTACTGTCAGCAAAATAATGAGGATCCTCC |

| | | | | | |
|---|---|---|---|---|---|
| IGKV1-135*01_S6159_LEWES | LEWES/EiJ | Musmus IGKV1-135*01 F | 95.58 | 301 | GATGTTGTGATGACTCAGACCCCACTCACTTTGTCGGTTACCATTGGACAACCAGCCTCCATCTCTTGCAAGTCAAGTCAGAGCCTCTTATATAGTGATGGAAAGACATATTTGGATTGGTTATTACAGAGGCCAGGTCAGTCTCCAAAGAGCCTAATCTATCTGGTGTCTAAACTGGAATCTGGAGTCCCTGACAGGTTCAGTGGCAGTGGATCAGGGACAGATTTCATACTGAAAATCAGCAGAGTGGAGGCTGAGGATTTGGGAGTTTATTACTGCGTGCAAGGTACACATTTTCCTC |
| IGKV5-39*01_LEWES | LEWES/EiJ | Musmus IGKV5-39*01 F | 100 | 287 | GACATTGTGATGACTCAGTCTCCAGCCACCCTGTCTGTGACTCCAGGAGATAGAGTCTCTCTTTCCTGCAGGGCCAGCCAGAGTATTAGCGACTACTTACACTGGTATCAACAAAAATCACATGAGTCTCCAAGGCTTCTCATCAAATATGCTTCCCAATCCATCTCTGGGATCCCCTCCAGGTTCAGTGGCAGTGGATCAGGGTCAGATTTCACTCTCAGTATCAACAGTGTGGAACCTGAAGATGTTGGAGTGTATTACTGTCAAAATGGTCACAGCTTTCCTCC |
| IGKV1-117*01_S7519_LEWES | LEWES/EiJ | Musmus IGKV1-117*01 F | 96.94 | 302 | GATGTTTTGATGACCCAAACTCCACTCTCCCTGCCTGTCAGTCTTGGAGATCAAGCCTCCATCTCTTGCAGATCTAGTCAGAGCCTTGTACACAGTAATGGAAACACCTATTTATATTGGTTACTGCAGAAGTCAGGCCAGTCTCCAAAGCTCCTGATCTACAAAGTTTCCAACCGATTTTCTGGGGTCCCAGACAGGTTCAGTGGCAATGGATCAGGGACAGATTTCACACTCAAGATCAGCAGAGTGGAGGCTGAGGATCTGGGAGTTTATTACTGCTTTCAAGGTTCACATGTTCCTCC |
| IGKV14-126*01_S4454_LEWES | LEWES/EiJ | Musmus IGKV14-126*01 F | 99.28 | 287 | GACATCAAGATGACCCAGTCTCCATCCTCCATGTATGCATCGCTGGGAGAGAGAGTCACTATCACTTGCAAGGCGAGTCAGGACATTAAAAGCTATTTAAGCTGGTACCAGCAGAAACCAGGGAAATCTCCTAAGACCCTGATCTATTATGCAACAAGCTTGGCAGATGGGGTCCCATCAAGATTCAGCGGCAGTGGATCTGGGCAAGATTATTCTCTAACCATCAGCAGCCTGGAGTCTGACGATACAGCAACTTATTACTGTCTACAGCATGGTGAGAGCCCTCC |
| IGKV4-79*01_LEWES | LEWES/EiJ | Musmus IGKV4-79*01 F | 100 | 292 | CAAATTGTTCTCACCCAGTCTCCAGCAATCATGTCTGCATCTCCTGGGGAGAAGGTCACCTTGACCTGCAGTGCCAGCTCAAGTGTAAGTTCCAGCTACTTGTACTGGTACCAGCAGAAGCCAGGATCCTCCCCCAAACTCTGGATTTATAGCACATCCAACCTGGCTTCTGGAGTCCCTGCTCGCTTCAGTGGCAGTGGGTCTGGGACCTCTTACTCTCTCACAATCAGCAGCATGGAGGCTGAAGATGCTGCCTCTTATTTCTGCCATCAGTGGAGTAGTTACCCACCCC |
| IGKV11-125*01_LEWES | LEWES/EiJ | Musmus IGKV11-125*01 F | 100 | 287 | GATGTCCAGATGATTCAGTCTCCATCCTCCCTGTCTGCATCTTTGGGAGACATAGTCACCATGACTTGCCAGGCAAGTCAGGGCACTAGCATTAATTTAAACTGGTTTCAGCAAAAACCAGGGAAAGCTCCTAAGCTCCTGATCTATGGTGCAAGCAACTTGGAAGATGGGGTCCCATCAAGGTTCAGTGGCAGTAGATATGGGACAGATTTCACTCTCACCATCAGCAGCCTGGAGGATGAAGATATGGCAACTTATTTCTGTCTACAGCATAGTTATCTCCCTCC |
| IGKV14-111*01_S1462_LEWES | LEWES/EiJ | Musmus IGKV14-111*01 F | 97.85 | 286 | GACATCAAGATGACCCAGTCTCCATCTTCCATGTATGCATCTCTAGGAGAGAGAGTCACTATCACTTGCAAGGCGAGTCAGGACATTAATAGCTATTTAAGCTGGTACCAGCAGAAACCAGGGAAATCTCCTAAGACCCTGATCTATTATGCAAACAGCTTGGTAGATGGGGTCCCATCAAGGTTCAGTGGCAGTGGATCTGGGCAAGATTGTTCTCTCACCATCAGCAGCCTGGAGTATGAAGATACGGGAATTTATTATTGTCTACAGTATGATGAGTTTCCTC |
| IGKV9-123*01_S5635_LEWES | LEWES/EiJ | Musmus IGKV9-123*01 F | 98.57 | 285 | GACATCCAGATGATTCAGTCTCCATCGTCCATGTTTGTCTCTCTGGGAGACAGAGTCAGTCTCTCTTGCCGGGCTAGTCAGGGCATTAGAGGTAATTTAGACTGGTATCAGCAGAAACCAGGTGGAACTATTAAACTCCTGATCTACTCCACATCCAATTTAAATTCTGGTGTCCCATCAAGGTTCAGTGGCAGTGGGTCTGGGTCAGATTATTCTCTCACCATCAGCAGCCTAGAGTCTGAAGATTTTGCAGACTATTACTGCCTACAGCGTAATGCATATCCT |

| | | | | | |
|---|---|---|---|---|---|
| IGKV1-110*02_S1372_LEWES | LEWES/EiJ | Musmus IGKV1-110*02 F, or Musmus IGKV1-117*01 F | 95.92 | 302 | GATGTTGTGATGACCCAAACTCCACTCTCCCTGCCTGTCAGTCTTGGAGATCAAGCCTCCATCTCTTGCAGATCTAGTCAGAGCATTGTACACAGTAATAGATACACCTATTTAGAATGGTACCTGCAGAAACCAGGCCAGTCTCTAAAGCTCCTGATATACGGGGTTTCCAACCGATTTTCTGGGGTCCCAGACAGGTTCAGTGGCAGTGGATCAAGGACAGATTTCACACTCAAGATCAGCAGAGTGGAGGCTGAGGATCTGGGAGTTTATTTCTGCTTTCAAGGTACACATGTTCCTCA |
| IGKV12-98*01_LEWES | LEWES/EiJ | Musmus IGKV12-98*01 F | 100 | 287 | GACATTCAGATGACCCAGTCTCCTGCCTCCCAGTCTGCATCTCTGGGAGAAAGTGTCACCATCACATGCCTGGCAAGTCAGACCATTGGTACATGGTTAGCATGGTATCAGCAGAAACCAGGGAAATCTCCTCAGCTCCTGATTTATGCTGCAACCAGCTTGGCAGATGGGGTCCCATCAAGGTTCAGTGGTAGTGGATCTGGCACAAAATTTTCTTTCAAGATCAGCAGCCTACAGGCTGAAGATTTTGTAAGTTATTACTGTCAACAACTTTACAGTACTCCTCT |
| IGKV14-130*01_LEWES | LEWES/EiJ | Musmus IGKV14-130*01 F | 100 | 287 | GAAATCCAGATGACCCAGTCTCCATCCTCTATGTCTGCATCTCTGGGAGACAGAATAACCATCACTTGCCAGGCAACTCAAGACATTGTTAAGAATTTAAACTGGTATCAGCAGAAACCAGGGAAACCCCCTTCATTCCTGATCTATTATGCAACTGAACTGGCAGAAGGGGTCCCATCAAGGTTCAGTGGCAGTGGGTCTGGGTCAGACTATTCTCTGACAATCAGCAACCTGGAGTCTGAAGATTTTGCAGACTATTACTGTCTACAGTTTTATGAGTTTCCTCC |
| IGKV5-45*01_LEWES | LEWES/EiJ | Musmus IGKV5-45*01 F | 100 | 287 | GATATTGTGCTAACTCAGTCTCCAGCCACCCTGTCTGTGACTCCAGGAGATAGAGTCAGTCTTTCCTGCAGGGCCAGTCAAAGTATTAGCAACTACCTACACTGGTATCAACAAAAATCACATGAGTCTCCAAGGCTTCTCATCAAGTATGCTTCCCAGTCCATCTCTGGGATCCCCTCCAGGTTCAGTGGCAGTGGATCAGGGACAGATTTCACTCTCAGTATCAACAGTGTGGAGACTGAAGATTTTGGAATGTATTTCTGTCAACAGAGTAACAGCTGGCCTCA |
| IGKV7-33*01_LEWES | LEWES/EiJ | Musmus IGKV7-33*01 F | 100 | 305 | GACATTGTGATGACTCAGTCTCCAACTTTCCTTGCTGTGACAGCAAGTAAGAAGGTCACCATTAGTTGCACGGCCAGTGAGAGCCTTTATTCAAGCAAACACAAGGTGCACTACTTGGCTTGGTACCAGAAGAAACCAGAGCAATCTCCTAAACTGCTGATATACGGGGCATCCAACCGATACATTGGGGTCCCTGATCGCTTCACAGGCAGTGGATCTGGGACAGATTTCACTCTGACCATCAGCAGTGTACAGGTTGAAGACCTCACACATTATTACTGTGCACAGTTTTACAGCTATCCTCT |
| IGKV1-88*01_LEWES | LEWES/EiJ | Musmus IGKV1-88*01 F | 100 | 302 | GATGTTGTGGTGACTCAAACTCCACTCTCCCTGCCTGTCAGCTTTGGAGATCAAGTTTCTATCTCTTGCAGGTCTAGTCAGAGTCTTGCAAACAGTTATGGGAACACCTATTTGTCTTGGTACCTGCACAAGCCTGGCCAGTCTCCACAGCTCCTCATCTATGGGATTTCCAACAGATTTTCTGGGGTGCCAGACAGGTTCAGTGGCAGTGGTTCAGGGACAGATTTCACACTCAAGATCAGCACAATAAAGCCTGAGGACTTGGGAATGTATTACTGCTTACAAGGTACACATCAGCCTCC |
| IGKV12-38*01_LEWES | LEWES/EiJ | Musmus IGKV12-38*01 F | 100 | 287 | GACATCCAGATGACTCAGTCTCCAGCCTCCCTGGCTGCATCTGTGGGAGAAACTGTCACCATCACATGTCGAGCAAGTGAGAACATTTACTACAGTTTAGCATGGTATCAGCAGAAGCAAGGGAAATCTCCTCAGCTCCTGATCTATAATGCAAACAGCTTGGAAGATGGTGTCCCATCGAGGTTCAGTGGCAGTGGATCTGGGACACAGTATTCTATGAAGATCAACAGCATGCAGCCTGAAGATACCGCAACTTATTTCTGTAAACAGGCTTATGACGTTCCTCC |
| IGKV4-69*01_S6643_LEWES | LEWES/EiJ | Musmus IGKV4-69*01 F | 99.64 | 281 | CAAATTCTTCTCACCCAGTCTCCAGCAATCATGTCTGCATCTCCAGGGGAGAAGGTCACCATGACCTGCAGTGCCAGCTCAAGTGTAAGTTACATGCACTGGTACCAGCAGAAGCCAGGATCCTCGCCCAAACCCTGGATTTATGACACATCCAACCTGGCTTCTGGATTCCCTGCTCGCTTCAGTGGCTGTGGGTCTGGGACCTCTTACTCTCTCATAATCAGCAGCATGGAGGCTGAAGATGCTGCCACTTATTACTGCCATCAGCGGAGTAGTTACCC |

| | | | | | |
|---|---|---|---|---|---|
| IGKV4-57*01_S9219_LEWES | LEWES/EiJ | Musmus IGKV4-57-1*01 F | 100 | 286 | CAAATTGTTCTCACCCAGTCTCCAGCAATCATGTCTGCATCTCCAGGGGAGAAGGTCACCATAACCTGCAGTGCCAGCTCAAGTGTAAGTTACATGCACTGGTTCCAGCAGAAGCCAGGCACTTCTCCCAAACTCTGGATTTATAGCACATCCAACCTGGCTTCTGGAGTCCCTGCTCGCTTCAGTGGCAGTGGATCTGGGACCTCTTACTCTCTCACAATCAGCCGAATGGAGGCTGAAGATGCTGCCACTTATTACTGCCAGCAAAGGAGTAGTTACCCACTCA |
| IGKV4-63*01_LEWES | LEWES/EiJ | Musmus IGKV4-63*01 F | 100 | 286 | GAAAATGTTCTCACCCAGTCTCCAGCAATCATGTCTGCATCTCCAGGGGAAAAGGTCACCATGACCTGCAGTGCCAGCTCAAGTGTAAGTTACATGCACTGGTACCAGCAGAAGTCAAGCACCTCCCCCAAACTCTGGATTTATGACACATCCAAACTGGCTTCTGGAGTCCCAGGTCGCTTCAGTGGCAGTGGGTCTGGAAACTCTTACTCTCTCACGATCAGCAGCATGGAGGCTGAAGATGTTGCCACTTATTACTGTTTTCAGGGGAGTGGGTACCCACTCA |
| IGKV2-109*03_S3211_LEWES | LEWES/EiJ | Musmus IGKV2-109*03 F, or Musmus IGKV2-109*04 F | 97.62 | 301 | GATATTGTGATGACACAGGCTGCATTCTCCAATCCAGTCACTCTTGGAACATCAGCTTCCATCTCCTGCAGGTCTAGTAAGAGTCTCCTAAATAGTAATGGCATCACTTCTTTGTATTGGTATCTGCAGAGGCCAGGCCAGTCTCCTCAGCTCCTGATATATCGGATGTCCAACCTTGCCTCAGGAGTCCCAGACAGGTTCAGTAGCAGTGGGTCAGGAACTGATTTCACACTGAGAATCAGCAGAGTGGAGGCTGAGGATGTGGGTGTTTATTACTGTGCTCAAAGGCTAGAACTCCCTC |
| IGKV11-125*01_S3440_LEWES | LEWES/EiJ | Musmus IGKV11-106*02 F | 96.06 | 287 | GATGTTCAAATGACCCAGTCTCCATCCTCCCTGTCTGCATCTTTGGGAGAGAGAGTCTCCCTGACTTGCCAGGCAAGTCAGAGCGTTAGCAATTATTTAAACTGGTATCAGCAAACACCAGGGAAAGCTCCTAGGCTCTTGATCTATGGTGCAAGCAAATTGGAAGATAGGGTCCCTTCAAGGTTCAGTGGAACTGGATATGGGACAGATTTCACTTTCACCATCAGAAGCCTGGAGGAAGAAGATGTGGCAACTTATTTCTGTCTACAGCATAGGTATCTCCCTCC |
| IGKV4-58*01_LEWES | LEWES/EiJ | Musmus IGKV4-58*01 F | 100 | 292 | GAAAATGTGCTCACCCAGTCTCCAGCAATAATGGCTGCCTCTCTGGGGCAGAAGGTCACCATGACCTGCAGTGCCAGCTCAAGTGTAAGTTCCAGTTACTTGCACTGGTACCAGCAGAAGTCAGGCGCTTCCCCCAAACCCTTGATTCATAGGACATCCAACCTGGCTTCTGGAGTCCCAGCTCGCTTCAGTGGCAGTGGGTCTGGGACCTCTTACTCTCTCACAATCAGCAGCGTGGAGGCTGAAGATGATGCAACTTATTACTGCCAGCAGTGGAGTGGTTACCCATTCA |
| IGKV4-90*01_LEWES | LEWES/EiJ | Musmus IGKV4-90*01 F | 100 | 285 | GAAATTTTGCTCACCCAGTCTCCAGCAATCATAGCTGCATCTCCTGGGGAGAAGGTCACCATCACCTGCAGTGCCAGCTCAAGTGTAAGTTACATGAACTGGTACCAGCAGAAACCAGGATCCTCCCCCAAAATATGGATTTATGGTATATCCAACCTGGCTTCTGGAGTTCCTGCTCGCTTCAGTGGCAGTGGGTCTGGGACATCTTTCTCTTTCACAATCAACAGCATGGAGGCTGAAGATGTTGCCACTTATTACTGTCAGCAAAGGAGTAGTTACCCACCC |
| IGKV2-109*03_S4495_LEWES | LEWES/EiJ | Musmus IGKV2-109*03 F, or Musmus IGKV2-109*04 F | 98.98 | 301 | GATATTGTGATGACGCAGGCTGCATTCTCCAATCCAGTCACTCTTGGAACATCAGCTTCCATCTCCTGCAGGTCTAGTAAGAGTCTCCTACATAGTAATGGCATCACTTATTTGTATTGGTATCTGCAGAGGCCAGGCCAGTCTCCTCAGCTCCTGATATATCGGATGTCCAACCTTGCCTCAGGAGTCCCAGACAGGTTCAGTAGCAGTGGGTCAGGAACTGATTTCACACTGAGAATCAGCAGAGTGGAGGCTGAGGATGTGGGTGTTTATTACTGTGCTCAAATGCTAGAACTCCCTC |
| IGKV4-51*01_LEWES | LEWES/EiJ | Musmus IGKV4-51*01 F | 100 | 292 | GAAAATGTGCTCACCCAGTCTCCAGCAATAATGGCTGCCTCTCTGGGGGAGAAGGTCACCATGACCTGCAGTGCCAGCTCAAGTGTAAGTTCCAGCTACTTGCACTGGTACCAGCAGAAGTCAGGCACTTCCCCCAAACTCTGGATTTATGGCACATCCAACCTGGCTTCTGGAGTCCCAGCTCGCTTCAGTGGCAGTGGGGCTGGGATCTCTTACTCTCTCACAATCAGCAGCATGGAGGCTGAAAATGATGCAACTTATTACTGCCAGCAGTGGAGTGGTTACCCATTCA |

| | | | | | |
|---|---|---|---|---|---|
| IGKV5-37*01_LEWES | LEWES/EiJ | Musmus IGKV5-37*01 F | 100 | 287 | GACATCCTGCTGACCCAGTCTCCAGCCACCCTGTCTGTGACTCCAGGAGAAACAGTCA<br>GTCTTTCCTGTAGGGCCAGCCAGAGTATTTACAAGAACCTACACTGGTATCAACAGAA<br>ATCACATCGGTCTCCAAGGCTTCTCATCAAGTATGCTTCTGATTCCATCTCTGGGATCC<br>CCTCCAGGTTCACTGGCAGTGGATCAGGGACAGATTACACTCTCAGTATCAACAGTGT<br>GAAGCCCGAAGATGAAGGAATATATTACTGTCTTCAAGGTTACAGCACACCTTC |
| IGKV4-50*01_S0901_LEWES | LEWES/EiJ | Musmus IGKV4-50*01 F | 100 | 286 | GAAAATGTGCTCACCCAGTCTCCAGCAATCATGTCTGCATCTCTAGGGGAGAAGGTCA<br>CCATGAGCTGCAGGGCCAGCTCAAGTGTAAATTACATGTACTGGTACCAGCAGAAGTC<br>AGATGCCTCCCCCAAACTATGGATTTATTACACATCCAACCTGGCTCCTGGAGTCCCA<br>GCTCGCTTCAGTGGCAGTGGGTCTGGGAACTCTTATTCTCTCACAATCAGCAGCATGG<br>AGGGTGAAGATGCTGCCACTTATTACTGCCAGCAGTTTACTAGTTCCCCATACA |
| IGKV6-15*01_S9105_MRL | MRL/MpJ | Musmus IGKV6-15*01 F | 97.13 | 287 | GACATTGTGATGACCCAGTCTCAAAAATTCATGTCCACATCAGTAGGAGACAGGGTCA<br>GCATCAGCTGCAAGGCCAGTCAGAATGTGGGTAATATTATAGCCTGGTATCAACAGA<br>AACCAGGGCAATCTCCTAAAGCACTGATTTACTTGGCATCCTACCGGTACAGTGGAGT<br>CCCTGATCGCTTCACAGGCAGTGGATCTGGGACAGATTTCACTCTCACCATTAGCAAT<br>GTGCAGTCTGAAGACTTGGCAGAGTATTTCTGTCAGCAATATAGCAGCTCTCCTCT |
| IGKV6-20*01_S2082_MRL | MRL/MpJ | Musmus IGKV6-20*01 F | 99.64 | 287 | AACATTGTAATGACCCAATCTCCCAAATCCATGTCCATGTCAGTAGGAGAGAGGGTCA<br>CCTTGAGCTGCAAGGCCAGTGAGAATGTGGGTACTTATGTATCCTGGTATCAACAGAA<br>ACCAGAGCAGTCTCCTAAACTGCTGATATACGGGGCATCCAACCGGTACACTGGGGTC<br>CCCGATCGCTTCACAGGCAGTGGATCTGCAACAGATTTCACTCTGACCATCAGCAGTG<br>TGCAGGCTGAAGACCTTGCAGATTATTACTGTGGACAGAGTTACAGCTATCCTCC |
| IGKV1-135*01_MRL | MRL/MpJ | Musmus IGKV1-135*01 F | 100 | 302 | GATGTTGTGATGACCCAGACTCCACTCACTTTGTCGGTTACCATTGGACAACCAGCCT<br>CCATCTCTTGCAAGTCAAGTCAGAGCCTCTTAGATAGTGATGGAAAGACATATTTGAA<br>TTGGTTGTTACAGAGGCCAGGCCAGTCTCCAAAGCGCCTAATCTATCTGGTGTCTAAA<br>CTGGACTCTGGAGTCCCTGACAGGTTCACTGGCAGTGGATCAGGGACAGATTTCACAC<br>TGAAAATCAGCAGAGTGGAGGCTGAGGATTTGGGAGTTTATTATTGCTGGCAAGGTAC<br>ACATTTTCCTCA |
| IGKV5-45*01_S0430_MRL | MRL/MpJ | Musmus IGKV5-45*01 F | 99.64 | 287 | GATATTGTGCTAACTCAGTCTCCAGCCACCCTGTCTGTGACTCCAGGAGATAGAGTCA<br>GTCTTTCCTGCAGGGCCAGCCAAAGTATTAGCAACTACCTACACTGGTATCAACAAAA<br>ATCACATGAGTCTCCAAGGCTTCTCATCAAGTATGCTTCCCAGTCCATCTCTGGGATCC<br>CCTCCAGGTTCAGTGGCAGTGGATCAGGGACAGATTTCACTCTCAGTATCAACAGTGT<br>GGAGACTGAAGATTTTGGAATGTATTTCTGTCAACAGAGTAACAGCTGGCCTCA |
| IGKV2-137*01_MRL | MRL/MpJ | Musmus IGKV2-137*01 F | 100 | 302 | GATATTGTGATGACTCAGGCTGCACCCTCTGTACCTGTCACTCCTGGAGAGTCAGTAT<br>CCATCTCCTGCAGGTCTAGTAAGAGTCTCCTGCATAGTAATGGCAACACTTACTTGTAT<br>TGGTTCCTGCAGAGGCCAGGCCAGTCTCCTCAGCTCCTGATATATCGGATGTCCAACC<br>TTGCCTCAGGAGTCCCAGACAGGTTCAGTGGCAGTGGGTCAGGAACTGCTTTCACACT<br>GAGAATCAGTAGAGTGGAGGCTGAGGATGTGGGTGTTTATTACTGTATGCAACATCTA<br>GAATATCCTTT |
| IGKV12-46*01_S4026_MRL | MRL/MpJ | Musmus IGKV12-46*01 F | 99.64 | 287 | GACATCCAGATGACTCAGTCTCCAGCCTCCCTATCTGTATCTGTGGGAGAAACTGTCA<br>CCATCACATGTCGAGCAAGTGAGAATATTTACAGTAATTTAGCATGGTATCAGCAGAA<br>ACAGGGAAAATCTCCTCAGCTCCTGGTCTATGCTGCAACAAACTTAGCAGATGGTGTT<br>CCATCAAGGTTCAGTGGCAGTGGATCAGGCACACAGTATTCCCTCAAGATCAACAGCC<br>TGCAGTCTGAAGATTTTGGGAGTTATTACTGTCAACATTTTTGGGGTACTCCTCC |

| | | | | | |
|---|---|---|---|---|---|
| IGKV8-21*01_S1419_MRL | MRL/MpJ | Musmus IGKV8-21*01 F | 99.33 | 302 | GACATTGTGATGTCACAGTCTCCATCCTCCCTGGCTGTGTCAGCAGGAGAGAAGGTCACTATGAGCTGCAAATCCAGTCAGAGTCTGTTCAACAGTAGAACCCGAAAGAACTACTTGGCTTGGTACCAGCAGAAACCAGGGCAGTCTCCTAAACTGCTGATCTACTGGGCATCCACTAGGGAATCTGGGGTCCCTGATCGCTTCACAGGCAGTGGATCTGGGACAGATTTCACTCTCACCATCAGCAGTGTGCAGGCTGAAGACCTGGCAGTTTATTACTGCAAGCAATCTTATTATCTTCC |
| IGKV12-46*01_S4541_MRL | MRL/MpJ | Musmus IGKV12-46*01 F | 97.13 | 287 | GACATCCAGATGACTCAGTCTCCAGCCTCCCTATCTGCATCTGTGGGAGAAACTGTCACCATCACATGTCGAGCAAGTGAGAATATTTACAGTAATTTAGCATGGTATCAGCAGAAACAGGGAAAATCTCCTCAGCTCCTGGTCTATGCTGCAACAAATTTAGCAGATGGTGTGCCATCAAGGTTCAGTGGCAGTGGATCAGGCACACAGTTTTCTCTGAAGATCAACAGCCTGCAGCCTGAAGATTTTGGGAGTTATTACTGTCAACATTTTTATGGTACTCCTCC |
| IGKV8-19*01_MRL | MRL/MpJ | Musmus IGKV8-19*01 F | 100 | 305 | GACATTGTGATGACACAGTCTCCATCCTCCCTGACTGTGACAGCAGGAGAGAAGGTCACTATGAGCTGCAAGTCCAGTCAGAGTCTGTTAAACAGTGGAAATCAAAAGAACTACTTGACCTGGTACCAGCAGAAACCAGGGCAGCCTCCTAAACTGTTGATCTACTGGGCATCCACTAGGGAATCTGGGGTCCCTGATCGCTTCACAGGCAGTGGATCTGGAACAGATTTCACTCTCACCATCAGCAGTGTGCAGGCTGAAGACCTGGCAGTTTATTACTGTCAGAATGATTATAGTTATCCTCC |
| IGKV12-46*01_S1254_MRL | MRL/MpJ | Musmus IGKV12-46*01 F | 94.62 | 287 | GACATCCAGATGACTCAGTCTCCAGCTTCCCTGTCTGCATCTGTGGGAGAAACTGTCACCATCACATGTCGAGCAAGTGAGAATATTGACAGTTATTTAGCATGGTATCAGCAGAAACAGGGAAAATCTCCTCAGCTCCTGGTCTATGCTGCAACACTCTTAGCAGATGGTGTGCCATCAAGGTTCAGTGGCAGTGGATCAGGCACACAGTATTCTCTCAAGATCAACAGCCTGCAGTCTGAAGATGTTGCGAGATATTACTGTCAACATTATTATAGTACTCCTCC |
| IGKV6-15*01_S9094_MRL | MRL/MpJ | Musmus IGKV6-15*01 F | 96.06 | 287 | GACATTGTGATGACCCAGTCTCAAAAATTCATGTCCACATCAGTAGGAGACAGGGTCAGCGTCACCTGCAAGGCCAGTCAGTATGTGGGTACTTATGTAGCCTGGTATCAACAGAAACCAGGACAATCTCCTAAAGCACTGATTTACTCGGCATCCACCCGGCACACTGGAGTCCCTGATCGCTTCACAGGCAGTGGATCTGGGACAGATTTCACTCTCACCATTAGCAATGTGCAGTCTGAAGACTTGGCAGAGTATTTCTGTGAGCAATACAGCAGCTCTCCTCT |
| IGKV1-110*01_MRL | MRL/MpJ | Musmus IGKV1-110*01 F | 100 | 302 | GATGTTGTGATGACCCAAACTCCACTCTCCCTGCCTGTCAGTCTTGGAGATCAAGCCTCCATCTCTTGCAGATCTAGTCAGAGCCTTGTACACAGTAATGGAAACACCTATTTACATTGGTACCTGCAGAAGCCAGGCCAGTCTCCAAAGCTCCTGATCTACAAAGTTTCCAACCGATTTTCTGGGGTCCCAGACAGGTTCAGTGGCAGTGGATCAGGGACAGATTTCACACTCAAGATCAGCAGAGTGGAGGCTGAGGATCTGGGAGTTTATTTCTGCTCTCAAAGTACACATGTTCCTCC |
| IGKV6-b*01_MRL | MRL/MpJ | Musmus IGKV6-b*01 F | 100 | 287 | AGTATTGTGATGACCCAGACTCCCAAATTCCTGCCTGTATCAGCAGGAGACAGGGTTACCATGACCTGCAAGGCCAGTCAGAGTGTGGGTAATAATGTAGCCTGGTACCAACAGAAGCCAGGACAGTCTCCTAAACTGCTGATATACTATGCATCCAATCGCTACACTGGAGTCCCTGATCGCTTCACTGGCAGTGGATCTGGGACAGATTTCACTTTCACCATCAGCAGTGTGCAGGTTGAAGACCTGGCAGTTTATTTCTGTCAGCAGCATTATAGCTCTCCTCC |
| IGKV12-44*01_MRL | MRL/MpJ | Musmus IGKV12-44*01 F | 100 | 287 | GACATCCAGATGACTCAGTCTCCAGCCTCCCTATCTGCATCTGTGGGAGAAACTGTCACCATCACATGTCGAGCAAGTGAGAATATTTACAGTTATTTAGCATGGTATCAGCAGAAACAGGGAAAATCTCCTCAGCTCCTGGTCTATAATGCAAAAACCTTAGCAGAAGGTGTGCCATCAAGGTTCAGTGGCAGTGGATCAGGCACACAGTTTTCTCTGAAGATCAACAGCCTGCAGCCTGAAGATTTTGGGAGTTATTACTGTCAACATCATTATGGTACTCCTCC |

| | | | | | |
|---|---|---|---|---|---|
| IGKV1-117*01_MRL | MRL/MpJ | Musmus IGKV1-117*01 F | 100 | 302 | GATGTTTTGATGACCCAAACTCCACTCTCCCTGCCTGTCAGTCTTGGAGATCAAGCCTCCATCTCTTGCAGATCTAGTCAGAGCATTGTACATAGTAATGGAAACACCTATTTAGAATGGTACCTGCAGAAACCAGGCCAGTCTCCAAAGCTCCTGATCTACAAAGTTTCCAACCGATTTTCTGGGGTCCCAGACAGGTTCAGTGGCAGTGGATCAGGGACAGATTTCACACTCAAGATCAGCAGAGTGGAGGCTGAGGATCTGGGAGTTTATTACTGCTTTCAAGGTTCACATGTTCCTCC |
| IGKV6-23*01_S4960_MRL | MRL/MpJ | Musmus IGKV6-23*01 F | 92.11 | 287 | GACATTGTGATGACCCAGTCTCAAAAATTCATGTCCACATCAGTAGGAGAGAGGGTCAGCATCACCTGCAAGGCCAGTCAGAATGTAGGTACTAATGTAGCCTGGTATCAGCAGAAAGCAGGGCAGTCTCTTGAACTGCTGATCTATGGGGCATCCAACCGGCACACTGGAGTCCCTGATCGCTTCACAGGCAGTGGATCTGGGACAGATTTCACTCTCACCATCACCAATGTGCAGTCTGAAGACATGACAAATTATTTCTGTGAGCAATATAGCAGCTATCCTCT |
| IGKV12-41*01_S6800_MRL | MRL/MpJ | Musmus IGKV12-41*01 F | 97.13 | 287 | GATATCCAGATGACTCAGTCTCCAGCCTCCCTATCTGCATCTGTGGGAGAAACTGTCACCATCACATGTCGAGCAAGTGGGAATATTCACAATTATTTAGCATGGTATCAGCAGAAACAGGGAAAATCTCCTCAGCTCCTGGTCTATAATGCAAAAACCTTAGCGGAAGGTGTGCCATCAAGGTTCAGTGGCAGTGGATCAGGAACACAATATTCTCTCAAGATCAACAGCCTGCAGCCTGAGGATTTTGGGAGTTATTACTGTCATCATTATTATAGTACTCCTCC |
| IGKV3-1*01_S7345_MRL | MRL/MpJ | Musmus IGKV3-1*01 F | 98.63 | 299 | GACATTGTGCTCACCCAATCTCCAGCTTCTTTGGCTGTGTCTCTAGGGCAGAGTGTCACCATCTCCTGCAGAGCCAGTGAAAGTGTTGAATATTATGGCACTAGTTTAATGCAGTGGTACCAACAGAAACCAGGACAGCCACCCAAACTCCTCATCTATGGTGCATCCAACGTAGAATCTGGGGTCCCTGCCAGGTTTAGTGGCAGTGGGTCTGGGACAGACTTCAGCCTCAACATCCATCCTGTGGAGGAGGATGATATTGCAATGTATTTCTGTCAGCAAAGTAGGAAGGTTCCTTC |
| IGKV5-43*01_S9511_MRL | MRL/MpJ | Musmus IGKV5-43*01 F | 99.64 | 287 | GATATTGTGCTAACTCAGTCTCCAGCCACCCTGTCTGTGACTCCAGGAGATAGCGTCAGTCTTTCCTGCAGGGCCAGCCAAAGTATTAGCAACAACCTACACTGGTATCAACAAAAATCACATGAGTCTCCAAGGCTTCTCATCAAGTATGCTTCCCAGTCCATCTCTGGGATCCCCTCCAAGTTCAGTGGCAGTGGATCAGGGACAGATTTCACTCTCAGTATCAACAGTGTGGAGACTGAAGATTTTGGAATGTATTTCTGTCAACAGAGTAACAGCTGGCCTCA |
| IGKV10-94*03_MRL | MRL/MpJ | Musmus IGKV10-94*03 F, or Musmus IGKV10-94*08 F | 100 | 287 | GATATCCAGATGACACAGACTACATCCTCCCTGTCTGCCTCTCTGGGAGACAGAGTCACCATCAGTTGCAGGGCAAGTCAGGATATTAGCAATTATTTAAACTGGTATCAGCAGAAACCAGATGGAACTGTTAAACTCCTGATCTACTACACATCAAGATTACACTCAGGAGTCCCATCAAGGTTCAGTGGCAGTGGGTCTGGGACAGATTATTCTCTCACCATCAGCAACCTGGAACCTGAAGATATTGCCACTTACTATTGTCAGCAGTATAGTAAGCTTCCTCC |
| IGKV4-74*01_S3301_MRL | MRL/MpJ | Musmus IGKV4-74*01 F | 94.68 | 292 | CAGATTGTTCTCACCCAGTCTCCAGCAATCATGTCTGCATCTCCAGGGGAGAAGGTCACCATGACCTGCAGGGCCAGCTCAAGTGTAAGTTCCAGTTACTTGCACTGGTACCAGCAGAAGCCAGGATCTTCCCCCAAACTCTGGATTTATAGCACATCCAACCTGGCTTCAGGAGTCCCAGCTCGCTTCAGTGGCAGTGGGTCTGGGACCTCTTACTCTCTCACAATCAGCAGTGTGGAGGCTGAGGATGCTGCCACTTATTACTGCCAGCAGTATGATAGTTCCCCACCCA |
| IGKV8-24*01_S8843_MRL | MRL/MpJ | Musmus IGKV8-24*01 F | 98.32 | 305 | GACATTGTGATGACACAGTCTCCATCCTCCCTGGCTATGTCAGTAGGACAGAAGGTCACTATGAGCTGCAAGTCCAGTCAGAGCCTTTTAAGTAGTAGCAATCAAAAGAACTATTTGGCCTGGTACCAGCAGAAACCAGGACAGTCTCCTAAACTTCTGGTATACTTTGCATCCACTAGGGAATCTGGGGTCCCTGATCGCTTCATAGGCAGTGGATCTGGGACAGATTTCACTCTTACCATCAGCAGTGTGCAGGCTGAAGACCTGGCACTTTATTACTGTCAGCAACATTATAGCACTCCTCC |

| | | | | | |
|---|---|---|---|---|---|
| IGKV4-56*01_S5696_MRL | MRL/MpJ | Musmus IGKV4-56*01 P | 96.38 | 286 | CAAATTGTTCTCACCCAGTCTCCAGCAATCATGTCTGCATCTCCAGGGCAGAAAGTCACCATAACCTGCAGTGCCAGCTCAAGTGTAAATTACATGCACTGGTACCAGCAGAAGCTAGGATCCTCCCCCAAACTCTGGATTTATGACACATCCAAACTGGCTCCTGGAGTCCCTGCTCGCTTCAGTGGCAGTGGGTCTGGGACCTCTTACTCTCTCACAATCAGCAGCATGGAGGCTGAAGATGCTGCCTCTTATTTCTGCCATCAGTGGAGTAGTTACCCACCCA |
| IGKV14-126*01_MRL | MRL/MpJ | Musmus IGKV14-126*01 F | 100 | 287 | GACATCAAGATGACCCAGTCTCCATCCTCCATGTATGCATCGCTGGGAGAGAGAGTCACTATCACTTGCAAGGCGAGTCAGGACATTAAAAGCTATTTAAGCTGGTACCAGCAGAAACCATGGAAATCTCCTAAGACCCTGATCTATTATGCAACAAGCTTGGCAGATGGGGTCCCATCAAGATTCAGTGGCAGTGGATCTGGGCAAGATTATTCTCTAACCATCAGCAGCCTGGAGTCTGACGATACAGCAACTTATTACTGTCTACAGCATGGTGAGAGCCCTCC |
| IGKV4-81*01_S7273_MRL | MRL/MpJ | Musmus IGKV4-81*01 F | 96.45 | 290 | GAAAATGTGCTGACCCAGTCTCCAGCAATCATGGCTGCATCTCCAGGGGAGAAGGTCACCATGACCTGCAGTGCCAGCTCAAGTGTAAGTTCTGGTAACTTTCACTGGTACCAGCAGAAGCCAGGCACTTCTCCCAAACTCTGGATTTATAGGACATCCAACCTGGCTTCTGGAGTCCCCGCTCGCTTCAGTGGCAGTGGGTCTGGGACCTCTTACTCTCTTACAATCAGCAGCATGGAGGCCGAAGATGCTGCCACTTATTACTGCCAGCAGTGGAGTGGTTACCCACC |
| IGKV3-4*01_S8857_MRL | MRL/MpJ | Musmus IGKV3-4*01 F | 97.25 | 299 | GACATTGTGCTGACCCAATCTCCAGCTTCTTTGGCTGTGTCTCTAGGGCAGAGGGCCACCATCTCCTGCAGAGCCAGCGAAAGTGTTGATTATGATGGCGATAGTTATATGAACTGGTACCAACAGAAACCAGGACAGCCACCGAAACTCCTCATCTATGCTGCATCCAATCTAGAATCTGGGATCCCTGCCAGGTTTAGTGGCAGTGGGTCTGGGACAGACTTCACCCTCAACATTCATCCTGTGGAGGAGGAGGATGCTGCAACCTATTACTGTCTGCAAAGTAATGAGGCTCCTCC |
| IGKV15-103*01_MRL | MRL/MpJ | Musmus IGKV15-103*01 ORF | 100 | 287 | GACATCCAGATGAACCAGTCTCCATCCAGTCTGTCTGCATCCCTTGGAGACACAATTACCATCACTTGCCATGCCAGTCAGAACATTAATGTTTGGTTAAGCTGGTACCAGCAGAAACCAGGAAATATTCCTAAACTATTGATCTATAAGGCTTCCAACTTGCACACAGGCGTCCCATCAAGGTTTAGTGGCAGTGGATCTGGAACAGGTTTCACATTAACCATCAGCAGCCTGCAGCCTGAAGACATTGCCACTTACTACTGTCAACAGGGTCAAAGTTATCCTCT |
| IGKV3-4*01_S3473_MRL | MRL/MpJ | Musmus IGKV3-4*01 F | 92.44 | 299 | GACATTGTGCTGACCCAATCTCCAGCTTCTTTGGCTGTGTCTCTAGGGCAGAGGGCCACCATATCCTGCAAGGCCAGCCAGAGTATCAATTATTACGGTGATAATTATATGCACTGGTTCCAACAGAAACCAGGACAGCCAACCAAACTCCTTATCTATGATGCATCCAACCTAGAATCTGGAGTTCCTGACAGGTTCAGTGGCAGTGGGTCTGGGACAGACTTCACCCTCAACATCCATCCTGTGGAGGAGGAGGATGCTGCAACCTATTACTGTCAGCAAAGCAAGGAGTTTCCTCC |
| IGKV4-57-1*01_S8603_MRL | MRL/MpJ | Musmus IGKV4-74*01 F | 95.04 | 292 | CAAATTGTTCTCACCCAGTCTCCAGCAATCATGTCTGCCTCTCCAGGGGAGAAGGTCACCATGACCTGCAGTGCCAGCTCAAGTGTAAGTTCCAGGTACTTGCACTGGTACCAGCAGAAGTCAGGAGCCTCCCCCAAACTCTGGATTTATGGCACATCCAACCTGGCTTCTGGAGTCCCTGCTCGCTTCAGTGGCAGTGGGTCTGGGACCTCTTACTCTCTCACAATCAGCAGCGTGGAGGCTGAAGATGCTGCCACTTATTACTGCCAGCAGTATCATAGTGACCCACTCA |
| IGKV19-93*01_S8964_MRL | MRL/MpJ | Musmus IGKV19-93*01 F | 98.21 | 287 | GACATCCAGATGACACAGTCTCCATCCTCACTGTCTGCATCTCTGGGAGGCAAAGTCACCATCACTTGCAAGGCAAGCCAAGACATTAACAAGAATATAGCTTGGTACCAACACAAGCCTGGAAAAGGTCCTAGGCTGCTCATATGGTACACATCTACATTACAGCCAGGCATCCCATCAAGGTTCAGTGGAAGTGGGTCTGGGAGAGATTATTCCTTCAGCATCAGCAACCTGGAGCCTGAAGATATTGCAACTTATTACTGTCTACAGTATGATAATCTTCCACC |

| | | | | | |
|---|---|---|---|---|---|
| IGKV4-53*01_S4588_MRL | MRL/MpJ | Musmus IGKV4-53*01 F | 98.94 | 292 | GAAATTGTGCTCACCCAGTCTCCAGCACTCATGGCTGCATCTCCAGGGGAGAAGGTCACCATCACCTGCAGTGTCAGCTCAAGTATAAGTTCCAGCAACTTACACTGGTACCAGCAGAAGTCAGGAACCTCCCCCAAACCCTGGATTTATGGCACATCCAACCTTGCTTCTGGAGTCCCTGTTCGCTTCAGTGGCAGTGGATCTGGGACCTCTTATTCTCTCACAATCAGCAGCATGGAGGCTGAAGATGCTGCCACTTATTACTGTCAACAGTGGAGTAGTTACCCGCTCA |
| IGKV5-39*01_S3142_MRL | MRL/MpJ | Musmus IGKV5-39*01 F | 98.92 | 287 | GACATTGTGATGACTCAGTCTCCAGCCACCCTGTCTGTGACTCCAGGAGATAGGGTCTCTCTTTCCTGCAGGGCCAGTCAGAGTATTAGCGACTACTTACACTGGTATCAACAAAAATCACATGAGTCTCCAAGGCTTCTCATCAAGTATGCTTCCCAATCCATCTCTGGGATCCCCTCCAGGTTCAGTGGCAGTGGATCAGGGTCAGATTTCACTCTCAGTATCAACAGTGTGGAACCTGAAGATGTTGGAGTGTATTACTGTCAAAATGGTCACAGCTTTCCTCC |
| IGKV4-50*01_S2188_MRL | MRL/MpJ | Musmus IGKV4-50*01 F | 97.1 | 286 | GAAAATGTGCTCACCCAGTCTCCAGCAATCATGTCTGCAACTCTAGGGGAGAAGGTCACCATGAGCTGCAGGGCCAGCTCAAATGTAAAGTACATGTACTGGTACCAGCAGAAGTCAGGTGCCTCCCCCAAACTATGGATTTATTACACATCCAACCTGGCTTCTGGAGTCCCAGCTCGCTTCAGTGGCAGTGGGTCTGGGACCTCTTATTCTCTCACAATCAGCAGCGTGGAGGCTGAAGATGCTGCCACTTATTACTGCCAGCAGTTTACTAGTTCCCCATCCA |
| IGKV8-34*01_S3615_MRL | MRL/MpJ | Musmus IGKV8-34*01 F | 99.33 | 305 | GACATTTTGATGACTCAGTCTCCATCCTCCCTGACTGTGTCAGCAGGAGAGAAGGTCACTATGAGCTGCAAGTCCAGTCAGAGTCTTTTAGCTAGTGCCAACCAAAATAACTACTTGGCCTGGCACCAGCAGAAACCAGGACGATCTCCTAAAATGCTGATAATTTGGGCATCCACTAGGGTATCTGGAGTCCCTGATCGCTTCATAGGCAGTGGATCTGGGACGGATTTCACTCTGACCATCAACAGTGTGCAGGCTGAAGATCTGGCTGTTTATTACTGTCAGCAGTCCTACAGCGCTCCTAC |
| IGKV14-100*01_S4794_MRL | MRL/MpJ | Musmus IGKV14-100*01 F | 99.64 | 287 | GACATCCTGATGACCCAATCTCCATCCTCCATGTCTGTATCTCTGGGAGACACAGTCAGCATCACTTGCCATGCAAGTCAGGGCATTAGCAGTAATATAGGGTGGTTGCAGCAGAAACCAGGGAAATCATTTAAGGGCCTGATCTATCATGCAACCAACTTGGAAGATGGAGTTCCATCAAGGTTCAGTGGCAGTGGATCTGGAGCAGATTATTCTCTCACCATCAGCAGCCTGGAATCTGAAGATTTTGCAGACTATTACTGTGTACAGTATGCTCAGTTTCCTCC |
| IGKV4-62*01_S2199_MRL | MRL/MpJ | Musmus IGKV4-62*01 ORF | 96.74 | 286 | CAAATTGTTCTCACCCAGTCTCCAGCAATCATGTCTGCATCTCCAGGGGAGAAGGTCACCATGACCTGCAGTGCCAGCTCAAGTGTAAGTTACATGTACTGGTACCAGCAGAAGCCAGGATCCTCCCCCAGACTCTGGATTTATGACACATCCAACCTGGCTTCTGGAGTCCCCGCTCGCTTCAGTGGCAGTAGGTCTGGGACCTCTTATTCTCTCACAATCAGCAGCATGGAGGCTGAAGATGCTGCCACTTATTACTGCCATCAGCGGAGTAGTTACCCATTCA |
| IGKV16-104*01_MRL | MRL/MpJ | Musmus IGKV16-104*01 F | 100 | 288 | GATGTCCAGATAACCCAGTCTCCATCTTATCTTGCTGCATCTCCTGGAGAAACCATTACTATTAATTGCAGGGCAAGTAAGAGCATTAGCAAATATTTAGCCTGGTATCAAGAGAAACCTGGGAAAACTAATAAGCTTCTTATCTACTCTGGATCCACTTTGCAATCTGGAATTCCATCAAGGTTCAGTGGCAGTGGATCTGGTACAGATTTCACTCTCACCATCAGTAGCCTGGAGCCTGAAGATTTTGCAATGTATTACTGTCAACAGCATAATGAATACCCGTAC |
| IGKV3-7*01_S6705A_MRL | MRL/MpJ | Musmus IGKV3-7*02 F | 97.59 | 299 | GACATTGTGCTGACACAGTCTCCTGCTTCCTTAGCTGTATCTCTGGGGCAGAGGGCCACCATCTCCTGCAGGGCCAGCAAAAGTGTCAGTACATCTAGCTATAGTTACATGCACTGGTACCAACAGAAACCAGGACAGCCACCCAAACTCCTCATCAAGTATGCATCCTACCTAGAATCTGGGGTTCCTGCCAGGTTCAGTGGCAGTGGGTCTGGGACAGACTTCACCCTCAACATCCATCCTGTGGAGGAGGAGGATGCTGCAACATATTACTGTCAGCACAGTAGGGAGTTTCCTCC |

| | | | | | |
|---|---|---|---|---|---|
| IGKV9-120*01_S2628_MRL | MRL/MpJ | Musmus IGKV9-120*02 F | 100 | 285 | GACATCCAGATGACCCAGTCTCCATCCTCCTTATCTGCCTCTCTGGGAGAAAGAGTCAGTCTCACTTGTCGGGCAAGTCAGGACATTGGTAGTAGCTTAAACTGGCTTCAGCAGGAACCAGATGGAACTATTAAACGCCTGATCTACGCCACATCCAGTTTAGATTCTGGTGTCCCCAAAAGGTTCAGTGGCAGTAGGTCTGGGTCAGATTATTCTCTCACCATCAGCAGCCTTGAGTCTGAAGATTTTGTAGACTATTACTGTCTACAATATGCTAGTTCTCCT |
| IGKV4-80*01_S4247_MRL | MRL/MpJ | Musmus IGKV4-80*01 F | 99.64 | 282 | CAAATTGTTCTCACCCAGTCTCCAGCAATCATGTCTGCATCTCTAGGGGAGGAGATCACCCTAACCTGCAGTGCCAGCTCGAGTGTAAGTTACATGCACTGGTTCCAGCAGAAGTCAGGCACTTCTCCCAAACTCTTGATTTATAGCACATCCAACCTGGCTTCTGGAGTCCCTTCTCGCTTCAGTGGCAGTGGGTCTGGGACCTTTTATTCTCTCACAATCAGCAGTGTGGAGGCTGAAGATGCTGCCGATTATTACTGCCATCAGTGGAGTAGTTATCCA |
| IGKV10-96*03_MRL | MRL/MpJ | Musmus IGKV10-96*03 F | 100 | 287 | GATATCCAGATGACACAGACTACATCCTCCCTGTCTGCCTCTCTGGGAGACAGAGTCACCATCAGTTGCAGGGCAAGTCAGGACATTAGCAATTATTTAAACTGGTATCAGCAGAAACCAGATGGAACTGTTAAACTCCTGATCTACTACACATCAAGATTACACTCAGGAGTCCCATCAAGGTTCAGTGGCAGTGGGTCTGGGACAGATTATTCTCTCACTATTAGCAACCTGGAACAAGAAGATATTGCCACTTACTTTTGCCAACAGGATAGTAAGCATCCTCC |
| IGKV4-59*01_S3489_MRL | MRL/MpJ | Musmus IGKV4-59*01 F | 98.19 | 286 | CAAATTGTTCTCACCCAGTCTCCAGCAATCATGTCTGCATCTCCAGGGGAGAAGGTCACCATGACATGCAGTGCCAGCTCAAGTGTAAGTTACATGCACTGGTACCAGCAGAAGTCAGGCACCTCCCCCAAAAGATGGATTTATGACACATCCAAACTGGATTCTGGAGTCCCTGCTCGCTTCAGTGGCAGTGGGTCTGGGACCTCTTATTCTCTCACAATCAGCAGCATGGAGGCTGAAGATGCTGCCACTTATTACTGCTATCAGTGGAGTAGTTACCCACTCA |
| IGKV8-30*01_S3861_MRL | MRL/MpJ | Musmus IGKV8-30*01 F | 95.62 | 302 | GACATTGTGATGTCACAGTCTCCATCCGCCCTAGCTGTGTCAGTTGGAGAGAAGGTCACTATGAGCTGCAAGTCCAGTCAGAGCCTTTTATATAGTAGCAATCAAAAGAACTACTTGGCCTGGTACCAGCAGAAACCAGGGCAGTCTCCTAAACTGTTAATCTACTGGGCATCCACTAGGGAATCTGGGGTCCCTGACCGCTTCACAGGCAGTAGATCAGGGACAGATTTCACTCTCACCATCAGCAGTGTGCAGGCTGAAGACCTGGCCGTTTATTACTGCAAGCAATCTTATAGCTATCC |
| IGKV2-112*01_MRL | MRL/MpJ | Musmus IGKV2-112*01 F | 100 | 300 | GATATTGTGATAACCCAGGATGAACTCTCCAATCCTGTCACTTCTGGAGAATCAGTTTCCATCTCCTGCAGGTCTAGTAAGAGTCTCCTATATAAGGATGGGAAGACATACTTGAATTGGTTTCTGCAGAGACCAGGACAATCTCCTCAGCTCCTGATCTATTTGATGTCCACCCGTGCATCAGGAGTCTCAGACCGGTTTAGTGGCAGTGGGTCAGGAACAGATTTCACCCTGGAAATCAGTAGAGTGAAGGCTGAGGATGTGGGTGTGTATTACTGTCAACAACTTGTAGAGTATCCT |
| IGKV17-127*01_MRL | MRL/MpJ | Musmus IGKV17-127*01 F | 100 | 287 | GAAACAACTGTGACCCAGTCTCCAGCATCCCTGTCCGTGGCTACAGGAGAAAAAGTCACTATCAGATGCATAACCAGCACTGATATTGATGATGATATGAACTGGTACCAGCAGAAGCCAGGGGAACCTCCTAAGCTCCTTATTTCAGAAGGCAATACTCTTCGTCCTGGAGTCCCATCCCGATTCTCCAGCAGTGGCTATGGCACAGATTTTGTTTTTACAATTGAAAACACGCTCTCAGAAGATGTTGCAGATTACTACTGTTTGCAAAGTGATAACATGCCTCT |
| IGKV14-111*01_MRL | MRL/MpJ | Musmus IGKV14-111*01 F | 100 | 287 | GACATCAAGATGACCCAGTCTCCATCTTCCATGTATGCATCTCTAGGAGAGAGAGTCACTATCACTTGCAAGGCGAGTCAGGACATTAATAGCTATTTAAGCTGGTTCCAGCAGAAACCAGGGAAATCTCCTAAGACCCTGATCTATCGTGCAAACAGATTGGTAGATGGGGTCCCATCAAGGTTCAGTGGCAGTGGATCTGGGCAAGATTATTCTCTCACCATCAGCAGCCTGGAGTATGAAGATATGGGAATTTATTATTGTCTACAGTATGATGAGTTTCCTCC |

| | | | | | |
|---|---|---|---|---|---|
| IGKV2-109*01_MRL | MRL/MpJ | Musmus IGKV2-109*01 F | 100 | 302 | GATATTGTGATGACGCAGGCTGCATTCTCCAATCCAGTCACTCTTGGAACATCAGCTTCCATCTCCTGCAGGTCTAGTAAGAGTCTCCTACATAGTAATGGCATCACTTATTTGTATTGGTATCTGCAGAAGCCAGGCCAGTCTCCTCAGCTCCTGATTTATCAGATGTCCAACCTTGCCTCAGGAGTCCCAGACAGGTTCAGTAGCAGTGGGTCAGGAACTGATTTCACACTGAGAATCAGCAGAGTGGAGGCTGAGGATGTGGGTGTTTATTACTGTGCTCAAAATCTAGAACTTCCTCC |
| IGKV4-78*01_S3978_MRL | MRL/MpJ | Musmus IGKV4-78*01 F | 97.87 | 292 | CAAATTGTTCTCACCCAGTCTCCAGCAATCATGTCTGCATCTCCTGGGGAACGGGTCACCATGACCTGCAGTGCCAGCTCAAGTGTAAGTTCCAGCTACTTGTACTGGTACCAGCAGAAGCCAGGATCCTCCCCCAAACTATGGATTTATAGCACATCCAACCTGGCTTCTGGAGTCCCTGCTCGCTTCAGTGGCAGTGGGTCTGGGACCTCTTATTCTCTCACAATCAGCAGCATGGAGGCTGAAGATGCTGCCACTTATTACTGCCAGCAGTACAGTGGTTACCCATCCA |
| IGKV3-10*01_S5437_MRL | MRL/MpJ | Musmus IGKV3-10*01 F | 98.97 | 299 | AAAATTGTGCTGACCCAATCTCCAGCTTCTTTGGCTGTGTCTCTAAGGCAGAGGGCCACCATATCCTGCAGAGCCAGTGAAAGTGTTGATAGTTATGGCAATAGTTTTATGCACTGGTACCAGCAGAAACCAGGACAGCCACCCAAACTCCTCATCTATCGTGCATCCAACCTAGAATCTGGGGTCCCTGCCAGGTTCAGTGGCAGTGGGTCTAGGACAGACTTCACCCTCACCATTGATCCTGTGGAGGCTGATGATGCTGCAACCTATTACTGTCAGCAAAATAATGAGGATCCTCC |
| IGKV17-121*01_MRL | MRL/MpJ | Musmus IGKV17-121*01 F | 100 | 287 | GAAACAACTGTGACCCAGTCTCCAGCATCCCTGTCCATGGCTATAGGAGAAAAAGTCACCATCAGATGCATAACCAGCACTGATATTGATGATGATATGAACTGGTACCAGCAGAAGCCAGGGGAACCTCCTAAGCTCCTTATTTCAGAAGGCAATACTCTTCGTCCTGGAGTCCCATCCCGATTCTCCAGCAGTGGCTATGGTACAGATTTTGTTTTTACAATTGAAAACATGCTCTCAGAAGATGTTGCAGATTACTACTGTTTGCAAAGTGATAACTTGCCTCT |
| IGKV4-91*01_S4811_MRL | MRL/MpJ | Musmus IGKV4-91*01 F | 95.74 | 292 | GAAATTGTGCTTACCCAGTCTCCAACCACCATGGCTGCATCTCCTGGGGAGAAGGTCACCATCACCTGCAGTGCCAGCTCAAGTATAAGTTCCAATTACTTGCACTGGTATCAGCAGAAGCCAGGATTCCCTCCTAAACTCTTGATATATAGGACATCCAATCTGGCTTCTGGAGTCCCAGCTCGCTTCAGTGGCAGTGGGTCTGGGACCTCTTACTCTCTCACAATTGGCACCATGGAGGCTGAAGGTGCTGCCACTTATTACTGCCAGCATGGTAGTAGTTTACTACGCA |
| IGKV4-53*01_S3575_MRL | MRL/MpJ | Musmus IGKV4-53*01 F | 98.94 | 292 | GAAATTGTGCTCACCCAGTCTCCAGCACTCATGGCTGCATCTCCAGGGGAGAAGGTCACCATCACCTGCAGTGTCAGCTCAAGTATAAGTTCCAGCAACTTACACTGGTACCAGCAGAAGTCAGGAACCTCCCCCAAACCCTGGATTTATGGCACATCCAACCTTGCTTCTGGAGTCCCTGTTCGCTTCAGTGGCAGTGGATCTGGGACCTCTTATTCTCTCACAATCAGCAGCATGGAGGCTGAAGATGCTGCCACTTATTACTGTCAACAGTGGAGTAGTTACCCACTCA |
| IGKV1-133*01_MRL | MRL/MpJ | Musmus IGKV1-133*01 F | 100 | 302 | GATGTTGTGATGACCCAGACTCCACTCACTTTGTCGGTTACCATTGGACAACCAGCCTCTATCTCTTGCAAGTCAAGTCAGAGCCTCTTATATAGTAATGGAAAAACCTATTTGAATTGGTTATTACAGAGGCCAGGCCAGTCTCCAAAGCGCCTAATCTATCTGGTGTCTAAACTGGACTCTGGAGTCCCTGACAGGTTCACTGGCAGTGGATCAGGAACAGATTTTACACTGAAAATCAGCAGAGTGGAGGCTGAGGATTTGGGAGTTTATTACTGCGTGCAAGGTACACATTTTCCTCA |

| | | | | | |
|---|---|---|---|---|---|
| IGKV4-74*01_S3912_MRL | MRL/MpJ | Musmus IGKV4-74*01 F | 95.04 | 292 | CAAATTGTTCTCACCCAGTCTCCAGCAATCATGTCTGCCTCTCCAGGGGAGAAGGTCACCATGACCTGCAGTGCCAGCTCAAGTGTAAGTTCCAGGTACTTGCACTGGTACCAGCAGAAGTCAGGAGCCTCCCCCAAACTCTGGATTTATGGCACATCCAACCTGGCTTCTGGAGTCCCTGCTCGCTTCAGTGGCAGTGGGTCTGGGACCTCTTACTCTCTCACAATCAGCAGCGTGGAGGCTGAAGATGCTGCCACTTATTACTGCCAGCAGTATCATAGTGACCCACCCA |
| IGKV11-125*01_MRL | MRL/MpJ | Musmus IGKV11-125*01 F | 100 | 287 | GATGTCCAGATGATTCAGTCTCCATCCTCCCTGTCTGCATCTTTGGGAGACATAGTCACCATGACTTGCCAGGCAAGTCAGGGCACTAGCATTAATTTAAACTGGTTTCAGCAAAAACCAGGGAAAGCTCCTAAGCTCCTGATCTATGGTGCAAGCAACTTGGAAGATGGGGTCCCATCAAGGTTCAGTGGCAGTAGATATGGGACAGATTTCACTCTCACCATCAGCAGCCTGGAGGATGAAGATATGGCAACTTATTTCTGTCTACAGCATAGTTATCTCCCTCC |
| IGKV13-85*01_S2277_MRL | MRL/MpJ | Musmus IGKV13-85*01 F | 97.13 | 286 | GACATCCAGATGACACAATCTTCATCCTCCTTGTCTGTATCTCTAGGAGACAGAGTCACCATTACTTGCAAGGCAAGTGAGCACATTAATAGTTGGTTAGCCTGGTATCAGCAAAAACCAGGAAATGCTCCTAGGCTCTTAATATCTGGTGCAACCAGTTTGGAAACTGGGGTTCCTTCAAGATTCAGTGGCAGTGCATCTGGAAAGGATTACACTCTCAGCATTACTAGTCTTCAGACTGAAGATGTTGCTACTTATTACTGTCAACAGTATTGGGGTACTCCTC |
| IGKV8-28*02_MRL | MRL/MpJ | Musmus IGKV8-28*02 F | 100 | 302 | GACATTGTGATGACACAGTCTCCATCCTCCCTGAGTGTGTCAGCAGGAGATAAGGTCACTATGAGCTGCAAGTCCAGTCAGAGTCTGTTAAACAGTAGAAACCAAAAGAACTACTTGGCCTGGTACCAGCAGAAACCATGGCAGCCTCCTAAACTGCTGATCTACGGGGCATCCACTAGGGAATCTGGGGTCCCTGATCGCTTCACAGGCAGTGGATCTGGAACAGATTTCACTCTCACCATCAGCAGTGTGCAGGCTGAAGACCTGGCAGTTTATTACTGTCAGAATGATTATAGTTATCC |
| IGKV12-38*01_S2611_MRL | MRL/MpJ | Musmus IGKV12-38*01 F | 98.21 | 287 | GACATCCAGATGACTCAGTCTCCAGCCTCCCTGGCTGCATCTGTGGGAGAAACCATCACCATCACATGTCAAGCAAGTGAGAACATTTACTTCAGTTTAGCATGGTATCAGCAGAAGCAAGGGAAATCTCCTCAGCTCCTGATCTATAATGCAAACAGCTTGGAAGATGGTGTCCCATCGAGGTTCAGTGGCAGTGGATCTGGGACACAGTATTCTATGAAGATCAACAGCATGCAGCCTGAAGATACTGCAACTTATTTCTGTAAACAGGCTTATGACTTTCCTCC |
| IGKV1-122*01_MRL | MRL/MpJ | Musmus IGKV1-122*01 F | 100 | 302 | GATGCTGTGATGACCCAAACTCCACTCTCCCTGCCTGTCAGTCTTGGAGATCAAGCCTCCATCTCTTGCAGGTCTAGTCAGAGCCTTGAAAACAGTAATGGAAACACCTATTTGAACTGGTACCTCCAGAAACCAGGCCAGTCTCCACAGCTCCTGATCTACAGGGTTTCCAACCGATTTTCTGGGGTCCTAGACAGGTTCAGTGGTAGTGGATCAGGGACAGATTTCACACTGAAAATCAGCAGAGTGGAGGCTGAGGATTTGGGAGTTTATTTCTGCCTCCAAGTTACACATGTCCCTCC |
| IGKV8-16*01_S1056_MRL | MRL/MpJ | Musmus IGKV8-16*01 F | 98.99 | 305 | GAAATTGTGTTGACACAGTCTATACCATCCCTGACTGTGTCAGCAGGAGAGAGGGTCACTATCAGCTGCAAATCCAATCAGAATCTTTTATGGAGTGGAAACCAACGGTACTGTTTGGTCTGGCACCAGTGGAAACCAGGGCAAACTCCTACACCGTTGATCACCTGGACATCTGATAGGTACTCTGGAGTCCCTGATCGTTTCATAGGCAGTGGATCTGTGACAGATTTCACTCTGACCATCAGCAGTGTGCAGGATGAAGATGTGGCAGTTTATTTCTGTCAGCAGCATTTACACATTCCTCC |

| | | | | | |
|---|---|---|---|---|---|
| IGKV8-16*01_S7495_MRL | MRL/MpJ | Musmus IGKV8-16*01 F | 99.33 | 305 | GAAATTGTGTTGACGCAGTCTATACCATCCCTGACTGTGTCAGCAGGAGAGAGGGTCACTATCAGCTGCAAATCCAATCAGAATCTTTTATGGAGTGGAAACCAAAGGTACTGTTTGGTCTGGCACCAGTGGAAACCAGGGCAAACTCCTACACCGTTGATCACCTGGACATCTGATAGATACTCTGGAGTCCCTGATCGTTTCATAGGCAGTGGATCTGTGACAGATTTCACTCTGACCATCAGCAGTGTGCAGGCTGAAGATGTGGCAGTTTATTTCTGTCAGCAGCATTTACACATTCCTCC |
| IGKV9-124*01_MRL | MRL/MpJ | Musmus IGKV9-124*01 F | 100 | 285 | GACATCCAGATGACCCAGTCTCCATCCTCCTTATCTGCCTCTCTGGGAGAAAGAGTCAGTCTCACTTGTCGGGCAAGTCAGGAAATTAGTGGTTACTTAAGCTGGCTTCAGCAGAAACCAGATGGAACTATTAAACGCCTGATCTACGCCGCATCCACTTTAGATTCTGGTGTCCCAAAAAGGTTCAGTGGCAGTAGGTCTGGGTCAGATTATTCTCTCACCATCAGCAGCCTTGAGTCTGAAGATTTTGCAGACTATTACTGTCTACAATATGCTAGTTATCCT |
| IGKV4-86*01_S4316_MRL | MRL/MpJ | Musmus IGKV4-86*01 F | 100 | 283 | GAAATTGTGCTCACTCAGTCTCCAGCCATCACAGCTGCATCTCTGGGGCAAAAGGTCACCATCACCTGCAGTGCCAGCTCAAGTGTAAGTTACATGCACTGGTACCAGCAGAAGTCAGGCACCTCCCCCAAACCATGGATTTATGAAATATCCAAACTGGCTTCTGGAGTCCCAGCTCGCTTCAGTGGCAGTGGGTCTGGGACCTCTTACTCTCTCACAATCAGCAGCATGGAGGCTGAAGATGCTGCCATTTATTACTGCCAGCAGTGGAATTATCCACTTA |
| IGKV5-37*01_S9729_MRL | MRL/MpJ | Musmus IGKV5-37*01 F | 97.13 | 285 | GACATCCTGATGACCCAGTCTCCAGCCACCCTGTCTGTGACTCCAGGAGAAACAGTCAGTCTTTCCTGTAGGGCCAGCCAGAATACTTACAAGAACCTACACTGGTATCAACAGAAATCACATGGGACTCCAAAGCTTCTCATCAAGTATGCATCTGATCCCATCTCTGGGATCCCCTCCAGGTTCACTGGCAGTGGATCAGGGACAGATTACACTCTCAGTATCAACAGTGTGAAGCCCGAAGATGAAGGAATATATTACTGTCTTCAAGGTTACAGCATGCCT |
| IGKV7-33*01_S2856_MRL | MRL/MpJ | Musmus IGKV7-33*01 F | 97.31 | 305 | GACATTGTGATGACTCAGTCTCCAACTTTCCTTGCTGTGACAGCAAGTAAGAAGGTCACCATTAGTTGCACGGCCAGTGAGAGCCTTTATTCAAGCAAACACAAGGTGAACTACTTAGCTTGGTACCAGAAGAAACCAGAGCAATCTCCTAAACTGCTGATTTATGGGGCATCCAACCGATTCACTGGGGTCCCTGATCGCTTCACAGGCAGTGGATCTGGGACAGATTTCACTCTGACCATCAGCAGTGTGCAGGCTGAAGACCTCACACATTATTACTGTGCACAGTTTTACAGCTATCCTCT |
| IGKV1-99*01_S3206_MRL | MRL/MpJ | Musmus IGKV1-99*01 F | 98.3 | 302 | GATGTTGTTCTGACCCAAACTCCACTCTCTCTGCCTGTCAATATTGGAGATCAAGCCTCTATCTCTTGCAAGTCTACTAAGAGCCTTCTGAATAGTGATGGATTCACTTATTTGGGCTGGTACCTGCAGAAGCCAGGCCAGTCTCCACAGCTCCTAATATATTTGGTTTCTAATCGATTTTCTGGAGTTCCAGACAGGTTCAGTGGTAGTGGGTCAGGGACAGATTTCACCCTCAAGATCAGCAGAGTGGAGGCTGAGGATTTGGGAGTTTATTATTGCTTCCAGAGTAACTATCTTCCTCT |
| IGKV6-25*01_S5447_MRL | MRL/MpJ | Musmus IGKV6-25*01 F | 98.21 | 286 | TGCATTGTGATGACCCAGTCTCACAAATTCATGTCCACATCAGTAGGAGACAGGGTCAGCATCACCTGCAAGGCCAGTCAGGATGTGAGTACTGCTGTAGCCTGGTATCAACAAAAACCAGGGCAATCTCCTAAACTACTGATTTACTGGGCATCCACCCGGCACACTGGAGTCCCTGATCGCTTCACAGGCAGTGGATCTGGGACAGATTTCACTCTTACCATCAGCAGTGTGCAGGCTGAAGACCTGGCACTTTATTACTGTCAGCAACATTATAGCACTCCTC |
| IGKV1-88*01_S0716_MRL | MRL/MpJ | Musmus IGKV1-88*01 F | 99.66 | 302 | GATGTTGTGGTGACTCAAACTCCACTCTCCCTGCCTGTCAGCTTTGGAGATCAAGTTTCTATCTCTTGCAGGTCTAGTCAGAGTCTTGCAAACAGTTATGGGAACACCTATTTGTCTTGGTACCTGCACAAGCCTGGCCAGTCTCCACAGCTCCTCATCTATGGGATTTCCAACAGATTTTCCGGGGTGCCAGACAGGTTCAGTGGCAGTGGTTCAGGGACAGATTTCACACTCAAGATCAGCACAATAAAGCCTGAGGACTTGGGAATGTATTACTGCTTACAAGGTACACATCAGCCTCC |

| | | | | | |
|---|---|---|---|---|---|
| IGKV1-132*01_MRL | MRL/MpJ | Musmus IGKV1-132*01 F | 100 | 302 | GATGTTGTGATGACCCAGACTCCACTGTCTTTGTCGGTTACCATTGGACAACCAGCCTCTATCTCTTGCAAGTCAAGTCAGAGCCTCTTATATAGTAATGGAAAGACATATTTGAATTGGTTACAACAGAGGCCTGGCCAGGCTCCAAAGCACCTAATGTATCAGGTGTCCAAACTGGACCCTGGCATCCCTGACAGGTTCAGTGGCAGTGGATCAGAAACAGATTTTACACTTAAAATCAGCAGAGTGGAGGCTGAAGATTTGGGAGTTTATTACTGCTTGCAAGGTACATATTATCCTCA |
| IGKV4-62*01_S8002_MRL | MRL/MpJ | Musmus IGKV4-62*01 ORF | 96.74 | 286 | CAAATTGTTCTCACCCAGTCTCCAGCAATCATGTCTGCATCTCCAGGGGAGAAGGTCACCATGACCTGCAGTGCCAGCTCAAGTGTAAGTTACATGTACTGGTACCAGCAGAAGCCAGGATCCTCCCCCAGACTCTGGATTTATGACACATCCAACCTGGCTTCTGGAGTCCCCGCTCGCTTCAGTGGCAGTAGGTCTGGGACCTCTTATTCTCTCACAATCAGCAGCATGGAGGCTGAAGATGCTGCCACTTATTACTGCCATCAGCGGAGTAGTTACCCATCCA |
| IGKV4-81*01_S9153_MRL | MRL/MpJ | Musmus IGKV4-81*01 F | 96.45 | 292 | GAAAATGTGCTGACCCAGTCTCCAGCAATCATGGCTGCATCTCCAGGGGAGAAGGTCACCATGACCTGCAGTGCCAGCTCAAGTGTAAGTTCTGGTAACTTTCACTGGTACCAGCAGAAGCCAGGCACTTCTCCCAAACTCTGGATTTATAGGACATCCAACCTGGCTTCTGGAGTCCCCGCTCGCTTCAGTGGCAGTGGGTCTGGGACCTCTTACTCTCTTACAATCAGCAGCATGGAGGCCGAAGATGCTGCCACTTATTACTGCCAGCAGTGGAGTGGTTACCCACTCA |
| IGKV8-27*01_MRL | MRL/MpJ | Musmus IGKV8-27*01 F | 100 | 300 | AACATTATGATGACACAGTCGCCATCATCTCTGGCTGTGTCTGCAGGAGAAAAGGTCACTATGAGCTGTAAGTCCAGTCAAAGTGTTTTATACAGTTCAAATCAGAAGAACTACTTGGCCTGGTACCAGCAGAAACCAGGGCAGTCTCCTAAACTGCTGATCTACTGGGCATCCACTAGGGAATCTGGTGTCCCTGATCGCTTCACAGGCAGTGGATCTGGGACAGATTTTACTCTTACCATCAGCAGTGTACAAGCTGAAGACCTGGCAGTTTATTACTGTCATCAATACCTCTCCTCG |
| IGKV8-30*01_S4505_MSM | MSM/MsJ | Musmus IGKV8-30*01 F | 97.31 | 305 | GACATTGTTATGTCACAGTCTCCATCCTCCCTGGCTGTGTCAGTTGGAGAGAAGGTTACTATGAGCTGCAAGTCCAGTCAGAGCCTTTTATATAGTAGCAATCAAAAGAACTACTTGGCTTGGTACCAGCAGAAACCTGGGCAGTCTCCTAAACTGTTGATCTACTGGGCATCCACTAGGGAATCTTGGGTCCCTGATCGCTTCACAGGCAGTGGATCTGGGACAGATTTCACTCTCACCATCAGCAGTGTGAAGGCTGAAGACCTGGCCGTTTATTACTGTCAGCAATATTATAGCGATCCTCC |
| IGKV6-20*01_S2863_MSM | MSM/MsJ | Musmus IGKV6-20*01 F | 99.28 | 287 | AACATTGTAATGACCCAATCTCCCAAATCCATGTCCATGTCAGTAGGAGAGAGGGTCACCTTGAGCTGCAAGGCCAGTGAGAATGTGGGTACTTATGTATCCTGGTATCAACAGAAACCAGAGCAGTCTCCTAAACTGCTGATATACGGGGCATCCAACCGGTACACTGGGGTCCCTGATCGCTTCACAGGCAGTGGATCTGCAACAGATTTCACTCTGACCATCAGCAGTGTGCAGGCTGAAGACTTTGCAGATTATCACTGTGGACAGAGTTACAGCTATCCTCC |
| IGKV6-14*01_S5086_MSM | MSM/MsJ | Musmus IGKV6-14*01 F | 95.7 | 287 | GACATTGTGATGACCCAGTCTCAAAAATTCATGTCCACATCAGTAGGAGAGAGGGTCATCATCACCTGCAAGGCCAGTCAGAATGTGGGTACTGCTGTAGCCTGGTATCAACAGAAACCAGGACAATCTCCTAAAGCACTGATTTACTCGGCATCCTACCGGTACACTGGAGTCCCTGATCGCTTCACAGGCAGTGGATCTAGGACAGATTTCACTCTCACCATTAGCAATGTGCAGTCTGAAGACCTGGCAGATTATTTCTGTCTGCAACATTGGAGTTATCCTCT |

| | | | | | |
|---|---|---|---|---|---|
| IGKV2-137*01_S0906_MSM | MSM/MsJ | Musmus IGKV2-137*01 F | 94.22 | 302 | GATATTGTGATGACTCAGGCTGCACCCTCTGTACCTGTCACTCCTGGAGAGTCAGTATCCATCTCCTGCAGATCTAGTAAGAGTCTCCTGCACAGTAGTGGCAAGCATAGGTTGTATTGGTTCCTACAGAGGCCAGGCCAGTCTCCTCAGCGCCTGATATATTATATGTCCAACCTTGCCTCAGGAGTCCCAGACAGGTTCAGTGGCAGTGGGTCAGGAACTGATTTCACACTGAGAATCAGTAGAGTGGAGGCTGAGGATGTGGGTGTTTATTACTGTATGCAAAGTCTAGAATATCCTTT |
| IGKV10-94*01_Mus_spretus_S4111_MSM | MSM/MsJ | Musspr IGKV10-94*01 F | 98.21 | 287 | GATATCCAGATGACACAGACTACATCCTCCCTGTCTGCCTCTCTGGGAGACAGAGTCACCATCAGTTGCAGGGCAAGTCAGGATATTAGCAATTATTTAAACTGGTATCAGCAGAAACCAGATGGAACTGTTAAACCCCTGATCTATTATGCGTCAAGTTTACACTCAGGAGTCCCATCAAGGTTCAGTGGCAGTGGGTCTGGGACAGATTATTCTCTCACCATCAGCAACCTGGAACCTGAAGATATTGCCACTTACTATTGTCAGCAGTATAGTAAGCTTCCTCC |
| IGKV6-13*01_S5013_MSM | MSM/MsJ | Musmus IGKV6-13*01 F | 99.28 | 285 | GACATTGTGATGCCCCAGTCTCAAAAATTCATGTCCACATCAGTAGGAGACAGGGTCAGCGTCACCTGCAAGGCCAGTCAGAATGTGGGTACTGCTGTAGCCTGGTATCAACAGAAACCAGGACAATCTCCTAAACTACTGATTTACTCGGCATCCAATCGGTACACTGGAGTCCCTGATCGCTTCACAGGCAGTGGATCTGGGACAGATTTCACTCTCACCATCAGCAATATGCAGTCTGAAGACCTGGCAGATTATTTCTGCCAGCAATATAGCAGCTATCCT |
| IGKV15-103*01_S4991_MSM | MSM/MsJ | Musmus IGKV15-103*01 ORF | 95.34 | 287 | GACATCCAAGTGAACCAGTCTCCATCCAGTCTGTCTGCATCCCTCGGAGACACAATTACCATCACTTGCCGTGCCAGTCAGGACATTAGTATTTGGTTAAGCTGGTACCAGCAGAAACCAGGAAATATTCCTAAACTATTGATCTATAAGGCTTCCAACTTGCACACAGGCGTCCCACCAAGGTTTAGTGGCAGTGGATCTGGGACAGATTTCACATTAACCATCAGCAGCCTACAGCCTGAAGACATTGCCACTTACTACTGTCTACAGAGTCAAAGTTATCCTCT |
| IGKV8-19*01_MSM | MSM/MsJ | Musmus IGKV8-19*01 F | 100 | 305 | GACATTGTGATGACACAGTCTCCATCCTCCCTGACTGTGACAGCAGGAGAGAAGGTCACTATGAGCTGCAAGTCCAGTCAGAGTCTGTTAAACAGTGGAAATCAAAAGAACTACTTGACCTGGTACCAGCAGAAACCAGGGCAGCCTCCTAAACTGTTGATCTACTGGGCATCCACTAGGGAATCTGGGGTCCCTGATCGCTTCACAGGCAGTGGATCTGGAACAGATTTCACTCTCACCATCAGCAGTGTGCAGGCTGAAGACCTGGCAGTTTATTACTGTCAGAATGATTATAGTTATCCTCC |
| IGKV6-23*01_S0027_MSM | MSM/MsJ | Musmus IGKV6-23*01 F | 93.91 | 287 | GACATTGTGATGACCCAGTCGCAAAAATTCATGTCCACATCAGTAGGAGAGAGGGTCGGCATCAGCTGCAAGGCCAGTCAGAATGTGGGTACTGCTGTAGCCTGGTATCAACAGAAACCAGGGCAGTCTCCTAAACTACTGATTTACTGGGCATCCACCCGGCACACTGAAGTCCCTGATCGCTTCACAGGCAGTGGATCTGGGACAGATTTCACTCTTACCATCAGCAGTATGCAGTCTGAAGACCTGGCAGATTATTTCTGTCAGCAACATTATAGCACTCCTCT |
| IGKV14-126*01_S6908_MSM | MSM/MsJ | Musmus IGKV14-126*01 F | 99.64 | 286 | GACATCAAGATGACCCAGTCTCCATCCTCCATGTATGCATCGCTGGGAGAGAGAGTCACTATCACTTGCAAGGCGAGTCAGGACATTAAAAGCTATTTAAGCTGGTACCAGCAGAAACCAGGGAAATCTCCTAAGACCCTGATCTATTATGCAACAAGCTTGGCAGATGGGGTCCCATCAAGATTCAGTGGCAGTGGATCTGGGCAAGATTATTCTCTAACCATCAGCAGCCTGGAGTCTGACGATACAGCAACTTATTACTGTCTACAGCATGGTGAGAGCCCTC |
| IGKV8-34*01_S0627_MSM | MSM/MsJ | Musmus IGKV8-34*01 F | 98.32 | 305 | GACATTTTGATGACTCAGTCTCCATCCTCCCTGGCTGTGTCAGCAGGAGAGAAGGTCACTATGAGCTGCAAGTCCAGTCAGAGTCTTTTAGCTAGTGCCAACCAAAATAACTACTTGGCCTGGCACCAGCAGAAACCAGGACGATCTCCTAAAATGCTGATAATTTGGGCATCCACTAGGGTATCTGGAGTCCCTGATCGCTTCATAGGCAGTGGATCTGGAACGGATTTCACTCTGACCATCAACAGTGTGCAGGCTGAAGATCTGGCTGTTTATTACTGTCAGCAGTCCTATAGCGCTCCTAC |

| | | | | | |
|---|---|---|---|---|---|
| IGKV10-94*01_S9482_MSM | MSM/MsJ | Musmus IGKV10-94*01 F | 98.21 | 287 | GATATCCAGATGACACAGAGTACATCCACCCTGTCTGCCTCTCTGGGAGACAGAGTCACCATCAGTTGCAGTGCAAGTCAGGGCATTAGCAATTATTTAAACTGGTATCAGCAGAAACCAGATGGAACTGTTAAACTCCTGATCTATTATACATCAAGTTTACACTCAGGAGTCCCATCAAGGTTCAGTGGCAGTGGGTCTGGAACAGATTATTCTCTCACCATCAGCAACCTGGAACCTGAAGATATTGCCACTTACTATTGTCAGCAGTATAGTAATTTTCCTCC |
| IGKV12-44*01_S9406_MSM | MSM/MsJ | Musmus IGKV12-44*01 F | 96.77 | 287 | GACATCCAGATGACTCAGTCTCCAGCTTCCCTATCTGCATCTGTGGGAGAAACTGTCACCATCACATGTCGAGCAAGTGGGAATATTTACAGTAATTTAGCATGGTATCAGCAGAAACAGGGAAAATCTCCTCAGCTCCTGGTCTATAATGCAAAAACCTTAGCAGAAGGTGTGCCATCAAGGTTCAGTGGCAGTGGATCAGGAACACAATATTCTCTCAAGATCAACAGCCTACAGCCTGAAGATTTTGGGAGTTATTACTGTCAACATTATTATGGTACTCCTCC |
| IGKV1-122*01_S2905_MSM | MSM/MsJ | Musmus IGKV1-122*01 F | 97.28 | 302 | GATGCTGTGATGACCCAAACTCCACTCTCCCTGCCTGTCAGTCTTGGAGATCAAGCCTCCATCTCTTGCAGATCTAGTCAGAGCCTTGTACACAGTAATGGAAACACCTATTTGAACTGGTACCTCCAGAAACCAGGCCAGTCTCCACAGCTCTTGATCTACAGGGTTTCCAACCGATTTTCTGGGGTCCTAGACAGGTTCAGTGGCAGTGGTTCAGGGACAGATTTCACACTCAAGATCAGCAGAGTGGAGGCTGAGGATTTGGGAGTTTATTTCTGCCTCCAAGTTACACATGTCCCTCC |
| IGKV12-40*01_S1894_MSM | MSM/MsJ | Musmus IGKV12-40*01 P | 98.57 | 287 | GACATCCAGATGACTCAGTCTCCAGCCTCCCTATCTGCATCTGTGGGAGAAACTGTCACCATCACATGTCAAGCAAGTGAGAATATTGCCAGTGATTTAGCATGGTATCAGCAGAAACAGGGAAAATCTCCTCAGCTCCTGGTCTATGATGCGAGAAACTTAGCAGATGGTGTGCCATCAAGGTTCAGTGGCAGTGGATCAGGCACACAGTATTCTCTCAATATCAACAGCCTGCAGTCTGAAGATGTTGCGAGATATTACTGTCAACATTATTATGGTACTCCTCC |
| IGKV2-137*01_S4057_MSM | MSM/MsJ | Musmus IGKV2-137*01 F | 94.22 | 302 | GATATTGTGATGACTCAGGCTGCACCCTCTGTACCTGTCACTCCTGGAGAGTCAGTATCCATCTCCTGCAGATCTAGTAAGAGTCTCCTGCACAGTAGTGGCAAGCATAGGTTGTATTGGTTCCTACAGAGGCCAGGCCAGTCTCCTCAGCGCCTGATATATTATATGTCCAACCTTGCCTCAGGAGTCCCAGACAGGTTCAGTGGCAGTGGGTCAGGAACTGATTTCACACTGAGAATCAGTAGAGTGGAGGCTGAGGATGTGGGTGTTTATTACTGTATGCAAAGTCTAGAATATCCATT |
| IGKV4-91*01_S4567_MSM | MSM/MsJ | Musmus IGKV4-91*01 F | 96.81 | 292 | GAAATTGTGCTCACCCAGTCTCCAACCACCATGGCTGCATCTCCCGGGGAGAAGGTCACCATCACCTGCAGTGCCAGCTCAAGTATAAGTTCCAATTACTTGCACTGGTATCAGCAGAAGCCAGGATTCCCTCCTAAACTCTTGATATATAGGACATCCAATCTGGCTTCTGGAGTCCCAGCTCGCTTCAGTGGCAGTGGGTCTGGGACCTCTTACTCTCTCACAATTGGCACCATGGAGGCTGAAGGTGCTGCCACTTATTACTGCCAGCAGGGTAGTAGTTTACCACACA |
| IGKV1-117*02_S1601_MSM | MSM/MsJ | Musmus IGKV1-117*01 F, or Musmus IGKV1-117*02 F | 99.32 | 302 | GATGTTGTGATGACCCAAACTCCACTCTCCCTGCCTGTCAGTCTTGGAGATCAAGCCTCCATCTCTTGCAGATCTAGTCAGAGCATTGTACATAGTAATGGAAACACCTATTTAGAATGGTACCTGCAGAAACCAGGCCAGTCTCCAAAGCTCCTGATCTACAAAGTTTCCAACCGATTTTCTGGGGTCCCAGACAGGTTCAGTGGCAGTGGATCAGGGACAGACTTCACACTCAAGATCAGCAGAGTGGAGGCTGAGGATCTGGGAGTTTATTACTGCTTTCAAGGTTCACATGTTCCTCC |
| IGKV4-86*01_S6393_MSM | MSM/MsJ | Musmus IGKV4-86*01 F | 94.93 | 281 | GAAATTGTGCTCACTCAGTCTCCAGCCATCATAGCTGCATCTCTGGGGCAAAAGGTCACCATGACCTGCAGTGCCAGCTCAAGTGTAAGTTACATGCACTGGTACCAGCAGAAGTCAGGCACCTCCCCCAAACTCTGGATTTACGGCACATCCAACCTGGCTTCTGGAGTCCCAGCTCGCTTCAGTGGCAGTGGGTCTGGGACCTCTTACTCTCTCACAATCAGCAGCGTGGAGGCTGAAGATGATGCAACTTATTACTGCCAGCAGTGGAATTACCCACT |

| | | | | | |
|---|---|---|---|---|---|
| IGKV12-46*01_S7378_MSM | MSM/MsJ | Musmus IGKV12-46*01 F | 99.64 | 287 | GACATCCAGATGACTCAGTCTCCAGCCTCCCTATCTGTATCTGTGGGAGAAACTGTCACCATCACATGTCGAGCAAGTGAGAATATTTACAGTAATTTAGCATGGTATCAGCAGAAACAGGGAAAATCTCCTCAGCTCCTGGTCTATGCTGCAACAAACTTAGCAGACGGTGTGCCATCAAGGTTCAGTGGCAGTGGATCAGGCACACAGTATTCCCTCAAGATCAACAGCCTGCAGTCTGAAGATTTTGGGAGTTATTACTGTCAACATTTTTGGGGTACTCCTCC |
| IGKV19-93*01_S0826_MSM | MSM/MsJ | Musmus IGKV19-93*01 F | 96.77 | 287 | GACATCCAGATGACACAGTCTCCATCCTCACTGTCTGCATCTCTGGGAGGCAAAGTCACCATCACTTGCAAGGCAAGCCAAGACATTAACAAGAATATAGCTTGGTACCAACACAAGCCTGGAAAAGGTCCTAGGCTGCTCATACGTTACACATCAATACTAGAGTCAGGCATCCCATCAAGGTTCAGTGGAAGTGGGTCTGGGAGAGATTATTCCTTCAGCATCAGCAACCTGGAGCCTGAAGATATTGCAACTTATTACTGTCTACAGTATTATAATCTTCCACC |
| IGKV2-137*01_S7910_MSM | MSM/MsJ | Musmus IGKV2-137*01 F | 96.6 | 300 | GATATTGTGATGACTCAGGCTGCACCCTCTGTACCTGTCACTCCTGGAGAGTCAGTATCCATCTCCTGCAGGTCTAGTAAGAGTCTCCTGCATAGTAATGGCAACCATTTGTTGTATTGGTTCCTGCAGAGGCCAGGCCAGTCTCCTCAGCTCCTGATATATTATATGTCCAACCTTGCCTCAGGAGTCCCAGACAGGTTCAGTGGCAGTGGGTCAGGAACTGATTTCACACTGAGAATCAGTAGAGTGGAGGCTGAGGATGTGGGTGTTTATTACTGTATGCAAAGTCTAGAATATCCT |
| IGKV17-127*01_S9169_MSM | MSM/MsJ | Musmus IGKV17-127*01 F | 95.7 | 287 | GAAACAACTGTGACCCAGTCTCCAGCATCCCTGTCCATGGCTACAGGAGAAAAAGTCACTATCAGTTGCATCACCAGCACTGATATTGATGATGATATGAACTGGTACCAGCAGAAGCCCGGAGAAGCTCCTAAGCTCCTTATTTCAGAAGGCAATACTCTTCGTCCTGGAGTCCCATCCCGGTTCTCCAGTAGTGGCTATGGCACAGATTTTGTTTTTACAATTGAAAACACTCTCTCAGAAGATGTTGCAGATTACTTCTGTCAGCAAAGTGATAACCTGCCGCT |
| IGKV4-80*01_S4247_MSM | MSM/MsJ | Musmus IGKV4-80*01 F | 99.64 | 282 | CAAATTGTTCTCACCCAGTCTCCAGCAATCATGTCTGCATCTCTAGGGGAGGAGATCACCCTAACCTGCAGTGCCAGCTCGAGTGTAAGTTACATGCACTGGTTCCAGCAGAAGTCAGGCACTTCTCCCAAACTCTTGATTTATAGCACATCCAACCTGGCTTCTGGAGTCCCTTCTCGCTTCAGTGGCAGTGGGTCTGGGACCTTTTATTCTCTCACAATCAGCAGTGTGGAGGCTGAAGATGCTGCCGATTATTACTGCCATCAGTGGAGTAGTTATCCA |
| IGKV4-55*01_S7109_MSM | MSM/MsJ | Musmus IGKV4-55*01 F | 97.46 | 286 | CAAATTGTTCTCACCCAGTCTCCAGCAATCATGTCTGCATCTCCAGGGGAGAAGGTCACCATGACCTGCAGTGCCAGCTCAAGTGTAAGTTACATGTACTGGTACCAGCAGAAGCCAGGATCCTCCCCCAGACTCTGGATTTATGACACATCCAACCTGGCTTCTGGAGTCCCCGCTCGCTTCAGTGGCAGTGGGTCTGGGACCTCTTATTCTCTCACAATCAGCAGCATGGAGGCTGAAGATGCTGCCACTTATTACTGCCAGCAGTGGAGTAGTAACCAACCCA |
| IGKV9-120*01_S5725_MSM | MSM/MsJ | Musmus IGKV9-120*02 F | 96.42 | 285 | GACATCCAGATGACCCAGTCTCCATCCTCCTTATCTGCCTCTCTGGGAGAAAGAATCAGTCTCACTTGCCGGGCAAGTCAGGACATTGGTGGTAGCTTAAACTGGTTCCAGCAGAAACCAGATGGAACTATTAAACGCCTGATCTACGGCACATCCAGTTTAGATTCTGGTGTCCCCAAAAGGTTCAGTGGCAGTAGGTCTGGGTCAGATTATTCTCTCACCATCAGCAGCCTTGAGTCTGAAGACTTTGCAGACTATTACTGTTTACAATATGCTAGTTATCCT |
| IGKV4-74*01_S8852_MSM | MSM/MsJ | Musmus IGKV4-74*01 F | 94.33 | 292 | CAAATTGTTCTCACCCAGTCTCCAGCAATCATGTCTGCATCTCCAGGGGAGAAGGTCACCATGACCTGCAGGGCCAGCTCAAGTGTACGTTCCAGTTACTTGCACTGGTACCAGCAGAAGCCAGGATCTTCCCCCAAACTCTGGATTTATAGAACATCCAACCTGGCTTCAGGAGTCCCAGCTCGCTTCAGTGGCAGTGGGTCTGGGACCTCTTACTCTCTCACAATCAGCAGTGTGGAGGCTGAGGATGCTGCCACTTATTACTGCCAGCAGTATGATAGTTCCCCATCCA |

| | | | | | |
|---|---|---|---|---|---|
| IGKV1-110*01_S5250_MSM | MSM/MsJ | Musmus IGKV1-110*01 F | 97.96 | 302 | GATGTTTTGATGACCCAAACTCCACTCTCCCTGCCTGTCAGTCTTGGAGATCAAGCCTCCATCTCTTGCAGATCTAGTCAGAGCCTTGTACACAGTAATGGAAACACCTATTTATATTGGTACCTGCAGAAGCCAGGCCAGTCTCCAAAGCTCCTGATCTACAAAGTTTCCAACCGATTTTCTGGGGTCCCAGACAGGTTCAGTGGCAGTGGATCAGGGACAGATTTCACACTCAAGATCAGCAGAGTGGAGGCTGAGGATATGGGAGTTTATTACTGCTTTCAAGGTACACATGTTCCTCC |
| IGKV4-62*01_S0389_MSM | MSM/MsJ | Musmus IGKV4-55*01 F | 97.46 | 286 | CAAATTGTTCTCACCCAGTCTCCAGCAATCATGTCTGCATCTCCAGGGGAGAAGGTCACCATGACCTGCAGTGCCAGCTCAAGTGTAAGTTACATGTACTGGTACCAGCAGAAGCCAGGATCCTCCCCCAGACTCTGGATTTATGACACATCCAACCTGGCTTCTGGAGTCCCCGCTCGCTTCAGTGGCAGTGGGTCTGGGACCTCTTATTCTCTCACAATCAGCAGCATGGAGGCTGAAGATGCTGCCACTTATTACTGCCAGCAGTGGAGTAGTAACCAATTCA |
| IGKV12-44*01_MSM | MSM/MsJ | Musmus IGKV12-44*01 F | 100 | 287 | GACATCCAGATGACTCAGTCTCCAGCCTCCCTATCTGCATCTGTGGGAGAAACTGTCACCATCACATGTCGAGCAAGTGAGAATATTTACAGTTATTTAGCATGGTATCAGCAGAAACAGGGAAAATCTCCTCAGCTCCTGGTCTATAATGCAAAAACCTTAGCAGAAGGTGTGCCATCAAGGTTCAGTGGCAGTGGATCAGGCACACAGTTTTCTCTGAAGATCAACAGCCTGCAGCCTGAAGATTTTGGGAGTTATTACTGTCAACATCATTATGGTACTCCTCC |
| IGKV10-96*04_MSM | MSM/MsJ | Musmus IGKV10-96*04 F | 100 | 287 | GATATCCAGATGACACAGACTACATCCTCCCTGTCTGCCTCTCTGGGAGACAGAGTCACCATCAGTTGCAGGGCAAGTCAGGACATTAGCAATTATTTAAACTGGTATCAGCAGAAACCAGATGGAACTGTTAAACTACTGATCTACTACACATCAAGATTACACTCAGGAGTCCCATCAAGGTTCAGTGGCAGTGGGTCTGGGACAGATTATTCTCTCACTATTAGCAACCTGGAACAAGAAGATATTGCCACTTACTTTTGCCAACAGGATAGTAAGCATCCTCC |
| IGKV5-48*01_MSM | MSM/MsJ | Musmus IGKV5-48*01 F | 100 | 287 | GACATCTTGCTGACTCAGTCTCCAGCCATCCTGTCTGTGAGTCCAGGAGAAAGAGTCAGTTTCTCCTGCAGGGCCAGTCAGAGCATTGGCACAAGCATACACTGGTATCAGCAAAGAACAAATGGTTCTCCAAGGCTTCTCATAAAGTATGCTTCTGAGTCTATCTCTGGGATCCCTTCCAGGTTTAGTGGCAGTGGATCAGGGACAGATTTTACTCTTAGCATCAACAGTGTGGAGTCTGAAGATATTGCAGATTATTACTGTCAACAAAGTAATAGCTGGCCAAC |
| IGKV12-89*01_S8177_MSM | MSM/MsJ | Musmus IGKV12-89*01 F | 98.92 | 287 | GACATCCAGATGACTCAGTCTCCAGCTTCACTGTCTGCATCTGTGGGAGAAACTGTCACCATCACATGTGGAGCAAGTGAGAATATTTACGGTGCTTTAAATTGGTATCAGCAGAAACAGGGAAAATCTCCTCAGCTCCTGATCTATGGTGCAACCAACTTGGCAGATGGAATGTCATCGAGGTTCAGTGGTAGTGGATCTGGTAGACAGTATTCTCTCAAGATCAGTAGCCTGCATCCTGACGATGTTGCAACGTATTACTGTCAAAATGTGTTAAGTACCCCTCC |
| IGKV1-133*01_S2477_MSM | MSM/MsJ | Musmus IGKV1-133*01 F | 95.58 | 302 | GATGTTGTGATGACTCAGACCCCACTCACTTTGTCGGTTACCATTGGACAACCAGCCTCCATCTCTTGCAAGTCAAGTCAGAGCCTCTTATATAGTGATGGAAAAACATATTTGGATTGGTTATTACAGAGGCCAGGTCAGTCTCCAAAGAGCCTAATCTATCTGGTGTCTAAACTGGAATCTGGAGTCCCTGACAGGTTCAGTGGCAGTGGATCAGGGACAGATTTCATACTGAAAATCAGCAGAGTGGAGGCTGAGGATTTGGGAGTTTATTACTGCGTGCAAGGTACACATTTTCCTCA |
| IGKV3-2*01_S2813_MSM | MSM/MsJ | Musmus IGKV3-2*01 F | 99.66 | 299 | GACATTGTGCTGACCCAATCTCCAGCTTCTTTGGCTGTGTCTCTAGGGCAGAGGGCCACCATCTCCTGCAGAGCCAGCGAAAGTGTTGATAATTATGGCATTAGTTTTATGAACTGGTTCCAACAGAAACCAGGACAGCCACCCAAACTCCTCATCTATGCTGCATCCAACCAAGGATCTGGGGTCCCTGCCAGGTTTAGTGGCAGTGGGTCTGGGACAGACTTCAGCCTCAACATCCATCCTATGGAGGAGGATGATACTGCAATGTATTTCTGTCAGCAAAGTAAGGAGGTTCCTCC |

| | | | | | |
|---|---|---|---|---|---|
| IGKV4-59*01_S6619_MSM | MSM/MsJ | Musmus IGKV4-59*01 F | 97.46 | 286 | CAAATTGTTCTCACCCAGTCTCCAGCAATCATGTCTGCATCTCCAGGGGAGAAGGTCACCATGACATGCAGTGCCAGCTCAAATGTAAATTATATGCACTGGTACCAGCAGAAGTCAGGCACCTCCCCCAAAAGATGGATTTATGACACATCCAAACTGGATTCTGGAGTCCCTGCTTGCTTCAGTGGCAGTGGGTCTGGGACCTCTTATTCTCTCACAATCAGCAGCATGGAGGCTGAAGATGCTGCCACTTATTACTGCCAGCAGTGGAGTAGTAACCCACCCA |
| IGKV1-135*01_S8190_MSM | MSM/MsJ | Musmus IGKV1-135*01 F | 96.26 | 302 | GATGTTGTGATGACTCAGACCCCACTCACTTTGTCGGTTACCATTGGACAACCAGCCTCCATCTCTTGCAAGTCAAGTCAGAGCCTCTTACATAGTAATGGAAAGACATATTTGAATTGGTTATTACAGAGGCCAGGCCAGTCTCCAAAGCTCCTAATCTATCTGGTGTCTAAACTGGAATCTGGAGTCCCTGACAGGTTCAGTGGCAGTGGATCAGGGACAGATTTCACACTGAAAATCAGCAGAGTGGAGGCTGAGGATTTGGGAGTTTATTACTGCTTGCAAGCTACACATTTTCCTCA |
| IGKV16-104*01_S7508_MSM | MSM/MsJ | Musmus IGKV16-104*01 F | 98.92 | 288 | GATGTCCAGATAACCCAGTCTCCATCTTATCTTGCTGCATCTCCTGGAGAAACCATTACTATTAATTGCAGGGCAAGTAAGAGCATTAGGACATATTTAGCCTGGTATCAAGAGAAACCTGGGAAAACTAATAAGCTTCTTATCTACTCTGGATCCACTTTGCAATCTGGAATTCCATCAAGGTTCAGTGGCAGTGGATCTGGTACAGATTTTACTCTCACCATCAGTAGCCTGGAGCCTGAAGATTTTGCAATGTATTACTGTCAACAGCATAATGAATACCCGTAC |
| IGKV4-68*01_S8363_MSM | MSM/MsJ | Musmus IGKV4-68*01 F | 96.74 | 286 | CAAATTGTTCTCACCCAGTCTCCAGCAATCATGTCTGCATCTCCAGGGGAGAAGGTCACCATGACCTGCAGTGCCAGCTCAAGTGTAAGTTATATGCACTGGTACCAGCAGAAGCCAGGATCCTCACCCAAACCCTGGATTTATCGCACATCCAACCTGGCTTCTGGAGTCCCTGCTCGCTTCAGTGGCAGTGGGTCTGGGACCTCTTACTCTCTCACAATTACCAGCATTGAGGCTGAAGATGCTGCCACTTATTACTGCCAGCAGTGGAGTAGTAACCCACCCA |
| IGKV4-81*01_S7470_MSM | MSM/MsJ | Musmus IGKV4-81*01 F | 96.81 | 292 | GAAAATGTGCTGACCCAGTCTCCAGCAATCATGGCTGCATCTCCAGGGGAGAAGGTCACCATGACCTGCAGTGCCAGCTCAAGTGTAAGTTCTGGTAACTTTCACTGGTACCAGCAGAAGCCAGGCACTTCTCCCAAACTCTGGATTTATAGGACATCCAACCTGGCTTCTGGAGTCCCAGCTCGCTTCAGTGGCAGTGGGTCTGGGACCTCTTACTCTCTTACAATCAGCAGCATGGAGGCCGAAGATGCTGCCACTTATTACTGCCAGCAGTGGAGTGGTTACCCACCCA |
| IGKV1-133*01_S9974_MSM | MSM/MsJ | Musmus IGKV1-133*01 F | 96.26 | 302 | GATGTTGTGATGACCCAGACTCCACTCACTTTGTCTGTTACCATTGGACAGCCAGCTTCCATCTCTTGCAAGTCAAGTCAGAGCCTCTTATATAGTGATGGAAAAACCTATTTGAATTGGTTATTACAGAGTCCAGGCCAGTCTCCAAAGCTCCTAATCTATCTGGTGTCTAAACTGGAATCTGGAGTCCCTGACAGATTCAGTGGCAGTGGATCAGGGACAGATTTTACACTGAAAATCAGCAGAGTGGAGGCTGAGGATTTGGGAGTTTATTACTGCGTGCAAGGTACACATTTCCCTCA |
| IGKV20-101-2*01_S8388_MSM | MSM/MsJ | Musmus IGKV20-101-2*01 F | 97.46 | 282 | AATATCCAGGTGATCCAGTCACCATTTCTGTCTGCATCTGTGGGAGAGAGGGTCACAATCAGCTGCAAGACACATCAGCATATTAACAGTTCCATAGCCTGGTACCAGCAAAAATTTGGAAAAGCTCCCAAACTCCTGATAAGAGATGGAAGTTTTTCTCTAACAGACACCCCATCAAGGTTCAGTGGGAATGGATTTGGCACAGATTTCACACTCAGCATCAGCAGTATGCAGCCTGAAGATGGTGCCACATATTTCTGCCAGCAGCATTTTAACTATTAC |
| IGKV9-124*01_S3726_MSM | MSM/MsJ | Musmus IGKV9-124*01 F | 98.57 | 285 | GACATCCAGATGACCCAGTCTCCATCCTCCTTATCTGTCTCTCTGGGAGAAAGAGTCAGTCTCACTTGTCGGGCAAGTCAGGAAATTAGTGGTTACTTAAGCTGGCTTCAGCAGAAACCAGATGGAACTATTAAACGCCTGATCTACAGCACATCCACTTTAGATTCTGGTGTCCCAAAAAGGTTCAGTGGCAGTAGGTCTGGGTCAGATTATTCTCTCACCATCAGCAGCCTTGAGTCTGAAGATTTTGCAGACTATTACTGTCTACAATATGCTAGTTCTCCT |

| | | | | | |
|---|---|---|---|---|---|
| IGKV4-62*01_S4904_MSM | MSM/MsJ | Musmus IGKV4-62*01 ORF, or Musmus IGKV4-68*01 F | 96.74 | 286 | CAAATTGTTCTCACCCAGTCTCCAGCAATCATGTCTGCATCTCCAGGGGAGAAGGTCACCATGACCTGCAGTGCCAGCTCAAGTGTAAGTTACATGTACTGGTACCAGCAGAAGCCAGGATCTTCCCCCAAACCCTGGATTTATGACACATCCAACCTGGCTTCTGGAGTCCCTGCTCGCTTCAGTGGCAGTGGGTCTGGGACCTCTTACTCTCTCACAATCAGCCGCATGGAGGCTGAAGATGCTGCCACTTATTACTGCCAGCAGTACAGTGGTTACCCATTCA |
| IGKV7-33*01_S0872_MSM | MSM/MsJ | Musmus IGKV7-33*01 F | 97.64 | 305 | GACATTGTGATGACTCAGTCTCCAACTTTCCTTGCTGTGACAGCAAGTAAGAAGGTCACCATTAGTTGCACGGCCAGTGAGAGCCTTTATTCAAGCAAACACAAGGTGAACTACTTGGCTTGGTACCAGAAGAAACCAGAGCAATCTCCTAAACTGCTGATTTATGGGGCATCCAACCGATTCACTGGGGTCCCTGATCGCTTCACAGGCAGTGGATCTGGGACAGATTTCACTCTGACCATCAGCAGTGTGCAGGCTGAAGACCTCACACATTATTACTGTGCACAGTTTTACAGCTATCCTCT |
| IGKV4-73*01_S7826_MSM | MSM/MsJ | Musmus IGKV4-73*01 ORF | 97.87 | 292 | CAAATTGTTCTCACCCAGTCTCCAGCAATCATGTCTGCATCTCCTGGGGAACGGGTCACCATGACCTGCAGTGCCAGCTCAAGTGTAAGTTCCAGCTACTTGCACTGGTACCAGCAGAAGCTAGGTTCCTCCCCAAAACTCTGGATTTATAGCACATCCAACCTGGCTTCTGGAGTCCCAGCTCGCTTCAGTGGCAGTGGGTCTGGGACCTCTTACTCTCTCACAATCAGCAGCATGGAGGCTGAAGATGCTGCCACTTATTACTGCCAGCAGTGGAGTAGTAACCAACTCA |
| IGKV4-57-1*01_S5694_MSM | MSM/MsJ | Musmus IGKV4-57*01 F | 95.65 | 292 | GAAAATGTGCTCACCCAGTCTCCAGCACTCATGGCTGCATCTCCAGGGGAGAAGGTCACCATGACCTGCAGTGCCAGCTCAAGTGTAAGTTCCAGTTACTTGCACTGGTACCAGCAGAAGTCAGGTGCCTCCCCCAAACTCTGGATTTACGGCACATCCAACCTAGCTTCTGGAGTCCCAGCTCGCTTCAGTGGCAGTGGGTCTGGGACCTCTTACTCTCTCACAATCAGCAGCGTGGAGGCTGAAGATGATGCAACTTATTACTGCCAGCAGGCGTGGAATTACCCATTCA |
| IGKV4-61*01_S1531_MSM | MSM/MsJ | Musmus IGKV4-61*01 F | 99.64 | 286 | CAAATTGTTCTCACCCAGTCTCCAGCAATCCTGTCTGCATCTCCAGGGGAGAAGGTCACCATATCCTGCAGTGCCAGCTCAAGTGTAAGTTACATGTACTGGTACCAGCAGAAGCCAGGATCCTCCCCCAAACCCTGGATTTATCGCACATCCAACCTGGCTTCTGGAGTCCCTGCTCGCTTCAGTGGCAGTGGGTCTGGGACCTCTTACTCTCTCACAATCAGCAGCATGGAGGCTGAAGATGCTGCCACTTATTACTGCCAGCAGTATCATAGTTACCCACCCA |
| IGKV1-99*01_S6030_MSM | MSM/MsJ | Musmus IGKV1-99*01 F | 97.96 | 302 | GATGTTGTTCTGACCCAAACTCCACTCTCTCTGCCTGTCAATATTGGAGATCAAGCCTCTATCTCTTGCAAGTCTACTAAGACCCTTCTGAATAGTGATGGATTCACTTATTTGGGCTGGTACCTGCAGAAGCCAGGCCAGTCTCCACAGCTCCTAATATATTTGGTTTCTAATCGATTTTCTGGAGTTCCAGACAGGTTCAGTGGTAGTGGGTCAGGGACAGATTTCACCCTCAAGATCAGCAGAGTGGAGGCTGAGGATTTGGGAGTTTATTATTGCTTCCAGAGTAACTATCTTCCTCT |
| IGKV4-50*01_S4730_MSM | MSM/MsJ | Musmus IGKV4-50*01 F | 97.46 | 286 | GAAAATGTGCTCACCCAGTCTCCAGCACTCATGTCTGCATCTCCAGGGGAGAAGGTCACCATGAGCTGCAGGGCCAGCTCAAGTGTAAATTACATGTACTGGTACCAGCAGAAGTCAGGTGCCTCCCCCAAACTATGGATTTATTACACATCCAACCTGGCTCCTGGAATCCCAGCTCGCTTCAGTGGCAGTGGGTCTGGGACCTCTTATTCTCTCACAATCAGCAGCGTGGAGGCTGAAGATGCTGCCACTTATTACTGCCAGCAGTTTACTAGTTCCCCATCCA |

| | | | | | |
|---|---|---|---|---|---|
| IGKV3-10*01_S9898_MSM | MSM/MsJ | Musmus IGKV3-10*01 F, or Musmus IGKV3-5*01 F | 98.97 | 299 | GACATTGTGCTGACCCAATCTCCAGCTTCTTTGGCTGTGTCTCTAGGGCAGAGGGCCACCATATCCTGCAGAGCCAGTGAAAGTGTTGATAGTTATGGCAATAGTTTTATGCACTGGTACCAGCAGAAACCAGGACAGCCACCCAAACTCCTCATCTATCTTGCATCCAACCTAGAATCTGGGGTCCCTGCCAGGTTCAGTGGCAGTGGGTCTAGGACAGACTTCACCCTCACCATTAATCCTGTGGAGGCTGATGATGTTGCAACCTATTACTGTCAGCAAAATAATGAGGATCCTCC |
| IGKV3-12*01_S4919_MSM | MSM/MsJ | Musmus IGKV3-12*01 F | 98.28 | 299 | GACATTGTGCTGACACAGTCTCCTGCTTCCTTAGCTGTATCTCTGGGGCAGAGGGCCACCATCTCATGCAGGGCCAGCCAAAGTGTCAGTACATCTAGCTATAGTTACATGCACTGGTACCAACAGAAACCAGGACAGCCACCCAAACTCCTCATCTATTTTGCATCCAACCTAGAATCTGGGGTCCCTGCCAGGTTCAGTGGCAGTGGGTCTGGGACAGACTTCACCCTCAACATCCATCCTGTGGAGGAGGAGGATGCTGCAACATATTACTGTCAGCACAGTAGGGAGTTTCCTCC |
| IGKV4-81*01_S9628_MSM | MSM/MsJ | Musmus IGKV4-81*01 F | 96.45 | 292 | GAAAATGTGCTGACCCAGTCTCCAGCAATCATGGCTGCATCTCCAGGGGAGAAGGTCACCATGACCTGCAGTGCCAGCTCAAGTGTAAGTTCTGATAACTTTCACTGGTACCAGCAGAAGTCAGGCACTTCTCCCAAACTCTGGATTTATAGGACATCCAAATTGGCTTCTGGAGTCCCAGCTCGCTTCAGTGGCAGTGGGTCTGGGACCTCTTACTCTCTTACAATCAGCAGCGTGGAGGCCGAAGATGCTGCCACTTATTACTGCCAGCAGTGGAGTGGTTACCCGCTCA |
| IGKV4-59*01_S5176_MSM | MSM/MsJ | Musmus IGKV4-59*01 F | 97.1 | 286 | CAAATTGTTCTCACCCAGTCTCCAGCAATCATGTCTGCATCTCCAGGGGAGAAGGTCACCATGACCTGCAGTGCCAGCTCAAGTGTAAGTTACATGTACTGGTACCAGCAGAAGCCAGGATCCTCCCCCAAAAAATGGATTTATGACACATCCAACCTGGCTTCTGGAGTCCCCGCTCGCTTCAGTGGCAGTGGGTCTGGGACCTCTTATTCTCTCACAATCAGCAGCATGGAGGCTGAAGATGCTGCCACTTATTACTGCCAGCAGTGGAGTAGTTACCCATCCA |
| IGKV12-98*01_S2656_MSM | MSM/MsJ | Musmus IGKV12-98*01 F | 98.92 | 287 | GACATTCAGATGACCCAGTCTCCTGCCTCCCAGTCTGCATCTCTGGGAGAAAGTGTCACCATCACATGCCTGGCAAGTCAGACCATTGGTACATGGTTAGCATGGTATCAGCAGAAACCAGGGAAATCTCCTCAGCTCCTGATTTATGCTGCAACCAGCTTGGCAGATGGGGTCCCATCAAGGTTCAGTGGTAGTGGATCTGGCACAAAGTTTTCTTTCAAGATCAGCAGCCTACAGGCTGAAGATTTTGCAAGTTATTACTGTCAACAACATTACAGTACTCCTCT |
| IGKV6-25*01_S3805_MSM | MSM/MsJ | Musmus IGKV6-23*01 F | 93.91 | 287 | GACATTGTGATGACCCAGTCGCAAAAATTCATGTCCACATCAGTAGGAGAGAGGGTCGGCATCAGCTGCAAGGCCAGTCAGAATGTGGGTACTGCTGTAGCCTGGTATCAACAGAAACCAGGGCAGTCTCCTAAACTACTGATTTACTGGGCATCCACCCGGCACACTGAAGTCCCTGATCGCTTCACAGGCAGTGGATCTGGGACAGATTTCACTCTTACCATCAGCAGTATGCAGTCTGAAGACCTGGCAGATTATTTCTGTCAGCAACATTATAGCACTCCTCC |
| IGKV4-50*01_S4988_MSM | MSM/MsJ | Musmus IGKV4-50*01 F | 96.74 | 286 | GAAAATGTGCTCACCCAGTCTCCAGCAATCATGTCTGCAACTCTAGGGGAGAAGGTCACCATGAGCTGCAGGGCCAGCTCAAATGTAAAGAACATGTACTGGTACCAGCAGAAGTCAGGTGCCTCCCCCAAACTATGGATTTATTACACATCCAACCTGGCTTCTGGAGTCCCAGCTCGCTTCAGTGGCAGTGGGTCTGGGACCTCTTATTCTCTCACAATCAGCAGCGTGGAGGCTGAAGATGCTGCCACTTATTACTGCCAGCAGTTTACTAGTTCCCCATACA |
| IGKV4-51*01_S7391_MSM | MSM/MsJ | Musmus IGKV4-51*01 F, or Musmus IGKV4-57-1*01 F | 92.55 | 292 | GAAAATGTGCTCACCCAGTCTCCAGCACTCATGGCTGCATCTCCAGGGGAGAAGGTCACCATGACCTGCAGTGCCAGCTCAAGTGTAAGTTCTAGCTACTTGCACTGGTACCAGCAGAAGTCAGGTGCCTCCCCCAAACTCTGGATTTACCGCACATCCAACCTGGCTTCCGGAGTCCCAGCTCGCTTCAGTGGCAGTGGGTCTGGGACCTCTTACTCTCTCACAATCAGCAACGTGGAGGCTGAAGATGATGCAACTTATTACTGCCAGCAGGGGTGGGATTACCCATTCA |

| | | | | | |
|---|---|---|---|---|---|
| IGKV13-85*01_S1899_MSM | MSM/MsJ | Musmus IGKV13-85*01 F | 97.49 | 287 | GACATCCAGATGACACAATCTTCATCCTCCTTGTCTGTATCTCTACGAGGCAGAGTCACCATTACGTGCAAGGCAAGTGACCACATTAATAATTGGTTAGCCTGGTATCAGCAGAAACCAGGAAATGCTTCTAGGCTCTTAATATCTAGTGCGACCAGTTTGGAAAATGGGGTTCCTTCAAGATTCAGTGGCAGTGGATCTGGAAAGGATTACACTCTCAGCATTACCAGTCTTCAGACTGAAGATGTTGCTACTTATTACTGTCAACAGTATTGGAGTACTCCTCC |
| IGKV4-60*01_S2188_MSM | MSM/MsJ | Musmus IGKV4-60*01 P | 98.58 | 292 | GAAATTGTGCTCACCCAGTCTCCAGCACTCATGGCTGCATCTCCAGGGGAGAAGGTCAGCATCACCTGCAGTGTCAGCTCAAGTATAAGTTCCAGCCACTTGCACTGGTACCAGCAGAAGTCAGGAACCTCCCCCAAACTCTGGATTTATGGCACATCCAACCTGGCTTCTGGAGTCCCTGCTCGCTTCAGTGGCAGTGGATCTGGGACCTCTTACTCTCTCACAATCAGCAGCATGGAGGCTGAAGATGCTGCCACTTATTACTGTCAACAGTGGAGTAGTTACCCAACCA |
| IGKV3-9*01_S0307_MSM | MSM/MsJ | Musmus IGKV3-9*01 F | 91.07 | 299 | GACATTGTGCTGACCCAATCTCCAGCTTCTTTGGCTGTGTCTCTAGGGCAGAGGGCCACCATCTCCTGCAGAGCCAGCGAAAGTGTCAGTATTCGTGGTACTAGTTTAATGCACTGGTACCAACAGAAACCAGGACAGCCACCCAAAGTCCTCATCTATCTTGCATCCAACCTAGAATCTGGGGTTCCTGCAAGGTTTAGTGGCAGTGGGTCTGGGACAGGCTTCACCCTCAATATCCATCCTGTGGAGGAAGATGATGCTGCAACCTATTACTGTCAGCAAAGTAGGGAATATCCTCC |
| IGKV4-50*01_S6568_MSM | MSM/MsJ | Musmus IGKV4-50*01 F | 96.74 | 286 | GAAAATGTGCTCACCCAGTCTCCAGCAATCATGTCTGCAACTCTAGGGGAGAAGGTCACCATGAGCTGCAGGGCCAGCTCAAATGTAAAGAACATGTACTGGTACCAGCAGAAGTCAGGTGCCTCCCCCAAACTATGGATTTATTACACATCCAACCTGGCTTCTGGAGTCCCAGCTCGCTTCAGTGGCAGTGGGTCTGGGACCTCTTATTCTCTCACAATCAGCAGCGTGGAGGCTGAAGATGCTGCCACTTATTACTGCCAGCAGTTTACTAGTTCCCCATCCA |
| IGKV9-123*01_S7058_MSM | MSM/MsJ | Musmus IGKV9-123*01 F | 99.28 | 285 | GACATCCAGATGATTCAGTCTCCATCGTCCATGTTTGCCTCTCTGGGAGACAGAGTCAGTCTCTCTTGCCGGGCTAGTCAGGGCATTAGAGGTAATTTAGACTGGTATCAGCAGAAACCAGGTGGAACTATTAAACTCCTGATCTACTCCACATCTAATTTAAATTCTGGTGTCCCATCAAGGTTCAGTGGCAGTGGGTCTGGGTCAGATTATTCTCTCACCATCAGCAGCCTAGAGTCTGAAGATTTTGCAGACTATTACTGTCTACAGCGTAATGCGTATCCT |
| IGKV11-125*01_MSM | MSM/MsJ | Musmus IGKV11-125*01 F | 100 | 287 | GATGTCCAGATGATTCAGTCTCCATCCTCCCTGTCTGCATCTTTGGGAGACATAGTCACCATGACTTGCCAGGCAAGTCAGGGCACTAGCATTAATTTAAACTGGTTTCAGCAAAAACCAGGGAAAGCTCCTAAGCTCCTGATCTATGGTGCAAGCAACTTGGAAGATGGGGTCCCATCAAGGTTCAGTGGCAGTAGATATGGGACAGATTTCACTCTCACCATCAGCAGCCTGGAGGATGAAGATATGGCAACTTATTTCTGTCTACAGCATAGTTATCTCCCTCC |
| IGKV1-133*01_S2808_MSM | MSM/MsJ | Musmus IGKV1-133*01 F | 99.66 | 302 | GATGTTGTGATGACCCAGACTCCACTCACTTTGTTGGTTACCATTGGACAACCAGCCTCTATCTCTTGCAAGTCAAGTCAGAGCCTCTTATATAGTAATGGAAAAACCTATTTGAATTGGTTATTACAGAGGCCAGGCCAGTCTCCAAAGCGCCTAATCTATCTGGTGTCTAAACTGGACTCTGGAGTCCCTGACAGGTTCACTGGCAGTGGATCAGGAACAGATTTTACACTGAAAATCAGCAGAGTGGAGGCTGAGGATTTGGGAGTTTATTACTGCGTGCAAGGTACACATTTTCCTCA |
| IGKV3-1*01_S2114_MSM | MSM/MsJ | Musmus IGKV3-1*01 F | 99.66 | 299 | GACATTGTGCTCACCCAATCTCCAGCTTCTTTGGCTGTGTCTCTAGGGCAGAGAGCCACCATCTCCTGCAGAGCCAGTGAAAGTGTTGAATATTATGGCACAAGTTTAATGCAGTGGTACCAACAGAAACCAGGACAGCCACCCAAAATCCTCATCTATGCTGCATCCAACGTAGAATCTGGGGTCCCTGCCAGGTTTAGTGGCAGTGGGTCTGGGACAGACTTCAGCCTCAACATCCATCCTGTGGAGGAGGATGATATTGCAATGTATTTCTGTCAGCAAAGTAGGAAGGTTCCTTC |

| | | | | | |
|---|---|---|---|---|---|
| IGKV4-92*01_S1372_MSM | MSM/MsJ | Musmus IGKV4-92*01 F | 97.52 | 291 | GAAATTGTTCTCACCCAGTCTCCAATATCCATAACTGCATCTCAAGGGGAGAAGGTCACCATCACCTGCCGTGCCAGCTCAAGTATAAGTTCCAATTACTTACACTGGTACCAGCAGAAGCCAGAATCCTCCCCTAAACTTTTGATTTATAGGACATCCATCCTGGCATCTGGAGTACTAGACAGCTTCAGTGGCAGTGGGTCTGAGAGCTCTTACACTCTGACAATCAGCTGCATGCAGGCCGAAGTTGCTTCCACTTACTATTGTCAGCAGGGGAGTAGTAGTCCACCA |
| IGKV3-12*01_S5721_MSM | MSM/MsJ | Musmus IGKV3-12*01 F | 97.59 | 299 | GACATTGTGCTGACACAGTCTCCTGCTTCCTTAGCTGTATCTCTGGAGCAGAGGGCCACCATCTCATGCAGGGCCAGCCAAAGTGTCAGTACATCTAGCTATAGTTACATGCACTGGTACCAACAGAAACCAGGACAGCCACCCAAACTCCTCATCTATTTTGCATCCAACCTAGAATCTGGGGTCCCTGCCAGGTTCAGTGGCAGTGGGTCTGGGACAGACTTCACCCTCAACATCCATCCTGTGGAGGAGGAGGATGCTGCAACATATTACTGTCAGCACAATAGGGAGATTCCTCC |
| IGKV4-57*01_S8859_MSM | MSM/MsJ | Musmus IGKV4-57-1*01 F | 93.26 | 286 | CAAATTGTTCTCACCCAGTCTCCAGCAATCATGGCTGCATCTCCAGGGGAGAAGTTCACCATAACCTGCAGTGCCAGCTCAAGTGTAAGTTACATGCACTGGTTCCAGCAGAAGCCAGGCACTTCTCCCAAACTCTGGATTTATAGGACATCGAACCTGGCTTCTGGAGTCCCTGCTCGCTTCAGTGGCAGTGGGTCTGGGACCTCTTATTCTCTCACAATCAGCAGCATGGAGGCTGAAGATGCTGCCACTTATTACTGCCTGCAGCAGAGTAGTTACCCACCCA |
| IGKV8-18*01_S2898_MSM | MSM/MsJ | Musmus IGKV8-18*01 ORF | 99.66 | 310 | GACATTGTGATGACCCAGTCTCCATCCTCCCTGGCTGTGACAGCAGGAGAGAAGGTCACTATGAGATGCAAGTCCAGTCAGAGTCTTTTGTGGAGTGTAAACCAAAAGAACTACTTATCCTGGTACCAGCAGAAACAAGGGCAGCCTCCTAAACTGCTTATCTATGGGGCATCCATTAGAGAATCTTGGGTCCCTGATCGATTCACAGGAAGTGGATCTGGGACAGACTTCACTCTCACCATTAGCAATGTGCATGCTGAAGACCTAGCAGTTTATTACTGTCAGCACAATCATGGCAGCTTTCTCCCCC |
| IGKV12-38*01_S3762_MSM | MSM/MsJ | Musmus IGKV12-38*01 F | 98.57 | 287 | GACATCCAGATGACTCAGTCTCCAGCCTCCCTGGCTGCATCTGTGGGAGAAACTATCACCATCACATGTCAAGCAAGTGAGAACATTTACTACAGTTTAGCATGGTATCAGCAGAAGCAAGGGAAATCTCCTCAGCTCCTGATCTATAATGCAAACAGCTTGGAAGATGGTGTCCCATCAAGGTTCAGTGGCAGTGGATCTGGGACACAGTATTCTATGAAGATCAACAGCATGCAGCCTGAAGATACTGCAACTTATTTCTGTAAACAGGCTTATGACTTTCCTCC |
| IGKV5-39*01_S0817_MSM | MSM/MsJ | Musmus IGKV5-39*01 F | 99.64 | 287 | GACATTGTGATGACTCAGTCTCCAGCCACCCTGTCTGTGACTCCAGGAGATAGAGTCTCTCTTTCCTGCAGGGCCAGCCAGAGTATTAGCAACTACTTACACTGGTATCAACAAAAATCACATGAGTCTCCAAGGCTTCTCATCAAATATGCTTCCCAATCCATCTCTGGGATCCCCTCCAGGTTCAGTGGCAGTGGATCAGGGTCAGATTTCACTCTCAGTATCAACAGTGTGGAACCTGAAGATGTTGGAGTGTATTACTGTCAAAATGGTCACAGCTTTCCTCC |
| IGKV5-45*01_S2588_MSM | MSM/MsJ | Musmus IGKV5-45*01 F | 98.92 | 287 | GATATTGTGCTAACTCAGTCTCCAGCCACCCTGTCTGTGACTCCAGGAGATAGAGTCAGTCTTTCCTGCAGGGCCAGCCAAAGTATTAGCAACTACCTACACAGGTATCAACAAAAATCACATGAGTCTCCAAGGCTTCTCATCAAGTATGCTTCCCAGTCCATCTCTGGGATCCCCTCCAGGTTCAGTGGCAGTGGATCAGGGACAGATTTCACTCTCAGTATCAACAGTGTGGAGACTGAAGATTTTGGAATGTATTTTTGTCAACAGAGTAACAGCTGGCCTCA |
| IGKV6-15*01_S8899_MSM | MSM/MsJ | Musmus IGKV6-15*01 F | 97.13 | 287 | GACATTGTGATGACCCAGTCTCAAAAATTCATGTCCACATCAGTAGGAGACAGGGTCAGCGTCACCTGCAAGGCCAGTCAGTATGTGGGTACTTATGTAGCCTGGTATCAACAGAAACCAGGGCAATCTCCTAAAGCACTGATTTACTCGGCATCCTACCGGTACACTGGAGTCACTGATCGCTTCACAGGCAGTGGATCTGGGACAGATTTCACTCTCACCATTAGCAATGTGCAGTCTGAAGACTTAGCAGATTATTTCTGTCAGCAATATAGCAGCTCTCCTCT |

| | | | | | |
|---|---|---|---|---|---|
| IGKV13-84*01_S4500_MSM | MSM/MsJ | Musmus IGKV13-84*01 F | 98.92 | 287 | GACATCCAGATGACACAATCTTCATCCTCCTTGTCTGTATCTCTAGGAGACAGAGTTACCATTACTTGCAAGGCAAGTGAGGACATATATAATCGGTTAGCCTCGTATCAGCAGAAACCAGGAAATGCTCCTAGGCTCTTAATATCTGGTGCAACCAGTTTGGAAACTGGGGTTCCTTCAAGATTCAGTGGCAGTGGATCTGGAAAGGATTACACTCTCAGCATTACCAGTCTTCAGACTGAAGATGTTGCTACTTATTACTGTCAACAGTATTGGAGTACTCCTCC |
| IGKV17-121*01_S5117_MSM | MSM/MsJ | Musmus IGKV17-127*01 F | 98.57 | 287 | GAAACAACTGTGACCCAGTCTCCAGCATCCCTGTCCATGGCTACAGGAGAAAAAGTCACTATCAGATGCATAACCAGCACTGATATTGATGATGATATGAACTGGTACCAGCAGAAGCCAGGGGAACCTCCTAAGCTCCTTATTTCAGAAGGCAATACTCTTCGTCCTGAAGTCCCATCCCGATTCTCCAGCAGTGGCTATGGTACAGATTTTGTTTTTACAATTGAAAACACTCTCTCAGAAGATGTTGCAGATTACTACTGTTTGCAAAGTGATAACTTGCCTCT |
| IGKV11-125*01_S1984_MSM | MSM/MsJ | Musmus IGKV11-106*02 F | 96.06 | 286 | GATGTTCAAATGACCCAGTCTCCATCCTCCCTGTCTGCATCTTTGGGAGAGAGAGTCTCCCTGACTTGCCAGGCAAGTCAGAGCATTAGCAATTATTTAAACTGGTTTCAGCAAACACCAGGGAAAGCTCCTAGGCTCTTGATCTATGATGCAAACAACTTGGAAGATGGGGTCCCTTCAAGGTTCAGTGGCACTGGATATGGGACAGATTTCACTTTCACCATCAGCAGCCTGGAGGAAGAAGATGTCGCAACTTATTTCTGTCTACAGCATAAGTATCTCCCTC |
| IGKV3-9*01_S4460_MSM | MSM/MsJ | Musmus IGKV3-9*01 F | 92.1 | 299 | GACATTGTGCTGACCCAATCTCCAGCTTCTTTGGCTGTGTCTCTAGGGCAGAGGGCCACCATCTCCTGCAAGGCCAGCCAAAGTGTCAGTTTTGCTGGTACTAGTTTAATGCACTGGTACCAACAGAAACCAGGACAGCCACCCAAAGTCCTCATCTATCGTGCATCCAACCTAGAATCTGGGGTTCCTACCAGGTTTAGTGGCAGTGGGTCTGGGACAGACTTCACCCTCAATATCCACCCTGTGGAGGAAGATGATGCTGCAACCTATTACTGTCAGCAAAGTAGGGAATATCCTCC |
| IGKV12-41*01_S5467_MSM | MSM/MsJ | Musmus IGKV12-41*01 F | 96.42 | 284 | GACATCCAGATGACTCAGTCTCCAGCTTCCCTATCTGCATCTGTGGGAGAAACTGTCACCATCACATGTCGAGCAAGTGGGAATATTTACAGTAATTTAGCATGGTATCAGCAGAAACAGGGAAAATCTCCTCAGCTCCTGGTCTATAATGCAAAAACCTTAGCAGAAGGTGTGCCATCAAGGTTCAGTGGCAGTGGATCAGGAACACAATATTCTCTCAAGATCAACAGCCTACAGCCTGAAGATTTTGGGAGTTATTACTGTCAACATTATTATAATACTCC |
| IGKV1-35*01_S0416_MSM | MSM/MsJ | Musmus IGKV1-35*01 ORF | 97.28 | 301 | GACATTGTGATGACCCAGACTCCACTCACTTTATCAGCTACCATTGGACAATCAGCCTCCATCTCTTGCAGGTCAAGTCAGAGTCTCTTACATAGTAATGGAAACACATACTTGAATTGGTTTCTACAGAGGCCAGGCCAATCTCCACAGCTTCTGATTTATGGGGTGTTTGAACGGGAATCTGGGGTTCCTGACAGGTTCAGTGGCAGTGGGTCAGGAACAGATTTCACACTCAAGATCAGCAGAGTGGAGGCTGAGGATTTGGGAGTTTATTACTGCATGCAAGCTACCTATGAACCTC |
| IGKV4-56*01_S4810_MSM | MSM/MsJ | Musmus IGKV4-59*01 F | 94.57 | 286 | CAAATTGTTCTCACCCAGTCTCCAGCAATCATTTCTGCATCTCCGGGGCAGAAAGTCACCATAACCTGCAGTGCCAGCTCCAGTGTAAATTACATGCACTGGTTCCAGCAGAAGCCAGGATCCTCCCCCAAACTCTGGATTTATGACACATCCAAACTGGCTCCTGGAGTCCCTGCTCGCTTCAGTGGCAGTGGGTCTGGGACCTCTTACTCTCTCACAATCAGCAGCATGGAGGCTGAAGATGCTGCCACTTATTACTGCCAGCAGTGGAGTAGTTACCCACCCA |
| IGKV4-56*01_S9438_MSM | MSM/MsJ | Musmus IGKV4-59*01 F | 94.57 | 286 | CAAATTGTTCTCACCCAGTCTCCAGCAATCATTTCTGCATCTCCGGGGCAGAAAGTCACCATAACCTGCAGTGCCAGCTCCAGTGTAAATTACATGCACTGGTTCCAGCAGAAGCCAGGATCCTCCCCCAAACTCTGGATTTATGACACATCCAAACTGGCTCCTGGAGTCCCTGCTCGCTTCAGTGGCAGTGGGTCTGGGACCTCTTACTCTCTCACAATCAGCAGCATGGAGGCTGAAGATGCTGCCACTTATTACTGCCAGCAGTGGAGTAGTTACCCGCTCA |

| | | | | | |
|---|---|---|---|---|---|
| IGKV4-59*01_S5858_MSM | MSM/MsJ | Musmus IGKV4-59*01 F | 97.1 | 286 | CAAATTGTTCTCACCCAGTCTCCAGCAATCATGTCTGCATCTCCAGGGGAGAAGGTCACCATGACCTGCAGTGCCAGCTCAAGTGTAAGTTACATGTACTGGTACCAGCAGAAGCCAGGATCCTCCCCCAAAAAATGGATTTATGACACATCCAACCTGGCTTCTGGAGTCCCCGCTCGCTTCAGTGGCAGTGGGTCTGGGACCTCTTATTCTCTCACAATCAGCAGCATGGAGGCTGAAGATGCTGCCACTTATTACTGCCAGCAGTGGAGTAGTTACCCACCCA |
| IGKV6-20*01_S2082_NOD | NOD/ShiLtJ | Musmus IGKV6-20*01 F | 99.64 | 287 | AACATTGTAATGACCCAATCTCCCAAATCCATGTCCATGTCAGTAGGAGAGAGGGTCACCTTGAGCTGCAAGGCCAGTGAGAATGTGGGTACTTATGTATCCTGGTATCAACAGAAACCAGAGCAGTCTCCTAAACTGCTGATATACGGGGCATCCAACCGGTACACTGGGGTCCCCGATCGCTTCACAGGCAGTGGATCTGCAACAGATTTCACTCTGACCATCAGCAGTGTGCAGGCTGAAGACCTTGCAGATTATTACTGTGGACAGAGTTACAGCTATCCTCC |
| IGKV8-21*01_S1419_NOD | NOD/ShiLtJ | Musmus IGKV8-21*01 F | 99.33 | 302 | GACATTGTGATGTCACAGTCTCCATCCTCCCTGGCTGTGTCAGCAGGAGAGAAGGTCACTATGAGCTGCAAATCCAGTCAGAGTCTGTTCAACAGTAGAACCCGAAAGAACTACTTGGCTTGGTACCAGCAGAAACCAGGGCAGTCTCCTAAACTGCTGATCTACTGGGCATCCACTAGGGAATCTGGGGTCCCTGATCGCTTCACAGGCAGTGGATCTGGGACAGATTTCACTCTCACCATCAGCAGTGTGCAGGCTGAAGACCTGGCAGTTTATTACTGCAAGCAATCTTATTATCTTCC |
| IGKV2-137*01_S0442_NOD | NOD/ShiLtJ | Musmus IGKV2-137*01 F | 93.54 | 302 | GATATTGTGATGACTCAGGCTGCACCCTCTGTACCTGTCACTCCTGGAGAGTCAGTATCCATCTCCTGCAGGTCTAGTACGAGTCTCCTGCACAGTAGTGGCAAGCATAGGTTGTATTGGTTCCTACAGAGGCCAGGCCAGTCTCCTCAGCTCCTGATATATTATATGTCCAACCTTGCCTCAGGAGTCCCAGACAGGTTCAGTGGCAGTGGGTCAGGAACTGATTTCACACTGAGAATCAGTAGAGTGGAGGCTGAGGATTTTGGTGTTTATTATTGTATGCAAAGTCTAGAATATCCTTT |
| IGKV12-46*01_S4541_NOD | NOD/ShiLtJ | Musmus IGKV12-46*01 F | 97.13 | 287 | GACATCCAGATGACTCAGTCTCCAGCCTCCCTATCTGCATCTGTGGGAGAAACTGTCACCATCACATGTCGAGCAAGTGAGAATATTTACAGTAATTTAGCATGGTATCAGCAGAAACAGGGAAAATCTCCTCAGCTCCTGGTCTATGCTGCAACAAATTTAGCAGATGGTGTGCCATCAAGGTTCAGTGGCAGTGGATCAGGCACACAGTTTTCTCTGAAGATCAACAGCCTGCAGCCTGAAGATTTTGGGAGTTATTACTGTCAACATTTTTATGGTACTCCTCC |
| IGKV12-46*01_S1254_NOD | NOD/ShiLtJ | Musmus IGKV12-46*01 F | 94.62 | 287 | GACATCCAGATGACTCAGTCTCCAGCTTCCCTGTCTGCATCTGTGGGAGAAACTGTCACCATCACATGTCGAGCAAGTGAGAATATTGACAGTTATTTAGCATGGTATCAGCAGAAACAGGGAAAATCTCCTCAGCTCCTGGTCTATGCTGCAACACTCTTAGCAGATGGTGTGCCATCAAGGTTCAGTGGCAGTGGATCAGGCACACAGTATTCTCTCAAGATCAACAGCCTGCAGTCTGAAGATGTTGCGAGATATTACTGTCAACATTATTATAGTACTCCTCC |
| IGKV6-15*01_S9105_NOD | NOD/ShiLtJ | Musmus IGKV6-15*01 F | 97.13 | 287 | GACATTGTGATGACCCAGTCTCAAAAATTCATGTCCACATCAGTAGGAGACAGGGTCAGCATCAGCTGCAAGGCCAGTCAGAATGTGGGTAATATTATAGCCTGGTATCAACAGAAACCAGGGCAATCTCCTAAAGCACTGATTTACTTGGCATCCTACCGGTACAGTGGAGTCCCTGATCGCTTCACAGGCAGTGGATCTGGGACAGATTTCACTCTCACCATTAGCAATGTGCAGTCTGAAGACTTGGCAGAGTATTTCTGTCAGCAATATAGCAGCTCTCCTCT |
| IGKV12-44*01_NOD | NOD/ShiLtJ | Musmus IGKV12-44*01 F | 100 | 287 | GACATCCAGATGACTCAGTCTCCAGCCTCCCTATCTGCATCTGTGGGAGAAACTGTCACCATCACATGTCGAGCAAGTGAGAATATTTACAGTTATTTAGCATGGTATCAGCAGAAACAGGGAAAATCTCCTCAGCTCCTGGTCTATAATGCAAAAACCTTAGCAGAAGGTGTGCCATCAAGGTTCAGTGGCAGTGGATCAGGCACACAGTTTTCTCTGAAGATCAACAGCCTGCAGCCTGAAGATTTTGGGAGTTATTACTGTCAACATCATTATGGTACTCCTCC |

| | | | | | |
|---|---|---|---|---|---|
| IGKV6-b*01_NOD | NOD/ShiLtJ | Musmus IGKV6-b*01 F | 100 | 287 | AGTATTGTGATGACCCAGACTCCCAAATTCCTGCCTGTATCAGCAGGAGACAGGGTTACCATGACCTGCAAGGCCAGTCAGAGTGTGGGTAATAATGTAGCCTGGTACCAACAGAAGCCAGGACAGTCTCCTAAACTGCTGATATACTATGCATCCAATCGCTACACTGGAGTCCCTGATCGCTTCACTGGCAGTGGATCTGGGACAGATTTCACTTTCACCATCAGCAGTGTGCAGGTTGAAGACCTGGCAGTTTATTTCTGTCAGCAGCATTATAGCTCTCCTCC |
| IGKV14-126*01_NOD | NOD/ShiLtJ | Musmus IGKV14-126*01 F | 100 | 287 | GACATCAAGATGACCCAGTCTCCATCCTCCATGTATGCATCGCTGGGAGAGAGAGTCACTATCACTTGCAAGGCGAGTCAGGACATTAAAAGCTATTTAAGCTGGTACCAGCAGAAACCATGGAAATCTCCTAAGACCCTGATCTATTATGCAACAAGCTTGGCAGATGGGGTCCCATCAAGATTCAGTGGCAGTGGATCTGGGCAAGATTATTCTCTAACCATCAGCAGCCTGGAGTCTGACGATACAGCAACTTATTACTGTCTACAGCATGGTGAGAGCCCTCC |
| IGKV5-45*01_S0430_NOD | NOD/ShiLtJ | Musmus IGKV5-45*01 F | 99.64 | 287 | GATATTGTGCTAACTCAGTCTCCAGCCACCCTGTCTGTGACTCCAGGAGATAGAGTCAGTCTTTCCTGCAGGGCCAGCCAAAGTATTAGCAACTACCTACACTGGTATCAACAAAAATCACATGAGTCTCCAAGGCTTCTCATCAAGTATGCTTCCCAGTCCATCTCTGGGATCCCCTCCAGGTTCAGTGGCAGTGGATCAGGGACAGATTTCACTCTCAGTATCAACAGTGTGGAGACTGAAGATTTTGGAATGTATTTCTGTCAACAGAGTAACAGCTGGCCTCA |
| IGKV10-94*03_NOD | NOD/ShiLtJ | Musmus IGKV10-94*03 F, or Musmus IGKV10-94*08 F | 100 | 287 | GATATCCAGATGACACAGACTACATCCTCCCTGTCTGCCTCTCTGGGAGACAGAGTCACCATCAGTTGCAGGGCAAGTCAGGATATTAGCAATTATTTAAACTGGTATCAGCAGAAACCAGATGGAACTGTTAAACTCCTGATCTACTACACATCAAGATTACACTCAGGAGTCCCATCAAGGTTCAGTGGCAGTGGGTCTGGGACAGATTATTCTCTCACCATCAGCAACCTGGAACCTGAAGATATTGCCACTTACTATTGTCAGCAGTATAGTAAGCTTCCTCC |
| IGKV12-46*01_S4026_NOD | NOD/ShiLtJ | Musmus IGKV12-46*01 F | 99.64 | 287 | GACATCCAGATGACTCAGTCTCCAGCCTCCCTATCTGTATCTGTGGGAGAAACTGTCACCATCACATGTCGAGCAAGTGAGAATATTTACAGTAATTTAGCATGGTATCAGCAGAAACAGGGAAAATCTCCTCAGCTCCTGGTCTATGCTGCAACAAACTTAGCAGATGGTGTTCCATCAAGGTTCAGTGGCAGTGGATCAGGCACACAGTATTCCCTCAAGATCAACAGCCTGCAGTCTGAAGATTTTGGGAGTTATTACTGTCAACATTTTTGGGGTACTCCTCC |
| IGKV8-30*01_S6795A_NOD | NOD/ShiLtJ | Musmus IGKV8-30*01 F | 95.59 | 295 | GACATTGTGATGTCACAGTCTCCATCCGCCCTAGCTGTGTCAGTTGGAGAGAAGGTCACTATGAGCTGCAAGTCCAGTCAGAGCCTTTTATATAGTAGCAATCAAAAGAACTACTTGGCCTGGTACCAGCAGAAACCAGGGCAGTCTCCTAAACTGTTAATCTACTGGGCATCCACTAGGGAATCTGGGGTCCCTGACCGCTTCACAGGCAGTAGATCAGGGACAGATTTCACTCTCACCATCAGCAGTGTGCAGGCTGAAGACCTGGCCGTTTATTACTGCAAGCAATCTTATA |
| IGKV1-135*01_S3001_NOD | NOD/ShiLtJ | Musmus IGKV1-135*01 F | 95.92 | 302 | GATGTTGTGATGACTCAGACCCCACTCACTTTGTCGGTTACCATTGGACAACCAGCCTCCATCTCTTGCAAATCAAGTCAGAGCCTCTTACATAGTAATGGAAAGACATATTTGAATTGGTTATTACAGAGGCCAGGCCAGTCTCCAAAGCTCCTAATCTATCTGGTGTCTAAACTGGAATCTGGAGTCCCTGACAGGTTCAGTGGCAGTGGATCAGGGACAGATTTCACACTGAAAATCAGCAGAGTGGAGGCTGAGGATTTGGGAGTTTATTACTGCTTGCAAGCTACACATTTTCCTCA |
| IGKV8-19*01_NOD | NOD/ShiLtJ | Musmus IGKV8-19*01 F | 100 | 305 | GACATTGTGATGACACAGTCTCCATCCTCCCTGACTGTGACAGCAGGAGAGAAGGTCACTATGAGCTGCAAGTCCAGTCAGAGTCTGTTAAACAGTGGAAATCAAAAGAACTACTTGACCTGGTACCAGCAGAAACCAGGGCAGCCTCCTAAACTGTTGATCTACTGGGCATCCACTAGGGAATCTGGGGTCCCTGATCGCTTCACAGGCAGTGGATCTGGAACAGATTTCACTCTCACCATCAGCAGTGTGCAGGCTGAAGACCTGGCAGTTTATTACTGTCAGAATGATTATAGTTATCCTCC |

| | | | | | |
|---|---|---|---|---|---|
| IGKV1-135*01_S7405_NOD | NOD/ShiLtJ | Musmus IGKV1-135*01 F | 95.24 | 302 | GATGTTGTGATGACCCAGACTCCACTGTCTTTGTCGGTTACCATTGGACAACCAGCCTCCATCTCTTGCAAGTCAAGTCAGAGCCTCTTACATAGTGATGGAAAGACATATTTGAATTGGTTATTACAGAGGCCAGGCCAGTCTCCAAAGCTCCTAATCTATCTGGTGTCTAAACTGGAATCTGGCATCCCTGACAGGTTCAGTGGCAGTGGATCAGGGACAGATTTCACACTGAAAATCAGCAGAGTGGAGGTTGAGGATTTGGGAGTTTATTACTGCTTGCAACATACACATTTTCCTCA |
| IGKV8-34*01_S3615_NOD | NOD/ShiLtJ | Musmus IGKV8-34*01 F | 99.33 | 305 | GACATTTTGATGACTCAGTCTCCATCCTCCCTGACTGTGTCAGCAGGAGAGAAGGTCACTATGAGCTGCAAGTCCAGTCAGAGTCTTTTAGCTAGTGCCAACCAAAATAACTACTTGGCCTGGCACCAGCAGAAACCAGGACGATCTCCTAAAATGCTGATAATTTGGGCATCCACTAGGGTATCTGGAGTCCCTGATCGCTTCATAGGCAGTGGATCTGGGACGGATTTCACTCTGACCATCAACAGTGTGCAGGCTGAAGATCTGGCTGTTTATTACTGTCAGCAGTCCTACAGCGCTCCTAC |
| IGKV15-103*01_S0817_NOD | NOD/ShiLtJ | Musmus IGKV15-103*01 ORF | 97.85 | 287 | GACATCCAGATGAACCAGTCTCCATCCAGTCTGTCTGCATCCCTCGGAGACACAATTACCATCACTTGCCGTGCCAGTCAGAACATTAATATTTGGTTAAGCTGGTACCAGCAGAAACCAGGAAATATTCCTAAACTATTGATCTATAAGGCTTCCAACTTGCACACAGGCGTCCCATCAAGGTTTAGTGGCAGTGGATCTGGAACAGATTTCACATTAACCATCAGCAGTCTGCAGCCTGAAGACATTGCCACTTACTACTGTCTACAGGGTCAAAGTTATCCTCT |
| IGKV12-41*01_S6800_NOD | NOD/ShiLtJ | Musmus IGKV12-41*01 F | 97.13 | 287 | GATATCCAGATGACTCAGTCTCCAGCCTCCCTATCTGCATCTGTGGGAGAAACTGTCACCATCACATGTCGAGCAAGTGGGAATATTCACAATTATTTAGCATGGTATCAGCAGAAACAGGGAAAATCTCCTCAGCTCCTGGTCTATAATGCAAAAACCTTAGCGGAAGGTGTGCCATCAAGGTTCAGTGGCAGTGGATCAGGAACACAATATTCTCTCAAGATCAACAGCCTGCAGCCTGAGGATTTTGGGAGTTATTACTGTCATCATTATTATAGTACTCCTCC |
| IGKV8-24*01_S8843_NOD | NOD/ShiLtJ | Musmus IGKV8-24*01 F | 98.32 | 305 | GACATTGTGATGACACAGTCTCCATCCTCCCTGGCTATGTCAGTAGGACAGAAGGTCACTATGAGCTGCAAGTCCAGTCAGAGCCTTTTAAGTAGTAGCAATCAAAAGAACTATTTGGCCTGGTACCAGCAGAAACCAGGACAGTCTCCTAAACTTCTGGTATACTTTGCATCCACTAGGGAATCTGGGGTCCCTGATCGCTTCATAGGCAGTGGATCTGGGACAGATTTCACTCTTACCATCAGCAGTGTGCAGGCTGAAGACCTGGCACTTTATTACTGTCAGCAACATTATAGCACTCCTCC |
| IGKV1-110*02_NOD | NOD/ShiLtJ | Musmus IGKV1-110*02 F | 100 | 302 | GATGTTGTGATGACCCAAACTCCACTCTCCCTGCCTGTCAGTCTTGGAGATCAAGCTTCCATCTCTTGCAGATCTAGTCAGAGCCTTGTACACAGCAATGGAAACACCTATTTATATTGGTACCTGCAGAAGCCAGGCCAGTCTCCAAAGCTCCTGATCTACAGGGTTTCCAACCGATTTTCTGGGGTCCCAGACAGGTTCAGTGGCAGTGGATCAGGGACAGATTTCACACTCAAGATCAGCAGAGTGGAGGCTGAGGATCTGGGAGTTTATTTCTGCTTTCAAGGTACACATGTTCCTCA |
| IGKV8-16*01_S7495_NOD | NOD/ShiLtJ | Musmus IGKV8-16*01 F | 99.33 | 305 | GAAATTGTGTTGACGCAGTCTATACCATCCCTGACTGTGTCAGCAGGAGAGAGGGTCACTATCAGCTGCAAATCCAATCAGAATCTTTTATGGAGTGGAAACCAAAGGTACTGTTTGGTCTGGCACCAGTGGAAACCAGGGCAAACTCCTACACCGTTGATCACCTGGACATCTGATAGATACTCTGGAGTCCCTGATCGTTTCATAGGCAGTGGATCTGTGACAGATTTCACTCTGACCATCAGCAGTGTGCAGGCTGAAGATGTGGCAGTTTATTTCTGTCAGCAGCATTTACACATTCCTCC |

| | | | | | |
|---|---|---|---|---|---|
| IGKV4-80*01_S5342_NOD | NOD/ShiLtJ | Musmus IGKV4-80*01 F | 99.64 | 281 | CAAATTGTTCTCACCCAGTCTCCAGCAATCATGTCTGCATCTCTAGGGGAGGAGATCACCCTAACCTGCAGTGCCAGCTCGAGTGTAAGTTACATGCACTGGTTCCAGCAGAAGTCAGGCACTTCTCCCAAACTCTTGATTTATAGCACATCCAACCTGGCTTCTGGAGTCCCTTCTCGCTTCAGTGGCAGTGGGTCTGGGACCTTTTATTCTCTCACAATCAGCAGTGTGGAGGCTGAAGATGCTGCCGATTATTACTGCCATCAGTGGAGTAGTTATCC |
| IGKV4-62*01_S2199_NOD | NOD/ShiLtJ | Musmus IGKV4-62*01 ORF | 96.74 | 286 | CAAATTGTTCTCACCCAGTCTCCAGCAATCATGTCTGCATCTCCAGGGGAGAAGGTCACCATGACCTGCAGTGCCAGCTCAAGTGTAAGTTACATGTACTGGTACCAGCAGAAGCCAGGATCCTCCCCCAGACTCTGGATTTATGACACATCCAACCTGGCTTCTGGAGTCCCCGCTCGCTTCAGTGGCAGTAGGTCTGGGACCTCTTATTCTCTCACAATCAGCAGCATGGAGGCTGAAGATGCTGCCACTTATTACTGCCATCAGCGGAGTAGTTACCCATTCA |
| IGKV5-43*01_S9511_NOD | NOD/ShiLtJ | Musmus IGKV5-43*01 F | 99.64 | 287 | GATATTGTGCTAACTCAGTCTCCAGCCACCCTGTCTGTGACTCCAGGAGATAGCGTCAGTCTTTCCTGCAGGGCCAGCCAAAGTATTAGCAACAACCTACACTGGTATCAACAAAAATCACATGAGTCTCCAAGGCTTCTCATCAAGTATGCTTCCCAGTCCATCTCTGGGATCCCCTCCAAGTTCAGTGGCAGTGGATCAGGGACAGATTTCACTCTCAGTATCAACAGTGTGGAGACTGAAGATTTTGGAATGTATTTCTGTCAACAGAGTAACAGCTGGCCTCA |
| IGKV3-1*01_S7345_NOD | NOD/ShiLtJ | Musmus IGKV3-1*01 F | 98.63 | 299 | GACATTGTGCTCACCCAATCTCCAGCTTCTTTGGCTGTGTCTCTAGGGCAGAGTGTCACCATCTCCTGCAGAGCCAGTGAAAGTGTTGAATATTATGGCACTAGTTTAATGCAGTGGTACCAACAGAAACCAGGACAGCCACCCAAACTCCTCATCTATGGTGCATCCAACGTAGAATCTGGGGTCCCTGCCAGGTTTAGTGGCAGTGGGTCTGGGACAGACTTCAGCCTCAACATCCATCCTGTGGAGGAGGATGATATTGCAATGTATTTCTGTCAGCAAAGTAGGAAGGTTCCTTC |
| IGKV4-74*01_S0897_NOD | NOD/ShiLtJ | Musmus IGKV4-74*01 F | 94.68 | 292 | CAGATTGTTCTCACCCAGTCTCCAGCAATCATGTCTGCATCTCCAGGGGAGAAGGTCACCATGACCTGCAGGGCCAGCTCAAGTGTAAGTTCCAGTTACTTGCACTGGTACCAGCAGAAGCCAGGATCTTCCCCCAAACTCTGGATTTATAGCACATCCAACCTGGCTTCAGGAGTCCCAGCTCGCTTCAGTGGCAGTGGGTCTGGGACCTCTTACTCTCTCACAATCAGCAGTGTGGAGGCTGAGGATGCTGCCACTTATTACTGCCAGCAGTATGATAGTTCCCCACTCA |
| IGKV1-117*01_S1348_NOD | NOD/ShiLtJ | Musmus IGKV1-117*01 F | 99.32 | 301 | GATGTTGTGATGACCCAAACTCCACTCTCCCTGCCTGTCAGTCTTGGAGATCAAGCCTCCATCTCTTGCAGATCTAGTCAGAGCATTGTACATAGTAATGGAAACACCTATTTAGAATGGTACCTGCAGAAGCCAGGCCAGTCTCCAAAGCTCCTGATCTACAAAGTTTCCAACCGATTTTCTGGGGTCCCAGACAGGTTCAGTGGCAGTGGATCAGGGACAGATTTCACACTCAAGATCAGCAGAGTGGAGGCTGAGGATCTGGGAGTTTATTACTGCTTTCAAGGTTCACATGTTCCTC |
| IGKV2-137*01_S6567_NOD | NOD/ShiLtJ | Musmus IGKV2-137*01 F | 98.64 | 302 | GATATTGTGATGACTCAGGCTGCACCCTCTGTACCTGTCACTCCTGGAGAGTCAGTATCCATCTCCTGCAGGTCTAGTAAGAGTCTCCTGCATAGTAATGGCAACACTTACTTGTATTGGTTCCTACAGAGGCCTGGCCAGTCTCCTCAGCTCCTGATATATCGGATGTCCAACCTTGCCTCAGGAGTCCCAGACAGGTTCAGTGGCAGTGGGTCAGGAACTGCTTTCACACTGAGAATCAGTAGAGTCGAGGCTGAGGATGTGGGTGTTTATTACTGTATGCAACATCTAGAGTATCCTTT |
| IGKV4-56*01_S5696_NOD | NOD/ShiLtJ | Musmus IGKV4-56*01 P | 96.38 | 286 | CAAATTGTTCTCACCCAGTCTCCAGCAATCATGTCTGCATCTCCAGGGCAGAAAGTCACCATAACCTGCAGTGCCAGCTCAAGTGTAAATTACATGCACTGGTACCAGCAGAAGCTAGGATCCTCCCCCAAACTCTGGATTTATGACACATCCAAACTGGCTCCTGGAGTCCCTGCTCGCTTCAGTGGCAGTGGGTCTGGGACCTCTTACTCTCTCACAATCAGCAGCATGGAGGCTGAAGATGCTGCCTCTTATTTCTGCCATCAGTGGAGTAGTTACCCACCCA |

| | | | | | |
|---|---|---|---|---|---|
| IGKV1-132*01_S5178_NOD | NOD/ShiLtJ | Musmus IGKV1-132*01 F | 97.96 | 302 | GATGTTGTGATGACCCAGACTCCACTGTCTTTGTCGGTTACCATTGGACAACCAGCCTCTATCTCTTGCAAGTCAAGTCAGAGCCTCTTATATAGTGATGGAAAGACATATTTGAATTGGTTACAACAGAGGCCAGGCCAGTCTCCAAAGCGCCTAATGTATCAGGTGTCCAAACTGGACCCTGGCATCCCTGACAGGTTCAGTGGCAGTGGATCAGAGACAGATTTTACACTTAAAATCAGCAGAGTGGAGGCTGAGGATTTGGGAGTTTATTACTGCTTGCAAGGTACATATTATCCTCA |
| IGKV4-59*01_S1025_NOD | NOD/ShiLtJ | Musmus IGKV4-59*01 F | 98.19 | 286 | CAAATTGTTCTCACCCAGTCTCCAGCAATCATGTCTGCATCTCCAGGGGAGAAGGTCACCATGACATGCAGTGCCAGCTCAAGTGTAAGTTACATGCACTGGTACCAGCAGAAGTCAGGCACCTCCCCCAAAAGATGGATTTATGACACATCCAAACTGGATTCTGGAGTCCCTGCTCGCTTCAGTGGCAGTGGGTCTGGGACCTCTTATTCTCTCACAATCAGCAGCATGGAGGCTGAAGATGCTGCCACTTATTACTGCTATCAGTGGAGTAGTTACCCACCCA |
| IGKV3-7*01_S6705_NOD | NOD/ShiLtJ | Musmus IGKV3-7*02 F | 97.59 | 299 | GACATTGTGCTGACACAGTCTCCTGCTTCCTTAGCTGTATCTCTGGGGCAGAGGGCCACCATCTCCTGCAGGGCCAGCAAAAGTGTCAGTACATCTAGCTATAGTTACATGCACTGGTACCAACAGAAACCAGGACAGCCACCCAAACTCCTCATCAAGTATGCATCCTACCTAGAATCTGGGGTTCCTGCCAGGTTCAGTGGCAGTGGGTCTGGGACAGACTTCACCCTCAACATCCATCCTGTGGAGGAGGAGGATGCTGCAACATATTACTGTCAGCACAGTAGGGAGTTTCCTCC |
| IGKV2-109*03_NOD | NOD/ShiLtJ | Musmus IGKV2-109*03 F | 100 | 302 | GATATTGTGATGACGCAGGCTGCATTCTCCAATCCAGTCACTCTTGGAACATCAGCTTCCATCTCTTGCAGTTCTAGTAAGAGTCTCCTACATAGTAATGGCATCACTTATTTGTATTGGTATCTGCAGAGGCCAGGCCAGTCTCCTCAGCTCCTGATATATCGGATGTCCAACCTTGCCTCAGGAGTCCCAGACAGGTTCAGTGGCAGTGGGTCAGGAACTGATTTCACACTGAGAATCAGCAGAGTGGAGGCTGAGGATGTGGGTGTTTATTACTGTGCTCAAATGCTAGAACGCCCTCC |
| IGKV3-4*01_S9382_NOD | NOD/ShiLtJ | Musmus IGKV3-4*01 F | 97.25 | 298 | GACATTGTGCTGACCCAATCTCCAGCTTCTTTGGCTGTGTCTCTAGGGCAGAGGGCCACCATCTCCTGCAGAGCCAGCGAAAGTGTTGATTATGATGGCGATAGTTATATGAACTGGTACCAACAGAAACCAGGACAGCCACCGAAACTCCTCATCTATGCTGCATCCAATCTAGAATCTGGGATCCCTGCCAGGTTTAGTGGCAGTGGGTCTGGGACAGACTTCACCCTCAACATTCATCCTGTGGAGGAGGAGGATGCTGCAACCTATTACTGTCTGCAAAGTAATGAGGCTCCTC |
| IGKV4-50*01_S2188_NOD | NOD/ShiLtJ | Musmus IGKV4-50*01 F | 97.1 | 286 | GAAAATGTGCTCACCCAGTCTCCAGCAATCATGTCTGCAACTCTAGGGGAGAAGGTCACCATGAGCTGCAGGGCCAGCTCAAATGTAAAGTACATGTACTGGTACCAGCAGAAGTCAGGTGCCTCCCCCAAACTATGGATTTATTACACATCCAACCTGGCTTCTGGAGTCCCAGCTCGCTTCAGTGGCAGTGGGTCTGGGACCTCTTATTCTCTCACAATCAGCAGCGTGGAGGCTGAAGATGCTGCCACTTATTACTGCCAGCAGTTTACTAGTTCCCCATCCA |
| IGKV4-53*01_S3575_NOD | NOD/ShiLtJ | Musmus IGKV4-53*01 F | 98.94 | 292 | GAAATTGTGCTCACCCAGTCTCCAGCACTCATGGCTGCATCTCCAGGGGAGAAGGTCACCATCACCTGCAGTGTCAGCTCAAGTATAAGTTCCAGCAACTTACACTGGTACCAGCAGAAGTCAGGAACCTCCCCCAAACCCTGGATTTATGGCACATCCAACCTTGCTTCTGGAGTCCCTGTTCGCTTCAGTGGCAGTGGATCTGGGACCTCTTATTCTCTCACAATCAGCAGCATGGAGGCTGAAGATGCTGCCACTTATTACTGTCAACAGTGGAGTAGTTACCCACTCA |

| | | | | | |
|---|---|---|---|---|---|
| IGKV1-133*01_S5139_NOD | NOD/ShiLtJ | Musmus IGKV1-133*01 F | 99.32 | 302 | GATGTTGTGATGACCCAGACTCCACTCACTTTGTCAGTTACCATTGGACAACCAGCCTCTATCTCTTGCAAGTCAAGTCAGAGCCTCTTATATAGTAATGGAAAAACCTATTTGAACTGGTTATTACAGAGGCCAGGCCAGTCTCCAAAGCGCCTAATCTATCTGGTGTCTAAACTGGACTCTGGAGTCCCTGACAGGTTCACTGGCAGTGGATCAGGAACAGATTTTACACTGAAAATCAGCAGAGTGGAGGCTGAGGATTTGGGAGTTTATTACTGCGTGCAAGGTACACATTTTCCTCA |
| IGKV2-109*04_NOD | NOD/ShiLtJ | Musmus IGKV2-109*04 F | 100 | 302 | GATATTGTGATGACGCAGGCTGCATTCTCCAATCCAGTCACTCTTGGAACGTCAGCTTCCATCTCCTGCAGGTCTAGTAAGAGTCTCCTACATAGTGATGGCATCACTTATTTGTATTGGTATCTGCAGAGGCCAGGCCAGTCTCCTCAGCTCCTGATATATCGGATGTCCAACCTTGCCTCAGGAGTCCCAGACAGGTTCAGTGGCAGTGGGTCAGGAACTGATTTCACACTGAGAATCAGCAGAGTGGAGGCTGAGGATGTGGGTGTTTATTACTGTGCTCAAATGCTAGAATTCCCTCC |
| IGKV19-93*01_S8964_NOD | NOD/ShiLtJ | Musmus IGKV19-93*01 F | 98.21 | 287 | GACATCCAGATGACACAGTCTCCATCCTCACTGTCTGCATCTCTGGGAGGCAAAGTCACCATCACTTGCAAGGCAAGCCAAGACATTAACAAGAATATAGCTTGGTACCAACACAAGCCTGGAAAAGGTCCTAGGCTGCTCATATGGTACACATCTACATTACAGCCAGGCATCCCATCAAGGTTCAGTGGAAGTGGGTCTGGGAGAGATTATTCCTTCAGCATCAGCAACCTGGAGCCTGAAGATATTGCAACTTATTACTGTCTACAGTATGATAATCTTCCACC |
| IGKV1-133*01_S3070_NOD | NOD/ShiLtJ | Musmus IGKV1-133*01 F | 95.92 | 302 | GATGTTGTGATGACCCAGACTCCACTCACTTTGTCTGTTACCATTGGACAGCCAGCTTCCATTTCTTGCAAGTCAAGTCAGAGCCTCTTATATAGTGATGGAAAAACCTATTTGAATTGGTTATTACAGAGTCCAGGCCAGTCTCCAAAGCTCCTAATCTATCTGGTGTCTAAACTGGAATCTGGAGTCCCTGACAGATTCAGTGGCAGTGGATCAGGGACAGATTTTACACTGAAAATCAGCAGAGTGGAGGCTGAGGATTTGGGAGTTTATTACTGCGTGCAAGGTACACATTTCCCTCA |
| IGKV6-23*01_S4960_NOD | NOD/ShiLtJ | Musmus IGKV6-23*01 F | 92.11 | 287 | GACATTGTGATGACCCAGTCTCAAAAATTCATGTCCACATCAGTAGGAGAGAGGGTCAGCATCACCTGCAAGGCCAGTCAGAATGTAGGTACTAATGTAGCCTGGTATCAGCAGAAAGCAGGGCAGTCTCTTGAACTGCTGATCTATGGGGCATCCAACCGGCACACTGGAGTCCCTGATCGCTTCACAGGCAGTGGATCTGGGACAGATTTCACTCTCACCATCACCAATGTGCAGTCTGAAGACATGACAAATTATTTCTGTGAGCAATATAGCAGCTATCCTCT |
| IGKV4-81*01_S2422_NOD | NOD/ShiLtJ | Musmus IGKV4-81*01 F | 96.45 | 292 | GAAAATGTGCTGACCCAGTCTCCAGCAATCATGGCTGCATCTCCAGGGGAGAAGGTCACCATGACCTGCAGTGCCAGCTCAAGTGTAAGTTCTGGTAACTTTCACTGGTACCAGCAGAAGCCAGGCACTTCTCCCAAACTCTGGATTTATAGGACATCCAACCTGGCTTCTGGAGTCCCCGCTCGCTTCAGTGGCAGTGGGTCTGGGACCTCTTACTCTCTTACAATCAGCAGCATGGAGGCCGAAGATGCTGCCACTTATTACTGCCAGCAGTGGAGTGGTTACCCACCCA |
| IGKV4-74*01_S3912_NOD | NOD/ShiLtJ | Musmus IGKV4-74*01 F | 95.04 | 292 | CAAATTGTTCTCACCCAGTCTCCAGCAATCATGTCTGCCTCTCCAGGGGAGAAGGTCACCATGACCTGCAGTGCCAGCTCAAGTGTAAGTTCCAGGTACTTGCACTGGTACCAGCAGAAGTCAGGAGCCTCCCCCAAACTCTGGATTTATGGCACATCCAACCTGGCTTCTGGAGTCCCTGCTCGCTTCAGTGGCAGTGGGTCTGGGACCTCTTACTCTCTCACAATCAGCAGCGTGGAGGCTGAAGATGCTGCCACTTATTACTGCCAGCAGTATCATAGTGACCCACCCA |

| | | | | | |
|---|---|---|---|---|---|
| IGKV10-96*03_NOD | NOD/ShiLtJ | Musmus IGKV10-96*03 F | 100 | 287 | GATATCCAGATGACACAGACTACATCCTCCCTGTCTGCCTCTCTGGGAGACAGAGTCACCATCAGTTGCAGGGCAAGTCAGGACATTAGCAATTATTTAAACTGGTATCAGCAGAAACCAGATGGAACTGTTAAACTCCTGATCTACTACACATCAAGATTACACTCAGGAGTCCCATCAAGGTTCAGTGGCAGTGGGTCTGGGACAGATTATTCTCTCACTATTAGCAACCTGGAACAAGAAGATATTGCCACTTACTTTTGCCAACAGGATAGTAAGCATCCTCC |
| IGKV8-16*01_S4105_NOD | NOD/ShiLtJ | Musmus IGKV8-16*01 F | 98.99 | 304 | GAAATTGTGTTGACACAGTCTATACCATCCCTGACTGTGTCAGCAGGAGAGAGGGTCACTATCAGCTGCAAATCCAATCAGAATCTTTTATGGAGTGGAAACCAACGGTACTGTTTGGTCTGGCACCAGTGGAAACCAGGGCAAACTCCTACACCGTTGATCACCTGGACATCTGATAGGTACTCTGGAGTCCCTGATCGTTTCATAGGCAGTGGATCTGTGACAGATTTCACTCTGACCATCAGCAGTGTGCAGGATGAAGATGTGGCAGTTTATTTCTGTCAGCAGCATTTACACATTCCTC |
| IGKV6-23*01_S5820_NOD | NOD/ShiLtJ | Musmus IGKV6-23*01 F | 92.11 | 287 | GACATTGTGATGACCCAGTCTCAAAAATTCATGTCCACATCAGTAGGAGAGAGGGTCAGCATCACCTGCAAGGCCAGTCAGAATGTAGGTACTAATGTAGCCTGGTATCAGCAGAAAGCAGGGCAGTCTCTTGAACTGCTGATCTATGGGGCATCCAACCGGCACACTGGAGTCCCTGATCGCTTCACAGGCAGTGGATCTGGGACAGATTTCACTCTCACCATCACCAATGTGCAGTCTGAAGACATGACAAATTATTTCTGTGAGCAATATAGCAGCTATCCGCT |
| IGKV4-78*01_S3978_NOD | NOD/ShiLtJ | Musmus IGKV4-78*01 F | 97.87 | 292 | CAAATTGTTCTCACCCAGTCTCCAGCAATCATGTCTGCATCTCCTGGGGAACGGGTCACCATGACCTGCAGTGCCAGCTCAAGTGTAAGTTCCAGCTACTTGTACTGGTACCAGCAGAAGCCAGGATCCTCCCCCAAACTATGGATTTATAGCACATCCAACCTGGCTTCTGGAGTCCCTGCTCGCTTCAGTGGCAGTGGGTCTGGGACCTCTTATTCTCTCACAATCAGCAGCATGGAGGCTGAAGATGCTGCCACTTATTACTGCCAGCAGTACAGTGGTTACCCATCCA |
| IGKV8-28*02_NOD | NOD/ShiLtJ | Musmus IGKV8-28*02 F | 100 | 302 | GACATTGTGATGACACAGTCTCCATCCTCCCTGAGTGTGTCAGCAGGAGATAAGGTCACTATGAGCTGCAAGTCCAGTCAGAGTCTGTTAAACAGTAGAAACCAAAAGAACTACTTGGCCTGGTACCAGCAGAAACCATGGCAGCCTCCTAAACTGCTGATCTACGGGGCATCCACTAGGGAATCTGGGGTCCCTGATCGCTTCACAGGCAGTGGATCTGGAACAGATTTCACTCTCACCATCAGCAGTGTGCAGGCTGAAGACCTGGCAGTTTATTACTGTCAGAATGATTATAGTTATCC |
| IGKV3-4*01_S4235_NOD | NOD/ShiLtJ | Musmus IGKV3-4*01 F | 92.44 | 298 | GACATTGTGCTGACCCAATCTCCAGCTTCTTTGGCTGTGTCTCTAGGGCAGAGGGCCACCATATCCTGCAAGGCCAGCCAGAGTATCAATTATTACGGTGATAATTATATGCACTGGTTCCAACAGAAACCAGGACAGCCAACCAAACTCCTTATCTATGATGCATCCAACCTAGAATCTGGAGTTCCTGACAGGTTCAGTGGCAGTGGGTCTGGGACAGACTTCACCCTCAACATCCATCCTGTGGAGGAGGAGGATGCTGCAACCTATTACTGTCAGCAAAGCAAGGAGTTTCCTC |
| IGKV4-57-1*01_S1729_NOD | NOD/ShiLtJ | Musmus IGKV4-57-1*01 F | 92.91 | 292 | GAAATTGTGCTCACCCAGTCTCCAGCACTCATGGCTGCATCTCCAGGGGAGAAGGTCACCATGACCTGCAGTGCCAGCTCAAGTGTAGGTTCCAGTTACTTGCACTGGTACCAGCAGAAGTCAGGAGCCTCCCCCAAACTCTGGATTTACGGCACATCCAACCTGGCTTCTGGAGTCCCTGCTTGCTTCAGTGGCAGTGGGTCTGGGACCTCTTACTCTCTCACAATCAGCAGCGTGGAGGCTGAAGATGATGCAACTTATTACTGCCAGCAGGGGTGGGATTACCCACTCA |

| | | | | | |
|---|---|---|---|---|---|
| IGKV4-57-1*01_S8603_NOD | NOD/ShiLtJ | Musmus IGKV4-74*01 F | 95.04 | 292 | CAAATTGTTCTCACCCAGTCTCCAGCAATCATGTCTGCCTCTCCAGGGGAGAAGGTCACCATGACCTGCAGTGCCAGCTCAAGTGTAAGTTCCAGGTACTTGCACTGGTACCAGCAGAAGTCAGGAGCCTCCCCCAAACTCTGGATTTATGGCACATCCAACCTGGCTTCTGGAGTCCCTGCTCGCTTCAGTGGCAGTGGGTCTGGGACCTCTTACTCTCTCACAATCAGCAGCGTGGAGGCTGAAGATGCTGCCACTTATTACTGCCAGCAGTATCATAGTGACCCACTCA |
| IGKV17-127*01_S4378_NOD | NOD/ShiLtJ | Musmus IGKV17-127*01 F | 99.64 | 287 | GAAACAACTGTGACCCAGTCTCCAGCATCCCTGTCCGTGGCTACAGGAGAAAAAGTCACTATCAGATGCATAACCAACACTGATATTGATGATGATATGAACTGGTACCAGCAGAAGCCAGGGGAACCTCCTAAGCTCCTTATTTCAGAAGGCAATACTCTTCGTCCTGGAGTCCCATCCCGATTCTCCAGCAGTGGCTATGGCACAGATTTTGTTTTTACAATTGAAAACACGCTCTCAGAAGATGTTGCAGATTACTACTGTTTGCAAAGTGATAACATGCCTCT |
| IGKV9-124*01_S2023_NOD | NOD/ShiLtJ | Musmus IGKV9-124*01 F | 98.92 | 285 | GACATCCAGATGACCCAGTCTCCATCCTCCTTATCTGCCTCTCTGGGAGAAAGAGTCAGTCTCACTTGTCGGGCAAGTCAGGAAATTAGTGGTTACTTAAGCTGGCTTCAGCAGAAACCAGATGGAACTATTAAACGCCTGATCTACAGCACATCCACTTTAGATTCTGGTGTCCCAAAAAGGTTCAGTGGCAGTAGGTCTGGGTCAGATTATTCTCTCACCATCAGCAGCCTTGAGTCTGAAGATTTTGCAGACTATTACTGTCTACAATATGCTAGTTCTCCT |
| IGKV1-122*01_S2138_NOD | NOD/ShiLtJ | Musmus IGKV1-122*01 F | 98.64 | 299 | GATGCTGTGATGACCCAAACTCCACTCTCCCTGCCTGTCAGTCTTGGAGATCAAGCCTCCATCTCTTGCAGGTCTAGTCAGAGCCTTGAAAACAGTAATGGAAACACCTATTTGAACTGGTACCTCCAGAAACCAGGCCAGTCTCCACAGCTCCTGATCTACAGGGTTTCCAACCGATTTTCTGGGGTCCTAGACAGGTTCAGTGGCAGTGGTTCAGGGACAGATTTCACACTCAAGATCAGCAGAGTGGAGGCTGAGGATTTGGGAGTTTATTTCTGCCTCCAAGTTACACATGTCCC |
| IGKV1-99*01_S3206_NOD | NOD/ShiLtJ | Musmus IGKV1-99*01 F | 98.3 | 302 | GATGTTGTTCTGACCCAAACTCCACTCTCTCTGCCTGTCAATATTGGAGATCAAGCCTCTATCTCTTGCAAGTCTACTAAGAGCCTTCTGAATAGTGATGGATTCACTTATTTGGGCTGGTACCTGCAGAAGCCAGGCCAGTCTCCACAGCTCCTAATATATTTGGTTTCTAATCGATTTTCTGGAGTTCCAGACAGGTTCAGTGGTAGTGGGTCAGGGACAGATTTCACCCTCAAGATCAGCAGAGTGGAGGCTGAGGATTTGGGAGTTTATTATTGCTTCCAGAGTAACTATCTTCCTCT |
| IGKV1-88*01_S0716_NOD | NOD/ShiLtJ | Musmus IGKV1-88*01 F | 99.66 | 302 | GATGTTGTGGTGACTCAAACTCCACTCTCCCTGCCTGTCAGCTTTGGAGATCAAGTTTCTATCTCTTGCAGGTCTAGTCAGAGTCTTGCAAACAGTTATGGGAACACCTATTTGTCTTGGTACCTGCACAAGCCTGGCCAGTCTCCACAGCTCCTCATCTATGGGATTTCCAACAGATTTTCCGGGGTGCCAGACAGGTTCAGTGGCAGTGGTTCAGGGACAGATTTCACACTCAAGATCAGCACAATAAAGCCTGAGGACTTGGGAATGTATTACTGCTTACAAGGTACACATCAGCCTCC |
| IGKV16-104*01_S8917_NOD | NOD/ShiLtJ | Musmus IGKV16-104*01 F | 99.64 | 288 | GATGTCCAGATAACCCAGTCTCCATCTTATCTTGCTGCATCTCCTGGAGAAACCATTACTATTAATTGCAGGGCAAGTAAGAGCATTAGCAAATATTTAGCCTGGTATCAAGAGAAACCTGGGAAAACTAATAAGCTTCTTATCTACTCTGGATCCACTTTGCAATCTGGAATTCCATCAAGGTTCAGTGGCAGTGGATCTGGTACAGATTTCACTCTCACCATCAGTAGCCTGGAGCCTGAAGATTTTGCAATGTATTACTGTCAACAACATAATGAATCCCCGTAC |
| IGKV16-104*01_S6011_NOD | NOD/ShiLtJ | Musmus IGKV16-104*01 F | 100 | 288 | GATGTCCAGATAACCCAGTCTCCATCTTATCTTGCTGCATCTCCTGGAGAAACCATTACTATTAATTGCAGGGCAAGTAAGAGCATTAGCAAATATTTAGCCTGGTATCAAGAGAAACCTGGGAAAACTAATAAGCTTCTTATCTACTCTGGATCCACTTTGCAATCTGGAATTCCATCAAGGTTCAGTGGCAGTGGATCTGGTACAGATTTCACTCTCACCATCAGTAGCCTGGAGCCTGAAGATTTTGCAATGTATTACTGTCAACAGCATAATGAATCCCCGTAC |

| | | | | | |
|---|---|---|---|---|---|
| IGKV4-74*01_S3301_NOD | NOD/ShiLtJ | Musmus IGKV4-74*01 F | 94.68 | 292 | CAGATTGTTCTCACCCAGTCTCCAGCAATCATGTCTGCATCTCCAGGGGAGAAGGTCACCATGACCTGCAGGGCCAGCTCAAGTGTAAGTTCCAGTTACTTGCACTGGTACCAGCAGAAGCCAGGATCTTCCCCCAAACTCTGGATTTATAGCACATCCAACCTGGCTTCAGGAGTCCCAGCTCGCTTCAGTGGCAGTGGGTCTGGGACCTCTTACTCTCTCACAATCAGCAGTGTGGAGGCTGAGGATGCTGCCACTTATTACTGCCAGCAGTATGATAGTTCCCCACCCA |
| IGKV4-78*01_S4828_NOD | NOD/ShiLtJ | Musmus IGKV4-78*01 F | 97.52 | 292 | CAAATTGTTCTCACCCAGTCTCCAGCAATCATGTCTGCATCTCCTGGGGAACGGGTCACCATGACCTGCAGTGCCAGCTCAAGTGTAAGTTCCAGCTACTTGTACTGGTACCAGCAGAAGCCAGGATCCTCCCCCAAACTATGGATTTATAGCACATCCAACCTGGCTTCTGGAGTCCCTGCTCGCTTCAGTGGCAGTGGGTCTGGGACCTCTTATTCTCTCACAATCAGCAGCATGGAGGCTGAAGATGCTGCCACTTATTACTGCCAGCAGTACAATGGTTACCCATCCA |
| IGKV8-27*01_NOD | NOD/ShiLtJ | Musmus IGKV8-27*01 F | 100 | 300 | AACATTATGATGACACAGTCGCCATCATCTCTGGCTGTGTCTGCAGGAGAAAAGGTCACTATGAGCTGTAAGTCCAGTCAAAGTGTTTTATACAGTTCAAATCAGAAGAACTACTTGGCCTGGTACCAGCAGAAACCAGGGCAGTCTCCTAAACTGCTGATCTACTGGGCATCCACTAGGGAATCTGGTGTCCCTGATCGCTTCACAGGCAGTGGATCTGGGACAGATTTTACTCTTACCATCAGCAGTGTACAAGCTGAAGACCTGGCAGTTTATTACTGTCATCAATACCTCTCCTCG |
| IGKV6-20*01_S2082_NOR | NOR/LtJ | Musmus IGKV6-20*01 F | 99.64 | 287 | AACATTGTAATGACCCAATCTCCCAAATCCATGTCCATGTCAGTAGGAGAGAGGGTCACCTTGAGCTGCAAGGCCAGTGAGAATGTGGGTACTTATGTATCCTGGTATCAACAGAAACCAGAGCAGTCTCCTAAACTGCTGATATACGGGGCATCCAACCGGTACACTGGGGTCCCCGATCGCTTCACAGGCAGTGGATCTGCAACAGATTTCACTCTGACCATCAGCAGTGTGCAGGCTGAAGACCTTGCAGATTATTACTGTGGACAGAGTTACAGCTATCCTCC |
| IGKV6-15*01_S9105_NOR | NOR/LtJ | Musmus IGKV6-15*01 F | 97.13 | 287 | GACATTGTGATGACCCAGTCTCAAAAATTCATGTCCACATCAGTAGGAGACAGGGTCAGCATCAGCTGCAAGGCCAGTCAGAATGTGGGTAATATTATAGCCTGGTATCAACAGAAACCAGGGCAATCTCCTAAAGCACTGATTTACTTGGCATCCTACCGGTACAGTGGAGTCCCTGATCGCTTCACAGGCAGTGGATCTGGGACAGATTTCACTCTCACCATTAGCAATGTGCAGTCTGAAGACTTGGCAGAGTATTTCTGTCAGCAATATAGCAGCTCTCCTCT |
| IGKV12-46*01_S4541_NOR | NOR/LtJ | Musmus IGKV12-46*01 F | 97.13 | 287 | GACATCCAGATGACTCAGTCTCCAGCCTCCCTATCTGCATCTGTGGGAGAAACTGTCACCATCACATGTCGAGCAAGTGAGAATATTTACAGTAATTTAGCATGGTATCAGCAGAAACAGGGAAAATCTCCTCAGCTCCTGGTCTATGCTGCAACAAATTTAGCAGATGGTGTGCCATCAAGGTTCAGTGGCAGTGGATCAGGCACACAGTTTTCTCTGAAGATCAACAGCCTGCAGCCTGAAGATTTTGGGAGTTATTACTGTCAACATTTTTATGGTACTCCTCC |
| IGKV12-46*01_S1254_NOR | NOR/LtJ | Musmus IGKV12-46*01 F | 94.62 | 287 | GACATCCAGATGACTCAGTCTCCAGCTTCCCTGTCTGCATCTGTGGGAGAAACTGTCACCATCACATGTCGAGCAAGTGAGAATATTGACAGTTATTTAGCATGGTATCAGCAGAAACAGGGAAAATCTCCTCAGCTCCTGGTCTATGCTGCAACACTCTTAGCAGATGGTGTGCCATCAAGGTTCAGTGGCAGTGGATCAGGCACACAGTATTCTCTCAAGATCAACAGCCTGCAGTCTGAAGATGTTGCGAGATATTACTGTCAACATTATTATAGTACTCCTCC |
| IGKV6-b*01_NOR | NOR/LtJ | Musmus IGKV6-b*01 F | 100 | 287 | AGTATTGTGATGACCCAGACTCCCAAATTCCTGCCTGTATCAGCAGGAGACAGGGTTACCATGACCTGCAAGGCCAGTCAGAGTGTGGGTAATAATGTAGCCTGGTACCAACAGAAGCCAGGACAGTCTCCTAAACTGCTGATATACTATGCATCCAATCGCTACACTGGAGTCCCTGATCGCTTCACTGGCAGTGGATCTGGGACAGATTTCACTTTCACCATCAGCAGTGTGCAGGTTGAAGACCTGGCAGTTTATTTCTGTCAGCAGCATTATAGCTCTCCTCC |

| | | | | | |
|---|---|---|---|---|---|
| IGKV1-117*01_S1348_NOR | NOR/LtJ | Musmus IGKV1-117*01 F | 99.32 | 301 | GATGTTGTGATGACCCAAACTCCACTCTCCCTGCCTGTCAGTCTTGGAGATCAAGCCTCCATCTCTTGCAGATCTAGTCAGAGCATTGTACATAGTAATGGAAACACCTATTTAGAATGGTACCTGCAGAAGCCAGGCCAGTCTCCAAAGCTCCTGATCTACAAAGTTTCCAACCGATTTTCTGGGGTCCCAGACAGGTTCAGTGGCAGTGGATCAGGGACAGATTTCACACTCAAGATCAGCAGAGTGGAGGCTGAGGATCTGGGAGTTTATTACTGCTTTCAAGGTTCACATGTTCCTC |
| IGKV6-15*01_S9094_NOR | NOR/LtJ | Musmus IGKV6-15*01 F | 96.06 | 287 | GACATTGTGATGACCCAGTCTCAAAAATTCATGTCCACATCAGTAGGAGACAGGGTCAGCGTCACCTGCAAGGCCAGTCAGTATGTGGGTACTTATGTAGCCTGGTATCAACAGAAACCAGGACAATCTCCTAAAGCACTGATTTACTCGGCATCCACCCGGCACACTGGAGTCCCTGATCGCTTCACAGGCAGTGGATCTGGGACAGATTTCACTCTCACCATTAGCAATGTGCAGTCTGAAGACTTGGCAGAGTATTTCTGTGAGCAATACAGCAGCTCTCCTCT |
| IGKV1-135*01_S3001_NOR | NOR/LtJ | Musmus IGKV1-135*01 F | 95.92 | 302 | GATGTTGTGATGACTCAGACCCCACTCACTTTGTCGGTTACCATTGGACAACCAGCCTCCATCTCTTGCAAATCAAGTCAGAGCCTCTTACATAGTAATGGAAAGACATATTTGAATTGGTTATTACAGAGGCCAGGCCAGTCTCCAAAGCTCCTAATCTATCTGGTGTCTAAACTGGAATCTGGAGTCCCTGACAGGTTCAGTGGCAGTGGATCAGGGACAGATTTCACACTGAAAATCAGCAGAGTGGAGGCTGAGGATTTGGGAGTTTATTACTGCTTGCAAGCTACACATTTTCCTCA |
| IGKV10-94*03_NOR | NOR/LtJ | Musmus IGKV10-94*03 F, or Musmus IGKV10-94*08 F | 100 | 287 | GATATCCAGATGACACAGACTACATCCTCCCTGTCTGCCTCTCTGGGAGACAGAGTCACCATCAGTTGCAGGGCAAGTCAGGATATTAGCAATTATTTAAACTGGTATCAGCAGAAACCAGATGGAACTGTTAAACTCCTGATCTACTACACATCAAGATTACACTCAGGAGTCCCATCAAGGTTCAGTGGCAGTGGGTCTGGGACAGATTATTCTCTCACCATCAGCAACCTGGAACCTGAAGATATTGCCACTTACTATTGTCAGCAGTATAGTAAGCTTCCTCC |
| IGKV12-44*01_NOR | NOR/LtJ | Musmus IGKV12-44*01 F | 100 | 287 | GACATCCAGATGACTCAGTCTCCAGCCTCCCTATCTGCATCTGTGGGAGAAACTGTCACCATCACATGTCGAGCAAGTGAGAATATTTACAGTTATTTAGCATGGTATCAGCAGAAACAGGGAAAATCTCCTCAGCTCCTGGTCTATAATGCAAAAACCTTAGCAGAAGGTGTGCCATCAAGGTTCAGTGGCAGTGGATCAGGCACACAGTTTTCTCTGAAGATCAACAGCCTGCAGCCTGAAGATTTTGGGAGTTATTACTGTCAACATCATTATGGTACTCCTCC |
| IGKV5-45*01_S0430_NOR | NOR/LtJ | Musmus IGKV5-45*01 F | 99.64 | 287 | GATATTGTGCTAACTCAGTCTCCAGCCACCCTGTCTGTGACTCCAGGAGATAGAGTCAGTCTTTCCTGCAGGGCCAGCCAAAGTATTAGCAACTACCTACACTGGTATCAACAAAAATCACATGAGTCTCCAAGGCTTCTCATCAAGTATGCTTCCCAGTCCATCTCTGGGATCCCCTCCAGGTTCAGTGGCAGTGGATCAGGGACAGATTTCACTCTCAGTATCAACAGTGTGGAGACTGAAGATTTTGGAATGTATTTCTGTCAACAGAGTAACAGCTGGCCTCA |
| IGKV2-137*01_S0442_NOR | NOR/LtJ | Musmus IGKV2-137*01 F | 93.54 | 302 | GATATTGTGATGACTCAGGCTGCACCCTCTGTACCTGTCACTCCTGGAGAGTCAGTATCCATCTCCTGCAGGTCTAGTACGAGTCTCCTGCACAGTAGTGGCAAGCATAGGTTGTATTGGTTCCTACAGAGGCCAGGCCAGTCTCCTCAGCTCCTGATATATTATATGTCCAACCTTGCCTCAGGAGTCCCAGACAGGTTCAGTGGCAGTGGGTCAGGAACTGATTTCACACTGAGAATCAGTAGAGTGGAGGCTGAGGATTTTGGTGTTTATTATTGTATGCAAAGTCTAGAATATCCTTT |
| IGKV5-43*01_S9511_NOR | NOR/LtJ | Musmus IGKV5-43*01 F | 99.64 | 287 | GATATTGTGCTAACTCAGTCTCCAGCCACCCTGTCTGTGACTCCAGGAGATAGCGTCAGTCTTTCCTGCAGGGCCAGCCAAAGTATTAGCAACAACCTACACTGGTATCAACAAAAATCACATGAGTCTCCAAGGCTTCTCATCAAGTATGCTTCCCAGTCCATCTCTGGGATCCCCTCCAAGTTCAGTGGCAGTGGATCAGGGACAGATTTCACTCTCAGTATCAACAGTGTGGAGACTGAAGATTTTGGAATGTATTTCTGTCAACAGAGTAACAGCTGGCCTCA |

| | | | | | |
|---|---|---|---|---|---|
| IGKV4-74*01_S2000_NOR | NOR/LtJ | Musmus IGKV4-74*01 F | 94.68 | 292 | CAGATTGTTCTCACCCAGTCTCCAGCAATCATGTCTGCATCTCCAGGGGAGAAGGTCACCATGACCTGCAGGGCCAGCTCAAGTGTAAGTTCCAGTTACTTGCACTGGTACCAGCAGAAGCCAGGATCTTCCCCCAAACTCTGGATTTATAGCACATCCAACCTGGCTTCAGGAGTCCCAGCTCGCTTCAGTGGCAGTGGGTCTGGGACCTCTTACTCTCTCACAATCAGCAGTGTGGAGGCTGAGGATGCTGCCACTTATTACTGCCAGCAGTATGATAGTTCCCCATCCA |
| IGKV8-19*01_NOR | NOR/LtJ | Musmus IGKV8-19*01 F | 100 | 305 | GACATTGTGATGACACAGTCTCCATCCTCCCTGACTGTGACAGCAGGAGAGAAGGTCACTATGAGCTGCAAGTCCAGTCAGAGTCTGTTAAACAGTGGAAATCAAAAGAACTACTTGACCTGGTACCAGCAGAAACCAGGGCAGCCTCCTAAACTGTTGATCTACTGGGCATCCACTAGGGAATCTGGGGTCCCTGATCGCTTCACAGGCAGTGGATCTGGAACAGATTTCACTCTCACCATCAGCAGTGTGCAGGCTGAAGACCTGGCAGTTTATTACTGTCAGAATGATTATAGTTATCCTCC |
| IGKV1-135*01_S7405_NOR | NOR/LtJ | Musmus IGKV1-135*01 F | 95.24 | 302 | GATGTTGTGATGACCCAGACTCCACTGTCTTTGTCGGTTACCATTGGACAACCAGCCTCCATCTCTTGCAAGTCAAGTCAGAGCCTCTTACATAGTGATGGAAAGACATATTTGAATTGGTTATTACAGAGGCCAGGCCAGTCTCCAAAGCTCCTAATCTATCTGGTGTCTAAACTGGAATCTGGCATCCCTGACAGGTTCAGTGGCAGTGGATCAGGGACAGATTTCACACTGAAAATCAGCAGAGTGGAGGTTGAGGATTTGGGAGTTTATTACTGCTTGCAACATACACATTTTCCTCA |
| IGKV12-46*01_S4026_NOR | NOR/LtJ | Musmus IGKV12-46*01 F | 99.64 | 287 | GACATCCAGATGACTCAGTCTCCAGCCTCCCTATCTGTATCTGTGGGAGAAACTGTCACCATCACATGTCGAGCAAGTGAGAATATTTACAGTAATTTAGCATGGTATCAGCAGAAACAGGGAAAATCTCCTCAGCTCCTGGTCTATGCTGCAACAAACTTAGCAGATGGTGTTCCATCAAGGTTCAGTGGCAGTGGATCAGGCACACAGTATTCCCTCAAGATCAACAGCCTGCAGTCTGAAGATTTTGGGAGTTATTACTGTCAACATTTTTGGGGTACTCCTCC |
| IGKV6-23*01_S4960_NOR | NOR/LtJ | Musmus IGKV6-23*01 F | 92.11 | 287 | GACATTGTGATGACCCAGTCTCAAAAATTCATGTCCACATCAGTAGGAGAGAGGGTCAGCATCACCTGCAAGGCCAGTCAGAATGTAGGTACTAATGTAGCCTGGTATCAGCAGAAAGCAGGGCAGTCTCTTGAACTGCTGATCTATGGGGCATCCAACCGGCACACTGGAGTCCCTGATCGCTTCACAGGCAGTGGATCTGGGACAGATTTCACTCTCACCATCACCAATGTGCAGTCTGAAGACATGACAAATTATTTCTGTGAGCAATATAGCAGCTATCCTCT |
| IGKV8-21*01_S1419_NOR | NOR/LtJ | Musmus IGKV8-21*01 F | 99.33 | 302 | GACATTGTGATGTCACAGTCTCCATCCTCCCTGGCTGTGTCAGCAGGAGAGAAGGTCACTATGAGCTGCAAATCCAGTCAGAGTCTGTTCAACAGTAGAACCCGAAAGAACTACTTGGCTTGGTACCAGCAGAAACCAGGGCAGTCTCCTAAACTGCTGATCTACTGGGCATCCACTAGGGAATCTGGGGTCCCTGATCGCTTCACAGGCAGTGGATCTGGGACAGATTTCACTCTCACCATCAGCAGTGTGCAGGCTGAAGACCTGGCAGTTTATTACTGCAAGCAATCTTATTATCTTCC |
| IGKV1-110*02_NOR | NOR/LtJ | Musmus IGKV1-110*02 F | 100 | 302 | GATGTTGTGATGACCCAAACTCCACTCTCCCTGCCTGTCAGTCTTGGAGATCAAGCTTCCATCTCTTGCAGATCTAGTCAGAGCCTTGTACACAGCAATGGAAACACCTATTTATATTGGTACCTGCAGAAGCCAGGCCAGTCTCCAAAGCTCCTGATCTACAGGGTTTCCAACCGATTTTCTGGGGTCCCAGACAGGTTCAGTGGCAGTGGATCAGGGACAGATTTCACACTCAAGATCAGCAGAGTGGAGGCTGAGGATCTGGGAGTTTATTTCTGCTTTCAAGGTACACATGTTCCTCA |

| | | | | | |
|---|---|---|---|---|---|
| IGKV14-126*01_NOR | NOR/LtJ | Musmus IGKV14-126*01 F | 100 | 287 | GACATCAAGATGACCCAGTCTCCATCCTCCATGTATGCATCGCTGGGAGAGAGAGTCACTATCACTTGCAAGGCGAGTCAGGACATTAAAAGCTATTTAAGCTGGTACCAGCAGAAACCATGGAAATCTCCTAAGACCCTGATCTATTATGCAACAAGCTTGGCAGATGGGGTCCCATCAAGATTCAGTGGCAGTGGATCTGGGCAAGATTATTCTCTAACCATCAGCAGCCTGGAGTCTGACGATACAGCAACTTATTACTGTCTACAGCATGGTGAGAGCCCTCC |
| IGKV4-56*01_S5696_NOR | NOR/LtJ | Musmus IGKV4-56*01 P | 96.38 | 286 | CAAATTGTTCTCACCCAGTCTCCAGCAATCATGTCTGCATCTCCAGGGCAGAAAGTCACCATAACCTGCAGTGCCAGCTCAAGTGTAAATTACATGCACTGGTACCAGCAGAAGCTAGGATCCTCCCCCAAACTCTGGATTTATGACACATCCAAACTGGCTCCTGGAGTCCCTGCTCGCTTCAGTGGCAGTGGGTCTGGGACCTCTTACTCTCTCACAATCAGCAGCATGGAGGCTGAAGATGCTGCCTCTTATTTCTGCCATCAGTGGAGTAGTTACCCACCCA |
| IGKV1-132*01_S5178_NOR | NOR/LtJ | Musmus IGKV1-132*01 F | 97.96 | 302 | GATGTTGTGATGACCCAGACTCCACTGTCTTTGTCGGTTACCATTGGACAACCAGCCTCTATCTCTTGCAAGTCAAGTCAGAGCCTCTTATATAGTGATGGAAAGACATATTTGAATTGGTTACAACAGAGGCCAGGCCAGTCTCCAAAGCGCCTAATGTATCAGGTGTCCAAACTGGACCCTGGCATCCCTGACAGGTTCAGTGGCAGTGGATCAGAGACAGATTTTACACTTAAAATCAGCAGAGTGGAGGCTGAGGATTTGGGAGTTTATTACTGCTTGCAAGGTACATATTATCCTCA |
| IGKV12-41*01_S6800_NOR | NOR/LtJ | Musmus IGKV12-41*01 F | 97.13 | 287 | GATATCCAGATGACTCAGTCTCCAGCCTCCCTATCTGCATCTGTGGGAGAAACTGTCACCATCACATGTCGAGCAAGTGGGAATATTCACAATTATTTAGCATGGTATCAGCAGAAACAGGGAAAATCTCCTCAGCTCCTGGTCTATAATGCAAAAACCTTAGCGGAAGGTGTGCCATCAAGGTTCAGTGGCAGTGGATCAGGAACACAATATTCTCTCAAGATCAACAGCCTGCAGCCTGAGGATTTTGGGAGTTATTACTGTCATCATTATTATAGTACTCCTCC |
| IGKV1-133*01_S3070_NOR | NOR/LtJ | Musmus IGKV1-133*01 F | 95.92 | 302 | GATGTTGTGATGACCCAGACTCCACTCACTTTGTCTGTTACCATTGGACAGCCAGCTTCCATTTCTTGCAAGTCAAGTCAGAGCCTCTTATATAGTGATGGAAAAACCTATTTGAATTGGTTATTACAGAGTCCAGGCCAGTCTCCAAAGCTCCTAATCTATCTGGTGTCTAAACTGGAATCTGGAGTCCCTGACAGATTCAGTGGCAGTGGATCAGGGACAGATTTTACACTGAAAATCAGCAGAGTGGAGGCTGAGGATTTGGGAGTTTATTACTGCGTGCAAGGTACACATTTCCCTCA |
| IGKV15-103*01_S0817_NOR | NOR/LtJ | Musmus IGKV15-103*01 ORF | 97.85 | 287 | GACATCCAGATGAACCAGTCTCCATCCAGTCTGTCTGCATCCCTCGGAGACACAATTACCATCACTTGCCGTGCCAGTCAGAACATTAATATTTGGTTAAGCTGGTACCAGCAGAAACCAGGAAATATTCCTAAACTATTGATCTATAAGGCTTCCAACTTGCACACAGGCGTCCCATCAAGGTTTAGTGGCAGTGGATCTGGAACAGATTTCACATTAACCATCAGCAGTCTGCAGCCTGAAGACATTGCCACTTACTACTGTCTACAGGGTCAAAGTTATCCTCT |
| IGKV8-34*01_S3615_NOR | NOR/LtJ | Musmus IGKV8-34*01 F | 99.33 | 305 | GACATTTTGATGACTCAGTCTCCATCCTCCCTGACTGTGTCAGCAGGAGAGAAGGTCACTATGAGCTGCAAGTCCAGTCAGAGTCTTTTAGCTAGTGCCAACCAAAATAACTACTTGGCCTGGCACCAGCAGAAACCAGGACGATCTCCTAAAATGCTGATAATTTGGGCATCCACTAGGGTATCTGGAGTCCCTGATCGCTTCATAGGCAGTGGATCTGGGACGGATTTCACTCTGACCATCAACAGTGTGCAGGCTGAAGATCTGGCTGTTTATTACTGTCAGCAGTCCTACAGCGCTCCTAC |
| IGKV4-80*01_S4247_NOR | NOR/LtJ | Musmus IGKV4-80*01 F | 99.64 | 282 | CAAATTGTTCTCACCCAGTCTCCAGCAATCATGTCTGCATCTCTAGGGGAGGAGATCACCCTAACCTGCAGTGCCAGCTCGAGTGTAAGTTACATGCACTGGTTCCAGCAGAAGTCAGGCACTTCTCCCAAACTCTTGATTTATAGCACATCCAACCTGGCTTCTGGAGTCCCTTCTCGCTTCAGTGGCAGTGGGTCTGGGACCTTTTATTCTCTCACAATCAGCAGTGTGGAGGCTGAAGATGCTGCCGATTATTACTGCCATCAGTGGAGTAGTTATCCA |

| | | | | | |
|---|---|---|---|---|---|
| IGKV17-127*01_S3263_NOR | NOR/LtJ | Musmus IGKV17-127*01 F | 99.64 | 287 | GAAACAACTGTGACCCAGTCTCCAGCATCCCTGTCCGTGGCTACAGGAGAAAAAGTCACTATCAGATGCATAACCAACACTGATATTGATGATGATATGAACTGGTACCAGCAGAAGCCAGGGGAACCTCCTAAGCTCCTTATTTCAGAAGGCAATACTCTTCGTCCTGGAGTCCCATCCCGATTCTCCAGCAGTGGCTATGGCACAGATTTTGTTTTTACAATTGAAAACACGCTCTCAGAAGATGTTGCAGATTACTACTGTTTGCAAAGTGATAACATGCCGCT |
| IGKV1-122*01_S7970_NOR | NOR/LtJ | Musmus IGKV1-122*01 F | 98.64 | 302 | GATGCTGTGATGACCCAAACTCCACTCTCCCTGCCTGTCAGTCTTGGAGATCAAGCCTCCATCTCTTGCAGGTCTAGTCAGAGCCTTGAAAACAGTAATGGAAACACCTATTTGAACTGGTACCTCCAGAAACCAGGCCAGTCTCCACAGCTCCTGATCTACAGGGTTTCCAACCGATTTTCTGGGGTCCTAGACAGGTTCAGTGGCAGTGGTTCAGGGACAGATTTCACACTCAAGATCAGCAGAGTGGAGGCTGAGGATTTGGGAGTTTATTTCTGCCTCCAAGTTACACATGTCCCTCC |
| IGKV3-7*01_S6705_NOR | NOR/LtJ | Musmus IGKV3-7*02 F | 97.59 | 299 | GACATTGTGCTGACACAGTCTCCTGCTTCCTTAGCTGTATCTCTGGGGCAGAGGGCCACCATCTCCTGCAGGGCCAGCAAAAGTGTCAGTACATCTAGCTATAGTTACATGCACTGGTACCAACAGAAACCAGGACAGCCACCCAAACTCCTCATCAAGTATGCATCCTACCTAGAATCTGGGGTTCCTGCCAGGTTCAGTGGCAGTGGGTCTGGGACAGACTTCACCCTCAACATCCATCCTGTGGAGGAGGAGGATGCTGCAACATATTACTGTCAGCACAGTAGGGAGTTTCCTCC |
| IGKV3-4*01_S3473_NOR | NOR/LtJ | Musmus IGKV3-4*01 F | 92.44 | 299 | GACATTGTGCTGACCCAATCTCCAGCTTCTTTGGCTGTGTCTCTAGGGCAGAGGGCCACCATATCCTGCAAGGCCAGCCAGAGTATCAATTATTACGGTGATAATTATATGCACTGGTTCCAACAGAAACCAGGACAGCCAACCAAACTCCTTATCTATGATGCATCCAACCTAGAATCTGGAGTTCCTGACAGGTTCAGTGGCAGTGGGTCTGGGACAGACTTCACCCTCAACATCCATCCTGTGGAGGAGGAGGATGCTGCAACCTATTACTGTCAGCAAAGCAAGGAGTTTCCTCC |
| IGKV3-1*01_S7345_NOR | NOR/LtJ | Musmus IGKV3-1*01 F | 98.63 | 299 | GACATTGTGCTCACCCAATCTCCAGCTTCTTTGGCTGTGTCTCTAGGGCAGAGTGTCACCATCTCCTGCAGAGCCAGTGAAAGTGTTGAATATTATGGCACTAGTTTAATGCAGTGGTACCAACAGAAACCAGGACAGCCACCCAAACTCCTCATCTATGGTGCATCCAACGTAGAATCTGGGGTCCCTGCCAGGTTTAGTGGCAGTGGGTCTGGGACAGACTTCAGCCTCAACATCCATCCTGTGGAGGAGGATGATATTGCAATGTATTTCTGTCAGCAAAGTAGGAAGGTTCCTTC |
| IGKV3-4*01_S8857_NOR | NOR/LtJ | Musmus IGKV3-4*01 F | 97.25 | 299 | GACATTGTGCTGACCCAATCTCCAGCTTCTTTGGCTGTGTCTCTAGGGCAGAGGGCCACCATCTCCTGCAGAGCCAGCGAAAGTGTTGATTATGATGGCGATAGTTATATGAACTGGTACCAACAGAAACCAGGACAGCCACCGAAACTCCTCATCTATGCTGCATCCAATCTAGAATCTGGGATCCCTGCCAGGTTTAGTGGCAGTGGGTCTGGGACAGACTTCACCCTCAACATTCATCCTGTGGAGGAGGAGGATGCTGCAACCTATTACTGTCTGCAAAGTAATGAGGCTCCTCC |
| IGKV16-104*01_S9985_NOR | NOR/LtJ | Musmus IGKV14-87-2*01 F | 99.64 | 287 | GACATCCAAATGACCCAGTCCTCATCATTTCTCTCTGCATCTCTAGGAGATCATCTTACAATCAACTGCAGGGCCAGTAAGGATATTAACAAGTATTTTGCTTGGGTTCAACAGAAGCCAGGGAAGGCTCCAAGGATGTTGATTCATTTTGCTTCCACCTTGCTACCTGGGGTTCCAGAAAAGTTCAGTGGGAGTGGATCTGGGACAGATTTTTCTCTCACTATCAGAAACATAGAGTCTGAAGATATTGCAATGTATTACTGTCTACAGTATTCTGAGCATCCGTA |

| | | | | | |
|---|---|---|---|---|---|
| IGKV17-127*01_S4378_NOR | NOR/LtJ | Musmus IGKV17-127*01 F | 99.64 | 287 | GAAACAACTGTGACCCAGTCTCCAGCATCCCTGTCCGTGGCTACAGGAGAAAAAGTCACTATCAGATGCATAACCAACACTGATATTGATGATGATATGAACTGGTACCAGCAGAAGCCAGGGGAACCTCCTAAGCTCCTTATTTCAGAAGGCAATACTCTTCGTCCTGGAGTCCCATCCCGATTCTCCAGCAGTGGCTATGGCACAGATTTTGTTTTTACAATTGAAAACACGCTCTCAGAAGATGTTGCAGATTACTACTGTTTGCAAAGTGATAACATGCCTCT |
| IGKV9-124*01_S2023_NOR | NOR/LtJ | Musmus IGKV9-124*01 F | 98.92 | 285 | GACATCCAGATGACCCAGTCTCCATCCTCCTTATCTGCCTCTCTGGGAGAAAGAGTCAGTCTCACTTGTCGGGCAAGTCAGGAAATTAGTGGTTACTTAAGCTGGCTTCAGCAGAAACCAGATGGAACTATTAAACGCCTGATCTACAGCACATCCACTTTAGATTCTGGTGTCCCAAAAAGGTTCAGTGGCAGTAGGTCTGGGTCAGATTATTCTCTCACCATCAGCAGCCTTGAGTCTGAAGATTTTGCAGACTATTACTGTCTACAATATGCTAGTTCTCCT |
| IGKV8-24*01_S8843_NOR | NOR/LtJ | Musmus IGKV8-24*01 F | 98.32 | 305 | GACATTGTGATGACACAGTCTCCATCCTCCCTGGCTATGTCAGTAGGACAGAAGGTCACTATGAGCTGCAAGTCCAGTCAGAGCCTTTTAAGTAGTAGCAATCAAAAGAACTATTTGGCCTGGTACCAGCAGAAACCAGGACAGTCTCCTAAACTTCTGGTATACTTTGCATCCACTAGGGAATCTGGGGTCCCTGATCGCTTCATAGGCAGTGGATCTGGGACAGATTTCACTCTTACCATCAGCAGTGTGCAGGCTGAAGACCTGGCACTTTATTACTGTCAGCAACATTATAGCACTCCTCC |
| IGKV1-133*01_S5139_NOR | NOR/LtJ | Musmus IGKV1-133*01 F | 99.32 | 302 | GATGTTGTGATGACCCAGACTCCACTCACTTTGTCAGTTACCATTGGACAACCAGCCTCTATCTCTTGCAAGTCAAGTCAGAGCCTCTTATATAGTAATGGAAAAACCTATTTGAACTGGTTATTACAGAGGCCAGGCCAGTCTCCAAAGCGCCTAATCTATCTGGTGTCTAAACTGGACTCTGGAGTCCCTGACAGGTTCACTGGCAGTGGATCAGGAACAGATTTTACACTGAAAATCAGCAGAGTGGAGGCTGAGGATTTGGGAGTTTATTACTGCGTGCAAGGTACACATTTTCCTCA |
| IGKV3-10*01_S5437_NOR | NOR/LtJ | Musmus IGKV3-10*01 F | 98.97 | 299 | AAAATTGTGCTGACCCAATCTCCAGCTTCTTTGGCTGTGTCTCTAAGGCAGAGGGCCACCATATCCTGCAGAGCCAGTGAAAGTGTTGATAGTTATGGCAATAGTTTTATGCACTGGTACCAGCAGAAACCAGGACAGCCACCCAAACTCCTCATCTATCGTGCATCCAACCTAGAATCTGGGGTCCCTGCCAGGTTCAGTGGCAGTGGGTCTAGGACAGACTTCACCCTCACCATTGATCCTGTGGAGGCTGATGATGCTGCAACCTATTACTGTCAGCAAAATAATGAGGATCCTCC |
| IGKV19-93*01_S8964_NOR | NOR/LtJ | Musmus IGKV19-93*01 F | 98.21 | 287 | GACATCCAGATGACACAGTCTCCATCCTCACTGTCTGCATCTCTGGGAGGCAAAGTCACCATCACTTGCAAGGCAAGCCAAGACATTAACAAGAATATAGCTTGGTACCAACACAAGCCTGGAAAAGGTCCTAGGCTGCTCATATGGTACACATCTACATTACAGCCAGGCATCCCATCAAGGTTCAGTGGAAGTGGGTCTGGGAGAGATTATTCCTTCAGCATCAGCAACCTGGAGCCTGAAGATATTGCAACTTATTACTGTCTACAGTATGATAATCTTCCACC |
| IGKV8-16*01_S7495_NOR | NOR/LtJ | Musmus IGKV8-16*01 F | 99.33 | 305 | GAAATTGTGTTGACGCAGTCTATACCATCCCTGACTGTGTCAGCAGGAGAGAGGGTCACTATCAGCTGCAAATCCAATCAGAATCTTTTATGGAGTGGAAACCAAAGGTACTGTTTGGTCTGGCACCAGTGGAAACCAGGGCAAACTCCTACACCGTTGATCACCTGGACATCTGATAGATACTCTGGAGTCCCTGATCGTTTCATAGGCAGTGGATCTGTGACAGATTTCACTCTGACCATCAGCAGTGTGCAGGCTGAAGATGTGGCAGTTTATTTCTGTCAGCAGCATTTACACATTCCTCC |
| IGKV4-86*01_S4316_NOR | NOR/LtJ | Musmus IGKV4-86*01 F | 100 | 283 | GAAATTGTGCTCACTCAGTCTCCAGCCATCACAGCTGCATCTCTGGGGCAAAAGGTCACCATCACCTGCAGTGCCAGCTCAAGTGTAAGTTACATGCACTGGTACCAGCAGAAGTCAGGCACCTCCCCCAAACCATGGATTTATGAAATATCCAAACTGGCTTCTGGAGTCCCAGCTCGCTTCAGTGGCAGTGGGTCTGGGACCTCTTACTCTCTCACAATCAGCAGCATGGAGGCTGAAGATGCTGCCATTTATTACTGCCAGCAGTGGAATTATCCACTTA |

| | | | | | |
|---|---|---|---|---|---|
| IGKV4-53*01_S3575_NOR | NOR/LtJ | Musmus IGKV4-53*01 F | 98.94 | 292 | GAAATTGTGCTCACCCAGTCTCCAGCACTCATGGCTGCATCTCCAGGGGAGAAGGTCACCATCACCTGCAGTGTCAGCTCAAGTATAAGTTCCAGCAACTTACACTGGTACCAGCAGAAGTCAGGAACCTCCCCCAAACCCTGGATTTATGGCACATCCAACCTTGCTTCTGGAGTCCCTGTTCGCTTCAGTGGCAGTGGATCTGGGACCTCTTATTCTCTCACAATCAGCAGCATGGAGGCTGAAGATGCTGCCACTTATTACTGTCAACAGTGGAGTAGTTACCCACTCA |
| IGKV4-74*01_S3912_NOR | NOR/LtJ | Musmus IGKV4-74*01 F | 95.04 | 292 | CAAATTGTTCTCACCCAGTCTCCAGCAATCATGTCTGCCTCTCCAGGGGAGAAGGTCACCATGACCTGCAGTGCCAGCTCAAGTGTAAGTTCCAGGTACTTGCACTGGTACCAGCAGAAGTCAGGAGCCTCCCCCAAACTCTGGATTTATGGCACATCCAACCTGGCTTCTGGAGTCCCTGCTCGCTTCAGTGGCAGTGGGTCTGGGACCTCTTACTCTCTCACAATCAGCAGCGTGGAGGCTGAAGATGCTGCCACTTATTACTGCCAGCAGTATCATAGTGACCCACCCA |
| IGKV4-57-1*01_S8603_NOR | NOR/LtJ | Musmus IGKV4-74*01 F | 95.04 | 292 | CAAATTGTTCTCACCCAGTCTCCAGCAATCATGTCTGCCTCTCCAGGGGAGAAGGTCACCATGACCTGCAGTGCCAGCTCAAGTGTAAGTTCCAGGTACTTGCACTGGTACCAGCAGAAGTCAGGAGCCTCCCCCAAACTCTGGATTTATGGCACATCCAACCTGGCTTCTGGAGTCCCTGCTCGCTTCAGTGGCAGTGGGTCTGGGACCTCTTACTCTCTCACAATCAGCAGCGTGGAGGCTGAAGATGCTGCCACTTATTACTGCCAGCAGTATCATAGTGACCCACTCA |
| IGKV4-50*01_S2188_NOR | NOR/LtJ | Musmus IGKV4-50*01 F | 97.1 | 286 | GAAAATGTGCTCACCCAGTCTCCAGCAATCATGTCTGCAACTCTAGGGGAGAAGGTCACCATGAGCTGCAGGGCCAGCTCAAATGTAAAGTACATGTACTGGTACCAGCAGAAGTCAGGTGCCTCCCCCAAACTATGGATTTATTACACATCCAACCTGGCTTCTGGAGTCCCAGCTCGCTTCAGTGGCAGTGGGTCTGGGACCTCTTATTCTCTCACAATCAGCAGCGTGGAGGCTGAAGATGCTGCCACTTATTACTGCCAGCAGTTTACTAGTTCCCCATCCA |
| IGKV2-109*03_NOR | NOR/LtJ | Musmus IGKV2-109*03 F | 100 | 302 | GATATTGTGATGACGCAGGCTGCATTCTCCAATCCAGTCACTCTTGGAACATCAGCTTCCATCTCTTGCAGTTCTAGTAAGAGTCTCCTACATAGTAATGGCATCACTTATTTGTATTGGTATCTGCAGAGGCCAGGCCAGTCTCCTCAGCTCCTGATATATCGGATGTCCAACCTTGCCTCAGGAGTCCCAGACAGGTTCAGTGGCAGTGGGTCAGGAACTGATTTCACACTGAGAATCAGCAGAGTGGAGGCTGAGGATGTGGGTGTTTATTACTGTGCTCAAATGCTAGAACGCCCTCC |
| IGKV4-62*01_S2199_NOR | NOR/LtJ | Musmus IGKV4-62*01 ORF | 96.74 | 286 | CAAATTGTTCTCACCCAGTCTCCAGCAATCATGTCTGCATCTCCAGGGGAGAAGGTCACCATGACCTGCAGTGCCAGCTCAAGTGTAAGTTACATGTACTGGTACCAGCAGAAGCCAGGATCCTCCCCCAGACTCTGGATTTATGACACATCCAACCTGGCTTCTGGAGTCCCCGCTCGCTTCAGTGGCAGTAGGTCTGGGACCTCTTATTCTCTCACAATCAGCAGCATGGAGGCTGAAGATGCTGCCACTTATTACTGCCATCAGCGGAGTAGTTACCCATTCA |
| IGKV10-96*03_NOR | NOR/LtJ | Musmus IGKV10-96*03 F | 100 | 287 | GATATCCAGATGACACAGACTACATCCTCCCTGTCTGCCTCTCTGGGAGACAGAGTCACCATCAGTTGCAGGGCAAGTCAGGACATTAGCAATTATTTAAACTGGTATCAGCAGAAACCAGATGGAACTGTTAAACTCCTGATCTACTACACATCAAGATTACACTCAGGAGTCCCATCAAGGTTCAGTGGCAGTGGGTCTGGGACAGATTATTCTCTCACTATTAGCAACCTGGAACAAGAAGATATTGCCACTTACTTTTGCCAACAGGATAGTAAGCATCCTCC |

| | | | | | |
|---|---|---|---|---|---|
| IGKV2-137*01_S6567_NOR | NOR/LtJ | Musmus IGKV2-137*01 F | 98.64 | 302 | GATATTGTGATGACTCAGGCTGCACCCTCTGTACCTGTCACTCCTGGAGAGTCAGTATCCATCTCCTGCAGGTCTAGTAAGAGTCTCCTGCATAGTAATGGCAACACTTACTTGTATTGGTTCCTACAGAGGCCTGGCCAGTCTCCTCAGCTCCTGATATATCGGATGTCCAACCTTGCCTCAGGAGTCCCAGACAGGTTCAGTGGCAGTGGGTCAGGAACTGCTTTCACACTGAGAATCAGTAGAGTCGAGGCTGAGGATGTGGGTGTTTATTACTGTATGCAACATCTAGAGTATCCTTT |
| IGKV14-100*01_S4794_NOR | NOR/LtJ | Musmus IGKV14-100*01 F | 99.64 | 287 | GACATCCTGATGACCCAATCTCCATCCTCCATGTCTGTATCTCTGGGAGACACAGTCAGCATCACTTGCCATGCAAGTCAGGGCATTAGCAGTAATATAGGGTGGTTGCAGCAGAAACCAGGGAAATCATTTAAGGGCCTGATCTATCATGCAACCAACTTGGAAGATGGAGTTCCATCAAGGTTCAGTGGCAGTGGATCTGGAGCAGATTATTCTCTCACCATCAGCAGCCTGGAATCTGAAGATTTTGCAGACTATTACTGTGTACAGTATGCTCAGTTTCCTCC |
| IGKV9-120*01_S7607_NOR | NOR/LtJ | Musmus IGKV9-120*02 F | 96.42 | 285 | GACATCCAGATGACCCAGTCTCCATCCTCCTTATCTGCCTCTCTGGGAGAAAGAATCAGTCTCACTTGCCGGGCAAGTCAGGACATTTATGGTAGCTTAAACTGGTTTCAGCAGAAACCAGATGGAACTATTAAACTCCTGATCTACGGCACATCCAGTTTAGATTCTGGTGTCCCCAAAAGGTTCAGTGGCAGTAGGTCTGGGTCAGATTATTCTCTCACCATCAGCAGCCTTGAGTCTGAAGATTTTGCAGACTATTACTGTCTACAATATGCTAGTTCTCCT |
| IGKV4-59*01_S9913_NOR | NOR/LtJ | Musmus IGKV4-59*01 F | 98.19 | 282 | CAAATTGTTCTCACCCAGTCTCCAGCAATCATGTCTGCATCTCCAGGGGAGAAGGTCACCATGACATGCAGTGCCAGCTCAAGTGTAAGTTACATGCACTGGTACCAGCAGAAGTCAGGCACCTCCCCCAAAAGATGGATTTATGACACATCCAAACTGGATTCTGGAGTCCCTGCTCGCTTCAGTGGCAGTGGGTCTGGGACCTCTTATTCTCTCACAATCAGCAGCATGGAGGCTGAAGATGCTGCCACTTATTACTGCTATCAGTGGAGTAGTTACCCA |
| IGKV7-33*01_S2856_NOR | NOR/LtJ | Musmus IGKV7-33*01 F | 97.31 | 305 | GACATTGTGATGACTCAGTCTCCAACTTTCCTTGCTGTGACAGCAAGTAAGAAGGTCACCATTAGTTGCACGGCCAGTGAGAGCCTTTATTCAAGCAAACACAAGGTGAACTACTTAGCTTGGTACCAGAAGAAACCAGAGCAATCTCCTAAACTGCTGATTTATGGGGCATCCAACCGATTCACTGGGGTCCCTGATCGCTTCACAGGCAGTGGATCTGGGACAGATTTCACTCTGACCATCAGCAGTGTGCAGGCTGAAGACCTCACACATTATTACTGTGCACAGTTTTACAGCTATCCTCT |
| IGKV8-28*02_NOR | NOR/LtJ | Musmus IGKV8-28*02 F | 100 | 302 | GACATTGTGATGACACAGTCTCCATCCTCCCTGAGTGTGTCAGCAGGAGATAAGGTCACTATGAGCTGCAAGTCCAGTCAGAGTCTGTTAAACAGTAGAAACCAAAAGAACTACTTGGCCTGGTACCAGCAGAAACCATGGCAGCCTCCTAAACTGCTGATCTACGGGGCATCCACTAGGGAATCTGGGGTCCCTGATCGCTTCACAGGCAGTGGATCTGGAACAGATTTCACTCTCACCATCAGCAGTGTGCAGGCTGAAGACCTGGCAGTTTATTACTGTCAGAATGATTATAGTTATCC |
| IGKV16-104*01_S6011_NOR | NOR/LtJ | Musmus IGKV16-104*01 F | 100 | 288 | GATGTCCAGATAACCCAGTCTCCATCTTATCTTGCTGCATCTCCTGGAGAAACCATTACTATTAATTGCAGGGCAAGTAAGAGCATTAGCAAATATTTAGCCTGGTATCAAGAGAAACCTGGGAAAACTAATAAGCTTCTTATCTACTCTGGATCCACTTTGCAATCTGGAATTCCATCAAGGTTCAGTGGCAGTGGATCTGGTACAGATTTCACTCTCACCATCAGTAGCCTGGAGCCTGAAGATTTTGCAATGTATTACTGTCAACAGCATAATGAATCCCCGTAC |
| IGKV2-109*04_NOR | NOR/LtJ | Musmus IGKV2-109*04 F | 100 | 302 | GATATTGTGATGACGCAGGCTGCATTCTCCAATCCAGTCACTCTTGGAACGTCAGCTTCCATCTCCTGCAGGTCTAGTAAGAGTCTCCTACATAGTGATGGCATCACTTATTTGTATTGGTATCTGCAGAGGCCAGGCCAGTCTCCTCAGCTCCTGATATATCGGATGTCCAACCTTGCCTCAGGAGTCCCAGACAGGTTCAGTGGCAGTGGGTCAGGAACTGATTTCACACTGAGAATCAGCAGAGTGGAGGCTGAGGATGTGGGTGTTTATTACTGTGCTCAAATGCTAGAATTCCCTCC |

| | | | | | |
|---|---|---|---|---|---|
| IGKV12-38*01_S2611_NOR | NOR/LtJ | Musmus IGKV12-38*01 F | 98.21 | 287 | GACATCCAGATGACTCAGTCTCCAGCCTCCCTGGCTGCATCTGTGGGAGAAACCATCACCATCACATGTCAAGCAAGTGAGAACATTTACTTCAGTTTAGCATGGTATCAGCAGAAGCAAGGGAAATCTCCTCAGCTCCTGATCTATAATGCAAACAGCTTGGAAGATGGTGTCCCATCGAGGTTCAGTGGCAGTGGATCTGGGACACAGTATTCTATGAAGATCAACAGCATGCAGCCTGAAGATACTGCAACTTATTTCTGTAAACAGGCTTATGACTTTCCTCC |
| IGKV4-78*01_S9188_NOR | NOR/LtJ | Musmus IGKV4-78*01 F | 97.87 | 292 | CAAATTGTTCTCACCCAGTCTCCAGCAATCATGTCTGCATCTCCTGGGGAACGGGTCACCATGACCTGCAGTGCCAGCTCAAGTGTAAGTTCCAGCTACTTGTACTGGTACCAGCAGAAGCCAGGATCCTCCCCCAAACTATGGATTTATAGCACATCCAACCTGGCTTCTGGAGTCCCTGCTCGCTTCAGTGGCAGTGGGTCTGGGACCTCTTATTCTCTCACAATCAGCAGCATGGAGGCTGAAGATGCTGCCACTTATTACTGCCAGCAGTACAGTGGTTACCCATTCA |
| IGKV8-30*01_S3861_NOR | NOR/LtJ | Musmus IGKV8-30*01 F | 95.62 | 302 | GACATTGTGATGTCACAGTCTCCATCCGCCCTAGCTGTGTCAGTTGGAGAGAAGGTCACTATGAGCTGCAAGTCCAGTCAGAGCCTTTTATATAGTAGCAATCAAAAGAACTACTTGGCCTGGTACCAGCAGAAACCAGGGCAGTCTCCTAAACTGTTAATCTACTGGGCATCCACTAGGGAATCTGGGGTCCCTGACCGCTTCACAGGCAGTAGATCAGGGACAGATTTCACTCTCACCATCAGCAGTGTGCAGGCTGAAGACCTGGCCGTTTATTACTGCAAGCAATCTTATAGCTATCC |
| IGKV2-112*02_NOR | NOR/LtJ | Musmus IGKV2-112*02 F | 100 | 301 | GATATTGTGATAACCCAGGATGAACTCTCCAATCCTGTCACTTCTGGAGAATCAGTTTCCATCTCCTGCAGGTCTAGTAAGAGTCTCCTATATAAGGATGGGAAGACATACTTGAATTGGTTTCTGCAGAGGCCAGGACAGTCTCCTCAGCTCCTGGTCTATTGGATGTCCACCCGTGCATCAGGAGTCTCAGACCGGTTTAGTGGCAGTGGGTCAGGAACAGATTTCACACTGGAAATCAGTAGAGTGAAGGCTGAGGATGTCGGTGTGTATTACTGTCAACAAGTTGTAGAGTATCCTC |
| IGKV14-130*01_NOR | NOR/LtJ | Musmus IGKV14-130*01 F | 100 | 287 | GAAATCCAGATGACCCAGTCTCCATCCTCTATGTCTGCATCTCTGGGAGACAGAATAACCATCACTTGCCAGGCAACTCAAGACATTGTTAAGAATTTAAACTGGTATCAGCAGAAACCAGGGAAACCCCCTTCATTCCTGATCTATTATGCAACTGAACTGGCAGAAGGGGTCCCATCAAGGTTCAGTGGCAGTGGGTCTGGGTCAGACTATTCTCTGACAATCAGCAACCTGGAGTCTGAAGATTTTGCAGACTATTACTGTCTACAGTTTTATGAGTTTCCTCC |
| IGKV4-81*01_S2422_NOR | NOR/LtJ | Musmus IGKV4-81*01 F | 96.45 | 292 | GAAAATGTGCTGACCCAGTCTCCAGCAATCATGGCTGCATCTCCAGGGGAGAAGGTCACCATGACCTGCAGTGCCAGCTCAAGTGTAAGTTCTGGTAACTTTCACTGGTACCAGCAGAAGCCAGGCACTTCTCCCAAACTCTGGATTTATAGGACATCCAACCTGGCTTCTGGAGTCCCCGCTCGCTTCAGTGGCAGTGGGTCTGGGACCTCTTACTCTCTTACAATCAGCAGCATGGAGGCCGAAGATGCTGCCACTTATTACTGCCAGCAGTGGAGTGGTTACCCACCCA |
| IGKV8-16*01_S1056_NOR | NOR/LtJ | Musmus IGKV8-16*01 F | 98.99 | 305 | GAAATTGTGTTGACACAGTCTATACCATCCCTGACTGTGTCAGCAGGAGAGAGGGTCACTATCAGCTGCAAATCCAATCAGAATCTTTTATGGAGTGGAAACCAACGGTACTGTTTGGTCTGGCACCAGTGGAAACCAGGGCAAACTCCTACACCGTTGATCACCTGGACATCTGATAGGTACTCTGGAGTCCCTGATCGTTTCATAGGCAGTGGATCTGTGACAGATTTCACTCTGACCATCAGCAGTGTGCAGGATGAAGATGTGGCAGTTTATTTCTGTCAGCAGCATTTACACATTCCTCC |

| | | | | | |
|---|---|---|---|---|---|
| IGKV1-99*01_S3206_NOR | NOR/LtJ | Musmus IGKV1-99*01 F | 98.3 | 302 | GATGTTGTTCTGACCCAAACTCCACTCTCTCTGCCTGTCAATATTGGAGATCAAGCCTCTATCTCTTGCAAGTCTACTAAGAGCCTTCTGAATAGTGATGGATTCACTTATTTGGGCTGGTACCTGCAGAAGCCAGGCCAGTCTCCACAGCTCCTAATATATTTGGTTTCTAATCGATTTTCTGGAGTTCCAGACAGGTTCAGTGGTAGTGGGTCAGGGACAGATTTCACCCTCAAGATCAGCAGAGTGGAGGCTGAGGATTTGGGAGTTTATTATTGCTTCCAGAGTAACTATCTTCCTCT |
| IGKV13-85*01_S2277_NOR | NOR/LtJ | Musmus IGKV13-85*01 F | 97.13 | 286 | GACATCCAGATGACACAATCTTCATCCTCCTTGTCTGTATCTCTAGGAGACAGAGTCACCATTACTTGCAAGGCAAGTGAGCACATTAATAGTTGGTTAGCCTGGTATCAGCAAAAACCAGGAAATGCTCCTAGGCTCTTAATATCTGGTGCAACCAGTTTGGAAACTGGGGTTCCTTCAAGATTCAGTGGCAGTGCATCTGGAAAGGATTACACTCTCAGCATTACTAGTCTTCAGACTGAAGATGTTGCTACTTATTACTGTCAACAGTATTGGGGTACTCCTC |
| IGKV5-37*01_S7203_NOR | NOR/LtJ | Musmus IGKV5-37*01 F | 97.13 | 286 | GACATCCTGATGACCCAGTCTCCAGCCACCCTGTCTGTGACTCCAGGAGAAACAGTCAGTCTTTCCTGTAGGGCCAGCCAGAATACTTACAAGAACCTACACTGGTATCAACAGAAATCACATGGGACTCCAAAGCTTCTCATCAAGTATGCATCTGATCCCATCTCTGGGATCCCCTCCAGGTTCACTGGCAGTGGATCAGGGACAGATTACACTCTCAGTATCAACAGTGTGAAGCCCGAAGATGAAGGAATATATTACTGTCTTCAAGGTTACAGCATGCCTT |
| IGKV5-39*01_S3142_NOR | NOR/LtJ | Musmus IGKV5-39*01 F | 98.92 | 287 | GACATTGTGATGACTCAGTCTCCAGCCACCCTGTCTGTGACTCCAGGAGATAGGGTCTCTCTTTCCTGCAGGGCCAGTCAGAGTATTAGCGACTACTTACACTGGTATCAACAAAAATCACATGAGTCTCCAAGGCTTCTCATCAAGTATGCTTCCCAATCCATCTCTGGGATCCCCTCCAGGTTCAGTGGCAGTGGATCAGGGTCAGATTTCACTCTCAGTATCAACAGTGTGGAACCTGAAGATGTTGGAGTGTATTACTGTCAAAATGGTCACAGCTTTCCTCC |
| IGKV1-115*01_S9289_NOR | NOR/LtJ | Musmus IGKV1-115*01 P | 95.58 | 302 | GATGTTCTGTTGACCCAAACTCCACTTTTCCTGCATGTCAGCCTTGGAGATCAAGCCTCTATCTCTTGCACATCTAGTCAGAGTCTTGTACACAGTAATGGAAATTCCTATTTGGATTGGCACCTGCAGAAGCCAGACCAGTCTCTACAACTCCTGATCTATGAGGTTTCCAAACGAAATTCTGGGGTTCCAGACAGGTTCAGTGGCAGTGGATCAGGAACAGATTTCACACTTAAGATCAGCAGAGTAGAGCCTGAGGATTTGGGAGTTTATTACTGCTTCCAACGTACACATTTACCTCC |
| IGKV6-25*01_S5447_NOR | NOR/LtJ | Musmus IGKV6-25*01 F | 98.21 | 286 | TGCATTGTGATGACCCAGTCTCACAAATTCATGTCCACATCAGTAGGAGACAGGGTCAGCATCACCTGCAAGGCCAGTCAGGATGTGAGTACTGCTGTAGCCTGGTATCAACAAAAACCAGGGCAATCTCCTAAACTACTGATTTACTGGGCATCCACCCGGCACACTGGAGTCCCTGATCGCTTCACAGGCAGTGGATCTGGGACAGATTTCACTCTTACCATCAGCAGTGTGCAGGCTGAAGACCTGGCACTTTATTACTGTCAGCAACATTATAGCACTCCTC |
| IGKV9-123*01_S4220_NOR | NOR/LtJ | Musmus IGKV9-123*01 F | 99.28 | 285 | GACATCCAGATGATTCAGTCTCCATCGTCCATGTTTGGCTCTCTGGGAGACAGAGTCAGTCTCTCTTGCCGGGCTAGTCAGGGCATTAGAGGTAATTTAGACTGGTATCAGCAGAAACCAGGTGGAACTATTAAACTCCTGATCTACTCCACATCCAATTTAAATTCTGGTGTCCCATCAAGGTTCAGTGGCAGTGGGTCTGGGTCAGATTATTCTCTCACCATCAGCAGCCTAGAGTCTGAAGATTTTGCAGACTATTACTGTCTACAGCGTAATGCGTATCCT |
| IGKV1-99*01_S7463_NOR | NOR/LtJ | Musmus IGKV1-99*01 F | 98.3 | 302 | GATGTTGTTCTGACCCAAACTCCACTCTCTCTGCCTGTCAATATTGGAGATCAAGCCTCTATCTCTTGCAAGTCTACTAAGAGCCTTCTGAATAGTGATGGATTCACTTATTTGGGCTGGTACCTGCAGAAGCCAGGCCAGTCTCCACAGCTCCTAATATATTTGGTTTCTAATCGATTTTCTGGAGTTCCAGACAGGTTCAGTGGTAGTGGGTCAGGGACAGATTTCACCCTCAAGATCAGCAGAGTGGAGGCTGAGGATTTGGGAGTTTATTATTGCTTCCAGAGTAACTATCTTCCGCT |

| | | | | | |
|---|---|---|---|---|---|
| IGKV11-125*01_NOR | NOR/LtJ | Musmus IGKV11-125*01 F | 100 | 287 | GATGTCCAGATGATTCAGTCTCCATCCTCCCTGTCTGCATCTTTGGGAGACATAGTCACCATGACTTGCCAGGCAAGTCAGGGCACTAGCATTAATTTAAACTGGTTTCAGCAAAAACCAGGGAAAGCTCCTAAGCTCCTGATCTATGGTGCAAGCAACTTGGAAGATGGGGTCCCATCAAGGTTCAGTGGCAGTAGATATGGGACAGATTTCACTCTCACCATCAGCAGCCTGGAGGATGAAGATATGGCAACTTATTTCTGTCTACAGCATAGTTATCTCCCTCC |
| IGKV8-30*01_NOR | NOR/LtJ | Musmus IGKV8-30*01 F | 100 | 305 | GACATTGTGATGTCACAGTCTCCATCCTCCCTAGCTGTGTCAGTTGGAGAGAAGGTTACTATGAGCTGCAAGTCCAGTCAGAGCCTTTTATATAGTAGCAATCAAAAGAACTACTTGGCCTGGTACCAGCAGAAACCAGGGCAGTCTCCTAAACTGCTGATTTACTGGGCATCCACTAGGGAATCTGGGGTCCCTGATCGCTTCACAGGCAGTGGATCTGGGACAGATTTCACTCTCACCATCAGCAGTGTGAAGGCTGAAGACCTGGCAGTTTATTACTGTCAGCAATATTATAGCTATCCTCC |
| IGKV8-27*01_NOR | NOR/LtJ | Musmus IGKV8-27*01 F | 100 | 300 | AACATTATGATGACACAGTCGCCATCATCTCTGGCTGTGTCTGCAGGAGAAAAGGTCACTATGAGCTGTAAGTCCAGTCAAAGTGTTTTATACAGTTCAAATCAGAAGAACTACTTGGCCTGGTACCAGCAGAAACCAGGGCAGTCTCCTAAACTGCTGATCTACTGGGCATCCACTAGGGAATCTGGTGTCCCTGATCGCTTCACAGGCAGTGGATCTGGGACAGATTTTACTCTTACCATCAGCAGTGTACAAGCTGAAGACCTGGCAGTTTATTACTGTCATCAATACCTCTCCTCG |
| IGKV12-40*01_S9191_NOR | NOR/LtJ | Musmus IGKV12-40*01 P, or Musmus IGKV12-46*01 F | 93.91 | 286 | GACATCCAGATGACTCAGTCTCCAGCTTCCCTGTCTGCATCTGTGGGAGAAACTGTCACCATCACATGTCGAGCAAGTGAGAATATTGACAGTTATTTAGCATGGTATCAGCAGAAACAGGGAAAATCTCCTCAGCTCCTGGTCTATGTTGCAACACTCTTAGCAGATGGTGTGCCATCAAGGTTCAGTGGCAGTGGATCAGGCACACAGTATTCTCTCAAGATCAACAGCCTGCAGTCTGAAGATGTTGCGAGATATTACTGTCAGCATTATTATAGTACTCCTC |
| IGKV4-55*01_NOR | NOR/LtJ | Musmus IGKV4-55*01 F | 100 | 286 | CAAATTGTTCTCACCCAGTCTCCAGCAATCATGTCTGCATCTCCAGGGGAGAAGGTCACCATGACCTGCAGTGCCAGCTCAAGTGTAAGTTACATGTACTGGTACCAGCAGAAGCCAGGATCCTCCCCCAGACTCCTGATTTATGACACATCCAACCTGGCTTCTGGAGTCCCTGTTCGCTTCAGTGGCAGTGGGTCTGGGACCTCTTACTCTCTCACAATCAGCCGAATGGAGGCTGAAGATGCTGCCACTTATTACTGCCAGCAGTGGAGTAGTTACCCACCCA |
| IGKV4-72*01_NOR | NOR/LtJ | Musmus IGKV4-72*01 F | 100 | 286 | CAAATTGTTCTCTCCCAGTCTCCAGCAATCCTGTCTGCATCTCCAGGGGAGAAGGTCACAATGACTTGCAGGGCCAGCTCAAGTGTAAGTTACATGCACTGGTACCAGCAGAAGCCAGGATCCTCCCCCAAACCCTGGATTTATGCCACATCCAACCTGGCTTCTGGAGTCCCTGCTCGCTTCAGTGGCAGTGGGTCTGGGACCTCTTACTCTCTCACAATCAGCAGAGTGGAGGCTGAAGATGCTGCCACTTATTACTGCCAGCAGTGGAGTAGTAACCCACCCA |
| IGKV4-91*01_S5997_NOR | NOR/LtJ | Musmus IGKV4-91*01 F | 100 | 292 | GAAATTGTGCTCACCCAGTCTCCAACCACCATGGCTGCATCTCCCGGGGAGAAGATCACTATCACCTGCAGTGCCAGCTCAAGTATAAGTTCCAATTACTTGCATTGGTATCAGCAGAAGCCAGGATTCTCCCCTAAACTCTTGATTTATAGGACATCCAATCTGGCTTCTGGAGTCCCAGCTCGCTTCAGTGGCAGTGGGTCTGGGACCTCTTACTCTCTCACAATTGGCACCATGGAGGCTGAAGATGTTGCCACTTACTACTGCCAGCAGGGTAGTAGTATACCACTCA |
| IGKV8-30*01_NZB | NZB/BlNJ | Musmus IGKV8-30*01 F | 100 | 305 | GACATTGTGATGTCACAGTCTCCATCCTCCCTAGCTGTGTCAGTTGGAGAGAAGGTTACTATGAGCTGCAAGTCCAGTCAGAGCCTTTTATATAGTAGCAATCAAAAGAACTACTTGGCCTGGTACCAGCAGAAACCAGGGCAGTCTCCTAAACTGCTGATTTACTGGGCATCCACTAGGGAATCTGGGGTCCCTGATCGCTTCACAGGCAGTGGATCTGGGACAGATTTCACTCTCACCATCAGCAGTGTGAAGGCTGAAGACCTGGCAGTTTATTACTGTCAGCAATATTATAGCTATCCTCC |

| | | | | | |
|---|---|---|---|---|---|
| IGKV12-89*01_NZB | NZB/BlNJ | Musmus IGKV12-89*01 F | 100 | 287 | GACATCCAGATGACTCAGTCTCCAGCTTCACTGTCTGCATCTGTGGGAGAAACTGTCACCATCACATGTGGAGCAAGTGAGAATATTTACGGTGCTTTAAATTGGTATCAGCGGAAACAGGGAAAATCTCCTCAGCTCCTGATCTATGGTGCAACCAACTTGGCAGATGGCATGTCATCGAGGTTCAGTGGCAGTGGATCTGGTAGACAGTATTCTCTCAAGATCAGTAGCCTGCATCCTGACGATGTTGCAACGTATTACTGTCAAAATGTGTTAAGTACTCCTCC |
| IGKV1-135*01_S3001A_NZB | NZB/BlNJ | Musmus IGKV1-135*01 F | 95.92 | 302 | GATGTTGTGATGACTCAGACCCCACTCACTTTGTCGGTTACCATTGGACAACCAGCCTCCATCTCTTGCAAATCAAGTCAGAGCCTCTTACATAGTAATGGAAAGACATATTTGAATTGGTTATTACAGAGGCCAGGCCAGTCTCCAAAGCTCCTAATCTATCTGGTGTCTAAACTGGAATCTGGAGTCCCTGACAGGTTCAGTGGCAGTGGATCAGGGACAGATTTCACACTGAAAATCAGCAGAGTGGAGGCTGAGGATTTGGGAGTTTATTACTGCTTGCAAGCTACACATTTTCCTCA |
| IGKV2-137*01_S0442_NZB | NZB/BlNJ | Musmus IGKV2-137*01 F | 93.54 | 302 | GATATTGTGATGACTCAGGCTGCACCCTCTGTACCTGTCACTCCTGGAGAGTCAGTATCCATCTCCTGCAGGTCTAGTACGAGTCTCCTGCACAGTAGTGGCAAGCATAGGTTGTATTGGTTCCTACAGAGGCCAGGCCAGTCTCCTCAGCTCCTGATATATTATATGTCCAACCTTGCCTCAGGAGTCCCAGACAGGTTCAGTGGCAGTGGGTCAGGAACTGATTTCACACTGAGAATCAGTAGAGTGGAGGCTGAGGATTTTGGTGTTTATTATTGTATGCAAAGTCTAGAATATCCTTT |
| IGKV6-15*01_NZB | NZB/BlNJ | Musmus IGKV6-15*01 F | 100 | 287 | GACATTGTGATGACCCAGTCTCAAAAATTCATGTCCACATCAGTAGGAGACAGGGTCAGCGTCACCTGCAAGGCCAGTCAGAATGTGGGTACTAATGTAGCCTGGTATCAACAGAAACCAGGGCAATCTCCTAAAGCACTGATTTACTCGGCATCCTACCGGTACAGTGGAGTCCCTGATCGCTTCACAGGCAGTGGATCTGGGACAGATTTCACTCTCACCATCAGCAATGTGCAGTCTGAAGACTTGGCAGAGTATTTCTGTCAGCAATATAACAGCTATCCTCT |
| IGKV6-20*01_NZB | NZB/BlNJ | Musmus IGKV6-20*01 F | 100 | 287 | AACATTGTAATGACCCAATCTCCCAAATCCATGTCCATGTCAGTAGGAGAGAGGGTCACCTTGAGCTGCAAGGCCAGTGAGAATGTGGGTACTTATGTATCCTGGTATCAACAGAAACCAGAGCAGTCTCCTAAACTGCTGATATACGGGGCATCCAACCGGTACACTGGGGTCCCCGATCGCTTCACAGGCAGTGGATCTGCAACAGATTTCACTCTGACCATCAGCAGTGTGCAGGCTGAAGACCTTGCAGATTATCACTGTGGACAGAGTTACAGCTATCCTCC |
| IGKV1-133*01_S5139_NZB | NZB/BlNJ | Musmus IGKV1-133*01 F | 99.32 | 302 | GATGTTGTGATGACCCAGACTCCACTCACTTTGTCAGTTACCATTGGACAACCAGCCTCTATCTCTTGCAAGTCAAGTCAGAGCCTCTTATATAGTAATGGAAAAACCTATTTGAACTGGTTATTACAGAGGCCAGGCCAGTCTCCAAAGCGCCTAATCTATCTGGTGTCTAAACTGGACTCTGGAGTCCCTGACAGGTTCACTGGCAGTGGATCAGGAACAGATTTTACACTGAAAATCAGCAGAGTGGAGGCTGAGGATTTGGGAGTTTATTACTGCGTGCAAGGTACACATTTTCCTCA |
| IGKV6-13*01_NZB | NZB/BlNJ | Musmus IGKV6-13*01 F | 100 | 285 | GACATTGTGATGACCCAGTCTCAAAAATTCATGTCCACATCAGTAGGAGACAGGGTCAGCATCACCTGCAAGGCCAGTCAGAATGTGGGTACTGCTGTAGCCTGGTATCAACAGAAACCAGGACAATCTCCTAAACTACTGATTTACTCGGCATCCAATCGGTACACTGGAGTCCCTGATCGCTTCACAGGCAGTGGATCTGGGACAGATTTCACTCTCACCATCAGCAATATGCAGTCTGAAGACCTGGCAGATTATTTCTGCCAGCAATATAGCAGCTATCCT |
| IGKV14-126*01_NZB | NZB/BlNJ | Musmus IGKV14-126*01 F | 100 | 287 | GACATCAAGATGACCCAGTCTCCATCCTCCATGTATGCATCGCTGGGAGAGAGAGTCACTATCACTTGCAAGGCGAGTCAGGACATTAAAAGCTATTTAAGCTGGTACCAGCAGAAACCATGGAAATCTCCTAAGACCCTGATCTATTATGCAACAAGCTTGGCAGATGGGGTCCCATCAAGATTCAGTGGCAGTGGATCTGGGCAAGATTATTCTCTAACCATCAGCAGCCTGGAGTCTGACGATACAGCAACTTATTACTGTCTACAGCATGGTGAGAGCCCTCC |

| | | | | | |
|---|---|---|---|---|---|
| IGKV8-27*01_NZB | NZB/BlNJ | Musmus IGKV8-27*01 F | 100 | 300 | AACATTATGATGACACAGTCGCCATCATCTCTGGCTGTGTCTGCAGGAGAAAAGGTCACTATGAGCTGTAAGTCCAGTCAAAGTGTTTTATACAGTTCAAATCAGAAGAACTACTTGGCCTGGTACCAGCAGAAACCAGGGCAGTCTCCTAAACTGCTGATCTACTGGGCATCCACTAGGGAATCTGGTGTCCCTGATCGCTTCACAGGCAGTGGATCTGGGACAGATTTTACTCTTACCATCAGCAGTGTACAAGCTGAAGACCTGGCAGTTTATTACTGTCATCAATACCTCTCCTCG |
| IGKV1-135*01_S7405_NZB | NZB/BlNJ | Musmus IGKV1-135*01 F | 95.24 | 302 | GATGTTGTGATGACCCAGACTCCACTGTCTTTGTCGGTTACCATTGGACAACCAGCCTCCATCTCTTGCAAGTCAAGTCAGAGCCTCTTACATAGTGATGGAAAGACATATTTGAATTGGTTATTACAGAGGCCAGGCCAGTCTCCAAAGCTCCTAATCTATCTGGTGTCTAAACTGGAATCTGGCATCCCTGACAGGTTCAGTGGCAGTGGATCAGGGACAGATTTCACACTGAAAATCAGCAGAGTGGAGGTTGAGGATTTGGGAGTTTATTACTGCTTGCAACATACACATTTTCCTCA |
| IGKV1-132*01_S5178_NZB | NZB/BlNJ | Musmus IGKV1-132*01 F | 97.96 | 302 | GATGTTGTGATGACCCAGACTCCACTGTCTTTGTCGGTTACCATTGGACAACCAGCCTCTATCTCTTGCAAGTCAAGTCAGAGCCTCTTATATAGTGATGGAAAGACATATTTGAATTGGTTACAACAGAGGCCAGGCCAGTCTCCAAAGCGCCTAATGTATCAGGTGTCCAAACTGGACCCTGGCATCCCTGACAGGTTCAGTGGCAGTGGATCAGAGACAGATTTTACACTTAAAATCAGCAGAGTGGAGGCTGAGGATTTGGGAGTTTATTACTGCTTGCAAGGTACATATTATCCTCA |
| IGKV12-44*01_NZB | NZB/BlNJ | Musmus IGKV12-44*01 F | 100 | 287 | GACATCCAGATGACTCAGTCTCCAGCCTCCCTATCTGCATCTGTGGGAGAAACTGTCACCATCACATGTCGAGCAAGTGAGAATATTTACAGTTATTTAGCATGGTATCAGCAGAAACAGGGAAAATCTCCTCAGCTCCTGGTCTATAATGCAAAAACCTTAGCAGAAGGTGTGCCATCAAGGTTCAGTGGCAGTGGATCAGGCACACAGTTTTCTCTGAAGATCAACAGCCTGCAGCCTGAAGATTTTGGGAGTTATTACTGTCAACATCATTATGGTACTCCTCC |
| IGKV8-19*01_NZB | NZB/BlNJ | Musmus IGKV8-19*01 F | 100 | 305 | GACATTGTGATGACACAGTCTCCATCCTCCCTGACTGTGACAGCAGGAGAGAAGGTCACTATGAGCTGCAAGTCCAGTCAGAGTCTGTTAAACAGTGGAAATCAAAAGAACTACTTGACCTGGTACCAGCAGAAACCAGGGCAGCCTCCTAAACTGTTGATCTACTGGGCATCCACTAGGGAATCTGGGGTCCCTGATCGCTTCACAGGCAGTGGATCTGGAACAGATTTCACTCTCACCATCAGCAGTGTGCAGGCTGAAGACCTGGCAGTTTATTACTGTCAGAATGATTATAGTTATCCTCC |
| IGKV4-53*01_NZB | NZB/BlNJ | Musmus IGKV4-53*01 F | 100 | 292 | GAAATTGTGCTCACCCAGTCTCCAGCACTCATGGCTGCATCTCCAGGGGAGAAGGTCACCATCACCTGCAGTGTCAGCTCAAGTATAAGTTCCAGCAACTTGCACTGGTACCAGCAGAAGTCAGAAACCTCCCCCAAACCCTGGATTTATGGCACATCCAACCTGGCTTCTGGAGTCCCTGTTCGCTTCAGTGGCAGTGGATCTGGGACCTCTTATTCTCTCACAATCAGCAGCATGGAGGCTGAAGATGCTGCCACTTATTACTGTCAACAGTGGAGTAGTTACCCACTCA |
| IGKV4-59*01_NZB | NZB/BlNJ | Musmus IGKV4-59*01 F | 100 | 286 | CAAATTGTTCTCACCCAGTCTCCAGCAATCATGTCTGCATCTCCAGGGGAGAAGGTCACCATGACCTGCAGTGCCAGCTCAAGTGTAAGTTACATGCACTGGTACCAGCAGAAGTCAGGCACCTCCCCCAAAAGATGGATTTATGACACATCCAAACTGGCTTCTGGAGTCCCTGCTCGCTTCAGTGGCAGTGGGTCTGGGACCTCTTACTCTCTCACAATCAGCAGCATGGAGGCTGAAGATGCTGCCACTTATTACTGCCAGCAGTGGAGTAGTAACCCACCCA |

| | | | | | |
|---|---|---|---|---|---|
| IGKV8-21*01_NZB | NZB/BlNJ | Musmus IGKV8-21*01 F | 100 | 302 | GACATTGTGATGTCACAGTCTCCATCCTCCCTGGCTGTGTCAGCAGGAGAGAAGGTCACTATGAGCTGCAAATCCAGTCAGAGTCTGCTCAACAGTAGAACCCGAAAGAACTACTTGGCTTGGTACCAGCAGAAACCAGGGCAGTCTCCTAAACTGCTGATCTACTGGGCATCCACTAGGGAATCTGGGGTCCCTGATCGCTTCACAGGCAGTGGATCTGGGACAGATTTCACTCTCACCATCAGCAGTGTGCAGGCTGAAGACCTGGCAGTTTATTACTGCAAGCAATCTTATAATCTTCC |
| IGKV12-46*01_NZB | NZB/BlNJ | Musmus IGKV12-46*01 F | 100 | 287 | GACATCCAGATGACTCAGTCTCCAGCCTCCCTATCTGTATCTGTGGGAGAAACTGTCACCATCACATGTCGAGCAAGTGAGAATATTTACAGTAATTTAGCATGGTATCAGCAGAAACAGGGAAAATCTCCTCAGCTCCTGGTCTATGCTGCAACAAACTTAGCAGATGGTGTGCCATCAAGGTTCAGTGGCAGTGGATCAGGCACACAGTATTCCCTCAAGATCAACAGCCTGCAGTCTGAAGATTTTGGGAGTTATTACTGTCAACATTTTTGGGGTACTCCTCC |
| IGKV5-43*01_NZB | NZB/BlNJ | Musmus IGKV5-43*01 F | 100 | 287 | GATATTGTGCTAACTCAGTCTCCAGCCACCCTGTCTGTGACTCCAGGAGATAGCGTCAGTCTTTCCTGCAGGGCCAGCCAAAGTATTAGCAACAACCTACACTGGTATCAACAAAAATCACATGAGTCTCCAAGGCTTCTCATCAAGTATGCTTCCCAGTCCATCTCTGGGATCCCCTCCAGGTTCAGTGGCAGTGGATCAGGGACAGATTTCACTCTCAGTATCAACAGTGTGGAGACTGAAGATTTTGGAATGTATTTCTGTCAACAGAGTAACAGCTGGCCTCA |
| IGKV8-28*01_NZB | NZB/BlNJ | Musmus IGKV8-28*01 F | 100 | 305 | GACATTGTGATGACACAGTCTCCATCCTCCCTGAGTGTGTCAGCAGGAGAGAAGGTCACTATGAGCTGCAAGTCCAGTCAGAGTCTGTTAAACAGTGGAAATCAAAAGAACTACTTGGCCTGGTACCAGCAGAAACCAGGGCAGCCTCCTAAACTGTTGATCTACGGGGCATCCACTAGGGAATCTGGGGTCCCTGATCGCTTCACAGGCAGTGGATCTGGAACCGATTTCACTCTTACCATCAGCAGTGTGCAGGCTGAAGACCTGGCAGTTTATTACTGTCAGAATGATCATAGTTATCCTCC |
| IGKV4-91*01_NZB | NZB/BlNJ | Musmus IGKV4-91*01 F | 100 | 292 | GAAATTGTGCTCACCCAGTCTCCAACCACCATGGCTGCATCTCCCGGGGAGAAGATCACTATCACCTGCAGTGCCAGCTCAAGTATAAGTTCCAATTACTTGCATTGGTATCAGCAGAAGCCAGGATTCTCCCCTAAACTCTTGATTTATAGGACATCCAATCTGGCTTCTGGAGTCCCAGCTCGCTTCAGTGGCAGTGGGTCTGGGACCTCTTACTCTCTCACAATTGGCACCATGGAGGCTGAAGATGTTGCCACTTACTACTGCCAGCAGGGTAGTAGTATACCACGCA |
| IGKV1-110*02_NZB | NZB/BlNJ | Musmus IGKV1-110*02 F | 100 | 302 | GATGTTGTGATGACCCAAACTCCACTCTCCCTGCCTGTCAGTCTTGGAGATCAAGCTTCCATCTCTTGCAGATCTAGTCAGAGCCTTGTACACAGCAATGGAAACACCTATTTATATTGGTACCTGCAGAAGCCAGGCCAGTCTCCAAAGCTCCTGATCTACAGGGTTTCCAACCGATTTTCTGGGGTCCCAGACAGGTTCAGTGGCAGTGGATCAGGGACAGATTTCACACTCAAGATCAGCAGAGTGGAGGCTGAGGATCTGGGAGTTTATTTCTGCTTTCAAGGTACACATGTTCCTCA |
| IGKV10-96*01_NZB | NZB/BlNJ | Musmus IGKV10-96*01 F | 100 | 286 | GATATCCAGATGACACAGACTACATCCTCCCTGTCTGCCTCTCTGGGAGACAGAGTCACCATCAGTTGCAGGGCAAGTCAGGACATTAGCAATTATTTAAACTGGTATCAGCAGAAACCAGATGGAACTGTTAAACTCCTGATCTACTACACATCAAGATTACACTCAGGAGTCCCATCAAGGTTCAGTGGCAGTGGGTCTGGAACAGATTATTCTCTCACCATTAGCAACCTGGAGCAAGAAGATATTGCCACTTACTTTTGCCAACAGGGTAATACGCTTCCTC |
| IGKV4-57*01_NZB | NZB/BlNJ | Musmus IGKV4-57*01 F | 100 | 286 | CAAATTGTTCTCACCCAGTCTCCAGCAATCATGTCTGCATCTCCAGGGGAGAAGGTCACCATAACCTGCAGTGCCAGCTCAAGTGTAAGTTACATGCACTGGTTCCAGCAGAAGCCAGGCACTTCTCCCAAACTCTGGATTTATAGCACATCCAACCTGGCTTCTGGAGTCCCTGCTCGCTTCAGTGGCAGTGGATCTGGGACCTCTTACTCTCTCACAATCAGCCGAATGGAGGCTGAAGATGCTGCCACTTATTACTGCCAGCAAAGGAGTAGTTACCCACCCA |

| | | | | | |
|---|---|---|---|---|---|
| IGKV1-117*01_S1348_NZB | NZB/BlNJ | Musmus IGKV1-117*01 F | 99.32 | 301 | GATGTTGTGATGACCCAAACTCCACTCTCCCTGCCTGTCAGTCTTGGAGATCAAGCCTCCATCTCTTGCAGATCTAGTCAGAGCATTGTACATAGTAATGGAAACACCTATTTAGAATGGTACCTGCAGAAGCCAGGCCAGTCTCCAAAGCTCCTGATCTACAAAGTTTCCAACCGATTTTCTGGGGTCCCAGACAGGTTCAGTGGCAGTGGATCAGGGACAGATTTCACACTCAAGATCAGCAGAGTGGAGGCTGAGGATCTGGGAGTTTATTACTGCTTTCAAGGTTCACATGTTCCTC |
| IGKV4-70*01_NZB | NZB/BlNJ | Musmus IGKV4-70*01 F | 100 | 282 | CAAATTGTTCTCACCCAGTCTCCAGCAATCATGTCTGCATCTCCAGGGGAGAAGGTCACCATGACCTGCAGTGCCAGCTCAAGTATAAGTTACATGCACTGGTACCAGCAGAAGCCAGGCACCTCCCCCAAAAGATGGATTTATGACACATCCAAACTGGCTTCTGGAGTCCCTGCTCGCTTCAGTGGCAGTGGGTCTGGGACCTCTTATTCTCTCACAATCAGCAGCATGGAGGCTGAAGATGCTGCCACTTATTACTGCCATCAGCGGAGTAGTTACCCA |
| IGKV6-23*01_NZB | NZB/BlNJ | Musmus IGKV6-23*01 F | 100 | 287 | GACATTGTGATGACCCAGTCTCACAAATTCATGTCCACATCAGTAGGAGACAGGGTCAGCATCACCTGCAAGGCCAGTCAGGATGTGGGTACTGCTGTAGCCTGGTATCAACAGAAACCAGGGCAATCTCCTAAACTACTGATTTACTGGGCATCCACCCGGCACACTGGAGTCCCTGATCGCTTCACAGGCAGTGGATCTGGGACAGATTTCACTCTCACCATTAGCAATGTGCAGTCTGAAGACTTGGCAGATTATTTCTGTCAGCAATATAGCAGCTATCCTCT |
| IGKV8-18*01_NZB | NZB/BlNJ | Musmus IGKV8-18*01 ORF | 100 | 310 | GACATTGTGATGACCCAGTCTCCATCCTCCCTGGCTGTGACAGCAGGAGAGAAGGTCACTATGAGATGCAAGTCCAGTCAGAGTCTTTTGTGGAGTGTAAACCAAAATAACTACTTATCCTGGTACCAGCAGAAACAAGGGCAGCCTCCTAAACTGCTTATCTATGGGGCATCCATTAGAGAATCTTGGGTCCCTGATCGATTCACAGGAAGTGGATCTGGGACAGACTTCACTCTCACCATTAGCAATGTGCATGCTGAAGACCTAGCAGTTTATTACTGTCAGCACAATCATGGCAGCTTTCTCCCCC |
| IGKV3-7*01_S5296_NZB | NZB/BlNJ | Musmus IGKV3-7*02 F | 100 | 299 | GACATTGTGCTGACACAGTCTCCTGCTTCCTTAGCTGTATCTCTGGGGCAGAGGGCCACCATCTCATGCAGGGCCAGCCAAAGTGTCAGTACATCTAGCTATAGTTATATGCACTGGTACCAACAGAAACCAGGACAGCCACCCAAACTCCTCATCAAGTATGCATCCAACCTAGAATCTGGGGTCCCTGCCAGGTTCAGTGGCAGTGGGTCTGGGACAGACTTCACCCTCAACATCCATCCTGTGGAGGAGGAGGATACTGCAACATATTACTGTCAGCACAGTTGGGAGATTCCTCC |
| IGKV4-55*01_NZB | NZB/BlNJ | Musmus IGKV4-55*01 F | 100 | 286 | CAAATTGTTCTCACCCAGTCTCCAGCAATCATGTCTGCATCTCCAGGGGAGAAGGTCACCATGACCTGCAGTGCCAGCTCAAGTGTAAGTTACATGTACTGGTACCAGCAGAAGCCAGGATCCTCCCCCAGACTCCTGATTTATGACACATCCAACCTGGCTTCTGGAGTCCCTGTTCGCTTCAGTGGCAGTGGGTCTGGGACCTCTTACTCTCTCACAATCAGCCGAATGGAGGCTGAAGATGCTGCCACTTATTACTGCCAGCAGTGGAGTAGTTACCCACCCA |
| IGKV4-57*01_S9219_NZB | NZB/BlNJ | Musmus IGKV4-57-1*01 F | 100 | 286 | CAAATTGTTCTCACCCAGTCTCCAGCAATCATGTCTGCATCTCCAGGGGAGAAGGTCACCATAACCTGCAGTGCCAGCTCAAGTGTAAGTTACATGCACTGGTTCCAGCAGAAGCCAGGCACTTCTCCCAAACTCTGGATTTATAGCACATCCAACCTGGCTTCTGGAGTCCCTGCTCGCTTCAGTGGCAGTGGATCTGGGACCTCTTACTCTCTCACAATCAGCCGAATGGAGGCTGAAGATGCTGCCACTTATTACTGCCAGCAAAGGAGTAGTTACCCACTCA |
| IGKV4-80*01_NZB | NZB/BlNJ | Musmus IGKV4-80*01 F | 100 | 282 | CAAATTGTTCTCACCCAGTCTCCAGCAATCATGTCTGCATCTCTAGGGGAGGAGATCACCCTAACCTGCAGTGCCAGCTCGAGTGTAAGTTACATGCACTGGTACCAGCAGAAGTCAGGCACTTCTCCCAAACTCTTGATTTATAGCACATCCAACCTGGCTTCTGGAGTCCCTTCTCGCTTCAGTGGCAGTGGGTCTGGGACCTTTTATTCTCTCACAATCAGCAGTGTGGAGGCTGAAGATGCTGCCGATTATTACTGCCATCAGTGGAGTAGTTATCCA |

| | | | | | |
|---|---|---|---|---|---|
| IGKV8-24*01_NZB | NZB/BlNJ | Musmus IGKV8-24*01 F | 100 | 305 | GACATTGTGATGACACAGTCTCCATCCTCCCTGGCTATGTCAGTAGGACAGAAGGTCACTATGAGCTGCAAGTCCAGTCAGAGCCTTTTAAATAGTAGCAATCAAAAGAACTATTTGGCCTGGTACCAGCAGAAACCAGGACAGTCTCCTAAACTTCTGGTATACTTTGCATCCACTAGGGAATCTGGGGTCCCTGATCGCTTCATAGGCAGTGGATCTGGGACAGATTTCACTCTTACCATCAGCAGTGTGCAGGCTGAAGACCTGGCAGATTACTTCTGTCAGCAACATTATAGCACTCCTCC |
| IGKV12-41*01_NZB | NZB/BlNJ | Musmus IGKV12-41*01 F | 100 | 287 | GACATCCAGATGACTCAGTCTCCAGCCTCCCTATCTGCATCTGTGGGAGAAACTGTCACCATCACATGTCGAGCAAGTGGGAATATTCACAATTATTTAGCATGGTATCAGCAGAAACAGGGAAAATCTCCTCAGCTCCTGGTCTATAATGCAAAAACCTTAGCAGATGGTGTGCCATCAAGGTTCAGTGGCAGTGGATCAGGAACACAATATTCTCTCAAGATCAACAGCCTGCAGCCTGAAGATTTTGGGAGTTATTACTGTCAACATTTTTGGAGTACTCCTCC |
| IGKV5-48*01_NZB | NZB/BlNJ | Musmus IGKV5-48*01 F | 100 | 287 | GACATCTTGCTGACTCAGTCTCCAGCCATCCTGTCTGTGAGTCCAGGAGAAAGAGTCAGTTTCTCCTGCAGGGCCAGTCAGAGCATTGGCACAAGCATACACTGGTATCAGCAAAGAACAAATGGTTCTCCAAGGCTTCTCATAAAGTATGCTTCTGAGTCTATCTCTGGGATCCCTTCCAGGTTTAGTGGCAGTGGATCAGGGACAGATTTTACTCTTAGCATCAACAGTGTGGAGTCTGAAGATATTGCAGATTATTACTGTCAACAAAGTAATAGCTGGCCAAC |
| IGKV4-57-1*01_NZB | NZB/BlNJ | Musmus IGKV4-57*01 F | 100 | 292 | GAAAATGTGCTCACCCAGTCTCCAGCAATCATGTCTGCATCTCCAGGGGAAAAGGTCACCATGACCTGCAGGGCCAGCTCAAGTGTAAGTTCCAGTTACTTGCACTGGTACCAGCAGAAGTCAGGTGCCTCCCCCAAACTCTGGATTTATAGCACATCCAACTTGGCTTCTGGAGTCCCTGCTCGCTTCAGTGGCAGTGGGTCTGGGACCTCTTACTCTCTCACAATCAGCAGTGTGGAGGCTGAAGATGCTGCCACTTATTACTGCCAGCAGTACAGTGGTTACCCACTCA |
| IGKV4-68*01_NZB | NZB/BlNJ | Musmus IGKV4-68*01 F | 100 | 286 | CAAATTGTTCTCACCCAGTCTCCAGCACTCATGTCTGCATCTCCAGGGGAGAAGGTCACCATGACCTGCAGTGCCAGCTCAAGTGTAAGTTACATGTACTGGTACCAGCAGAAGCCAAGATCCTCCCCCAAACCCTGGATTTATCTCACATCCAACCTGGCTTCTGGAGTCCCTGCTCGCTTCAGTGGCAGTGGGTCTGGGACCTCTTACTCTCTCACAATCAGCAGCATGGAGGCTGAAGATGCTGCCACTTATTACTGCCAGCAGTGGAGTAGTAACCCACCCA |
| IGKV15-103*01_S0817_NZB | NZB/BlNJ | Musmus IGKV15-103*01 ORF | 97.85 | 287 | GACATCCAGATGAACCAGTCTCCATCCAGTCTGTCTGCATCCCTCGGAGACACAATTACCATCACTTGCCGTGCCAGTCAGAACATTAATATTTGGTTAAGCTGGTACCAGCAGAAACCAGGAAATATTCCTAAACTATTGATCTATAAGGCTTCCAACTTGCACACAGGCGTCCCATCAAGGTTTAGTGGCAGTGGATCTGGAACAGATTTCACATTAACCATCAGCAGTCTGCAGCCTGAAGACATTGCCACTTACTACTGTCTACAGGGTCAAAGTTATCCTCT |
| IGKV8-16*01_NZB | NZB/BlNJ | Musmus IGKV8-16*01 F | 100 | 305 | GAAATTGTGTTGACCCAGTCTATACCATCCCTGACTGTGTCAGCAGGAGAGAGGGTCACTATCAGCTGCAAATCCAATCAGAATCTTTTATGGAGTGGAAACCAAAGGTACTGTTTGGTCTGGCACCAGTGGAAACCAGGGCAAACTCCTACACCGTTGATCACCTGGACATCTGATAGGTACTCTGGAGTCCCTGATCGTTTCATAGGCAGTGGATCTGTGACAGATTTCACTCTGACCATCAGCAGTGTGCAGGCTGAAGATGTGGCAGTTTATTTCTGTCAGCAGCATTTACACATTCCTCC |
| IGKV4-69*01_NZB | NZB/BlNJ | Musmus IGKV4-69*01 F | 100 | 282 | CAAATTCTTCTCACCCAGTCTCCAGCAATCATGTCTGCATCTCCAGGGGAGAAGGTCACCATGACCTGCAGTGCCAGCTCAAGTGTAAGTTACATGCACTGGTACCAGCAGAAGCCAGGATCCTCGCCCAAACCCTGGATTTATGACACATCCAACCTGGCTTCTGGATTCCCTGCTCGCTTCAGTGGCAGTGGGTCTGGGACCTCTTACTCTCTCATAATCAGCAGCATGGAGGCTGAAGATGCTGCCACTTATTACTGCCATCAGCGGAGTAGTTACCCA |

| | | | | | |
|---|---|---|---|---|---|
| IGKV19-93*01_NZB | NZB/BlNJ | Musmus IGKV19-93*01 F | 100 | 287 | GACATCCAGATGACACAGTCTCCATCCTCACTGTCTGCATCTCTGGGAGGCAAAGTCA<br>CCATCACTTGCAAGGCAAGCCAAGACATTAACAAGTATATAGCTTGGTACCAACACA<br>AGCCTGGAAAAGGTCCTAGGCTGCTCATACATTACACATCTACATTACAGCCAGGCAT<br>CCCATCAAGGTTCAGTGGAAGTGGGTCTGGGAGAGATTATTCCTTCAGCATCAGCAAC<br>CTGGAGCCTGAAGATATTGCAACTTATTATTGTCTACAGTATGATAATCTTCTACC |
| IGKV6-17*01_NZB | NZB/BlNJ | Musmus IGKV6-17*01 F | 100 | 287 | GACATTGTGATGACCCAGTCTCACAAATTCATGTCCACATCAGTAGGAGACAGGGTCA<br>GCATCACCTGCAAGGCCAGTCAGGATGTGAGTACTGCTGTAGCCTGGTATCAACAGAA<br>ACCAGGACAATCTCCTAAACTACTGATTTACTCGGCATCCTACCGGTACACTGGAGTC<br>CCTGATCGCTTCACTGGCAGTGGATCTGGGACGGATTTCACTTTCACCATCAGCAGTG<br>TGCAGGCTGAAGACCTGGCAGTTTATTACTGTCAGCAACATTATAGTACTCCTCC |
| IGKV6-29*01_NZB | NZB/BlNJ | Musmus IGKV6-29*01 F | 100 | 287 | AACATTGTAATGACCCAATCTCCCAAATCCATGTCCATGTCAGTAGGAGAGAGGGTCA<br>CCTTGAGCTGCAAGGCCAGTGAGAATGTGGGTACTTATGTATCCTGGTATCAACAGAA<br>ACCAGAGCAGTCTCCTAAACTGCTGATATACGGGGCATCCAACCGGTACCCTGGGGTC<br>CCTGATCGCTTCACAGGCAGTGGATCTGCAACAGATTTCACTCTGACCATCAGCAGTC<br>TGCAGGCTGAAGACCTTGCAGATTATCACTGTGGACAGGGTTACAGCTATCTTCC |
| IGKV12-98*01_NZB | NZB/BlNJ | Musmus IGKV12-98*01 F | 100 | 287 | GACATTCAGATGACCCAGTCTCCTGCCTCCCAGTCTGCATCTCTGGGAGAAAGTGTCA<br>CCATCACATGCCTGGCAAGTCAGACCATTGGTACATGGTTAGCATGGTATCAGCAGAA<br>ACCAGGGAAATCTCCTCAGCTCCTGATTTATGCTGCAACCAGCTTGGCAGATGGGGTC<br>CCATCAAGGTTCAGTGGTAGTGGATCTGGCACAAAATTTTCTTTCAAGATCAGCAGCC<br>TACAGGCTGAAGATTTTGTAAGTTATTACTGTCAACAACTTTACAGTACTCCTCT |
| IGKV3-10*01_NZB | NZB/BlNJ | Musmus IGKV3-10*01 F | 100 | 299 | AACATTGTGCTGACCCAATCTCCAGCTTCTTTGGCTGTGTCTCTAGGGCAGAGGGCCA<br>CCATATCCTGCAGAGCCAGTGAAAGTGTTGATAGTTATGGCAATAGTTTTATGCACTG<br>GTACCAGCAGAAACCAGGACAGCCACCCAAACTCCTCATCTATCTTGCATCCAACCTA<br>GAATCTGGGGTCCCTGCCAGGTTCAGTGGCAGTGGGTCTAGGACAGACTTCACCCTCA<br>CCATTGATCCTGTGGAGGCTGATGATGCTGCAACCTATTACTGTCAGCAAAATAATGA<br>GGATCCTCC |
| IGKV4-72*01_NZB | NZB/BlNJ | Musmus IGKV4-72*01 F | 100 | 286 | CAAATTGTTCTCTCCCAGTCTCCAGCAATCCTGTCTGCATCTCCAGGGGAGAAGGTCA<br>CAATGACTTGCAGGGCCAGCTCAAGTGTAAGTTACATGCACTGGTACCAGCAGAAGC<br>CAGGATCCTCCCCCAAACCCTGGATTTATGCCACATCCAACCTGGCTTCTGGAGTCCCT<br>GCTCGCTTCAGTGGCAGTGGGTCTGGGACCTCTTACTCTCTCACAATCAGCAGAGTGG<br>AGGCTGAAGATGCTGCCACTTATTACTGCCAGCAGTGGAGTAGTAACCCACCCA |
| IGKV6-32*01_NZB | NZB/BlNJ | Musmus IGKV6-32*01 F | 100 | 287 | AGTATTGTGATGACCCAGACTCCCAAATTCCTGCTTGTATCAGCAGGAGACAGGGTTA<br>CCATAACCTGCAAGGCCAGTCAGAGTGTGAGTAATGATGTAGCTTGGTACCAACAGA<br>AGCCAGGGCAGTCTCCTAAACTGCTGATATACTATGCATCCAATCGCTACACTGGAGT<br>CCCTGATCGCTTCACTGGCAGTGGATATGGGACGGATTTCACTTTCACCATCAGCACT<br>GTGCAGGCTGAAGACCTGGCAGTTTATTTCTGTCAGCAGGATTATAGCTCTCCTCC |
| IGKV10-94*01_NZB | NZB/BlNJ | Musmus IGKV10-94*01 F | 100 | 287 | GATATCCAGATGACACAGACTACATCCTCCCTGTCTGCCTCTCTGGGAGACAGAGTCA<br>CCATCAGTTGCAGTGCAAGTCAGGGCATTAGCAATTATTTAAACTGGTATCAGCAGAA<br>ACCAGATGGAACTGTTAAACTCCTGATCTATTACACATCAAGTTTACACTCAGGAGTC<br>CCATCAAGGTTCAGTGGCAGTGGGTCTGGGACAGATTATTCTCTCACCATCAGCAACC<br>TGGAACCTGAAGATATTGCCACTTACTATTGTCAGCAGTATAGTAAGCTTCCTCC |

| | | | | | |
|---|---|---|---|---|---|
| IGKV14-100*01_NZB | NZB/BlNJ | Musmus IGKV14-100*01 F | 100 | 287 | GACATCCTGATGACCCAATCTCCATCCTCCATGTCTGTATCTCTGGGAGACACAGTCAGCATCACTTGCCATGCAAGTCAGGGCATTAGCAGTAATATAGGGTGGTTGCAGCAGAAACCAGGGAAATCATTTAAGGGCCTGATCTATCATGGAACCAACTTGGAAGATGGAGTTCCATCAAGGTTCAGTGGCAGTGGATCTGGAGCAGATTATTCTCTCACCATCAGCAGCCTGGAATCTGAAGATTTTGCAGACTATTACTGTGTACAGTATGCTCAGTTTCCTCC |
| IGKV4-63*01_NZB | NZB/BlNJ | Musmus IGKV4-63*01 F | 100 | 286 | GAAAATGTTCTCACCCAGTCTCCAGCAATCATGTCTGCATCTCCAGGGGAAAAGGTCACCATGACCTGCAGTGCCAGCTCAAGTGTAAGTTACATGCACTGGTACCAGCAGAAGTCAAGCACCTCCCCCAAACTCTGGATTTATGACACATCCAAACTGGCTTCTGGAGTCCCAGGTCGCTTCAGTGGCAGTGGGTCTGGAAACTCTTACTCTCTCACGATCAGCAGCATGGAGGCTGAAGATGTTGCCACTTATTACTGTTTTCAGGGGAGTGGGTACCCACTCA |
| IGKV4-74*01_NZB | NZB/BlNJ | Musmus IGKV4-74*01 F | 100 | 292 | CAAATTGTTCTCACCCAGTCTCCAGCAATCATGTCTGCATCTCTAGGGGAACGGGTCACCATGACCTGCACTGCCAGCTCAAGTGTAAGTTCCAGTTACTTGCACTGGTACCAGCAGAAGCCAGGATCCTCCCCCAAACTCTGGATTTATAGCACATCCAACCTGGCTTCTGGAGTCCCAGCTCGCTTCAGTGGCAGTGGGTCTGGGACCTCTTACTCTCTCACAATCAGCAGCATGGAGGCTGAAGATGCTGCCACTTATTACTGCCACCAGTATCATCGTTCCCCACCCA |
| IGKV8-34*01_NZB | NZB/BlNJ | Musmus IGKV8-34*01 F | 100 | 305 | GACATTTTGATGACCCAGTCTCCATCCTCCCTGACTGTGTCAGCAGGAGAGAAGGTCACTATGAGCTGCAAGTCCAGTCAGAGTCTTTTAGCTAGTGGCAACCAAAATAACTACTTGGCCTGGCACCAGCAGAAACCAGGACGATCTCCTAAAATGCTGATAATTTGGGCATCCACTAGGGTATCTGGAGTCCCTGATCGCTTCATAGGCAGTGGATCTGGGACGGATTTCACTCTGACCATCAACAGTGTGCAGGCTGAAGATCTGGCTGTTTATTACTGTCAGCAGTCCTACAGCGCTCCTAC |
| IGKV1-122*01_S7970_NZB | NZB/BlNJ | Musmus IGKV1-122*01 F | 98.64 | 302 | GATGCTGTGATGACCCAAACTCCACTCTCCCTGCCTGTCAGTCTTGGAGATCAAGCCTCCATCTCTTGCAGGTCTAGTCAGAGCCTTGAAAACAGTAATGGAAACACCTATTTGAACTGGTACCTCCAGAAACCAGGCCAGTCTCCACAGCTCCTGATCTACAGGGTTTCCAACCGATTTTCTGGGGTCCTAGACAGGTTCAGTGGCAGTGGTTCAGGGACAGATTTCACACTCAAGATCAGCAGAGTGGAGGCTGAGGATTTGGGAGTTTATTTCTGCCTCCAAGTTACACATGTCCCTCC |
| IGKV9-124*01_S2023_NZB | NZB/BlNJ | Musmus IGKV9-124*01 F | 98.92 | 285 | GACATCCAGATGACCCAGTCTCCATCCTCCTTATCTGCCTCTCTGGGAGAAAGAGTCAGTCTCACTTGTCGGGCAAGTCAGGAAATTAGTGGTTACTTAAGCTGGCTTCAGCAGAAACCAGATGGAACTATTAAACGCCTGATCTACAGCACATCCACTTTAGATTCTGGTGTCCCAAAAAGGTTCAGTGGCAGTAGGTCTGGGTCAGATTATTCTCTCACCATCAGCAGCCTTGAGTCTGAAGATTTTGCAGACTATTACTGTCTACAATATGCTAGTTCTCCT |
| IGKV16-104*01_S6011_NZB | NZB/BlNJ | Musmus IGKV16-104*01 F | 100 | 288 | GATGTCCAGATAACCCAGTCTCCATCTTATCTTGCTGCATCTCCTGGAGAAACCATTACTATTAATTGCAGGGCAAGTAAGAGCATTAGCAAATATTTAGCCTGGTATCAAGAGAAACCTGGGAAAACTAATAAGCTTCTTATCTACTCTGGATCCACTTTGCAATCTGGAATTCCATCAAGGTTCAGTGGCAGTGGATCTGGTACAGATTTCACTCTCACCATCAGTAGCCTGGAGCCTGAAGATTTTGCAATGTATTACTGTCAACAGCATAATGAATCCCCGTAC |
| IGKV13-84*01_NZB | NZB/BlNJ | Musmus IGKV13-84*01 F | 100 | 287 | GACATCCAGATGACACAATCTTCATCCTCCTTTTCTGTATCTCTAGGAGACAGAGTCACCATTACTTGCAAGGCAAGTGAGGACATATATAATCGGTTAGCCTGGTATCAGCAGAAACCAGGAAATGCTCCTAGGCTCTTAATATCTGGTGCAACCAGTTTGGAAACTGGGGTTCCTTCAAGATTCAGTGGCAGTGGATCTGGAAAGGATTACACTCTCAGCATTACCAGTCTTCAGACTGAAGATGTTGCTACTTATTACTGTCAACAGTATTGGAGTACTCCTCC |

| | | | | | |
|---|---|---|---|---|---|
| IGKV5-39*01_NZB | NZB/BlNJ | Musmus IGKV5-39*01 F | 100 | 287 | GACATTGTGATGACTCAGTCTCCAGCCACCCTGTCTGTGACTCCAGGAGATAGAGTCTCTCTTTCCTGCAGGGCCAGCCAGAGTATTAGCGACTACTTACACTGGTATCAACAAAATCACATGAGTCTCCAAGGCTTCTCATCAAATATGCTTCCCAATCCATCTCTGGGATCCCCTCCAGGTTCAGTGGCAGTGGATCAGGGTCAGATTTCACTCTCAGTATCAACAGTGTGGAACCTGAAGATGTTGGAGTGTATTACTGTCAAAATGGTCACAGCTTTCCTCC |
| IGKV1-115*01_S3051_NZB | NZB/BlNJ | Musmus IGKV1-115*01 P | 95.58 | 301 | GATGTTCTGTTGACCCAAACTCCACTTTTCCTGCATGTCAGCCTTGGAGATCAAGCCTCTATCTCTTGCACATCTAGTCAGAGTCTTGTACACAGTAATGGAAATTCCTATTTGGATTGGCACCTGCAGAAGCCAGACCAGTCTCTACAACTCCTGATCTATGAGGTTTCCAAACGAAATTCTGGGGTTCCAGACAGGTTCAGTGGCAGTGGATCAGGAACAGATTTCACACTTAAGATCAGCAGAGTAGAGCCTGAGGATTTGGGAGTTTATTACTGCTTCCAACGTACACATTTACCTC |
| IGKV3-2*01_NZB | NZB/BlNJ | Musmus IGKV3-2*01 F | 100 | 299 | GACATTGTGCTGACCCAATCTCCAGCTTCTTTGGCTGTGTCTCTAGGGCAGAGGGCCACCATCTCCTGCAGAGCCAGCGAAAGTGTTGATAATTATGGCATTAGTTTTATGAACTGGTTCCAACAGAAACCAGGACAGCCACCCAAACTCCTCATCTATGCTGCATCCAACCAAGGATCCGGGGTCCCTGCCAGGTTTAGTGGCAGTGGGTCTGGGACAGACTTCAGCCTCAACATCCATCCTATGGAGGAGGATGATACTGCAATGTATTTCTGTCAGCAAAGTAAGGAGGTTCCTCC |
| IGKV6-25*01_NZB | NZB/BlNJ | Musmus IGKV6-25*01 F | 100 | 287 | GACATTGTGATGACCCAGTCTCACAAATTCATGTCCACATCAGTAGGAGACAGGGTCAGCATCACCTGCAAGGCCAGTCAGGATGTGAGTACTGCTGTAGCCTGGTATCAACAAAAACCAGGGCAATCTCCTAAACTACTGATTTACTGGGCATCCACCCGGCACACTGGAGTCCCTGATCGCTTCACAGGCAGTGGATCTGGGACAGATTATACTCTCACCATCAGCAGTGTGCAGGCTGAAGACCTGGCACTTTATTACTGTCAGCAACATTATAGCACTCCTCC |
| IGKV14-130*01_NZB | NZB/BlNJ | Musmus IGKV14-130*01 F | 100 | 287 | GAAATCCAGATGACCCAGTCTCCATCCTCTATGTCTGCATCTCTGGGAGACAGAATAACCATCACTTGCCAGGCAACTCAAGACATTGTTAAGAATTTAAACTGGTATCAGCAGAAACCAGGGAAACCCCCTTCATTCCTGATCTATTATGCAACTGAACTGGCAGAAGGGGTCCCATCAAGGTTCAGTGGCAGTGGGTCTGGGTCAGACTATTCTCTGACAATCAGCAACCTGGAGTCTGAAGATTTTGCAGACTATTACTGTCTACAGTTTTATGAGTTTCCTCC |
| IGKV2-109*03_NZB | NZB/BlNJ | Musmus IGKV2-109*03 F | 100 | 302 | GATATTGTGATGACGCAGGCTGCATTCTCCAATCCAGTCACTCTTGGAACATCAGCTTCCATCTCTTGCAGTTCTAGTAAGAGTCTCCTACATAGTAATGGCATCACTTATTTGTATTGGTATCTGCAGAGGCCAGGCCAGTCTCCTCAGCTCCTGATATATCGGATGTCCAACCTTGCCTCAGGAGTCCCAGACAGGTTCAGTGGCAGTGGGTCAGGAACTGATTTCACACTGAGAATCAGCAGAGTGGAGGCTGAGGATGTGGGTGTTTATTACTGTGCTCAAATGCTAGAACGCCCTCC |
| IGKV3-12*01_NZB | NZB/BlNJ | Musmus IGKV3-12*01 F | 100 | 299 | GACATTGTGCTGACACAGTCTCCTGCTTCCTTAGCTGTATCTCTGGGGCAGAGGGCCACCATCTCATGCAGGGCCAGCAAAAGTGTCAGTACATCTGGCTATAGTTATATGCACTGGTACCAACAGAAACCAGGACAGCCACCCAAACTCCTCATCTATCTTGCATCCAACCTAGAATCTGGGGTCCCTGCCAGGTTCAGTGGCAGTGGGTCTGGGACAGACTTCACCCTCAACATCCATCCTGTGGAGGAGGAGGATGCTGCAACCTATTACTGTCAGCACAGTAGGGAGCTTCCTCC |
| IGKV4-50*01_NZB | NZB/BlNJ | Musmus IGKV4-50*01 F | 100 | 286 | GAAAATGTGCTCACCCAGTCTCCAGCAATCATGTCTGCATCTCTAGGGGAGAAGGTCACCATGAGCTGCAGGGCCAGCTCAAGTGTAAATTACATGTACTGGTACCAGCAGAAGTCAGATGCCTCCCCCAAACTATGGATTTATTACACATCCAACCTGGCTCCTGGAGTCCCAGCTCGCTTCAGTGGCAGTGGGTCTGGGAACTCTTATTCTCTCACAATCAGCAGCATGGAGGGTGAAGATGCTGCCACTTATTACTGCCAGCAGTTTACTAGTTCCCCATCCA |

| | | | | | |
|---|---|---|---|---|---|
| IGKV4-79*01_NZB | NZB/BlNJ | Musmus IGKV4-79*01 F | 100 | 292 | CAAATTGTTCTCACCCAGTCTCCAGCAATCATGTCTGCATCTCCTGGGGAGAAGGTCACCTTGACCTGCAGTGCCAGCTCAAGTGTAAGTTCCAGCTACTTGTACTGGTACCAGCAGAAGCCAGGATCCTCCCCCAAACTCTGGATTTATAGCACATCCAACCTGGCTTCTGGAGTCCCTGCTCGCTTCAGTGGCAGTGGGTCTGGGACCTCTTACTCTCTCACAATCAGCAGCATGGAGGCTGAAGATGCTGCCTCTTATTTCTGCCATCAGTGGAGTAGTTACCCACCCC |
| IGKV4-61*01_NZB | NZB/BlNJ | Musmus IGKV4-61*01 F | 100 | 286 | CAAATTGTTCTCACCCAGTCTCCAGCAATCATGTCTGCATCTCCAGGGGAGAAGGTCACCATATCCTGCAGTGCCAGCTCAAGTGTAAGTTACATGTACTGGTACCAGCAGAAGCCAGGATCCTCCCCCAAACCCTGGATTTATCGCACATCCAACCTGGCTTCTGGAGTCCCTGCTCGCTTCAGTGGCAGTGGGTCTGGGACCTCTTACTCTCTCACAATCAGCAGCATGGAGGCTGAAGATGCTGCCACTTATTACTGCCAGCAGTATCATAGTTACCCACCCA |
| IGKV9-120*01_S7607_NZB | NZB/BlNJ | Musmus IGKV9-120*02 F | 96.42 | 285 | GACATCCAGATGACCCAGTCTCCATCCTCCTTATCTGCCTCTCTGGGAGAAAGAATCAGTCTCACTTGCCGGGCAAGTCAGGACATTTATGGTAGCTTAAACTGGTTTCAGCAGAAACCAGATGGAACTATTAAACTCCTGATCTACGGCACATCCAGTTTAGATTCTGGTGTCCCCAAAAGGTTCAGTGGCAGTAGGTCTGGGTCAGATTATTCTCTCACCATCAGCAGCCTTGAGTCTGAAGATTTTGCAGACTATTACTGTCTACAATATGCTAGTTCTCCT |
| IGKV17-127*01_S4378_NZB | NZB/BlNJ | Musmus IGKV17-127*01 F | 99.64 | 287 | GAAACAACTGTGACCCAGTCTCCAGCATCCCTGTCCGTGGCTACAGGAGAAAAAGTCACTATCAGATGCATAACCAACACTGATATTGATGATGATATGAACTGGTACCAGCAGAAGCCAGGGGAACCTCCTAAGCTCCTTATTTCAGAAGGCAATACTCTTCGTCCTGGAGTCCCATCCCGATTCTCCAGCAGTGGCTATGGCACAGATTTTGTTTTTACAATTGAAAACACGCTCTCAGAAGATGTTGCAGATTACTACTGTTTGCAAAGTGATAACATGCCTCT |
| IGKV2-112*02_NZB | NZB/BlNJ | Musmus IGKV2-112*02 F | 100 | 301 | GATATTGTGATAACCCAGGATGAACTCTCCAATCCTGTCACTTCTGGAGAATCAGTTTCCATCTCCTGCAGGTCTAGTAAGAGTCTCCTATATAAGGATGGGAAGACATACTTGAATTGGTTTCTGCAGAGGCCAGGACAGTCTCCTCAGCTCCTGGTCTATTGGATGTCCACCCGTGCATCAGGAGTCTCAGACCGGTTTAGTGGCAGTGGGTCAGGAACAGATTTCACACTGGAAATCAGTAGAGTGAAGGCTGAGGATGTCGGTGTGTATTACTGTCAACAAGTTGTAGAGTATCCTC |
| IGKV3-5*01_NZB | NZB/BlNJ | Musmus IGKV3-5*01 F | 100 | 299 | GACATTGTGCTGACCCAATCTCCAGCTTCTTTGGCTGTGTCTCTAGGGCAGAGGGCCACCATATCCTGCAGAGCCAGTGAAAGTGTTGATAGTTATGGCAATAGTTTTATGCACTGGTACCAGCAGAAACCAGGACAGCCACCCAAACTCCTCATCTATCGTGCATCCAACCTAGAATCTGGGATCCCTGCCAGGTTCAGTGGCAGTGGGTCTAGGACAGACTTCACCCTCACCATTAATCCTGTGGAGGCTGATGATGTTGCAACCTATTACTGTCAGCAAAGTAATGAGGATCCTCC |
| IGKV1-88*01_NZB | NZB/BlNJ | Musmus IGKV1-88*01 F | 100 | 302 | GATGTTGTGGTGACTCAAACTCCACTCTCCCTGCCTGTCAGCTTTGGAGATCAAGTTTCTATCTCTTGCAGGTCTAGTCAGAGTCTTGCAAACAGTTATGGGAACACCTATTTGTCTTGGTACCTGCACAAGCCTGGCCAGTCTCCACAGCTCCTCATCTATGGGATTTCCAACAGATTTTCTGGGGTGCCAGACAGGTTCAGTGGCAGTGGTTCAGGGACAGATTTCACACTCAAGATCAGCACAATAAAGCCTGAGGACTTGGGAATGTATTACTGCTTACAAGGTACACATCAGCCTCC |
| IGKV13-85*01_NZB | NZB/BlNJ | Musmus IGKV13-85*01 F | 100 | 287 | GACATCCAGATGACACAATCTTCATCCTACTTGTCTGTATCTCTAGGAGGCAGAGTCACCATTACTTGCAAGGCAAGTGACCACATTAATAATTGGTTAGCCTGGTATCAGCAGAAACCAGGAAATGCTCCTAGGCTCTTAATATCTGGTGCAACCAGTTTGGAAACTGGGGTTCCTTCAAGATTCAGTGGCAGTGGATCTGGAAAGGATTACACTCTCAGCATTACCAGTCTTCAGACTGAAGATGTTGCTACTTATTACTGTCAACAGTATTGGAGTACTCCTCC |

| | | | | | |
|---|---|---|---|---|---|
| IGKV9-123*01_S4220_NZB | NZB/BlNJ | Musmus IGKV9-123*01 F | 99.28 | 285 | GACATCCAGATGATTCAGTCTCCATCGTCCATGTTTGGCTCTCTGGGAGACAGAGTCAGTCTCTCTTGCCGGGCTAGTCAGGGCATTAGAGGTAATTTAGACTGGTATCAGCAGAAACCAGGTGGAACTATTAAACTCCTGATCTACTCCACATCCAATTTAAATTCTGGTGTCCCATCAAGGTTCAGTGGCAGTGGGTCTGGGTCAGATTATTCTCTCACCATCAGCAGCCTAGAGTCTGAAGATTTTGCAGACTATTACTGTCTACAGCGTAATGCGTATCCT |
| IGKV8-23-1*01_NZB | NZB/BlNJ | Musmus IGKV8-16*01 F, or Musmus IGKV8-23-1*01 ORF | 100 | 284 | GAAATTGTGCTGACTGTGTCAGCAGGAGAGAGGGTCACTATCAGCTGCAAATCCAATCAGAATCTTTTATGGAGTGGAAACCAAAGGTACTGTTTGGTCTGGCACCAGTGGAAACCAGGGCAAACTCCTACACCGTTGATCACCTGGACATCTGATAGGTACTCTGGAGTCCCTGATCGTTTCATAGGCAGTGGATCTGTGACAGATTTCACTCTGACCATCAGCAGTGTGCAGGCTGAAGATGTGGCAGTTTATTTCTGTCAGCAGCATTTACACATTCCTCC |
| IGKV3-4*01_NZB | NZB/BlNJ | Musmus IGKV3-4*01 F | 100 | 299 | GACATTGTGCTGACCCAATCTCCAGCTTCTTTGGCTGTGTCTCTAGGGCAGAGGGCCACCATCTCCTGCAAGGCCAGCCAAAGTGTTGATTATGATGGTGATAGTTATATGAACTGGTACCAACAGAAACCAGGACAGCCACCCAAACTCCTCATCTATGCTGCATCCAATCTAGAATCTGGGATCCCAGCCAGGTTTAGTGGCAGTGGGTCTGGGACAGACTTCACCCTCAACATCCATCCTGTGGAGGAGGAGGATGCTGCAACCTATTACTGTCAGCAAAGTAATGAGGATCCTCC |
| IGKV5-45*01_NZB | NZB/BlNJ | Musmus IGKV5-45*01 F | 100 | 287 | GATATTGTGCTAACTCAGTCTCCAGCCACCCTGTCTGTGACTCCAGGAGATAGAGTCAGTCTTTCCTGCAGGGCCAGTCAAAGTATTAGCAACTACCTACACTGGTATCAACAAAAATCACATGAGTCTCCAAGGCTTCTCATCAAGTATGCTTCCCAGTCCATCTCTGGGATCCCCTCCAGGTTCAGTGGCAGTGGATCAGGGACAGATTTCACTCTCAGTATCAACAGTGTGGAGACTGAAGATTTTGGAATGTATTTCTGTCAACAGAGTAACAGCTGGCCTCA |
| IGKV2-109*04_NZB | NZB/BlNJ | Musmus IGKV2-109*04 F | 100 | 302 | GATATTGTGATGACGCAGGCTGCATTCTCCAATCCAGTCACTCTTGGAACGTCAGCTTCCATCTCCTGCAGGTCTAGTAAGAGTCTCCTACATAGTGATGGCATCACTTATTTGTATTGGTATCTGCAGAGGCCAGGCCAGTCTCCTCAGCTCCTGATATATCGGATGTCCAACCTTGCCTCAGGAGTCCCAGACAGGTTCAGTGGCAGTGGGTCAGGAACTGATTTCACACTGAGAATCAGCAGAGTGGAGGCTGAGGATGTGGGTGTTTATTACTGTGCTCAAATGCTAGAATTCCCTCC |
| IGKV4-86*01_NZB | NZB/BlNJ | Musmus IGKV4-86*01 F | 100 | 283 | GAAATTGTGCTCACTCAGTCTCCAGCCATCACAGCTGCATCTCTGGGGCAAAAGGTCACCATCACCTGCAGTGCCAGCTCAAGTGTAAGTTACATGCACTGGTACCAGCAGAAGTCAGGCACCTCCCCCAAACCATGGATTTATGAAATATCCAAACTGGCTTCTGGAGTCCCAGCTCGCTTCAGTGGCAGTGGGTCTGGGACCTCTTACTCTCTCACAATCAGCAGCATGGAGGCTGAAGATGCTGCCATTTATTACTGCCAGCAGTGGAATTATCCTCTTA |
| IGKV7-33*01_NZB | NZB/BlNJ | Musmus IGKV7-33*01 F | 100 | 305 | GACATTGTGATGACTCAGTCTCCAACTTTCCTTGCTGTGACAGCAAGTAAGAAGGTCACCATTAGTTGCACGGCCAGTGAGAGCCTTTATTCAAGCAAACACAAGGTGCACTACTTGGCTTGGTACCAGAAGAAACCAGAGCAATCTCCTAAACTGCTGATATACGGGGCATCCAACCGATACATTGGGGTCCCTGATCGCTTCACAGGCAGTGGATCTGGGACAGATTTCACTCTGACCATCAGCAGTGTACAGGTTGAAGACCTCACACATTATTACTGTGCACAGTTTTACAGCTATCCTCT |
| IGKV1-99*01_NZB | NZB/BlNJ | Musmus IGKV1-99*01 F | 100 | 302 | GATGTTGTTCTGACCCAAACTCCACTCTCTCTGCCTGTCAATATTGGAGATCAAGCCTCTATCTCTTGCAAGTCTACTAAGAGTCTTCTGAATAGTGATGGATTCACTTATTTGGACTGGTACCTGCAGAAGCCAGGCCAGTCTCCACAGCTCCTAATATATTTGGTTTCTAATCGATTTTCTGGAGTTCCAGACAGGTTCAGTGGCAGTGGGTCAGGAACAGATTTCACACTCAAGATCAGCAGAGTGGAGGCTGAGGATTTGGGAGTTTATTATTGCTTCCAGAGTAACTATCTTCCTCT |

| | | | | | |
|---|---|---|---|---|---|
| IGKV10-95*01_NZB | NZB/BlNJ | Musmus IGKV10-95*01 F | 100 | 287 | GATATCCAGATGACACAGACTACTTCCTCCCTGTCTGCCTCTCTGGGAGACAGAGTCACCATCAGTTGCAGGGCAAGTGAGGACATTAGCACTTATTTAAACTGGTATCAGCAGAAACCAGATGGAACTGTTAAACTCCTGATCTATTACACATCAGGATTACACTCAGGAGTCCCATCAAGGTTCAGTGGCAGTGGGTCTGGGGCAGATTATTCTCTCACCATCAGCAACCTGGAACCTGAAGATATTGCCACTTACTATTGTCAGCAGTATAGTAAGCTTCCTCC |
| IGKV4-58*01_NZB | NZB/BlNJ | Musmus IGKV4-58*01 F | 100 | 292 | GAAAATGTGCTCACCCAGTCTCCAGCAATAATGGCTGCCTCTCTGGGGCAGAAGGTCACCATGACCTGCAGTGCCAGCTCAAGTGTAAGTTCCAGTTACTTGCACTGGTACCAGCAGAAGTCAGGCGCTTCCCCCAAACCCTTGATTCATAGGACATCCAACCTGGCTTCTGGAGTCCCAGCTCGCTTCAGTGGCAGTGGGTCTGGGACCTCTTACTCTCTCACAATCAGCAGCGTGGAGGCTGAAGATGATGCAACTTATTACTGCCAGCAGTGGAGTGGTTACCCATTCA |
| IGKV4-71*01_NZB | NZB/BlNJ | Musmus IGKV4-71*01 F | 100 | 286 | CAAATTGTTCTCACCCAGTCTCCAGCAATCATGTCTGCATCTCCAGGGGAGAAGGTCACCATGACCTGCAGTGCCAGCTCAAGTGTAAGTTACATGCACTGGTACCAGCAGAAGCCAGGATCCTCCCCCAGACTCTGGATTTATTTAACATTCAACTTGGCTTCTGGAGTCCCTGCTCGCTTCAGTGGCAGTGGGTCTGGGACCTCTTACTCTCTCTCAATCAGCAGCATGGAGGCTGAAGATGCTGCCACTTATTACTGCCAGCAGTGGAGTAGTAACCCACCCA |
| IGKV11-125*01_NZB | NZB/BlNJ | Musmus IGKV11-125*01 F | 100 | 287 | GATGTCCAGATGATTCAGTCTCCATCCTCCCTGTCTGCATCTTTGGGAGACATAGTCACCATGACTTGCCAGGCAAGTCAGGGCACTAGCATTAATTTAAACTGGTTTCAGCAAAAACCAGGGAAAGCTCCTAAGCTCCTGATCTATGGTGCAAGCAACTTGGAAGATGGGGTCCCATCAAGGTTCAGTGGCAGTAGATATGGGACAGATTTCACTCTCACCATCAGCAGCCTGGAGGATGAAGATATGGCAACTTATTTCTGTCTACAGCATAGTTATCTCCCTCC |
| IGKV18-36*01_NZB | NZB/BlNJ | Musmus IGKV18-36*01 F | 100 | 287 | ACTGGAGAAACAACACAGGCTCCAGCTTCTCTGAGTTTTCTCTTGGTGAAACAGCAACACTGTCATGCAGGTCCAGTGAGAGTGTTGGCAGCTACTTAGCCTGGTACCAGCAGAAAGCAGAGCAAGTTCCCCGGCTCCTTATCCATAGTGCCTCCACTAGGGCCGGTGGTGTCCCAGTCCGGTTCAGTGGCACTGGGTCTGGGACAGACTTCACTCTCACCATCAGCAGTCTAGAACCTGAAGATGCTGCAGTTTACTACTGTCAACCTTTCAAAAGTTGGTCATA |
| IGKV4-52*01_NZB | NZB/BlNJ | Musmus IGKV4-52*01 ORF, or Musmus IGKV4-54*01 ORF | 100 | 286 | CAAATTGTTCTCACCCAGTCTCCAGCAATCCTGTCTGCATCTCCAGGGGAGAAGGTCACCATGACCTGCAGTGCCAGCTCAAGTGTAAGTTACATGTACAGGTACCAGCAGAAGCCAGGATCCTCACCCAAACCCTGGATTTATGGCACATCCAACCTGGCTTCTGGAGTCCCTGCTCGCTTCAGTGGCAGTGGATCTGGGACCTCTTATTCTCTCACAATCAGCAGCATGGAGGCTGAAGATGCTGCCACTTATTACTGCCAGCAATATCATAGTTACCCACCCA |
| IGKV4-90*01_NZB | NZB/BlNJ | Musmus IGKV4-90*01 F | 100 | 285 | GAAATTTTGCTCACCCAGTCTCCAGCAATCATAGCTGCATCTCCTGGGGAGAAGGTCACCATCACCTGCAGTGCCAGCTCAAGTGTAAGTTACATGAACTGGTACCAGCAGAAACCAGGATCCTCCCCCAAAATATGGATTTATGGTATATCCAACCTGGCTTCTGGAGTTCCTGCTCGCTTCAGTGGCAGTGGGTCTGGGACATCTTTCTCTTTCACAATCAACAGCATGGAGGCTGAAGATGTTGCCACTTATTACTGTCAGCAAAGGAGTAGTTACCCACCC |
| IGKV4-92*01_NZB | NZB/BlNJ | Musmus IGKV4-92*01 F | 100 | 291 | GAAATGGTTCTCACCCAGTCTCCAGTATCCATAACTGCATCTCGAGGGGAGAAGGTCACCATCACCTGCCGTGCCAGCTCAAGTATAAGTTCCAATTACTTACACTGGTACCAGCAGAAGCCAGGATCCTCCCCTAAACTTTTGATTTATAGGACATCCATCCTGGCATCTGGAGTCCTAGACAGCTTCAGTGGCAGTGGGTCTGAGAGCTCTTACACTCTGACAATCAGCTGCATGCAGGACGAAGTTGCTGCCACTTACTATTGTCAGCAGGGGAGTAGTAGCCCACCA |

| | | | | | |
|---|---|---|---|---|---|
| IGKV11-125*01_S8154_NZB | NZB/BlNJ | Musmus IGKV11-106*02 F | 95.34 | 287 | GATGTTCAAATGACCCAGTCTCCATCCTCCCTGTCTGCATCTTTGGGAGAAAGAGTCTCCCTGACCTGCCAGGCAAGTCAGAGCATTAGCAATTATTTAAACTGGTATCAGCAAACACTAGGGAAAGCTGCTAGGCTCTTGATCTATGGTGCAAGCAAATTGGAAGATGGGGTCCCTTCAAGGTTCAGTGGAACTGGATATGGGACAGATTTCACTTTCACCATCAGCAGCCAGGAGGAAGAAGATGTGGCAACTTATTTCTGTCTACAGCATAGGTATCTCCCTCC |
| IGKV3-1*01_NZB | NZB/BlNJ | Musmus IGKV3-1*01 F | 100 | 299 | GACATTGTGCTCACCCAATCTCCAGCTTCTTTGGCTGTGTCTCTAGGGCAGAGAGCCACCATCTCCTGCAGAGCCAGTGAAAGTGTTGAATATTATGGCACAAGTTTAATGCAGTGGTACCAACAGAAACCAGGACAGCCACCCAAACTCCTCATCTATGCTGCATCCAACGTAGAATCTGGGGTCCCTGCCAGGTTTAGTGGCAGTGGGTCTGGGACAGACTTCAGCCTCAACATCCATCCTGTGGAGGAGGATGATATTGCAATGTATTTCTGTCAGCAAAGTAGGAAGGTTCCTTC |
| IGKV4-52*01_S5019_NZB | NZB/BlNJ | Musmus IGKV4-52*01 ORF, or Musmus IGKV4-54*01 ORF | 100 | 286 | CAAATTGTTCTCACCCAGTCTCCAGCAATCCTGTCTGCATCTCCAGGGGAGAAGGTCACCATGACCTGCAGTGCCAGCTCAAGTGTAAGTTACATGTACAGGTACCAGCAGAAGCCAGGATCCTCACCCAAACCCTGGATTTATGGCACATCCAACCTGGCTTCTGGAGTCCCTGCTCGCTTCAGTGGCAGTGGATCTGGGACCTCTTATTCTCTCACAATCAGCAGCATGGAGGCTGAAGATGCTGCCACTTATTACTGCCAGCAATATCATAGTTACCCACTCA |
| IGKV4-61*01_S2955_NZB | NZB/BlNJ | Musmus IGKV4-61*01 F | 100 | 286 | CAAATTGTTCTCACCCAGTCTCCAGCAATCATGTCTGCATCTCCAGGGGAGAAGGTCACCATATCCTGCAGTGCCAGCTCAAGTGTAAGTTACATGTACTGGTACCAGCAGAAGCCAGGATCCTCCCCCAAACCCTGGATTTATCGCACATCCAACCTGGCTTCTGGAGTCCCTGCTCGCTTCAGTGGCAGTGGGTCTGGGACCTCTTACTCTCTCACAATCAGCAGCATGGAGGCTGAAGATGCTGCCACTTATTACTGCCAGCAGTATCATAGTTACCCACTCA |
| IGKV12-38*01_NZB | NZB/BlNJ | Musmus IGKV12-38*01 F | 100 | 287 | GACATCCAGATGACTCAGTCTCCAGCCTCCCTGGCTGCATCTGTGGGAGAAACTGTCACCATCACATGTCGAGCAAGTGAGAACATTTACTACAGTTTAGCATGGTATCAGCAGAAGCAAGGGAAATCTCCTCAGCTCCTGATCTATAATGCAAACAGCTTGGAAGATGGTGTCCCATCGAGGTTCAGTGGCAGTGGATCTGGGACACAGTATTCTATGAAGATCAACAGCATGCAGCCTGAAGATACCGCAACTTATTTCTGTAAACAGGCTTATGACGTTCCTCC |
| IGKV14-111*01_S2608_NZB | NZB/BlNJ | Musmus IGKV14-111*01 F | 97.85 | 287 | GACATCAAGATGACCCAGTCTCCATCTTCCATGTATGCATCTCTAGGAGAGAGAGTCACTATCACTTGCAAGGCGAGTCAGGACATTAATAGCTATTTACGCTGGTACCAGCAGAAACCAGGAAAATCTCCTAAGACCCTGATCTATGGTGCAAACAGCTTGGTAGATGGGGTCCCATCAAGGTTCAGTGGCAGTGGATCTGGGCAAGATTATTCTCTCACCATCAGCAGTCTGGAGTATGAAGATATGGGAATTTATTATTGTCTACAGTATGATGAGTTTCCTCC |
| IGKV4-50*01_S0901_NZB | NZB/BlNJ | Musmus IGKV4-50*01 F | 100 | 286 | GAAAATGTGCTCACCCAGTCTCCAGCAATCATGTCTGCATCTCTAGGGGAGAAGGTCACCATGAGCTGCAGGGCCAGCTCAAGTGTAAATTACATGTACTGGTACCAGCAGAAGTCAGATGCCTCCCCCAAACTATGGATTTATTACACATCCAACCTGGCTCCTGGAGTCCCAGCTCGCTTCAGTGGCAGTGGGTCTGGGAACTCTTATTCTCTCACAATCAGCAGCATGGAGGGTGAAGATGCTGCCACTTATTACTGCCAGCAGTTTACTAGTTCCCCATACA |
| IGKV4-56*01_NZB | NZB/BlNJ | Musmus IGKV4-56*01 P | 100 | 286 | CAAATTGTTCTCACCCAGTCTCCAGCAATCATGTCTGCATCTCCAGGGCAGAAAGTCACCATAACCTGCAGTGCCATCTCAAGTGTAAATTACATGCACTGGTACCAGCAGAAGCCAGGATCCTCCCCCAAACTCTGGATTTATGCAACATCCAAACTGGCTCTTGGAGTCCCTGCTTGCTTCAGTGGCAGTGGGTCTGGGACCTCTTACTCTCTCACAATCAGCAGCATGGTGGCTGAAGATGCCACCTCTTATTTCTGTCATCAGTGGAGTAGTTACCCACCCA |

| | | | | | |
|---|---|---|---|---|---|
| IGKV4-73*01_S9912_NZB | NZB/BlNJ | Musmus IGKV4-73*01 ORF | 100 | 292 | CAAATTGTTCTCACCCAGTCTCCAGCAATCATGTCTGCATCTCCTGGGGAACGGGTCACCATGACCTGCAGTGCCAGCTCAAGTGTAAGTTCCAGCTACTTGTACTGGTACCAGCAGAAGTCAGGATCCTCCCCAAAACTCTGGATTTATAGCATATCCAACCTGGCTTCTGGAGTCCCAGCTCGCTTCAGTGGCAGTGGGTCTGGGACCTCTTACTCTCTCACAATCAACAGCATGGAGGCTGAAGATGCTGCCACTTATTACTGCCAGCAGTGGAGTAGTAACCCGCTCA |
| IGKV4-81*01_NZB | NZB/BlNJ | Musmus IGKV4-81*01 F | 100 | 292 | GAAAATGTGCTGACCCAGTCTCCAGCAATCATGGCTGCATCTCCAGGGGAGAAGGTCACCATGACCTGCAGTGCCAGCTCAAGTGTAAGTTCTAGTAACTTGCACTGGTACCAGCAGAAGTCAGGCACTTCTACCAAATTCTGGATTTATAGGACATCCAACCTGGCTTCAGAAGTCCCAGCTCCCTTCAGTGGCAGTGGGTCTGGGACCTCTTACTCTCTTACAATCAGCAGCGTGGAGGCCGAAGATGCTGCCACTTATTACTGCCAGCAGTGGAGTGGTTACCCACCCA |
| IGKV5-37*01_NZB | NZB/BlNJ | Musmus IGKV5-37*01 F | 100 | 287 | GACATCCTGCTGACCCAGTCTCCAGCCACCCTGTCTGTGACTCCAGGAGAAACAGTCAGTCTTTCCTGTAGGGCCAGCCAGAGTATTTACAAGAACCTACACTGGTATCAACAGAAATCACATCGGTCTCCAAGGCTTCTCATCAAGTATGCTTCTGATTCCATCTCTGGGATCCCCTCCAGGTTCACTGGCAGTGGATCAGGGACAGATTACACTCTCAGTATCAACAGTGTGAAGCCCGAAGATGAAGGAATATATTACTGTCTTCAAGGTTACAGCACACCTTC |
| IGKV6-14*01_NZB | NZB/BlNJ | Musmus IGKV6-14*01 F | 100 | 287 | GACATTGTGATGACCCAGTCTCAAAAATTCATGTCCACATCAGTAGGAGACAGGGTCAGCATCACCTGCAAGGCCAGTCAGAATGTTCGTACTGCTGTAGCCTGGTATCAACAGAAACCAGGGCAGTCTCCTAAAGCACTGATTTACTTGGCATCCAACCGGCACACTGGAGTCCCTGATCGCTTCACAGGCAGTGGATCTGGGACAGATTTCACTCTCACCATTAGCAATGTGCAATCTGAAGACCTGGCAGATTATTTCTGTCTGCAACATTGGAATTATCCTCT |
| IGKV8-26*01_NZB | NZB/BlNJ | Musmus IGKV8-26*01 ORF | 100 | 311 | GACATTGTGATGACCCAGTCTCCATCCTCCCTGGCTGTGACAGCAGGAGAGAAGGTCACTATGAGCTGCAAGTCCAGTCAGAGTCTTTTGTGGAGTGTAAACCAAAAGAACTACTTGTCCTGGTACCAGCAGAAACAAAGGCAGCCTCCTAAACTGCTTATCTATGGGGCATCCATTAGAGAATCTTGGGTCCCTGATCGGTTCACAGGAAGTGGATCTGGGACAGACTTCACTCTCACCATTAGCAATGTGCATGCTGAAGACCTAGCAGTTTATTACTGTCAACACAATCATGGCAGCTTTCTCCCCCC |
| IGKV9-123*01_S4862_NZB | NZB/BlNJ | Musmus IGKV9-123*01 F | 89.61 | 285 | GACATCCAGATGACCCAGTCTCCATCCTCCATGTCTGCCTCTCTGGGAGAAAGAGTCAGTCTCACTTGCCGGGCCAGTCAGGGCATTAACGGTAATTTACACTGGTTTCAGCAGAAGTCAGGTGGAACTCTTAAACGCCTGATCTACTCCACGTCCAATTTAGATTCTGGTGTTCCATCAAGGTTCAGTGGCAGCGGGTCTGGGTCAGATTATTCTCTCACCATCAGCAGCCTGGAGTCTGAAGATTTTGCAATCTATTACTATCTACAGTATGATGAACATCCT |
| IGKV3-9*01_NZB | NZB/BlNJ | Musmus IGKV3-9*01 F | 100 | 299 | GACATTGTGCTGACCCAATCTCCAGCTTCTTTGGCTGTGTCTCTAGGACAGAGGGCCACCATATCCTGCCAAGCCAGCGAAAGTGTCAGTTTTGCTGGTACAAGTTTAATGCACTGGTACCAACAGAAACCAGGACAGCCACCCAAACTCCTCATCTATCGTGCATCCAACCTAGAATCTGGAGTCCCTGCCAGGTTCAGTGGCAGTGGGTCTGAGTCAGACTTCACTCTCACCATCGATCCTGTGGAGGAAGATGATGCTGCAATGTATTACTGTATGCAAAGTATGGAAGATCCTCC |

| | | | | | |
|---|---|---|---|---|---|
| IGKV4-60*01_NZB | NZB/BlNJ | Musmus IGKV4-60*01 P | 100 | 292 | GAAATTGTGCTCACCCAGTCTCCAGCACTCATGGCTGCATCTCCAGGGGAGAAGGTCACCATCACCTGCAGTGTCAGCTCAAGTATAAGTTCCAGCTACTTGCACTGGTACCAGCAGAAGTCAGGAATCTCCCCCAAACCCTGGATTTATGGCACATCCAACCTGGCTTCTGGAGTCCCTGCTCGCTTCAGTGGCAGTGGATCTGGGACCTCTTACTCTCTCACAATCAGCAGCATGGAGGCTGAAGATGCTGCCACTTATTACTGTCAACAGTGGAGTAGTTCCCCACCCA |
| IGKV4-73*01_NZB | NZB/BlNJ | Musmus IGKV4-73*01 ORF | 100 | 292 | CAAATTGTTCTCACCCAGTCTCCAGCAATCATGTCTGCATCTCCTGGGGAACGGGTCACCATGACCTGCAGTGCCAGCTCAAGTGTAAGTTCCAGCTACTTGTACTGGTACCAGCAGAAGTCAGGATCCTCCCCAAAACTCTGGATTTATAGCATATCCAACCTGGCTTCTGGAGTCCCAGCTCGCTTCAGTGGCAGTGGGTCTGGGACCTCTTACTCTCTCACAATCAACAGCATGGAGGCTGAAGATGCTGCCACTTATTACTGCCAGCAGTGGAGTAGTAACCCACCCA |
| IGKV4-74*01_S8852_PWD | PWD/PhJ | Musmus IGKV4-74*01 F | 94.33 | 292 | CAAATTGTTCTCACCCAGTCTCCAGCAATCATGTCTGCATCTCCAGGGGAGAAGGTCACCATGACCTGCAGGGCCAGCTCAAGTGTACGTTCCAGTTACTTGCACTGGTACCAGCAGAAGCCAGGATCTTCCCCCAAACTCTGGATTTATAGAACATCCAACCTGGCTTCAGGAGTCCCAGCTCGCTTCAGTGGCAGTGGGTCTGGGACCTCTTACTCTCTCACAATCAGCAGTGTGGAGGCTGAGGATGCTGCCACTTATTACTGCCAGCAGTATGATAGTTCCCCATCCA |
| IGKV6-15*01_S9094_PWD | PWD/PhJ | Musmus IGKV6-15*01 F | 96.06 | 287 | GACATTGTGATGACCCAGTCTCAAAAATTCATGTCCACATCAGTAGGAGACAGGGTCAGCGTCACCTGCAAGGCCAGTCAGTATGTGGGTACTTATGTAGCCTGGTATCAACAGAAACCAGGACAATCTCCTAAAGCACTGATTTACTCGGCATCCACCCGGCACACTGGAGTCCCTGATCGCTTCACAGGCAGTGGATCTGGGACAGATTTCACTCTCACCATTAGCAATGTGCAGTCTGAAGACTTGGCAGAGTATTTCTGTGAGCAATACAGCAGCTCTCCTCT |
| IGKV6-13*01_S1558_PWD | PWD/PhJ | Musmus IGKV6-13*01 F | 99.64 | 285 | GACATTGTGATGACCCAGTCTCAAAAATTCATGTCCACATCAGTAGGAGACAGGGTCAGCGTCACCTGCAAGGCCAGTCAGAATGTGGGTACTGCTGTAGCCTGGTATCAACAGAAACCAGGACAATCTCCTAAACTACTGATTTACTCGGCATCCAATCGGTACACTGGAGTCCCTGATCGCTTCACAGGCAGTGGATCTGGGACAGATTTCACTCTCACCATCAGCAATATGCAGTCTGAAGACCTGGCAGATTATTTCTGCCAGCAATATAGCAGCTATCCT |
| IGKV6-15*01_S1998_PWD | PWD/PhJ | Musmus IGKV6-15*01 F | 96.42 | 287 | GACATTGTGATGACCCAGTCTCAAAAATTCATGTCCACATCAGTAGGAGACAGGGTCAGCGTCACCTGCAAGGCCAGTCAGAATGTGGGTACTTATGTAGCCTGGTATCAACAGAAACCAGGGCAATCTCCTAAACTACTGATTTACTCGGCATCCAATCGGCACACTGGATTCCCTGATCGCTTCACAGGCAGTGGATCTGGGACAGATTTCACTCTCACCATCAGCAATGTGCAGTCTGAAGACTTGGCAGAGTATTTCTGTGAGCAATATAGCAGCTATCCTCT |
| IGKV6-20*01_S7621_PWD | PWD/PhJ | Musmus IGKV6-20*01 F | 99.64 | 287 | AACATTGTAATGACCCAATCTCCCAAATCCATGTCCATGTCAGTAGGAGAGAGGGTCACCTTGAGCTGCAAGGCCAGTGAGAATGTGGGTACTTATGTATCCTGGTATCAACAGAAACCAGAGCAGTCTCCTAAACTGCTGATATACGGGGCATCCAACCGGTACACTGGGGTCCCCGATCGCTTCACAGGCAGTGGATCTGCAACAGATTTCACTCTGACCATCAGCAGTGTGCAGGCTGAAGACCTTGCAGATTATCACTGTGCACAGAGTTACAGCTATCCTCC |
| IGKV6-14*01_S4666_PWD | PWD/PhJ | Musmus IGKV6-14*01 F | 96.42 | 287 | GACATTGTGATGACCCAGTCTCAAAAATTCATGTCCACATCAGTAGAAGACAGGGTCAGCATCACCTGCAAGGCCAGTCAGAATGTAGGTACTGCTGTAGCCTGGTATCAACAGAAACCAGGGCAGTCTCCTAAACTACTGATTTACTTGGCATCCAATCGGCACACTGGAGTCCCTGATCGCTTCACAGGCAGTGGATCTGGGACAGATTTCACTCTCACCATCAGCAATGTGCAGTCTGAAGACCTGGCAGATTATTTTTGTCTGCAACATTGGAGTTATCCTCT |

| | | | | | |
|---|---|---|---|---|---|
| IGKV8-28*01_S2077_PWD | PWD/PhJ | Musmus IGKV8-28*01 F | 99.33 | 305 | GACATTGTGATGACACAGTCTCCATCCTCCCTGAGTGTGTCAGCAGGAGAGAAGGTCACTATGAGCTGCAAGTCCAGTCAGAGTCTGTTAAACAGTGGAAACCAAAAGAACTACTTGGCCTGGTACCAGCAGAAACCAGGGCAGCCTCCTAAACTGTTGATCTACGGGGCATCCACTAGGGAATCTGGGGTCCCTGATCGCTTCACAGGCAGTGGATCTGGAACCGATTTCACTCTTACCATCAGCAGTGTGCAGGCTGAAGACCTGGCAGTTTATTACTGTCAGAATGATTATAGTTATCCTCC |
| IGKV2-137*01_S9859_PWD | PWD/PhJ | Musmus IGKV2-137*01 F | 94.56 | 302 | GATATTGTGATGACTCAGGCTGCACCCTCTGTACCTGTCACTCCTGGAGAGTCAGTATCCATCTCCTGCAGGTCTAGTAAGAGTCTCCTGCACAGTAGTGGCAAGCATAGGTTGTATTGGTTCCTACAGAGGCCAGGCCAGTCTCCTCAGCGCCTGATATATTATATGTCCAACCTTGCCTCAGGAGTCCCAGACAGGTTCAGTGGCAGTGGGTCAGGAACTGATTTCACACTGAGAATCAGTAGAGTGGAGGCTGAGGATGTGGGTGTTTATTACTGTATGCAAAGTCTAGAATATCCATT |
| IGKV6-25*01_S2065_PWD | PWD/PhJ | Musmus IGKV6-13*01 F, or Musmus IGKV6-14*01 F | 92.11 | 287 | GACATTGTGATGACCCAGTCACAAAAATTCATGTCCACATCAGTAGGAGAGAGGGTCAGCATCAGCTGCAAGGCCAGTCAGAATGTGGGTAATATTATAGCCTGGTATCAACAGAAACCAGGGCAGTCTCCTAAACCACTGATTTACTTGGCTTCCAATCGGCACACTGGAGTCCCTGATCGCTTCACAGGCAGTGGATCTGGGACAGATTTCACTCTGACCATCAGCAGTGTGCAGGCTGAAGACCTGGCAGATTATTTCTGTCAGCAACATTATAGCACTCCTCC |
| IGKV10-94*01_Mus_spretus_S4447_PWD | PWD/PhJ | Musspr IGKV10-94*01 F | 97.49 | 287 | GATATCCAGATGACACAGACTACATCCTCCCTGTCTGCCTCTCTAGGAGACAGAGTCACCATCAGTTGCAGGGCAAGTCAGGATATTAGCAATTATTTAAACTGGTATCAGCAGAAACCAGATGGAACTGTTAAACCCCTGATCTATTATGCGTCAAGTTTACACTCAGGAGTCCCATCAAGGTTCAGTGGCAGTGGGTCTGGGACAGATTATTCTCTCACCATCAGCAACCTGGAACCTGAAGATATTGCCACTTATTATTGTCAGCAGTATAGTAAGCTTCCTCC |
| IGKV12-40*01_S1894_PWD | PWD/PhJ | Musmus IGKV12-40*01 P | 98.57 | 287 | GACATCCAGATGACTCAGTCTCCAGCCTCCCTATCTGCATCTGTGGGAGAAACTGTCACCATCACATGTCAAGCAAGTGAGAATATTGCCAGTGATTTAGCATGGTATCAGCAGAAACAGGGAAAATCTCCTCAGCTCCTGGTCTATGATGCGAGAAACTTAGCAGATGGTGTGCCATCAAGGTTCAGTGGCAGTGGATCAGGCACACAGTATTCTCTCAATATCAACAGCCTGCAGTCTGAAGATGTTGCGAGATATTACTGTCAACATTATTATGGTACTCCTCC |
| IGKV4-55*01_S7109_PWD | PWD/PhJ | Musmus IGKV4-55*01 F | 97.46 | 286 | CAAATTGTTCTCACCCAGTCTCCAGCAATCATGTCTGCATCTCCAGGGGAGAAGGTCACCATGACCTGCAGTGCCAGCTCAAGTGTAAGTTACATGTACTGGTACCAGCAGAAGCCAGGATCCTCCCCCAGACTCTGGATTTATGACACATCCAACCTGGCTTCTGGAGTCCCCGCTCGCTTCAGTGGCAGTGGGTCTGGGACCTCTTATTCTCTCACAATCAGCAGCATGGAGGCTGAAGATGCTGCCACTTATTACTGCCAGCAGTGGAGTAGTAACCAACCCA |
| IGKV4-80*01_S9832_PWD | PWD/PhJ | Musmus IGKV4-80*01 F | 99.28 | 282 | CAAATTGTTCTCACCCAGTCTCCAGCAATCATGTCTGCATCTCTAGGGGAGGAGATCACCCTAACTTGCAGTGCCAGCTCGAGTGTAAGTTACATGCACTGGTTCCAGCAGAAGTCAGGCACTTCTCCCAAACTCTTGATTTATAGCACATCCAACCTGGCTTCTGGAGTCCCTTCTCGCTTCAGTGGCAGTGGGTCTGGGACCTTTTATTCTCTCACAATCAGCAGTGTGGAGGCTGAAGATGCTGCCGATTATTACTGCCATCAGTGGAGTAGTTATCCA |
| IGKV15-103*01_S9355_PWD | PWD/PhJ | Musmus IGKV15-103*01 ORF | 94.62 | 287 | GACATCCAAATGAACCAGTCTCCATCCAGTCTGTCTGCATCCCTCGGAGACACAATTACCATCACTTGCCGTGCCAGTCAGGGCATTAGTATTTGGTTAAGCTGGTACCAGCAGAAACCAGGAAATATTCCTAAACTATTGATCTATAAGGCTTCCAACTTGCACACAGGAGTCCCACCAAGGTTTAGTGGCAGTGGATCTGGTACAGATTTCACATTAACTATCAGCAGCCTACAGCCTGAAGACATTGCCACTTACTACTGTCTACAGAGTCAAAGTTATCCTCT |

| | | | | | |
|---|---|---|---|---|---|
| IGKV8-30*01_S8873_PWD | PWD/PhJ | Musmus IGKV8-30*01 F | 96.61 | 295 | GACATTGTGATGTCACAGTCTCCATCCTCCCTGGCTGTGTCAGTTGGAGAGAAGGTTACTATGAGCTGCAAATCCAGTCAGAGCCTTTTATATAGTAGCAATCAAAAGAACTACTTGGCCTGGTACCAGCAGAAACCTGGGCAGTCTCCTAAACTGTTGATCTACTGGGCATCCACTAGGGAATCTGGGGTCCCTGATCGCTTCACAGGCAGTGGATCTGGGACAGATTTCACTCTCACCATCAGCAGTGTGAAGGCTGAAGACCTGGCCGTTTATTACTGCAAGCAATCTTATG |
| IGKV12-44*01_S2900_PWD | PWD/PhJ | Musmus IGKV12-44*01 F | 96.77 | 287 | GACATCCAGATGACTCAGTCTCCAGCTTCCCTATCTGCATCTGTGGGAGAAACTGTCACCATCACATGTCGAGCAACTGGGAATATTTACAGTAATTTAGCATGGTATCAGCAGAAACAGGGAAAATCTCCTCAGCTCCTGGTCTATAATGCAAAAACCTTAGCAGAAGGTGTGCCATCAAGGTTCAGTGGCAGTGGATCAGGAACACAATATTCTCTCAAGATCAACAGCCTGCAGCCTGAAGATTTTGGGAGTTATTACTGTCAACATTATTATGGTACTCCTCC |
| IGKV6-13*01_S4184_PWD | PWD/PhJ | Musmus IGKV6-13*01 F | 89.61 | 285 | GACATTGTGATGACCCAGTCACAAAAATTCATGTCCACATCAGTAGGAGAGAGGGTCAGCATCAGCTGCAAGGCCAGTCAGAATGTGGGTAATATTATAGCCTGGTATCAACAGAAACCAGGACAGTCTCCTAAAGCACTGATTTACTGGGCTTCCAATCGGCACACTGGAGTCCCTGATCGCTTCACAGGCAGTGGATCTGGGACAGATTTCACTCTGACCATCAGCAGTGTGCAGGCTGAAGACCTTGCAGATTATCACTGTGGACAGAGTTACAGCTATCCT |
| IGKV1-122*01_S7970_PWD | PWD/PhJ | Musmus IGKV1-122*01 F | 98.64 | 302 | GATGCTGTGATGACCCAAACTCCACTCTCCCTGCCTGTCAGTCTTGGAGATCAAGCCTCCATCTCTTGCAGGTCTAGTCAGAGCCTTGAAAACAGTAATGGAAACACCTATTTGAACTGGTACCTCCAGAAACCAGGCCAGTCTCCACAGCTCCTGATCTACAGGGTTTCCAACCGATTTTCTGGGGTCCTAGACAGGTTCAGTGGCAGTGGTTCAGGGACAGATTTCACACTCAAGATCAGCAGAGTGGAGGCTGAGGATTTGGGAGTTTATTTCTGCCTCCAAGTTACACATGTCCCTCC |
| IGKV4-62*01_S7295_PWD | PWD/PhJ | Musmus IGKV4-62*01 ORF, or Musmus IGKV4-68*01 F | 96.74 | 286 | CAAATTGTTCTCACCCAGTCTCCAGCAATCATGTCTGCATCTCCAGGGGAGAAGGTCACCATGACCTGCAGTGCCAGCTCAAGTGTAAGTTACATGTACTGGTACCAGCAGAAGCCAGGATCTTCCCCCAAACCCTGGATTTATGACACATCCAACCTGGCTTCTGGAGTCCCTGCTCGCTTCAGTGGCAGTGGGTCTGGGACCTCTTACTCTCTCACAATCAGCCGCATGGAGGCTGAAGATGCTGCCACTTATTACTGCCAGCAGTACAGTGGTTACCCATACA |
| IGKV1-135*01_S0376_PWD | PWD/PhJ | Musmus IGKV1-135*01 F | 95.92 | 302 | GATGTTGTGATGACTCAGACCCCACTCACTTTGTCGGTTACCATTGGACAACCAGCTTCCATCTCTTGCAAGTCAAGTCAGAGCCTCTTACATAGTAATGGAAAGACATATTTGAATTGGTTATTACAGAGGCCAGGCCAGTCTCCAAAGCTCCTAATCTATCTGGTGTCTAAACTGGAATCTGGAGTCCCTGACAGGTTCAGTGGCAGTGGATCAGGGACAGATTTCACACTGAAAATCAGCAGAGTGGAGGCTGAGGATTTGGGAGTTTATTACTGCTTGCAAGCTACACATTTTCCTCA |
| IGKV12-46*01_S2796_PWD | PWD/PhJ | Musmus IGKV12-46*01 F | 98.21 | 287 | GACATCCAGATGACTCAGTCTCCAGCCTCCCTATCTGCATCTGTAGGAGAAACTGTCACCATCACATGTCGAGCAAGTGAGAATATTTACAGTTATTTAGCATGGTATCAGCAGAAACAGGGAAAATCTCCTCAGCTCCTGGTCTATGCTGCAACAAACTTAGCAGATGGTGTGCCATCAAGGTTCAGTGGCAGTGGATCAGGCACACAGTATTCCCTCAAGATCAACAGCCTGCAGCCTGAAGATTTTGGGAGTTATTACTGTCAACATTTTTGGGTACTCCTCC |
| IGKV14-126*01_PWD | PWD/PhJ | Musmus IGKV14-126*01 F | 100 | 287 | GACATCAAGATGACCCAGTCTCCATCCTCCATGTATGCATCGCTGGGAGAGAGAGTCACTATCACTTGCAAGGCGAGTCAGGACATTAAAAGCTATTTAAGCTGGTACCAGCAGAAACCATGGAAATCTCCTAAGACCCTGATCTATTATGCAACAAGCTTGGCAGATGGGGTCCCATCAAGATTCAGTGGCAGTGGATCTGGGCAAGATTATTCTCTAACCATCAGCAGCCTGGAGTCTGACGATACAGCAACTTATTACTGTCTACAGCATGGTGAGAGCCCTCC |

| | | | | | |
|---|---|---|---|---|---|
| IGKV1-117*02_S1951_PWD | PWD/PhJ | Musmus IGKV1-117*02 F | 99.66 | 302 | GATGTTGTGATGACCCAAACTCCACTCTCCCTGCCTGTCAGTCTTGGAGATCAAGCCTCCATCTCTTGCAGATCTAGTCAGAGCATTGTACATAGTAATGGAAACACCTATTTAGAATGGTACCTGCAGAAACCAGGCCAGTCTCCAAAGCTCCTGATCTACAAAGTTTCCAACCGATTGTCTGGGGTCCCAGACAGGTTCAGTGGCAGTGGATCAGGGACAGACTTCACACTCAAGATCAGCAGAGTGGAGGCTGAGGATCTGGGAGTTTATTACTGCTTTCAAGGTTCACATGTTCCTCC |
| IGKV8-16*01_PWD | PWD/PhJ | Musmus IGKV8-16*01 F | 100 | 305 | GAAATTGTGTTGACCCAGTCTATACCATCCCTGACTGTGTCAGCAGGAGAGAGGGTCACTATCAGCTGCAAATCCAATCAGAATCTTTTATGGAGTGGAAACCAAAGGTACTGTTTGGTCTGGCACCAGTGGAAACCAGGGCAAACTCCTACACCGTTGATCACCTGGACATCTGATAGGTACTCTGGAGTCCCTGATCGTTTCATAGGCAGTGGATCTGTGACAGATTTCACTCTGACCATCAGCAGTGTGCAGGCTGAAGATGTGGCAGTTTATTTCTGTCAGCAGCATTTACACATTCCTCC |
| IGKV10-94*01_S9482_PWD | PWD/PhJ | Musmus IGKV10-94*01 F | 98.21 | 287 | GATATCCAGATGACACAGAGTACATCCACCCTGTCTGCCTCTCTGGGAGACAGAGTCACCATCAGTTGCAGTGCAAGTCAGGGCATTAGCAATTATTTAAACTGGTATCAGCAGAAACCAGATGGAACTGTTAAACTCCTGATCTATTATACATCAAGTTTACACTCAGGAGTCCCATCAAGGTTCAGTGGCAGTGGGTCTGGAACAGATTATTCTCTCACCATCAGCAACCTGGAACCTGAAGATATTGCCACTTACTATTGTCAGCAGTATAGTAATTTTCCTCC |
| IGKV1-135*01_S6159_PWD | PWD/PhJ | Musmus IGKV1-135*01 F | 95.58 | 301 | GATGTTGTGATGACTCAGACCCCACTCACTTTGTCGGTTACCATTGGACAACCAGCCTCCATCTCTTGCAAGTCAAGTCAGAGCCTCTTATATAGTGATGGAAAGACATATTTGGATTGGTTATTACAGAGGCCAGGTCAGTCTCCAAAGAGCCTAATCTATCTGGTGTCTAAACTGGAATCTGGAGTCCCTGACAGGTTCAGTGGCAGTGGATCAGGGACAGATTTCATACTGAAAATCAGCAGAGTGGAGGCTGAGGATTTGGGAGTTTATTACTGCGTGCAAGGTACACATTTTCCTC |
| IGKV14-111*01_S2944_PWD | PWD/PhJ | Musmus IGKV14-111*01 F | 98.21 | 287 | GACATCAAGATGACCCAGTCTCCATCTTCCATGTATGCATCTCTAGGAGAGAGAGTCACTATCACTTGCAAGGCGAGTCAGGACATTAATAGCTATTTAAGCTGGTACCAGCAGAAACCAGGGAAATCTCCTAAGACCCTGATCTATTATGCAAACAGCTTGGTAGATGGGGTCCCATCAAGGTTCAGTGGCAGTGGATCTGGGCAAGATTATTCTCTCACCATCAGCAGCCTGGAGTATGAAGATACGGGAATTTATTATTGTCTACAGTATGATGAGTTTCCTCC |
| IGKV4-68*01_S8814_PWD | PWD/PhJ | Musmus IGKV4-68*01 F | 96.74 | 286 | CAAATTGTTCTCACCCAGTCTCCAGCAATCATGTCTGCATCTCCAGGGGAGAAGGTCACCATGACCTGCAGTGCCAGCTCAAGTGTAAGTTATATGCACTGGTACCAGCAGAAGCCAGGATCCTCACCCAAACCCTGGATTTATCGCACATCCAACCTGGCTTCTGGAGTCCCTGCTCGCTTCAGTGGCAGTGGGTCTGGGACCTCTTACTCTCTCACAATTACCAGCATTGAGGCTGAAGATGCTGCCACTTATTACTGCCAGCAGTGGAGTAGTAACCCACTCA |
| IGKV3-2*01_PWD | PWD/PhJ | Musmus IGKV3-2*01 F | 100 | 299 | GACATTGTGCTGACCCAATCTCCAGCTTCTTTGGCTGTGTCTCTAGGGCAGAGGGCCACCATCTCCTGCAGAGCCAGCGAAAGTGTTGATAATTATGGCATTAGTTTTATGAACTGGTTCCAACAGAAACCAGGACAGCCACCCAAACTCCTCATCTATGCTGCATCCAACCAAGGATCCGGGGTCCCTGCCAGGTTTAGTGGCAGTGGGTCTGGGACAGACTTCAGCCTCAACATCCATCCTATGGAGGAGGATGATACTGCAATGTATTTCTGTCAGCAAAGTAAGGAGGTTCCTCC |
| IGKV12-44*01_PWD | PWD/PhJ | Musmus IGKV12-44*01 F | 100 | 287 | GACATCCAGATGACTCAGTCTCCAGCCTCCCTATCTGCATCTGTGGGAGAAACTGTCACCATCACATGTCGAGCAAGTGAGAATATTTACAGTTATTTAGCATGGTATCAGCAGAAACAGGGAAAATCTCCTCAGCTCCTGGTCTATAATGCAAAAACCTTAGCAGAAGGTGTGCCATCAAGGTTCAGTGGCAGTGGATCAGGCACACAGTTTTCTCTGAAGATCAACAGCCTGCAGCCTGAAGATTTTGGGAGTTATTACTGTCAACATCATTATGGTACTCCTCC |

| | | | | | |
|---|---|---|---|---|---|
| IGKV19-93*01_S9010_PWD | PWD/PhJ | Musmus IGKV19-93*01 F | 96.42 | 287 | GACATCCAGATGACACAGTCTCCATCCTCACTGTCTGCATCTCTGGGAGGCAAAGTCACCATCACTTGCAAGGCAAGCCAGGACATTAACAAGAATATAGCTTGGTACCAACACAAGCCTGGAAAAGGTCCTAGGCTGCTCATACGTTACACATCAATACTAGAGTCAGGCATCCCATCAAGGTTCAGTGGAAGTGGGTCTGGGAGAGATTATTCCTTCAGCATCAGCAACCTGGAGCCTGAAGATATTGCAACATATTACTGTCTACAGTATGATAATCTTCCACC |
| IGKV4-59*01_S5176_PWD | PWD/PhJ | Musmus IGKV4-59*01 F | 97.1 | 286 | CAAATTGTTCTCACCCAGTCTCCAGCAATCATGTCTGCATCTCCAGGGGAGAAGGTCACCATGACCTGCAGTGCCAGCTCAAGTGTAAGTTACATGTACTGGTACCAGCAGAAGCCAGGATCCTCCCCCAAAAAATGGATTTATGACACATCCAACCTGGCTTCTGGAGTCCCCGCTCGCTTCAGTGGCAGTGGGTCTGGGACCTCTTATTCTCTCACAATCAGCAGCATGGAGGCTGAAGATGCTGCCACTTATTACTGCCAGCAGTGGAGTAGTTACCCATCCA |
| IGKV4-61*01_S1531_PWD | PWD/PhJ | Musmus IGKV4-61*01 F | 99.64 | 286 | CAAATTGTTCTCACCCAGTCTCCAGCAATCCTGTCTGCATCTCCAGGGGAGAAGGTCACCATATCCTGCAGTGCCAGCTCAAGTGTAAGTTACATGTACTGGTACCAGCAGAAGCCAGGATCCTCCCCCAAACCCTGGATTTATCGCACATCCAACCTGGCTTCTGGAGTCCCTGCTCGCTTCAGTGGCAGTGGGTCTGGGACCTCTTACTCTCTCACAATCAGCAGCATGGAGGCTGAAGATGCTGCCACTTATTACTGCCAGCAGTATCATAGTTACCCACCCA |
| IGKV5-48*01_S4867_PWD | PWD/PhJ | Musmus IGKV5-48*01 F | 99.64 | 287 | GACATCTTGCTGACTCAGTCTCCAGCCATCCTGTCTGTGAGTCCAGGAGAAAGAGTCAGTTTCTCCTGCAGGGCCAGTCAGAGCATTGGCACAAGCATACACTGGTATCAGCAAAGAACAAATGGTTCTCCAAGGCTTCTCATAAAGTATGCTTCTGAGTCTATCTCTGGGATCCCTTCCAGGTTTAGTGGCAGTGGATCAGGGTCAGATTTTACTCTTAGCATCAACAGTGTGGAGTCTGAAGATATTGCAGATTATTACTGTCAACAAAGTAATAGCTGGCCAAC |
| IGKV4-73*01_S3184_PWD | PWD/PhJ | Musmus IGKV4-73*01 ORF | 97.87 | 292 | CAAATTGTTCTCACCCAGTCTCCAGCAATCATGTCTGCATCTCCTGGGGAACGGGTCACCATGACCTGCAGTGCCAGCTCAAGTGTAAGTTCCAGCTACTTGTACTGGTACCAGCAGAAGCCAGGATCCTCCCCCAAACTCTGGATTTATAGCACATCCAACCTGGCTTCTGGAGTCCCTGCTCGCTTCAGTGGCAGTGGGTCTGGGACCTCTTATTCTCTCACAATCAGCAGCATGGAGGCTGAAGATGCTGCCACTTATTACTGCCAGCAGTGGAGTAGTAACCAACCCA |
| IGKV1-133*01_S9974_PWD | PWD/PhJ | Musmus IGKV1-133*01 F | 96.26 | 302 | GATGTTGTGATGACCCAGACTCCACTCACTTTGTCTGTTACCATTGGACAGCCAGCTTCCATCTCTTGCAAGTCAAGTCAGAGCCTCTTATATAGTGATGGAAAAACCTATTTGAATTGGTTATTACAGAGTCCAGGCCAGTCTCCAAAGCTCCTAATCTATCTGGTGTCTAAACTGGAATCTGGAGTCCCTGACAGATTCAGTGGCAGTGGATCAGGGACAGATTTTACACTGAAAATCAGCAGAGTGGAGGCTGAGGATTTGGGAGTTTATTACTGCGTGCAAGGTACACATTTCCCTCA |
| IGKV16-104*01_S2488_PWD | PWD/PhJ | Musmus IGKV14-87-2*01 F | 99.28 | 288 | GACATCCAAATGACCCAGTCCTCATCATTTCTCTCTGCATCTCTAGGAGATCATCTTACAATCAACTGCAGGGCCAGTAAGGATATTAACAAGTATTTTGCTTGGGTTCAACAGAAGCCAGGGAAGGCTCCAAGGATGTTGATTCATTTTGCTTCCACCTTGCTACCTGGGGTTCCAGAAAAGTTCAGTGGGAGTGGATCTGGGACAGATTTTTCTCTCACTATCAGAAACATAGAGTCCGAAGATATTGCAATGTATTACTGTCTACAGTATTCTGAGCATCCGTAC |
| IGKV4-86*01_S6393_PWD | PWD/PhJ | Musmus IGKV4-86*01 F | 94.93 | 281 | GAAATTGTGCTCACTCAGTCTCCAGCCATCATAGCTGCATCTCTGGGGCAAAAGGTCACCATGACCTGCAGTGCCAGCTCAAGTGTAAGTTACATGCACTGGTACCAGCAGAAGTCAGGCACCTCCCCCAAACTCTGGATTTACGGCACATCCAACCTGGCTTCTGGAGTCCCAGCTCGCTTCAGTGGCAGTGGGTCTGGGACCTCTTACTCTCTCACAATCAGCAGCGTGGAGGCTGAAGATGATGCAACTTATTACTGCCAGCAGTGGAATTACCCACT |

| | | | | | |
|---|---|---|---|---|---|
| IGKV16-104*01_S7081_PWD | PWD/PhJ | Musmus IGKV16-104*01 F | 98.21 | 288 | GATGTCCAGATAACCCAGTCTCCATCTTATCTTGCTGCATCTCCTGGAGAAACCATTACTATTAATTGCAGGGCAAGTAAGAGCATTAACAAATATTTAGCCTGGTATCAAGAGAAACCTGGGAAAACTAATAAGCTTCTTATCTACTCTGGATCCACCTTGCAATCTGGAATTCCATCAAGGTTCAGTGGCAGTGGATCTGGTACAGATTTTACTCTCACCATCAGTAGCTTGGAGCCTGAAGATTTTGCAATGTATCACTGTCAACAGCATAATGAATACCCGTAC |
| IGKV3-12*01_S8984_PWD | PWD/PhJ | Musmus IGKV3-12*01 F | 98.28 | 299 | GACATTGTGCTGACACAGTCTCCTGCTTCCTTAGCTGTATCTCTGGGGCAGAGGGCCACCATCTCCTGCAGGGCCAGCAAAAGTGTCAGTACATCTAGCTATAGTTACATGCACTGGTACCAACAGAAACCAGGACAGCCACCCAAACTCCTCATCTATTTTGCATCCAACCTAGAATCTGGGGTCCCTGCCAGGTTCAGTGGCAGTGGGTCTGGGACAGACTTCACCCTCAACATCCATCCTGTGGAGGAGGAGGATGCTGCAACATATTACTGTCAGCACAGTAGGGAGTTTCCTCC |
| IGKV1-110*01_S5755_PWD | PWD/PhJ | Musmus IGKV1-110*01 F | 97.96 | 301 | GATGTTTTGATGACCCAAACTCCACTCTCCCTGCCTGTCAGTCTTGGAGATCAAGCCTCCATCTCTTGCAGATCTAGTCAGAGCCTTGTACACAGTAATGGAAACACCTATTTATATTGGTACCTGCAGAAGCCAGGCCAGTCTCCAAAGCTCCTGATCTACAAAGTTTCCAACCGATTTTCTGGGGTCCCAGACAGGTTCAGTGGCAGTGGATCAGGGACAGATTTCACACTCAAGATCAGCAGAGTGGAGGCTGAGGATATGGGAGTTTATTACTGCTTTCAAGGTACACATGTTCCTC |
| IGKV4-53*01_S8918_PWD | PWD/PhJ | Musmus IGKV4-60*01 P | 98.23 | 292 | GAAATTGTGCTCACCCAGTCTCCAGCACTCATGGCTGCATCTCCAGGGGAGAAGGTCAGCATCACCTGCAGTGTCAGCTCAAGTATAAGTTCCAGCCACTTACACTGGTACCAGCAGAAGTCAGGAACCTCCCCCAAACTCTGGATTTATGGCACATCCAACCTGGCTTCTGGAGTCCCTGCTCGCTTCAGTGGCAGTGGATCTGGGACCTCTTACTCTCTCACAATCAGCAGCATGGAGGCTGAAGATGCTGCCACTTATTACTGTCAACAGTGGAGTAGTTACCCACTCA |
| IGKV5-45*01_S0430_PWD | PWD/PhJ | Musmus IGKV5-45*01 F | 99.64 | 287 | GATATTGTGCTAACTCAGTCTCCAGCCACCCTGTCTGTGACTCCAGGAGATAGAGTCAGTCTTTCCTGCAGGGCCAGCCAAAGTATTAGCAACTACCTACACTGGTATCAACAAAAATCACATGAGTCTCCAAGGCTTCTCATCAAGTATGCTTCCCAGTCCATCTCTGGGATCCCCTCCAGGTTCAGTGGCAGTGGATCAGGGACAGATTTCACTCTCAGTATCAACAGTGTGGAGACTGAAGATTTTGGAATGTATTTCTGTCAACAGAGTAACAGCTGGCCTCA |
| IGKV12-89*01_S6231_PWD | PWD/PhJ | Musmus IGKV12-89*01 F | 99.28 | 287 | GACATCCAGATGACTCAGTCTCCAGCTTCACTGTCTGCATCTGTGGGAGAAAGTGTCTCCATCACATGTGGAGCAAGTGAGAATATTTACGGTGCTTTAAATTGGTATCAGCGGAAACAGGGAAAATCTCCTCAGCTCCTGATCTATGGTGCAACCAACTTGGCAGATGGCATGTCATCGAGGTTCAGTGGCAGTGGATCTGGTAGACAGTATTCTCTCAAGATCAGTAGCCTGCATCCTGACGATGTTGCAACGTATTACTGTCAAAATGTGTTAAGTACCCCTCC |
| IGKV9-124*01_S5062_PWD | PWD/PhJ | Musmus IGKV9-124*01 F | 99.64 | 285 | GACATCCAGATGACCCAGTCTCCATCCTCCTTATCTGCCTCTCTGGGAGAAAGAGTCAGTCTCACTTGTCGGGCAAGTCAGGAAATTAGTGGTTACTTAAGCTGGCTTCAGCAGAAACCAGATGGAACTATTAAACGCCTGATCTACGCTGCATCCACTTTAGATTCTGGTGTCCCAAAAAGGTTCAGTGGCAGTAGGTCTGGGTCAGATTATTCTCTCACCATCAGCAGCCTTGAGTCTGAAGATTTTGCAGACTATTACTGTCTACAATATGCTAGTTATCCT |
| IGKV4-57-1*01_S1667_PWD | PWD/PhJ | Musmus IGKV4-57*01 F | 95.65 | 292 | GAAAATGTGCTCACCCAGTCTCCAGCACTCATGGCTGCATCTCCAGGGGAGAAGGTCACCATGACCTGCAGTGCCAGCTCAAGTGTAAGTTCCAGTTACTTGCACTGGTACCAGCAGAAGTCAGGTGCCTCCCCCAAACTCTGGATTTACGGCACATCCAACCTGGCTTCTGGAGTCCCAGCTCGCTTCAGTGGCAGTGGGTCTGGGACCTCTTACTCTCTCACAATCAGCAGCGTGGAGGCTGAAGATGATGCAACTTATTACTGCCAGCAGGCGTGGAATTACCCATTCA |

| | | | | | |
|---|---|---|---|---|---|
| IGKV4-57*01_S0701_PWD | PWD/PhJ | Musmus IGKV4-57-1*01 F | 93.62 | 286 | CAAATTGTTCTCACCCAGTCTCCAGCAATCATGGCTGCATCTCCAGGGGAGAAGTTCACCATAACCTGCAGTGCCAGCTCAAGTGTAAGTTACATGCACTGGTTCCAGCAGAAGCCAGGCACTTCTCCCAAACTCTGGATTTATAGGACATCGAACCTGGCTTCTGGAGTCCCTGCTCGCTTCAGTGGCAGTGGGTCTGGGACCTCTTATTCTCTCACAATCAGCAGCATGGATGCTGAAGATGCTGCCACTTATTACTGCCTGCAGCGGAGTAGTTACCCACCCA |
| IGKV3-10*01_S9898_PWD | PWD/PhJ | Musmus IGKV3-10*01 F, or Musmus IGKV3-5*01 F | 98.97 | 299 | GACATTGTGCTGACCCAATCTCCAGCTTCTTTGGCTGTGTCTCTAGGGCAGAGGGCCACCATATCCTGCAGAGCCAGTGAAAGTGTTGATAGTTATGGCAATAGTTTTATGCACTGGTACCAGCAGAAACCAGGACAGCCACCCAAACTCCTCATCTATCTTGCATCCAACCTAGAATCTGGGGTCCCTGCCAGGTTCAGTGGCAGTGGGTCTAGGACAGACTTCACCCTCACCATTAATCCTGTGGAGGCTGATGATGTTGCAACCTATTACTGTCAGCAAAATAATGAGGATCCTCC |
| IGKV12-38*01_S6950_PWD | PWD/PhJ | Musmus IGKV12-38*01 F | 98.57 | 287 | GACATCCAGATGACTCAGTCTCCAGCCTCCCTGGCTGCATCTGTGGGAGAAACCATCACCATCACATGTCAAGCAAGTGAGAACATTTACTACAGTTTAGCATGGTATCAGCAGAAGCAAGGGAAATCTCCTCAGCTCCTGATCTATAATGCAAACAGCTTGGAAGATGGTGTCCCATCGAGGTTCAGTGGCAGTGGATCTGGGACACAGTATTCTATGAAGATCAACAGCATGCAGCCTGAAGATACTGCAACTTATTTCTGTAAACAGGCTTATGACTTTCCTCC |
| IGKV4-81*01_S1013_PWD | PWD/PhJ | Musmus IGKV4-81*01 F | 96.1 | 292 | GAAAATGTGCTGACCCAGTATCCAGCAATCATGGCTGCATCTCCAGGGGAGAAGGTCACCATGACCTGCAGTGCCAGCTCAAGTGTAAGTTCTGATAACTTTCACTGGTACCAGCAGAAGTCAGGCACTTCTCCCAAACTCTGGATTTATAGGACATCCAAATTGGCTTCTGGAGTCCCAGCTCGCTTCAGTGGCAGTGGGTCTGGGACCTCTTACTCTCTTACAATCAGCAGCGTGGAGGCCGAAGATGCTGCCACTTATTACTGCCAGCAGTGGAGTGGTTACCCACTCA |
| IGKV10-96*03_S5777_PWD | PWD/PhJ | Musmus IGKV10-96*03 F | 99.28 | 287 | GATATCCAGATGACACAGACTACATCCTCCCTGTCTGCCTCTCTGGGAGACAGAGTCACCATCAGTTGCAGGGCAAGTCAGGACATTAGCAATTATTTAAACTGGTATCAGCAGAAACCAGATGGAACTGTTAAACTCCTGATCTACTACACATCAAGATTACACTCAGGAGTCCCATCAAGGTTCAGTGGCAGTGGGTCTGGGACAGATTATTCTCTCATCATTAGCAACCTGGAACAAGAAGATATTGCCACTTACTTTTGCCAACAGGATAGTAAGCGTCCTCC |
| IGKV4-91*01_S4567_PWD | PWD/PhJ | Musmus IGKV4-91*01 F | 96.81 | 292 | GAAATTGTGCTCACCCAGTCTCCAACCACCATGGCTGCATCTCCCGGGGAGAAGGTCACCATCACCTGCAGTGCCAGCTCAAGTATAAGTTCCAATTACTTGCACTGGTATCAGCAGAAGCCAGGATTCCCTCCTAAACTCTTGATATATAGGACATCCAATCTGGCTTCTGGAGTCCCAGCTCGCTTCAGTGGCAGTGGGTCTGGGACCTCTTACTCTCTCACAATTGGCACCATGGAGGCTGAAGGTGCTGCCACTTATTACTGCCAGCAGGGTAGTAGTTTACCACACA |
| IGKV1-110*02_S1372_PWD | PWD/PhJ | Musmus IGKV1-110*02 F, or Musmus IGKV1-117*01 F | 95.92 | 302 | GATGTTGTGATGACCCAAACTCCACTCTCCCTGCCTGTCAGTCTTGGAGATCAAGCCTCCATCTCTTGCAGATCTAGTCAGAGCATTGTACACAGTAATAGATACACCTATTTAGAATGGTACCTGCAGAAACCAGGCCAGTCTCTAAAGCTCCTGATATACGGGGTTTCCAACCGATTTTCTGGGGTCCCAGACAGGTTCAGTGGCAGTGGATCAAGGACAGATTTCACACTCAAGATCAGCAGAGTGGAGGCTGAGGATCTGGGAGTTTATTTCTGCTTTCAAGGTACACATGTTCCTCA |
| IGKV4-50*01_S7633_PWD | PWD/PhJ | Musmus IGKV4-50*01 F | 96.74 | 286 | GAAAATGTGCTCACCCAGTCTCCAGCAATCATGTCTGCAACTCTAGGGGAGAAGGTCACCATGAGCTGCAGGGCCAGCTCAAATGTAAAGTACATGTACTGGTACCAGCAGAAGTCAGGTGCCTCCCCCAAACTATGGATTTATTACACATTCAACCTGGCTTCTGGAGTCCCAGCTCGCTTCAGTGGCAGTGGGTCTGGGACCTCTTATTCTCTCACAATCAGCAGCGTGGAGGCTGAAGATGCTGCCACTTATTACTGCCAGCAGTTTACTAGTTCCCCATCCA |

| | | | | | |
|---|---|---|---|---|---|
| IGKV4-51*01_S7391_PWD | PWD/PhJ | Musmus IGKV4-51*01 F, or Musmus IGKV4-57-1*01 F | 92.55 | 292 | GAAAATGTGCTCACCCAGTCTCCAGCACTCATGGCTGCATCTCCAGGGGAGAAGGTCACCATGACCTGCAGTGCCAGCTCAAGTGTAAGTTCTAGCTACTTGCACTGGTACCAGCAGAAGTCAGGTGCCTCCCCCAAACTCTGGATTTACCGCACATCCAACCTGGCTTCCGGAGTCCCAGCTCGCTTCAGTGGCAGTGGGTCTGGGACCTCTTACTCTCTCACAATCAGCAACGTGGAGGCTGAAGATGATGCAACTTATTACTGCCAGCAGGGGTGGGATTACCCATTCA |
| IGKV4-81*01_S7470_PWD | PWD/PhJ | Musmus IGKV4-81*01 F | 96.81 | 292 | GAAAATGTGCTGACCCAGTCTCCAGCAATCATGGCTGCATCTCCAGGGGAGAAGGTCACCATGACCTGCAGTGCCAGCTCAAGTGTAAGTTCTGGTAACTTTCACTGGTACCAGCAGAAGCCAGGCACTTCTCCCAAACTCTGGATTTATAGGACATCCAACCTGGCTTCTGGAGTCCCAGCTCGCTTCAGTGGCAGTGGGTCTGGGACCTCTTACTCTCTTACAATCAGCAGCATGGAGGCCGAAGATGCTGCCACTTATTACTGCCAGCAGTGGAGTGGTTACCCACCCA |
| IGKV7-33*01_S5071_PWD | PWD/PhJ | Musmus IGKV7-33*01 F | 99.66 | 305 | GACATTGTGATGACTCAGTCTCCAACTTTCCTTGCTGTGACAGCAAGTAAGAAGGTCACCATTAGTTGCACGGCCAGTGAGAGCCTTTATTCAAGCAAACACAAGGTGCACTACTTGGCTTGGTACCAGAAGAAACCAGAGCAATCTCCTAAACTGCTGATATACGGGGCATCCAACCGATACATTGGGGTCCCTGATCGCTTCACAGGCAGTGGATCTGGGACAGATTTCACTCTGACCATCAGCAGTGTACAGGCTGAAGACCTCACACATTATTACTGTGCACAGTTTACAGCTATCCTCT |
| IGKV3-12*01_S3543_PWD | PWD/PhJ | Musmus IGKV3-12*01 F | 96.56 | 299 | GACATTGTGCTGACACAGTCTCCTGCTTCCTTAGCTGTATCTCTGGAGCAGAGGGCCACCATCTCATGCAGGGCCAGCCAAAGTGTCAGTACATCTAGCTATAGTTACATGCACTGGTACCAACAGAAACCAGGAAAGCCACCCAAACTCCTCATCTATGGTGCATCCAACCTAGAATCTGGGGTGCCTGCCAGGTTCAGTGGCAGTGGGTCTGGGACAGACTTCACCCTCAACATCCATCCTGTGGAGGAGGAGGATGCTGCAACATATTACTGTCAGCACAATAGGGAGATTCCTCC |
| IGKV4-56*01_S4810_PWD | PWD/PhJ | Musmus IGKV4-59*01 F | 94.57 | 286 | CAAATTGTTCTCACCCAGTCTCCAGCAATCATTTCTGCATCTCCGGGGCAGAAAGTCACCATAACCTGCAGTGCCAGCTCCAGTGTAAATTACATGCACTGGTTCCAGCAGAAGCCAGGATCCTCCCCCAAACTCTGGATTTATGACACATCCAAACTGGCTCCTGGAGTCCCTGCTCGCTTCAGTGGCAGTGGGTCTGGGACCTCTTACTCTCTCACAATCAGCAGCATGGAGGCTGAAGATGCTGCCACTTATTACTGCCAGCAGTGGAGTAGTTACCCACCCA |
| IGKV8-24*01_S6188_PWD | PWD/PhJ | Musmus IGKV8-24*01 F | 98.32 | 305 | GACATTGTGATGACACAGTCTCCATCCTCCCTGGCTATGTCAGTAGGACAGAAGGTCACTATGAGCTGCAAGTCCAGTCAGAGCCTTTTAAATAGTAGCAATCAAAAGAACTATTTGGCCTGGTACCAGCAGAAACCAGGACAGTCTCCTAAACTTCTGGTATACTTTGCATCCACTAGGGAATCTGGGGTCCCTGATCGCTTCACAGGCAGTGGATCTGGGACAGATTTCACTCTTGCCATCAGCAGTGTGCAGGCTGAAGACCTGGCAGTTTATTACTGTCAGCAACATTATAGCACTCCTCC |
| IGKV20-101-2*01_S8388_PWD | PWD/PhJ | Musmus IGKV20-101-2*01 F | 97.46 | 282 | AATATCCAGGTGATCCAGTCACCATTTCTGTCTGCATCTGTGGGAGAGAGGGTCACAATCAGCTGCAAGACACATCAGCATATTAACAGTTCCATAGCCTGGTACCAGCAAAAATTTGGAAAAGCTCCCAAACTCCTGATAAGAGATGGAAGTTTTTCTCTAACAGACACCCCATCAAGGTTCAGTGGGAATGGATTTGGCACAGATTTCACACTCAGCATCAGCAGTATGCAGCCTGAAGATGGTGCCACATATTTCTGCCAGCAGCATTTTAACTATTAC |

| | | | | | |
|---|---|---|---|---|---|
| IGKV9-120*01_S0946_PWD | PWD/PhJ | Musmus IGKV9-120*02 F | 97.13 | 285 | GACATCCAGATGACCCAGTCTCCATCCTCCTTATCTGCCTCTCTGGGAGAAAGAATCAGTCTCACTTGCCGGGCAAGTCAGGACATTGGTGGTAGCTTAAACTGGTTTCAGCAGAAACCAGATGGAACTTTTAAACGCCTGATCTACGCCACATCCAGTTTAGATTCTGGTGTCCCCAAAAGGTTCAGTGGCAGTAGGTCTGGGTCAGATTATTCTCTCACCATCAGCAGCCTTGAGTCTGAAGATTTTGCAGACTATTACTGTTTACAATATGCTAGTTATCCT |
| IGKV4-57-1*01_S4182_PWD | PWD/PhJ | Musmus IGKV4-57-1*01 F | 93.62 | 292 | GAAATTGTGCTCACCCAGTCTCCAACACTCATGACTGTATCTCCAGGGGAGAAGGTCACCATGACCTGCAGTGCCAGCTCAAGTGTAAGTTCCAGTTACTTGTTCTGGTACCAGCAGAAGTCAGGTGCCACCCCTAAACTCTGGATTTACGGCACATCCAACCTGGCTTCTGGAGTCCCAGCTCGCTTCAGTGGCAGTGGGTCTGGGACCTCTTACTCTCTCACAATCAGCAGCATGGAGGCTGAAGATGCTGCCACTTATTACTGCCAGCAGTGGAATGGTGCCCCACTCA |
| IGKV4-60*01_S2341_PWD | PWD/PhJ | Musmus IGKV4-60*01 P | 98.23 | 292 | GAAATTGTGCTCACCCAGTCTCCAGCACTCATGGCTGCATCTCCAGGGGAGAAGGTCAGCATCACCTGCAGTGTCAGCTCAAGTATAAGTTCCAGCCACTTACACTGGTACCAGCAGAAGTCAGGAACCTCCCCCAAACTCTGGATTTATGGCACATCCAACCTGGCTTCTGGAGTCCCTGCTCGCTTCAGTGGCAGTGGATCTGGGACCTCTTACTCTCTCACAATCAGCAGCATGGAGGCTGAAGATGCTGCCACTTATTACTGTCAACAGTGGAGTAGTTACCCACCCA |
| IGKV3-1*01_S7023_PWD | PWD/PhJ | Musmus IGKV3-1*01 F | 99.31 | 299 | GACATTGTGCTGACCCAATCTCCAGCTTCTTTGGCTGTGTCTCTAGGGCAGAGAGCCACCATCTCCTGCAGAGCCAGTGAAAGTGTTGAATATTATGGCACAAGTTTAATGCAGTGGTACCAACAGAAACCAGGACAGCCACCCAAAATCCTCATCTATGCTGCATCCAACGTAGAATCTGGGGTCCCTGCCAGGTTTAGTGGCAGTGGGTCTGGGACAGACTTCAGCCTCAACATCCATCCTGTGGAGGAGGATGATATTGCAATGTATTTCTGTCAGCAAAGTAGGAAGGTTCCTTC |
| IGKV3-9*01_S0307_PWD | PWD/PhJ | Musmus IGKV3-9*01 F | 91.07 | 299 | GACATTGTGCTGACCCAATCTCCAGCTTCTTTGGCTGTGTCTCTAGGGCAGAGGGCCACCATCTCCTGCAGAGCCAGCGAAAGTGTCAGTATTCGTGGTACTAGTTTAATGCACTGGTACCAACAGAAACCAGGACAGCCACCCAAAGTCCTCATCTATCTTGCATCCAACCTAGAATCTGGGGTTCCTGCAAGGTTTAGTGGCAGTGGGTCTGGGACAGGCTTCACCCTCAATATCCATCCTGTGGAGGAAGATGATGCTGCAACCTATTACTGTCAGCAAAGTAGGGAATATCCTCC |
| IGKV16-104*01_S2563_PWD | PWD/PhJ | Musmus IGKV14-87-2*01 F | 98.21 | 288 | GACATCCAAATGACCCAGTCCTCATCATTTCTCTCTGCATCTCTAGGAGATCATCTTACAATCAACTGCAGGGCCAGTAAGGATATTAACAAGTATTTTGCTTGGGTTCAACAGAAGCCAGGGAAGGCTCCAAGGATGTTGATTCATTTTGCTTCCACCTTGCTACCTGGGGTTCCAGAAAAGTTCAGTGGGAGTGGATCTGGGACAGATTTTTCTCTCACTGTCAGAAACATAGAGTCCGAAGATATTGCAATGTATTACTGTCTACAGTTTTCTGAACATCCGTAC |
| IGKV4-81*01_S3230_PWD | PWD/PhJ | Musmus IGKV4-81*01 F | 96.1 | 292 | GAAAATGTGCTGACCCAGTATCCAGCAATCATGGCTGCATCTCCAGGGGAGAAGGTCACCATGACCTGCAGTGCCAGCTCAAGTGTAAGTTCTGATAACTTTCACTGGTACCAGCAGAAGTCAGGCACTTCTCCCAAACTCTGGATTTATAGGACATCCAAATTGGCTTCTGGAGTCCCAGCTCGCTTCAGTGGCAGTGGGTCTGGGACCTCTTACTCTCTTACAATCAGCAGCGTGGAGGCCGAAGATGCTGCCACTTATTACTGCCAGCAGTGGAGTGGTTACCCACCCA |

| | | | | | |
|---|---|---|---|---|---|
| IGKV4-90*01_S4752_PWD | PWD/PhJ | Musmus IGKV4-90*01 F | 99.64 | 285 | GAAATTTTGCTCACCCAGTCTCCAGCAATCATAGCTGCATCTCCTGGGGAGAAGGTCA<br>CCATCACCTGCAGTGCCAGCTCAAGTGTAAGTTACATGAACTGGTACCAGCAGAAACC<br>AGGATCCTCTCCCAAAATATGGATTTATGGTATATCCAACCTGGCTTCTGGAGTTCCTG<br>CTCGCTTCAGTGGCAGTGGGTCTGGGACATCTTTCTCTTTCACAATCAACAGCATGGA<br>GGCTGAAGATGTTGCCACTTATTACTGTCAGCAAAGGAGTAGTTACCCACCC |
| IGKV2-109*03_S5201_PWD | PWD/PhJ | Musmus IGKV2-109*03 F, or Musmus IGKV2-109*04 F | 98.3 | 302 | GATATTGTGATGACACAGGCTGCATTCTCCAATCCAGTCACTCTTGGAACATCAGCTT<br>CCATCTCCTGCAGGTCTAGTAAGAGTCTCCTACATAGTAATGGCGTCACTTATTTGTAT<br>TGGTATCTGCAGAGGCCAGGCCAGTCTCCTCAGCTCCTGATATATCGGATGTCCAACC<br>TTGCCTCAGGAGTCCCAGACAGGTTCAGTAGCAGTGGGTCAGGAACTGATTTCACACT<br>GAGAATCAGCAGAGTGGAGGCTGAGGATGTGGGTGTTTATTACTGTGCTCAAATGCTA<br>GAACTCCCTCC |
| IGKV3-9*01_S8945_PWD | PWD/PhJ | Musmus IGKV3-9*01 F | 92.1 | 299 | GACATTGTGCTGACCCAATCTCCAGCTTCTTTGACTGTGTCTCTAGGGCAGAGAGCCA<br>CCATCTCCTGCAGAGCCAGTGAAAGTGTCAGTTTTGCTGGTACTAGTTTAATGCACTG<br>GTACCAACAGAAACCAGGACAGCCACCCAAACTCCTCATCTATCTTGCATCCAACCTA<br>GAATCTGGGGTTCCTGCCAGGTTTAGTGGCAGTGGGTCTGGGACAGACTTCACCCTCA<br>ATATCCATCCTGTGGAGGAAGATGATGCTGCAACCTATTACTGTCAGCAAAGTAGGGA<br>ATATCCTCC |
| IGKV11-125*01_PWD | PWD/PhJ | Musmus IGKV11-125*01 F | 100 | 287 | GATGTCCAGATGATTCAGTCTCCATCCTCCCTGTCTGCATCTTTGGGAGACATAGTCAC<br>CATGACTTGCCAGGCAAGTCAGGGCACTAGCATTAATTTAAACTGGTTTCAGCAAAAA<br>CCAGGGAAAGCTCCTAAGCTCCTGATCTATGGTGCAAGCAACTTGGAAGATGGGGTCC<br>CATCAAGGTTCAGTGGCAGTAGATATGGGACAGATTTCACTCTCACCATCAGCAGCCT<br>GGAGGATGAAGATATGGCAACTTATTTCTGTCTACAGCATAGTTATCTCCCTCC |
| IGKV13-85*01_S5594_PWD | PWD/PhJ | Musmus IGKV13-85*01 F | 97.85 | 287 | GACATCCAGATGACACAATCTTCATCCTCCTTGTCTGTATCTCTACGAGGCAGAGTCA<br>CCATTACGTGCAAGGCAAGTGACCACATTAATAATTGGTTAGCCTGGTATCAGCAGAA<br>ACCAGGAAATGCTTCTAGGCTCTTAATATCTGGTGCGACCAGTTTGGAAAATGGGGTT<br>CCTTCAAGATTCAGTGGCAGTGGATCTGGAAAGGATTACACTCTCAGCATTACCAGTC<br>TTCAGACTGAAGATGTTGCTACTTATTACTGTCAACAGTATTGGAGTACTCCTCC |
| IGKV5-39*01_S0817_PWD | PWD/PhJ | Musmus IGKV5-39*01 F | 99.64 | 287 | GACATTGTGATGACTCAGTCTCCAGCCACCCTGTCTGTGACTCCAGGAGATAGAGTCT<br>CTCTTTCCTGCAGGGCCAGCCAGAGTATTAGCAACTACTTACACTGGTATCAACAAAA<br>ATCACATGAGTCTCCAAGGCTTCTCATCAAATATGCTTCCCAATCCATCTCTGGGATCC<br>CCTCCAGGTTCAGTGGCAGTGGATCAGGGTCAGATTTCACTCTCAGTATCAACAGTGT<br>GGAACCTGAAGATGTTGGAGTGTATTACTGTCAAAATGGTCACAGCTTTCCTCC |
| IGKV6-d*01_S0533_PWD | PWD/PhJ | Musmus IGKV6-d*01 F | 98.2 | 287 | AGCATTGTGATGACCCAGTCTCCCAAATCCCTGCCTGTATCAGCAGGAGACAGGGTTA<br>CCATGACCTGCAAGGCCAGTCAGAGTGTGAGTAATGATGTAGCCTGGTACCAACAGA<br>AGCCAGGGCAGTCTCCTAAACTGGTGATATACTATGCATCCAATCGGTACACTGGAGT<br>CCCTGATTGCTTCACTGGCAGTGGATCTGGGACGGATTTCACTTTCACCAGCAGCAGT<br>GTGCAGGCTGAAGACCTGGCAGTTTATTTCTGTCAGCAGCATTATACTACTCCTCC |
| IGKV11-125*01_S1984_PWD | PWD/PhJ | Musmus IGKV11-106*02 F | 96.06 | 286 | GATGTTCAAATGACCCAGTCTCCATCCTCCCTGTCTGCATCTTTGGGAGAGAGAGTCTC<br>CCTGACTTGCCAGGCAAGTCAGAGCATTAGCAATTATTTAAACTGGTTTCAGCAAACA<br>CCAGGGAAAGCTCCTAGGCTCTTGATCTATGATGCAAACAACTTGGAAGATGGGGTCC<br>CTTCAAGGTTCAGTGGCACTGGATATGGGACAGATTTCACTTTCACCATCAGCAGCCT<br>GGAGGAAGAAGATGTCGCAACTTATTTCTGTCTACAGCATAAGTATCTCCCTC |

| | | | | | |
|---|---|---|---|---|---|
| IGKV3-9*01_S4460_PWD | PWD/PhJ | Musmus IGKV3-9*01 F | 92.1 | 299 | GACATTGTGCTGACCCAATCTCCAGCTTCTTTGGCTGTGTCTCTAGGGCAGAGGGCCACCATCTCCTGCAAGGCCAGCCAAAGTGTCAGTTTTGCTGGTACTAGTTTAATGCACTGGTACCAACAGAAACCAGGACAGCCACCCAAAGTCCTCATCTATCGTGCATCCAACCTAGAATCTGGGGTTCCTACCAGGTTTAGTGGCAGTGGGTCTGGGACAGACTTCACCCTCAATATCCACCCTGTGGAGGAAGATGATGCTGCAACCTATTACTGTCAGCAAAGTAGGGAATATCCTCC |
| IGKV4-91*01_S5853_PWD | PWD/PhJ | Musmus IGKV4-91*01 F | 96.81 | 292 | GAAATTGTGCTCACCCAGTCTCCAACCACCATGGCTGCATCTCCCGGGGAGAAGGTCACCATCACCTGCAGTGCCAGCTCAAGTATAAGTTCCAATTACTTGCACTGGTATCAGCAGAAGCCAGGATTCCCTCCTAAACTCTTGATATATAGGACATCCAATCTGGCTTCTGGAGTCCCAGCTCGCTTCAGTGGCAGTGGGTCTGGGACCTCTTACTCTCTCACAATTGGCACCATGGAGGCTGAAGGTGCTGCCACTTATTACTGCCAGCAGGGTAGTAGTTTACCACTCA |
| IGKV9-123*01_S4220_PWD | PWD/PhJ | Musmus IGKV9-123*01 F | 99.28 | 285 | GACATCCAGATGATTCAGTCTCCATCGTCCATGTTTGGCTCTCTGGGAGACAGAGTCAGTCTCTCTTGCCGGGCTAGTCAGGGCATTAGAGGTAATTTAGACTGGTATCAGCAGAAACCAGGTGGAACTATTAAACTCCTGATCTACTCCACATCCAATTTAAATTCTGGTGTCCCATCAAGGTTCAGTGGCAGTGGGTCTGGGTCAGATTATTCTCTCACCATCAGCAGCCTAGAGTCTGAAGATTTTGCAGACTATTACTGTCTACAGCGTAATGCGTATCCT |
| IGKV13-84*01_S4500_PWD | PWD/PhJ | Musmus IGKV13-84*01 F | 98.92 | 287 | GACATCCAGATGACACAATCTTCATCCTCCTTGTCTGTATCTCTAGGAGACAGAGTTACCATTACTTGCAAGGCAAGTGAGGACATATATAATCGGTTAGCCTCGTATCAGCAGAAACCAGGAAATGCTCCTAGGCTCTTAATATCTGGTGCAACCAGTTTGGAAACTGGGGTTCCTTCAAGATTCAGTGGCAGTGGATCTGGAAAGGATTACACTCTCAGCATTACCAGTCTTCAGACTGAAGATGTTGCTACTTATTACTGTCAACAGTATTGGAGTACTCCTCC |
| IGKV18-36*01_S9375_PWD | PWD/PhJ | Musmus IGKV18-36*01 F | 98.21 | 287 | ACTGGAGAAACAACACAGTCTCCAGCCTCTCTGAGTTTTTCTCTTGGTGAAACAGCAACCCTGACATGCAGGTCCAGTGAGAGTGTTGGCAGCTACTTAGCCTGGTACCAGCAGAAAGCAGAGCAAGTTCCCCGGCTCCTTATCCATAGTGCTTCCACTAGGGCCGGTGGTGTCCCAGTCCGGTTCAGTGGCACTGGGTCTGGGACAGACTTCACTCTCACCATCAGCAGTCTAGAACCTGAAGATGCTGCAGTTTACTACTGTCAACCTTTCAAAAGTTGGTCATA |
| IGKV8-18*01_S3987_PWD | PWD/PhJ | Musmus IGKV8-18*01 ORF, or Musmus IGKV8-26*01 ORF | 97.98 | 310 | GACATTGTGATGACCCAGTCTCCATCCTCCCTGGCTGTGACAGCAGGAGAGAAGGTCACTATGAGCTGCAAGTCCAGTCAGAGTCTTTTGTGGAGTGTAAACCAAAAGAACTACTTGTCCTGGTACCAGCAGAAACAAGGGAAGCCTCCTAAACTCCTTATCTATGGGGCATCCATTAGAGAATCTTGGGTCCCTGATCGATTCACAGGAAGTGGATCTGGGACAGACTTCACTCTCACCATTAGCAATGTGCATGCCGAAGACCTAGCAGTTTATTACTGTCAGCACAATCATGGCAGCTTTCTCCCCC |
| IGKV9-120*01_S9262_PWD | PWD/PhJ | Musmus IGKV9-120*02 F | 96.42 | 284 | GACATCCAGATGACCCAGTCTCCATCCTCCTTATCTGCCTCTCTGGGAGAAAGAATCAGTCTCACTTGCCGGGCAAGTCAGGACATTGGTGGTAGCTTAAACTGGTTTCAGCAGAAACCAGATGGAACTTTTAAACGCCTGATCTACGCCATATCCAGTTTAGATTCTGGTGTCCCCAAAAGGTTCAGTGGCAGTAGGTCTGGGTCAGATTATTCTCTCACCATCAGCAGCCTTGAGTCTGAAGATTTTGCAGACTATTACTGTTTACAATATGCAAGTTATCC |
| IGKV4-57-1*01_S8265_PWD | PWD/PhJ | Musmus IGKV4-57-1*01 F | 92.91 | 292 | GAAAATGTGCTCACCCAGTCTCCAGCACTCATGGCTGCATCTCCAGGGGAGAAGGTCACCATGACCTGCAGTGCCAGCTCAAGTGTAAGTTCCAGTTACTTGCACTGGTACCAGCAGAAGTCAGGTGCCTCCCCCAAACTCTGGATTTACGGCACATCCAACCTGGCTTCTGGAGTCCCAGCTCGCTTCAGTGGCAGTGGGTCTGGGACCTCTTACTCTCTCACAATCAGCAGCGTGGAGGCTGAAGATGATGCAACTTATTACTGCCAGCAGGCGTGGAATTACCCACTCA |

| | | | | | |
|---|---|---|---|---|---|
| IGKV4-92*01_S5510_PWD | PWD/PhJ | Musmus IGKV4-92*01 F | 99.65 | 291 | GAAATGGTTCTCACCCAGTCTCCAGTATCCATAACTGCATCTCGAGGGGAGAAGGTCACCATCACCTGCCGTGCCAGCTCAAGTATAAGTTCCAATTACTTACACTGGTACCAGCAGAAGCCAGGATCCTCCCCTAAACTTTTGATTTATAGGACATCCATCCTGGCATCTGGAGTCCTAGACAGCTTCAGTGGCAGTGGGTCTGAGAGCTCTTACACTCTGACAATCAGCTGCATGCAGGACGAAGTTGCTGCCACTTACTATTATCAGCAGGGGAGTAGTAGCCCACCA |
| IGKV12-98*01_S4885_PWD | PWD/PhJ | Musmus IGKV12-98*01 F | 98.21 | 287 | GACATTCAGATGACCCAGTCTCCTGCCTCCCAGTCTGCATCTCTGGGAGAAAGTGTCACCATCACATGCCTGGCAAGTCAGCCCATTGGTGCATGGTTAGCATGGTATCAGCAGAAACCAGGGAAATCTCCTCAGCTCCTGATTTATGCTGCAACCAGCTTGGCAGATGGGGTCCCATCAAGGTTCAGTGGTAGTGGATCTGGCACAAAGTTTTCTTTCAAGATCAGCAGCCTACAGGCTGAAGATTTTGCAAGTTATTACTGTCAACAACGTTACAGTACTCCTCT |
| IGKV4-59*01_S5858A_PWD | PWD/PhJ | Musmus IGKV4-59*01 F | 97.1 | 286 | CAAATTGTTCTCACCCAGTCTCCAGCAATCATGTCTGCATCTCCAGGGGAGAAGGTCACCATGACCTGCAGTGCCAGCTCAAGTGTAAGTTACATGTACTGGTACCAGCAGAAGCCAGGATCCTCCCCCAAAAAATGGATTTATGACACATCCAACCTGGCTTCTGGAGTCCCCGCTCGCTTCAGTGGCAGTGGGTCTGGGACCTCTTATTCTCTCACAATCAGCAGCATGGAGGCTGAAGATGCTGCCACTTATTACTGCCAGCAGTGGAGTAGTTACCCACCCA |
| IGKV4-79*01_S6901_PWD | PWD/PhJ | Musmus IGKV4-79*01 F | 95.39 | 291 | CAAATTGTTCTCACCCAGTCTCCAGCAATCATGTCTGCATCTCCAGGGGAGAAGGTCACCATGACCTGCAGGGCCAGCTCAAGTGTACGTTCCAGTTACTTGCACTGGTACCAGCAGAAGCCAGGATCTTCCCCCAAACTCTGGATTTATAGAACATCCAACCTGGCTTCAGGAGTCCCAGCTCGCTTCAGTGGCAGTGGGTCTGGGACCTCTTACTCTCTCACAATCAGCAGCATGGAGGCTGAAGATGCTGCCACTTATTACTGCCAGCAGTGGAGTAGTTACCCACCC |
| IGKV8-27*01_PWD | PWD/PhJ | Musmus IGKV8-27*01 F | 100 | 300 | AACATTATGATGACACAGTCGCCATCATCTCTGGCTGTGTCTGCAGGAGAAAAGGTCACTATGAGCTGTAAGTCCAGTCAAAGTGTTTTATACAGTTCAAATCAGAAGAACTACTTGGCCTGGTACCAGCAGAAACCAGGGCAGTCTCCTAAACTGCTGATCTACTGGGCATCCACTAGGGAATCTGGTGTCCCTGATCGCTTCACAGGCAGTGGATCTGGGACAGATTTTACTCTTACCATCAGCAGTGTACAAGCTGAAGACCTGGCAGTTTATTACTGTCATCAATACCTCTCCTCG |
| IGKV6-13*01_S9207_SJL | SJL/J | Musmus IGKV6-13*01 F | 98.57 | 285 | GACATTGTGATGACCCAGTCTCAAAAATTCATGTCCACAACAGTAGGAGACAGGGTCAGCATCACCTGCAAGGCCAGTCAGAATGTGGGTACTGCTGTAGCCTGGTATCAACAGAAACCAGGACAATCTCCTAAACTACTGATTTACTCAGCATCCAATCGGTACACTGGAGTCCCTGATCGCTTCACAGGCAGTGGATCTGGGACAGATTTCACTCTCACCATTAGCAATATGCAGTCTGAAGACCTGGCAGATTATTTCTGTCAGCAATATAGCAGCTATCCT |
| IGKV6-15*01_SJL | SJL/J | Musmus IGKV6-15*01 F | 100 | 287 | GACATTGTGATGACCCAGTCTCAAAAATTCATGTCCACATCAGTAGGAGACAGGGTCAGCGTCACCTGCAAGGCCAGTCAGAATGTGGGTACTAATGTAGCCTGGTATCAACAGAAACCAGGGCAATCTCCTAAAGCACTGATTTACTCGGCATCCTACCGGTACAGTGGAGTCCCTGATCGCTTCACAGGCAGTGGATCTGGGACAGATTTCACTCTCACCATCAGCAATGTGCAGTCTGAAGACTTGGCAGAGTATTTCTGTCAGCAATATAACAGCTATCCTCT |
| IGKV8-30*01_SJL | SJL/J | Musmus IGKV8-30*01 F | 100 | 305 | GACATTGTGATGTCACAGTCTCCATCCTCCCTAGCTGTGTCAGTTGGAGAGAAGGTTACTATGAGCTGCAAGTCCAGTCAGAGCCTTTTATATAGTAGCAATCAAAAGAACTACTTGGCCTGGTACCAGCAGAAACCAGGGCAGTCTCCTAAACTGCTGATTTACTGGGCATCCACTAGGGAATCTGGGGTCCCTGATCGCTTCACAGGCAGTGGATCTGGGACAGATTTCACTCTCACCATCAGCAGTGTGAAGGCTGAAGACCTGGCAGTTTATTACTGTCAGCAATATTATAGCTATCCTCC |

| | | | | | |
|---|---|---|---|---|---|
| IGKV4-59*01_SJL | SJL/J | Musmus IGKV4-59*01 F | 100 | 286 | CAAATTGTTCTCACCCAGTCTCCAGCAATCATGTCTGCATCTCCAGGGGAGAAGGTCACCATGACCTGCAGTGCCAGCTCAAGTGTAAGTTACATGCACTGGTACCAGCAGAAGTCAGGCACCTCCCCCAAAAGATGGATTTATGACACATCCAAACTGGCTTCTGGAGTCCCTGCTCGCTTCAGTGGCAGTGGGTCTGGGACCTCTTACTCTCTCACAATCAGCAGCATGGAGGCTGAAGATGCTGCCACTTATTACTGCCAGCAGTGGAGTAGTAACCCACCCA |
| IGKV8-27*01_SJL | SJL/J | Musmus IGKV8-27*01 F | 100 | 300 | AACATTATGATGACACAGTCGCCATCATCTCTGGCTGTGTCTGCAGGAGAAAAGGTCACTATGAGCTGTAAGTCCAGTCAAAGTGTTTTATACAGTTCAAATCAGAAGAACTACTTGGCCTGGTACCAGCAGAAACCAGGGCAGTCTCCTAAACTGCTGATCTACTGGGCATCCACTAGGGAATCTGGTGTCCCTGATCGCTTCACAGGCAGTGGATCTGGGACAGATTTTACTCTTACCATCAGCAGTGTACAAGCTGAAGACCTGGCAGTTTATTACTGTCATCAATACCTCTCCTCG |
| IGKV10-96*01_SJL | SJL/J | Musmus IGKV10-96*01 F | 100 | 286 | GATATCCAGATGACACAGACTACATCCTCCCTGTCTGCCTCTCTGGGAGACAGAGTCACCATCAGTTGCAGGGCAAGTCAGGACATTAGCAATTATTTAAACTGGTATCAGCAGAAACCAGATGGAACTGTTAAACTCCTGATCTACTACACATCAAGATTACACTCAGGAGTCCCATCAAGGTTCAGTGGCAGTGGGTCTGGAACAGATTATTCTCTCACCATTAGCAACCTGGAGCAAGAAGATATTGCCACTTACTTTTGCCAACAGGGTAATACGCTTCCTC |
| IGKV12-44*01_S4418_SJL | SJL/J | Musmus IGKV12-44*01 F | 100 | 288 | GACATCCAGATGACTCAGTCTCCAGCCTCCCTATCTGCATCTGTGGGAGAAACTGTCACCATCACATGTCGAGCAAGTGAGAATATTTACAGTTATTTAGCATGGTATCAGCAGAAACAGGGAAAATCTCCTCAGCTCCTGGTCTATAATGCAAAAACCTTAGCAGAAGGTGTGCCATCAAGGTTCAGTGGCAGTGGATCAGGCACACAGTTTTCTCTGAAGATCAACAGCCTGCAGCCTGAAGATTTTGGGAGTTATTACTGTCAACATCATTATGGTACTCCTCCT |
| IGKV6-23*01_SJL | SJL/J | Musmus IGKV6-23*01 F | 100 | 287 | GACATTGTGATGACCCAGTCTCACAAATTCATGTCCACATCAGTAGGAGACAGGGTCAGCATCACCTGCAAGGCCAGTCAGGATGTGGGTACTGCTGTAGCCTGGTATCAACAGAAACCAGGGCAATCTCCTAAACTACTGATTTACTGGGCATCCACCCGGCACACTGGAGTCCCTGATCGCTTCACAGGCAGTGGATCTGGGACAGATTTCACTCTCACCATTAGCAATGTGCAGTCTGAAGACTTGGCAGATTATTTCTGTCAGCAATATAGCAGCTATCCTCT |
| IGKV12-41*01_SJL | SJL/J | Musmus IGKV12-41*01 F | 100 | 287 | GACATCCAGATGACTCAGTCTCCAGCCTCCCTATCTGCATCTGTGGGAGAAACTGTCACCATCACATGTCGAGCAAGTGGGAATATTCACAATTATTTAGCATGGTATCAGCAGAAACAGGGAAAATCTCCTCAGCTCCTGGTCTATAATGCAAAAACCTTAGCAGATGGTGTGCCATCAAGGTTCAGTGGCAGTGGATCAGGAACACAATATTCTCTCAAGATCAACAGCCTGCAGCCTGAAGATTTTGGGAGTTATTACTGTCAACATTTTTGGAGTACTCCTCC |
| IGKV4-55*01_SJL | SJL/J | Musmus IGKV4-55*01 F | 100 | 286 | CAAATTGTTCTCACCCAGTCTCCAGCAATCATGTCTGCATCTCCAGGGGAGAAGGTCACCATGACCTGCAGTGCCAGCTCAAGTGTAAGTTACATGTACTGGTACCAGCAGAAGCCAGGATCCTCCCCCAGACTCCTGATTTATGACACATCCAACCTGGCTTCTGGAGTCCCTGTTCGCTTCAGTGGCAGTGGGTCTGGGACCTCTTACTCTCTCACAATCAGCCGAATGGAGGCTGAAGATGCTGCCACTTATTACTGCCAGCAGTGGAGTAGTTACCCACCCA |
| IGKV4-57-1*01_SJL | SJL/J | Musmus IGKV4-57*01 F | 100 | 292 | GAAAATGTGCTCACCCAGTCTCCAGCAATCATGTCTGCATCTCCAGGGGAAAAGGTCACCATGACCTGCAGGGCCAGCTCAAGTGTAAGTTCCAGTTACTTGCACTGGTACCAGCAGAAGTCAGGTGCCTCCCCCAAACTCTGGATTTATAGCACATCCAACTTGGCTTCTGGAGTCCCTGCTCGCTTCAGTGGCAGTGGGTCTGGGACCTCTTACTCTCTCACAATCAGCAGTGTGGAGGCTGAAGATGCTGCCACTTATTACTGCCAGCAGTACAGTGGTTACCCACTCA |

| | | | | | |
|---|---|---|---|---|---|
| IGKV4-80*01_SJL | SJL/J | Musmus IGKV4-80*01 F | 100 | 282 | CAAATTGTTCTCACCCAGTCTCCAGCAATCATGTCTGCATCTCTAGGGGAGGAGATCACCCTAACCTGCAGTGCCAGCTCGAGTGTAAGTTACATGCACTGGTACCAGCAGAAGTCAGGCACTTCTCCCAAACTCTTGATTTATAGCACATCCAACCTGGCTTCTGGAGTCCCTTCTCGCTTCAGTGGCAGTGGGTCTGGGACCTTTTATTCTCTCACAATCAGCAGTGTGGAGGCTGAAGATGCTGCCGATTATTACTGCCATCAGTGGAGTAGTTATCCA |
| IGKV4-70*01_SJL | SJL/J | Musmus IGKV4-70*01 F | 100 | 282 | CAAATTGTTCTCACCCAGTCTCCAGCAATCATGTCTGCATCTCCAGGGGAGAAGGTCACCATGACCTGCAGTGCCAGCTCAAGTATAAGTTACATGCACTGGTACCAGCAGAAGCCAGGCACCTCCCCCAAAAGATGGATTTATGACACATCCAAACTGGCTTCTGGAGTCCCTGCTCGCTTCAGTGGCAGTGGGTCTGGGACCTCTTATTCTCTCACAATCAGCAGCATGGAGGCTGAAGATGCTGCCACTTATTACTGCCATCAGCGGAGTAGTTACCCA |
| IGKV2-137*01_SJL | SJL/J | Musmus IGKV2-137*01 F | 100 | 302 | GATATTGTGATGACTCAGGCTGCACCCTCTGTACCTGTCACTCCTGGAGAGTCAGTATCCATCTCCTGCAGGTCTAGTAAGAGTCTCCTGCATAGTAATGGCAACACTTACTTGTATTGGTTCCTGCAGAGGCCAGGCCAGTCTCCTCAGCTCCTGATATATCGGATGTCCAACCTTGCCTCAGGAGTCCCAGACAGGTTCAGTGGCAGTGGGTCAGGAACTGCTTTCACACTGAGAATCAGTAGAGTGGAGGCTGAGGATGTGGGTGTTTATTACTGTATGCAACATCTAGAATATCCTTT |
| IGKV4-91*01_S5997_SJL | SJL/J | Musmus IGKV4-91*01 F | 100 | 292 | GAAATTGTGCTCACCCAGTCTCCAACCACCATGGCTGCATCTCCCGGGGAGAAGATCACTATCACCTGCAGTGCCAGCTCAAGTATAAGTTCCAATTACTTGCATTGGTATCAGCAGAAGCCAGGATTCTCCCCTAAACTCTTGATTTATAGGACATCCAATCTGGCTTCTGGAGTCCCAGCTCGCTTCAGTGGCAGTGGGTCTGGGACCTCTTACTCTCTCACAATTGGCACCATGGAGGCTGAAGATGTTGCCACTTACTACTGCCAGCAGGGTAGTAGTATACCACTCA |
| IGKV8-19*01_SJL | SJL/J | Musmus IGKV8-19*01 F | 100 | 305 | GACATTGTGATGACACAGTCTCCATCCTCCCTGACTGTGACAGCAGGAGAGAAGGTCACTATGAGCTGCAAGTCCAGTCAGAGTCTGTTAAACAGTGGAAATCAAAAGAACTACTTGACCTGGTACCAGCAGAAACCAGGGCAGCCTCCTAAACTGTTGATCTACTGGGCATCCACTAGGGAATCTGGGGTCCCTGATCGCTTCACAGGCAGTGGATCTGGAACAGATTTCACTCTCACCATCAGCAGTGTGCAGGCTGAAGACCTGGCAGTTTATTACTGTCAGAATGATTATAGTTATCCTCC |
| IGKV4-74*01_SJL | SJL/J | Musmus IGKV4-74*01 F | 100 | 292 | CAAATTGTTCTCACCCAGTCTCCAGCAATCATGTCTGCATCTCTAGGGGAACGGGTCACCATGACCTGCACTGCCAGCTCAAGTGTAAGTTCCAGTTACTTGCACTGGTACCAGCAGAAGCCAGGATCCTCCCCCAAACTCTGGATTTATAGCACATCCAACCTGGCTTCTGGAGTCCCAGCTCGCTTCAGTGGCAGTGGGTCTGGGACCTCTTACTCTCTCACAATCAGCAGCATGGAGGCTGAAGATGCTGCCACTTATTACTGCCACCAGTATCATCGTTCCCCACCCA |
| IGKV1-117*01_SJL | SJL/J | Musmus IGKV1-117*01 F | 100 | 302 | GATGTTTTGATGACCCAAACTCCACTCTCCCTGCCTGTCAGTCTTGGAGATCAAGCCTCCATCTCTTGCAGATCTAGTCAGAGCATTGTACATAGTAATGGAAACACCTATTTAGAATGGTACCTGCAGAAACCAGGCCAGTCTCCAAAGCTCCTGATCTACAAAGTTTCCAACCGATTTTCTGGGGTCCCAGACAGGTTCAGTGGCAGTGGATCAGGGACAGATTTCACACTCAAGATCAGCAGAGTGGAGGCTGAGGATCTGGGAGTTTATTACTGCTTTCAAGGTTCACATGTTCCTCC |

| | | | | | |
|---|---|---|---|---|---|
| IGKV15-103*01_SJL | SJL/J | Musmus IGKV15-103*01 ORF | 100 | 287 | GACATCCAGATGAACCAGTCTCCATCCAGTCTGTCTGCATCCCTTGGAGACACAATTACCATCACTTGCCATGCCAGTCAGAACATTAATGTTTGGTTAAGCTGGTACCAGCAGAAACCAGGAAATATTCCTAAACTATTGATCTATAAGGCTTCCAACTTGCACACAGGCGTCCCATCAAGGTTTAGTGGCAGTGGATCTGGAACAGGTTTCACATTAACCATCAGCAGCCTGCAGCCTGAAGACATTGCCACTTACTACTGTCAACAGGGTCAAAGTTATCCTCT |
| IGKV4-50*01_S0901_SJL | SJL/J | Musmus IGKV4-50*01 F | 100 | 286 | GAAAATGTGCTCACCCAGTCTCCAGCAATCATGTCTGCATCTCTAGGGGAGAAGGTCACCATGAGCTGCAGGGCCAGCTCAAGTGTAAATTACATGTACTGGTACCAGCAGAAGTCAGATGCCTCCCCCAAACTATGGATTTATTACACATCCAACCTGGCTCCTGGAGTCCCAGCTCGCTTCAGTGGCAGTGGGTCTGGGAACTCTTATTCTCTCACAATCAGCAGCATGGAGGGTGAAGATGCTGCCACTTATTACTGCCAGCAGTTTACTAGTTCCCCATACA |
| IGKV12-46*01_SJL | SJL/J | Musmus IGKV12-46*01 F | 100 | 287 | GACATCCAGATGACTCAGTCTCCAGCCTCCCTATCTGTATCTGTGGGAGAAACTGTCACCATCACATGTCGAGCAAGTGAGAATATTTACAGTAATTTAGCATGGTATCAGCAGAAACAGGGAAAATCTCCTCAGCTCCTGGTCTATGCTGCAACAAACTTAGCAGATGGTGTGCCATCAAGGTTCAGTGGCAGTGGATCAGGCACACAGTATTCCCTCAAGATCAACAGCCTGCAGTCTGAAGATTTTGGGAGTTATTACTGTCAACATTTTTGGGGTACTCCTCC |
| IGKV1-135*01_SJL | SJL/J | Musmus IGKV1-135*01 F | 100 | 302 | GATGTTGTGATGACCCAGACTCCACTCACTTTGTCGGTTACCATTGGACAACCAGCCTCCATCTCTTGCAAGTCAAGTCAGAGCCTCTTAGATAGTGATGGAAAGACATATTTGAATTGGTTGTTACAGAGGCCAGGCCAGTCTCCAAAGCGCCTAATCTATCTGGTGTCTAAACTGGACTCTGGAGTCCCTGACAGGTTCACTGGCAGTGGATCAGGGACAGATTTCACACTGAAAATCAGCAGAGTGGAGGCTGAGGATTTGGGAGTTTATTATTGCTGGCAAGGTACACATTTTCCTCA |
| IGKV4-63*01_SJL | SJL/J | Musmus IGKV4-63*01 F | 100 | 286 | GAAAATGTTCTCACCCAGTCTCCAGCAATCATGTCTGCATCTCCAGGGGAAAAGGTCACCATGACCTGCAGTGCCAGCTCAAGTGTAAGTTACATGCACTGGTACCAGCAGAAGTCAAGCACCTCCCCCAAACTCTGGATTTATGACACATCCAAACTGGCTTCTGGAGTCCCAGGTCGCTTCAGTGGCAGTGGGTCTGGAAACTCTTACTCTCTCACGATCAGCAGCATGGAGGCTGAAGATGTTGCCACTTATTACTGTTTTCAGGGGAGTGGGTACCCACTCA |
| IGKV8-21*01_SJL | SJL/J | Musmus IGKV8-21*01 F | 100 | 302 | GACATTGTGATGTCACAGTCTCCATCCTCCCTGGCTGTGTCAGCAGGAGAGAAGGTCACTATGAGCTGCAAATCCAGTCAGAGTCTGCTCAACAGTAGAACCCGAAAGAACTACTTGGCTTGGTACCAGCAGAAACCAGGGCAGTCTCCTAAACTGCTGATCTACTGGGCATCCACTAGGGAATCTGGGGTCCCTGATCGCTTCACAGGCAGTGGATCTGGGACAGATTTCACTCTCACCATCAGCAGTGTGCAGGCTGAAGACCTGGCAGTTTATTACTGCAAGCAATCTTATAATCTTCC |
| IGKV3-7*01_S5296_SJL | SJL/J | Musmus IGKV3-7*02 F | 100 | 299 | GACATTGTGCTGACACAGTCTCCTGCTTCCTTAGCTGTATCTCTGGGGCAGAGGGCCACCATCTCATGCAGGGCCAGCCAAAGTGTCAGTACATCTAGCTATAGTTATATGCACTGGTACCAACAGAAACCAGGACAGCCACCCAAACTCCTCATCAAGTATGCATCCAACCTAGAATCTGGGGTCCCTGCCAGGTTCAGTGGCAGTGGGTCTGGGACAGACTTCACCCTCAACATCCATCCTGTGGAGGAGGAGGATACTGCAACATATTACTGTCAGCACAGTTGGGAGATTCCTCC |
| IGKV5-43*01_SJL | SJL/J | Musmus IGKV5-43*01 F | 100 | 287 | GATATTGTGCTAACTCAGTCTCCAGCCACCCTGTCTGTGACTCCAGGAGATAGCGTCAGTCTTTCCTGCAGGGCCAGCCAAAGTATTAGCAACAACCTACACTGGTATCAACAAAAATCACATGAGTCTCCAAGGCTTCTCATCAAGTATGCTTCCCAGTCCATCTCTGGGATCCCCTCCAGGTTCAGTGGCAGTGGATCAGGGACAGATTTCACTCTCAGTATCAACAGTGTGGAGACTGAAGATTTTGGAATGTATTTCTGTCAACAGAGTAACAGCTGGCCTCA |

| | | | | | |
|---|---|---|---|---|---|
| IGKV8-28*01_SJL | SJL/J | Musmus IGKV8-28*01 F | 100 | 305 | GACATTGTGATGACACAGTCTCCATCCTCCCTGAGTGTGTCAGCAGGAGAGAAGGTCACTATGAGCTGCAAGTCCAGTCAGAGTCTGTTAAACAGTGGAAATCAAAAGAACTACTTGGCCTGGTACCAGCAGAAACCAGGGCAGCCTCCTAAACTGTTGATCTACGGGGCATCCACTAGGGAATCTGGGGTCCCTGATCGCTTCACAGGCAGTGGATCTGGAACCGATTTCACTCTTACCATCAGCAGTGTGCAGGCTGAAGACCTGGCAGTTTATTACTGTCAGAATGATCATAGTTATCCTCC |
| IGKV6-32*01_SJL | SJL/J | Musmus IGKV6-32*01 F | 100 | 287 | AGTATTGTGATGACCCAGACTCCCAAATTCCTGCTTGTATCAGCAGGAGACAGGGTTACCATAACCTGCAAGGCCAGTCAGAGTGTGAGTAATGATGTAGCTTGGTACCAACAGAAGCCAGGGCAGTCTCCTAAACTGCTGATATACTATGCATCCAATCGCTACACTGGAGTCCCTGATCGCTTCACTGGCAGTGGATATGGGACGGATTTCACTTTCACCATCAGCACTGTGCAGGCTGAAGACCTGGCAGTTTATTTCTGTCAGCAGGATTATAGCTCTCCTCC |
| IGKV12-89*01_SJL | SJL/J | Musmus IGKV12-89*01 F | 100 | 287 | GACATCCAGATGACTCAGTCTCCAGCTTCACTGTCTGCATCTGTGGGAGAAACTGTCACCATCACATGTGGAGCAAGTGAGAATATTTACGGTGCTTTAAATTGGTATCAGCGGAAACAGGGAAAATCTCCTCAGCTCCTGATCTATGGTGCAACCAACTTGGCAGATGGCATGTCATCGAGGTTCAGTGGCAGTGGATCTGGTAGACAGTATTCTCTCAAGATCAGTAGCCTGCATCCTGACGATGTTGCAACGTATTACTGTCAAAATGTGTTAAGTACTCCTCC |
| IGKV4-57*01_S9219_SJL | SJL/J | Musmus IGKV4-57-1*01 F | 100 | 286 | CAAATTGTTCTCACCCAGTCTCCAGCAATCATGTCTGCATCTCCAGGGGAGAAGGTCACCATAACCTGCAGTGCCAGCTCAAGTGTAAGTTACATGCACTGGTTCCAGCAGAAGCCAGGCACTTCTCCCAAACTCTGGATTTATAGCACATCCAACCTGGCTTCTGGAGTCCCTGCTCGCTTCAGTGGCAGTGGATCTGGGACCTCTTACTCTCTCACAATCAGCCGAATGGAGGCTGAAGATGCTGCCACTTATTACTGCCAGCAAAGGAGTAGTTACCCACTCA |
| IGKV10-94*01_SJL | SJL/J | Musmus IGKV10-94*01 F | 100 | 287 | GATATCCAGATGACACAGACTACATCCTCCCTGTCTGCCTCTCTGGGAGACAGAGTCACCATCAGTTGCAGTGCAAGTCAGGGCATTAGCAATTATTTAAACTGGTATCAGCAGAAACCAGATGGAACTGTTAAACTCCTGATCTATTACACATCAAGTTTACACTCAGGAGTCCCATCAAGGTTCAGTGGCAGTGGGTCTGGGACAGATTATTCTCTCACCATCAGCAACCTGGAACCTGAAGATATTGCCACTTACTATTGTCAGCAGTATAGTAAGCTTCCTCC |
| IGKV4-53*01_SJL | SJL/J | Musmus IGKV4-53*01 F | 100 | 292 | GAAATTGTGCTCACCCAGTCTCCAGCACTCATGGCTGCATCTCCAGGGGAGAAGGTCACCATCACCTGCAGTGTCAGCTCAAGTATAAGTTCCAGCAACTTGCACTGGTACCAGCAGAAGTCAGAAACCTCCCCCAAACCCTGGATTTATGGCACATCCAACCTGGCTTCTGGAGTCCCTGTTCGCTTCAGTGGCAGTGGATCTGGGACCTCTTATTCTCTCACAATCAGCAGCATGGAGGCTGAAGATGCTGCCACTTATTACTGTCAACAGTGGAGTAGTTACCCACTCA |
| IGKV4-68*01_S3092_SJL | SJL/J | Musmus IGKV4-68*01 F | 100 | 286 | CAAATTGTTCTCACCCAGTCTCCAGCACTCATGTCTGCATCTCCAGGGGAGAAGGTCACCATGACCTGCAGTGCCAGCTCAAGTGTAAGTTACATGTACTGGTACCAGCAGAAGCCAAGATCCTCCCCCAAACCCTGGATTTATCTCACATCCAACCTGGCTTCTGGAGTCCCTGCTCGCTTCAGTGGCAGTGGGTCTGGGACCTCTTACTCTCTCACAATCAGCAGCATGGAGGCTGAAGATGCTGCCACTTATTACTGCCAGCAGTGGAGTAGTAACCCACTCA |
| IGKV3-5*01_S5428_SJL | SJL/J | Musmus IGKV3-5*01 F | 97.94 | 299 | GACATTGTGCTGACCCAATCTCCAGCTTCTTTGGCTGTGTCTCTAGGGCAGAGGGCCACCATCTCCTGCAGAGCCAGCGAAAGTGTTGATAATTATGGCATTAGTTTTATGCACTGGTACCAGCAGAAACCAGGACAGCCACCCAAACTCCTCATCTATCGTGCATCCAACCTAGAATCTGGGATCCCTGCCAGGTTCAGTGGCAGTGGGTCTAGGACAGACTTCACCCTCACCATTAATCCTGTGGAGACTGATGATGTTGCAACCTATTACTGTCAGCAAAGTAATAAGGATCCTCC |

| | | | | | |
|---|---|---|---|---|---|
| IGKV6-20*01_SJL | SJL/J | Musmus IGKV6-20*01 F | 100 | 287 | AACATTGTAATGACCCAATCTCCCAAATCCATGTCCATGTCAGTAGGAGAGAGGGTCACCTTGAGCTGCAAGGCCAGTGAGAATGTGGGTACTTATGTATCCTGGTATCAACAGAAACCAGAGCAGTCTCCTAAACTGCTGATATACGGGGCATCCAACCGGTACACTGGGGTCCCCGATCGCTTCACAGGCAGTGGATCTGCAACAGATTTCACTCTGACCATCAGCAGTGTGCAGGCTGAAGACCTTGCAGATTATCACTGTGGACAGAGTTACAGCTATCCTCC |
| IGKV1-110*01_SJL | SJL/J | Musmus IGKV1-110*01 F | 100 | 302 | GATGTTGTGATGACCCAAACTCCACTCTCCCTGCCTGTCAGTCTTGGAGATCAAGCCTCCATCTCTTGCAGATCTAGTCAGAGCCTTGTACACAGTAATGGAAACACCTATTTACATTGGTACCTGCAGAAGCCAGGCCAGTCTCCAAAGCTCCTGATCTACAAAGTTTCCAACCGATTTTCTGGGGTCCCAGACAGGTTCAGTGGCAGTGGATCAGGGACAGATTTCACACTCAAGATCAGCAGAGTGGAGGCTGAGGATCTGGGAGTTTATTTCTGCTCTCAAAGTACACATGTTCCTCC |
| IGKV13-85*01_SJL | SJL/J | Musmus IGKV13-85*01 F | 100 | 287 | GACATCCAGATGACACAATCTTCATCCTACTTGTCTGTATCTCTAGGAGGCAGAGTCACCATTACTTGCAAGGCAAGTGACCACATTAATAATTGGTTAGCCTGGTATCAGCAGAAACCAGGAAATGCTCCTAGGCTCTTAATATCTGGTGCAACCAGTTTGGAAACTGGGGTTCCTTCAAGATTCAGTGGCAGTGGATCTGGAAAGGATTACACTCTCAGCATTACCAGTCTTCAGACTGAAGATGTTGCTACTTATTACTGTCAACAGTATTGGAGTACTCCTCC |
| IGKV6-17*01_SJL | SJL/J | Musmus IGKV6-17*01 F | 100 | 287 | GACATTGTGATGACCCAGTCTCACAAATTCATGTCCACATCAGTAGGAGACAGGGTCAGCATCACCTGCAAGGCCAGTCAGGATGTGAGTACTGCTGTAGCCTGGTATCAACAGAAACCAGGACAATCTCCTAAACTACTGATTTACTCGGCATCCTACCGGTACACTGGAGTCCCTGATCGCTTCACTGGCAGTGGATCTGGGACGGATTTCACTTTCACCATCAGCAGTGTGCAGGCTGAAGACCTGGCAGTTTATTACTGTCAGCAACATTATAGTACTCCTCC |
| IGKV2-109*01_SJL | SJL/J | Musmus IGKV2-109*01 F | 100 | 302 | GATATTGTGATGACGCAGGCTGCATTCTCCAATCCAGTCACTCTTGGAACATCAGCTTCCATCTCCTGCAGGTCTAGTAAGAGTCTCCTACATAGTAATGGCATCACTTATTTGTATTGGTATCTGCAGAAGCCAGGCCAGTCTCCTCAGCTCCTGATTTATCAGATGTCCAACCTTGCCTCAGGAGTCCCAGACAGGTTCAGTAGCAGTGGGTCAGGAACTGATTTCACACTGAGAATCAGCAGAGTGGAGGCTGAGGATGTGGGTGTTTATTACTGTGCTCAAAATCTAGAACTTCCTCC |
| IGKV8-16*01_SJL | SJL/J | Musmus IGKV8-16*01 F | 100 | 305 | GAAATTGTGTTGACCCAGTCTATACCATCCCTGACTGTGTCAGCAGGAGAGAGGGTCACTATCAGCTGCAAATCCAATCAGAATCTTTTATGGAGTGGAAACCAAAGGTACTGTTTGGTCTGGCACCAGTGGAAACCAGGGCAAACTCCTACACCGTTGATCACCTGGACATCTGATAGGTACTCTGGAGTCCCTGATCGTTTCATAGGCAGTGGATCTGTGACAGATTTCACTCTGACCATCAGCAGTGTGCAGGCTGAAGATGTGGCAGTTTATTTCTGTCAGCAGCATTTACACATTCCTCC |
| IGKV8-24*01_SJL | SJL/J | Musmus IGKV8-24*01 F | 100 | 305 | GACATTGTGATGACACAGTCTCCATCCTCCCTGGCTATGTCAGTAGGACAGAAGGTCACTATGAGCTGCAAGTCCAGTCAGAGCCTTTTAAATAGTAGCAATCAAAAGAACTATTTGGCCTGGTACCAGCAGAAACCAGGACAGTCTCCTAAACTTCTGGTATACTTTGCATCCACTAGGGAATCTGGGGTCCCTGATCGCTTCATAGGCAGTGGATCTGGGACAGATTTCACTCTTACCATCAGCAGTGTGCAGGCTGAAGACCTGGCAGATTACTTCTGTCAGCAACATTATAGCACTCCTCC |
| IGKV4-61*01_SJL | SJL/J | Musmus IGKV4-61*01 F | 100 | 286 | CAAATTGTTCTCACCCAGTCTCCAGCAATCATGTCTGCATCTCCAGGGGAGAAGGTCACCATATCCTGCAGTGCCAGCTCAAGTGTAAGTTACATGTACTGGTACCAGCAGAAGCCAGGATCCTCCCCCAAACCCTGGATTTATCGCACATCCAACCTGGCTTCTGGAGTCCCTGCTCGCTTCAGTGGCAGTGGGTCTGGGACCTCTTACTCTCTCACAATCAGCAGCATGGAGGCTGAAGATGCTGCCACTTATTACTGCCAGCAGTATCATAGTTACCCACCCA |

| | | | | | |
|---|---|---|---|---|---|
| IGKV6-25*01_SJL | SJL/J | Musmus IGKV6-25*01 F | 100 | 287 | GACATTGTGATGACCCAGTCTCACAAATTCATGTCCACATCAGTAGGAGACAGGGTCAGCATCACCTGCAAGGCCAGTCAGGATGTGAGTACTGCTGTAGCCTGGTATCAACAAAAACCAGGGCAATCTCCTAAACTACTGATTTACTGGGCATCCACCCGGCACACTGGAGTCCCTGATCGCTTCACAGGCAGTGGATCTGGGACAGATTATACTCTCACCATCAGCAGTGTGCAGGCTGAAGACCTGGCACTTTATTACTGTCAGCAACATTATAGCACTCCTCC |
| IGKV4-79*01_SJL | SJL/J | Musmus IGKV4-79*01 F | 100 | 292 | CAAATTGTTCTCACCCAGTCTCCAGCAATCATGTCTGCATCTCCTGGGGAGAAGGTCACCTTGACCTGCAGTGCCAGCTCAAGTGTAAGTTCCAGCTACTTGTACTGGTACCAGCAGAAGCCAGGATCCTCCCCCAAACTCTGGATTTATAGCACATCCAACCTGGCTTCTGGAGTCCCTGCTCGCTTCAGTGGCAGTGGGTCTGGGACCTCTTACTCTCTCACAATCAGCAGCATGGAGGCTGAAGATGCTGCCTCTTATTTCTGCCATCAGTGGAGTAGTTACCCACCCC |
| IGKV4-72*01_S3540_SJL | SJL/J | Musmus IGKV4-72*01 F | 100 | 286 | CAAATTGTTCTCTCCCAGTCTCCAGCAATCCTGTCTGCATCTCCAGGGGAGAAGGTCACAATGACTTGCAGGGCCAGCTCAAGTGTAAGTTACATGCACTGGTACCAGCAGAAGCCAGGATCCTCCCCCAAACCCTGGATTTATGCCACATCCAACCTGGCTTCTGGAGTCCCTGCTCGCTTCAGTGGCAGTGGGTCTGGGACCTCTTACTCTCTCACAATCAGCAGAGTGGAGGCTGAAGATGCTGCCACTTATTACTGCCAGCAGTGGAGTAGTAACCCACTCA |
| IGKV9-120*01_S2628_SJL | SJL/J | Musmus IGKV9-120*02 F | 100 | 285 | GACATCCAGATGACCCAGTCTCCATCCTCCTTATCTGCCTCTCTGGGAGAAAGAGTCAGTCTCACTTGTCGGGCAAGTCAGGACATTGGTAGTAGCTTAAACTGGCTTCAGCAGGAACCAGATGGAACTATTAAACGCCTGATCTACGCCACATCCAGTTTAGATTCTGGTGTCCCCAAAAGGTTCAGTGGCAGTAGGTCTGGGTCAGATTATTCTCTCACCATCAGCAGCCTTGAGTCTGAAGATTTTGTAGACTATTACTGTCTACAATATGCTAGTTCTCCT |
| IGKV17-121*01_SJL | SJL/J | Musmus IGKV17-121*01 F | 100 | 287 | GAAACAACTGTGACCCAGTCTCCAGCATCCCTGTCCATGGCTATAGGAGAAAAAGTCACCATCAGATGCATAACCAGCACTGATATTGATGATGATATGAACTGGTACCAGCAGAAGCCAGGGGAACCTCCTAAGCTCCTTATTTCAGAAGGCAATACTCTTCGTCCTGGAGTCCCATCCCGATTCTCCAGCAGTGGCTATGGTACAGATTTTGTTTTTACAATTGAAAACATGCTCTCAGAAGATGTTGCAGATTACTACTGTTTGCAAAGTGATAACTTGCCTCT |
| IGKV19-93*01_SJL | SJL/J | Musmus IGKV19-93*01 F | 100 | 287 | GACATCCAGATGACACAGTCTCCATCCTCACTGTCTGCATCTCTGGGAGGCAAAGTCACCATCACTTGCAAGGCAAGCCAAGACATTAACAAGTATATAGCTTGGTACCAACACAAGCCTGGAAAAGGTCCTAGGCTGCTCATACATTACACATCTACATTACAGCCAGGCATCCCATCAAGGTTCAGTGGAAGTGGGTCTGGGAGAGATTATTCCTTCAGCATCAGCAACCTGGAGCCTGAAGATATTGCAACTTATTATTGTCTACAGTATGATAATCTTCTACC |
| IGKV13-84*01_SJL | SJL/J | Musmus IGKV13-84*01 F | 100 | 287 | GACATCCAGATGACACAATCTTCATCCTCCTTTTCTGTATCTCTAGGAGACAGAGTCACCATTACTTGCAAGGCAAGTGAGGACATATATAATCGGTTAGCCTGGTATCAGCAGAAACCAGGAAATGCTCCTAGGCTCTTAATATCTGGTGCAACCAGTTTGGAAACTGGGGTTCCTTCAAGATTCAGTGGCAGTGGATCTGGAAAGGATTACACTCTCAGCATTACCAGTCTTCAGACTGAAGATGTTGCTACTTATTACTGTCAACAGTATTGGAGTACTCCTCC |
| IGKV8-34*01_SJL | SJL/J | Musmus IGKV8-34*01 F | 100 | 305 | GACATTTTGATGACCCAGTCTCCATCCTCCCTGACTGTGTCAGCAGGAGAGAAGGTCACTATGAGCTGCAAGTCCAGTCAGAGTCTTTTAGCTAGTGGCAACCAAAATAACTACTTGGCCTGGCACCAGCAGAAACCAGGACGATCTCCTAAAATGCTGATAATTTGGGCATCCACTAGGGTATCTGGAGTCCCTGATCGCTTCATAGGCAGTGGATCTGGGACGGATTTCACTCTGACCATCAACAGTGTGCAGGCTGAAGATCTGGCTGTTTATTACTGTCAGCAGTCCTACAGCGCTCCTAC |

| | | | | | |
|---|---|---|---|---|---|
| IGKV4-92*01_SJL | SJL/J | Musmus IGKV4-92*01 F | 100 | 291 | GAAATGGTTCTCACCCAGTCTCCAGTATCCATAACTGCATCTCGAGGGGAGAAGGTCACCATCACCTGCCGTGCCAGCTCAAGTATAAGTTCCAATTACTTACACTGGTACCAGCAGAAGCCAGGATCCTCCCCTAAACTTTTGATTTATAGGACATCCATCCTGGCATCTGGAGTCCTAGACAGCTTCAGTGGCAGTGGGTCTGAGAGCTCTTACACTCTGACAATCAGCTGCATGCAGGACGAAGTTGCTGCCACTTACTATTGTCAGCAGGGGAGTAGTAGCCCACCA |
| IGKV2-112*01_SJL | SJL/J | Musmus IGKV2-112*01 F | 100 | 300 | GATATTGTGATAACCCAGGATGAACTCTCCAATCCTGTCACTTCTGGAGAATCAGTTTCCATCTCCTGCAGGTCTAGTAAGAGTCTCCTATATAAGGATGGGAAGACATACTTGAATTGGTTTCTGCAGAGACCAGGACAATCTCCTCAGCTCCTGATCTATTTGATGTCCACCCGTGCATCAGGAGTCTCAGACCGGTTTAGTGGCAGTGGGTCAGGAACAGATTTCACCCTGGAAATCAGTAGAGTGAAGGCTGAGGATGTGGGTGTGTATTACTGTCAACAACTTGTAGAGTATCCT |
| IGKV16-104*01_SJL | SJL/J | Musmus IGKV16-104*01 F | 100 | 288 | GATGTCCAGATAACCCAGTCTCCATCTTATCTTGCTGCATCTCCTGGAGAAACCATTACTATTAATTGCAGGGCAAGTAAGAGCATTAGCAAATATTTAGCCTGGTATCAAGAGAAACCTGGGAAAACTAATAAGCTTCTTATCTACTCTGGATCCACTTTGCAATCTGGAATTCCATCAAGGTTCAGTGGCAGTGGATCTGGTACAGATTTCACTCTCACCATCAGTAGCCTGGAGCCTGAAGATTTTGCAATGTATTACTGTCAACAGCATAATGAATACCCGTAC |
| IGKV14-100*01_SJL | SJL/J | Musmus IGKV14-100*01 F | 100 | 287 | GACATCCTGATGACCCAATCTCCATCCTCCATGTCTGTATCTCTGGGAGACACAGTCAGCATCACTTGCCATGCAAGTCAGGGCATTAGCAGTAATATAGGGTGGTTGCAGCAGAAACCAGGGAAATCATTTAAGGGCCTGATCTATCATGGAACCAACTTGGAAGATGGAGTTCCATCAAGGTTCAGTGGCAGTGGATCTGGAGCAGATTATTCTCTCACCATCAGCAGCCTGGAATCTGAAGATTTTGCAGACTATTACTGTGTACAGTATGCTCAGTTTCCTCC |
| IGKV5-48*01_SJL | SJL/J | Musmus IGKV5-48*01 F | 100 | 287 | GACATCTTGCTGACTCAGTCTCCAGCCATCCTGTCTGTGAGTCCAGGAGAAAGAGTCAGTTTCTCCTGCAGGGCCAGTCAGAGCATTGGCACAAGCATACACTGGTATCAGCAAAGAACAAATGGTTCTCCAAGGCTTCTCATAAAGTATGCTTCTGAGTCTATCTCTGGGATCCCTTCCAGGTTTAGTGGCAGTGGATCAGGGACAGATTTTACTCTTAGCATCAACAGTGTGGAGTCTGAAGATATTGCAGATTATTACTGTCAACAAAGTAATAGCTGGCCAAC |
| IGKV14-111*01_SJL | SJL/J | Musmus IGKV14-111*01 F | 100 | 287 | GACATCAAGATGACCCAGTCTCCATCTTCCATGTATGCATCTCTAGGAGAGAGAGTCACTATCACTTGCAAGGCGAGTCAGGACATTAATAGCTATTTAAGCTGGTTCCAGCAGAAACCAGGGAAATCTCCTAAGACCCTGATCTATCGTGCAAACAGATTGGTAGATGGGGTCCCATCAAGGTTCAGTGGCAGTGGATCTGGGCAAGATTATTCTCTCACCATCAGCAGCCTGGAGTATGAAGATATGGGAATTTATTATTGTCTACAGTATGATGAGTTTCCTCC |
| IGKV14-126*01_SJL | SJL/J | Musmus IGKV14-126*01 F | 100 | 287 | GACATCAAGATGACCCAGTCTCCATCCTCCATGTATGCATCGCTGGGAGAGAGAGTCACTATCACTTGCAAGGCGAGTCAGGACATTAAAAGCTATTTAAGCTGGTACCAGCAGAAACCATGGAAATCTCCTAAGACCCTGATCTATTATGCAACAAGCTTGGCAGATGGGGTCCCATCAAGATTCAGTGGCAGTGGATCTGGGCAAGATTATTCTCTAACCATCAGCAGCCTGGAGTCTGACGATACAGCAACTTATTACTGTCTACAGCATGGTGAGAGCCCTCC |
| IGKV11-125*01_SJL | SJL/J | Musmus IGKV11-125*01 F | 100 | 287 | GATGTCCAGATGATTCAGTCTCCATCCTCCCTGTCTGCATCTTTGGGAGACATAGTCACCATGACTTGCCAGGCAAGTCAGGGCACTAGCATTAATTTAAACTGGTTTCAGCAAAAACCAGGGAAAGCTCCTAAGCTCCTGATCTATGGTGCAAGCAACTTGGAAGATGGGGTCCCATCAAGGTTCAGTGGCAGTAGATATGGGACAGATTTCACTCTCACCATCAGCAGCCTGGAGGATGAAGATATGGCAACTTATTTCTGTCTACAGCATAGTTATCTCCCTCC |

| | | | | | |
|---|---|---|---|---|---|
| IGKV17-127*01_SJL | SJL/J | Musmus IGKV17-127*01 F | 100 | 287 | GAAACAACTGTGACCCAGTCTCCAGCATCCCTGTCCGTGGCTACAGGAGAAAAAGTCACTATCAGATGCATAACCAGCACTGATATTGATGATGATATGAACTGGTACCAGCAGAAGCCAGGGGAACCTCCTAAGCTCCTTATTTCAGAAGGCAATACTCTTCGTCCTGGAGTCCCATCCCGATTCTCCAGCAGTGGCTATGGCACAGATTTTGTTTTTACAATTGAAAACACGCTCTCAGAAGATGTTGCAGATTACTACTGTTTGCAAAGTGATAACATGCCTCT |
| IGKV5-39*01_SJL | SJL/J | Musmus IGKV5-39*01 F | 100 | 287 | GACATTGTGATGACTCAGTCTCCAGCCACCCTGTCTGTGACTCCAGGAGATAGAGTCTCTCTTTCCTGCAGGGCCAGCCAGAGTATTAGCGACTACTTACACTGGTATCAACAAAAATCACATGAGTCTCCAAGGCTTCTCATCAAATATGCTTCCCAATCCATCTCTGGGATCCCCTCCAGGTTCAGTGGCAGTGGATCAGGGTCAGATTTCACTCTCAGTATCAACAGTGTGGAACCTGAAGATGTTGGAGTGTATTACTGTCAAAATGGTCACAGCTTTCCTCC |
| IGKV1-133*01_SJL | SJL/J | Musmus IGKV1-133*01 F | 100 | 302 | GATGTTGTGATGACCCAGACTCCACTCACTTTGTCGGTTACCATTGGACAACCAGCCTCTATCTCTTGCAAGTCAAGTCAGAGCCTCTTATATAGTAATGGAAAAACCTATTTGAATTGGTTATTACAGAGGCCAGGCCAGTCTCCAAAGCGCCTAATCTATCTGGTGTCTAAACTGGACTCTGGAGTCCCTGACAGGTTCACTGGCAGTGGATCAGGAACAGATTTTACACTGAAAATCAGCAGAGTGGAGGCTGAGGATTTGGGAGTTTATTACTGCGTGCAAGGTACACATTTTCCTCA |
| IGKV4-69*01_SJL | SJL/J | Musmus IGKV4-69*01 F | 100 | 282 | CAAATTCTTCTCACCCAGTCTCCAGCAATCATGTCTGCATCTCCAGGGGAGAAGGTCACCATGACCTGCAGTGCCAGCTCAAGTGTAAGTTACATGCACTGGTACCAGCAGAAGCCAGGATCCTCGCCCAAACCCTGGATTTATGACACATCCAACCTGGCTTCTGGATTCCCTGCTCGCTTCAGTGGCAGTGGGTCTGGGACCTCTTACTCTCTCATAATCAGCAGCATGGAGGCTGAAGATGCTGCCACTTATTACTGCCATCAGCGGAGTAGTTACCCA |
| IGKV12-38*01_SJL | SJL/J | Musmus IGKV12-38*01 F | 100 | 287 | GACATCCAGATGACTCAGTCTCCAGCCTCCCTGGCTGCATCTGTGGGAGAAACTGTCACCATCACATGTCGAGCAAGTGAGAACATTTACTACAGTTTAGCATGGTATCAGCAGAAGCAAGGGAAATCTCCTCAGCTCCTGATCTATAATGCAAACAGCTTGGAAGATGGTGTCCCATCGAGGTTCAGTGGCAGTGGATCTGGGACACAGTATTCTATGAAGATCAACAGCATGCAGCCTGAAGATACCGCAACTTATTTCTGTAAACAGGCTTATGACGTTCCTCC |
| IGKV3-9*01_S7901_SJL | SJL/J | Musmus IGKV3-9*01 F | 91.41 | 299 | GACATTGTGCTGACCCAATCTCCAGCTTCTTTGGCTGTGTCTCTAGGGCAGAGGGCCATCATCTCCTGCAAGGCCAGCCAAAGTGTCAGTTTTGCTGGTACTAGTTTAATGCACTGGTACCAACAGAAACCAGGACAGCAACCCAAACTCCTCATCTATCGTGCATCCAACCTAGAAGCTGGGGTTCCTACCAGGTTTAGTGGCAGTGGGTCTAGGACAGACTTCACCCTCAATATCCATCCTGTGGAGGAAGATGATGCTGCAACCTATTACTGTCAGCAAAGTAGGGAATATCCTCC |
| IGKV4-52*01_S5019_SJL | SJL/J | Musmus IGKV4-52*01 ORF, or Musmus IGKV4-54*01 ORF | 100 | 286 | CAAATTGTTCTCACCCAGTCTCCAGCAATCCTGTCTGCATCTCCAGGGGAGAAGGTCACCATGACCTGCAGTGCCAGCTCAAGTGTAAGTTACATGTACAGGTACCAGCAGAAGCCAGGATCCTCACCCAAACCCTGGATTTATGGCACATCCAACCTGGCTTCTGGAGTCCCTGCTCGCTTCAGTGGCAGTGGATCTGGGACCTCTTATTCTCTCACAATCAGCAGCATGGAGGCTGAAGATGCTGCCACTTATTACTGCCAGCAATATCATAGTTACCCACTCA |
| IGKV8-23-1*01_SJL | SJL/J | Musmus IGKV8-16*01 F, or Musmus IGKV8-23-1*01 ORF | 100 | 284 | GAAATTGTGCTGACTGTGTCAGCAGGAGAGAGGGTCACTATCAGCTGCAAATCCAATCAGAATCTTTTATGGAGTGGAAACCAAAGGTACTGTTTGGTCTGGCACCAGTGGAAACCAGGGCAAACTCCTACACCGTTGATCACCTGGACATCTGATAGGTACTCTGGAGTCCCTGATCGTTTCATAGGCAGTGGATCTGTGACAGATTTCACTCTGACCATCAGCAGTGTGCAGGCTGAAGATGTGGCAGTTTATTTCTGTCAGCAGCATTTACACATTCCTCC |

| | | | | | |
|---|---|---|---|---|---|
| IGKV4-52*01_SJL | SJL/J | Musmus IGKV4-52*01 ORF, or Musmus IGKV4-54*01 ORF | 100 | 286 | CAAATTGTTCTCACCCAGTCTCCAGCAATCCTGTCTGCATCTCCAGGGGAGAAGGTCA CCATGACCTGCAGTGCCAGCTCAAGTGTAAGTTACATGTACAGGTACCAGCAGAAGCC AGGATCCTCACCCAAACCCTGGATTTATGGCACATCCAACCTGGCTTCTGGAGTCCCT GCTCGCTTCAGTGGCAGTGGATCTGGGACCTCTTATTCTCTCACAATCAGCAGCATGG AGGCTGAAGATGCTGCCACTTATTACTGCCAGCAATATCATAGTTACCCACCCA |
| IGLV1*01_129S6 | 129S1/SvImJ | Musmus IGLV1*01 F | 100 | 294 | CAGGCTGTTGTGACTCAGGAATCTGCACTCACCACATCACCTGGTGAAACAGTCACAC TCACTTGTCGCTCAAGTACTGGGGCTGTTACAACTAGTAACTATGCCAACTGGGTCCA AGAAAAACCAGATCATTTATTCACTGGTCTAATAGGTGGTACCAACAACCGAGCTCCA GGTGTTCCTGCCAGATTCTCAGGCTCCCTGATTGGAGACAAGGCTGCCCTCACCATCA CAGGGGCACAGACTGAGGATGAGGCAATATATTTCTGTGCTCTATGGTACAGCAACCA TTTC |
| IGLV2*02_129S6 | 129S1/SvImJ | Musmus IGLV2*02 F | 100 | 294 | CAGGCTGTTGTGACTCAGGAATCTGCACTCACCACATCACCTGGTGGAACAGTCATAC TCACTTGTCGCTCAAGTACTGGGGCTGTTACAACTAGTAACTATGCCAACTGGGTCCA AGAAAAACCAGATCATTTATTCACTGGTCTAATAGGTGGTACCAGCAACCGAGCTCCA GGTGTTCCTGTCAGATTCTCAGGCTCCCTGATTGGAGACAAGGCTGCCCTCACCATCA CAGGGGCACAGACTGAGGATGATGCAATGTATTTCTGTGCTCTATGGTACAGCACCCA TTTC |
| IGLV3*01_129S6 | 129S1/SvImJ | Musmus IGLV3*01 F | 100 | 313 | CAACTTGTGCTCACTCAGTCATCTTCAGCCTCTTTCTCCCTGGGAGCCTCAGCAAAACT CACGTGCACCTTGAGTAGTCAGCACAGTACGTACACCATTGAATGGTATCAGCAACAG CCACTCAAGCCTCCTAAGTATGTGATGGAGCTTAAGAAAGATGGAAGCCACAGCACA GGTGATGGGATTCCTGATCGCTTCTCTGGATCCAGCTCTGGTGCTGATCGCTACCTTAG CATTTCCAACATCCAGCCTGAAGATGAAGCAATATACATCTGTGGTGTGGGTGATACA ATTAAGGAACAATTTGTGTAAC |
| IGLV1*01_AJ | A/J | Musmus IGLV1*01 F | 100 | 294 | CAGGCTGTTGTGACTCAGGAATCTGCACTCACCACATCACCTGGTGAAACAGTCACAC TCACTTGTCGCTCAAGTACTGGGGCTGTTACAACTAGTAACTATGCCAACTGGGTCCA AGAAAAACCAGATCATTTATTCACTGGTCTAATAGGTGGTACCAACAACCGAGCTCCA GGTGTTCCTGCCAGATTCTCAGGCTCCCTGATTGGAGACAAGGCTGCCCTCACCATCA CAGGGGCACAGACTGAGGATGAGGCAATATATTTCTGTGCTCTATGGTACAGCAACCA TTTC |
| IGLV2*02_AJ | A/J | Musmus IGLV2*02 F | 100 | 294 | CAGGCTGTTGTGACTCAGGAATCTGCACTCACCACATCACCTGGTGGAACAGTCATAC TCACTTGTCGCTCAAGTACTGGGGCTGTTACAACTAGTAACTATGCCAACTGGGTCCA AGAAAAACCAGATCATTTATTCACTGGTCTAATAGGTGGTACCAGCAACCGAGCTCCA GGTGTTCCTGTCAGATTCTCAGGCTCCCTGATTGGAGACAAGGCTGCCCTCACCATCA CAGGGGCACAGACTGAGGATGATGCAATGTATTTCTGTGCTCTATGGTACAGCACCCA TTTC |
| IGLV3*01_AJ | A/J | Musmus IGLV3*01 F | 100 | 313 | CAACTTGTGCTCACTCAGTCATCTTCAGCCTCTTTCTCCCTGGGAGCCTCAGCAAAACT CACGTGCACCTTGAGTAGTCAGCACAGTACGTACACCATTGAATGGTATCAGCAACAG CCACTCAAGCCTCCTAAGTATGTGATGGAGCTTAAGAAAGATGGAAGCCACAGCACA GGTGATGGGATTCCTGATCGCTTCTCTGGATCCAGCTCTGGTGCTGATCGCTACCTTAG CATTTCCAACATCCAGCCTGAAGATGAAGCAATATACATCTGTGGTGTGGGTGATACA ATTAAGGAACAATTTGTGTAAC |
| IGLV1*01_AKR | AKR/J | Musmus IGLV1*01 F | 100 | 294 | CAGGCTGTTGTGACTCAGGAATCTGCACTCACCACATCACCTGGTGAAACAGTCACAC TCACTTGTCGCTCAAGTACTGGGGCTGTTACAACTAGTAACTATGCCAACTGGGTCCA AGAAAAACCAGATCATTTATTCACTGGTCTAATAGGTGGTACCAACAACCGAGCTCCA |

| | | | | | |
|---|---|---|---|---|---|
| | | | | | GGTGTTCCTGCCAGATTCTCAGGCTCCCTGATTGGAGACAAGGCTGCCCTCACCATCA CAGGGGCACAGACTGAGGATGAGGCAATATATTTCTGTGCTCTATGGTACAGCAACCA TTTC |
| IGLV2*02 _AKR | AKR/J | Musmus IGLV2*02 F | 100 | 294 | CAGGCTGTTGTGACTCAGGAATCTGCACTCACCACATCACCTGGTGGAACAGTCATAC TCACTTGTCGCTCAAGTACTGGGGCTGTTACAACTAGTAACTATGCCAACTGGGTCCA AGAAAAACCAGATCATTTATTCACTGGTCTAATAGGTGGTACCAGCAACCGAGCTCCA GGTGTTCCTGTCAGATTCTCAGGCTCCCTGATTGGAGACAAGGCTGCCCTCACCATCA CAGGGGCACAGACTGAGGATGATGCAATGTATTTCTGTGCTCTATGGTACAGCACCCA TTTC |
| IGLV3*01 _AKR | AKR/J | Musmus IGLV3*01 F | 100 | 313 | CAACTTGTGCTCACTCAGTCATCTTCAGCCTCTTTCTCCCTGGGAGCCTCAGCAAAACT CACGTGCACCTTGAGTAGTCAGCACAGTACGTACACCATTGAATGGTATCAGCAACAG CCACTCAAGCCTCCTAAGTATGTGATGGAGCTTAAGAAAGATGGAAGCCACAGCACA GGTGATGGGATTCCTGATCGCTTCTCTGGATCCAGCTCTGGTGCTGATCGCTACCTTAG CATTTCCAACATCCAGCCTGAAGATGAAGCAATATACATCTGTGGTGTGGGTGATACA ATTAAGGAACAATTTGTGTAAC |
| IGLV1*01 _B6 | C57BL/6J | Musmus IGLV1*01 F | 100 | 294 | CAGGCTGTTGTGACTCAGGAATCTGCACTCACCACATCACCTGGTGAAACAGTCACAC TCACTTGTCGCTCAAGTACTGGGGCTGTTACAACTAGTAACTATGCCAACTGGGTCCA AGAAAAACCAGATCATTTATTCACTGGTCTAATAGGTGGTACCAACAACCGAGCTCCA GGTGTTCCTGCCAGATTCTCAGGCTCCCTGATTGGAGACAAGGCTGCCCTCACCATCA CAGGGGCACAGACTGAGGATGAGGCAATATATTTCTGTGCTCTATGGTACAGCAACCA TTTC |
| IGLV2*02 _B6 | C57BL/6J | Musmus IGLV2*02 F | 100 | 294 | CAGGCTGTTGTGACTCAGGAATCTGCACTCACCACATCACCTGGTGGAACAGTCATAC TCACTTGTCGCTCAAGTACTGGGGCTGTTACAACTAGTAACTATGCCAACTGGGTCCA AGAAAAACCAGATCATTTATTCACTGGTCTAATAGGTGGTACCAGCAACCGAGCTCCA GGTGTTCCTGTCAGATTCTCAGGCTCCCTGATTGGAGACAAGGCTGCCCTCACCATCA CAGGGGCACAGACTGAGGATGATGCAATGTATTTCTGTGCTCTATGGTACAGCACCCA TTTC |
| IGLV3*01 _B6 | C57BL/6J | Musmus IGLV3*01 F | 100 | 313 | CAACTTGTGCTCACTCAGTCATCTTCAGCCTCTTTCTCCCTGGGAGCCTCAGCAAAACT CACGTGCACCTTGAGTAGTCAGCACAGTACGTACACCATTGAATGGTATCAGCAACAG CCACTCAAGCCTCCTAAGTATGTGATGGAGCTTAAGAAAGATGGAAGCCACAGCACA GGTGATGGGATTCCTGATCGCTTCTCTGGATCCAGCTCTGGTGCTGATCGCTACCTTAG CATTTCCAACATCCAGCCTGAAGATGAAGCAATATACATCTGTGGTGTGGGTGATACA ATTAAGGAACAATTTGTGTAAC |
| IGLV1*01 _BALB | BALB/cByJ | Musmus IGLV1*01 F | 100 | 294 | CAGGCTGTTGTGACTCAGGAATCTGCACTCACCACATCACCTGGTGAAACAGTCACAC TCACTTGTCGCTCAAGTACTGGGGCTGTTACAACTAGTAACTATGCCAACTGGGTCCA AGAAAAACCAGATCATTTATTCACTGGTCTAATAGGTGGTACCAACAACCGAGCTCCA GGTGTTCCTGCCAGATTCTCAGGCTCCCTGATTGGAGACAAGGCTGCCCTCACCATCA CAGGGGCACAGACTGAGGATGAGGCAATATATTTCTGTGCTCTATGGTACAGCAACCA TTTC |
| IGLV2*02 _BALB | BALB/cByJ | Musmus IGLV2*02 F | 100 | 294 | CAGGCTGTTGTGACTCAGGAATCTGCACTCACCACATCACCTGGTGGAACAGTCATAC TCACTTGTCGCTCAAGTACTGGGGCTGTTACAACTAGTAACTATGCCAACTGGGTCCA AGAAAAACCAGATCATTTATTCACTGGTCTAATAGGTGGTACCAGCAACCGAGCTCCA GGTGTTCCTGTCAGATTCTCAGGCTCCCTGATTGGAGACAAGGCTGCCCTCACCATCA |

| | | | | | |
|---|---|---|---|---|---|
| | | | | | CAGGGGCACAGACTGAGGATGATGCAATGTATTTCTGTGCTCTATGGTACAGCACCCA TTTC |
| IGLV3*01 _BALB | BALB/cByJ | Musmus IGLV3*01 F | 100 | 313 | CAACTTGTGCTCACTCAGTCATCTTCAGCCTCTTTCTCCCTGGGAGCCTCAGCAAAACT CACGTGCACCTTGAGTAGTCAGCACAGTACGTACACCATTGAATGGTATCAGCAACAG CCACTCAAGCCTCCTAAGTATGTGATGGAGCTTAAGAAAGATGGAAGCCACAGCACA GGTGATGGGATTCCTGATCGCTTCTCTGGATCCAGCTCTGGTGCTGATCGCTACCTTAG CATTTCCAACATCCAGCCTGAAGATGAAGCAATATACATCTGTGGTGTGGGTGATACA ATTAAGGAACAATTTGTGTAAC |
| IGLV1*01 _C3H | C3H/HeJ | Musmus IGLV1*01 F | 100 | 294 | CAGGCTGTTGTGACTCAGGAATCTGCACTCACCACATCACCTGGTGAAACAGTCACAC TCACTTGTCGCTCAAGTACTGGGGCTGTTACAACTAGTAACTATGCCAACTGGGTCCA AGAAAAACCAGATCATTTATTCACTGGTCTAATAGGTGGTACCAACAACCGAGCTCCA GGTGTTCCTGCCAGATTCTCAGGCTCCCTGATTGGAGACAAGGCTGCCCTCACCATCA CAGGGGCACAGACTGAGGATGAGGCAATATATTTCTGTGCTCTATGGTACAGCAACCA TTTC |
| IGLV2*02 _C3H | C3H/HeJ | Musmus IGLV2*02 F | 100 | 294 | CAGGCTGTTGTGACTCAGGAATCTGCACTCACCACATCACCTGGTGGAACAGTCATAC TCACTTGTCGCTCAAGTACTGGGGCTGTTACAACTAGTAACTATGCCAACTGGGTCCA AGAAAAACCAGATCATTTATTCACTGGTCTAATAGGTGGTACCAGCAACCGAGCTCCA GGTGTTCCTGTCAGATTCTCAGGCTCCCTGATTGGAGACAAGGCTGCCCTCACCATCA CAGGGGCACAGACTGAGGATGATGCAATGTATTTCTGTGCTCTATGGTACAGCACCCA TTTC |
| IGLV3*01 _C3H | C3H/HeJ | Musmus IGLV3*01 F | 100 | 313 | CAACTTGTGCTCACTCAGTCATCTTCAGCCTCTTTCTCCCTGGGAGCCTCAGCAAAACT CACGTGCACCTTGAGTAGTCAGCACAGTACGTACACCATTGAATGGTATCAGCAACAG CCACTCAAGCCTCCTAAGTATGTGATGGAGCTTAAGAAAGATGGAAGCCACAGCACA GGTGATGGGATTCCTGATCGCTTCTCTGGATCCAGCTCTGGTGCTGATCGCTACCTTAG CATTTCCAACATCCAGCCTGAAGATGAAGCAATATACATCTGTGGTGTGGGTGATACA ATTAAGGAACAATTTGTGTAAC |
| IGLV5*01 _CAST | CAST/EiJ | Musmus IGLV5*01 [F] | 93.88 | 245 | TTACTCAGCCAAGCTCTGTGTCTACGTCTCTAGGAAGCACAGCCAAACTGCCTTGCAA GGCCAGCACTGGTAACATTGGAGACAGCTATGTGAACTGGTACCAGCAGTACATGGG AAGATCTCCCACTAATATAATCTATGGAGATGATCTCCGACCATCTGGAGTTTCTGAT AGGTTCTCTGGCTCCATTGACAGCTCTTCCAACTCAGCCTTCCTGACAATCCAAAATGT GCAGGCTGATGAT |
| IGLV3*01 _CAST | CAST/EiJ | Musspr IGLV3*01 F | 96.26 | 313 | CAACCTGTGCTCACTCAGTCATCTTCAGCCTCTTTCTCCCTGGGAGCCTCAGCAAAACT CACATGCACCTTGAGTAGTGAGCACAGTACGTACATCATTGAATGGTACCAGCAACAG CCACTCAAGCCTCCTAAGTATGTGATGCAGCTTAAGAAAGATGGAAGCCACAGCAAG GGTGATGGAATCCCTGATCGCTTCTCTGGATCCAGCTCTGGTGCTGACCGCTACCTTAG CATCTCCAACATCCAGCCTGAAGATGAAGCAATATACATCTGTGGTGTGAGTGGAACA ATTAAGGAACAATTCGTGTAAC |
| IGLV1*01 _CAST_2 | CAST/EiJ | Musspr IGLV2*01 F | 97.57 | 294 | CAGGCTGTTGTGACTCAGGAGTCTGTTCTCACCACATCACCTGGTGGTACAGTCATAA TCACTTGTCGCTCAAGTACTGGGGCTGTTACAACTAGTAACTATGCCATTTGGGTCCA AGAAAAAGCAGATCATTTATTCGCTGGTGTAATAGGTGATACAAGCAACCGAGCTCC AGGTGTTCCTGCCAGATTCTCAGGCTCCTTGATTGGAGACAAGGCTGCCCTCACCATC ACAGGGGCACAGACTGAGGATGAGGCAATATATTTCTGTGCTCTATGGTACAGCAACC ATTTC |

| | | | | | |
|---|---|---|---|---|---|
| IGLV1*01 _CAST | CAST/EiJ | Musmus IGLV1*01 F | 97.92 | 294 | CAGGCTGTCGTGACTCAGGAATCTGCACTCACCACATCACCTGGTGAAACAGTCACACTCACTTGTCGCTCAAGTACTGGGGCTGTTACAACTAGTAACTATGCCAACTGGGTCCAAGAAAAAACAGACCATTTATTCACTGGTCTAATAGGTGGTACCAACAACCGAGCTCCAGGTGTTCCTGTCAGATTCTCAGGCTCCCTGATTGGAGACAAGGCTGCCCTCACCATCACAGGGGCACAGACTGAGGATGATGCAATGTATTTCTGTGCTCTATGGTACAGCAACCATTTC |
| IGLV6*01 _CAST | CAST/EiJ | Musmus IGLV6*01 [F] | 98.78 | 245 | TTACTCAGCCAAGCTCTGTGTCTACATCTCTAGGAAGCACAGTCAAACTGCCTTGCAAGCGCAGCACTGATAATATTGGAAGCTACTATGTGCATTGGTACCAGCAACATATGGGAAGATCTCCTACCAATATGATCCATAGTGATGATCAGCGACCATCTGGAGTTTCTGATAGGTTCTCTGGCTCCATTGACAGCTCTTCCAACTCAGCCTTCCTGACAATCAATAATGTTCAGGCTGAGGAT |
| IGLV2*02 _CAST | CAST/EiJ | Musmus IGLV2*02 F | 99.65 | 294 | CAGGCTGTTGTGACTCAGGAATCTGCACTCACCACATCACCTGGTGGAACAGTCATACTCACTTGTCGCTCAAGTACTGGGGCTGTTACAACTAGTAACTATGCCAACTGGGTCCAAGAAAAACCAGATCATTTATTCACTGGTCTAATAGGTGGTACCAGCAACCGAGCTCCAGGTGTTCCAGTCAGATTCTCAGGCTCCCTGATTGGAGACAAGGCTGCCCTCACCATCACAGGGGCACAGACTGAGGATGATGCAATGTATTTCTGTGCTCTATGGTACAGCACCCATTTC |
| IGLV4*01 _CAST | CAST/EiJ | Musmus IGLV4*01 [F] | 100 | 245 | TTACTCAGCCAAGCTCTGTGTCTACGTCTCTAGGAAGCACAGTCAAACTGTCTTGCAAGCGCAGCACTGGTAACATTGGAAACAACTATGTGCACTGGTACCAGCAGTACATGGGAAGATCTCCCACCAATATGATCTATGATGATAATAAGCGACCATCTGGAGTTTCTGATAGGTTCTCTGGCTCCATTGACAGCTCTTCCAACTCAGCCTTCCTGACAATCAATAATGTGCAGGCTGAGGAT |
| IGLV8*01 _CAST | CAST/EiJ | Musmus IGLV8*01 [F] | 100 | 245 | TTACTCAGCCAAGCTCTGTGTCTACGTCTCTAGGAAGCACAGTCAAACTGCCTTGCAAGCGCAGCACTGGTAACATTGGAAACGACTATGTGCACTGGTACCAGCAACACATGGGAAGATCTCCCACCAATATGATCTATAGAGATGATCAGCGACCATCTGGAGTTTCTGATAGGTTCTCTGGCTCCATTGACAGCTCTTCCAACTCAGCCTTCCTGACAATCAATAATGTGCAGGCTGAGGAT |
| IGLV7*01 _CAST | CAST/EiJ | Musmus IGLV7*01 [F] | 100 | 245 | TTACTCAGCCAAGCTCTGTGTCTACGTCTCTAGGAAGCACAGTCAAACTGTCTTGCAAGCCCAGCACTGGTAAAATTGGAAATTACTTTATGAGCTGGTACCAGCAACACATGGGAAGATCTCCCACAAATATGATCTATAGAGATGATCTCCGACCATCTGGAGTTTCTGATAGGTTCTCTGGCTCCATTGACAGCTCTTCCAACTCAGCCTTCCTGACAATCAATAATGTGCAGGCTGAGGAT |
| IGLV3*01 _CBA | CBA/J | Musmus IGLV3*01 F | 100 | 313 | CAACTTGTGCTCACTCAGTCATCTTCAGCCTCTTTCTCCCTGGGAGCCTCAGCAAAACTCACGTGCACCTTGAGTAGTCAGCACAGTACGTACACCATTGAATGGTATCAGCAACAGCCACTCAAGCCTCCTAAGTATGTGATGGAGCTTAAGAAAGATGGAAGCCACAGCACAGGTGATGGGATTCCTGATCGCTTCTCTGGATCCAGCTCTGGTGCTGATCGCTACCTTAGCATTTCCAACATCCAGCCTGAAGATGAAGCAATATACATCTGTGGTGTGGGTGATACAATTAAGGAACAATTTGTGTAAC |
| IGLV1*01 _CBA | CBA/J | Musmus IGLV1*01 F | 100 | 294 | CAGGCTGTTGTGACTCAGGAATCTGCACTCACCACATCACCTGGTGAAACAGTCACACTCACTTGTCGCTCAAGTACTGGGGCTGTTACAACTAGTAACTATGCCAACTGGGTCCAAGAAAAACCAGATCATTTATTCACTGGTCTAATAGGTGGTACCAACAACCGAGCTCCAGGTGTTCCTGCCAGATTCTCAGGCTCCCTGATTGGAGACAAGGCTGCCCTCACCATCACAGGGGCACAGACTGAGGATGAGGCAATATATTTCTGTGCTCTATGGTACAGCAACCATTTC |

| | | | | | |
|---|---|---|---|---|---|
| IGLV2*02 _CBA | CBA/J | Musmus IGLV2*02 F | 100 | 294 | CAGGCTGTTGTGACTCAGGAATCTGCACTCACCACATCACCTGGTGGAACAGTCATAC TCACTTGTCGCTCAAGTACTGGGGCTGTTACAACTAGTAACTATGCCAACTGGGTCCA AGAAAAACCAGATCATTTATTCACTGGTCTAATAGGTGGTACCAGCAACCGAGCTCCA GGTGTTCCTGTCAGATTCTCAGGCTCCCTGATTGGAGACAAGGCTGCCCTCACCATCA CAGGGGCACAGACTGAGGATGATGCAATGTATTTCTGTGCTCTATGGTACAGCACCCA TTTC |
| IGLV1*01 _DBA1 | DBA/1J | Musmus IGLV1*01 F | 100 | 294 | CAGGCTGTTGTGACTCAGGAATCTGCACTCACCACATCACCTGGTGAAACAGTCACAC TCACTTGTCGCTCAAGTACTGGGGCTGTTACAACTAGTAACTATGCCAACTGGGTCCA AGAAAAACCAGATCATTTATTCACTGGTCTAATAGGTGGTACCAACAACCGAGCTCCA GGTGTTCCTGCCAGATTCTCAGGCTCCCTGATTGGAGACAAGGCTGCCCTCACCATCA CAGGGGCACAGACTGAGGATGAGGCAATATATTTCTGTGCTCTATGGTACAGCAACCA TTTC |
| IGLV2*02 _DBA1 | DBA/1J | Musmus IGLV2*02 F | 100 | 294 | CAGGCTGTTGTGACTCAGGAATCTGCACTCACCACATCACCTGGTGGAACAGTCATAC TCACTTGTCGCTCAAGTACTGGGGCTGTTACAACTAGTAACTATGCCAACTGGGTCCA AGAAAAACCAGATCATTTATTCACTGGTCTAATAGGTGGTACCAGCAACCGAGCTCCA GGTGTTCCTGTCAGATTCTCAGGCTCCCTGATTGGAGACAAGGCTGCCCTCACCATCA CAGGGGCACAGACTGAGGATGATGCAATGTATTTCTGTGCTCTATGGTACAGCACCCA TTTC |
| IGLV3*01 _DBA1 | DBA/1J | Musmus IGLV3*01 F | 100 | 313 | CAACTTGTGCTCACTCAGTCATCTTCAGCCTCTTTCTCCCTGGGAGCCTCAGCAAAACT CACGTGCACCTTGAGTAGTCAGCACAGTACGTACACCATTGAATGGTATCAGCAACAG CCACTCAAGCCTCCTAAGTATGTGATGGAGCTTAAGAAAGATGGAAGCCACAGCACA GGTGATGGGATTCCTGATCGCTTCTCTGGATCCAGCTCTGGTGCTGATCGCTACCTTAG CATTTCCAACATCCAGCCTGAAGATGAAGCAATATACATCTGTGGTGTGGGTGATACA ATTAAGGAACAATTTGTGTAAC |
| IGLV1*01 _DBA2 | DBA/2J | Musmus IGLV1*01 F | 100 | 294 | CAGGCTGTTGTGACTCAGGAATCTGCACTCACCACATCACCTGGTGAAACAGTCACAC TCACTTGTCGCTCAAGTACTGGGGCTGTTACAACTAGTAACTATGCCAACTGGGTCCA AGAAAAACCAGATCATTTATTCACTGGTCTAATAGGTGGTACCAACAACCGAGCTCCA GGTGTTCCTGCCAGATTCTCAGGCTCCCTGATTGGAGACAAGGCTGCCCTCACCATCA CAGGGGCACAGACTGAGGATGAGGCAATATATTTCTGTGCTCTATGGTACAGCAACCA TTTC |
| IGLV2*02 _DBA2 | DBA/2J | Musmus IGLV2*02 F | 100 | 294 | CAGGCTGTTGTGACTCAGGAATCTGCACTCACCACATCACCTGGTGGAACAGTCATAC TCACTTGTCGCTCAAGTACTGGGGCTGTTACAACTAGTAACTATGCCAACTGGGTCCA AGAAAAACCAGATCATTTATTCACTGGTCTAATAGGTGGTACCAGCAACCGAGCTCCA GGTGTTCCTGTCAGATTCTCAGGCTCCCTGATTGGAGACAAGGCTGCCCTCACCATCA CAGGGGCACAGACTGAGGATGATGCAATGTATTTCTGTGCTCTATGGTACAGCACCCA TTTC |
| IGLV3*01 _DBA2 | DBA/2J | Musmus IGLV3*01 F | 100 | 313 | CAACTTGTGCTCACTCAGTCATCTTCAGCCTCTTTCTCCCTGGGAGCCTCAGCAAAACT CACGTGCACCTTGAGTAGTCAGCACAGTACGTACACCATTGAATGGTATCAGCAACAG CCACTCAAGCCTCCTAAGTATGTGATGGAGCTTAAGAAAGATGGAAGCCACAGCACA GGTGATGGGATTCCTGATCGCTTCTCTGGATCCAGCTCTGGTGCTGATCGCTACCTTAG CATTTCCAACATCCAGCCTGAAGATGAAGCAATATACATCTGTGGTGTGGGTGATACA ATTAAGGAACAATTTGTGTAAC |
| IGLV3*01 _LEWES | LEWES/EiJ | Musspr IGLV3*01 F | 97.28 | 313 | CAACCTGTGCTCACTCAGTCATCTTCAGCCTCTTTCTCCCTGGGAGCCTCAGCAAAACT CACATGCACCTTGAGTAGTGAGCACAGTACGTACTTCATTGAATGGTACCAGCAACAG |

| | | | | | |
|---|---|---|---|---|---|
| | | | | | CCACTCAAGCCTCCTAAGTATGTGATGCAGCTTAAGAAAGATGGAAGCCACAGCAAG<br>GGTGATGGGATCCCTGATCGCTTCTCTGGATCCAGCTCTGGTGCTGATCGCTACCTTAG<br>CATCTCCAACATCCAGCCTGAAGATGAAGCAATATACATCTGTGGTGTGGGTGAAACA<br>ATTAAGGAACGATACGTGTAAC |
| IGLV1*01<br>_LEWES_<br>2 | LEWES/EiJ | Musspr IGLV2*01 F | 97.92 | 294 | CAGGCTGTTGTGACTCAGGAGTCTGTACTCACCACATCACCTGGTGGTACAGTCATAA<br>TCACTTGTCGCTCAAGTACTGGGGCTGTTACAACTAGTAACTATGCCATTTGGGTCCA<br>AGAAAAAGCAGATCATTTATTCGCTGGTGTAATAGGTGATACAAGCAACCGAGCTCC<br>AGGTGTTCCTGCCAGATTCTCAGGCTCCTTGATTGGAGACAAGGCTGCCCTCACCATC<br>ACAGGGGCACAGACTGAGGATGAGGCAATATATTTCTGTGCTCTATGGTACAGCAACC<br>ATTTC |
| IGLV1*01<br>_LEWES | LEWES/EiJ | Musmus IGLV1*01 F | 99.65 | 294 | CAGGCTGTTGTGACTCAGGAATCTGCACTCACCACATCACCTGGTGAAACAGTCACAC<br>TCACTTGTCGCTCAAGTACTGGGGCTGTTACAACTAGTAACTATGCCAACTGGGTCCA<br>AGAAAAACCAGATCATTTATTCACTGGTCTAATAGGTGGTACCAACAACCGAGCTCCA<br>GGTGTTCCTGCCAGATTCTCAGGCTCCCTGATTGGAGACAAGGCTGCCCTCACCATCA<br>CAGGGGCACAGACTGAGGATGAAGCAATATATTTCTGTGCTCTATGGTACAGCAACCA<br>TTTC |
| IGLV2*02<br>_LEWES | LEWES/EiJ | Musmus IGLV2*02 F | 100 | 294 | CAGGCTGTTGTGACTCAGGAATCTGCACTCACCACATCACCTGGTGGAACAGTCATAC<br>TCACTTGTCGCTCAAGTACTGGGGCTGTTACAACTAGTAACTATGCCAACTGGGTCCA<br>AGAAAAACCAGATCATTTATTCACTGGTCTAATAGGTGGTACCAGCAACCGAGCTCCA<br>GGTGTTCCTGTCAGATTCTCAGGCTCCCTGATTGGAGACAAGGCTGCCCTCACCATCA<br>CAGGGGCACAGACTGAGGATGATGCAATGTATTTCTGTGCTCTATGGTACAGCACCCA<br>TTTC |
| IGLV3*01<br>_MRL | MRL/MpJ | Musmus IGLV3*01 F | 100 | 313 | CAACTTGTGCTCACTCAGTCATCTTCAGCCTCTTTCTCCCTGGGAGCCTCAGCAAAACT<br>CACGTGCACCTTGAGTAGTCAGCACAGTACGTACACCATTGAATGGTATCAGCAACAG<br>CCACTCAAGCCTCCTAAGTATGTGATGGAGCTTAAGAAAGATGGAAGCCACAGCACA<br>GGTGATGGGATTCCTGATCGCTTCTCTGGATCCAGCTCTGGTGCTGATCGCTACCTTAG<br>CATTTCCAACATCCAGCCTGAAGATGAAGCAATATACATCTGTGGTGTGGGTGATACA<br>ATTAAGGAACAATTTGTGTAAC |
| IGLV1*01<br>_MRL | MRL/MpJ | Musmus IGLV1*01 F | 100 | 294 | CAGGCTGTTGTGACTCAGGAATCTGCACTCACCACATCACCTGGTGAAACAGTCACAC<br>TCACTTGTCGCTCAAGTACTGGGGCTGTTACAACTAGTAACTATGCCAACTGGGTCCA<br>AGAAAAACCAGATCATTTATTCACTGGTCTAATAGGTGGTACCAACAACCGAGCTCCA<br>GGTGTTCCTGCCAGATTCTCAGGCTCCCTGATTGGAGACAAGGCTGCCCTCACCATCA<br>CAGGGGCACAGACTGAGGATGAGGCAATATATTTCTGTGCTCTATGGTACAGCAACCA<br>TTTC |
| IGLV2*02<br>_MRL | MRL/MpJ | Musmus IGLV2*02 F | 100 | 294 | CAGGCTGTTGTGACTCAGGAATCTGCACTCACCACATCACCTGGTGGAACAGTCATAC<br>TCACTTGTCGCTCAAGTACTGGGGCTGTTACAACTAGTAACTATGCCAACTGGGTCCA<br>AGAAAAACCAGATCATTTATTCACTGGTCTAATAGGTGGTACCAGCAACCGAGCTCCA<br>GGTGTTCCTGTCAGATTCTCAGGCTCCCTGATTGGAGACAAGGCTGCCCTCACCATCA<br>CAGGGGCACAGACTGAGGATGATGCAATGTATTTCTGTGCTCTATGGTACAGCACCCA<br>TTTC |
| IGLV5*01<br>_MSM | MSM/MsJ | Musmus IGLV5*01 [F] | 93.88 | 245 | TTACTCAGCCAAGCTCTGTGTCTACGTCTCTAGGAAGCACAGCCAAACTGCCTTGCAA<br>GGCCAGCACTGGTAACATTGGAGACAGCTATGTGAACTGGTACCAGCAGTACATGGG<br>AAGATCTCCCACTAATATGATCTATGGAGATGATCTCCGACCATCTGGAGTTTCTGAT |

| | | | | | |
|---|---|---|---|---|---|
| | | | | | AGGTTCTCTGGCTCCATTCACAGCTCTTCCAACTCAGCCTTCCTGACAATCCAAAATGT GCAGGCTGATGAT |
| IGLV6*03 _MSM | MSM/MsJ | Musmus IGLV6*03 [F] | 99.59 | 245 | TTACTCAGCCAAGCTCTGTGTCTACATCTCTAGGAAGCACAGTCAAACTGCCTTGCAA GCGCAGCACTGGAAATATTGGAAGCTACTATGTGCATTGGTACCAGCAACATATGGG AAGATCTCCCACCAATATGATCCATAGTGATGATCAGCGACCATCTGGAGTTTCTGAT AGGTTCTCTGGCTCCATTGACAGCTCTTCCAACTCAGCCTTCCTGACAATCAATAATGT TCAGGCTGAGGAT |
| IGLV3*01 _MSM | MSM/MsJ | Musmus IGLV3*01 F | 99.66 | 313 | CAACTTGTGCTCACTCAGTCATCTTCAGCCTCTTTCTCCCTGGGAGCCTCAGCAAAACT CACGTGCACCTTGAGTAGTCAGCACAGTACGTACACCATTGAATGGTATCAGCAACAG CCACTCAAGCCTCCTAAGTATGTGATGGAGCTTAAGAAAGATGGAAGCCACAGCACA GGTGATGGGATCCCTGATCGCTTCTCTGGATCCAGCTCTGGTGCTGATCGCTACCTTAG CATTTCCAACATCCAGCCTGAAGATGAAGCAATATACATCTGTGGTGTGGGTGATACA ATTAAGGAACAATTTGTGTAAC |
| IGLV2*02 _MSM | MSM/MsJ | Musmus IGLV2*02 F | 100 | 294 | CAGGCTGTTGTGACTCAGGAATCTGCACTCACCACATCACCTGGTGGAACAGTCATAC TCACTTGTCGCTCAAGTACTGGGGCTGTTACAACTAGTAACTATGCCAACTGGGTCCA AGAAAAACCAGATCATTTATTCACTGGTCTAATAGGTGGTACCAGCAACCGAGCTCCA GGTGTTCCTGTCAGATTCTCAGGCTCCCTGATTGGAGACAAGGCTGCCCTCACCATCA CAGGGGCACAGACTGAGGATGATGCAATGTATTTCTGTGCTCTATGGTACAGCACCCA TTTC |
| IGLV1*01 _MSM | MSM/MsJ | Musmus IGLV1*01 F | 100 | 294 | CAGGCTGTTGTGACTCAGGAATCTGCACTCACCACATCACCTGGTGAAACAGTCACAC TCACTTGTCGCTCAAGTACTGGGGCTGTTACAACTAGTAACTATGCCAACTGGGTCCA AGAAAAACCAGATCATTTATTCACTGGTCTAATAGGTGGTACCAACAACCGAGCTCCA GGTGTTCCTGCCAGATTCTCAGGCTCCCTGATTGGAGACAAGGCTGCCCTCACCATCA CAGGGGCACAGACTGAGGATGAGGCAATATATTTCTGTGCTCTATGGTACAGCAACCA TTTC |
| IGLV1*01 _NOD | NOD/ShiLtJ | Musmus IGLV1*01 F | 100 | 294 | CAGGCTGTTGTGACTCAGGAATCTGCACTCACCACATCACCTGGTGAAACAGTCACAC TCACTTGTCGCTCAAGTACTGGGGCTGTTACAACTAGTAACTATGCCAACTGGGTCCA AGAAAAACCAGATCATTTATTCACTGGTCTAATAGGTGGTACCAACAACCGAGCTCCA GGTGTTCCTGCCAGATTCTCAGGCTCCCTGATTGGAGACAAGGCTGCCCTCACCATCA CAGGGGCACAGACTGAGGATGAGGCAATATATTTCTGTGCTCTATGGTACAGCAACCA TTTC |
| IGLV2*02 _NOD | NOD/ShiLtJ | Musmus IGLV2*02 F | 100 | 294 | CAGGCTGTTGTGACTCAGGAATCTGCACTCACCACATCACCTGGTGGAACAGTCATAC TCACTTGTCGCTCAAGTACTGGGGCTGTTACAACTAGTAACTATGCCAACTGGGTCCA AGAAAAACCAGATCATTTATTCACTGGTCTAATAGGTGGTACCAGCAACCGAGCTCCA GGTGTTCCTGTCAGATTCTCAGGCTCCCTGATTGGAGACAAGGCTGCCCTCACCATCA CAGGGGCACAGACTGAGGATGATGCAATGTATTTCTGTGCTCTATGGTACAGCACCCA TTTC |
| IGLV3*01 _NOD | NOD/ShiLtJ | Musmus IGLV3*01 F | 100 | 313 | CAACTTGTGCTCACTCAGTCATCTTCAGCCTCTTTCTCCCTGGGAGCCTCAGCAAAACT CACGTGCACCTTGAGTAGTCAGCACAGTACGTACACCATTGAATGGTATCAGCAACAG CCACTCAAGCCTCCTAAGTATGTGATGGAGCTTAAGAAAGATGGAAGCCACAGCACA GGTGATGGGATTCCTGATCGCTTCTCTGGATCCAGCTCTGGTGCTGATCGCTACCTTAG CATTTCCAACATCCAGCCTGAAGATGAAGCAATATACATCTGTGGTGTGGGTGATACA ATTAAGGAACAATTTGTGTAAC |

| | | | | | |
|---|---|---|---|---|---|
| IGLV1*01 _NOR | NOR/LtJ | Musmus IGLV1*01 F | 100 | 294 | CAGGCTGTTGTGACTCAGGAATCTGCACTCACCACATCACCTGGTGAAACAGTCACAC TCACTTGTCGCTCAAGTACTGGGGCTGTTACAACTAGTAACTATGCCAACTGGGTCCA AGAAAAACCAGATCATTTATTCACTGGTCTAATAGGTGGTACCAACAACCGAGCTCCA GGTGTTCCTGCCAGATTCTCAGGCTCCCTGATTGGAGACAAGGCTGCCCTCACCATCA CAGGGGCACAGACTGAGGATGAGGCAATATATTTCTGTGCTCTATGGTACAGCAACCA TTTC |
| IGLV2*02 _NOR | NOR/LtJ | Musmus IGLV2*02 F | 100 | 294 | CAGGCTGTTGTGACTCAGGAATCTGCACTCACCACATCACCTGGTGGAACAGTCATAC TCACTTGTCGCTCAAGTACTGGGGCTGTTACAACTAGTAACTATGCCAACTGGGTCCA AGAAAAACCAGATCATTTATTCACTGGTCTAATAGGTGGTACCAGCAACCGAGCTCCA GGTGTTCCTGTCAGATTCTCAGGCTCCCTGATTGGAGACAAGGCTGCCCTCACCATCA CAGGGGCACAGACTGAGGATGATGCAATGTATTTCTGTGCTCTATGGTACAGCACCCA TTTC |
| IGLV3*01 _NOR | NOR/LtJ | Musmus IGLV3*01 F | 100 | 313 | CAACTTGTGCTCACTCAGTCATCTTCAGCCTCTTTCTCCCTGGGAGCCTCAGCAAAACT CACGTGCACCTTGAGTAGTCAGCACAGTACGTACACCATTGAATGGTATCAGCAACAG CCACTCAAGCCTCCTAAGTATGTGATGGAGCTTAAGAAAGATGGAAGCCACAGCACA GGTGATGGGATTCCTGATCGCTTCTCTGGATCCAGCTCTGGTGCTGATCGCTACCTTAG CATTTCCAACATCCAGCCTGAAGATGAAGCAATATACATCTGTGGTGTGGGTGATACA ATTAAGGAACAATTTGTGTAAC |
| IGLV1*01 _NZB | NZB/BlNJ | Musmus IGLV1*01 F | 100 | 294 | CAGGCTGTTGTGACTCAGGAATCTGCACTCACCACATCACCTGGTGAAACAGTCACAC TCACTTGTCGCTCAAGTACTGGGGCTGTTACAACTAGTAACTATGCCAACTGGGTCCA AGAAAAACCAGATCATTTATTCACTGGTCTAATAGGTGGTACCAACAACCGAGCTCCA GGTGTTCCTGCCAGATTCTCAGGCTCCCTGATTGGAGACAAGGCTGCCCTCACCATCA CAGGGGCACAGACTGAGGATGAGGCAATATATTTCTGTGCTCTATGGTACAGCAACCA TTTC |
| IGLV2*02 _NZB | NZB/BlNJ | Musmus IGLV2*02 F | 100 | 294 | CAGGCTGTTGTGACTCAGGAATCTGCACTCACCACATCACCTGGTGGAACAGTCATAC TCACTTGTCGCTCAAGTACTGGGGCTGTTACAACTAGTAACTATGCCAACTGGGTCCA AGAAAAACCAGATCATTTATTCACTGGTCTAATAGGTGGTACCAGCAACCGAGCTCCA GGTGTTCCTGTCAGATTCTCAGGCTCCCTGATTGGAGACAAGGCTGCCCTCACCATCA CAGGGGCACAGACTGAGGATGATGCAATGTATTTCTGTGCTCTATGGTACAGCACCCA TTTC |
| IGLV3*01 _NZB | NZB/BlNJ | Musmus IGLV3*01 F | 100 | 313 | CAACTTGTGCTCACTCAGTCATCTTCAGCCTCTTTCTCCCTGGGAGCCTCAGCAAAACT CACGTGCACCTTGAGTAGTCAGCACAGTACGTACACCATTGAATGGTATCAGCAACAG CCACTCAAGCCTCCTAAGTATGTGATGGAGCTTAAGAAAGATGGAAGCCACAGCACA GGTGATGGGATTCCTGATCGCTTCTCTGGATCCAGCTCTGGTGCTGATCGCTACCTTAG CATTTCCAACATCCAGCCTGAAGATGAAGCAATATACATCTGTGGTGTGGGTGATACA ATTAAGGAACAATTTGTGTAAC |
| IGLV3*01 _PWD | PWD/PhJ | Musspr IGLV3*01 F | 95.92 | 313 | CAACCTGTGCTCACTCAGTCATCTTCAGCCTCTTTCTCCCTGGGAGCCTCAGCAAAACT CACATGCACCTTGAGTAGTGAGCACAGTACGTACATCATTGAATGGTACCAGCAACAG CCACTCAAGCCTCCTAAGTATGTGATGCAGCTTAAGAAAGATGGAAGCCACAGCAAG GGTGATGGAATCCCTGATCGCTTCTCTGGATCCAGCTCTGGTGCTGACCGCTACCTTAA CATCTCCAACATCCAGCCTGAAGATGAAGCAATATACATCTGTGGTGTGAGTGGAACA ATTAAGGAACAATTCGTGTAAC |
| IGLV2*02 _PWD | PWD/PhJ | Musmus IGLV2*02 F | 99.65 | 294 | CAGGCTGTTGTGACTCAGGAATCTGCACTCACCACATCACCTGGTGGAACAGTCATAC TCACTTGTCGCTCAAGTACTGGGGCTGTTACAACTAGTAACTATGCCAACTGGGTCCA |

| | | | | | |
|---|---|---|---|---|---|
| | | | | | AGAAAAACCAGATCATTTATTCACTGGTCTAATAGGTGGTACCAGCAACCGAGCTCCAGGTGTTCCAGTCAGATTCTCAGGCTCCCTGATTGGAGACAAGGCTGCCCTCACCATCACAGGGGCACAGACTGAGGATGATGCAATGTATTTCTGTGCTCTATGGTACAGCACCCATTTC |
| IGLV1*01_PWD | PWD/PhJ | Musmus IGLV1*01 F | 100 | 294 | CAGGCTGTTGTGACTCAGGAATCTGCACTCACCACATCACCTGGTGAAACAGTCACACTCACTTGTCGCTCAAGTACTGGGGCTGTTACAACTAGTAACTATGCCAACTGGGTCCAAGAAAAACCAGATCATTTATTCACTGGTCTAATAGGTGGTACCAACAACCGAGCTCCAGGTGTTCCTGCCAGATTCTCAGGCTCCCTGATTGGAGACAAGGCTGCCCTCACCATCACAGGGGCACAGACTGAGGATGAGGCAATATATTTCTGTGCTCTATGGTACAGCAACCATTTC |
| IGLV2*02_SJL | SJL/J | Musmus IGLV2*02 F | 100 | 294 | CAGGCTGTTGTGACTCAGGAATCTGCACTCACCACATCACCTGGTGGAACAGTCATACTCACTTGTCGCTCAAGTACTGGGGCTGTTACAACTAGTAACTATGCCAACTGGGTCCAAGAAAAACCAGATCATTTATTCACTGGTCTAATAGGTGGTACCAGCAACCGAGCTCCAGGTGTTCCTGTCAGATTCTCAGGCTCCCTGATTGGAGACAAGGCTGCCCTCACCATCACAGGGGCACAGACTGAGGATGATGCAATGTATTTCTGTGCTCTATGGTACAGCACCCATTTC |
| IGLV1*01_SJL | SJL/J | Musmus IGLV1*01 F | 100 | 294 | CAGGCTGTTGTGACTCAGGAATCTGCACTCACCACATCACCTGGTGAAACAGTCACACTCACTTGTCGCTCAAGTACTGGGGCTGTTACAACTAGTAACTATGCCAACTGGGTCCAAGAAAAACCAGATCATTTATTCACTGGTCTAATAGGTGGTACCAACAACCGAGCTCCAGGTGTTCCTGCCAGATTCTCAGGCTCCCTGATTGGAGACAAGGCTGCCCTCACCATCACAGGGGCACAGACTGAGGATGAGGCAATATATTTCTGTGCTCTATGGTACAGCAACCATTTC |
| IGLV3*01_SJL | SJL/J | Musmus IGLV3*01 F | 100 | 313 | CAACTTGTGCTCACTCAGTCATCTTCAGCCTCTTTCTCCCTGGGAGCCTCAGCAAAACTCACGTGCACCTTGAGTAGTCAGCACAGTACGTACACCATTGAATGGTATCAGCAACAGCCACTCAAGCCTCCTAAGTATGTGATGGAGCTTAAGAAAGATGGAAGCCACAGCACAGGTGATGGGATTCCTGATCGCTTCTCTGGATCCAGCTCTGGTGCTGATCGCTACCTTAGCATTTCCAACATCCAGCCTGAAGATGAAGCAATATACATCTGTGGTGTGGGTGATACAATTAAGGAACAATTTGTGTAAC |
